# Oncogenesis and Aging by Isotopic Functionalizations of the Proteins and Nucleic Acids

**DOI:** 10.1101/2020.06.27.175703

**Authors:** Reginald B. Little, Orit Uziel

## Abstract

Although the dynamics of telomeres during the life expectancy of normal cells have been extensively studied, there are still some unresolved issues regarding this research field. For example, the conditions required for telomere shortening leading to malignant transformations are not fully understood. In this work, we mass analyzed DNA of normal and cancer cells for comparing telomere isotopic compositions of white blood cells and cancer cells. We have found that the 1327 Da and 1672 Da characteristic telomere mass to charges cause differential mass distributions of about 1 Da for determining isotopic variations among normal cells relative to cancer cells. These isotopic differences are consistent with a prior theory that replacing primordial, common isotopes of ^1^H, ^12^C, ^14^N, ^16^O, ^24^Mg, ^31^P and/or ^32^S by nonprimordial, uncommon isotopes of ^2^D, ^13^C, ^15^N, ^17^O, ^25^Mg and/or ^33^S leads to altered enzymatic dynamics for modulating DNA and telomere codons towards transforming normal cells to cancer cells. The prior theory and current data are consistent also with a recently observed non-uniform methylation in DNA of cancer cells relative to more uniform methylation in DNA of normal cells. We observe further evidence of nonprimordial isotopic accelerations of acetylations, methylations, hydroxylations and aminations of nucleosides with alterations of phosphorylations of nucleotides for possibly explaining the induced mutations of DNA, RNA and proteins leading to cancer and more general alterations of DNA associated with aging. The different mass spectra of normal and cancer DNA may be reasoned by different functionalizations and isotopic enrichments as causing different motionally induced atomic and nucleotide orders by different nuclear magnetic moments (NMMs); many motionally induced oligonucleotides causing nanoscale disorder and chaos; and the many such motionally induced nanoscale chaoses of different genes causing order in macroscopic DNA for organelles organizations.

## Introduction

The functions of the DNA replications, RNA transcriptions and protein translations are altered intrinsically due to possible mutations, including the abnormal replacements of nucleotides. In normal cells, mutations are avoided by DNA repair mechanisms and due to the functions of telomeres with appropriate lengths. When telomeres are shortened they can no longer fulfill their protective functions and therefore mutations accumulate [1]. If some of these mutations are cancerous and telomerase (the enzyme that prevents telomere shortening) is upregulated, a tumor might develop [2]. Currently, it is unknown what leads to the accumulations of mutations in the DNA and the assumption is that this is these are random processes. Recent observations of international team of scientists correlated huge arrays of mutations for causing driver genes for genesis of cancer [3].

In general, random mutations that lead to constant activities of certain oncogenes products or the down regulations of other tumor suppressor gene products are the main causes of cancer. In this work, it is reasoned that isotopic induced functionalizations with clumping of isotopes on nonzero NMMs cause upregulations and/or down regulations of genes [4,5]. Cancer cells divide constantly in unregulated ways and develop tumor masses. At an advanced lethal stage, tumors develop metastasis to other organs, usually to the liver, brain and lungs.

A prior theory has proposed another explanation for the formations of mutations. According to this theory, isotopes in enzymes can manifest different chemical and enzymatic properties due to many body interactions, magnetism and nano, molecular and atomic sizes [4, 5]. By the theory [4,5] the isotopic enrichments and clumping can induce functionalizations and thereby cause upregulations and/or downregulations of genes. Thus, isotopic replacements and clumping in proteins and associated nucleic acids may lead to altered properties of the biomolecules, accounting for diseases such as cancer as by causing nonrandom mutations. On this basis, we propose that isotopic replacements of possible (common, primordial) ^1^H, ^12^C, ^14^N, ^16^O, ^24^Mg, ^31^P, ^32^S, ^43^Ca and/or others for (uncommon, nonprimordial) ^2^D, ^13^C, ^15^N, ^17^O, ^25^Mg, ^33^S, ^43^Ca and/or others may be a basis for alterations of telomere lengths and lengths and mutations of other driver genes, leading to malignant transformations of normal cells. Since isotopes (atoms of the same elements) have the same number of protons but different number of neutrons, it has been thought that isotopes have the same chemical properties for a given element. But based on the prior theory [4,5], the interior parts of enzymes were proposed to distinguish between isotopes by nuclear magnetic moments (NMM); as clumped non-zero NMM(s) were proposed to transiently alter enzymes structures, properties and dynamics affecting their activities. In this work, it is demonstrated by mass spectrometry that isotopic replacements and clumping are different in DNA of cancer cells versus DNA of normal white blood cells. This work determines that in addition to enrichments of nonprimordial isotopes of nonzero NMMs, the clumping of these nonzero-NMMs by normal cells even globally in relative abundance but clumping locally in multiple sites in enzymes and nucleic acids cause dramatic alterations of the biomolecules for causing cancer, other diseases and aging in general ways. Moreover isotopic replacements and clumping in cancer cells are shown to alter functionalizations of nucleosides, nucleotides, oligonucleotides and the telomeric gene by altering patterns of methylation, acetylation, amination, hydroxylation and phosphorylation of DNA in cancer cells relative to DNA in normal cells for causing altered upregulations, downregulations and mutations for different properties of the cancer DNA relative to the normal DNA.

On this basis, our work provides further confirmation of recent theory [4,5] and the observed selective sensitivity of cancer cells to the nonprimordial isotopes: ^25^Mg, ^43^Ca and ^67^Zn versus normal cells sensitivity. This may explain also the lack of sensitivity to the nonprimordial isotopes of ^24^Mg, ^40^Ca, and ^64^Zn of cancer and normal cells [6]. Additionally, this work may explain recent twin experiments of NASA [7] where telomeres of prolonged, space orbiting twin brother were elongated relatively to earth bound twin brother. According to this theory [4,5], cosmic rays may transmute ^1^H, ^12^C, ^14^N, ^16^O, ^24^Mg, ^32^S, and/or ^42^Ca to possibly ^2^D, ^13^C, ^15^N, ^17^O, ^25^Mg, and/or ^43^Ca, thus affecting telomerase activity and elongation of the telomeres of the twin brother in orbit relative to the brother on the surface of earth. This work further gives a theoretical basis for explaining the massive DNA sequencing and the observed nonrandom common multiple mutations oncogenesis observed among variety of many cancers in vivo [3]. Furthermore this data and the prior theory [4,5] offer theoretical basis for the experimentally observed gravitational selective killing of cancer cells relative to normal cells [8]. The data reported here and prior theory [4,5] also are consistent with observations regarding higher extent of nonrandom methylations of DNA in cancer cells [9] versus that of normal cells and [10]. The data further point to isotopic induced acetylation as important in causing the massive mutations in DNA as well as histones for ocogenesis.

## Results

The results of the mass spectroscopy analysis are shown in the following Figures I, II and III and Suppementary Materials I-XV. Figures I and II are the original data. Figure III gives specific mass intensities of importance from Figures I and II. New data {in Supplementary Figures III-VIII from a variety of different cancer DNA and more normal DNA are given} have been obtained for further substantiating Figures I and II. The new data are provided to reveal the general manifestations of these isotopic enrichment phenomena in many different cancers with emphasis on smaller oligonucleotides composing telomeres and mating telomeres (base pairs to telomere pieces). The observed masses correspond to fragmented pieces of the DNA of the cancer cells, normal B lymphocytes and total white blood cells. The observed fragmentations are induced by the energy of laser ablating the DNA in the mass spectrometer; such laser fragmentation is in analog to enzymatic and gravitational induced alterations. By analyzing the masses that are identical from the cancer (DNA-K562) to the other white (DNA-wbc) and normal B lymphocytes (DNA-skw) blood cells some fragments are assigned to the telomere codon of the DNAs. The telomere fragments were assigned to masses 1327 Da and 1672 Da on basis of prior mass analyses as in reference [11]. However, other fragments may also represent telomere codons T_2_AG_3_. These include fragments of various mass to charges ranging from −1 to - 6. Also the new data reveal mass spectral differences for normal and cancer DNA by isotopically different oligonucleotides of pentameric, tetrameric, trimeric, dimeric and monomeric nucleotides with emphasis on pieces composing telomeric codon TTAGGG and its pairing (mating) codon AATCCC. Cancer cells in general show smaller full width half maximum (FWHM) values, manifesting fewer molecular fragments and isotopic differences. This smaller FWHM of cancer are reasoned by more isotopic clumping and more nonrandom isotopic clustering in the cancer DNA. Smaller masses tend to be nucleotides and nucleosides from the DNA and some are from the telomeres. Medium mass pieces are larger segments of telomere domains and there are some heavier masses of telomeres themselves.

**Figure I.**
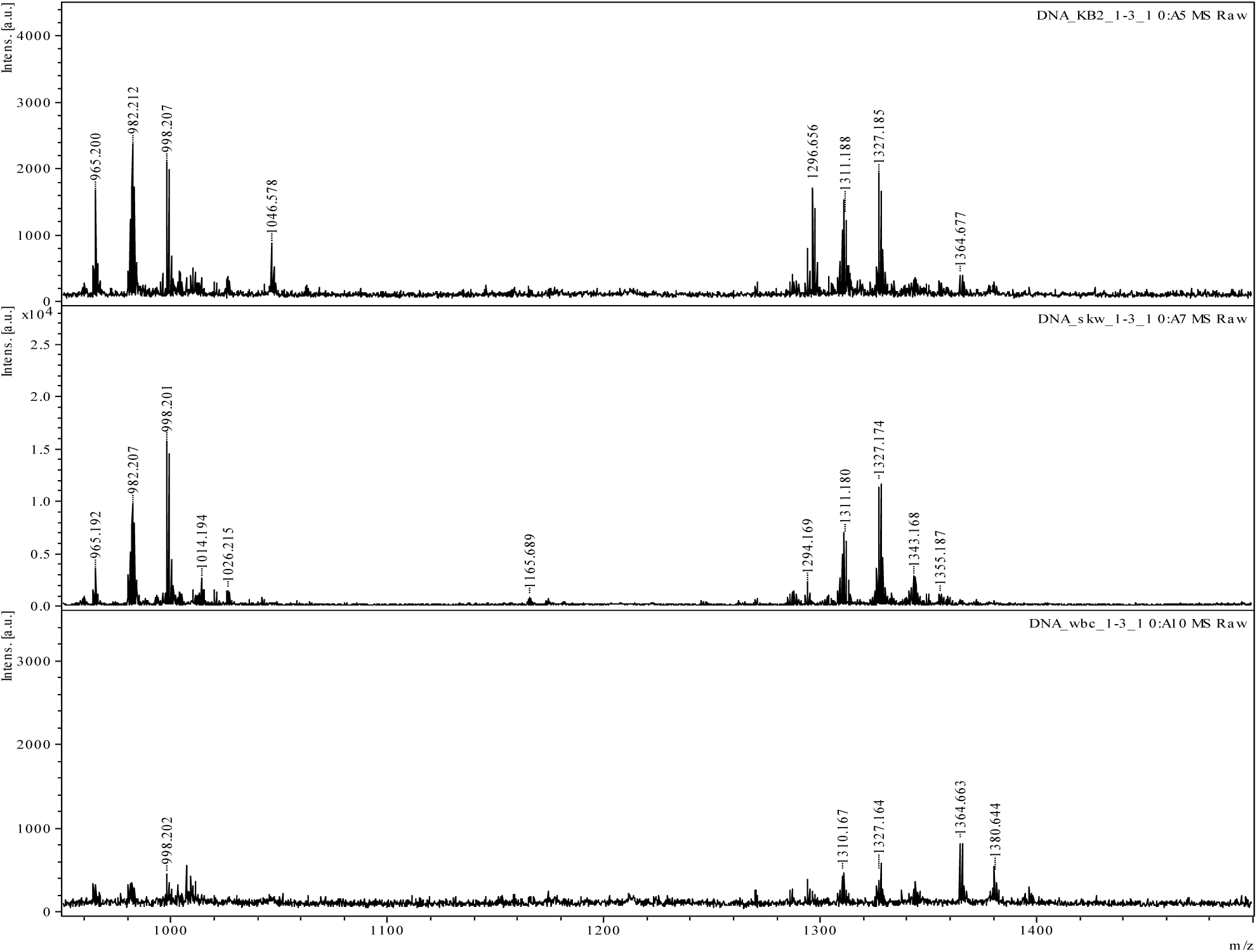
Mass Spectra from 975 Da to 1500 Da: The DNA K562 is mass spectrum of the cancer cells; the DNA SKW is the mass spectrum of the lymphocytes cells; and the DNA WBC is the mass spectrum of the white blood cells.

**Figure II.**
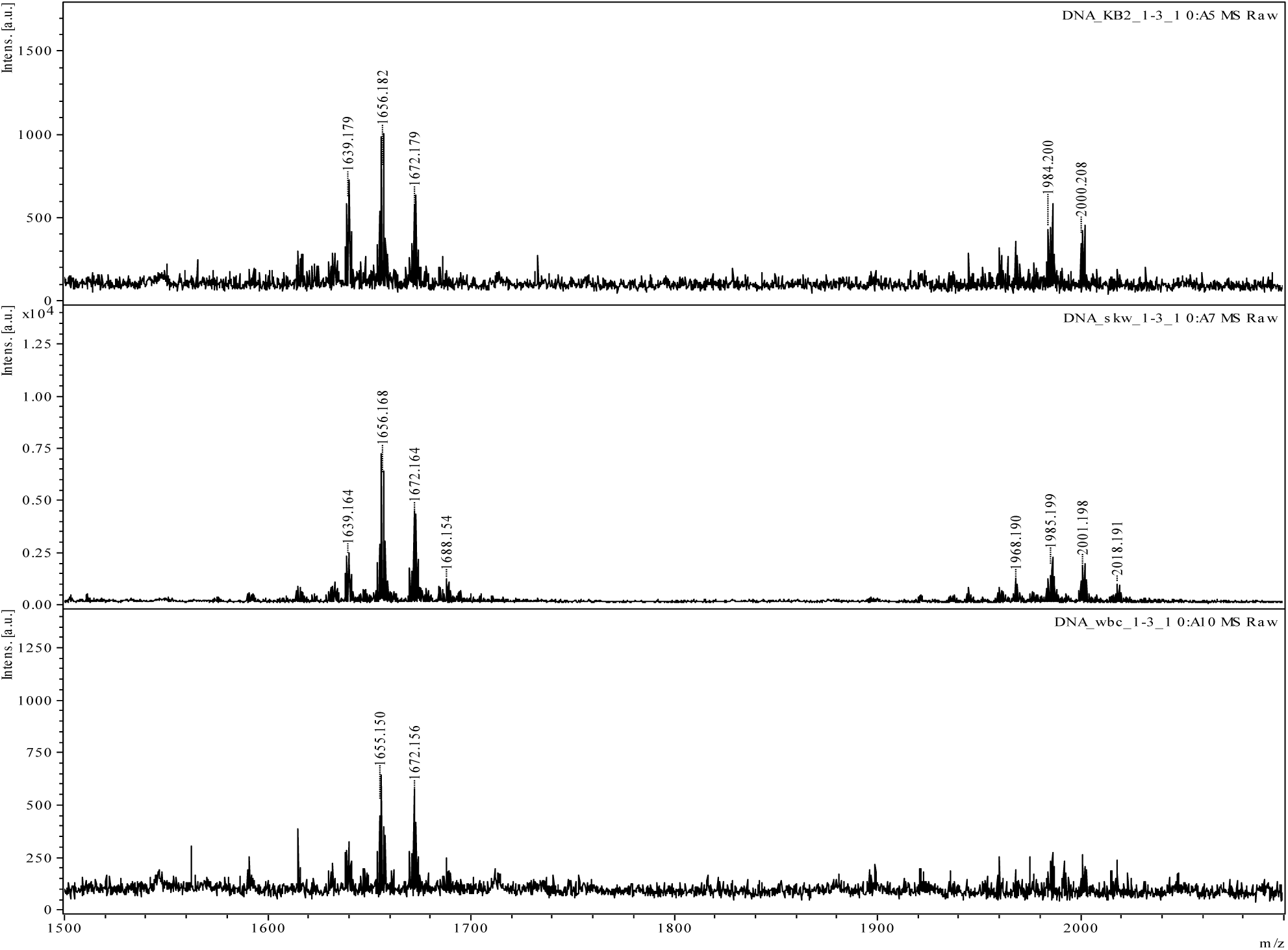
Mass Spectra from 1500 Da to 2100 Da. : The DNA K562 is mass spectrum for the cancer cells; the DNA SKW is the mass spectrum for the lymphocytes cells; and the DNA WBC is the mass spectrum for the white blood cells.

**Figure III.**
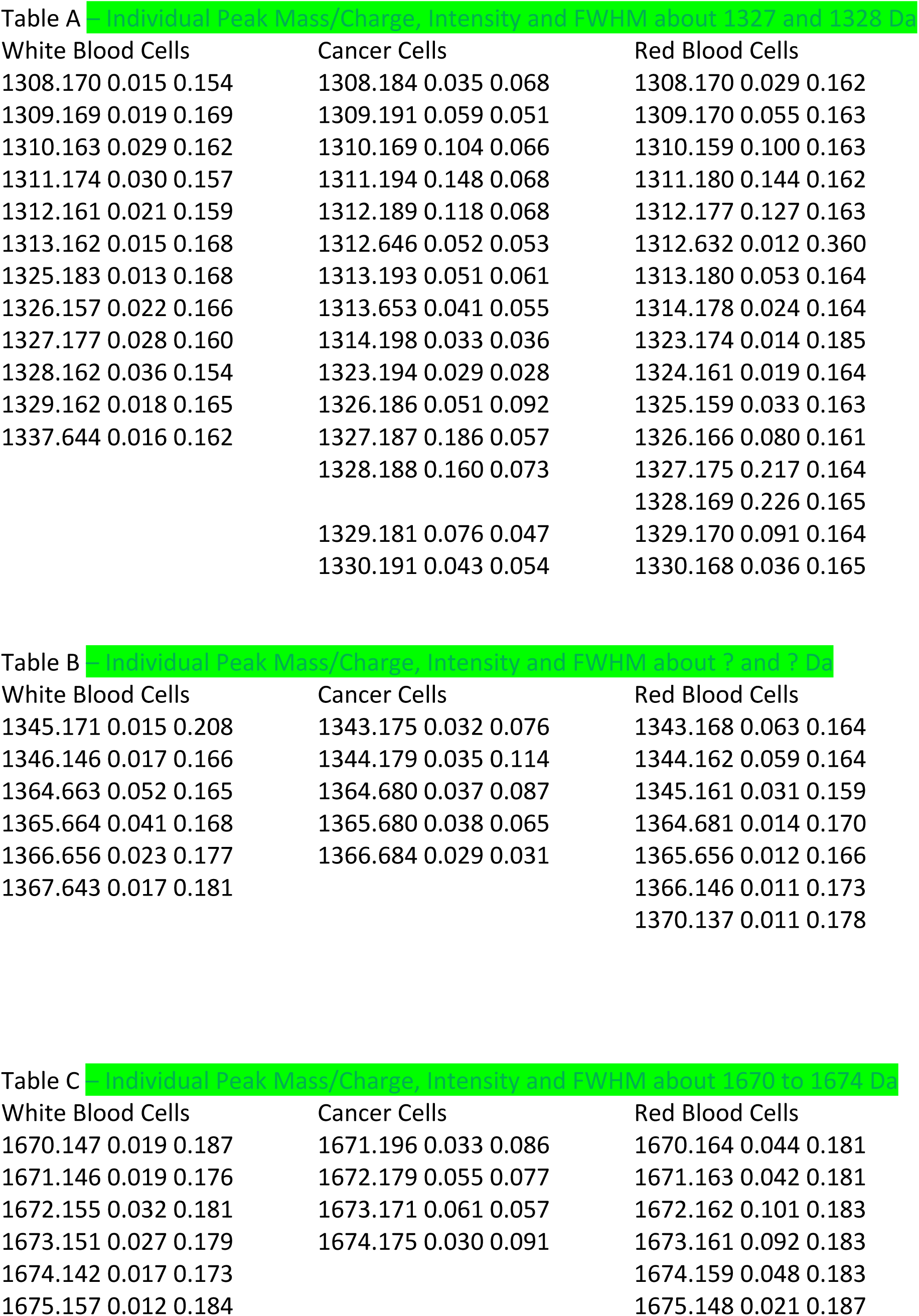

Figure I and II show mass spectra of DNA from three different cells: cancer cells, white blood cells and lymphocytes. The different peaks correspond to various fragments of the DNA having different possible masses and charges (mass to charge). According to reference [11] the masses at 1327 to 1328 Da, 1672 and 1673 Da correspond to telomere fragments. Although many other fragments are observed, the signals at 1327 Da and 1672 Da enable comparison of telomeres from the cancer cells, WBC and lymphocytes. The difference signals express different relative intensities of the DNA fragments. The different signals also express different FWHM values in the DNA of the cancer cells, WBC and lymphocytes. In general, cancer cells possess smaller FWHM relative to those of the WBC and lymphocytes. The smaller FWHMs values reflect the signal involving fewer molecular fragments, fewer isotopic differences and more isotopic clumping of nonprimordial isotopes. The 1327 Da and 1672 Da telomere signals with nearby peaks correlate with de-functionalizations (heavier) and functionalizations (less massive peaks) of these 1327 Da and 1672 Da telomeric (oligonucleotide) peaks, respectively, by possible isotopes of CH_3_, NH_2_, OH, CH3,C(O)(O) and even PO_3_^-^. More data have been measured, observed and compared for DNA from different cancer cells and more normal cells with emphasis on smaller oligonucleotides of telomeres and mating telomeres for substantiating these initial mass spectra of the whole telomere and demonstrating precision and replication of these prior measured result. See Supplementary Materials [I-XV].

In particular, the 1327 Da (mass/charge) corresponds to the telomere codon [G_3_(T_2_ AG_3_)_3_]^5-^ as reported in reference [11]. See Figure II. For this [G_3_(T_2_ AG_3_)_3_]^5-^ codon the relative intensity is greater in the cancer cell DNA at 1327 Da, but the relative intensity is greater at 1328 Da for the WBC and lymphocytes. The larger intensity at 1327 for cancer maybe due to ^15^N enrichment in the cancer telomere relative to ^14^N in the normal telomere as the smaller mass to charge can cause greater fragmenting of the ^14^N containing telomere, but the ^15^N containing telomere pieces is more stable in the larger charge. (In ^14^N, the e^-^ charge and mass are less as the e^-^ has to fiss to compensate for the nuclei fissed positive nuclear magnetic moment (NMMs). But in ^15^N the electron is more particulate as the nuclei fiss negative NMM the e^-^ fuses the negative excess and experience e^-^ e^-^ repulsion. The external electric field pushes and pulls e^-^ particle in ^15^N and electric field acts on relativistic fissed e^-^ in ^14^N, so the electric field coupled to the tiny e^-^ in the ^14^N more strongly for greater fissing and bond breaking. But the heavier e^-^ in ^15^N is less accelerated by the electric field. So the negative NMMs give nuclei less mass but the electron more charge and mass. So the positive NMMs give the e^-^ less charge and mass and the nuclei are heavier by the additional neutron. So in large positive NMMs, the e^-^ charge can be reduced so it is zero and even positive! Positive NMMs make less chemical basicity and negative NMMs make for greater chemical basicity by increasing negative charge!)

The difference of mass of 1 Da between the signals correlates with isotopic differences of elements (H, C, N, and O) in the telomeres, See Table I. At 1327 Da the cancer cells seem to be enriched in nonprimordial ^13^C or ^15^N isotopes and at 1328 Da the lymphocytes and WBC appear to be enriched in the primordial ^14^N and nonprimordial isotopes of ^17^O. The FWHM of 1327 Da for the cancerous oligonucleotide is smaller relative to FWHM at 1328 Da of WBC and lymphocytic telomeres on basis of more clumping of nonprimordial isotopes. The 1327 Da and 1328 Da mass/charge have nearby signals at 1313 Da to 1309 Da for mass differences ranging from 14 Da to 16 Da. The mass differences (Δm) of WBC are narrower with range of (1327 – 1312 Da) = 15 Da to (1328 – 1310 Da)= 18 Da. This is consistent with new data (Supplementary Materials I – XV) showing less ^17^O in normal cells as ^17^OH is heavier mass of 18 - 19 Da relative to ^13^CH_3_ and ^15^NH_2_ of 16 Da and 17 Da, respectively. The Δm of the cancer and lymphocytes are larger, ranging from (1327 – 1313 Da) = 14 Da to (1328 – 1309 Da) = 19 Da. This is consistent with new data showing more ^17^OH in cancer cells. These mass differences may involve the 1327 to 1328 Da defunctionalization by loss of CH_3_, NH_2_ and/or OH groups to masses 1309 to 1313 Da. It is also observed that less massive signals at 1303 – 1309 Da for mass differences (Supplementary Materials I – XV) of (1327 – 1304 Da)= 23 Da to (1328 – 1303 Da) = 25 Da, which may involve loss of ^24^Mg^2+^ or ^25^Mg^2+^ from the telomere pieces. The Mg^2+^ is known to bind DNA and are associated with many enzymes associated with replications of nucleic acids [4]. In addition to the less massive signals about 1317 Da to 1328 Da, more massive signals are observed. Heavier signals at 1333 to 1334 Da involve Δm of 5-7 Da which are possible for mass to charge range of 10 - 14 Da demonstrating possible methylation/demethylation of the telomeres. Heavier signals in range 1341 to 1345 Da involve Δm ranging from (1341-1328 Da) = 13Da to (1345 – 1327 Da) = 18 Da or Δm = 13-18 Da explaining range of possible methylations, aminations and hydroxylations of the telomere, See Table II (below). Heavier signals in range 1349 to 1350 Da involve Δm of (1349 −1328 Da) = 21 Da to (1350 – 1327 Da) = 23 Da that may explain possible hydroxylation or Na^+^ and/or Mg^2+^ ions functionalizations of the telomeres.

The new data for 11 different cancerous DNA and 4 normal DNA (Supplementary Materials I – XV) give reproductions and precisions of these prior data. The new DNA from the 11 different cancerous DNA from different types of cancers were also observed to have different relative intensities at 1127 Da and 1128 Da in consistency to prior cancer DNA in Figures I, II, and II. The new data of the 11 different cancer DNA and 4 normal DNA are also observed to fragment by defunctionalizations to smaller pieces (of oligotelomeric pentanucleotides, tetranucleotides, trinucleotides, dinucleotides and mononucleotides) in different ways. These oligonucleotides from fragmentations of these 11 different cancer DNA and 4 different normal DNA also were observed to have different functionalizations of heavier masses by methylations, acetylations, and phosphrylations. Furthermore, the new data give consistent observations on fragmented pieces of the telomeres and mating telomeres themselves with observations of all possible oligonucleotides of pentanucleotides, tetranucleotides, trinucleotides, dinucleotides and mononucleotides of the telomere with isotopic enrichments and functionalizations that vary between cancer and normal telomeres. In assessing the mass spectra (See Supplementary Materials I-XV.), it is reasoned by the model that the laser ablation energizes the fragmentation of the DNA in analogous way as the enzymatic cleaving and binding of the DNA in the cancer and normal cell biology. See Tables I-V and Supplementary Materials I-XV.

The 1670-1674 Da (mass to charge) corresponds to the telomere codon [G_3_(T_2_AG_3_)_3_]^4-^, See Figure II. For this [G_3_(T_2_AG_3_)_3_]^4-^ codon, the relative intensity is greater for the cancer cell at 1674 Da, but at 1672 Da the relative intensity is greater for the WBC and lymphocytes. See Table III. The difference in mass of 1Da between the samples correlates with isotopic differences of elements (H, C, N, and O) in the telomeres. The FWHM of the 1673 Da signal of cancer telomeres is smaller than the FWHM of the 1672 Da of the WBC and lymphocyte telomeres. The 1672 to 1673 Da mass to charges have nearby signals of lesser mass to charges at 1655 to 1657 Da for mass differences ranging (1672-1657 Da) = 15 Da to (1673-1655 Da) = 18 Da demonstrating possible aminations or hydroxylations. At 1657 Da, the cancer is enriched in nonprimordial isotopes and WBC and lymphocytes are enriched in the primordial isotopes. Such aminations of the cancer telomere as observed by the 1657 Da in the cancer DNA are consistent with new data of 11 different cancer DNA revealing enriched ^15^N in C, and A nucleosides relative to C and A nucleosides in normal DNA! (See Supplementary Materials I-XV). The aminations manifest this negative mass effect as by the negative NMM of ^15^N with consequent losses of mass, relativistically. In addition to the less massive signal about the 1672 to 1673 Da, more massive signal at 1688 to 1689 Da is observed. The lymphocytes and WBC are enriched at 1688 Da and cancer is enriched at 1689 Da. Heavier signal at 1688 and 1689 Da involves Δm (1688 – 1673 Da) = 15 Da to (1689 −1672 Da) = 17 Da (15 to 17 Da) reflecting possible aminations (15 to 16 Da) and/or hydroxylations (16-17 Da). Such greater nonprimordial aminations and hydroxylations of the telomeric codon in cancer DNA is consistent with observed greater enrichments of G nucleosides from 11 different cancer DNA relative to 4 different normal DNA! See Supplementary Materials I-XV. The FWHM at 1672 to 1673 Da is bigger or smaller than the FWHM at 1688 to 1689 Da.

The new data of the 11 cancer DNA and 4 normal DNA reveal similar trends in telomeric peaks 1327 and 1672 Da to further confirm the isotopic differences of normal vs cancer, where the whole telomere is measured. The new data provides more observations of the pieces of the telomere as by measuring masses of oligonucleotides, nucleotides and nucleosides (A, G, T, C) composing telomere with various possible methylations, acetylations, aminations, hydroxylations, and phosphorylations. The results determine distinct differences in masses of the A, G, T and C nucleotides and oligonocleotides that compose the telomere and mating telomere (base pair of the telomere). The observed spectra are further different with more isotopic substitution and clumping in G and T nucleotides relative to A and C nucleotides. The intensities, FWHM, functionalizations and isotopic clumping are observe to consistently manifest in the pentanucleotides and tetranucleotides (composing the telomere, [G_3_(T_2_AG_3_)_3_]_n_^4-^ and mating telomere [C_3_(A_2_TC_3_)]_n_^4-^) and also all possible trinucleotides, dinucleotides and mononucleotides (composing the telomere) for the different cancer cell lines. However the pattern of intensities, FWHM, functionalizations, and isotopic clumping differs in the new four normal pentanucleotides, tetranucleotides, trinucleotides, and dinucleotides and mononucleotides as fragmented from the normal telomeres. The cancer has more (or less) intensities of G, T and C and perhaps normal has more A. See Supplementary Materials I-XV. The dinucleotide possibilities from the telomere include: GG, GT, TA, TT, AG. The mates or base pairs of the telomere include: CC, CA, AT, AA, TC. See Table II. For the dinucleotides the normal reveals larger fragmentations with broader FWHM. The functionalizations vary for normal verse cancer. It seems that more of these pieces are fragmented from cancer DNA and more isotopic enriched in cancer. The dephosphorylations of the cancer was lesser for cancer dinucleotides relative to normal dinucleotide fragments from the telomere. The trinucleotides of telomeres include: GGG, GGT, GTT, TTA, TAG, and AGG and their mates (base pairs) telomere pieces: CCC, CCA, CAA, AAT, ATC, and TCC. See Supplementary Materials I-XV. The cancer and normal have unique patterns of these trinucleotides as well as possible tetranucleotides and pentanucleotides. The possible tetra-nucleotides of telomeres include: TTAG, TAGG, AGGG, GGGT and GGTT and their mates or base pairs telomeric pieces: AATC, ATCC, TCCC, CCCA and CCAA. See Supplementary Materials I-XV. These tetranucleotides also manifested different patterns of stability in cancer relative to normal DNA sources. The possible pentanucleotides include: GGGAT, GGATT, TTGGG. And the mating pentanucleotide pieces include: CCCTA, CCTAC and AACCC. See Supplementary Materials I-XV. These variations of intensities, functionalizations, phosphorylations and isotopic clumping of the pieces of the telomere follow the prior data of the whole telomere gene.

### Pentanucleotides

In Table V, it is observed that the highly phosphorylated telomeres are more stable with heavy isotopic enrichments. Highly phosphorylated mating telomere are stable with heavy isotopic enrichments. Methylations of 15P tends to stabilize cancer in telomere but destabilize mating telomere, both with heavy and some light isotopic enrichments. Acetylations more strongly stabilize cancer in mating telomeres and telomeres. Dephosphorylations from 15 to 14 phosphates and destabilizations of telomeres are observed with less isotopic enrichments. The mass spectra are in Supplementary XV. Methylations of 14P stabilize telomeric cancer and normal but destabilize mating telomeres. Acetylations more strongly stabilizes normal telomere and cancer mating telomere with heavy and light isotopic enrichments. Dephosphorylations 14P to 13P stabilize both cancer and normal telomere with isotopic enrichments but destabilize mating telomere. The mass spectra are in Supplementaries XIV and V. Functionalizations by methyl and acetyl do not much alter the dephoshosphorylations to 13P stabilities. Dephosphorylations 13P to 12P destabilize normal and slightly destabilize cancer in telomere and mating telomere with negligible isotopic enrichments and negligible change in stabilities with methylations. The mass spectra are in Supplementary XIV. The acetylations with the dephosphorylations 13P to 12P do isotopically enrich with light nuclei the mating. Dephosphorylations 12P to 11P do not much alter the instabilities and the tiny stabilities are for trace amounts of cancer pentanucleotides. The mass spectra are in Supplementary XIV. Methylations and acetylations show very little increase in stabilities and no dramatic effects of acetylations relative to methylations. Dephosphorylations 11P to 10P cause slightly less stabilities of the cancer telomeres with negligible enhancements in stabilities from isotopic enrichments methylations and acetylations. The mass spectra are in Supplementary II and XIV. But the dephosphorylations 11P to 10P of the mating telomeres reveal dramatic increased stabilities of the normal mating telomeres with increased stabilities of normal mating telomeres upon methylations and contrary increased stabilities of the cancer mating telomeres with isotopic enrichments by light nuclei by acetylations. These last data for 11P to 10P of the pentanucleotides overall the masses of 12P to 10P for tetranucleotides so the observed masses may not truly represent the pentanucleotides.

So continuing with the pentanucleotides mixed with the tetranucleotides, then the dephosphorylations from 10P to 9P causes increased stabilities of the cancer telomeres with isotopic enrichments of heavy and light nuclei and methylations and acetylations isotopically enriching the cancer telomeres. The mass spectra are in Supplementary II. But the dephosphorylations 10P to 9P of the mating telomere slightly stabilizes the normal with the methylations and more strongly the acetylations stabilizing the normal mating telomeres. It is important to consider that for the dephosphorylations 10P to 9P, the number of phosphates per nucleotide in the pentanucleotides is less than 2 and this correlates with changes in dynamics. The changes in dynamics from such may reflect overlap with the denucleotiding to fewer nucleotides in the oligonucleotides. The dephosphorylations from 9P to 8P cause increased stabilities of the normal telomeric nucleotides with methylations enhancing normal nucleotide telomere stabilities and acetylations enhancing cancer nucleotide telomeric stabilities with isotopic enrichment. The mass spectra are in Supplementaries XIV and II. Unlike the 13P to 10P dynamics, now the methylations and acetylations cause distinct differences and this may point to dinucleotides to smaller oligonucleotides and overlap in masses of oligonucleotides and smaller oligonucleotides. The dephosphorylations of the mating telomeres from 9P to 8P causes the mixed cancer and normal to be trace amounts with negligible alterations upon methylation or acetylations for the mating telomeres from 9P to 8P. So as dephosphorylate 8P to 7P, the cancer telomeres become more stable; normal less stable with heavy isotopic enrichments and the methylations slightly stabilize cancer but acetylations stabilize the normal telomeres for distinct differences in methylations and acetylations on nucleotide stabilities. The dephosphorylations of the telomere from 8P to 7P cause cancer stabilities of both telomeres and mating telomeres with methylations stabilizing the cancer and both the dephosphosphorylations and methylation enhancing isotopic enrichments of heavy isotopes. The mass spectra are in Supplementaries III and II. The acetylations upon the dephosphophorylations 8P to 7P cause more stable normal telomeres and mating telomeres. The dephosphorylations from 7P to 6P cause negligible increase in stabilities for trace stabilities with isotopic enrichments of the cancer telomeres and mating telomeres. The methylations of the telomeres slightly stabilize the cancer telomeres and negligibly changes the trace stabilities of the mating telomere. The mass spectra are in Supplementaries IV, III and II. The acetylations of the negligibly stabilize the trace cancer telomeres and mating telomeres with isotopic enrichments. The dephosphorylations of 6P to 5P causes the cancer telomeres and mating telomeres to be more stable and isotopically enriched with the methylations more stabilizing the telomeres and mating telomeres of cancer and isotopically enriching the cancer mating and telomeres. The mass spectra are in Supplementaries V, IV and III. But the acetylations with the dephosphorylations from 6P to 5P causes the negligible stabilizations of the telomere but increased stabilization of the normal mating telomeres. The dephosphorylations from 5P to 4P caused slight increases in stabilities of normal telomeres and mating telomeres with the methylations reinforcing the stabilities of the normal telomeres and mating telomeres, but the acetylations stabilize the cancer telomeres with heavy isotopic enrichments and negligible effect of acetylations of mating telomeres. The mass spectra are in Supplementaries III, IV and V. It is important to note that these observations for the heavily dephosphorylated pentanucleotides at 4P overlap with not only the tetranucleotides but also overlap with the highly phophosphorylated trinucleotides.

### Tetranucleotides

The dephosphorylations from 12P to 11P cause destabilizations of cancer telomeres with isotopic enrichments. See Table IV. This is characteristic of highly phosphorylated oligonucleotides of specific number of nucleotides dephosphorylating. The mass spectra are in Supplementaries II and III. But the dephosphorylations from 12P to 11P cause stabilizations of cancer mating telomeres with reductions in isotopic enrichments. The telomeres at 12P are stable for cancer but unstable for normal and the methylations slightly destabilize the cancer telomeres and likewise the acetylations slightly destabilize the cancer telomeres. The mating telomeres are more stable for normal pieces and the methylations and acetylations destabilize the normal mating telomeres causing more cancer stabilities. The dephosphorylations to 11P and methylations cause the telomeres to be more stable for normal than more stable for cancer with slight isotopic enrichments. The acetylations of the 11P cases more stable mixed cancer and normal telomeres. But the dephosphorylations and methylations of mating telomeres to 11P causes the cancer to be stable than the mixed cancer and normal mating telomeres to be stable with less isotopic enrichments. The mating telomeres at 11P become more stable for cancer upon acetylations.

The dephosphorylations from 11P to 10P cause the stabilizations of normal telomeres with isotopic enrichments. This is characteristic of highly phosphorylated oligonucleotides of specific number of nucleotides dephosphorylating. The mass spectra are in Supplementaries III and II. But the dephosphorylations from 11p to 10P cause stabilizations of cancer mating telomeres with huge increase in isotopic enrichments. The telomeres at 10P are stable for mixed cancer and normal tetranucleotides with huge isotopic enrichments and the methylations cause cancer stabilities but the acetylations cause normal stabilities with isotopic enrichments. The mating telomeres are more stable for cancer at 10P with huge isotopic enrichments and the methylations cause normal stabilities with isotopic enrichments and acetylations cause mixed cancer and normal with isotopic enrichments.

The dephosphorylations from 10P to 9P cause the stabilizations of cancer telomeres with isotopic enrichments. This is characteristic of highly phosphorylated oligonucleotides of specific number of nucleotides dephosphorylating. The mass spectra are in Supplementaries II, III and IV. But the dephosphorylations from 10P to 9P of the mating telomeres with loss of isotopic enrichments do not much alter the large stabilities of the mating telomeres. The telomeres at 9P is stable for cancer and of high isotopic enrichment and the methylations do increase the cancer stabilities and isotopic enrichments but the acetylations cause more mix of normal and cancer stabilities of the telomeres. The mating telomeres are slightly more cancer stable with not as much isotopic enrichments at 9P and the both methylations and acetylations cause cancer stabilities of trace amounts with less isotopic enrichments.

The dephosphorylations of the telomeres from 9P to 8P cause the slightly increased cancer stabilities with diminished isotopic enrichments. This is characteristic of highly phosphorylated oligonucleotides of specific number of nucleotides dephosphorylating. The mass spectra are in Supplementaries III and IV. But the dephosphorylations from 9P to 8P of the mating telomeres with loss of isotopic enrichments causes slight stabilizations of normal mating telomeres and diminish stabilities of cancer mating telomeres. The telomeres at 8P is more stable for cancer relative to 9P and less isotopically enriched and methylations and acetylations further increase the stabilities of the cancer telomeres with isotopic enrichments. The mating telomeres at 8P is slightly more stable in normal tetranucleotides with less change relative to 9P phosphorylated and less isotopically enriched and methylations and acetylations increase normal and cancer mating telomeres, respectively, with slight isotopic enrichments.

The dephophorylations of the telomeres from 8P to 7P cause increase on normal stabilities with no isotopic enrichments. This change in type of telomeres and loss of enrichment sare not characteristic of highly phosphorylated oligonucleotides of specific number of nucleotides dephosphorylating. The change from cancer to normal (and the loss of isotopic enrichments) under such large number of dephosphorylations reflect overlap in chemistry of the tetranucleotides and trinucleotides as the trinucleotides begin to produces masses in this range. The loss of enrichments driving the dynamics points to more trinucleotide contributions to the masses as the trinucleotides relative to tetranucleotides have less probable clumping of nonprimordial isotopes. The mass spectra are in Supplementaries III and IV. But the dephosphorylations from 8P to 7P of mating telomeres cause no change in relative normal to cancer mating telomeres with increased intensities of both and no isotopic enrichments. Such mixed contributions of the mating telomeres reflect overlap and masses of both tetranucleotides and trinucleotides. Very little changes in the stability patterns of the mating telomeres for the dephosphorylations point to different nucleotide numbers; oligonucleotides as the trinucleotides have less probable clumping of nonprimordial isotopes relative to tetranucleotides for less isotopic driven clumping with dephosphorylations. The telomeres are 7P are stable for the normal nucleotides with no isotopic enrichments and methylations further stabilizes the normal telomeres (likely reflecting the contributions of the trinucleotides with 9P) but the acetylations causing more stabilities of the cancer telomeres. The mating telomeres at 7P are mixed in stabilities of many cancer and many normal mating telomeres with no isotopic enrichments and the methylations and acetylations cause enhanced normal mating telomeric stabilities without isotopic enrichments. The dephosphorylations of the telomeres from 7P to 6P cause increase in stabilities of the normal telomeres with no isotopic enrichments. This change in type of telomeres and loss of enrichments are characteristic of highly phosphorylated oligonucleotides of specific number of nucleotides dephosphorylating. The change from cancer to normal (and the loss of isotopic enrichments) under such large number of dephosphorylations reflect overlap in chemistry of the tetranucleotides and trinucleotides as the trinucleotides begin to produce masses in this range. The loss of enrichments driving the dynamics points to more trinucleotide contributions to the masses as the trinucleotides relative to tetranucleotides have less probable clumping of nonprimordial isotopes. The mass spectra are in Supplementaries IV, V, and VI. But the dephosphorylations from 7P to 6P cause no change in the mixed nature of the mating telomeres. Such mixed contributions of the mating telomeres reflect overlaps and masses of both tetranucleotides and trinucleotides. Very little changes in the stability patterns of the mating telomeres for the dephosphorylations point to different nucleotide numbers; oligonucleotides as the trinucleotides have less probable clumping of nonprimordial isotopes relative to tetranucleotides for less isotopic driven clumping with dephosphorylations. The telomeres are 6P are more stable for the normal nucleotides with no isotopic enrichments and methylations further stabilize the normal telomeres (likely reflecting the contribution of the trinucleotides with 9P) but the acetylations causing more stabilities of the cancer telomeres.The mating telomeres at 6P are mixed in stabilities of many cancer and many normal mating telomeres with no isotopic enrichments and methylations cause enhanced normal mating telomeric stabilities without isotopic enrichments; but the acetylations cause more cancer mating telomeres with greater isotopic enrichments. These effects of the acetylations for changing normal to cancer at 6P reflect the trinucleotide nature and the greater tendency of acetyl to introduce nonprimordial isotopes relative to methyl as the trinucleotides have fewer probable nonprimordial isotopes relative to tetranucleotides and pentanucleotides and the nonprimordial functionalizations by the acetyl are better detected here in the trinucleotides.

The dephosphorylations of the telomeres from 6P to 5P cause increase in stabilities of the cancer telomeres and diminish stabilities of normal for mixed normal and cancer with slight isotopic enrichments. This change in type of telomeres and loss of enrichments are characteristic of highly phosphorylated oligonucleotides of specific number of nucleotides dephosphorylating. The change from normal to more cancer stabilities (and the loss of isotopic enrichments) under such large number of dephosphorylations reflect less overlap in chemistry of the tetranucleotides and trinucleotides as the tetranucleotides begin to less produce masses in this range. The gain of enrichments driving the dynamics points to more trinucleotide contributions to the masses as the trinucleotides relative to tetranucleotides have less probable clumping of nonprimordial isotopes and the low nonprimordial isotopes in trinucleotides are more sensitive to loss of positive NMMs of ^31^PO_3_^-^. The mass spectra are in Supplementaries V, and VI. But the dephosphorylations from 6P to 5P cause no change in the mixed nature of the mating telomeres. Such mixed contributions of the mating telomeres reflect overlaps and masses of both tetranucleotides and trinucleotides. Very little changes in the stability patterns of the mating telomeres for the dephosphorylations point to different nucleotide numbers; oligonucleotides as the trinucleotides have less probable clumping of nonprimordial isotopes relative to tetranucleotides for less isotopic driven clumping with dephosphorylations. The telomeres are 5P are less stable for the normal nucleotides relative to 6P with no isotopic enrichments and methylations further stabilize the 5P cancer telomeres (likely reflecting the contributions of the trinucleotides with 9P) but the acetylations causing more stabilities of the normal telomeres at 5P. The mating telomeres at 6P are mixed in stabilities of many cancer and many normal mating telomeres with no isotopic enrichments and the methylations cause enhanced cancer mating telomeric stabilities with isotopic enrichments; but the acetylations cause more cancer mating telomeres with greater isotopic enrichments. These effects of the acetylations and methylations for changing normal to cancer at 5P reflect the trinucleotide nature and the greater tendencies of acetyl and methyl to introduce nonprimordial isotopes in the trinucleotides (relative to tetranucleotides and pentanucleotides) as the trinucleotides have fewer probable nonprimordial isotopes and the nonprimordial functionalizations by the acetyl and methyl are better detected here in the trinucleotides.

The dephosphorylations of the telomeres from 5P to 4P causes even greater increase in stabilities of the cancer telomeres and even greater diminish stabilities of normal for cancer with isotopic enrichments. This lesser changes in type of telomere and greater of enrichments are characteristic of lesser phosphorylated oligonucleotides of specific number of nucleotides dephosphorylating. The change from normal to more cancer stabilities (and the loss of isotopic enrichments) under such large number of dephosphorylations reflect even lesser overlaps in chemistry of the tetranucleotides and trinucleotides as the tetranucleotides do not produce masses in this range. The slight gains of enrichments driving the dynamics points to more trinucleotide contributions to the masses as the trinucleotides relative to tetranucleotides have less probable clumping of nonprimordial isotopes and the low nonprimordial isotopes in trinucleotides are more sensitive to loss of positive NMMs of ^31^PO_3_^-^. The mass spectra are in Supplementaries V, VI, and VII. But the dephosphorylations from 5P to 4P causes no change in the mixed nature of the mating telomeres with more intense cancer and normal mating telomeres. Such mixed contributions of the mating telomeres reflect less overlaps and masses of both tetranucleotides and trinucleotides as the trinucleotides contribute more to these 4p trinucleotides. Slight changes in the stability patterns of the mating telomeres for the dephosphorylation point to different nucleotide numbers; oligonucleotides as the trinucleotides have less probable clumping of nonprimordial isotopes relative to tetranucleotides for more sensitive isotopic driven clumping with dephosphorylations. The telomeres are 4P are even lesser stable for the normal nucleotides relative to 5P with more isotopic enrichments and methylations further more stabilize the 4P normal telomeres (likely reflecting the contributions of the trinucleotides with 8P) but the acetylations causing more stabilities of the normal telomeres at 4P with greater isotopic enrichments. The mating telomeres at 4P are mixed in stabilities of many cancer and many normal mating telomeres with larger isotopic enrichments and the methylations cause enhanced cancer mating telomeric stabilities with less isotopic enrichments; but the acetylations cause mixed normal and cancer mating telomeres with lesser isotopic enrichments. These effects of the acetylations and methylations for changing normal to cancer at 4P reflect the trinucleotide natures and the greater tendencies of acetyl and methyl to introduce nonprimordial isotopes in the trinucleotides (relative to tetranucleotides and pentanucleotides) as the trinucleotides have fewer probable nonprimordial isotopes and the nonprimordial functionalizations by the acetyl and methyl are better detected here in the trinucleotides.

### Trinucleoside

The fully saturated (9P) telomeric trinucleotides manifested stable mixed intensely stable normal and cancer telomeres with more stable normal peaks with isotopic enrichments by light isotopes. See Table III. But the mating telomeres of full phosphorylations manifest stable cancer telomers with less isotopic enrichments. The dephosphorylations from 9P to 8P caused the change for stable cancer telomeres and mating telomeres. This is characteristic of highly phosphorylated oligonucleotides of specific number of nucleotides dephosphorylating. The mass spectra are in Supplementaries V and VI. The telomeres at 9P fully phosphorylated undergo increased cancer stabilities by methylations with isotopic enrichments but the acetylations cause mixed cancer and normal with isotopic enrichments. The mating telomeres at 9P full phosphorylations undergo methylations to manifest mix of stable normal and cancer mating telomeres with slight isotopic enrichments. The telomeres at 8P undergo slight drop in cancer stabilities and isotopic enrichments but manifest continued cancer stabilities by acetylations with slightly less isotopic enrichments. The mating telomeres at 8P undergo negligible change in cancer stabilities with isotopic enrichments during methylations, whereas the acetylations slightly changes the cancer stabilities with greater isotopic enrichments by light isotopes.

The dephosphorylations from 8P to 7P caused the lower stabilities of cancer telomeres with mix of stable normal and cancer nucleotides and less isotopic enrichments. This is characteristic of highly phosphorylated oligonucleotides of specific number of nucleotides dephosphorylating. The mass spectra are in Supplementaries VI and V. The dephosphorylations from 8P to 7P of mating telomeres caused a mixed of stable cancer and normal mating telomeres of large intensities and change from heavy isotopic enrichments to light isotopic enrichments. The methylations and acetylations of telomeres at 7P caused stabilizations of cancer telomeres with high intensities and isotopic enrichments. But the methylations and acetylations of mating telomeres caused stabilizations and increased intensities of cancer mating telomeres and mixed normal and cancer telomeres, respectively.

The dephosphorylations from 7P to 6P caused the stabilizations of normal telomeres with greater isotopic enrichments by light isotopes of ^15^N. This is characteristic of highly phosphorylated oligonucleotides of specific number of nucleotides dephosphorylating. The mass spectra are in Supplementaries VI, VII, and VIII. The dephosphorylations from 7P to 6P of mating telomeres caused less changes in stabilities with maintaining mix of normal and cancer mating telomeres. The methylations and acetylations at 6P maintain the stabilizations of the normal telomeres and slightly stabilize the cancer telomeres with isotopic enrichments, respectively. The methylations and acetylations at 6P of the mating telomeres caused stabilizations of cancer meting telomeres.

The dephosphorylations from 6P to 5P caused the even greater stabilizations of normal telomeres with less isotopic enrichments. This is characteristic of highly phosphorylated oligonucleotides of specific number of nucleotides dephosphorylating. The mass spectra are in Supplementaries VI, VII, and VIII. The dephosphorylations from 6P to 5P of mating telomeres caused negligible changes in relative stabilities of cancer and normal mating telomeres with increased intensities of both normal and mating telomeres and more isotopic enrichments by lighter isotopes. The methylations and acetylations at 5P of the telomeres caused similar patterns of stable normal telomeres with isotopic enrichments and increased stabilities of cancer telomeres with slight isotopic enrichments, respectively. The methylations and acetylations at 5P of the mating telomeres caused stabilizations of the normal mating telomeres with isotopic enrichments of mating telomeres and stabilizations of cancer mating telomeres with less isotopic enrichments, respectively, by acetylations.

The dephosphorylations from 5P to 4P caused the diminished stabilities of the normal telomeres with increased in isotopic enrichments of lighter isotopes. This is not characteristic of highly phosphorylated oligonucleotides of specific number of nucleotides dephosphorylating. The mass spectra are in Supplementaries VII, VIII, and IX. The dephosphorylations from 5P to 4P of mating telomeres caused decreased stabilities of the normal mating telomeres with isotopic enrichments of lighter isotopes. The methylations and acetylations at 4P of telomeres caused negligible change in stabilizations of mixed normal and cancer telomeres and stabilizations of normal telomeres with enrichments of light isotopes and heavy isotopes, respectively by acetylations. The methylations and acetylations at 4P of mating telomeres caused negligible changes in cancer stabilities with light isotopic enrichments and decreased cancer stabilities for mixed normal and cancer mating telomeres, respectively, by acetylations.

### Dinucleotides

The fully saturated (6P) telomeric dinucleotides manifested stable, intensely stable cancer telomeres with isotopic enrichments by heavy isotopes. See Table II. But the mating telomeres of full phosphorylations manifest stable mixed normal and cancer mating telomeres with heavy isotopic enrichments. The dephosphorylations from 6P to 5P caused the change for more stable normal telomeres and changes for more stable cancer mating telomeres. This is characteristic of highly phosphorylated oligonucleotides of specific number of nucleotides dephosphorylating. The mass spectra are in Supplementaries VIII and IX. The telomeres at 6P fully phosphorylated undergo increased mixed normal and cancer stabilities by methylations with isotopic enrichments but the acetylations cause increased cancer stabilities with isotopic enrichments. The mating telomeres at 6P full phosphorylations undergo methylations to manifest mix of stable cancer mating telomeres with isotopic enrichments, but the mating telomeres undergo stabilizations of mixed normal and cancer telomeres at 6P with heavy isotopic enrichments. The telomeres at 5P undergo changes in stabilities from cancer to mixed normal and cancer stabilities and isotopic enrichments for methylations, but the acetylations cause changes from cancer stabilities to normal stabilities with isotopic enrichments.

The dephosphorylation from 5P to 4P caused the changes for more stable cancer telomeres and changes for more stable and more intense cancer mating telomeres. This is characteristic of highly phosphorylated oligonucleotides of specific number of nucleotides dephosphorylating. The mass spectra are in Supplementaries IX and XI. The telomeres at 4P fully phosphorylated undergo slightly decreased cancer stabilities by methylations and acetylations with isotopic enrichments. The mating telomeres at 4P full phosphorylations undergo methylations to manifest negligible changes in stabilities of cancer mating telomeres with isotopic enrichments, but the mating telomeres undergo slight lower stabilizations of cancer telomeres at 4P with heavy isotopic enrichments.

The dephosphorylations from 4P to 3P caused the negligible changes in stabilities of stable cancer telomeres and mating telomeres. This is characteristic of highly phosphorylated oligonucleotides of specific number of nucleotides dephosphorylating. The mass spectra are in Supplementaries IX and X. The telomeres at 3P phosphorylated undergo slightly decreased cancer stabilities by methylations and acetylations with isotopic enrichments. The mating telomeres at 3P phosphorylations undergo methylations to manifest slightly lower stabilities of cancer mating telomeres with isotopic enrichments, and the mating telomeres undergo negligible changes in stabilizations of cancer telomeres at 3P with heavy isotopic enrichments.

The dephosphorylations from 3P to 2P caused the destabilizations of stable cancer telomeres and mating telomeres for more stable normal telomeres and mixed telomeres for mixed cancer and normal telomeres and mating telomeres. This is characteristic of highly phosphorylated oligonucleotides of specific number of nucleotides dephosphorylating. The mass spectra are in Supplementaries IX and X. The telomeres at 2P phosphorylated undergo negligible changes in cancer stabilities by methylations with isotopic enrichments and slightly increased cancer stabilities for acetylations with isotopic enrichments. The mating telomeres at 2P phosphorylation undergo methylations to manifest negligible change in stability of mixed normal and cancer mating telomeres with isotopic enrichments, and the mating telomeres undergo slight increase in stabilizations of cancer telomeres at 2P with heavy isotopic enrichment by acetylations.

The dephosphorylations from 2P to 1P caused the destabilizations of stable normal for stabilizing cancer for both telomeres and mating telomeres. This is not characteristic of highly phosphorylated oligonucleotides of specific number of nucleotides dephosphorylating. There may be overlaps with nucleotides. The mass spectra are in Supplementaries X and XI. The telomeres at 1P phosphorylated undergo negligible changes in cancer stabilities by methylations and acetylations with isotopic enrichments. The mating telomeres at 1P phosphorylations undergo methylations and acetylations to manifest negligible changes in stabilities of cancer mating telomeres with isotopic enrichments.

### Mononucleotides

The fully saturated (3P) telomeric mononucleotides manifested stable mixed cancer and normal, with more intense normal telomeres with isotopic enrichments by heavy isotopes. But the mating telomeres of full phosphorylations manifest stable cancer mating telomeres with light isotopic enrichments. See Supplementary Materials I-XV. The dephosphorylations from 3P to 2P caused the changes for more stable cancer telomeres and negligible changes for more stable cancer mating telomeres. This is characteristic of highly phosphorylated oligonucleotides of specific number of nucleotides dephosphorylating. The mass spectra are in Supplementaries XII and XI. The telomeres at 3P fully phosphorylated undergo increased mixed normal and cancer stabilities by methylations with isotopic enrichments but the acetylations cause increased normal stabilities with isotopic enrichments. The mating telomeres at 3P full phosphorylations undergo methylations to manifest increased normal mating telomeres with isotopic enrichments, but the mating telomeres undergo stabilizations increased cancer mating telomeres at 3P with heavy isotopic enrichments. The telomeres at 2P undergo change in stabilities from cancer to mixed normal and cancer stabilities and isotopic enrichments for methylations and acetylations.

The dephosphorylations from 2P to 1P caused the changes for less stable cancer telomeres for mixed normal and cancer telomeres and negligible changes for more stable cancer mating telomeres. This is characteristic of highly phosphorylated oligonucleotides of specific number of nucleotides dephosphorylating. The mass spectra are in Supplementaries XI and XII. The telomeres at 1P fully phosphorylated undergo increased cancer stabilities by methylations and acetylations with isotopic enrichments. The mating telomeres at 1P full phosphorylations undergo methylations to manifest negligible increased normal mating telomeres with isotopic enrichments, but the mating telomeres undergo negligible increase stabilizations of cancer mating telomeres at 3P with heavy isotopic enrichments.

## Discussion

In this study we describe for the first time the differences in mass distribution isotopes in telomeres from cancer versus normal cells. The observed difference is about 1327 to 1328 Da mass to charge. This difference may stem from different isotopic compositions, clumping and distributions in the telomeric codons in cancer cells relative to the normal cells. Telomeres of cancer cells are enriched and clumped in the primordial isotopes (^12^C at 1327 Da) possibly due to the more dynamical chemical transformations of the clustered, clumped non-primordial isotopic (^13^C) in the cancer DNA for 1328 Da mass to charge. Since the clustered, clumped nonprimordial (^13^C) isotopes in the 1328 Da masses in cancer telomeres are more rapidly defunctionalized and degrade to smaller mass pieces of 1309 to 1313 Da, ^13^C is depleted from cancer telomeres. The observed clumped enrichments of the nonprimordial isotopes at 1311 Da in the cancer cells versus the enrichments of primordial isotopes of ^14^N and nonprimordial ^13^C and ^17^O in the white blood cells at 1311 Da are consistent with this view. The greater variations of the FWHM in the cancer telomere at 1309-1313 Da is consistent with these clustered, clumped ^13^C and ^15^N nonprimordial isotopes which induced variation in the stability of 1328 Da leading to defunctionalizations of fragments with masses of 1309-1313 Da. Normal cells having primordial ^12^C and ^14^N isotopes with more random methylations lack the ^13^C --- ^13^C in the DNA internal interactions resulting in accelerated defunctionalizations. The large −5 charge induces ^12^CH_3_ defunctionalizations in the normal cell telomeres. The defunctionalizations of the 1328 Da masses in the cancer telomere are more induced (relative to the defunctionalizations of the 1672 and 1673 Da masses) due to larger −5 charge of this codon {[G_3_(T_2_ AG_3_)_3_]^5-^} of the 1328 Da mass. Larger negative charging may be explained by the present of more ^13^CH_3_ in the telomeres as the ^13^CH_3_ has a positive nuclear magnetic moment (NMM) and therefore tends to stabilize greater negative charge fragments. The ^15^N in the cancer telomeres does not as well stabilize the negative charge as ^17^O in normal telomeres, so the ^13^C and ^15^NH_2_ in cancer telomeres are more subject to fragmentations and defunctionalizations. Based on the theory explained in ref [4,5] the larger negative charge (and indirectly the clustered ^13^CH_3_ in the cancer cells) induces higher rates of rapid defunctionalizations of the ^13^C containing telomeres relative to the ^12^CH_3_ containing telomeres.

The larger negative charge may induce a higher loss of ^13^CH_3_ from the original 1328 Da telomeric pieces to form masses of smaller range from 1309 to 1313 Da. The negative charge probably increases demethylations due to the larger negative charge, which increases the probability of rehybridizations of ^13^CH_3_ during defunctionalizations. The presence of other nonprimordial isotopes (like ^15^NH_2_) with the ^13^CH_3_ in the telomeres may increase defunctionalizations of CH_3_. The opposite is observed in the normal white blood cells as they are less able to defunctionalize since they contain less clumped nonprimodial ^13^C isotopes with random distributions in 1328 Da pieces. Thereby (after defunctionalizations) they have more residual random nonprimordials at 1328 Da but less clumped nonprimordials at the defunctionalized products with smaller masses of 1309-1313 Da.

The observed difference in mass distributions in the 1672-1673 Da mass to charge may stem from different isotopic compositions of the telomeric codon in the cancer cells relative to the normal cells. The cancer telomeres seem to be enriched in the primordial isotopes (of ^12^C at 1327 Da) possibly due to the more dynamical chemical transformations of the nonprimordial isotopic (^13^C and ^15^N) telomeres having 1328 Da mass to charge; however the clumped nonprimordial (^13^C) containing 1673 Da defunctionalize less rapidly to smaller mass pieces of 1655 - 1657 Da and the smaller pieces have heavier enrichments at 1656 Da. The normal cells manifest more random, less nonprimordial (^13^C) isotopes in their DNA and are less enriched in nonprimordial ^13^C isotopes at 1673 Da; therefore, the telomeres of normal cells have higher peak at 1672 Da of enriched primordials and normal telomeres that do defunctionalize have mass distributions of less massive primordial pieces at 1655 Da. The observed clumped enrichments of the nonprimordial ^13^C isotopes at 1656 Da in the cancer cells versus the enrichment of primordial ^12^C isotopes in the white blood cells at 1655 Da is consistent with this view. The greater variation of the FWHM in cancer telomere at 1672 - 1673 Da is consistent with this clumped, clustered, nonprimordial ^13^C isotopic essence of the cancer telomeres and the induced variations in stabilities of 1673 Da to defunctionalize fragments of masses 1655-1657 Da. These defunctionalizations of the ^13^C clustered nonprimordial 1673 Da are less induced relative to the ^13^C clustered nonprimordial 1327 Da telomeric piece due the smaller charge of −4 {on codon [G_3_(T_2_AG_3_)_3_]^4-^} of 1672-1673 Da pieces relative to the larger charge of larger −5 of 1327 – 1328 Da pieces. The larger negative charge at 1327 to 1328 Da may cause more functionalizations and defunctionalizations of ^13^CH_3_ (with clumped ^15^NH_2_) in the telomeres of mass to charge 1327 to 1328 Da relative to their defunctionalizations of the (−4) decreased charge 1672 to 1673 Da fragments. The ^13^CH_3_ has a positive nuclear magnetic moment and therefore tends to stabilize greater negative charged fragments; as a result, the larger negative charges tend to form better complexes in telomeres with more CH_3_. The resulting smaller negative charge of 1672 – 1673 Da inhibits a loss of ^13^CH_3_ (with ^15^NH_2_) from the original 1673 Da fragments thus there will be less number of smaller mass of 1655 – 1657 Da. The smaller negative charge in cancer telomeres probably decreases demethylations due to the smaller negative charge decreasing probability of rehybridizations of ^13^CH_3_ during the defunctionalizations. The opposite is observed in the normal white blood cells as they are more able to defunctionalize their random primordial ^12^C isotopes in the 1672 Da fragments as there is less ^13^C to stabilize the negative charge (as in the cancer telomeres). Thereby the normal cells have more residual ^12^C primordials at 1672 Da and more ^12^C primordials at the defunctionalized products with smaller masses of 1655 to 1657 Da.

This observed enrichments and clusterings of ^13^CH_3_ in cancer cell DNA are consistent with higher methylation rate of DNA in cancer cells [9] and more nonrandom clusterings of methyl groups in DNA of cancer cells relative to normal cells [10]. The ^13^CH_3_ may provide an explanation for the higher clustered methylations in the DNA in cancer cells as the ^13^CH_3_ may be considered a stronger nucleophile than the ^12^CH_3_ [4,5]. The positive NMM of ^13^CH_3_ relative to the null (0) NMM of the ^12^CH_3_ causes valence electrons in the ^13^C nucleophile to be pulled more strongly to nuclei to form higher rehybridizations and nucleophilicity of the ^13^CH_3_ and as a consequence a higher stabilities of the resulting R-^13^CH_3_ methylated groups relative to ^12^CH_3_ nucleophiles and methylated groups. The observed greater clustered ^13^CH_3_ distributions in the DNA of cancer cells and the observed higher stabilities of negative charge accumulations inducing greater fragmentations of the ^13^CH_3_ enriched telomeres are consistent with stronger and differing possible interactions of ^13^CH_3_ enriched telomeres in cancer cells with ^25^Mg (- NMM) relative to ^24^Mg (null (0) NMM) for - NMM. These interactions result in adverse dynamics and alterations of the bindings and replications of DNA in cancer cells relative to normal cells (having less clustered^13^CH_3_ and less negative charge). The observations of this work are further consistent with study of isotopes in twins in space and earth [7]. The authors of this study have found that the elongation of DNA of orbiting twin astronauts relative to earth bound twin as the orbiting astronaut may experience neutrons in cosmic rays that enrich methyl groups with ^13^C and increase the nucleophilic attack of the DNA by ^13^CH_3_ thus elongating their telomeres. The ^13^CH_3_ in the orbiting twin’s telomeres may alter the unraveling of the telomere during reproductions as the ^13^CH_3_ is a stronger base than ^12^CH_3_ so the ^13^CH_3_ containing telomeres may not frazzle their ends as much for increasing stabilities. The functionalizations by ^13^CH_3_ may induce more additions to the telomeres rather than their decompositions and frazzlings.

The observe differences in telomeres and mating telomeres point to different biochemistry during replications as the mating telomeres have to fragment and the telomeres remain in tact and the observations here point to how normal telomeres mutate by isotopic replacements and/or functionalizations by methylations, aminations, hydroxylations, phosphorylations and acetylations for altering the abilities for the telomere to remain intact and the mating telomeres to break up and refuse to replicate the DNA for cancer manifestations. This also points to the discovery here of the aging process in addition to the cancer as these data points to youth of an organism due to random distributions of isotopes of ^13^C, ^15^N, ^17^O, ^25^Mg, and ^33^S and with aging (and disease onset) these data and the model point to clumping of these nonprimordial isotopes of ^13^C, ^15^N, ^17^O, ^25^Mg, and ^33^S in the enzymes and nucleic acids for altering the biochemistry and biology of the host for age related tissue failure and decline in organ functions and other manifestations of aging.

Thereby the data of the telomeres and its nucleotides, dinucleotides, trinucleotides, tetranucleotides and pentanucleotides and their fragments may be reasoned by the ordered nucleotides in constant motions for fractional fissing and fusing to many disordered nanoregions and domains. The many motions of such nanodomains fragment to bigger pieces of order on macroscale. Normal oligonucleotides and nucleotides have one pattern of fractional, reversible fissing and fusing and subatomic motions, atomic motions and nanoscale motions for time crystal normal patterns. But the cancer oligonucleotides and nucleotides have different patterns as due to different NMMs and some different chemical functionalizations. Such differences in isotopic compositions, functionalizations and motions of the cancerous verses the normal telomeres and their fragments cause the different mass spectra. So by reasoning in this way the cancer and normal DNAs can be explained and the data and life processes can be explained. These particle to wave phenonema as by matter to space dynamics explain and unify transportations, transformations and transmutations on leptonic/quark scales, hadronic scales, nuclear scale, atomic scales, molecular scales, nanoscales and macroscopic scales interdependently in new ways.

Fractional, reversible fissing and fusing of electrons and nuclei (with positive and negative NMMs out of rhythm in fractionally, reversibly fissing and fusing) for novel introduction of transmutations with affecting transformations (as ionic, covalent, magnetic and metallic bonds) mix for novel chemical dynamics of life. Such novel transmutations and transformations fissing out into macroscales manifest living transport phenomena for explaining organisms in new ways as by this model [4,5]. The transformations can partially shield the nuclei and electrons and the nuclei and electrons can transmute to drive the transformations and transportations as by partially shielding so nuclei form macroscale phenomena. Thereby the dynamics manifest a perpetual, agitating in-flow and out-flow of spatial and temporal irrationals into nuclei, electrons, atoms, molecules, and nanodomains into and from macroscopic objects. By this model [4,5] the macro-objects agitate many nanodomains to many, many subatomic particles and reversibly the many subatomics particles agitate the macroscale. Unlike conventional chemical dynamics such new dynamics explain the mass spectral data and life itself as by the NMMs inducing superluminous, fractional, reversible fissing and fusing of nuclei and electrons for altering the kinetic controls on macro-phenomena to thermodynamic controls at low temperature even below room temperature as the potentials drive dynamics unlike the conventional kinetics driving of dynamics by the new phenomena of this model [4,5] in high potential fields. But due to the dense potentials, the slightest thermal agitations of space (even down to absolute zero) can agitate the potentials to huge thermals reversibly to activate dynamics.

Cancer cells by different isotopic compositions possess different internal potentials than normal cells. The normal cells have random distributions of isotopes and random consequent internal potentials of the different elements by this model [4,5] in life. The model [4,5] proposes and explains the mass spectral data as the normal cells over time under the surroundings and the internal dynamics clump the nonprimordial isotopes to alter isotopic distributions within biomolecules for altering kinetic controls to thermodynamic controls of life processes and cancer cells are one disease as the outcome. By this model [4,5] a new basis for aging may be the clumping of nonprimordial isotopes in certain positions in biological molecules over time under the influence of environmental radio frequency waves, static magnetic fields, thermal agitations, gravity, electric fields and quanta fields, strong fields and weak fields for causing biological molecules to malfunction to cause diseases as cancer to originate. In the case here the clumpings of the nonprimordial isotopes are observed upon the telomeric genes for affecting aging and diseases like cancer.

On the basis of this model [4,5], DNA is discussed specifically to explain the data. In considering DNA and its parts of genes and the parts of genes of oligonucleotides and the parts of oligonucleotides of nucleotides, such model [4,5] can be applied as the 4 types of nucleotides take on different patterns for oligonucleotides and different oligonucleotides take different patterns for genes and different patterns of genes make up DNA. There are variations of nucleotides as T, C, A, and G with each having different nitrogen contents. G and T have more ^16^O and ^17^O (oxygen), ^12^C and ^13^C (carbon) and ^14^N and ^15^N (nitrogen) are greater. During biological processes as T → A with ^15^N becoming greater for clumping; and T → C with greater ^15^N from clumping in A and C. Thereby a new basis for aggressive cancer is introduced. The biology of DNA can thereby be reasoned by this model [4,5] and changes in biology of DNA by clumping NMMs can be reasoned for normal DNA to transform to cancerous DNA by isotopic replacements and clumpings among the genes, oligonucleotides and the nucleotides. The biology of aging can also by this model [4,5] be reasoned as due to more random distributions of nonprimordial isotopes during fertilization, fetal formation and birth and development of more and more clumpings of nonprimordial isotopes into childhood, adolescence, adult and senior years to natural death. By this model [4,5] the NMMs of normal biomolecules are proposed for normal activities of enzymes thereby acting to alter the clumpings of NMMs about DNA (for instance) as the proteins have clumped NMMs as ^1^H and ^31^P to thereby alter clumping of NMMs about the DNA to induce dynamics in the DNA. The proteins and nucleic acids in normal cells are heterogeneous mixtures on nanoscales and subnanoscales and unlike conventional chemistry such can extract thermal energy from surroundings to violate the Second Law of Thermodynamics. But the genesis of cancer and clumping of NMMs causes loss of accumulating thermal energy for more dissipative of organized energy to thermal energy.

In the normal cells with random distributions of isotopes and NMMs, the disorders manifest on nanoscales; so the nanoscales in normal cells can exist in the thermal space for extracting energy from the thermal space. The DNA also can be transformed as by functionalizations having enriched isotopic contents. Phosphorylations impart ^31^P and its + NMMs; methylation imparts ^13^C and its + NMM; amination imparts ^15^N and its – NMM (and associated and hydrogen bonding, hydroxylation); hydroxylation imparts ^17^O and its – NMM (and associated hydrogen bonding); and acetylation imparts ^13^C and two ^17^O for mix its – and + NMMs (and associated hydrogen bonding). Normal and cancerous DNA would be activated enzymatically in different ways by the protein and isotopically altered clumped proteins would cause different enzymatics upon cancer DNA relative to normal DNA. These altered isotopic clumpings of proteins and DNA explain diseases like cancer and are exemplified by the mass spectral data Figures I, II and III and Supplementary Materials I-XV. The natural isotopic clumpings in normal DNA (of ^1^H, ^14^N and ^31^P) and altered clumped isotopic patterns in cancer DNA (of ^13^C, ^15^N, ^17^O, ^25^Mg, and ^33^S) further explain the mass spectroscopy as the laser activates the DNA in analog to how the enzyme in living organisms activate the DNA to fragment to replicate and transcribe. Patterns for normal DNA arise and different patterns for cancer DNA manifest in the mass spectra and in analog to patterns during enzymatic operations on normal verse cancer DNA; functionalizations alter these patterns electronically and isotopically by clumping NMMs to alter the mass spectra and to alter the functions of the cancer DNA relative to normal DNA and related enzymes to manifest cancer biology relative to normal biology. Just as proteins have clumped NMMs to induce dynamics, RF can induce intrinsic clumpings of NMMs from prior nonclumped random distributions of isotopes, other fields also can alter clumping like altered electric and magnetic and gravity fields.

The laser in mass spectrometer is an example of an electromagnetic wave in analog to enzymes providing quantum fields for alterations of clumping and inducing chemical transportations and transformations by reversible transmutations. The laser alters clumpings of nonprimordial isotopes in the DNA to alter the chemical binding to explain the fragmentations of the DNA, genes, oligonucleotide and nucleotides and explain differences between cancer and normal DNA patterns of peaks in mass spectra in Supplementary Materials I-XV. The normal DNA has clumping and dynamics due to ^1^H, ^14^N and ^31^P in the DNA and proteins of various enzymes so the laser and enzymes act on these positive NMMs to induce kinetic controlled dynamics of biomolecules in the normal cells. But as the DNA and proteins enrich and clumpings of nonprimordials like ^15^N, ^17^O and ^13^C for cancerous DNA, then the DNA is altered in patterns of reversible, fractional transmutations of clumped NMMs for the consequent altered biological dynamics in the alteration of DNA unraveling, cleaving and fusing for different DNA, RNA and protein replications, transcriptions and translations and the manifestations of diseases. These differences due to different isotopic clumpings with ^15^N, ^17^O and ^13^C in normal and cancer DNA also lead to different laser induced fragmenting of DNA of cancer relative to normal DNA as revealed by the mass spectral data in these many Suppimentary Materials I-XV. The cancer originates as the clumpings of positive and negative NMMs in cancer DNA cause shift from kinetic controls to thermodynamic control of chemical dynamics of life for death as the outcome of cancer. But the normal cells having random distributions and more positive clumped NMMS operate under kinetic control and slow the equilibration for life cycle before death.

This is so for life and it could be that certain genes are more subject to clumping isotopes having negative NMMs and the telomeres may be more subject to clumping isotopes of negative NMMs so the telomeres act as detectors for gauging and limiting DNA replications on basis of clumpings of nonprimordial isotopes of clumped NMMs and sensitizing the cell so the cell knows when to stop reproducing due to excessive clumpings of nonprimordial isotopes. Without such limitation of replications by the telomeres the other important genes of life start to excessively clump nonprimordials of ^13^C, ^17^O, ^15^N, ^25^Mg and ^33^S to alter the cellular dynamics for causing diseases of the cell as the clumpings of positive and negative NMMs occur to alter the normal dynamics to cause cancer cellular dynamics. These effects are observed in the mass spectral data. The telomeres are high G and T contents and the resulting many oxygens cause large bond dipoles and H bonding resonances for inducing bond breakings. The A in G_3_AT_2_ gives some ^15^N, so the telomeres sense when ^15^N is getting into DNA so it signals the breakages and terminations of replications and cell life. But the oxygens in T cause more ^17^O replacements and these ^17^O(s) are seed for ^13^C and ^15^N substitutions. Theses replacements cause different oligonucleotides stabilities for observed mononucleotides, dinucleotides, trinucleotides, tetranucleotides and pentanucleotides as observed in Supplementary Materials I-XV. The replaced ^17^O in phosphates destabilize ATP and methylations by ^13^C with ^17^O stabilizing acetyls in acetates. There are more ^17^O in G and T nucleotides. ^17^O stabilizes as causing lack of GC as GC is less stable. Replacing ^12^C by ^13^C causes more stabilities by + charge and + NMMs. Telomeres are lacking CG and are more stable by lacking GC. CG changes to stabilize. So it is missing in telomeres so telomeres can be unstable to unravel and to alarm the cell to apoptosis after so many normal replications.

These effects of NMMs on the genes are in analog to effects of NMMs in normal cellular dynamics like those of ^31^P driving ATP/ADP/AMP dynamics for accumulating and storing chemical energy. These functionalizations of DNA and RNA and proteins by methylations, aminations, hydroxylations, acetylations and phosphorylations (and the reverse defunctionalizations) proposed in this paper occur in analog to ATP/ADP/AMP transformations and in this model it is suggested a new mechanism of ATP to ADP to AMP as by the NMMs of ^1^H, ^14^N and ^31^P as the ^31^P has positive nuclear magnetic moment (NMMs); ^14^N has positive NMM and ^1^H has positive NMMs for fractional reversible fissing fusing to drive and catalyze the chemical bond breakages and bond formations during ATP to ADP to AMP and vice versa. So ^31^P catalyzes formation ATP, but ^17^O or ^15^N may destabilize such catalysis of ^31^P in AMP/ADP/ATP to inter-transform these phosphates. So ^14^N may in enzymes form ATP but ^15^Mg and/or ^17^O may induce ATP breakage to release energy. So it is that nuclei of nonzero NMMs like ^31^P can break the second law of thermodynamics by transient hidden fractional, reversible fissing and fusing to cause some atoms to accumulate energy spontaneously just as they transmute to space momentarily to accumulate surrounding thermal energy of space. Just as ^1^H, ^14^N, and ^31^P by there intrinsic nonzero-NMMs drive phosphorylational processes of ATP to ADP to AMP, in this work other biochemistry of functionalizations of methylations, mainations, hyodroxylations, acetylations and as well phosphorylations are proposed and observed to be driven and altered by uncommon nonprimordial isotopes of ^13^C, ^15^N, ^17^O, ^25^Mg, ^33^S and isotopes of some cationic essential minerals. Therefore, by such transient reversible transmutations of positive to negative NMMs, the magnetic fields → electric fields for dramatic difference between ^14^N and ^15^N isotopes in these biomolecules and consequent altered biochemical dynamics as they clump.

The variations of these NMMs in T, C, A, and G having different nitrogen contents cause variations in the functions of nucleotides, oligonucleotides, genes and DNA. For instance, G and T nucleotides have more O for possibly more ^17^O and possibly more ^13^C and ^15^N; as T → A with clumping of ^15^N there are greater changes in bio-dynamics and as T → C with more clumped ^15^N the biodynamics can change severely for explaining altered biodynamics for more aggressive cancers. So just as phosphorus drives ATP biochemistry in the DNA, by clumping ^31^P, so also can phosphates drive methylations, aminations, hydroxylations and acetylations in DNA can differently drive and altered biodynamics (for ^12^C, ^14^N, ^16^O and ^26^Mg relative to ^13^C, ^15^N, ^17^O and ^25^Mg)) for normal DNA to become cancerous DNA. For instance, as phosphate chains shorten the basicities decrease as the tri-base units with reduced numbers; such tri-bases provide more proton orbitals to couple and modulate more ^31^P centers for altered biodynamics. The proton and proton orbitals by this model have NMMs as by ^1^H and no shielding core electrons for strongest effects of the fractional fissing and fusing of the bare proton upon the electron clouds for the novel magnetic phenomena discovered in this work. Other nuclei as by this model [4,5] also act as the proton of 1H but the nucleonic orbitals and NMMs upon the electronic clouds are weaker as the core electrons shield the fractional fissed fields of the nuclei of atomic numbers >1. But ^13^C atoms cannot be as stimulated by such protons as they have no lone electrons unlike the O in PO_3_ ^3-^. The CH_3_-C(O)(O)^-^ can be stimulated by many protons or 6 protons as three protons per oxygen. NH_2_ can be stimulated by 1 proton, OH can be stimulated by 2 protons. The surrounding acidity can induce chemistry by the protons stimulating the lone pairs as by this theory [4,5]. Such is a basis for phosphate stimulating novel biodynamics as it has 9 lone pairs and can be stimulated by 9 protons so the phosphate can store and release energy via the proton stimulating these lone pairs: lone pairs of the three oxygens on ^31^PO_3_ ^3-^. The proton stimulations also help ^17^OH replace ^16^O. Longer phos chain and more proton stimulations stabilize the nucleotide linkages as the bonds break and reform under protons fissing and fussing. The proton and positive NMM fiss to pull in electrons and this stabilize the PO_3_^3-^ to stabilize the nucleotide linkages. ^17^O and its negative NMM would destabilize the linkages. These effects of reduce phosphate numbers in oligonucleotides explain the patterns of the mass spectra of the Supplementary Materials I-XV.

## Materials and Method

### Cell growth

The experimental system consisted of three different cell types: total white blood cells, K562 (chronic myeloid leukemia) and SKW6.4 (B lymphocyte cell line). Total white blood cells were isolated by lysis of the red blood erythrocyte by using the Red Blood Cells lysis solution (Biological Industries, Israel) according to the provided manual. Briefly, cells were mixed with the lysis solution, agitated for 10 minutes and centrifuged for 2 minutes at 3125 RPM. The supernatant was then discarded. The lysis step was repeated twice and the pellet was used for DNA isolation. K562 cells were cultured in the presence of RPMI-1640 growth medium containing 10% FCS supplemented with 2mM L-Glutamine, 100units/ml penicillin and 100µg/ml streptomycin (Biological Industries Beit Haemek, Israel). Cells were grown in a 95% humidity incubator with 5% CO_2_. SKW6.4 cells were cultured in the presence of RPMI-1640, 10% FCS, 2mM L-Glutamine and 10mM HEPES, 100units/ml penicillin and 100µg/ml streptomycin (Biological Industries Beit Haemek, Israel). Cells were grown in a 95% humidity incubator with 5% CO_2_.

### DNA isolation

Cells were harvested and DNA was isolated by using the QIAamp DNA Mini Kit (Qiagen, MD, USA) according to the manufacturer’s instructions. Basically, samples were first lysed using proteinase K. The lysate in buffering conditions was loaded onto the mini spin column. During centrifugation, DNA was selectively bound to the column membrane. The remaining contaminants and enzyme inhibitors were removed in two wash steps and the NA was then eluted in TE buffer.

### MALDI

DNA samples of K562, SKW, and WBC were analyzed. 20 µL of water were added to each sample tube to dissolve the dried samples. Sample water solution was mixed with THAP (2′,4′,6′-Trihydroxyacetophenone monohydrate) matrix (saturated in 25 mM ammonium citrate in ACN/Water 50/50) at 1:3 (v/v) sample to matrix ratio. The mass measurements were performed on a Bruker Rapiflex MALDI TOF/TOF mass spectrometer (Bruker Daltonics, U.S.A.) in positive linear mode and with a 355 nm Bruker scanning smartbeam™ 3D laser.

## Conclusion

The isolation of DNA from different normal cells and many cancer cells of different type results in different patterns of decompositions of the cancer DNA and normal DNA as they are ablated in the laser of a mass spectrometer. Many patterns of the ablated pieces from cancer DNA are similar; many patterns from the ablated normal DNA are similar; and ablated patterns from normal and cancer DNA are different in functionalizations, intensities, full width half maxima and isotopic contents. These differences in the patterns of the ablated pieces from the cancer and normal DNA are explained on the basis of isotopic differences of ^13^C/^12^C, ^2^D/^1^H,^15^N/^14^N,^17^O/^16^O, ^25^Mg/^24^Mg, and ^33^S/^32^S. On the basis of such model [4,5] the different, nonzero nuclear magnetic moments of these nonprimordial uncommon stable isotopes (^2^D, ^13^C, ^15^N, ^17^O, ^25^Mg and ^33^S) relative to the primordial common stable isotopes (^1^H, ^12^C, ^14^N, ^16^O, ^24^Mg and ^32^S) cause difference internal multi-body chemical dynamics and enzymatics for forming DNA and this ablation of the DNA and the ocogenesis by the isotopic functionalizations of the proteins and nucleic acids. Similar differences in enzymatics of replications, transcriptions, translations and other biological biochemistry are suggested for causing disease and aging.

## Supporting information

Supplementary Materials III

Supplementary Materials VII

Supplementary Materials VIII

Supplementary Materials XII

Supplementary Materials XIV

## Acknowledgement

This research is made possible by Stillman College, University of Alabama and Tel-Aviv University. The mass analyses are provided by Mass Spectrometry Facility with help of Dr. Qiaoli Liang and director Professor Dr. Carolyn J. Cassady.

## Supplimentary Figures

**Supplimentary Materials I –** Tables of Mass Peaks and Nucleotide Assignments

**Supplimentary Materials II -** Mass Spectra from 1800 Da to 2000 Da

**Supplimentary Materials III -** Mass Spectra from 1700 Da – 1800 Da

**Supplimentary Materials IV -** Mass Spectra from 1600 Da −1700 Da

**Supplimentary Materials V -** Mass Spectra from 1400 Da −1600 Da

**Supplimentary Materials VI -** Mass Spectra from 1300 Da −1400 Da

**Supplimentary Materials VII -** Mass Spectra from 1200 Da – 1300 Da

**Supplimentary Materials VIII –** Mass Spectra from 1000 Da – 1200 Da

**Supplimentary Materials IX -** Mass Spectra from 800 Da −1000 Da

**Supplimentary Materials X -** Mass Spectra from 600 Da −800 Da

**Supplimentary Materials XI -** Mass Spectra from 400 Da −600 Da

**Supplimentary Materials XII -** Mass Spectra from 100 Da - 400 Da

**Supplimentary Materials XIV -** Mass Spectra from 2000 Da - 2300 D

**Supplimentary Materials XV -** Mass Spectra from 2300 Da - 2600 Da

