## Supplementary Materials I for "Oncogenesis and Aging by Isotopic Functionalizations of the Proteins and Nucleic Acids"

Table V
PentaNucleotides

| | | |
|---|---|---|
| | 4-phos | methyl |
| GGGAT | N1627GGGAT4p | C1641GGGAT4pMet |
| GGATT | Missing | C1613GGATT4pMet |
| TTGGG | C1613TTGGG4p | N1627TTGGG4pMet |
| CCCTA | N1487CCCTA4p | N1505cccta4pMet |
| CCTAC | C1513CCTAC4pH* | N1527CCTAC4pMet |
| AACCC | C1500AACCC4pL* | C1513AACCC5pMetH* |
| | 6-phos | methyl |
| GGGAT | C1782GGGAT6p | N1798GGGAT6pMet |
| GGATT | C1754GGATT6pH* | C1772GGATT6pMet |
| TTGGG | C1772TTGGG6pH* | C1791TTGGG6pMet |
| CCCTA | C1643CCCTA6pH* | C1658CCCTA6pMetH* |
| CCTAC | C1672CCTAC6pH* | C1682CCTAC6pMetH* |
| AACCC | C1656AACCC6pH* | C1672AACCC6pMetH* |
| | 8-phos | methyl |
| GGGAT | N1941GGGAT8p | N1957GGGAT8pMet |
| GGATT | N1914GGATT8P | NC1933GGATT8pMet |
| TTGGG | NC1929TTGGG8p | N1945TTGGG8pMet |
| CCCTA | CN1800CCCTA8p | CN1815CCCTA8pMet |
| CCTAC | CN1827CCTAC8p | CN1843CCTAC8pMet |
| AACCC | N1807AACCC8p | CN1827AACCC8pMet |
| | 10-phos | methyl |
| GGGAT | C2095GGGAT10P | Missing |

| | | |
|---|---|---|
| GGATT | Missing | C2084GGATT10pMet |
| TTGGG | C2084TTGGG10P | Missing |
| CCCTA | N1957CCCTA10P | N1975CCCTA 10pMet |
| CCTAC | N1975CCTAC10P | C1994CCTAC10pMet |
| AACCC | N1964AACCC10P | N1979AACCC10pMet |
| | 12phos | methyl |
| GGGAT | C2253gggat12P | c2266gggat12PMet |
| GGATT | C2228GGATT12P***L*** | c2240ggatt12P |
| TTGGG | c2240ttggg12P | C2254TTGGG12PMet |
| CCCTA | C2115CCCTA12P | Missing |
| CCTAC | Missing | c2150cctac12pMet |
| AACCC | C2120AACC12P | Missing |
| | 14-phos | methyl |
| GGGAT | c2407gggat14P***H*** | CN2419GGGAT14P***H*** |
| GGATT | C2383GGATT14p | C2395GGATT14pMet***H*** |
| TTGGG | C2395ttggg14p***H*** | C2413ttggg14pMet |
| CCCTA | C2267CCCTA14P | C2282cccta14pMet |
| CCTAC | c2292aatac14P | Missing |
| AACCC | Missing | c2291aaccc14pMet |

Table IV
TetraNucleotides

| | | |
|---|---|---|
| | 3-Phos | Methyl |
| GGGA | C1289GGGA3P | |
| GGGT | C1279GGGT3P | C1296gggt3pmet |

| | | |
|---|---|---|
| GGAT | N1262GGAT3P | c1279ggat3pmet |
| GGTT | C1255GGTT3P | N1270GGTT3pmet |
| GATT | C1233GATT3P | C1255gatt3pmet |
| TCCC | 1155TCCC3P | |
| ATCC | 1181ATCC3P | |
| ACCC | | |
| AACC | C1189aacc3p | N1203AACC3pMet |
| AATC | N1203AATC3P | C1222aatc3pmet |
| | 5=Phos | Methyl |
| GGGA | N1443ggga5p***H*** | c!1456ggga5pmet***H*** |
| GGGT | C!1434gggt5p | c1447gggt5pmet***H*** |
| GGAT | C!1418ggat5p* | c!1433ggat5pmet***H*** |
| GGTT | N1408ggtt5p* | c!1424ggtt5pmet***H*** |
| GATT | N1395gatt5p | C1408gatt5pmetH* |
| TCCC | c1316tccc5p | c1328tccc5pmet* |
| ATCC | c1338atcc5p | n1352atcc5pmet* |
| ACCC | c1323accc5p | C1334accc5pmet |
| AACC | n1346aacc5p* | c1361aacc5pmet* |
| AATC | c1361aatc5p | c1377aatc5pmet* |
| | 7-Phos | Methyl |
| GGGA | N1596ggga7p | c!1613ggga7pmet |
| GGGT | n1586gggt7p | n1608gggt7pmet |
| GGAT | c1577ggat7p | n$1586ggat7pmet |
| GGTT | n$1566ggtt7p | n$1583ggtt7pmet |
| GATT | n1551gatt7p | n$1564gatt7pmet |
| TCCC | c1469tccc7p | n!1484tccc7pmet |
| ATCC | c!1491atcc7p | n!1506atcc7pmet |
| ACCC | c!1477accc7p | C!1491accc7pmet |
| AACC | n!1506aacc7p | n!1521aacg7pmet |
| AATC | n!1521aatc7p | n$1529aatc7pmet |
| | 9-Phos | Methyl |
| GGGA | c!1757ggga9p***H*** | c!1770ggga9pmet***H*** |
| GGGT | c!1747gggt9p***H*** | n1761gggt9pmet* |
| GGAT | c1732ggat9p***H*** | c!1747ggat9pmet***H*** |

| | | | |
|---|---|---|---|
| GGTT | n1721ggtt9p* | c!1735ggtt9pmet***H*** | |
| GATT | c!1704gatt9p***H*** | n1722gatt9pmet* | |
| TCCC | n1624tccc9p* | c1640tccc9pmet | |
| ATCC | n1649atcc9p | c1662atcc9pmet | |
| ACCC | C!1637accc9p | n1649accc9pmet | |
| AACC | C!1659aacc9p* | c!1673aacc9pmet | |
| AATC | c!1673aatc9p | c1685aatc9pmet | |
| | 11-Phos | Methyl | |
| GGGA | n1914ggga11p | c1927ggga11pmet | |
| GGGT | c1901gggt11p | c1917gggt11pmet | |
| GGAT | n1888ggat11p* | c1901ggat11pmet | |
| GGTT | c!1879ggtt11p | c1894ggtt11pmet | |
| GATT | n1864gatt11p* | c!1879gatt11pmet | |
| | | 3 | 0 |
| TCCC | n1782tccc11p | n1797tccc11pmet | |
| ATCC | c1805atcc11p | c1821atcc11pmet | |
| ACCC | c!1791accc11p***H*** | c1805accc11pmet | |
| AACC | c!1814aacc11p | n1831aacc11pmet | |
| AATC | n1831aatc11p* | n1842aatc11pmet* | |

Table III
TriNucleotides

| | | |
|---|---|---|
| | | 2 Methyl |
| GGG | C!957ggg2p | N$967ggg2pmetHN |
| TGG | C!928tgg2pH | N$946tgg2pmetN |
| TGT | N$907tgt2pH | N$924tgt2pmetH |
| TAG | C916tag2p | C!928tag2pmetH |
| TAT | N$885tat2pN | N$907tat2pmetH |
| AGG | C!940agg2pHN | C!954agg2pmet |
| TAA | C!897taa2p | C!911taa2pmet |
| TCC | C!852tcc2pHN | C864tcc2pmetHN |
| TCA | C!875tca2pH | C893tca2pmet |

| | | |
|---|---|---|
| CAA | N$852pN | C!897caa2pmet |
| CCC | C!840ccc2p | C!852ccc2pmetHN |
| CCA | ~~~C877 | C!875cca2pmetH |
| | | 4 Methyl |
| GGG | n1112ggg4pL | N1127ggg4pmet |
| TGG | n1089tgg4pH | C1100tgg4pmet |
| TGT | ~~~~~ | C!1075tgt4pmetL |
| TAG | c1075tag4pL | N1089tag4pmetH |
| TAT | n1046tat4p | ~~~~~ |
| AGG | C!1098agg4p | N$1112agg4pmetL |
| | Below 4 then dinucleotides contribute pieces | |
| TAA | C!1054taa (CAA) 4pL | ~~~~~ |
| TCC | C!1009tcc4pL | n1023tcc4pmetH |
| TCA | L | n1047tca4pmet |
| CAA | ~~~~~~ 1057 | C!1054caa4pmet L |
| CCC | 993 | C!1009ccc4pmetL |
| CCA | C!1017cca4pH | C!1032cca4pmetL |
| | | 6 Methyl |
| GGG | N$1269ggg6pN | n1285ggg6pmet |
| TGG | C1244tgg6p* | n1259tgg6pmetL |
| TGT | N!1222tgt6pN | c!1234tgt6pmetL |
| TAG | C1229tag6p*N | N$1246tag6pmetN |
| TAT | N$1204tat6pN | N!1222tat6pmetN |
| AGG | C!1254agg6pL | N$1269agg6pmetN |
| TAA | C!1212taa6p | n1229taa6pmetN |
| TCC | c1164tcc6p* | c1179tcc6pmet* |
| TCA | c1189tca6p* | N$1204tca6pmetN |
| CAA | n1195caa6p* | C!1212caa6pmet |
| CCC | N1149ccc6p*H | C1164ccc6pmet* |
| CCA | N1173cca6p* | C1189cca6pmet |

| | | 8 Methyl |
|---|---|---|
| GGG | C!1425ggg8p***H*** | N$1440ggg8pmet |
| TGG | N$1400TGG8pN | C1415tgg8pmetH |
| TGT | C!1377TGT8p*N | C!1389tgt8pmet*L |
| TAG | C!1389TAG8p*L | C1399tag8pmet |
| TAT | C!1360TAT8p*H | c!1377tat8pmet* |
| AGG | C!1409agg8p***H*** | C!1424agg8pmet*H* |
| TAA | C!1371TAA8p*N | C!1382taa8pmet*H |
| TCC | C!1324TCC8p*H | C!1333tcc8pmet |
| TCA | C!1345tca8p*H | C!1360tca8pmetH |
| CAA | c!1356caa8p* | C!1371caa8pmet*N |
| CCC | C!1311ccc8p*H | C!1324ccc8pmet*H |
| CCA | C!1333cca8p* | C!1345cca8pmet*H |

Table II
Di Nucleotides

| | 1-Phos | Methyl |
|---|---|---|
| TT | c559tt1p | C!573tt1pmetH |
| AG | C!595ag1pL | C610ag1pmetH |
| GG | C!617gg1pH | N$629gg1pmet |
| TA | C!573ta1pH | c586ta1pmetH |
| TG | c587tg1p | C603tg1pmetH |
| CC | C532cc1p | C!551cc1pmetH |
| CA | C!559ca1pH | C!573ca1pmetH |
| AA | C580aa1p | C!595aa1pmetL |

| | | |
|---|---|---|
| TC | C!551tc1pH | C559tc1pmet |
| TA | C!573ta1pH | c586ta1pmetH |
| | 3-Phos | Methyl |
| TT | C720tt3pL!!! | C!739tt3pmet |
| AG | C!757ag3pH | C!771ag3pmetH |
| GG | C!771gg3pH | C!784gg3pmetH |
| TA | C!726ta3pH | C742ta3pmet |
| TG | C743tg3pH | C753TG3pmetH |
| | Cancerous | |
| CC | C!688cc3pH | C705cc3pmetL |
| CA | C!715ca3pLH!!! | C!726ca3pmetH |
| AA | C!739aa3pH | N753aa3pmetH |
| TC | C705tc3pL | C!718tc3pmetHL!!! |
| TA | C!726ta3pH | C742ta3pmet |
| | 5-Phos | Methyl |
| TT | C!875tt5pH | N$885tt5pmetN |
| AG | N$907ag5pH | N$924ag5pmetH |
| GG | N$924gg5pH | c940gg5pmetH |
| TA | n$885ta5pN | C!897ta5pmet |
| TG | C!897tg5p | N$907gt5pmetH |
| | Cancerous | |
| CC | C844cc5pH | c860cc5pmetH |
| CA | c868ca5p | N$885ca5pmetN |
| AA | C!897aa5p | N$907aa5pmetH |
| TC | c859tc5p | c!875tc5pmetH |
| TA | n$885ta5pN | C!897ta5pmet |

Table I

| Nucleotides | Monophos | |
|---|---|---|
| | | met |
| G | C270G | C349g1PH |
| A | N253A | C326a1pH |
| T | C243T | C!315t1p |
| C | C227C | C304c1pL |
| | Triphos | |
| | | met |
| G | C500g3pL | c517g3pmetL |
| A | N$489a3pNL T3pmet | c500a3pmet |
| T | N$473t3pN | N$494t3pmetN |
| C | C459c3pN | n$474c3pmetN |

| | | |
|---|---|---|
| acetyl | 5-phos | methyl |
| C1688GGGAT4pAcetH* | C1704GGGAT5PH* | C1723GGGAT5pMetH* |
| C1658GGATT4pAcetH* | C1679GGATT5pH* | C1695GGATT5pMet |
| C1679TTGGG4pAcetH* | C1695TTGGG5pH* | C1709TTGGG5pMetH* |
| N1549CCCTA4pAcet | C1569CCCTA5p | Missing |
| C1569CCTAC4pAcet | C1592CCTAC5p | Missing |
| Missing | Missing | C1591AACCC5pMet |
| acetyl | 7-phos | methyl |
| C1841GGGAT6pAcet | C1857GGGAT7P***H*** | C1879GGGAT7pMet |
| C1820GGATT6pAcet | C1835GGATT7p***H*** | C1850GGATT7pMet |
| C1835TTGGG6pAcet***H*** | C1850TTGGG7p | Missing |
| C1705CCCTA6pAcetH* | C1723CCCTA7pH* | C1735CCCTA7pMetH* |
| C1726CCTAC6pAcetH* | C1747CCTAC7pH* | C1757CCTAC7pMetH* |
| C1711AACCC6pAcetH* | C1735AACC7pH* | C1747AACCC7pMetH* |
| acetyl | 9-phos | methyl |
| C!2002GGGAT8pacet***L*** | C!2018GGGAT9P***L*** | C2030GGGAT9pMet***H*** |
| N1975GGATT8pAcet | C1993GGATT9p | C2008ggatt9pmet***H*** |
| C1986TTGGG8pAcet | C2008TTGGG9p***H*** | C2024TTGGG9pMet***L*** |
| CN1857CCCTA8pAcet | N1880CCCTA9p | CN1895CCTA9pMet |
| CN1882CCTAC8pAcet | N1901CCTAC9p | N1915CCTAC9pMet |
| Missing | Missing | N1901AACCC9pMet |
| acetyl | 11phos | methyl |
| C2152gggat10pAcet | c2172gggat11P | c2190GGGAT11PMet |

| | | |
|---|---|---|
| Missing | c2152ggatt11P | C2160ggatt11pMet |
| Missing | C2160ttggg11p | C2180ttggg11pMet |
| C2018CCCTA10Pacet***L*** | C2030CCCTA11P***H*** | C2045CCCTA11Pmet***L*** |
| C2040CCTAC10PAcet | C2055CCTAC11P | Missing |
| C2024AACCC10PAcet***L*** | C2045AACCC11P***L*** | Missing |
| acetyl | 13-phos | methyl |
| Missing | C2331GGGT13P***H**** | C2345GGGAT13pMet***H**** |
| c2282ggatt12pAect | Missing | N2322GGATT13PmetH |
| cN2300ttggg12pacet | N2322ttggg13P | C2331ttggg13pMet***H**** |
| C2172cccta12pAcet***L*** | C2190cccta13P***L*** | C2204CCCTA13PMet |
| C2194cctac12Pacet***L*** | C2214aaTAC13P***L*** | C2228aatac13PMet***L*** |
| C2183aaccc1pAcet***L*** | C2200aaccc13p | C2216aaccc13pMet |
| acetyl | 15-phos | methyl |
| N2468GGGAT14pAcet***L*** | N2488GGGAT15p | C2495GGGAT15pmet***L**** |
| C2438GGATT14pAcet | N2454GGATT15p***HL**** | CN2472GGATT15pMet***H**** |
| N2453ttggg14pAcet***HL**** | CN2472ttggg15p***L**** | N2488ttggg15Pmet |
| CN2330CCCTA14pAcet**L*** | C2346CCCTA15pL | C2363cccta15pMet |
| C2352aatac14pAcet***H*** | C2373aatac15p***H*** | C2380AATAC15pMet |
| C2335AACC14pAcet***L*** | C2356aaccc15P***H*** | C2373aaccc15PMet***H*** |

| | | |
|---|---|---|
| Acetyl | 4-Phos | Methyl |
| | c1368ggga4p* | C1377ggga4pmet |
| | c1354gggt4p* | N1369GGGT4pmet |

| | | |
|---|---|---|
| | c1341ggat4p | C1354ggat4pmet |
| c1296ggtt3pacet | C1335ggtt4p* | C1346ggtt4pmet* |
| c1279gatt3pacet | c1316gatt4p | N1330GATT4pmet* |
| N1203tccc3pacet | c1234tccc4p* | N1248TCCC4pmet |
| n1225atcc3pacet | n1263atcc4p* | C1277atcc4pmet |
| c1211accc3pacet | C1243accc4p* | C1255accc4pmet |
| c1234aacc3pacet | n1269aacc4p* | C1289aacc4pmet |
| N1248aatc3pacet* | C1289aatc4p | C1299aatc4pmet* |
| Acetyl | 6-Phos | Methyl |
| n$1483ggga5pacet | n$1521ggga6p | n1537ggga6pmet |
| c!1477gggt5pacet | n$1506gggt6p | n$1527gggt6pmet |
| N!1461ggat5pacet | c!1499ggat6p | n$1506ggat6pmet |
| c!1457ggtt5pacet | n$1483ggtt6p | n$1504ggtt6pmet |
| c!1436gatt5pacet* | c1469gatt6p | n$1483gatt6pmet |
| c!1355tccc5pacet* | n1395tccc6p* | n1408tccc6pmet |
| c1381atcc5pacet* | c!1419atcc6p | n1429atcc6pmet |
| c1368accc5pacet* | n$1401accc6p | C!1418accc6pmet* |
| n1390aacc5pacet* | c1424aacc6p | n1442aacc6pmet |
| c!1405aatc5pacet* | n$1439aatc6p | c!1456aatc6pmet |
| Acetyl | 8-Phos | Methyl |
| c1640ggga7pacet | c!1678ggga8p | c!1696ggga8pmet |
| C!1637gggt7pacet | c!1671gggt8p | C!1681gggt8pmet* |
| c!1613ggat7pacet | c!1656ggat8p | c!1671ggat8pmet |
| n1608ggtt7pacet | c1640gaac8p | C!1659ggtt8pmet* |
| n1596gatt7pacet | n1627gatt8p* | c1642gatt8pmet |
| Normal | | 0/5 |
| c1513tccc7pacet | n1551tccc8p | n$1564tccc8pmet |
| n1538atcc7pacet | c1569atcc8p | n$1586atcc8pmet |
| n$1524accc7pacet | n1551accc8p | c1569accc8pmet |
| n1542aacc7pacet | c1580aacc8p | n1596aacc8pmet |
| n$1564aatc7pacet | n1596aatc8p | C!1613aatc8pmet* |
| Acetyl | 10-Phos | Methyl |
| n1799ggga9pacet* | C1833ggga10p***H*** | c1849ggga10pmet* |
| n1785gggt9pacet* | n1824gggt10p | C!1839gggt10pmet |
| c!1772ggat9pacet***H*** | c1808ggat10p***H*** | c1821ggat10pmet |

| | | |
|---|---|---|
| n1765ggtt9pacet* | n1796ggtt10p* | c!1814ggtt10pmet***H*** |
| c!1747gatt9pacet***H*** | n1783gatt10p* | n1799gatt10p*met |
| 3 | 4 | 2 |
| c!1673tccc9pacet | c!1704tccc10p***H*** | n1722tccc10p*met |
| c!1694atcc9pacet | c1726atcc10p***H*** | n1743atcc10p*met |
| c!1677accc9pacet | c!1711accc10p***H*** | n1725accc10p*met |
| c!1701aacc9pacet***H*** | c!1735aacc10p***H*** | n1752aacc10p*met |
| c!1717aatc9pacet***H*** | n1752aatc10p* | c1766aatc10p*met |

| Acetyl | 12-Phos | Methyl |
|---|---|---|
| n1954ggga11pacet* | c!1993ggga12P | ???? |
| n1945gggt11pacet* | c!1985gggt12p | c1999gggt12pmet |
| c1931ggat11pacet | c1865ggat12p | c1982ggat12pmet |
| c1921ggtt11pacet | c1959ggtt12p***H*** | c1970ggtt12pmet |
| c1904gatt11pacet | c1938gatt12p | c1959gatt12pmet |
| 2 | 0 | 0 |
| c1821tccc11pacet | n1862tccc12p* | c!1879tccc12pmet |
| c1849atcc11pacet | n1885atcc12p* | c1898atcc12pmet |
| n1833accc11pacet* | n1868accc12p* | n1885accc12pmet* |
| c!1857aacc11pacet | c1894aacc12p | n1908aacc12pmet* |
| c1873aatc11pacet | n1908aatc12p* | c1927aatc12pmet |

| Acetyl | | 3 Methyl |
|---|---|---|
| C!1004ggg2pacetH | C!1031GGG3pL | C!1050ggg3pmetL |
| c!978tgg2pacet | N$1010tgg3pL | N$1023tgg3pmetH |
| C!954tgt2pacet | C!982tgt3pH | C!999tgt3pmetLH |
| C!954tag2pacet | C!997tag3pLH | C!1009tag3pmetL |
| C935tat2pacet | N$967tat3pHN | c!984tat3pmetHN |
| C!982agg2pacetH | C!1021agg3pH | C!1031agg3pmetL |
| N$946taa2pacetN | C!978taa3p | C!993taa3pmet |
| C!897tcc2ptcc | C!928tcc3pH | N$946tcc3pmetN |
| couple to C935 | C!954tca3p | N$967tca3pmetHN |

| | | |
|---|---|---|
| C!928caa2pacetH | n$967caa3pHN | C!978caa3pmet |
| N$885ccc2pacetN | C!911ccc3pH | C!928ccc3pmetH |
| N$907cca2pacetH | C!940cca3pH | C!954cca3pmet |
| Acetyl | | 5 Methyl |
| n1155ggg4pacetH | c1191ggg5p | n1204ggg5pmetN |
| N$1133tgg4pacetL | n1166tgg5p | c1181tgg5pmet |
| n1105tgt4pacet | n1143tgt5pL | n1155tgt5pmetH |
| C!1119tag4pacetLH | n1161tag5p | n1169tag5pmet |
| n$1089tat4pacetH | n1127tat5p | n1143tat5pmetL |
| N$1143agg4pacetL | n1176agg5p | C1190agg5pmet |
| C!1098taa4pacetL | n1133taa5pL | n1149taa5pmetH |
| C!1054tcc4pacetL | n1089tcc5p | n1103tcc5pmet |
| c!1075tca4pacetL | n1112tca5pL | n1127tca5pmet |
| N$1089caa4pacetH | c1119caa5pLH | n1134caa5pmetL |
| c1036ccc4pacet* | c1069ccc5pH | c1086ccc5pmetH |
| n1061cca4pacet* | c1097cca5pL | n1110cca5pmetL |
| Acetyl | | 7 Methyl |
| c!1312ggg6pacetH | C1350GGG7pH | C1365GGG7pmet*N |
| c!1290tgg6pAcetL | C!1324TGG7pH | C1343TGG7pmet*H |
| c1261tgt6pacet | c1299tgt7p | C!1312TGT7pmet* |
| n1271tag6pscetN | N$1303TAG7p | C!1324TAG7pmet*H |
| n1246tat6pacetN | n1280tat7p | c1296tat7pmet |
| c1294agg6pacet* | C!1333agg7p | C!1350agg7pmet*H |
| c!1254taa6pacetL | c1290taa7pL | C1311TAA7pmet*H |
| N$1204tcc6pacetN | n$1246tcc7pN | C!1257tcc7pmetL |
| c1234tca6pacetL | n$1269tca7pN | C1281tca7pmet |
| c1241caa6pacet | C!1276caa7p | C!1290caa7pmet |
| c1191ccc6pacet* | C!1324CCC7p | C1242CCC7pmet |
| C!1212acc6pacet | C!1254cca7pL | C!1266acc7pmet |

| Acetyl | | 9 Methyl |
|---|---|---|
| C!1468ggg8pacet***H*** | N$1505ggg9pN | c!1521ggg9pmetH |
| C!1447tgg8pacet | C!1477tgg9pL | c1496tgg9pmetL |
| C1418tgt8pacetH | C!1456tgt9pL | c!1468tgt9pmet |
| C!1427tag8pacet | N$1461tag9pN | C!1477tag9pmetL |
| C!1404tat8pacet | N$1439tat9p | c1456tat9pmetL |
| C!1456agg8pacetL | N$1486agg9pN | N$1502agg9pmetN |
| C!1411taa8pacetL | C!1447taa9p | N$1461TAA9pmetN |
| C!1365TCC8pacet | N$1401tcc9p | C1413tcc9pmetLH |
| C!1389tca8pacetL | C!1426tca9p | n$1439tcg9pmet |
| N$1392caa8pacetL | C!1434caa9pL | c!1447caa9pmet |
| c1350ccc8pacet*H | C!1389CCC9pL | N$1401ccc9pmetN |
| C!1371cca8pacetN | C!1411cca9p | C!1426cca9pmet |

| Acetyl | 2-Phos | Methyl |
|---|---|---|
| c603tt1pacetH | C!640tt2pH | C!654tt2pmetH |
| C!639ag1pacetH | ***C!673 ag2p H*** | N$688ag2pmetN |
| C!654gg1pacetH | N$689gg2pN | N$710gg2pmetN |
| C617ta1pacetH | N!649ta2pN | n669ta2pmetN |
| C630tg1pacetH | N!665tg2pNL | C!682tg2pmetH |
| C573cc1pacetH | C610cc2pH | N$625cc2pmetN |
| C!595ca1pacetL | C!636ca2pH | N$649ca2pmetN |
| C624aa1pacetH | C!658aa2pH | ***C!673 aa2pmet H*** |

| | | |
|---|---|---|
| C!590tc1pacetL | N$629tc2pN | C!640tc2pmetH |
| C617ta1pacetH | N!649ta2pN | n669ta2pmetN |
| Acetyl | 4-Phos | Methyl |
| C$761tt3pAcetH | C!796tt4pH | C814tt4pmet |
| C794AG3pACETH | C!830ag4pLN | N$849ag4pmetN |
| C809gg3pacetHN | c846gg4pH | c861gg4pmetH |
| C771ta3pACETH | C!807ta4pH | C!822ta4pmetH |
| C785tg3pACETH | C!821tg4pH | C!835tg4pmetHL |
| | Cancerous | |
| C!739cc3pAcetH | C!770cc4pH | C!784cc4pmetH |
| C!757ca3pacetH | C!794ca4pH | C!807ca4pmetHN |
| C!777aa3pacetH | c814aa4pHN | C!830aa4pmetL |
| C749tc3pacet | C!784tc4pH | c!796tc4pmetH |
| C771ta3pACETH | C!807ta4pH | C!822ta4pmetH |
| Acetyl | 6-Phos | Methyl |
| c917tt5pacetH | C!954tt6pH | N$967tt6pmetHN |
| c!954ag5pacetH | N$990ag6pHN | C!1004ag6pmetH |
| N$967gg5pacetH | C!999gg6pL | C!1021gg6pmetH |
| N$924ta5pacetH | c960ta6pH | C!978ta6pmetH |
| N$946tg5pacetHN | c977tg6p | N$992tg6pmet |
| | (Mixed | |
| N$885cc5pacetN | N$924cc6pH | C!940cc6pmetHN |
| N$907ca5pacet | N$946ca6pHN | c960ca6pmetH |
| C!940aa5pacetH | n970aa6p | C982aa6pmetH |
| N$907tc5pacetH | C!940tc6pH | C!954tc6pmetH |
| N$924ta5pacetH | c960ta6pH | C!978ta6pmetH |

| | | |
|---|---|---|
| | DiPhos | |
| acetyl | | met |
| C!360g1pmetH | C!391G1pacetH | c428g2pL |
| ~~~~~ | C372a1pacetH | c409a2pH |
| C!335t1metH | 359t1pacetH | C398t2p |
| ~~~~~ | C349c1pacetH | C383c2pH |

Acetyl

C544g3pacetH

n528a3pacetH

n517t3pacetH

C!508c3pacetH

Legend:

Purple = Only in Cancer DNA

Green = Only in Normal DNA

H = Heavy Isotopes (13C, 17

L = Light Isotopes (15N)

N = Isotopes in Normal

* = Isotopically Clumped

Black = in Cancer and Norma

! = Most Intense Peaks

Red = Only in Cancer DNA

Gold = Only in Normal DNA

acetyl
Missing
C1738GGATT5pAcetH* Telomeres
C1757TTGGG5pAcetH*

N1624CCCTA5pAcet
N1649CCTAC5pAcet Mating Telomeres
C1635AACCC5pAcet

acetyl
N1923GGGAT7pAcet
C1893GGATT7pAcet Telomeres
N1914TTGGG7pAcet

N1778CCCTA7pAcet
CN1807CCTAC7pAcet Mating Telomeres
C1791AACCC7pAcet

acetyl
Missing
C2052GGATT9pAcet***L*** Telomeres
C2062TTGGG9pAcet***L***

N1941CCCTA9pAcet
N1964CCTAC9pAcet Mating Telomeres
N1945AACCC9pAcet

acetyl
C2232gggat11pAcet

Legend:
Purple = Only in Cancer DNA
Green = Only in Normal DNA
H = Heavy Isotopes (13C, 17O)
L = Light Isotopes (15N)
N = Isotopes in Normal
* = Isotopically Clumped
Black = in Cancer and Normal DNA
! = Most Intense Peaks
Red = Only in Cancer DNA

C2204GGATT11pAcet		Telomeres		Gold = Only in Normal DNA
Missing

C2094CCCTA11pAcet
C2016CCTAC11pAcet		Mating Telomeres
C2095AACCC11pAcet

acetyl
C2384gggat13pAcet
C2363GGATT13PAcet		Telomeres
C2373ttggg13pAcet***H***

C2248cccta13pAcet
c2273aatac13pAcet		Mating Telomeres
C2254aaccc13PAcet

acetyl
C2445GGGAT15pAcet***H****
C2519GGATT15pACET***L****		Telomeres
C2535TTGGG15pAcet***L***

C2407CCCTA15pAcet***H***
CN2429aatac15pAcet***L****		Mating Telomeres
C2417aaccc15pAcet***H***

Acetyl
N1428ggga4pacet*
N1399gggt4pact*

c1381ggat4pacet*    Telomeres
C1377ggtt4pacet*
C1354gatt4pacet

C1277TCCC4pacet
n1203atcc4pacet
C1287ACCC4pacet
C1316aacc4pacet    Mating Telomeres
N1331aaTC4pacet*

Acetyl
c1566ggga6pacet
n1551gggt6pacet    Telomeres
n1538ggat6pacet
n1530ggtt6pacet
C!1515gatt6pacet

c!1435tccc6pacet*
c!1457atcc6pacet*
N1443accc6pacet*    Mating Telomeres
c1469aacc6pacet
n$1484aatc6pacet

Acetyl
n1722ggga8pacet***H***
c!1711gggt8pacet***H***    Telomeres
c!1696ggat8pacet
c!1687ggtt8pacet
c!1671gatt8pacet*

n$1586tccc8pacet
c!1613atcc8pacet
n1596accc8pacet    Mating Telomeres
n1623aacc8pacet*
C!1637aatc8pacet*

Acetyl
c!1879ggga10pacet
n1869gggt10p*acet    Telomeres
n1851ggat10p*acet

Legend:
Purple = Only in Cancer DNA
Green = Only in Normal DNA
H = Heavy Isotopes (13C, 17O)
L = Light Isotopes (15N)
N = Isotopes in Normal
* = Isotopically Clumped
Black = in Cancer and Normal DNA
! = Most Intense Peaks
Red = Only in Cancer DNA
Gold = Only in Normal DNA

n1842ggtt10p*acet

n1827gatt10p*acet

4

c!1747tccc10p*acet

c!1770atcc10p*acet

n1752accc10p*acet Mating Telomeres

n1779aacc10p*acet

n1796aatc10pacet

Acetyl

????

???? Telomeres

????

c1999ggtt12pacet

c!1985gatt12pacet

0

c1901tccc12pacet

c1927atcc12pacet

n1911accc12pacet* Mating Telomeres

c1938aacc12pacet

c1950aatc12pacet

Acetyl

~~~~~ C1096

c!1054tgg3pacetL

C!1031tgt3pacetL

~~~~~~ Telomeres

N1012tat3pacetL

~~~~~

c!1021taa3pacetH

C978tcc3pacet

C!997tca3pacetLH
~~~~~

| | |
|---|---|
| c!1009caa3pacetL | Mating Telomeres |
| C!958ccc3pacet | |
| C!982cca3pacetH | |

Acetyl

| | |
|---|---|
| C!1234ggg5pacetL | |
| C!1212tgg5paet | |
| c1184tgt5pacet | |
| c1193tag5pacet* | Telomeres |
| n1169tat5pacet* | |
| N!1222agg5pacetN | |

| | |
|---|---|
| C1177taa5pacet | |
| n1129tcc5pacet | |
| n1155tca5pacetH | |
| c1162caa5pacet | Mating Telomeres |
| c!1119ccc5pacetH | |
| C!1138cca5pacet | |

Acetyl

| | |
|---|---|
| C!1389GGG7pacet*N | |
| C!1367TGG7pacet*N | |
| C!1343TGT7pAcet*H | |
| C!1350TAG7pAcet*H | Telomeres |
| C!1328TAT7pAcet*H | |
| C!1377agg7pacetN | |

| | |
|---|---|
| C!1333TAA7pacet | |
| N1285tcc7pacetL | |
| C!1311tca7pacet*H | |
| N$1318CAA7pACET | Mating Telomeres |
| N1270ccc7pacetN | |
| c1294aac7pacet | |

Legend:

Purple = Only in Cancer DNA

Green = Only in Normal DNA

H = Heavy Isotopes (13C, 17O)

L = Light Isotopes (15N)

N = Isotopes in Normal

* = Isotopically Clumped

Acetyl

N1543ggg9pacet

C!1521tgg9pacetH

C!1499tgt9pacetL

N$1505tag9pacetN Telomeres

N$1483tat9pacetN

C1530agg9pacet

n1480taa9pacet

N$1439TCC9pacetL

C!1469TCA9pacet

C!1477caa9pacetL Mating Telomeres

C!1426ccc9pacet

C1451cca9pacetL

Black = in Cancer and Normal DNA

! = Most Intense Peaks

Red = Only in Cancer DNA

Gold = Only in Normal DNA

Acetyl

C!686tt2pacetH

c720ag2pacetL!!!

N$732gg2pacetN Telomeres

***C!695 ta2pacet L***

N$710tg2pacetN

C!654cc2pacetH

C!680ca2pacetH

***C!695 aa2pacet L*** Mating Telomeres

N!669tc2pacetNL

***C695 ta2pacet L***

Acetyl

C!840tt4pacetH

C!875ag4pacetH

N$885gg4pacetN Telomeres

C!853ta4pacetHN

c864tg4pacetH

C!807cc4pacetGN

C!835ca4pacetHN

N!853aa4pacetH Mating Telomeres

C!822tc4pacetH

C!853ta4pacetHN

***Acetyl***

C!995tt6pacetL

C!1031ag6pacetL

n1046gg6pacet Telomeres

C!1004ta6pacetH

C!1021tg6pacetH

N$967cc6pacetH

N$990ca6pacetN

c1013aa6pacetL Mating Telomeres

C!982tc6pacetH

C!1004ta6pacetH

Legend:
Purple = Only in Cancer DNA
Green = Only in Normal DNA
H = Heavy Isotopes (13C, 17O)
L = Light Isotopes (15N)
N = Isotopes in Normal
* = Isotopically Clumped
Black = in Cancer and Normal DNA
! = Most Intense Peaks
Red = Only in Cancer DNA
Gold = Only in Normal DNA

Acetyl

| | |
|---|---|
| C441g2pmet | C469g2pacetN |
| C422a2pmet | N$449a2pacetNH |
| N$413t2pmetLN | C441t2pacet |
| C404c2pmetNH | N$429c2pacetN |

A

A

es (13C, 17O)

and Normal DNA
