## Supplementary Materials II for "Oncogenesis and Aging by Isotopic Functionalizations of the Proteins and Nucleic Acids"

II Supplementary Materials – New Mass Spectra from 1800 Da -2000 Da

Results for Supplementary 2

So for the tetranucleotides from $(phos)_{12}$ to $(phos)_{11}$, the peaks for cancer decrease in number and the peaks for normal increase in number for more stable normal tetranucleotides and more unstable cancerous tetranucleotides with these dephosphorylations. See Supplementary Figures II for graphs. These results and observations point specific differences between cancerous and normal oligonucleotides chemically and isotopically. The dephosphorylations caused the more fragmenting and expressions of resulting more stable normal tetranucleotides to be more telomeric and isotopically enriched and the more stable cancer tetranucleotides to be mating telomeric and not enriched. The dephosphorylations also caused the nontelomeric tetranucleotides to be more from normal DNA and isotopically enriched. It seems these dephosphorylations are related and induced by the isotopic enrichments. The nontelomeric tetranucleotides of cancer tended to not be enriched so it seems the cancer instabilities of the nontelomeric tetranucleotides are related to there non isotopic enrichments. Isotopic enrichments strengthen the nontelomeric cancer tetranucleotides. The more intense cancer telomeric tetranucleotides were also less isotopically enriched. The dephosphorylations from $(phos)_{12}$ to $(phos)_{11}$ increased the isotopic enrichments of the normal relative to the cancer for the telomeric and nontelomeric tetranucleotides. But the mating telomeric tetranucleotides were less enriched from the dephosphorylations for less enrichments and stronger more stable cancer tetranucleotides for mating telomeres. Such seems compatible for replications and transcriptions as stable telomere seems compatible to unstable mating telomere for the unravelings and constructions of DNA mating telomeres onto the building templates of the telomeres. The dephosphorylations caused methylations to cause more cancer stabilities and intensities but less isotopic enrichments for telomeres and nontelomer tetranucleotides. The dephosphorylations caused acetylations to cause more cancer stabilities and more normal instabilities for telomeres and nontelomer tetranucleotides with more enrichments relative to methylations but the mating telomeres are more cancer stable and more normal enriched in nonprimordials. Thereby this dephosphorylations $(phos)_{12}$ to $(phos)_{11}$ and their varying consequences with respect to other dephosphorylations determines patterns of dephosphorylations and isotopic enrichment patterns in normal DNA which mutate and change for different patterns in DNA of cells that become cancerous. These dephosphorylations $(phos)_{12}$ to $(phos)_{11}$ induced isotopically dynamics from of varying methylations and acetylations in cancer DNA relative to normal DNA couple to other isotopically induced varied phosphorylations, methylations, and acetylations throughout the DNA for emerging order from nanodisorder.

So for the tetranucleotides from $(phos)_{11}$ to $(phos)_{10}$, the peaks for cancer decrease in number and the peaks for normal increase in number for more stable normal tetranucleotides and more unstable cancerous tetranucleotides with these dephosphorylations. See Supplementary Figures 2 for graphs. These results and observations point specific differences between cancerous and normal oligonucleotides chemically and isotopically. The dephosphorylations from $(phos)_{11}$ to $(phos)_{10}$ caused the resulting more stable normal tetranucleotides to be more telomeric and nontelomeric and isotopically enriched. The mating telomeres are not as stabilized with normal but both normal and cancerous mating telomeric tetranucleotides are highly isotopic enriched. The dephosphorylations from $(phos)_{11}$ to $(phos)_{10}$ also caused the nontelomeric tetranucleotides to be more from normal DNA and isotopically enriched. It seems the dephosphorylations are related to the isotopic enrichments. The nontelomeric, telomeric and mating telomeric tetranucleotides of cancer tended to also be enriched so it seems the cancer instabilities and normal stabilities of the nontelomeric, telomeric and mating telomeric tetranucleotides are related and induced by their isotopic enrichments. Enrichments strengthen the nontelomeric cancer tetranucleotides. The more intense cancer telomeric tetranucleotides were also less isotopically enriched. The dephosphorylations from $(phos)_{11}$ to $(phos)_{10}$ increased the isotopic enrichments of the cancer relative to the normal for the telomeric, mating telomeric and nontelomeric tetranucleotides. But the mating telomeric tetranucleotides were more enriched from the dephosphorylations for more enrichments and stronger more stable cancer tetranucleotides for mating telomeres. Such seems compatible for replications and transcriptions as stable telomere seems compatible to unstable mating telomeres for the unravelings and constructions of DNA mating telomeres onto the building templetes of the telomeres. The dephosphorylations from $(phos)_{11}$ to $(phos)_{10}$ caused methylations to cause more cancer stabilities and intensities but slightly less isotopic enrichments for telomeres and nontelomer tetranucleotides. But the methylations of the mating telomeres increased normal stabilities and isotopic enrichments for contrary effects on mating telomeres compared to telomeres. The dephosphorylations from $(phos)_{11}$ to $(phos)_{10}$ caused acetylations to cause more less cancer stabilities and more normal stabilities for telomeres with more enrichments relative to methylations; the nontelomeres are slightly more normal stable and less isotopically enriched; but the mating is more cancer stable and more normal enriched in nonprimordials. Thereby this dephosphorylations $(phos)_{11}$ to $(phos)_{10}$ and its varying consequence with respect to other dephosphrylations determine patterns of dephosphorylations and isotopic enrichment patterns in normal DNA which mutate and change for different patterns in DNA of cells that become cancerous. These dephosphorylations $(phos)_{11}$ to $(phos)_{10}$ induced isotopic dynamics from of varying methylations and acetylations in cancer DNA relative to normal DNA couple to other isotopically induced varied phosphorylations, methylations and acetylations throughout the DNA for emerging order from nanodisorder.

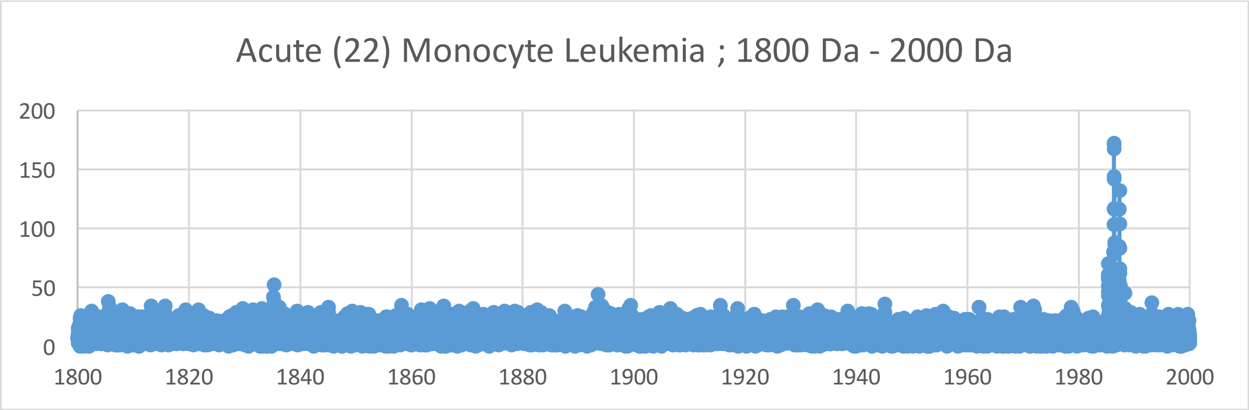
Acute (22) Monocyte Leukemia ; 1800 Da - 2000 Da
200
150
100
50
0
1800
1820
1840
1860
1880
1900
1920
1940
1960
1980
2000

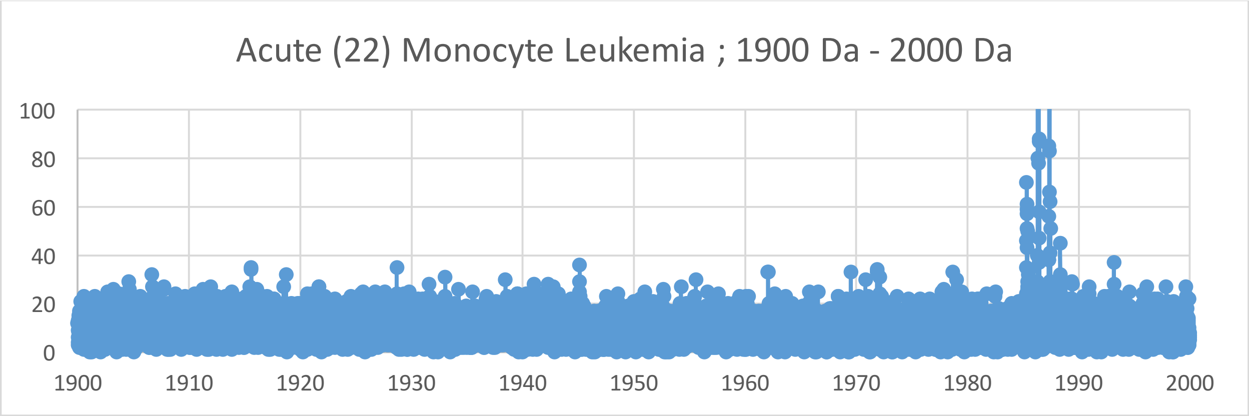
Acute (22) Monocyte Leukemia ; 1900 Da - 2000 Da
100
80
60
40
20
0
1900
1910
1920
1930
1940
1950
1960
1970
1980
1990
2000

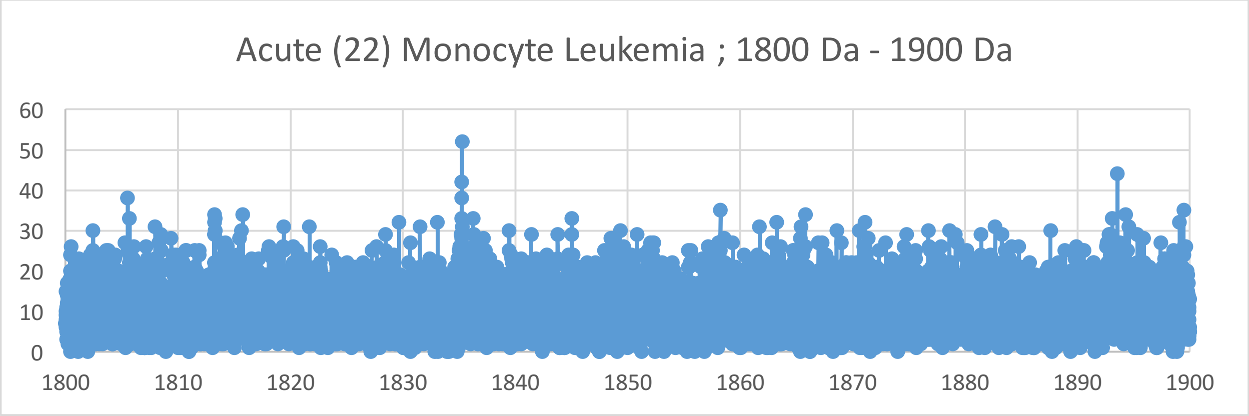
Acute (22) Monocyte Leukemia ; 1800 Da - 1900 Da
60
50
40
30
20
10
0
1800
1810
1820
1830
1840
1850
1860
1870
1880
1890
1900

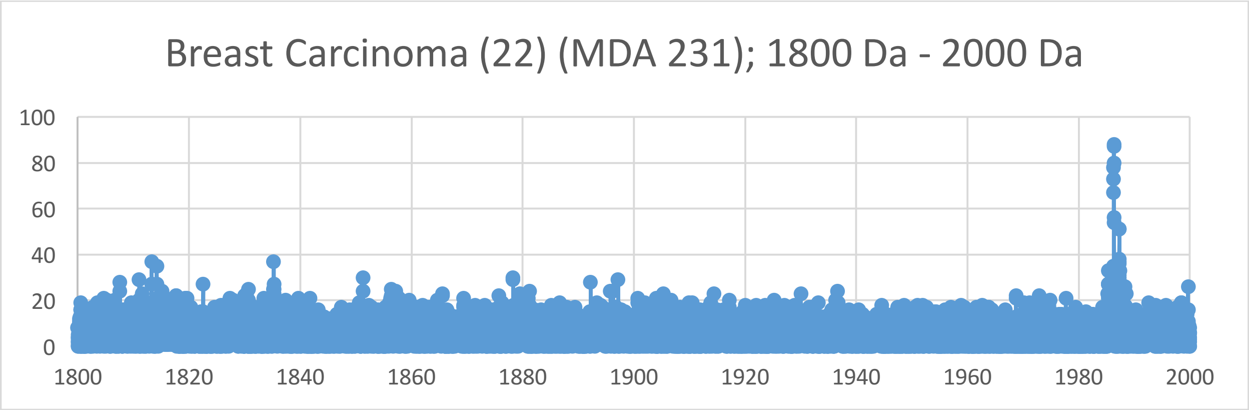
Breast Carcinoma (22) (MDA 231); 1800 Da - 2000 Da
100
80
60
40
20
0
1800
1820
1840
1860
1880
1900
1920
1940
1960
1980
2000

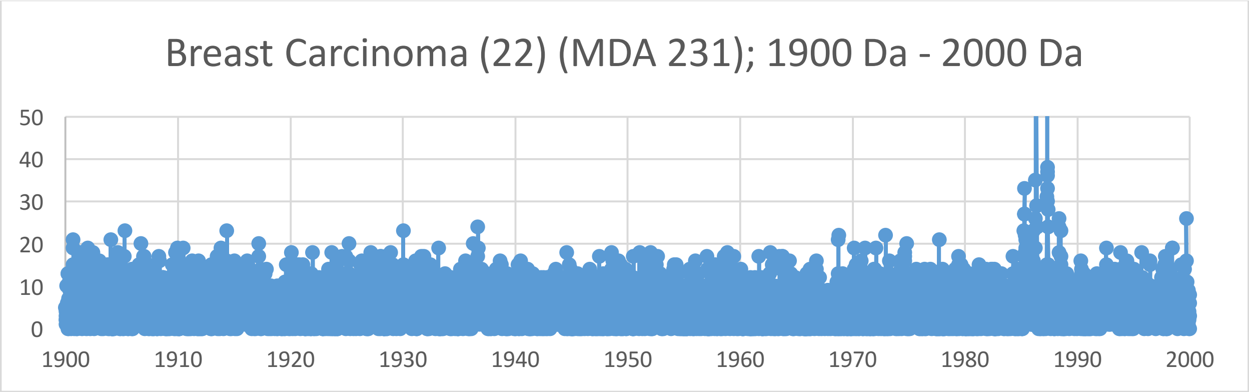
Breast Carcinoma (22) (MDA 231); 1900 Da - 2000 Da
50
40
30
20
10
0
1900
1910
1920
1930
1940
1950
1960
1970
1980
1990
2000

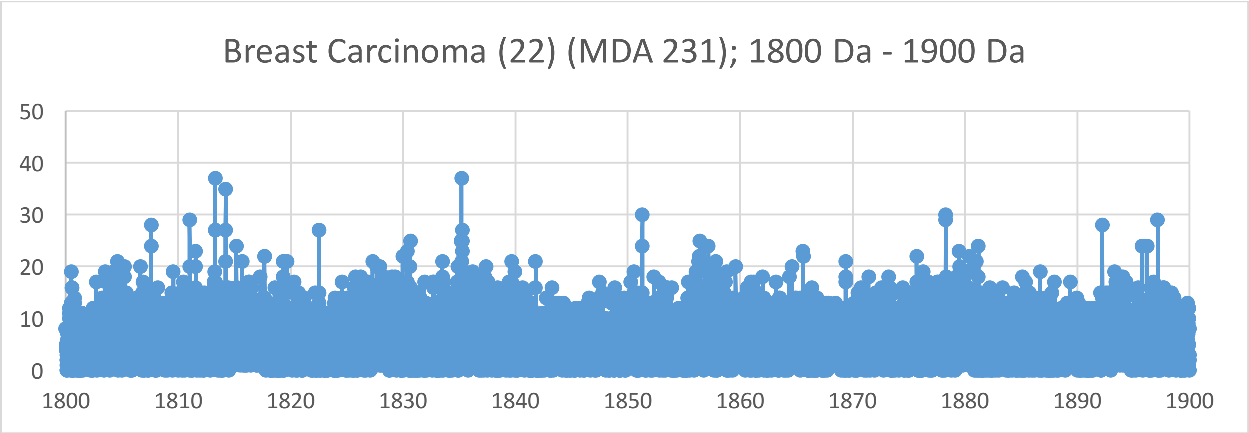
Breast Carcinoma (22) (MDA 231); 1800 Da - 1900 Da
50
40
30
20
10
0
1800
1810
1820
1830
1840
1850
1860
1870
1880
1890
1900

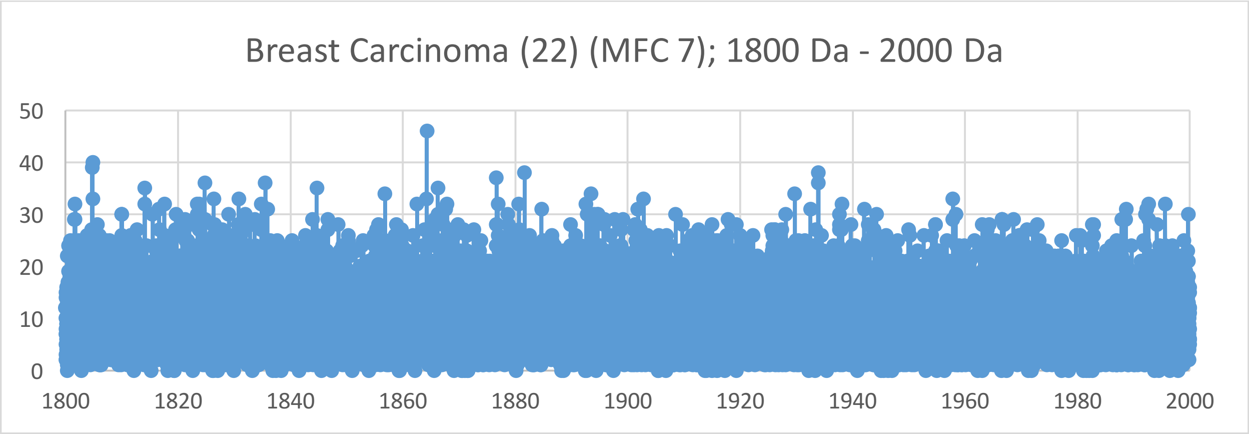
Breast Carcinoma (22) (MFC 7); 1800 Da - 2000 Da
50
40
30
20
10
0
1800
1820
1840
1860
1880
1900
1920
1940
1960
1980
2000

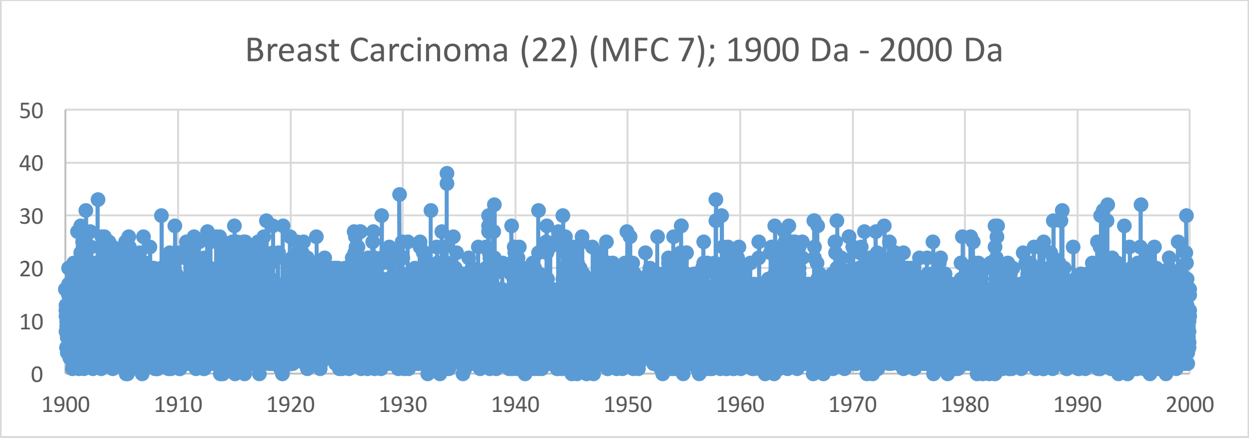
Breast Carcinoma (22) (MFC 7); 1900 Da - 2000 Da
50
40
30
20
10
0
1900
1910
1920
1930
1940
1950
1960
1970
1980
1990
2000

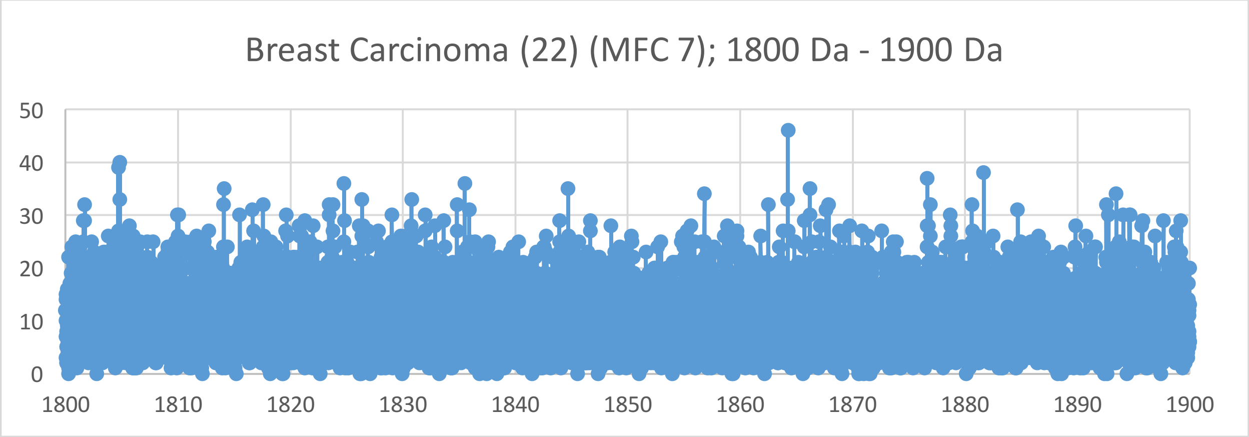
Breast Carcinoma (22) (MFC 7); 1800 Da - 1900 Da
50
40
30
20
10
0
1800
1810
1820
1830
1840
1850
1860
1870
1880
1890
1900

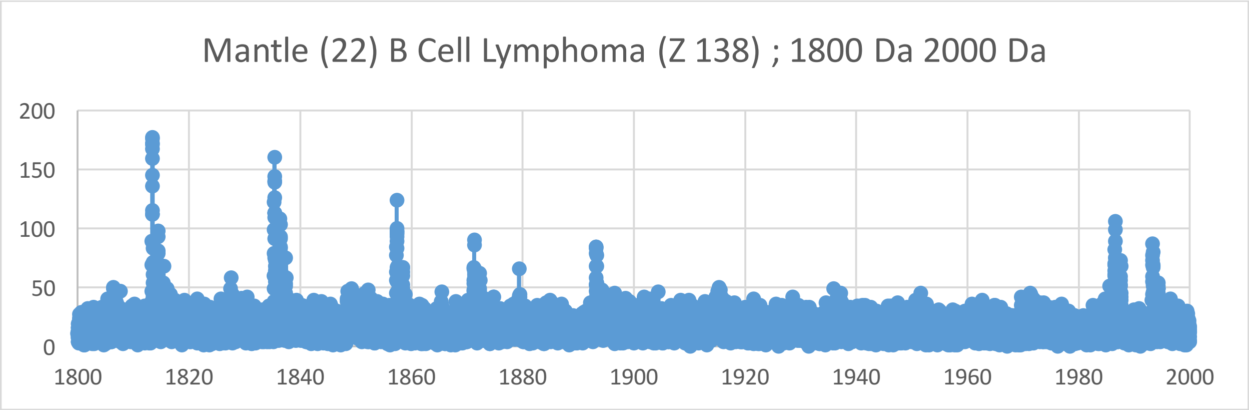
Mantle (22) B Cell Lymphoma (Z 138) ; 1800 Da 2000 Da
200
150
100
50
0
1800
1820
1840
1860
1880
1900
1920
1940
1960
1980
2000

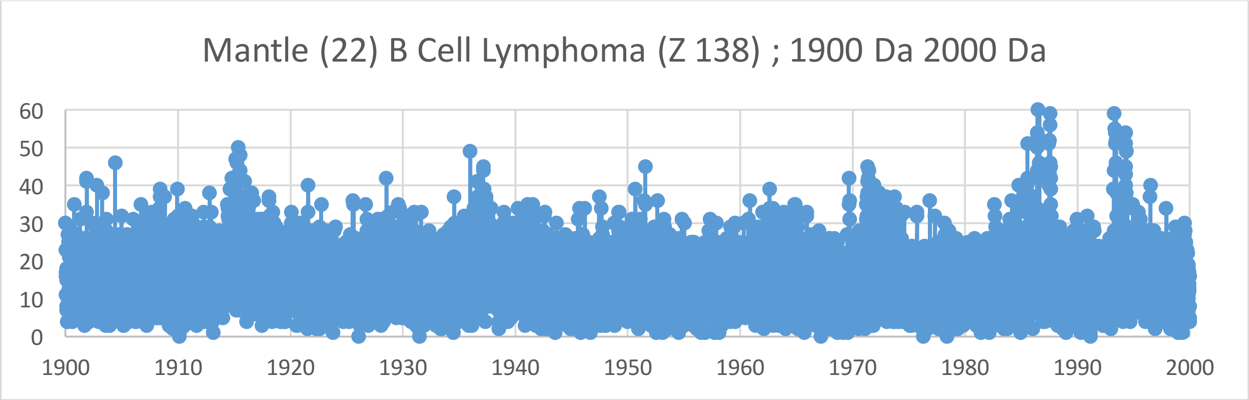
Mantle (22) B Cell Lymphoma (Z 138) ; 1900 Da 2000 Da
60
50
40
30
20
10
0
1900
1910
1920
1930
1940
1950
1960
1970
1980
1990
2000

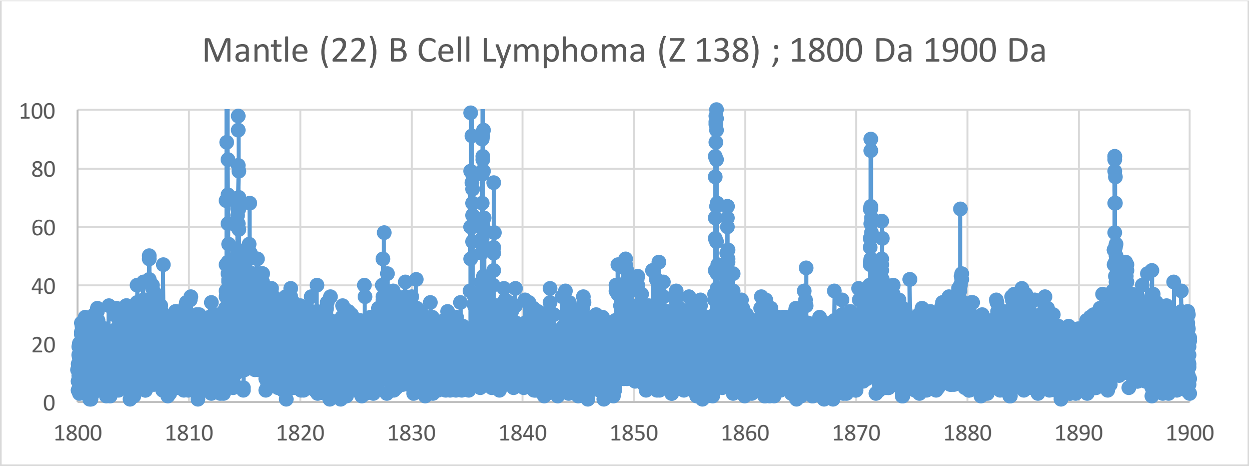
Mantle (22) B Cell Lymphoma (Z 138) ; 1800 Da 1900 Da
100
80
60
40
20
0
1800
1810
1820
1830
1840
1850
1860
1870
1880
1890
1900

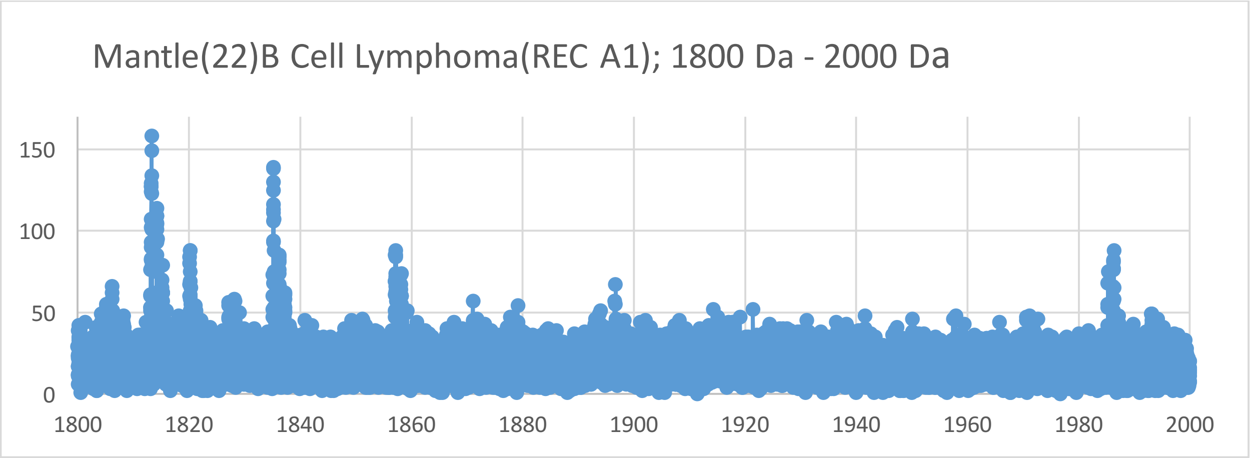
Mantle(22)B Cell Lymphoma(REC A1); 1800 Da - 2000 Da
0
50
100
150
1800
1820
1840
1860
1880
1900
1920
1940
1960
1980
2000

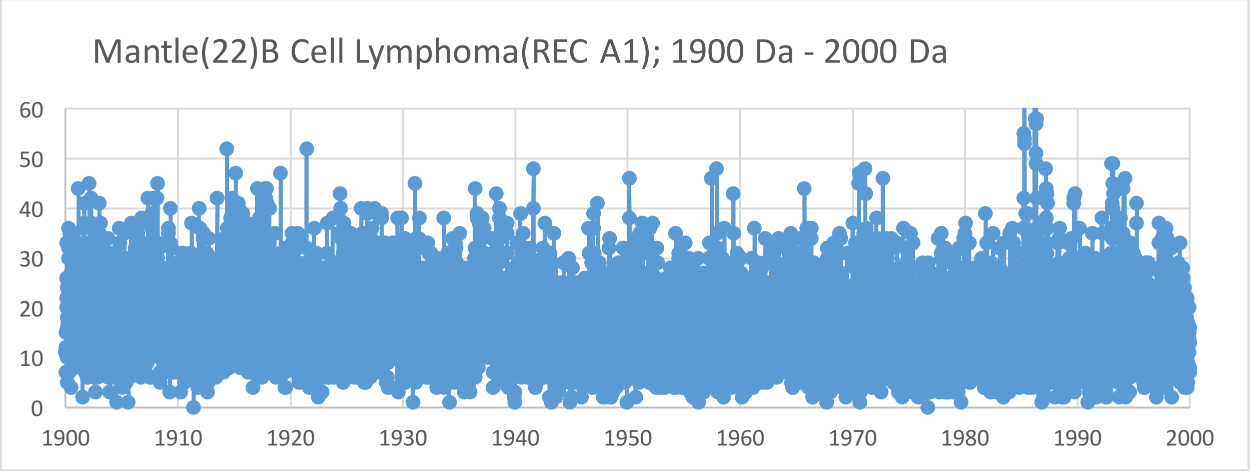
Mantle(22)B Cell Lymphoma(REC A1); 1900 Da - 2000 Da
0
10
20
30
40
50
60
1900
1910
1920
1930
1940
1950
1960
1970
1980
1990
2000

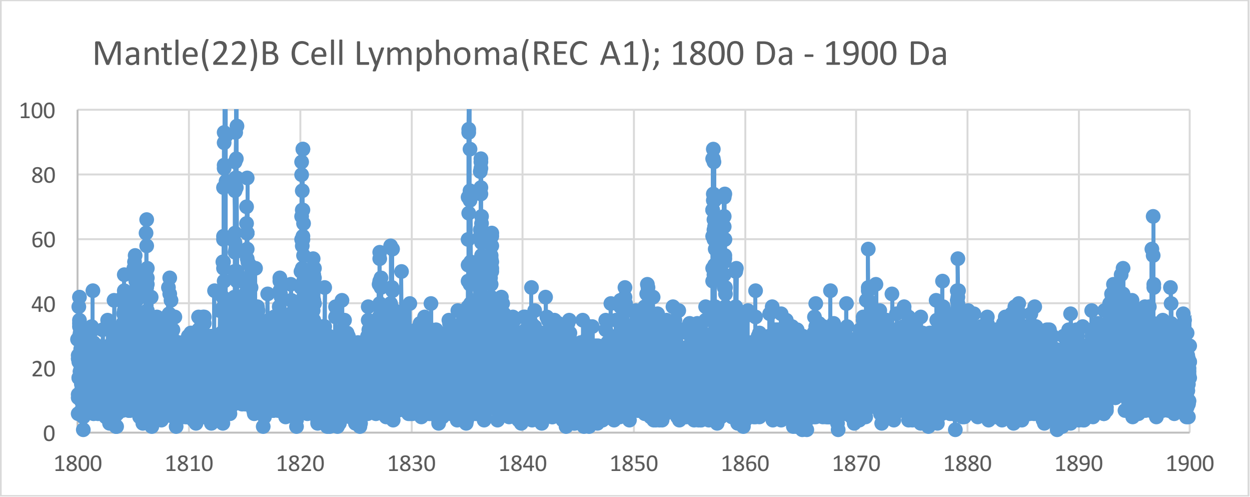
Mantle(22)B Cell Lymphoma(REC A1); 1800 Da - 1900 Da
0
20
40
60
80
100
1800
1810
1820
1830
1840
1850
1860
1870
1880
1890
1900

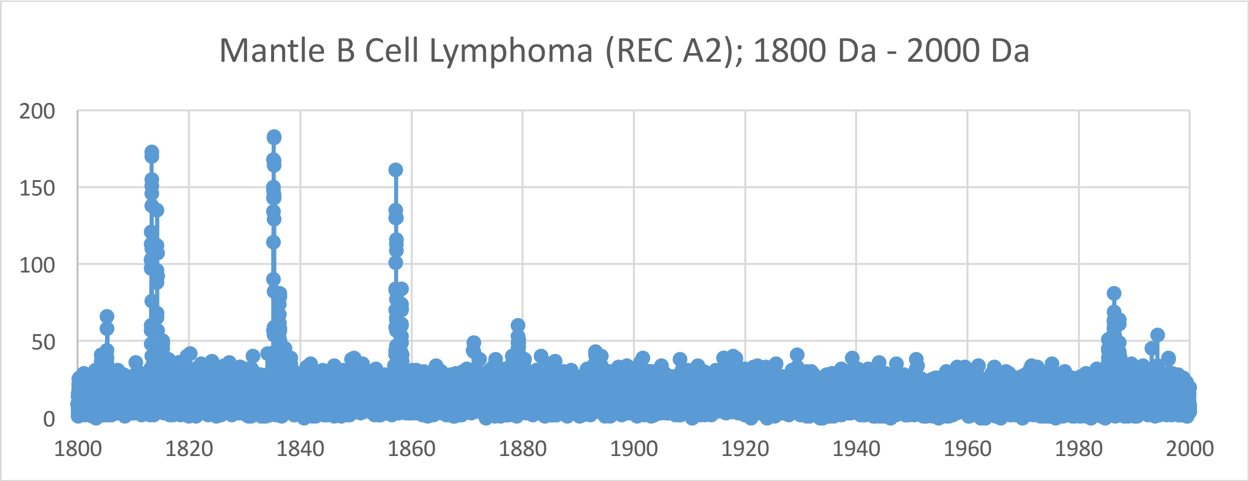
Mantle B Cell Lymphoma (REC A2); 1800 Da - 2000 Da
200
150
100
50
0
1800
1820
1840
1860
1880
1900
1920
1940
1960
1980
2000

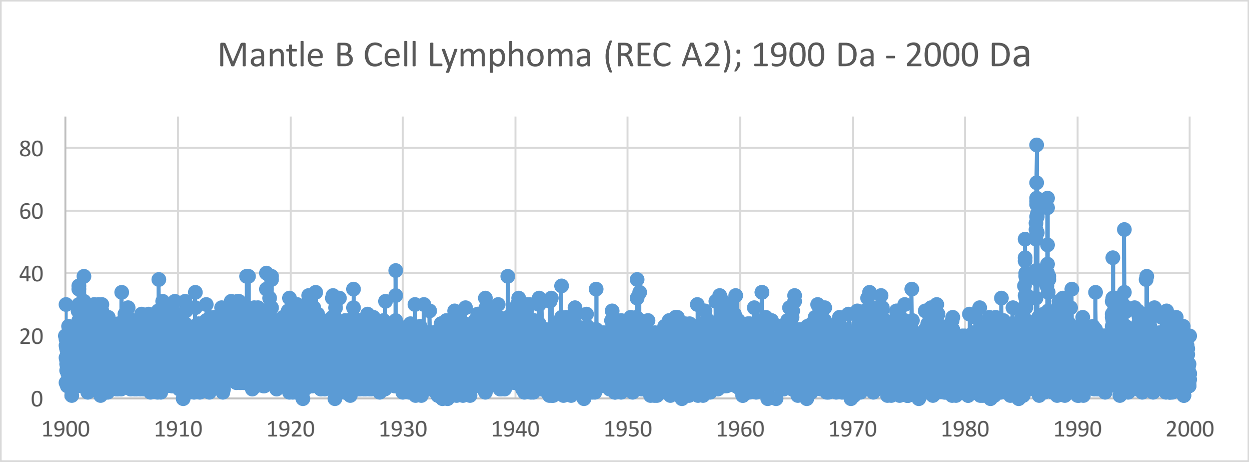
Mantle B Cell Lymphoma (REC A2); 1900 Da - 2000 Da
80
60
40
20
0
1900
1910
1920
1930
1940
1950
1960
1970
1980
1990
2000

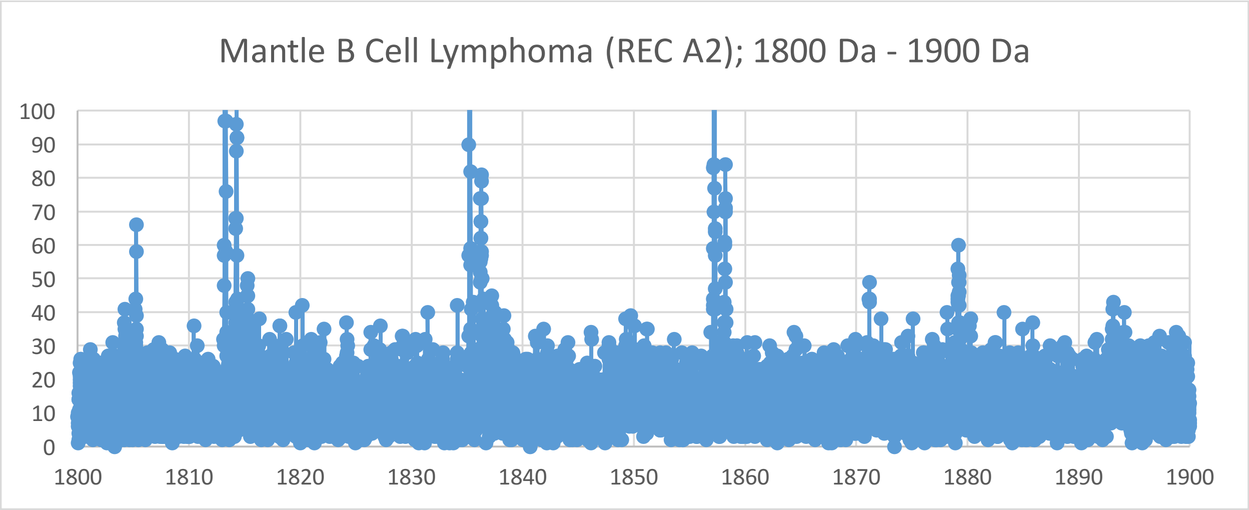
Mantle B Cell Lymphoma (REC A2); 1800 Da - 1900 Da
100
90
80
70
60
50
40
30
20
10
0
1800
1810
1820
1830
1840
1850
1860
1870
1880
1890
1900

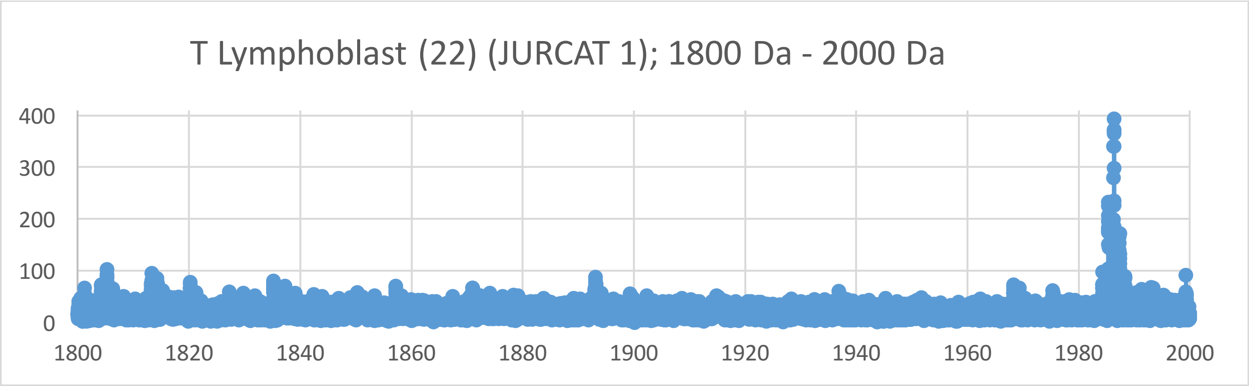
T Lymphoblast (22) (JURCAT 1); 1800 Da - 2000 Da
400
300
200
100
0
1800
1820
1840
1860
1880
1900
1920
1940
1960
1980
2000

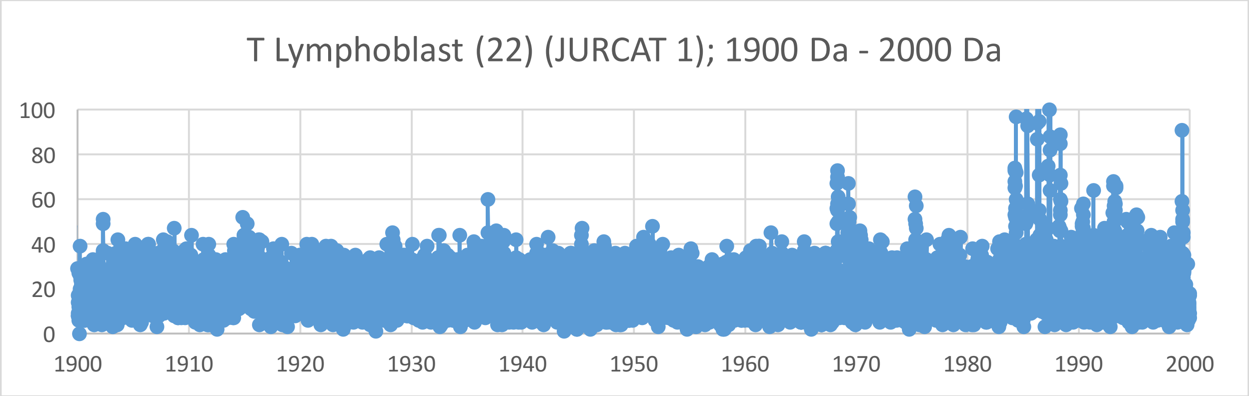
T Lymphoblast (22) (JURCAT 1); 1900 Da - 2000 Da
100
80
60
40
20
0
1900
1910
1920
1930
1940
1950
1960
1970
1980
1990
2000

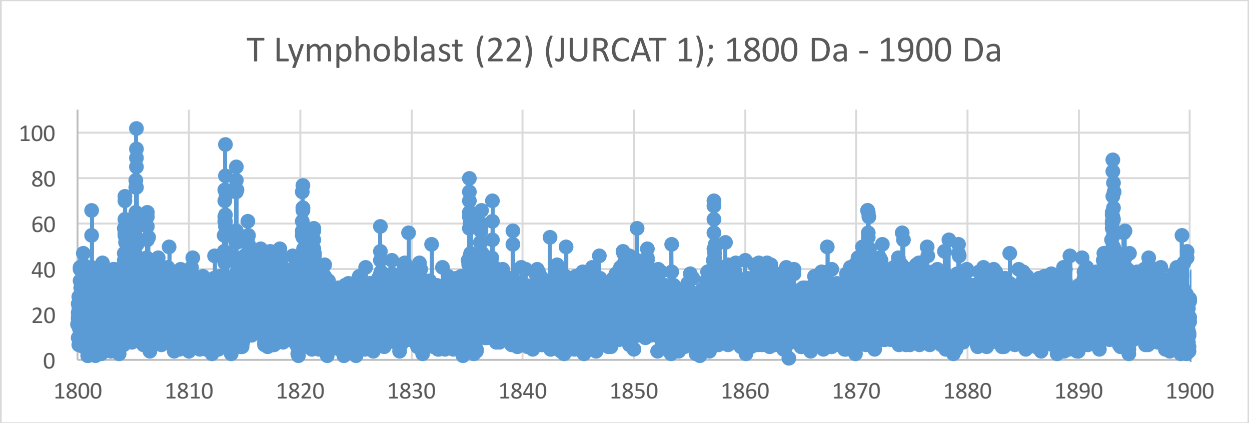
T Lymphoblast (22) (JURCAT 1); 1800 Da - 1900 Da
100
80
60
40
20
0
1800
1810
1820
1830
1840
1850
1860
1870
1880
1890
1900

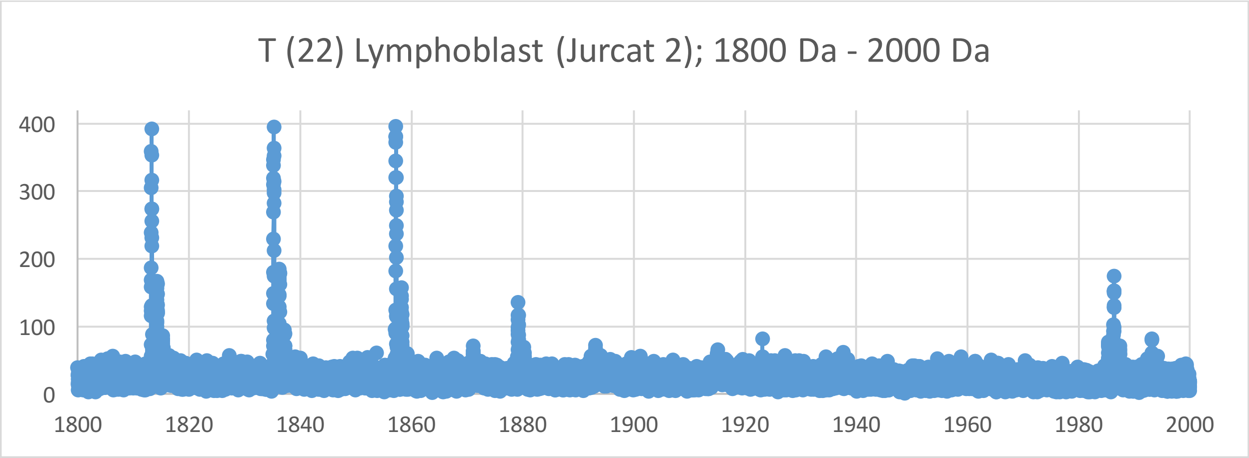
T (22) Lymphoblast (Jurcat 2); 1800 Da - 2000 Da
400
300
200
100
0
1800
1820
1840
1860
1880
1900
1920
1940
1960
1980
2000

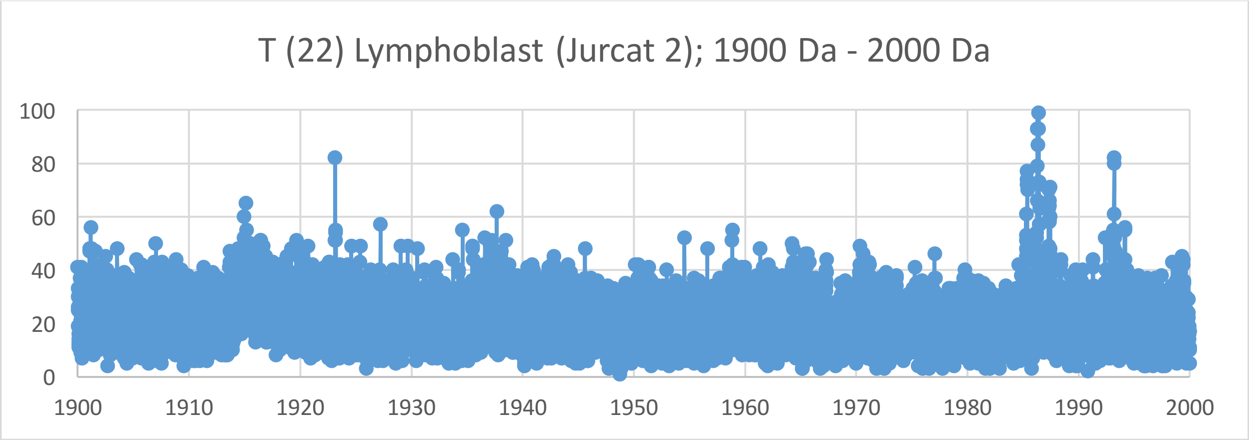
T (22) Lymphoblast (Jurcat 2); 1900 Da - 2000 Da
100
80
60
40
20
0
1900
1910
1920
1930
1940
1950
1960
1970
1980
1990
2000

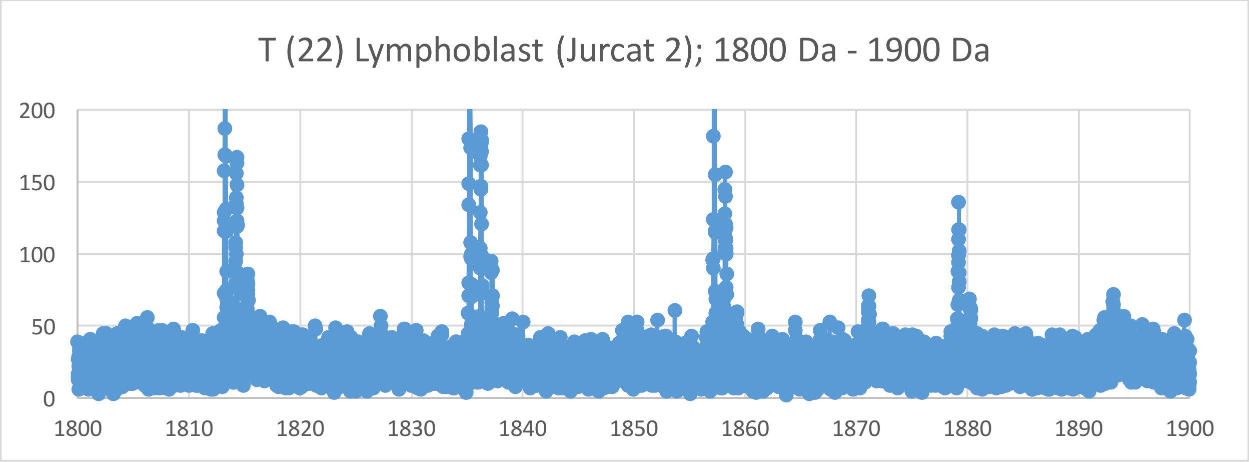
T (22) Lymphoblast (Jurcat 2); 1800 Da - 1900 Da
200
150
100
50
0
1800
1810
1820
1830
1840
1850
1860
1870
1880
1890
1900

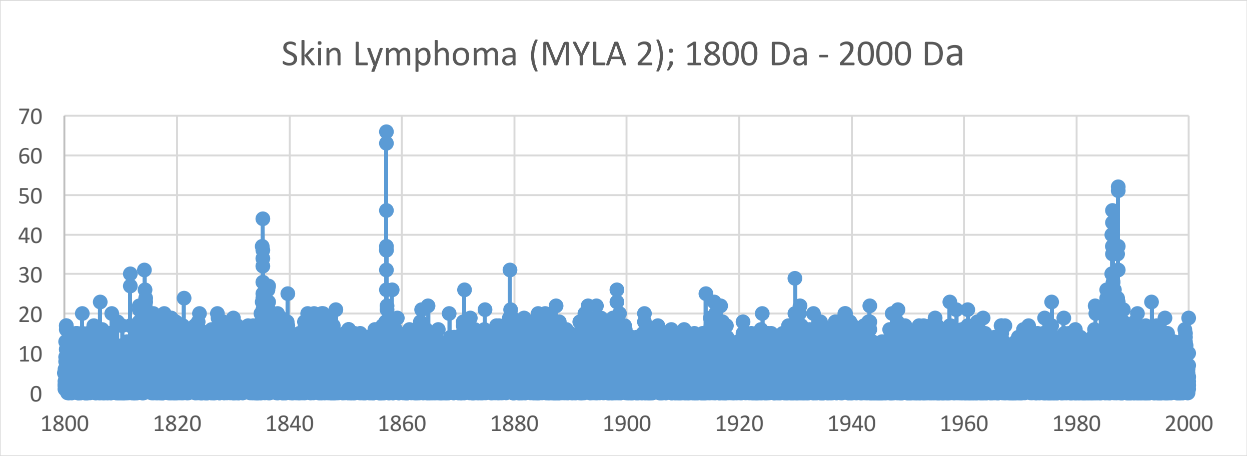
Skin Lymphoma (MYLA 2); 1800 Da - 2000 Da
70
60
50
40
30
20
10
0
1800
1820
1840
1860
1880
1900
1920
1940
1960
1980
2000

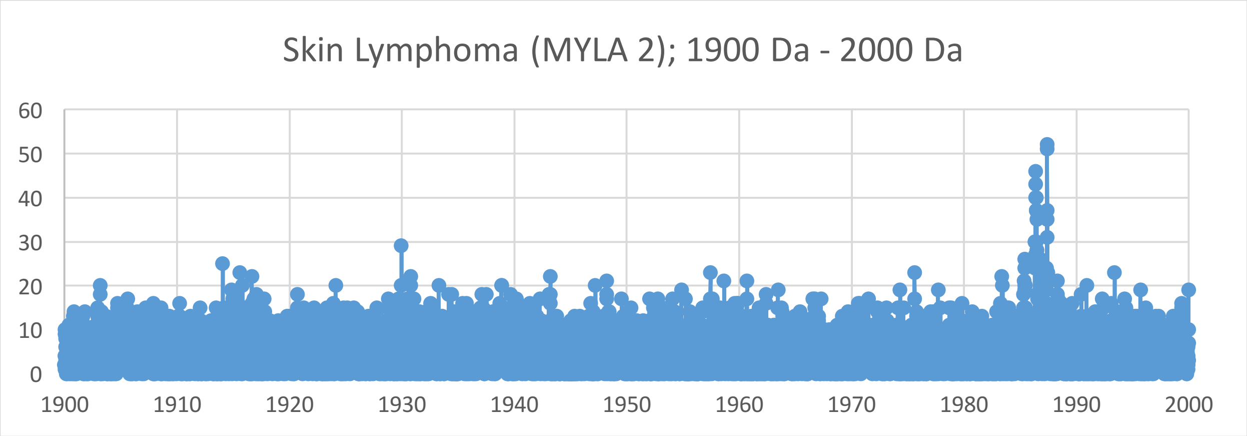
Skin Lymphoma (MYLA 2); 1900 Da - 2000 Da
60
50
40
30
20
10
0
1900
1910
1920
1930
1940
1950
1960
1970
1980
1990
2000

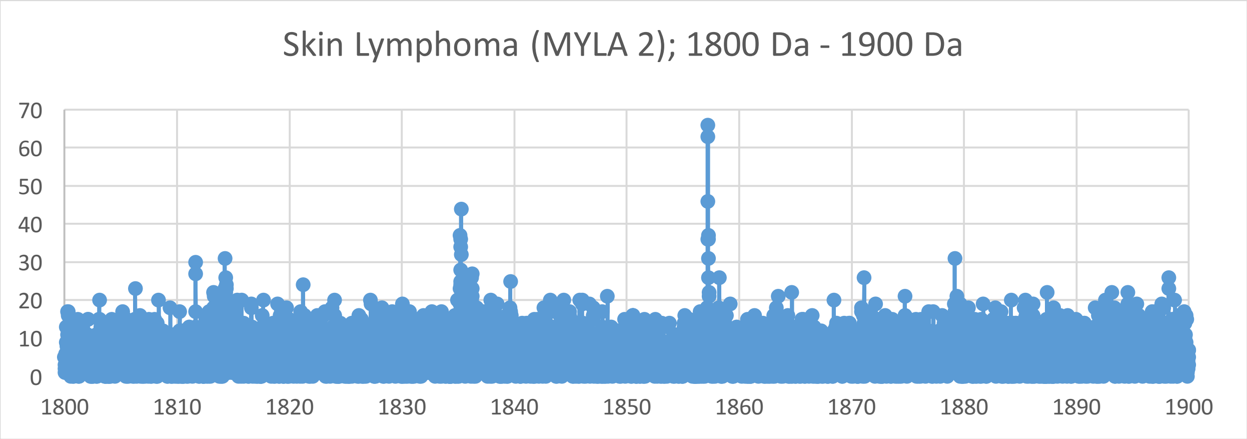
Skin Lymphoma (MYLA 2); 1800 Da - 1900 Da
70
60
50
40
30
20
10
0
1800
1810
1820
1830
1840
1850
1860
1870
1880
1890
1900

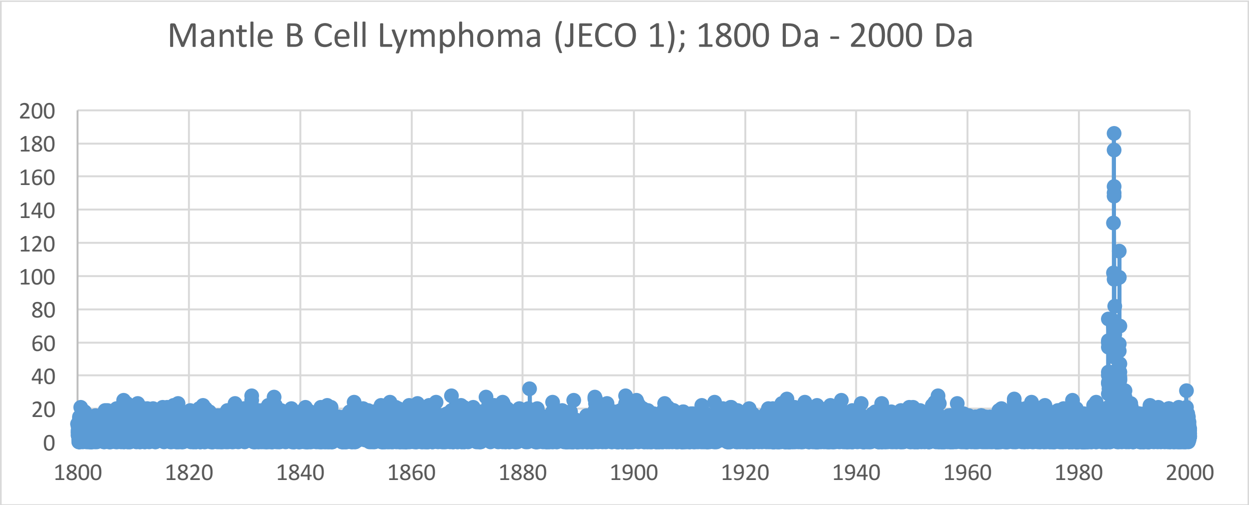
Mantle B Cell Lymphoma (JECO 1); 1800 Da - 2000 Da
200
180
160
140
120
100
80
60
40
20
0
1800
1820
1840
1860
1880
1900
1920
1940
1960
1980
2000

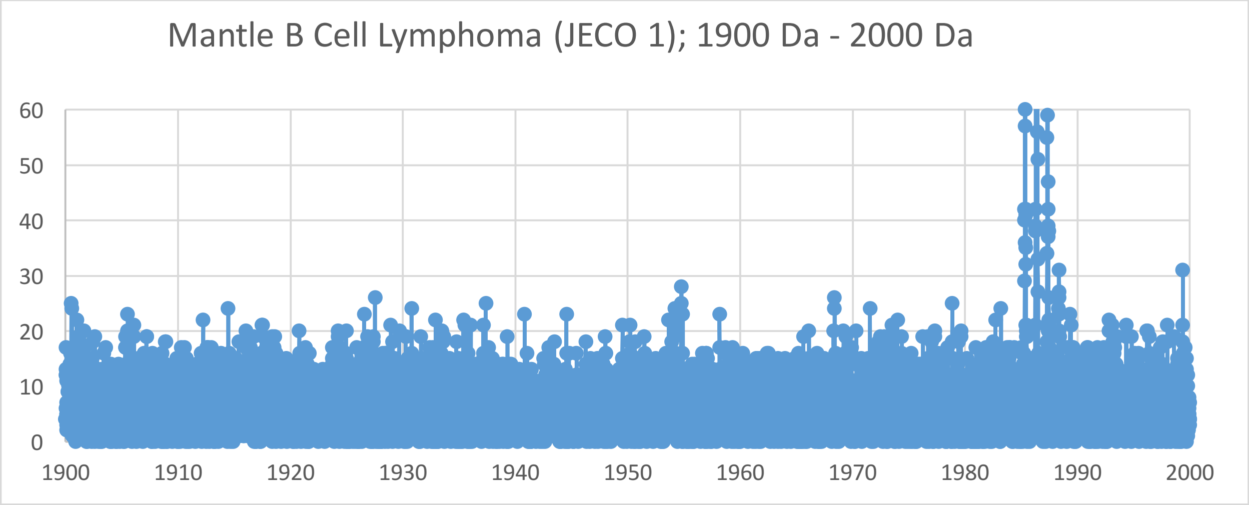
Mantle B Cell Lymphoma (JECO 1); 1900 Da - 2000 Da
60
50
40
30
20
10
0
1900
1910
1920
1930
1940
1950
1960
1970
1980
1990
2000

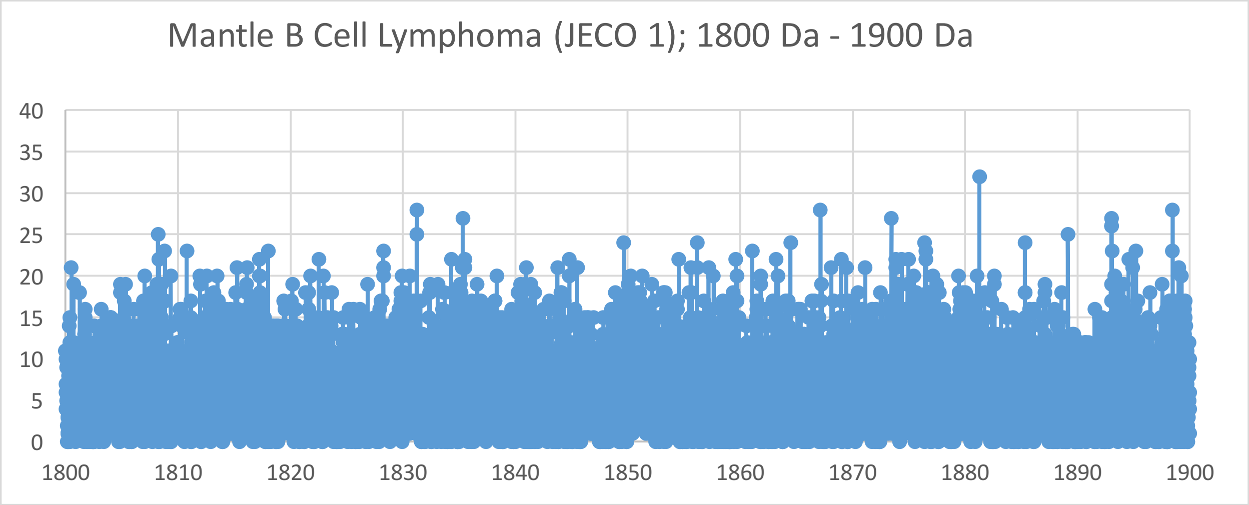
Mantle B Cell Lymphoma (JECO 1); 1800 Da - 1900 Da
40
35
30
25
20
15
10
5
0
1800
1810
1820
1830
1840
1850
1860
1870
1880
1890
1900

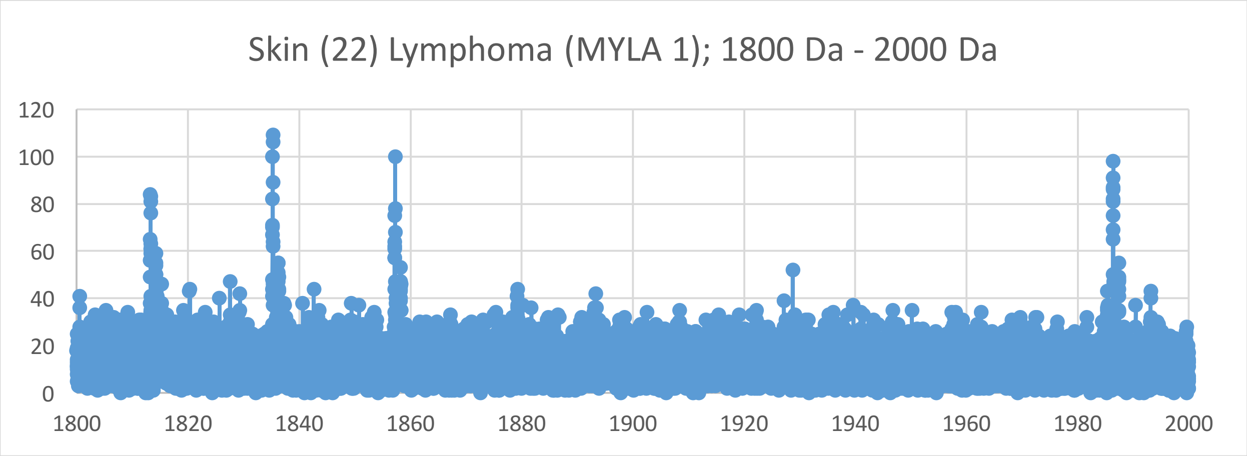
Skin (22) Lymphoma (MYLA 1); 1800 Da - 2000 Da
120
100
80
60
40
20
0
1800
1820
1840
1860
1880
1900
1920
1940
1960
1980
2000

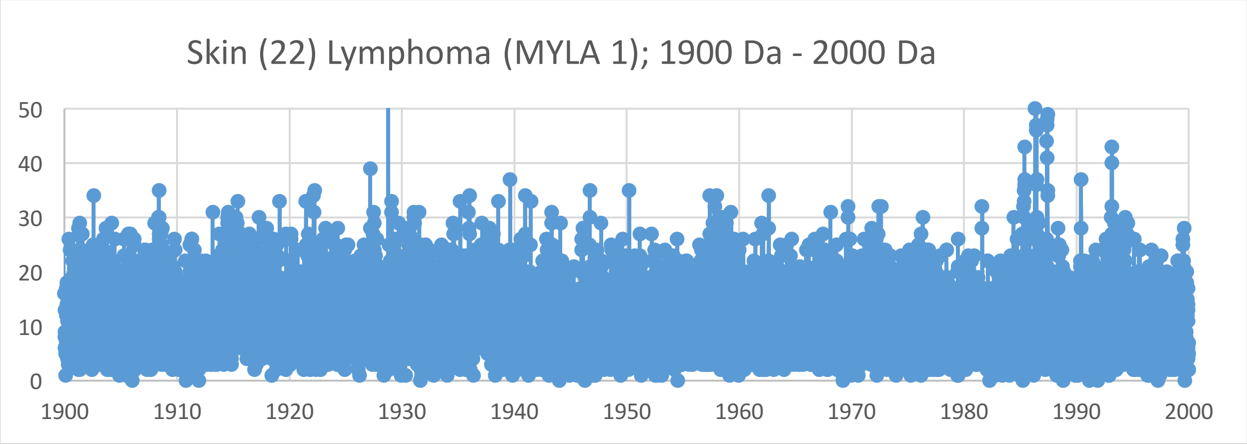
Skin (22) Lymphoma (MYLA 1); 1900 Da - 2000 Da
50
40
30
20
10
0
1900
1910
1920
1930
1940
1950
1960
1970
1980
1990
2000

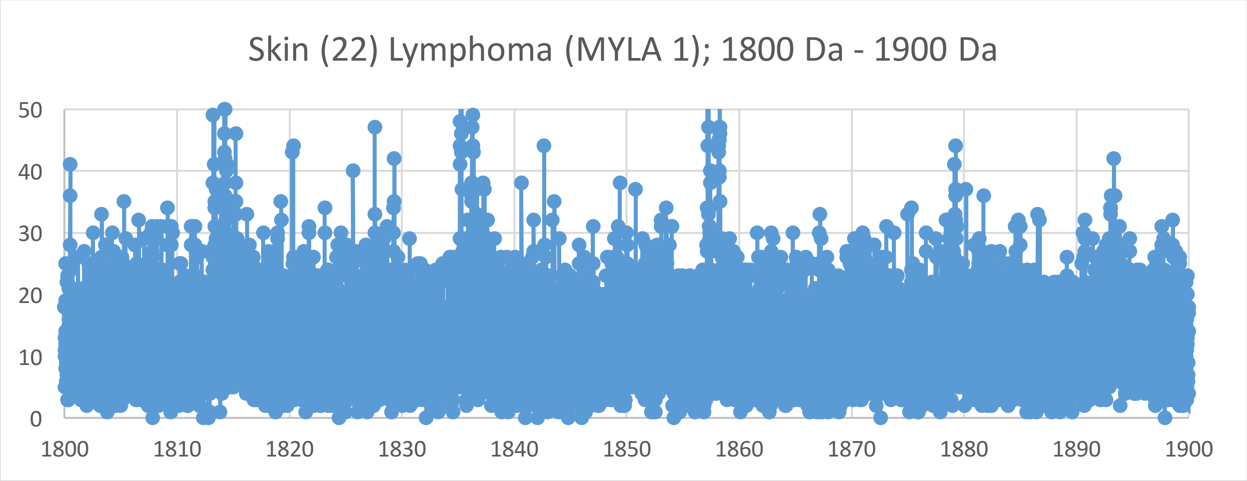
Skin (22) Lymphoma (MYLA 1); 1800 Da - 1900 Da
50
40
30
20
10
0
1800
1810
1820
1830
1840
1850
1860
1870
1880
1890
1900

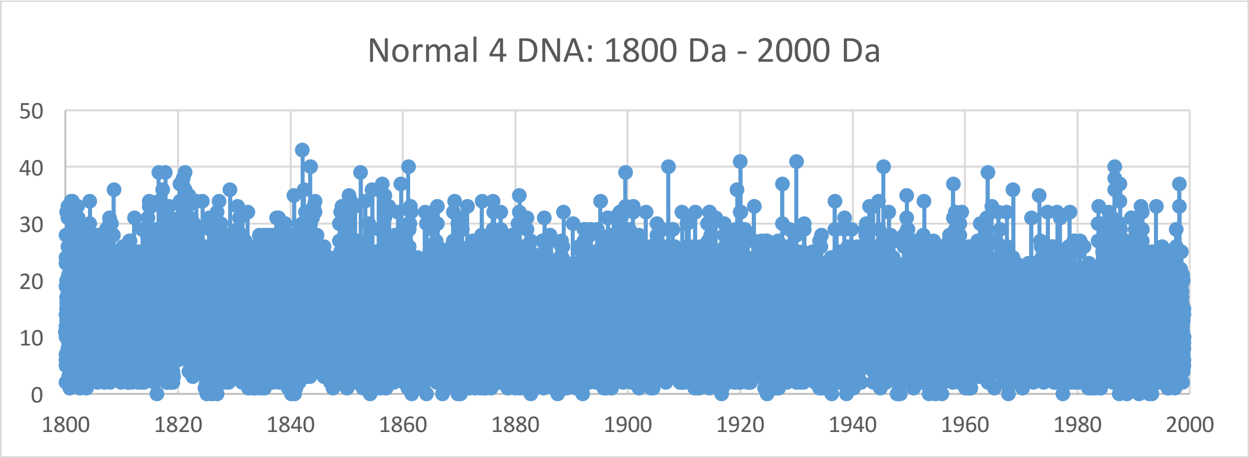
Normal 4 DNA: 1800 Da - 2000 Da
50
40
30
20
10
0
1800
1820
1840
1860
1880
1900
1920
1940
1960
1980
2000

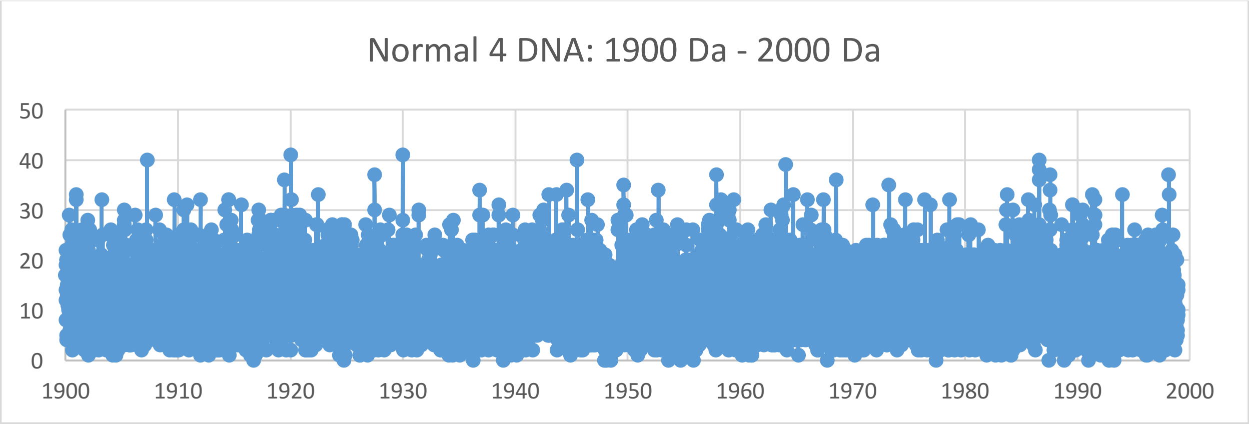
Normal 4 DNA: 1900 Da - 2000 Da
50
40
30
20
10
0
1900
1910
1920
1930
1940
1950
1960
1970
1980
1990
2000

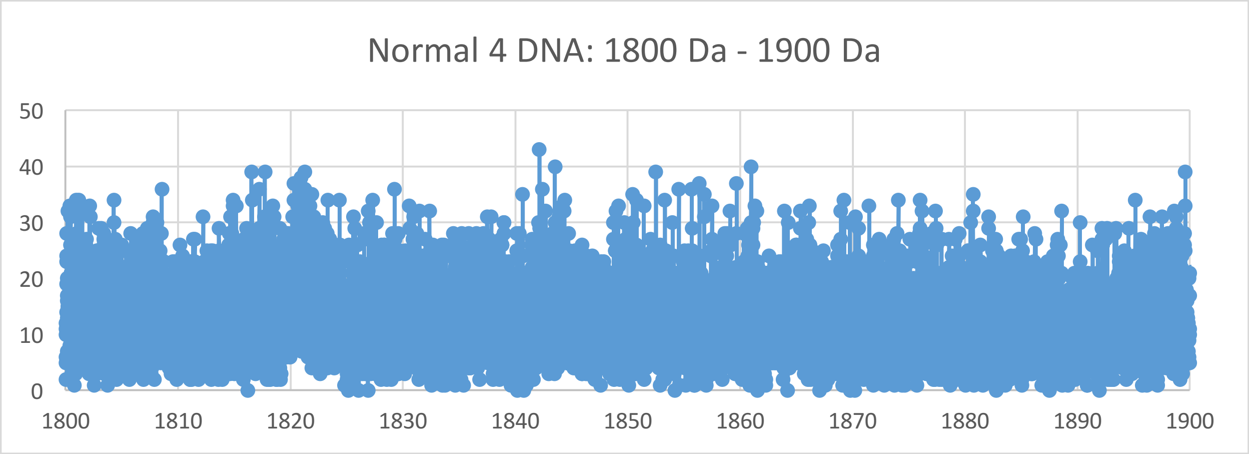
Normal 4 DNA: 1800 Da - 1900 Da
50
40
30
20
10
0
1800
1810
1820
1830
1840
1850
1860
1870
1880
1890
1900

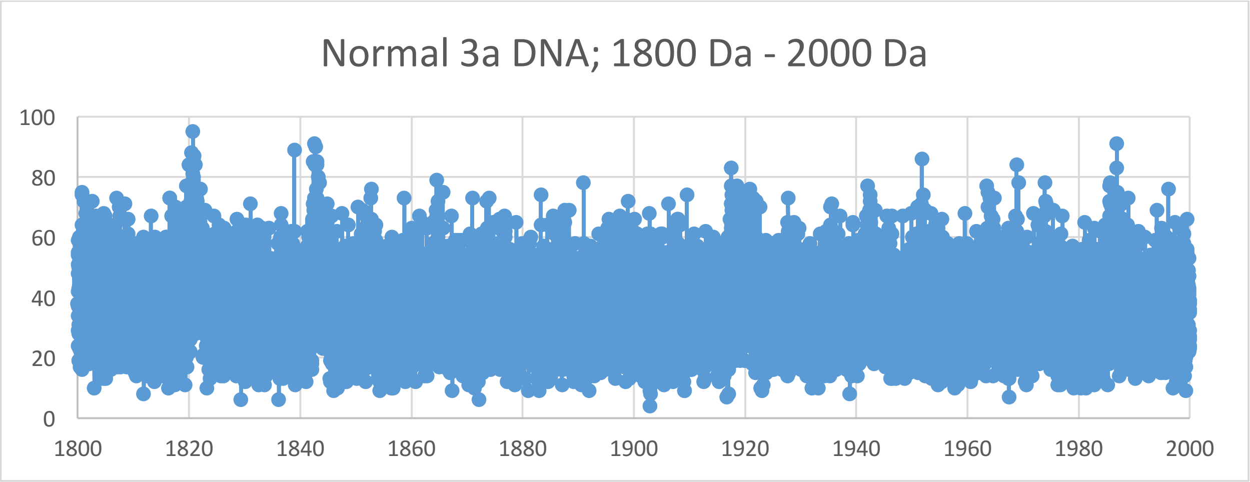
Normal 3a DNA; 1800 Da - 2000 Da
100
80
60
40
20
0
1800
1820
1840
1860
1880
1900
1920
1940
1960
1980
2000

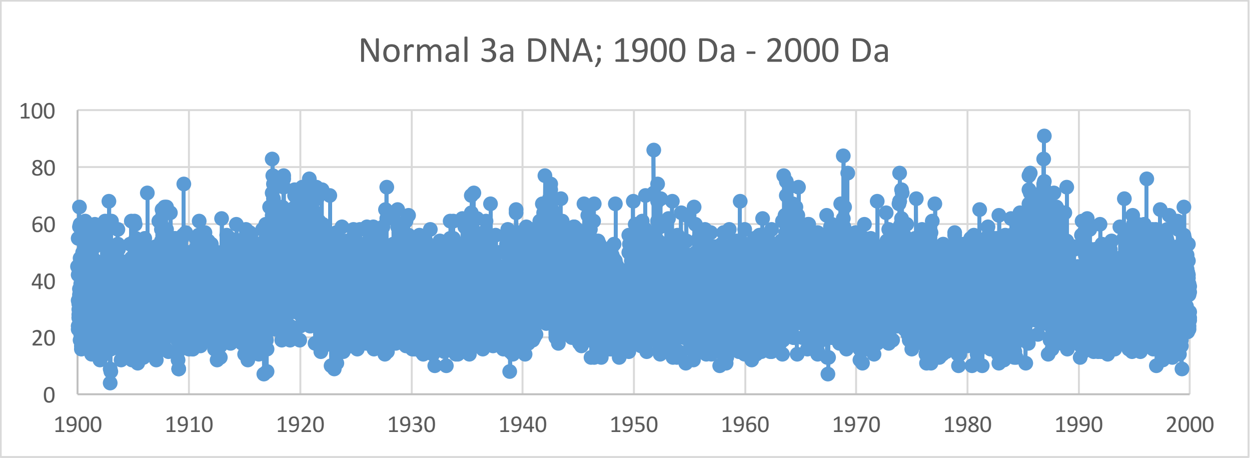
Normal 3a DNA; 1900 Da - 2000 Da
100
80
60
40
20
0
1900
1910
1920
1930
1940
1950
1960
1970
1980
1990
2000

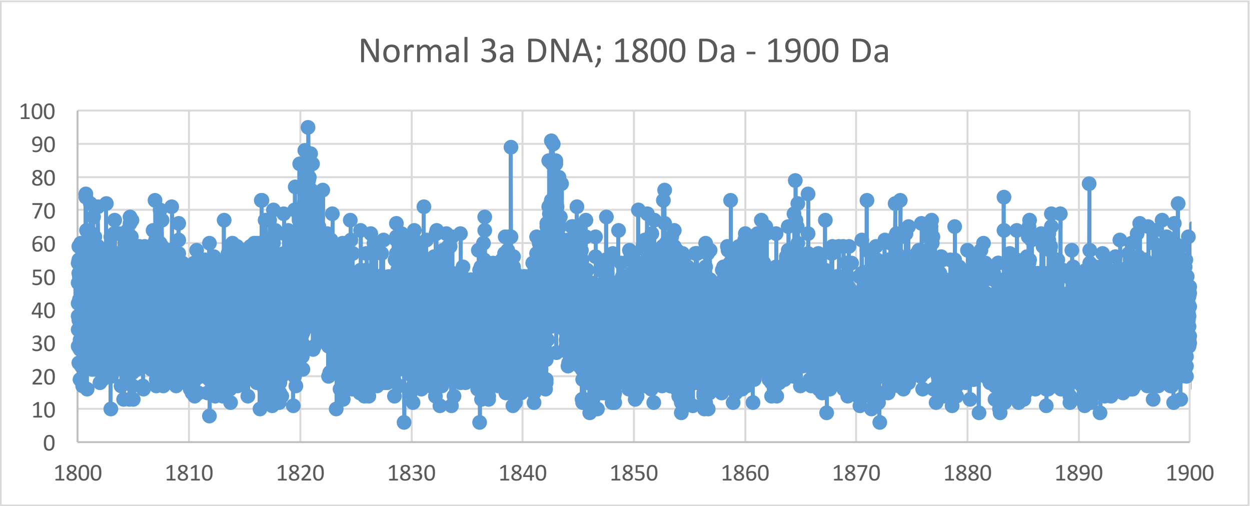
Normal 3a DNA; 1800 Da - 1900 Da
100
90
80
70
60
50
40
30
20
10
0
1800
1810
1820
1830
1840
1850
1860
1870
1880
1890
1900

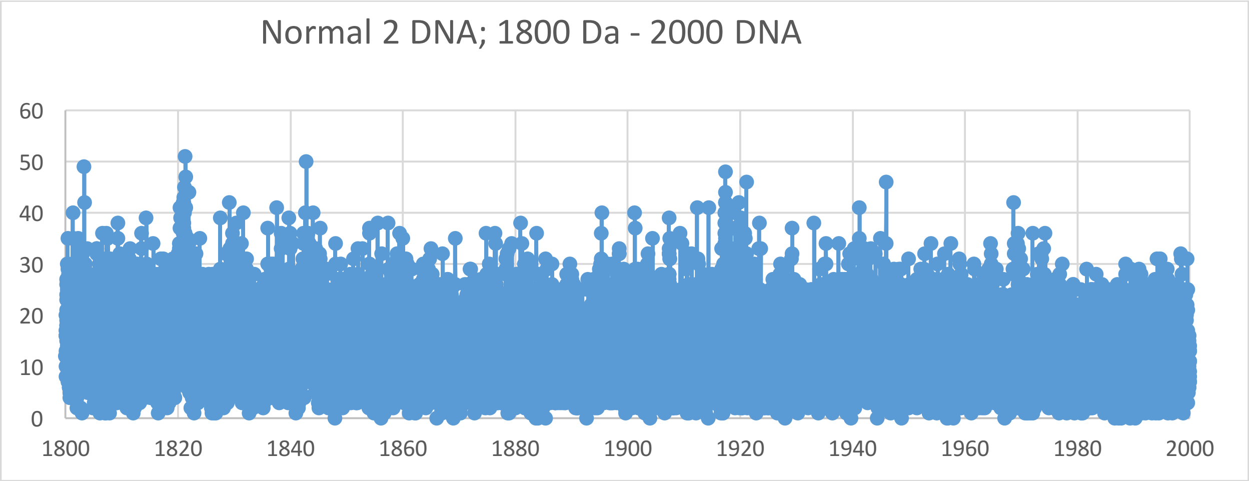
Normal 2 DNA; 1800 Da - 2000 DNA
0
10
20
30
40
50
60
1800
1820
1840
1860
1880
1900
1920
1940
1960
1980
2000

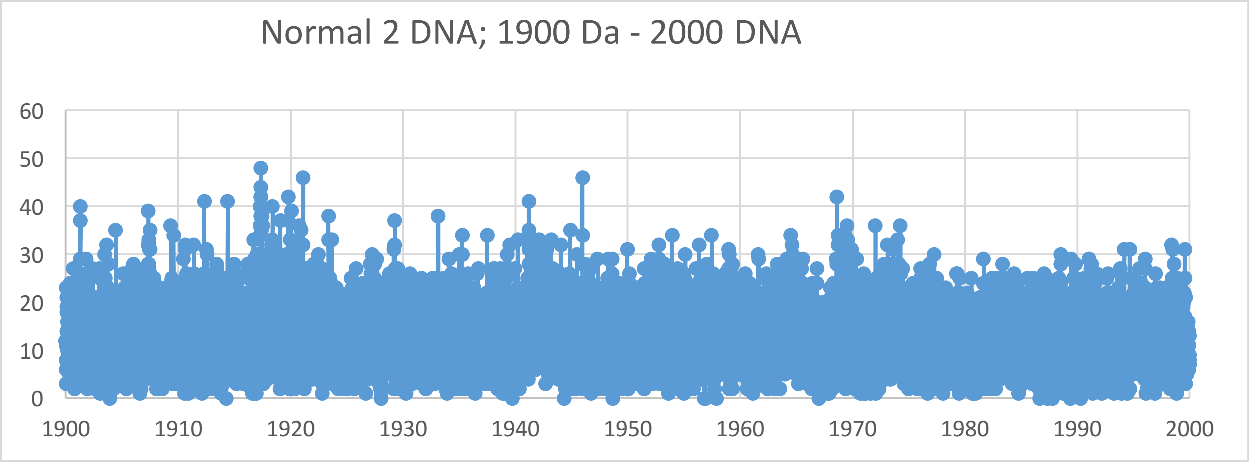
Normal 2 DNA; 1900 Da - 2000 DNA
0
10
20
30
40
50
60
1900
1910
1920
1930
1940
1950
1960
1970
1980
1990
2000

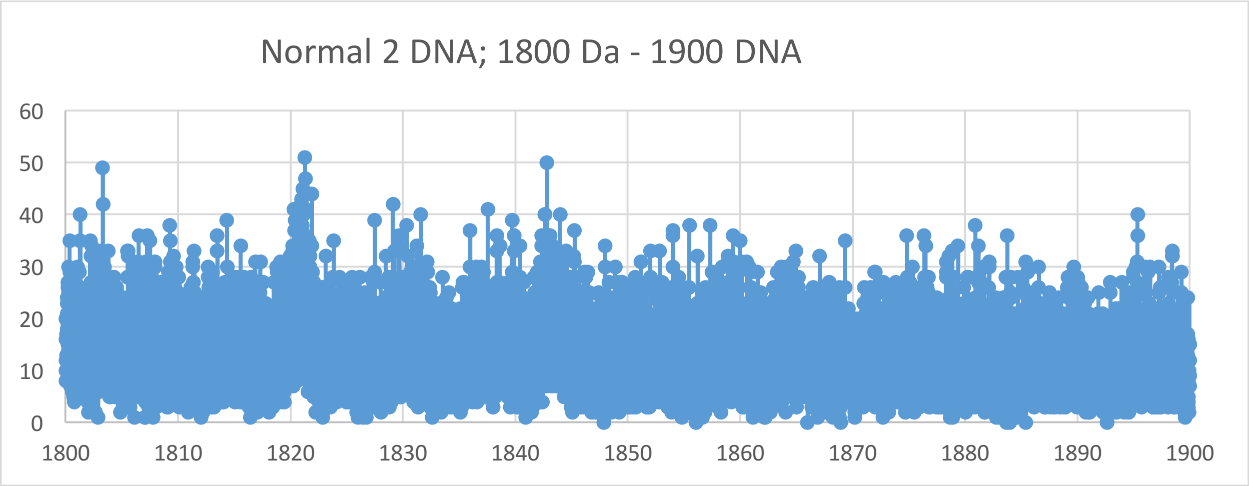
Normal 2 DNA; 1800 Da - 1900 DNA
0
10
20
30
40
50
60
1800
1810
1820
1830
1840
1850
1860
1870
1880
1890
1900

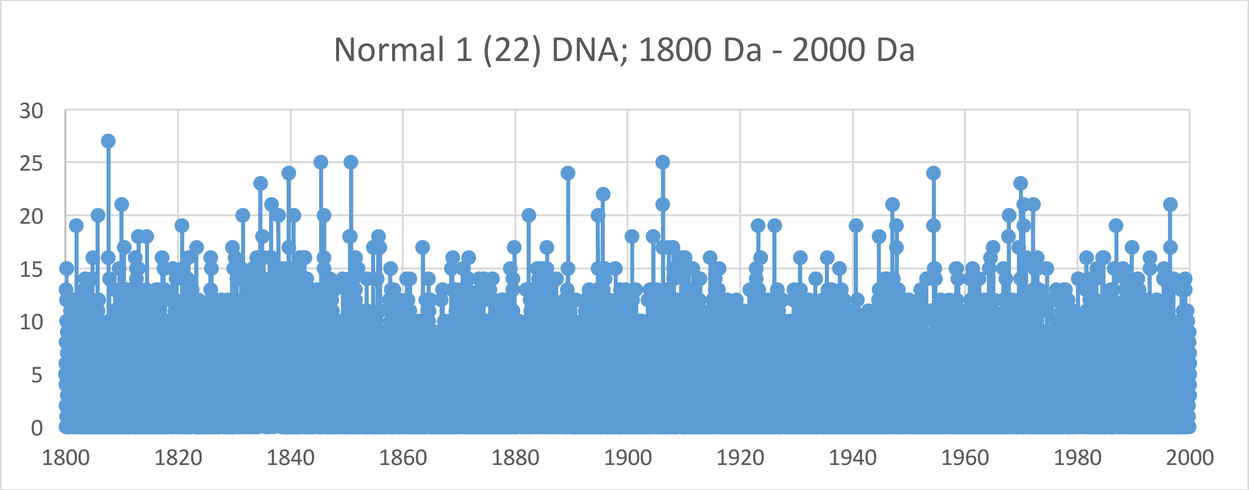
Normal 1 (22) DNA; 1800 Da - 2000 Da
0
5
10
15
20
25
30
1800
1820
1840
1860
1880
1900
1920
1940
1960
1980
2000

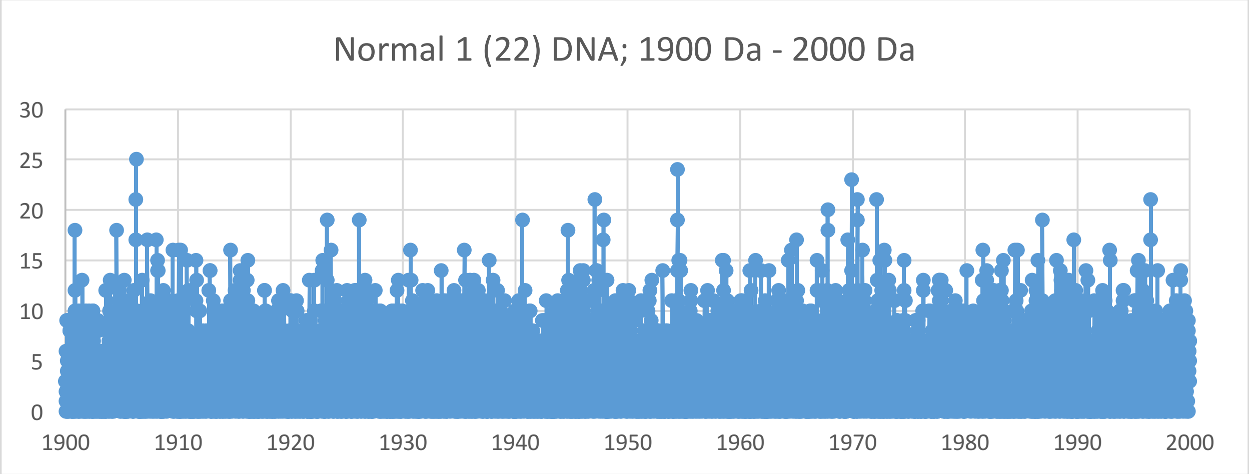
Normal 1 (22) DNA; 1900 Da - 2000 Da
0
5
10
15
20
25
30
1900
1910
1920
1930
1940
1950
1960
1970
1980
1990
2000

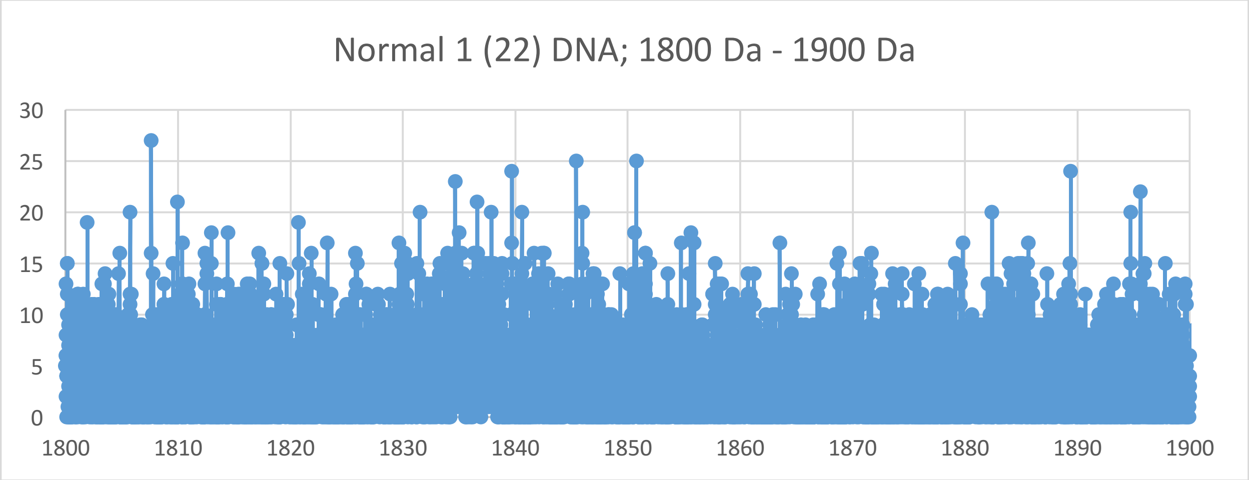
Normal 1 (22) DNA; 1800 Da - 1900 Da
0
5
10
15
20
25
30
1800
1810
1820
1830
1840
1850
1860
1870
1880
1890
1900
