## Supplementary Materials III for "Oncogenesis and Aging by Isotopic Functionalizations of the Proteins and Nucleic Acids"

III Supplementary Materials – New Mass Spectra from 1700 Da -1800 Da

Results for Supplementary 3

So for the tetranucleotides from $(phos)_{10}$ to $(phos)_9$, the peaks for cancer increase in number and the peaks for normal decrease in number for less stable normal tetranucleotides and more stable cancerous tetranucleotides with these dephosphorylations. The masses range from 1624 Da to 1879 Da. At $(phos)_{10}$ the saturate phosphorylations manifest stable cancerous normal tetranucleotides isotopically enriched, but the mating telomeres manifest cancerous isotopically enriched tetranucleotides. See Supplementary Figures III for graphs. These results and observations point specific differences between cancerous and normal oligonucleotides chemically and isotopically. The dephosphorylations from $(phos)_{10}$ to $(phos)_9$ caused the resulting more stable cancerous tetranucleotides to be more telomeric and nontelomeric and isotopically enriched for the telomeres but not for nontelomeres. The mating telomeres are not as slightly more stabilized with normal but both mixed normal and cancerous mating telomeric tetranucleotides manifest with less isotopic enrichments by these dephosphorylations. These dephosphorylations from $(phos)_{10}$ to $(phos)_9$ also caused the nontelomeric tetranucleotides to be more from cancer DNA and less isotopically enriched. It seems the dephosphorylations are related and induces by the isotopic enrichments and depletions; but the telomeres are more intense in cancer and more isotopically enriched whereas the intensities of the mating telomeres are not really altered but the isotopic enrichments are less. The telomeric tetranucleotides of cancer and normal pieces tended to also be enriched so it seems the cancer stabilities and normal stabilities of the telomeric and mating tetranucleotides are related to their isotopic enrichments for the dephosphorylations. The isotopic enrichments strengthen the telomeric cancer tetranucleotides while slightly weakening cancerous mating telomeric tetranucleotides. The more intense cancer telomeric tetranucleotides were also more isotopically enriched. These dephosphorylations from $(phos)_{10}$ to $(phos)_9$ increased the isotopic enrichments of the cancer relative to the normal for the nontelomeric tetranucleotides. But the mating telomeric tetranucleotides were less isotopically enriched from these dephosphorylations for less enrichments and stronger more stable cancer tetranucleotides for mating telomeres. Such seems compatible for replications and transcriptions as stable telomeres seem compatible to unstable mating telomeres for the unravelings and constructions of DNA mating telomeres onto the building templates of the telomeres. The dephosphorylations from $(phos)_{10}$ to $(phos)_9$ caused methylations to cause more cancer stabilities and intensities but dramatically more isotopic enrichments for telomeres. But the methylations of the mating telomeres increased cancer stabilities and less isotopic enrichments for contrary effects on mating telomeres compared to telomeres. This dephosphorylations from $(phos)_{10}$ to $(phos)_9$ caused acetylations to cause less cancer stabilities and more normal stabilities for mating telomeres with more enrichments relative to methylations; the telomeres and nontelomers are slightly more normal stable and more isotopically enriched; but the acetylations cause the mating telomers to be more cancer stable and less enriched in nonprimordials. Thereby these dephosphorylations $(phos)_{10}$ to $(phos)_9$ and their varying consequences with respect to other dephosphrylations determine patterns of dephosphorylations and isotopic enrichment patterns in normal DNA which mutate and change for different patterns in DNA of cells that become cancerous. These dephosphorylations (phos)$_{10}$ to (phos)$_9$ induced isotopically dynamics from of varying methylations and acetylations in cancer DNA relative to normal DNA couple to other isotopically induced varied phosphorylations, methylations and acetylations throughout the DNA for emerging order from nanodisorder.

So for the tetranucleotides from (phos)$_9$ to (phos)$_8$, the telomeric peaks for cancer increase in number and the peaks for normal decrease in number for less stable normal tetranucleotides and more stable cancerous tetranucleotides with these dephosphorylations. The masses range from 1551 Da to 1800 Da. See Supplementary Figures III for graphs. These results and observations point specific differences between cancerous and normal oligonucleotides chemically and isotopically. The nontelomeric and mating telomeric peaks for cancer increase with this dephosphorylations of the tetranucleotides. These dephosphorylations from (phos)$_9$ to (phos)$_8$ caused the resulting more stable normal tetranucleotides to be more nontelomeric and mating telomeric and less isotopically enriched; but the telomeres are less intense in normal tetranucleotides and less enriched isotopically. The mating telomeric are not as stabilized with cancer and normal mating telomeric tetranucleotides are less isotopic enriched by the dephosphorylations. These dephosphorylations from (phos)$_9$ to (phos)$_8$ also caused the nontelomeric tetranucleotides to be more from normal DNA and less isotopically enriched. It seems these dephosphorylations are related to the isotopic enrichments and depletions; but the telomeres are more intense in cancer and less isotopically enriched whereas the intensities of the mating telomeres are less intense in cancer and the isotopic enrichments are less. The telomeric tetranucleotides of cancer tended to also be less enriched so it seems the cancer stabilities of the telomeric tetranucleotides are related to there less isotopic enrichments for these dephosphorylations. Enrichments weaken the telomeric cancer tetranucleotides while strengthening cancerous mating telomeric teranucleotides. The more intense cancer telomeric tetranucleotides were also less isotopically enriched. These dephosphorylations from (phos)$_9$ to (phos)$_8$ decreased the isotopic enrichments of the cancer relative to the normal for the nontelomeric tetranucleotides. But the mating telomeric tetranucleotides were less enriched from the dephosphorylations for less enrichments and stronger more stable normal tetranucleotides for mating telomeres. Such seems compatible for replications and transcriptions as stable telomeres seem compatible to unstable mating telomeres for the unravelings and constructions of DNA mating telomeres onto the building templates of the telomeres. These dephosphorylations from (phos)$_9$ to (phos)$_8$ caused methylations to cause more cancer stabilities and intensities but slightly more isotopic enrichments for telomeres and (slightly for nontelomer) with large effects of the methylations at (phos)$_8$ on mating telomeric tetranucleotide as the intensities of normal increased. But the methylations of the mating telomeres increased cancer stabilities and less isotopic enrichments for contrary effects on mating telomeres compared to telomeres. These dephosphorylations from (phos)$_9$ to (phos)$_8$ caused acetylations to cause more normal stabilities and less cancer stabilities for mating non-telomeres with more enrichments relative to methylations; the telomeres and nontelomers are more cancer stable and more isotopically enriched. Thereby this dephosphorylations (phos)$_9$ to (phos)$_8$ and their varying consequences with respect to other dephosphorylations determine patterns of dephosphorylations and isotopic enrichment patterns in normal DNA which mutate and change for different patterns in DNA of cells that become cancerous. These dephosphorylations $(phos)_9$ to $(phos)_8$ induced isotopic dynamics from of varying methylations and acetylations in cancer DNA relative to normal DNA couple to other isotopically induced varied phosphorylations, methylations and acetylations throughout the DNA for emerging order from nanodisorder.

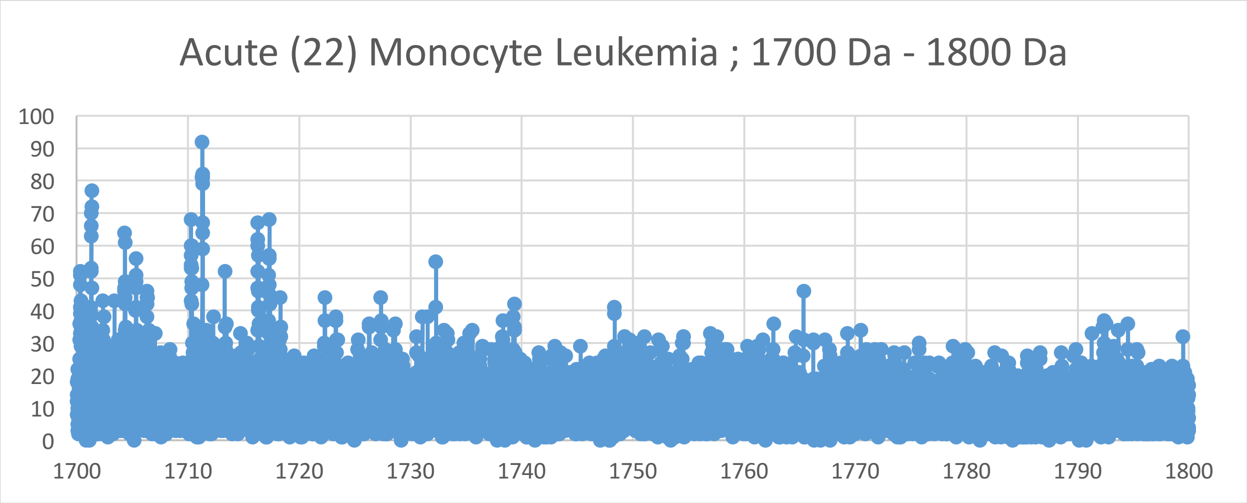
Acute (22) Monocyte Leukemia ; 1700 Da - 1800 Da
100
90
80
70
60
50
40
30
20
10
0
1700
1710
1720
1730
1740
1750
1760
1770
1780
1790
1800

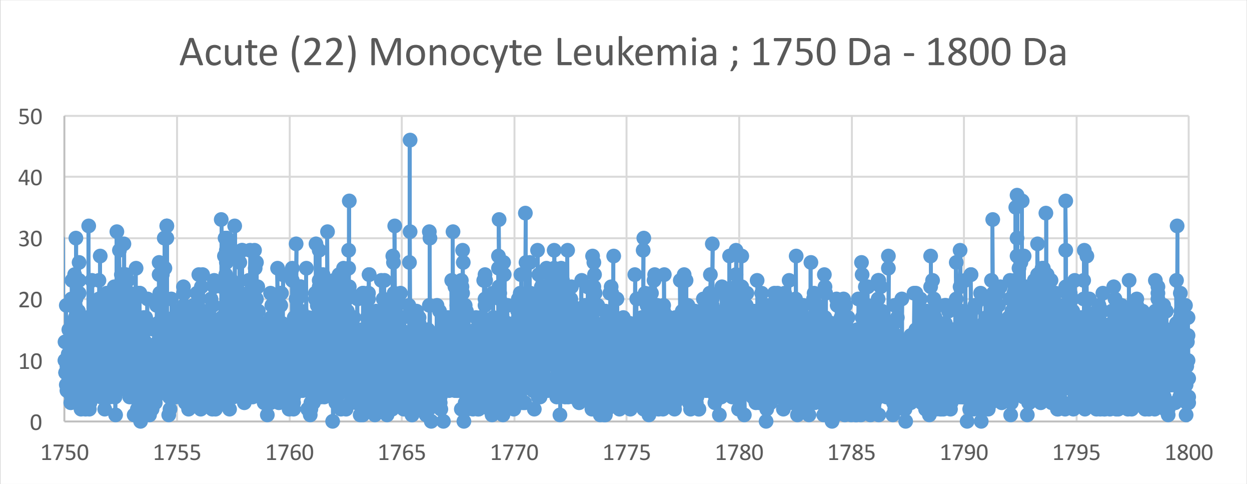
Acute (22) Monocyte Leukemia ; 1750 Da - 1800 Da
50
40
30
20
10
0
1750
1755
1760
1765
1770
1775
1780
1785
1790
1795
1800

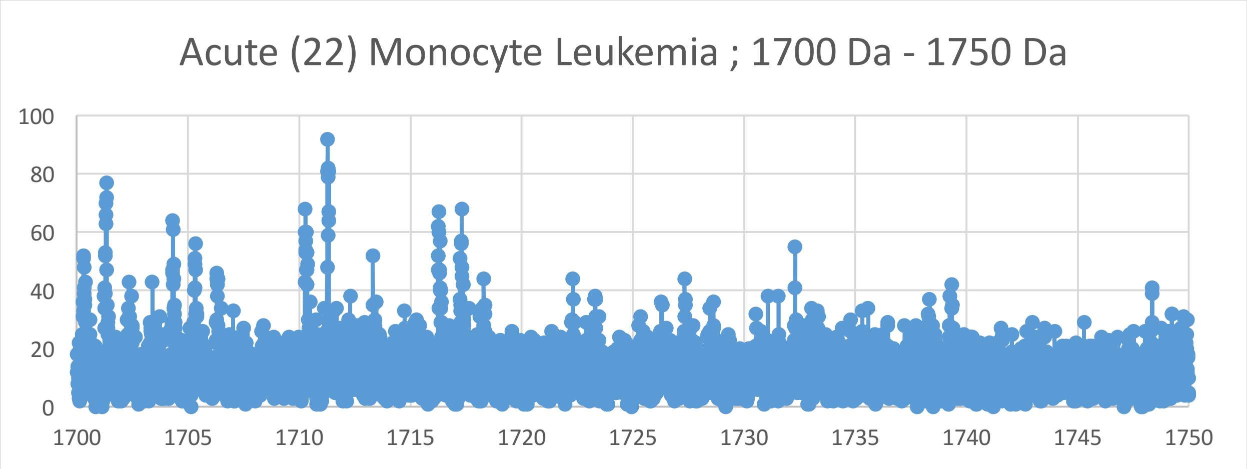
Acute (22) Monocyte Leukemia ; 1700 Da - 1750 Da
100
80
60
40
20
0
1700
1705
1710
1715
1720
1725
1730
1735
1740
1745
1750

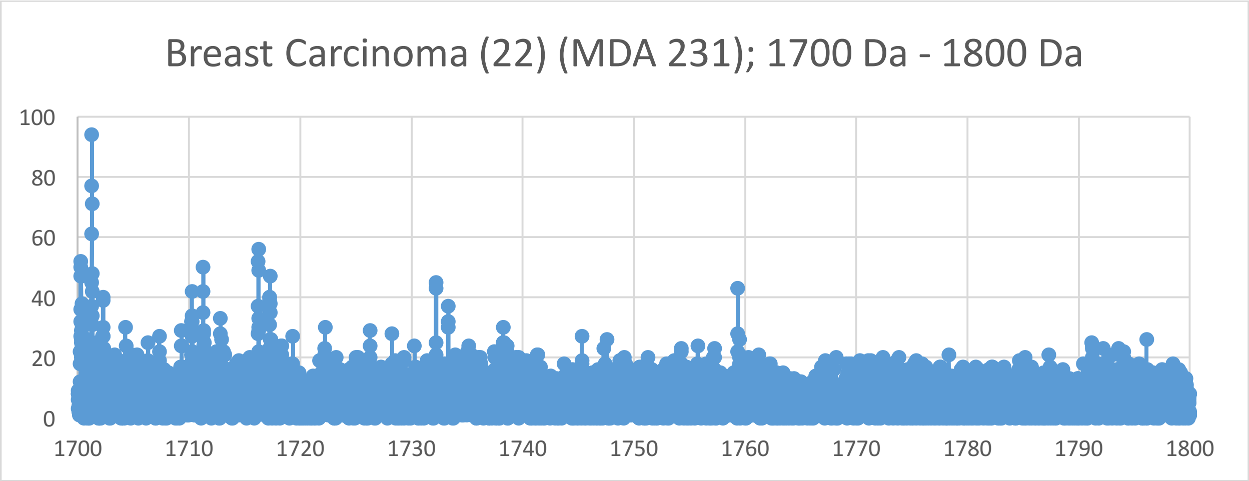
Breast Carcinoma (22) (MDA 231); 1700 Da - 1800 Da
100
80
60
40
20
0
1700
1710
1720
1730
1740
1750
1760
1770
1780
1790
1800

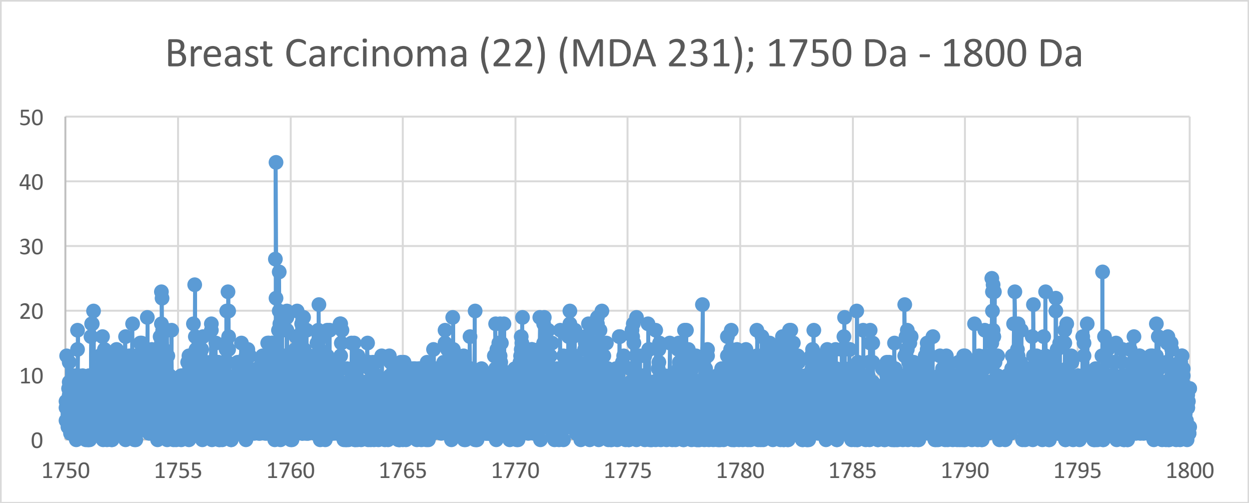
Breast Carcinoma (22) (MDA 231); 1750 Da - 1800 Da
50
40
30
20
10
0
1750
1755
1760
1765
1770
1775
1780
1785
1790
1795
1800

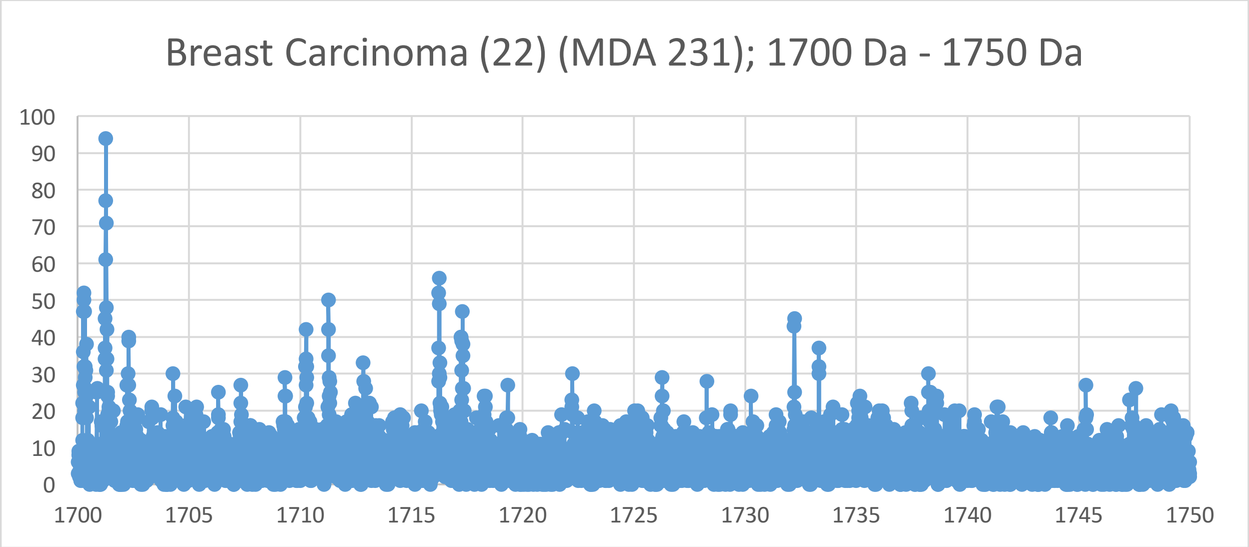
Breast Carcinoma (22) (MDA 231); 1700 Da - 1750 Da
100
90
80
70
60
50
40
30
20
10
0
1700
1705
1710
1715
1720
1725
1730
1735
1740
1745
1750

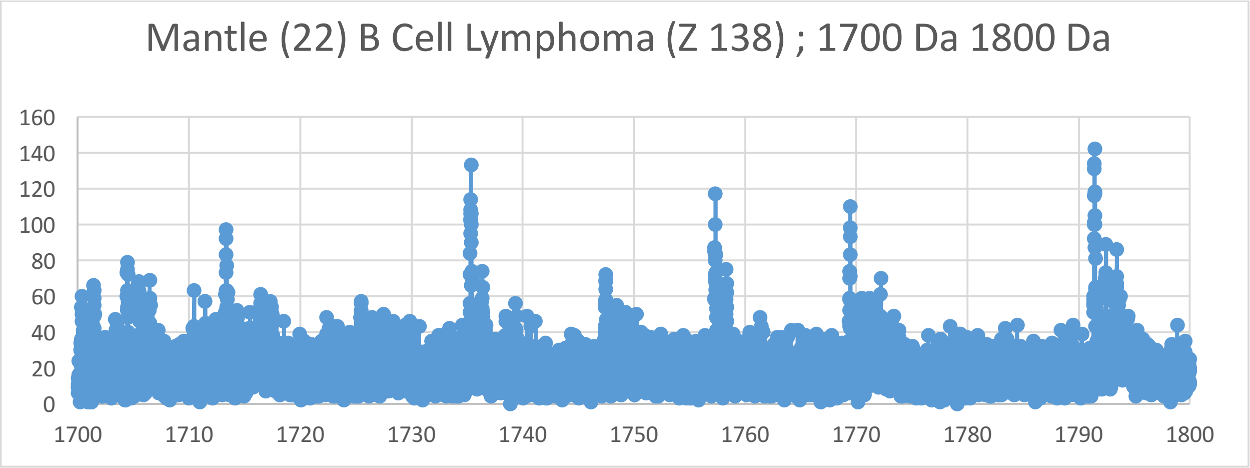
Mantle (22) B Cell Lymphoma (Z 138) ; 1700 Da 1800 Da
160
140
120
100
80
60
40
20
0
1700
1710
1720
1730
1740
1750
1760
1770
1780
1790
1800

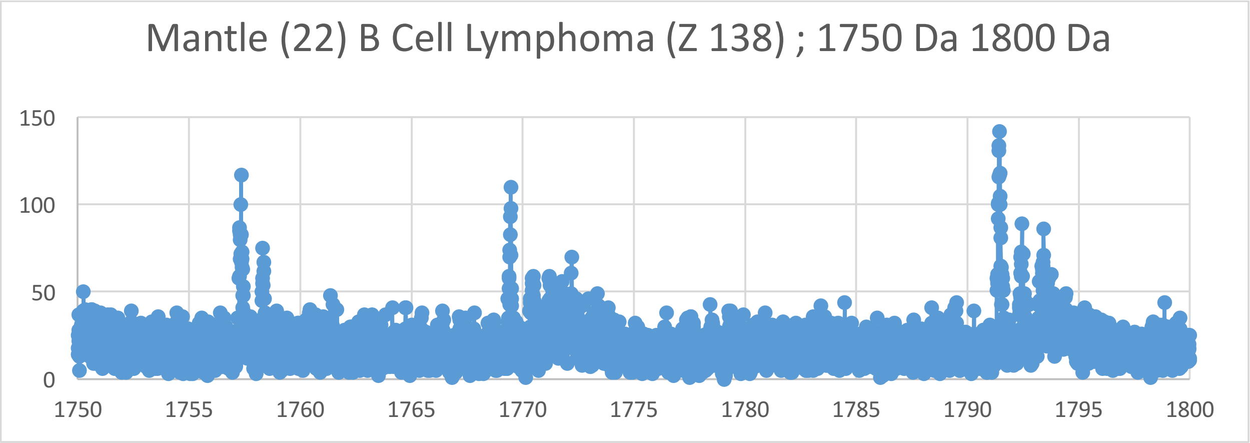
Mantle (22) B Cell Lymphoma (Z 138) ; 1750 Da 1800 Da
150
100
50
0
1750
1755
1760
1765
1770
1775
1780
1785
1790
1795
1800

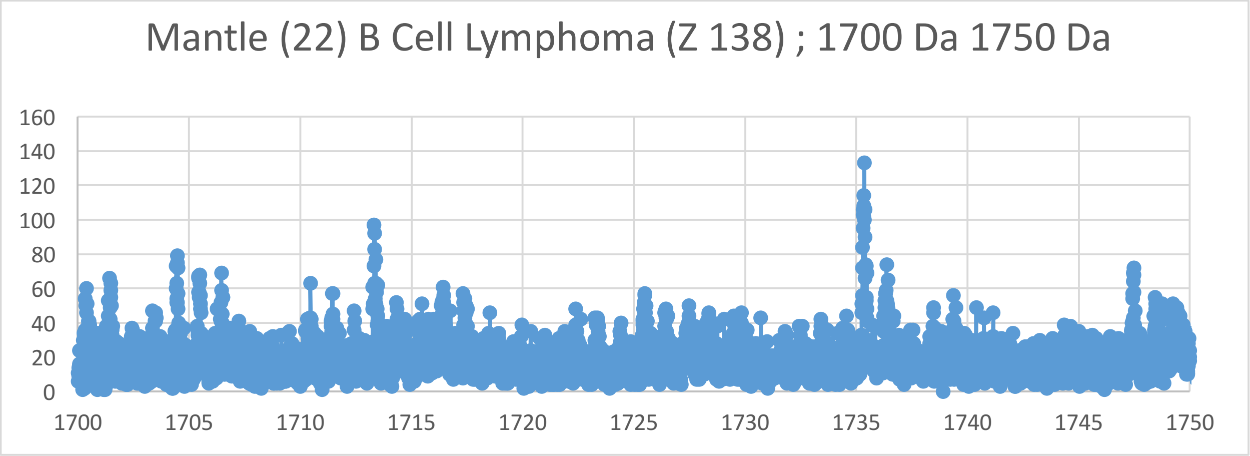
Mantle (22) B Cell Lymphoma (Z 138) ; 1700 Da 1750 Da
160
140
120
100
80
60
40
20
0
1700
1705
1710
1715
1720
1725
1730
1735
1740
1745
1750

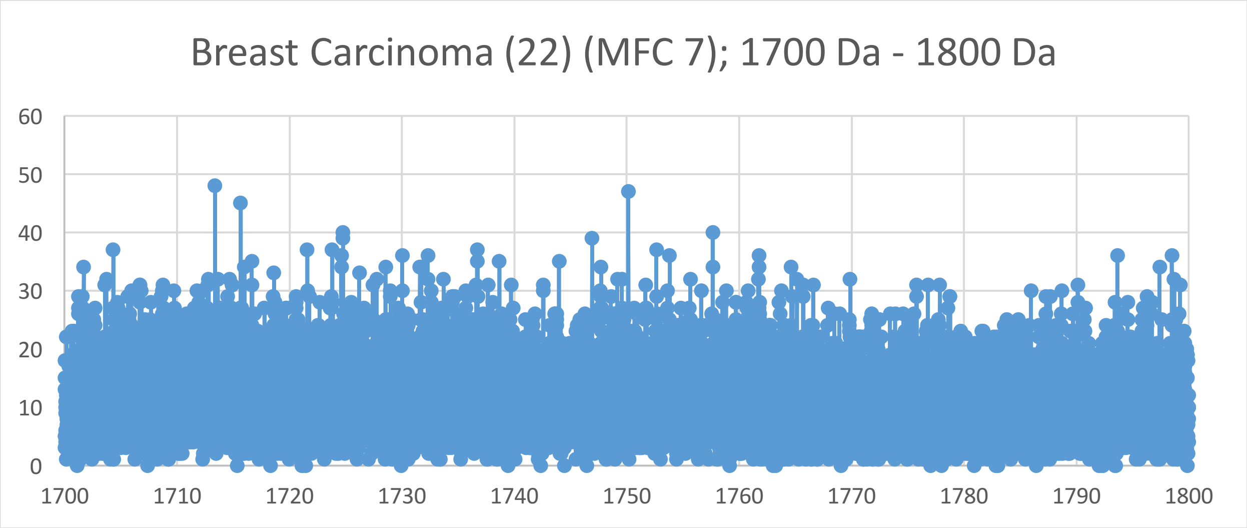
Breast Carcinoma (22) (MFC 7); 1700 Da - 1800 Da
60
50
40
30
20
10
0
1700
1710
1720
1730
1740
1750
1760
1770
1780
1790
1800

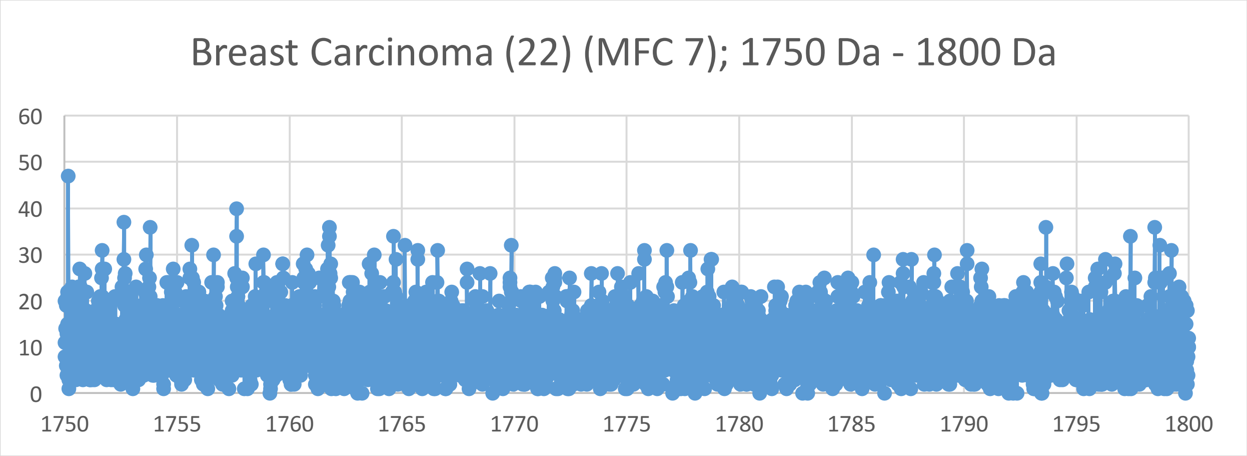
Breast Carcinoma (22) (MFC 7); 1750 Da - 1800 Da
60
50
40
30
20
10
0
1750
1755
1760
1765
1770
1775
1780
1785
1790
1795
1800

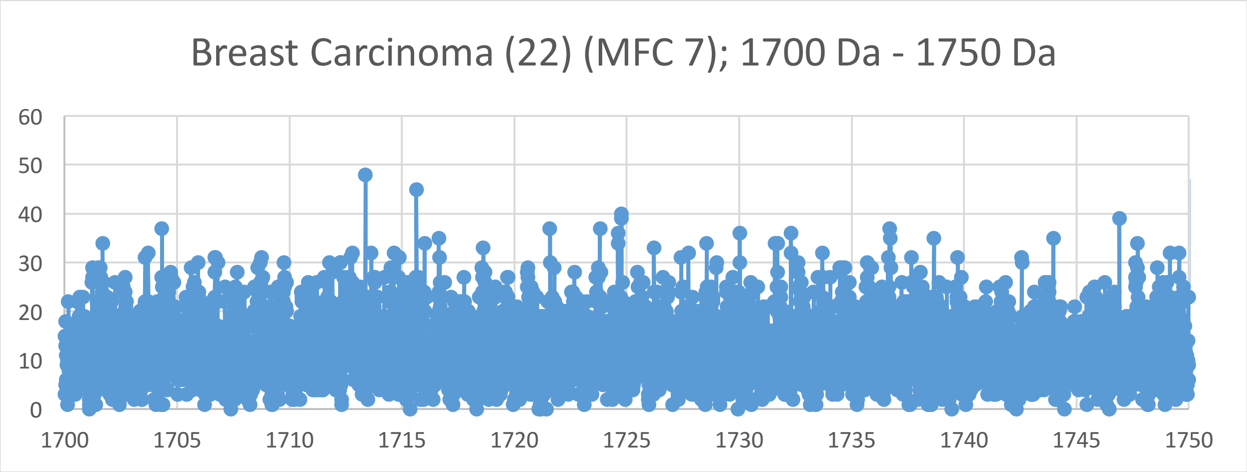
Breast Carcinoma (22) (MFC 7); 1700 Da - 1750 Da
60
50
40
30
20
10
0
1700
1705
1710
1715
1720
1725
1730
1735
1740
1745
1750

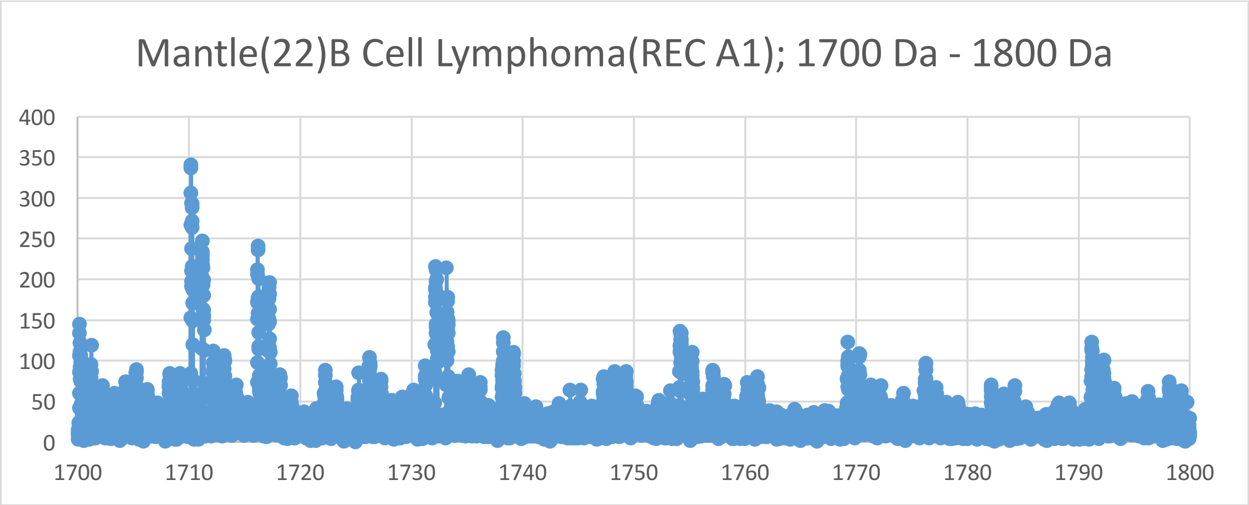
Mantle(22)B Cell Lymphoma(REC A1); 1700 Da - 1800 Da
400
350
300
250
200
150
100
50
0
1700
1710
1720
1730
1740
1750
1760
1770
1780
1790
1800

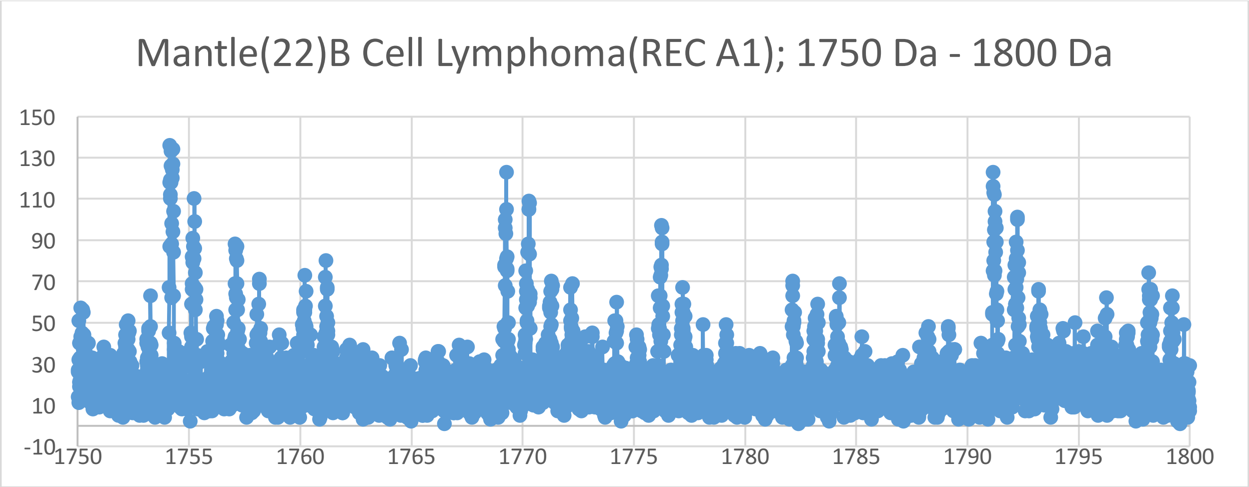
Mantle(22)B Cell Lymphoma(REC A1); 1750 Da - 1800 Da
150
130
110
90
70
50
30
10
-10
1750
1755
1760
1765
1770
1775
1780
1785
1790
1795
1800

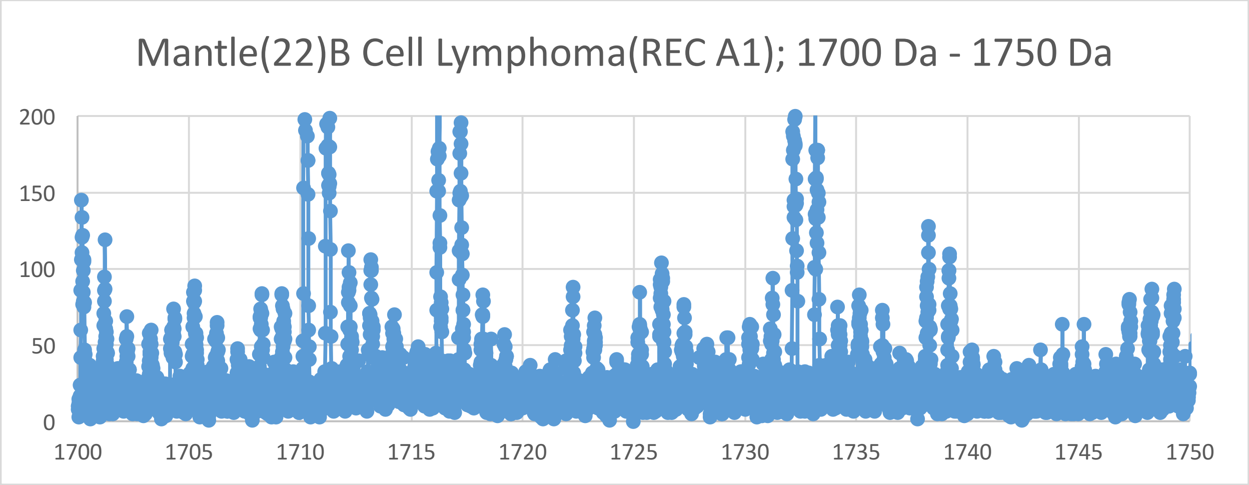
Mantle(22)B Cell Lymphoma(REC A1); 1700 Da - 1750 Da
200
150
100
50
0
1700
1705
1710
1715
1720
1725
1730
1735
1740
1745
1750

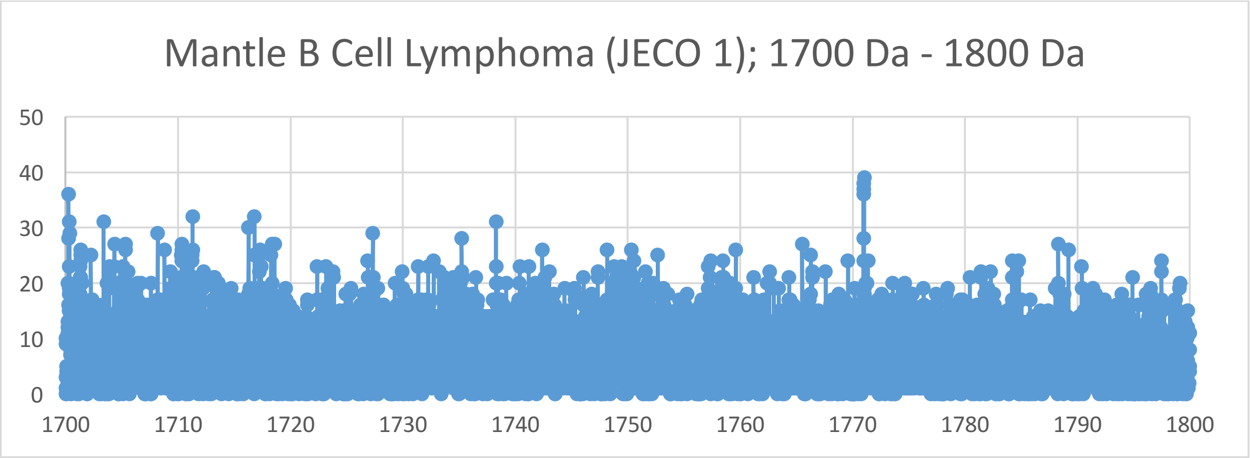
Mantle B Cell Lymphoma (JECO 1); 1700 Da - 1800 Da
50
40
30
20
10
0
1700
1710
1720
1730
1740
1750
1760
1770
1780
1790
1800

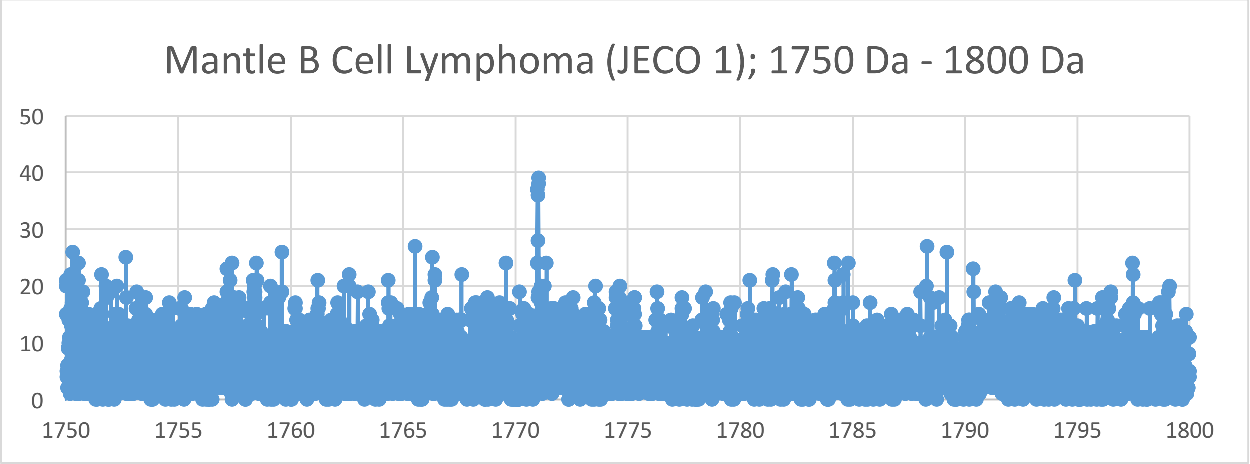
Mantle B Cell Lymphoma (JECO 1); 1750 Da - 1800 Da
50
40
30
20
10
0
1750
1755
1760
1765
1770
1775
1780
1785
1790
1795
1800

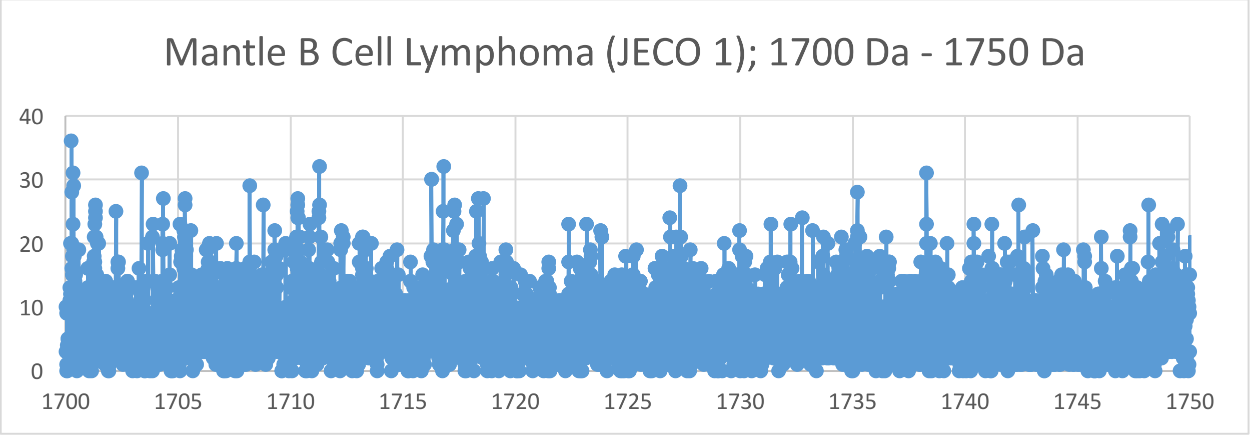
Mantle B Cell Lymphoma (JECO 1); 1700 Da - 1750 Da
40
30
20
10
0
1700
1705
1710
1715
1720
1725
1730
1735
1740
1745
1750

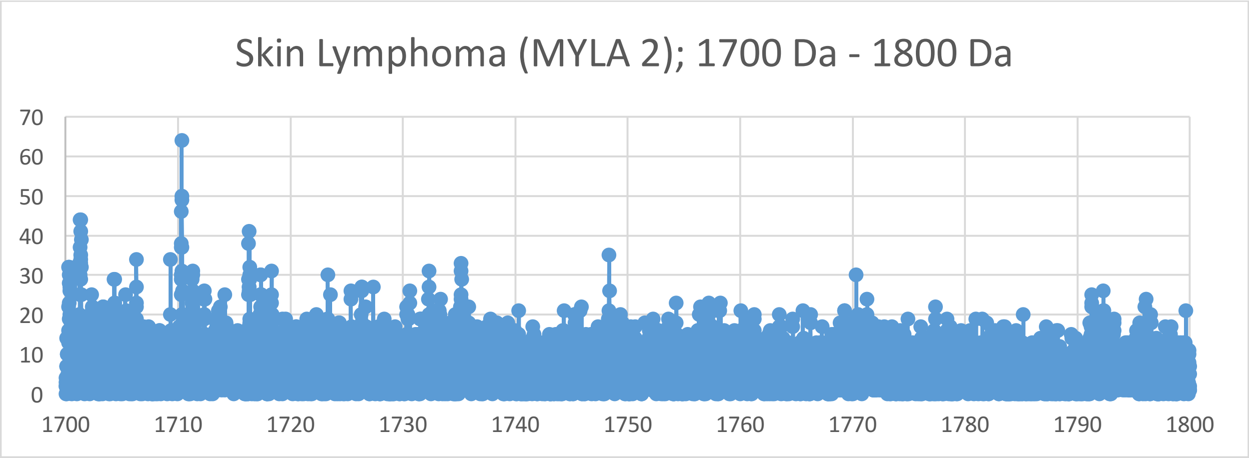
Skin Lymphoma (MYLA 2); 1700 Da - 1800 Da
70
60
50
40
30
20
10
0
1700
1710
1720
1730
1740
1750
1760
1770
1780
1790
1800

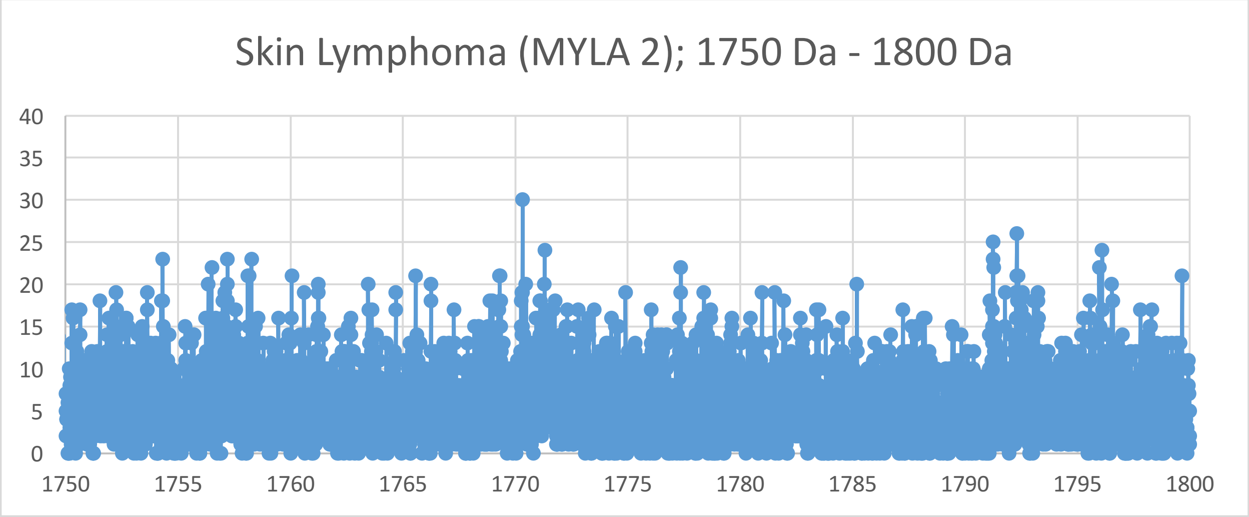
Skin Lymphoma (MYLA 2); 1750 Da - 1800 Da
40
35
30
25
20
15
10
5
0
1750
1755
1760
1765
1770
1775
1780
1785
1790
1795
1800

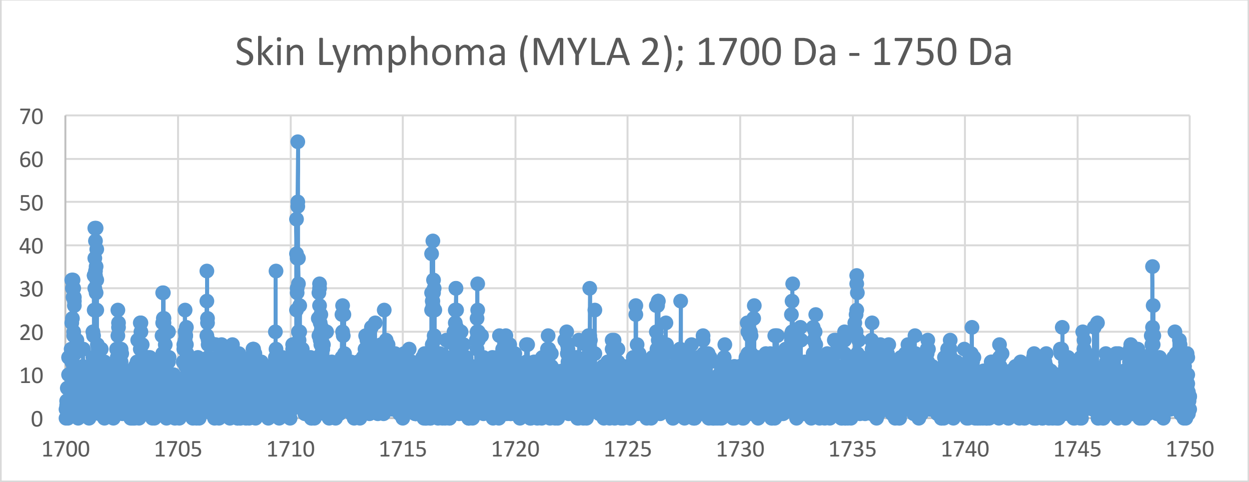
Skin Lymphoma (MYLA 2); 1700 Da - 1750 Da
70
60
50
40
30
20
10
0
1700
1705
1710
1715
1720
1725
1730
1735
1740
1745
1750

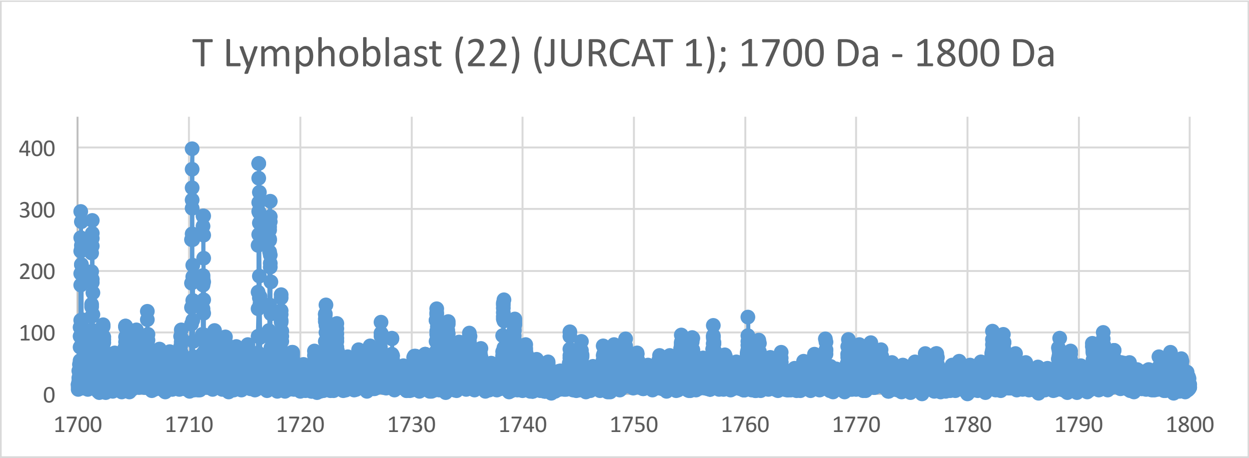
T Lymphoblast (22) (JURCAT 1); 1700 Da - 1800 Da
0
100
200
300
400
1700
1710
1720
1730
1740
1750
1760
1770
1780
1790
1800

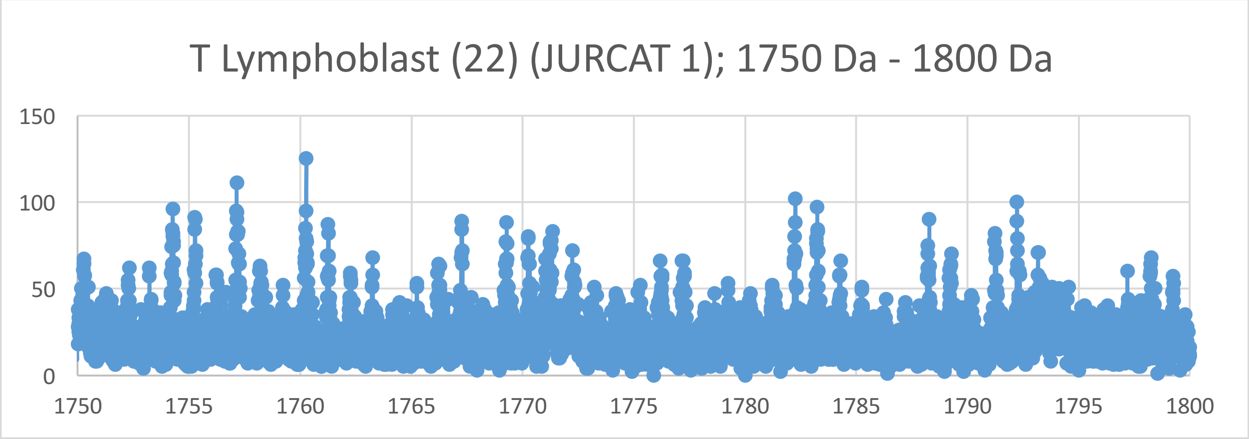
T Lymphoblast (22) (JURCAT 1); 1750 Da - 1800 Da
0
50
100
150
1750
1755
1760
1765
1770
1775
1780
1785
1790
1795
1800

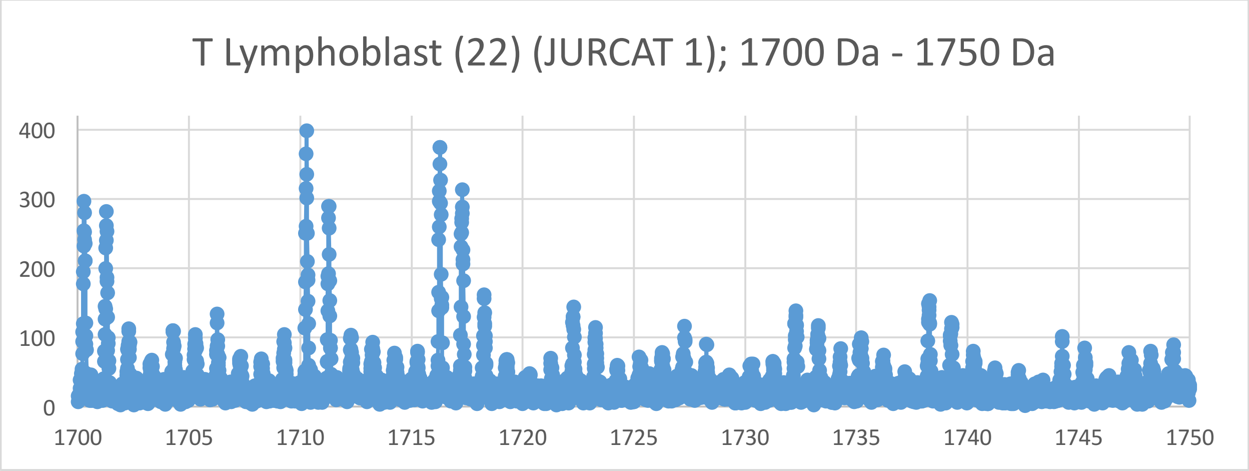
T Lymphoblast (22) (JURCAT 1); 1700 Da - 1750 Da
0
100
200
300
400
1700
1705
1710
1715
1720
1725
1730
1735
1740
1745
1750

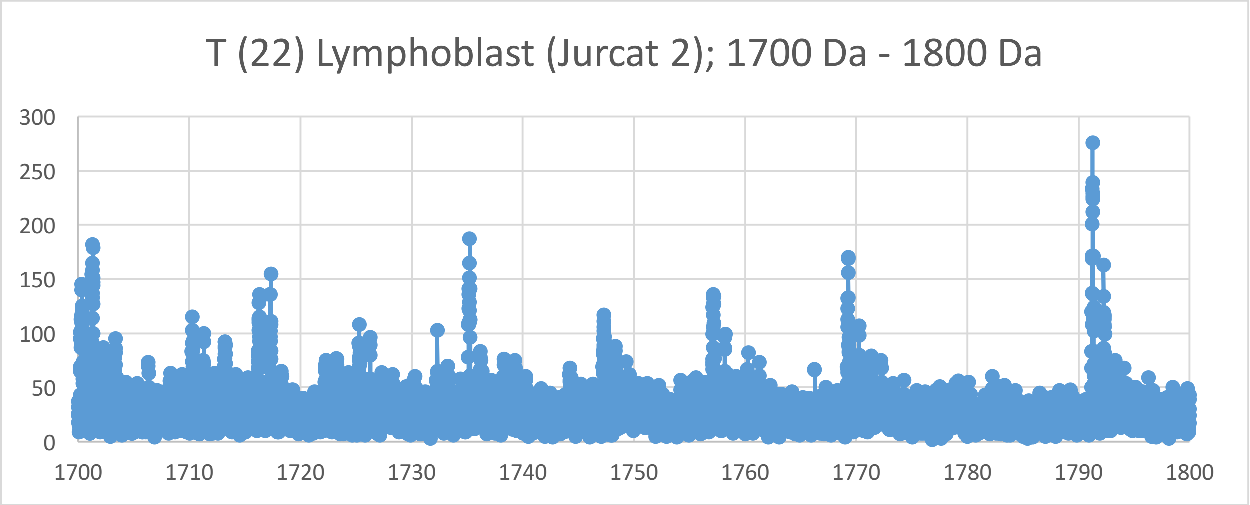
T (22) Lymphoblast (Jurcat 2); 1700 Da - 1800 Da
300
250
200
150
100
50
0
1700
1710
1720
1730
1740
1750
1760
1770
1780
1790
1800

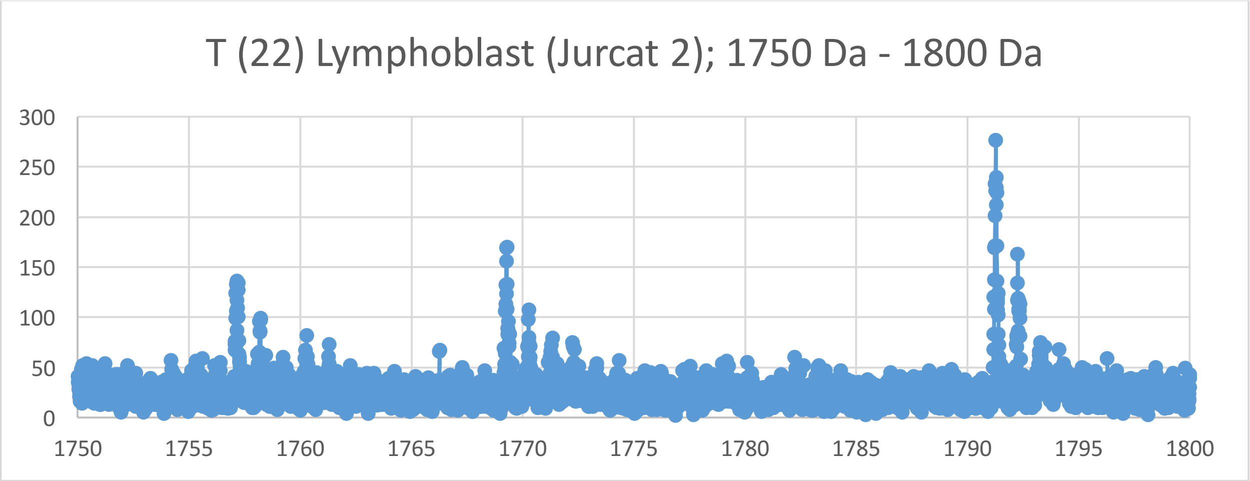
T (22) Lymphoblast (Jurcat 2); 1750 Da - 1800 Da
300
250
200
150
100
50
0
1750
1755
1760
1765
1770
1775
1780
1785
1790
1795
1800

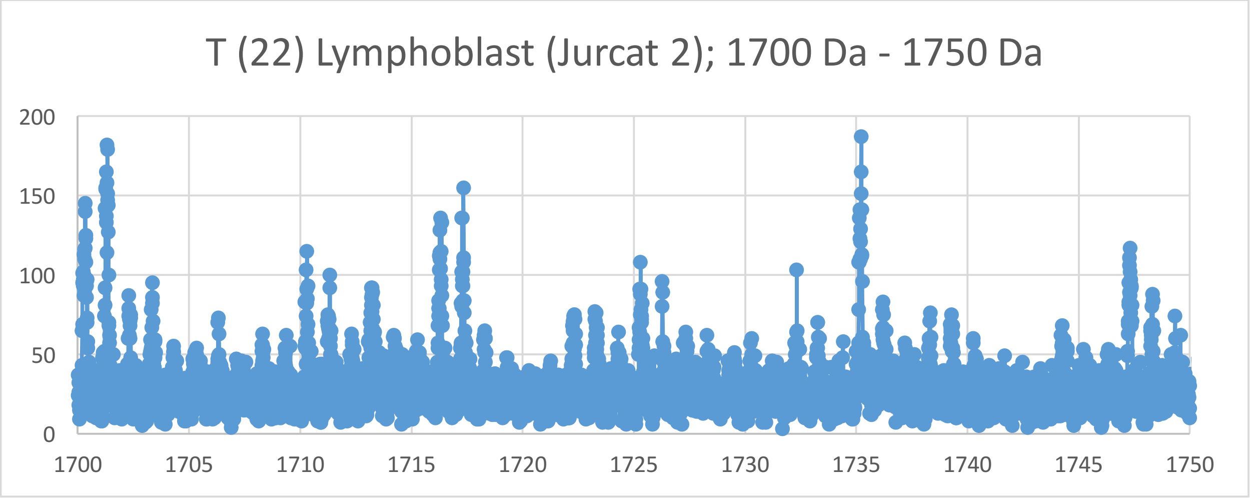
T (22) Lymphoblast (Jurcat 2); 1700 Da - 1750 Da
200
150
100
50
0
1700
1705
1710
1715
1720
1725
1730
1735
1740
1745
1750

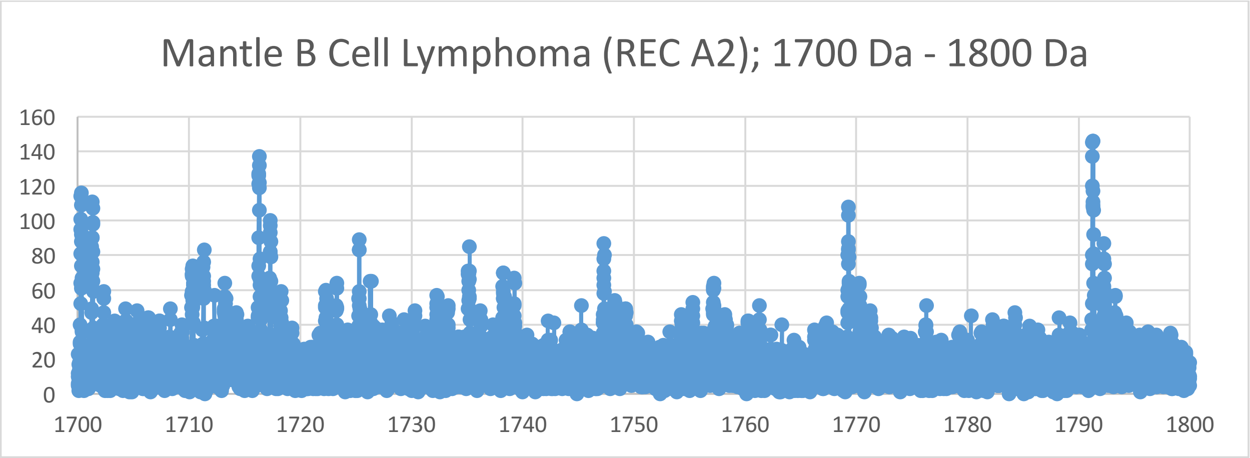
Mantle B Cell Lymphoma (REC A2); 1700 Da - 1800 Da
160
140
120
100
80
60
40
20
0
1700
1710
1720
1730
1740
1750
1760
1770
1780
1790
1800

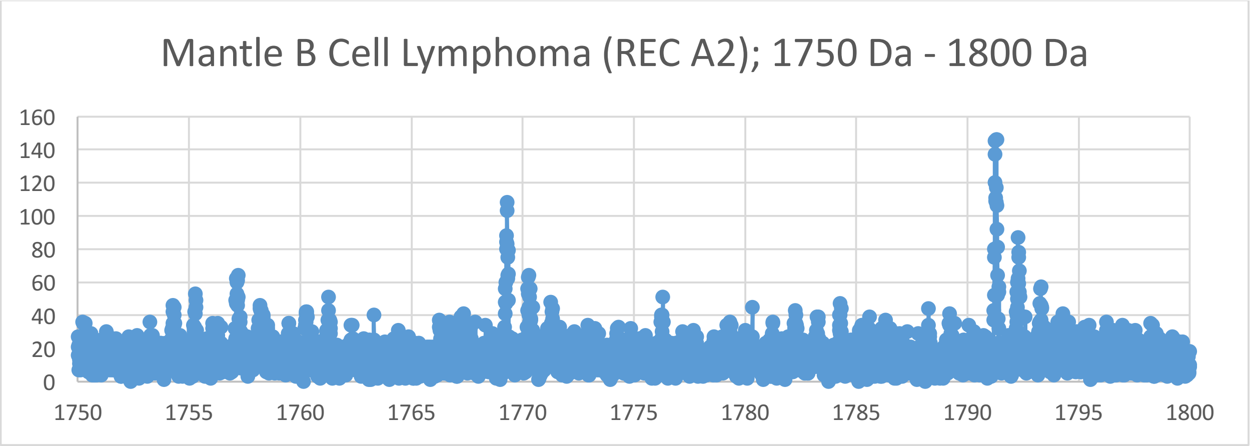
Mantle B Cell Lymphoma (REC A2); 1750 Da - 1800 Da
160
140
120
100
80
60
40
20
0
1750
1755
1760
1765
1770
1775
1780
1785
1790
1795
1800

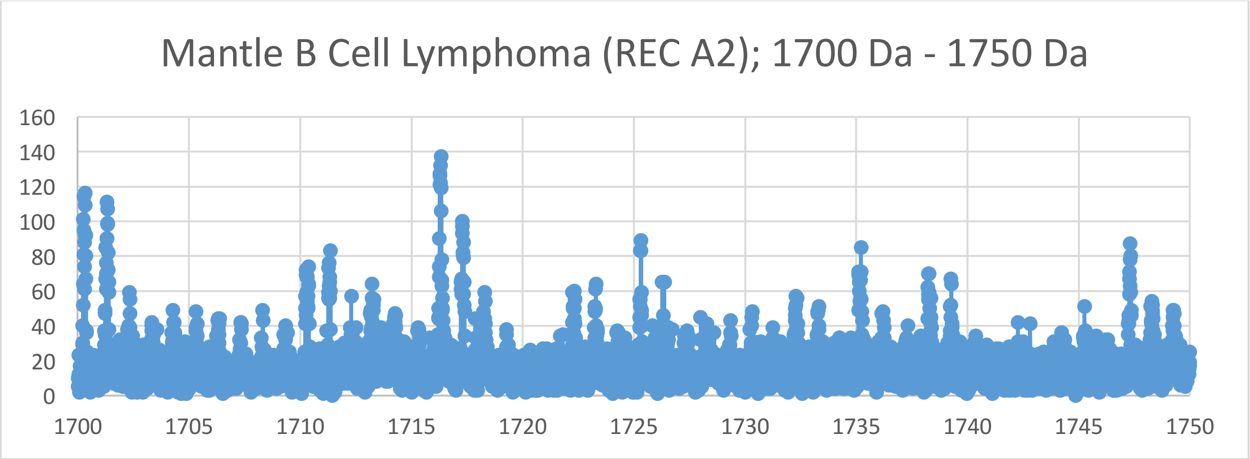
Mantle B Cell Lymphoma (REC A2); 1700 Da - 1750 Da
160
140
120
100
80
60
40
20
0
1700
1705
1710
1715
1720
1725
1730
1735
1740
1745
1750

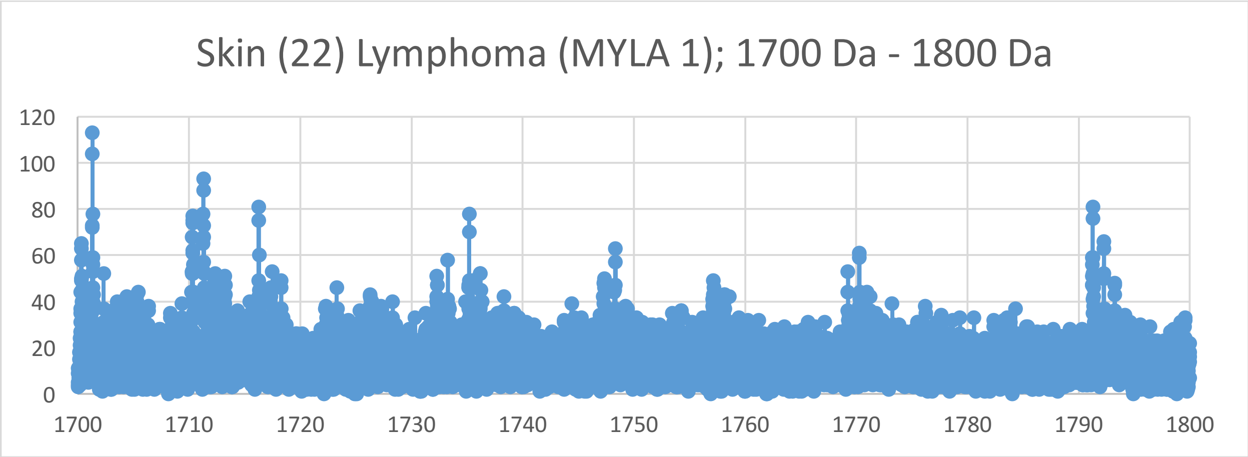
Skin (22) Lymphoma (MYLA 1); 1700 Da - 1800 Da
120
100
80
60
40
20
0
1700
1710
1720
1730
1740
1750
1760
1770
1780
1790
1800

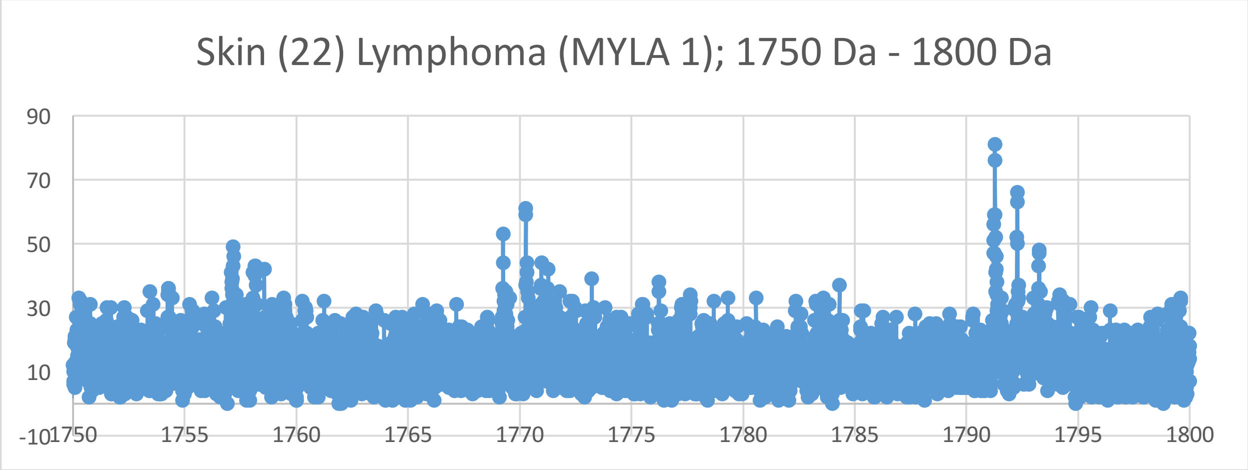
Skin (22) Lymphoma (MYLA 1); 1750 Da - 1800 Da
90
70
50
30
10
-10
1750
1755
1760
1765
1770
1775
1780
1785
1790
1795
1800

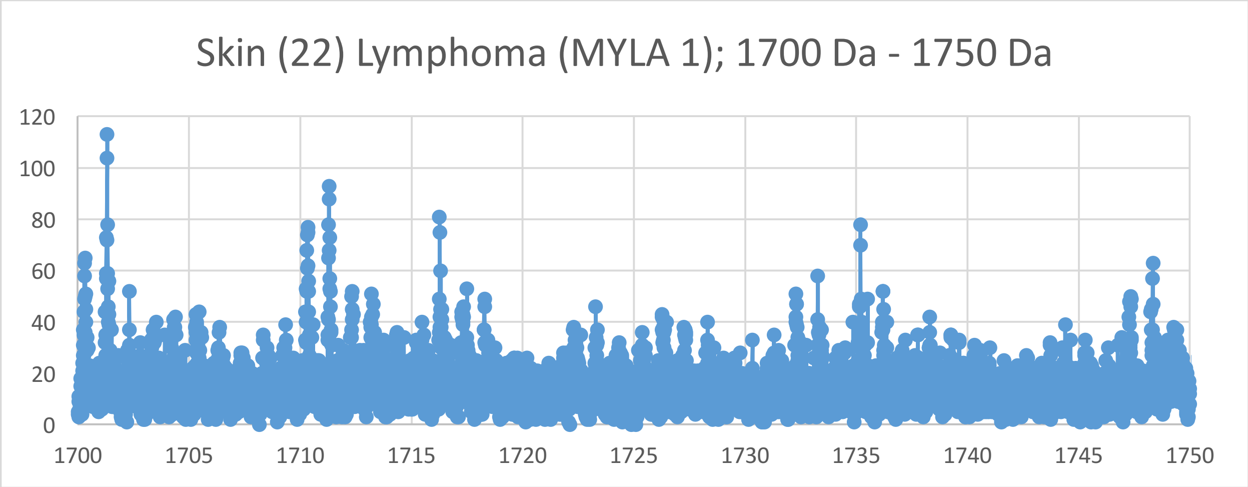
Skin (22) Lymphoma (MYLA 1); 1700 Da - 1750 Da
120
100
80
60
40
20
0
1700
1705
1710
1715
1720
1725
1730
1735
1740
1745
1750

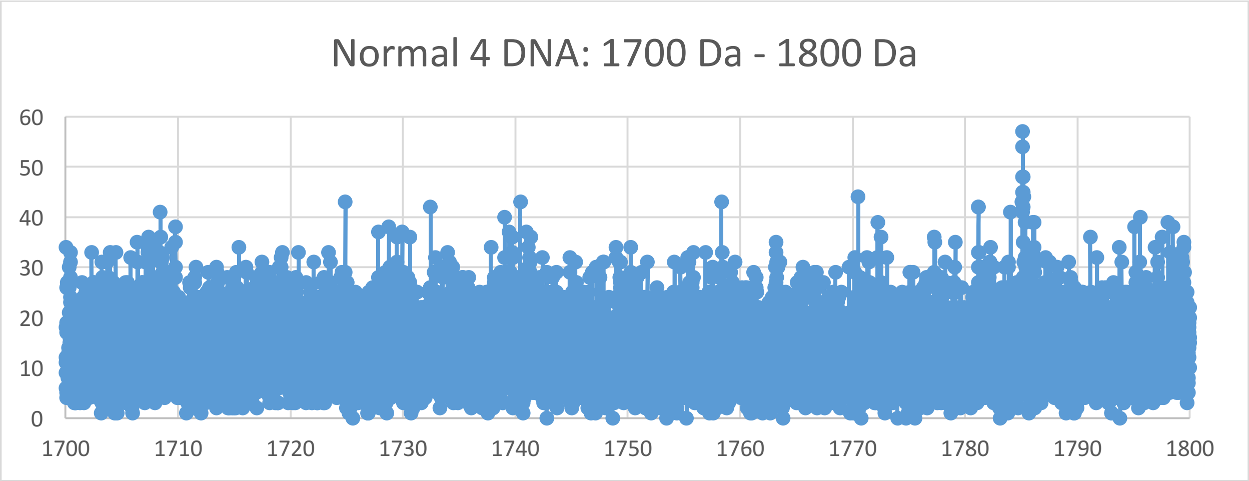
Normal 4 DNA: 1700 Da - 1800 Da
60
50
40
30
20
10
0
1700
1710
1720
1730
1740
1750
1760
1770
1780
1790
1800

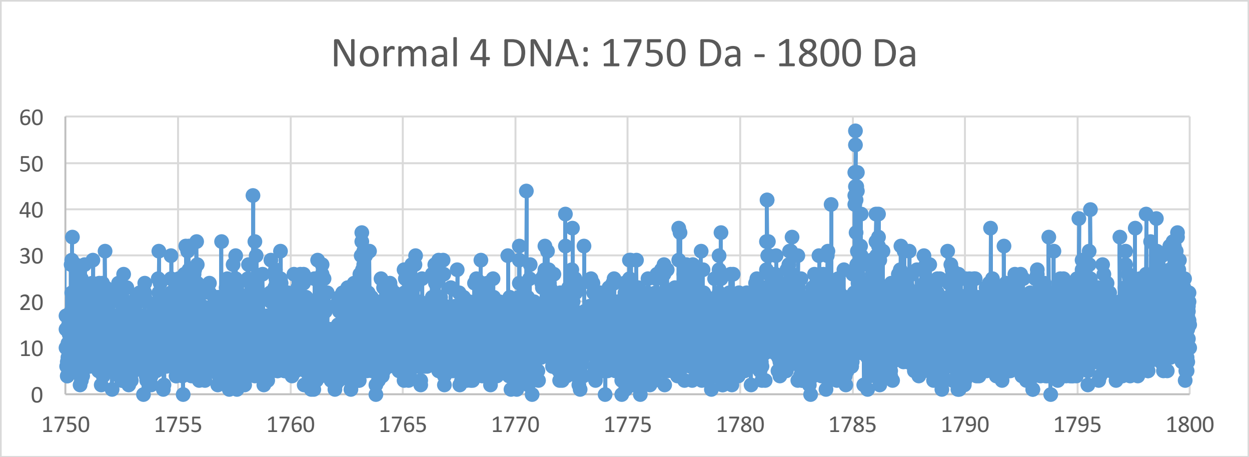
Normal 4 DNA: 1750 Da - 1800 Da
60
50
40
30
20
10
0
1750
1755
1760
1765
1770
1775
1780
1785
1790
1795
1800

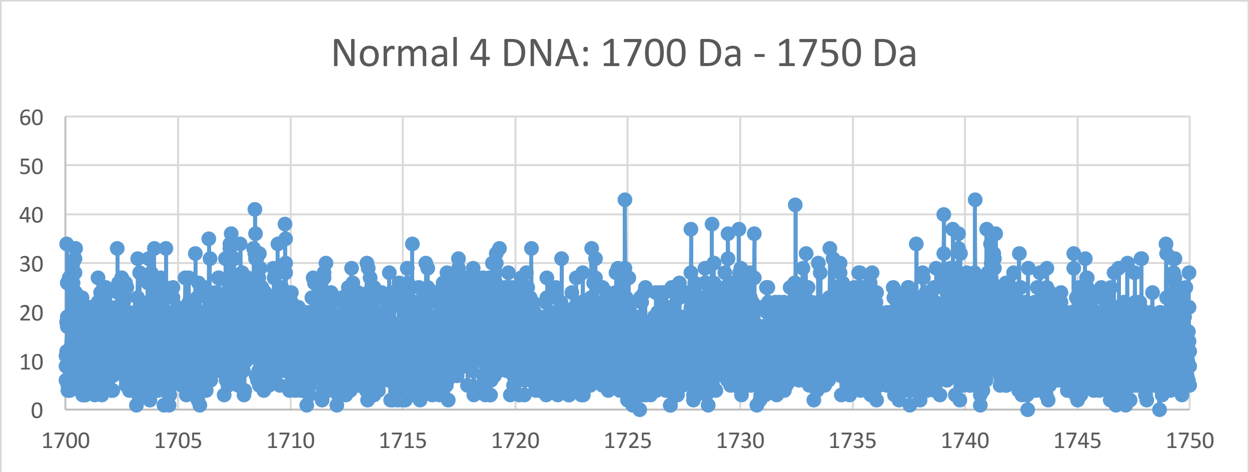
Normal 4 DNA: 1700 Da - 1750 Da
60
50
40
30
20
10
0
1700
1705
1710
1715
1720
1725
1730
1735
1740
1745
1750

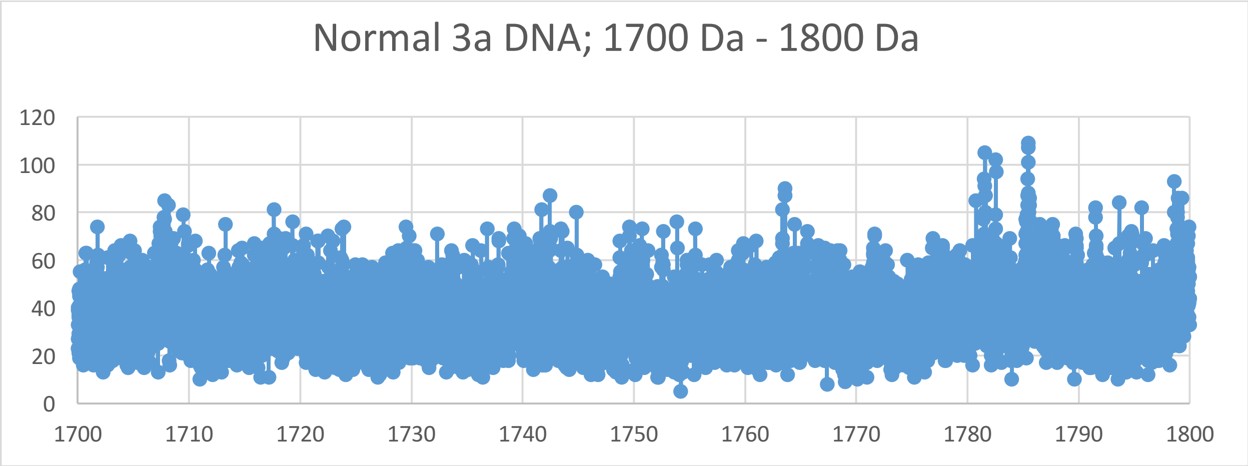
Normal 3a DNA; 1700 Da - 1800 Da
120
100
80
60
40
20
0
1700
1710
1720
1730
1740
1750
1760
1770
1780
1790
1800

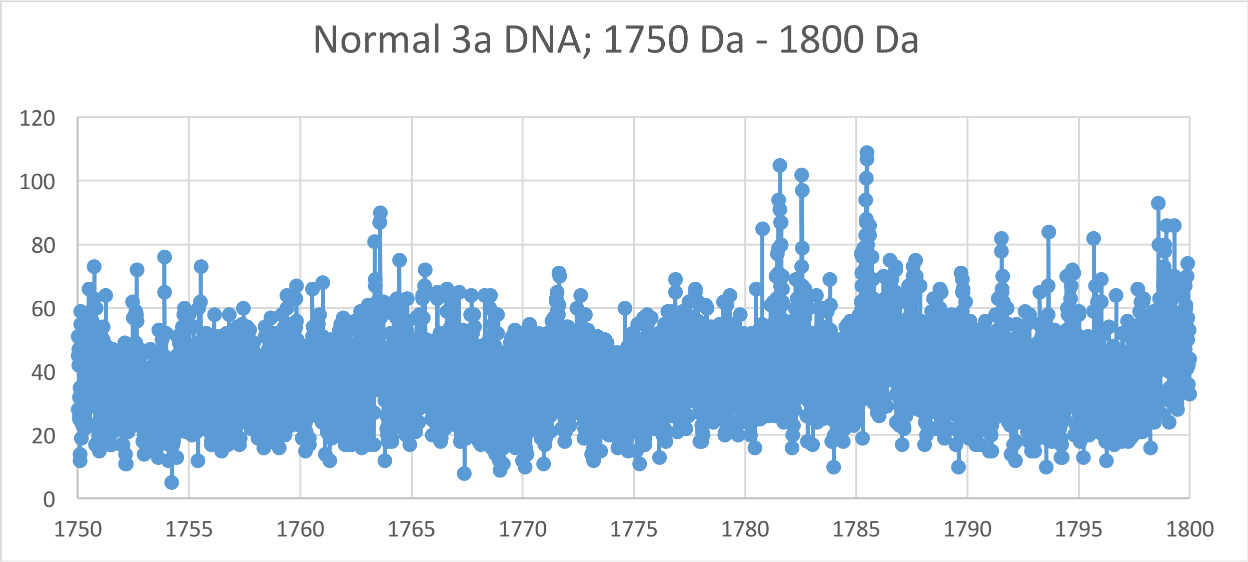
Normal 3a DNA; 1750 Da - 1800 Da
120
100
80
60
40
20
0
1750
1755
1760
1765
1770
1775
1780
1785
1790
1795
1800

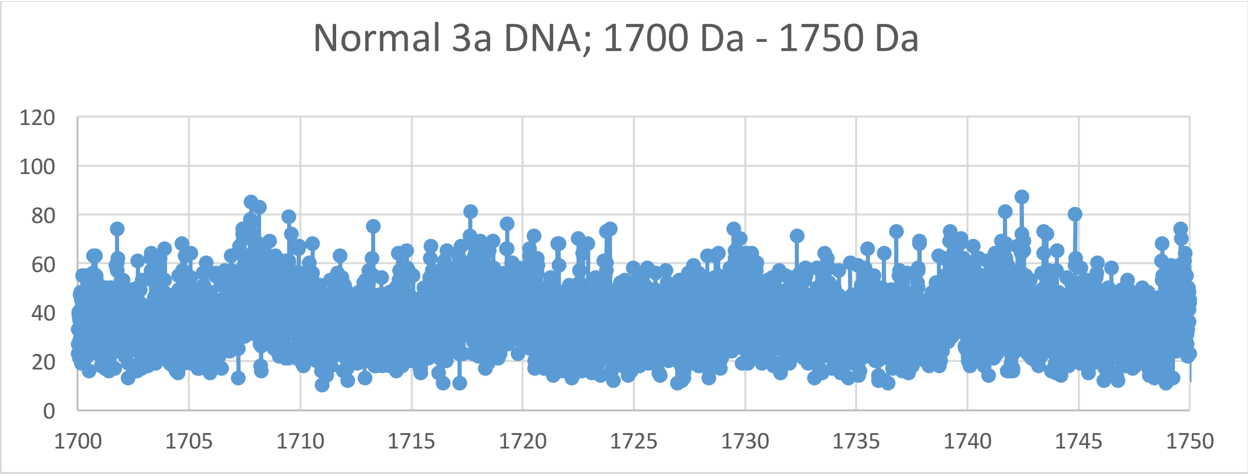
Normal 3a DNA; 1700 Da - 1750 Da
120
100
80
60
40
20
0
1700
1705
1710
1715
1720
1725
1730
1735
1740
1745
1750

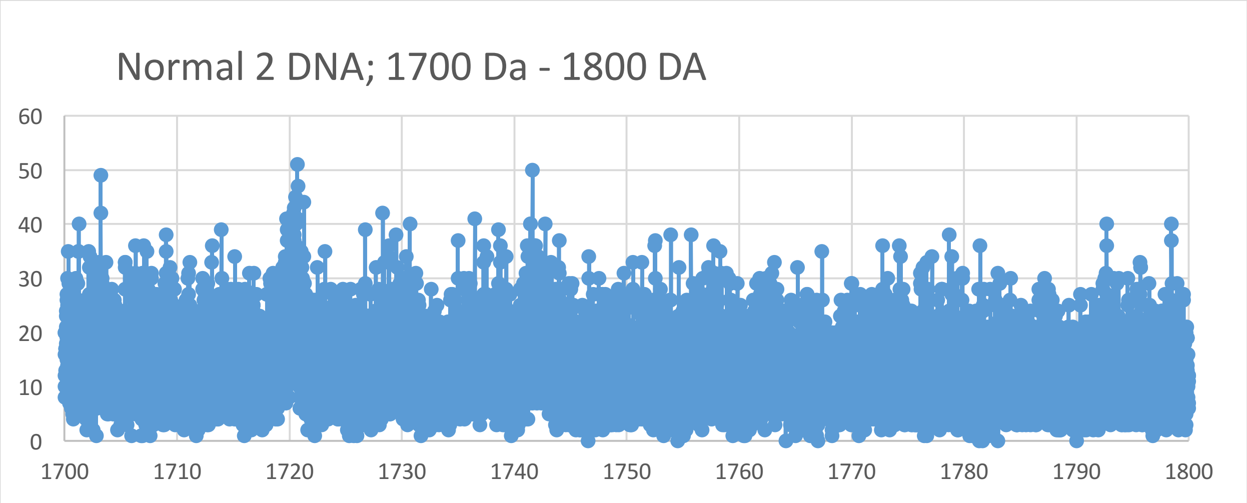
Normal 2 DNA; 1700 Da - 1800 DA
0
10
20
30
40
50
60
1700
1710
1720
1730
1740
1750
1760
1770
1780
1790
1800

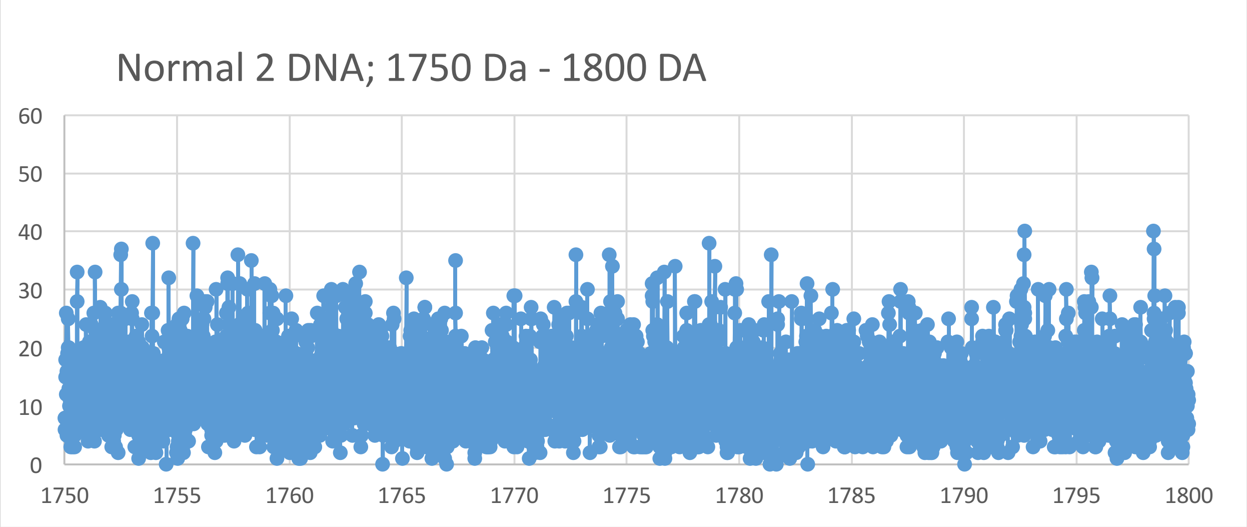
Normal 2 DNA; 1750 Da - 1800 DA
0
10
20
30
40
50
60
1750
1755
1760
1765
1770
1775
1780
1785
1790
1795
1800

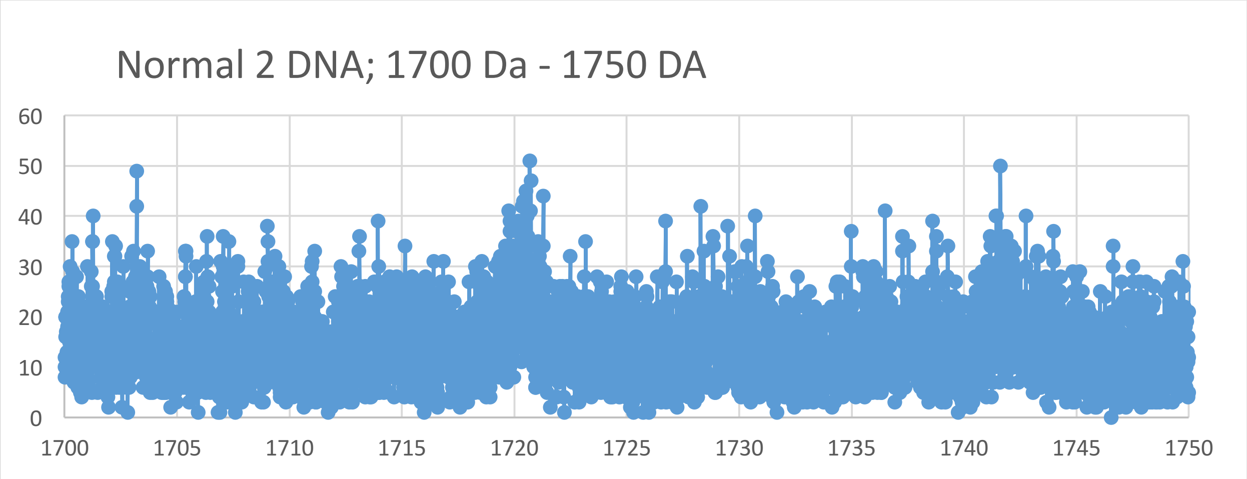
Normal 2 DNA; 1700 Da - 1750 DA
0
10
20
30
40
50
60
1700
1705
1710
1715
1720
1725
1730
1735
1740
1745
1750

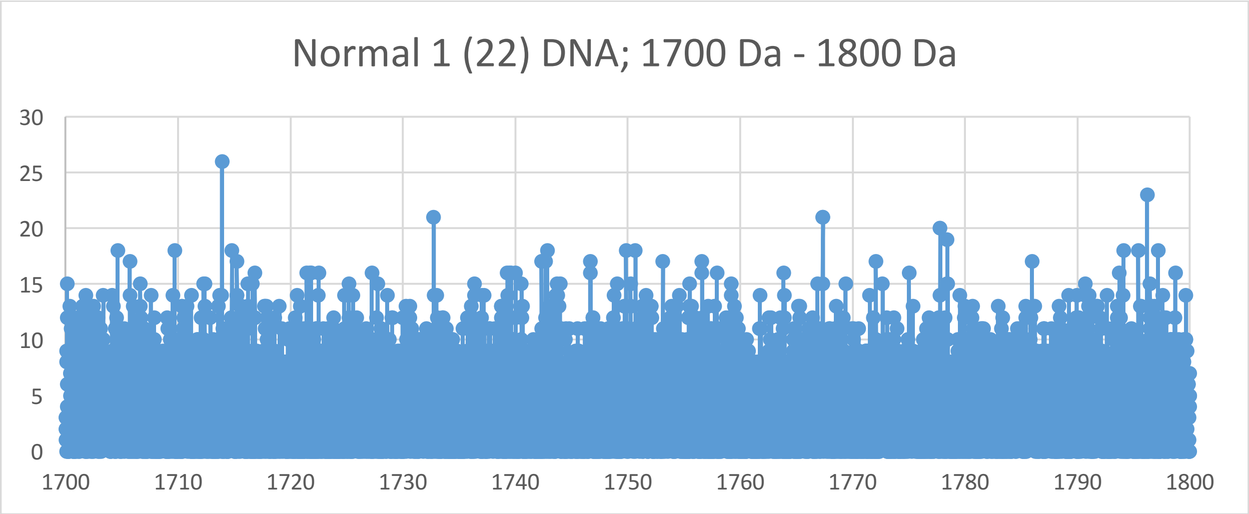
Normal 1 (22) DNA; 1700 Da - 1800 Da
30
25
20
15
10
5
0
1700
1710
1720
1730
1740
1750
1760
1770
1780
1790
1800

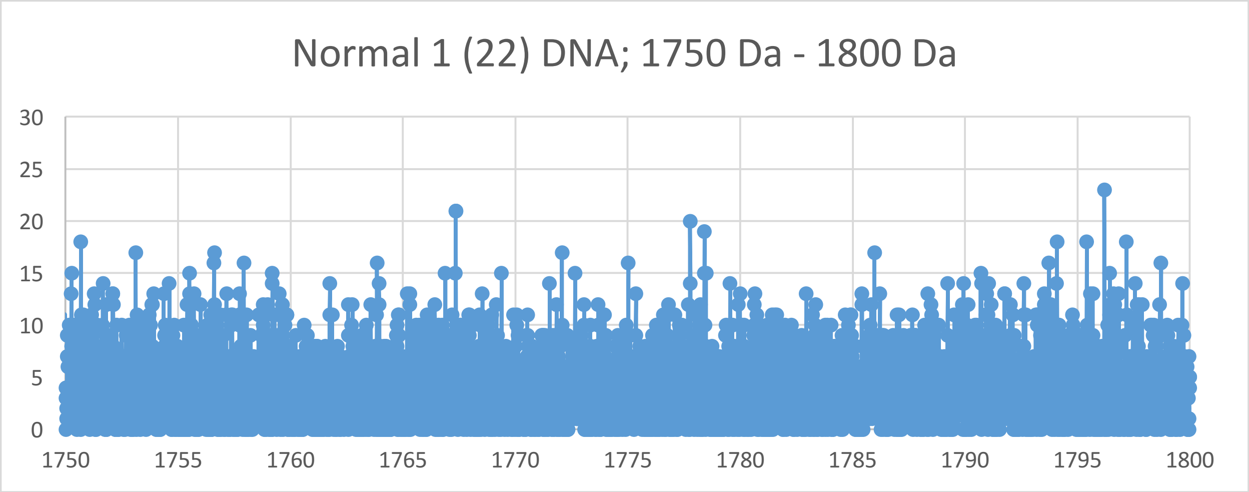
Normal 1 (22) DNA; 1750 Da - 1800 Da
30
25
20
15
10
5
0
1750
1755
1760
1765
1770
1775
1780
1785
1790
1795
1800

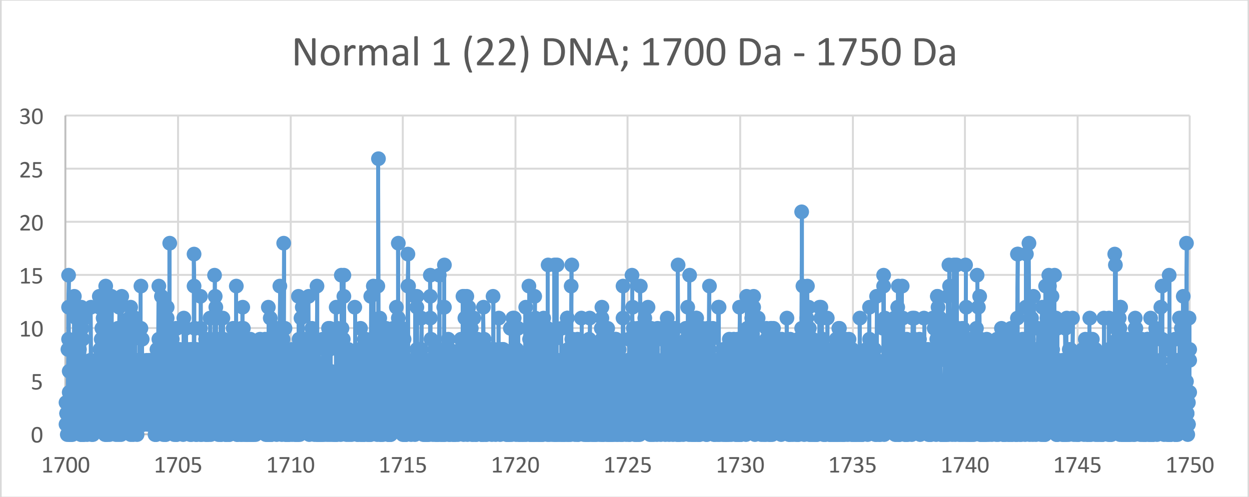
Normal 1 (22) DNA; 1700 Da - 1750 Da
30
25
20
15
10
5
0
1700
1705
1710
1715
1720
1725
1730
1735
1740
1745
1750
