## Supplementary Materials IV for "Oncogenesis and Aging by Isotopic Functionalizations of the Proteins and Nucleic Acids"

IV Supplementary Materials – New Mass Spectra from 1600 Da -1700 Da

Results for Supplementary 4

So for the tetranucleotides from $(phos)_8$ to $(phos)_7$, the telomeric and nontelomeic peaks for normal increase in number and the peaks for cancer decrease in number for more stable normal tetranucleotides and less stable cancerous tetranucleotides with this dephosphorylation. The masses range from 1324 Da to 1470 Da. See Supplementary Figures IV for graphs. These results and observations point specific differences between cancerous and normal oligonucleotides chemically and isotopically. The mating telomeric peaks for normal increase with this dephosphorylations of the tetranucleotides. This dephosphorylations from $(phos)_8$ to $(phos)_7$ caused the resulting more intense and stable normal tetranucleotides to be more nontelomeric, mating telomeric and telomeric and less isotopically enriched. These dephosphorylations from $(phos)_8$ to $(phos)_7$ also caused the nontelomeric tetranucleotides to be more from intense in normal DNA and less isotopically enriched. It seems these dephosphorylations are related to the isotopic enrichments and depletions; but the telomeres are more intense in normal and less isotopically enriched whereas the intensities of the mating telomeres are more intense in normal altered and the isotopic enrichments are less. The telomeric tetranucleotides of normal tended to also be less enriched so it seems the normal stabilities of the telomeric tetranucleotides are related to there less isotopic enrichments for these dephosphorylations. Enrichments weaken the telomeric and nontelomeric normal tetranucleotides while weakening cancerous mating telomeric tetranucleotides. The more intense normal telomeric tetranucleotides were also less isotopically enriched. These dephosphorylations from $(phos)_8$ to $(phos)_7$ decreased the isotopic enrichments of the cancer relative to the normal for the nontelomeric tetranucleotides. But the mating cancerous telomeric tetranucleotides were less enriched from these dephosphorylations for less enrichments; but the normal mating telomeric tetranucleotides are stronger and more isotopically enriched. Such seems compatible for replications and transcriptions as stable telomeres seem compatible to unstable mating telomeres for the unravelings and constructions of DNA mating telomeres onto the building templates of the telomeres. These dephosphorylations from $(phos)_8$ to $(phos)_7$ caused methylations to cause more cancer intensities and stabilities but more isotopic enrichments for telomeres, mating telomeres and nontelomeres and (slightly for nontelomeres) with large effects on mating telomeric tetranucleotides as the intensities of normal increased. But the methylations of the mating telomeres increased cancer stabilities and less isotopic enrichments for contrary effects on mating telomeres compared to telomeres. These dephosphorylations from $(phos)_8$ to $(phos)_7$ caused acetylations to cause more cancer stabilities and less normal stabilities for mating telomeres and nontelomeres with more enrichments relative to methylations; but the telomeres are more cancer stable and negligibly isotopically enriched by the acetylations. Thereby these dephosphorylations $(phos)_8$ to $(phos)_7$ and there varying consequences with respect to other dephosphorylations determine patterns of dephosphorylations and isotopic enrichment patterns in normal DNA which mutate and change for different patterns in DNA of cells that become cancerous. These dephosphorylations $(phos)_8$ to $(phos)_7$ induced isotopic dynamics from varying methylations and acetylations in cancer DNA relative to normal DNA couple to other isotopically induced varied phosphorylations, methylations and acetylations throughout the DNA for emerging order from nanodisorder.

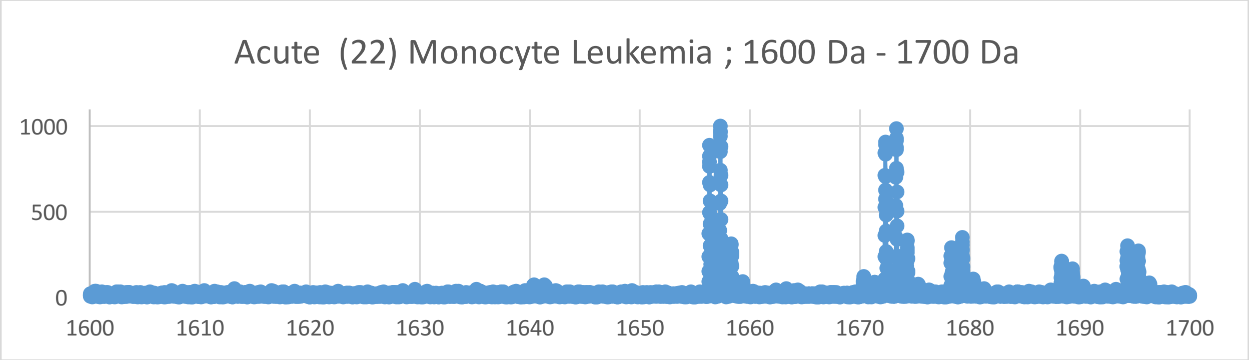
Acute (22) Monocyte Leukemia ; 1600 Da - 1700 Da
1000
500
0
1600
1610
1620
1630
1640
1650
1660
1670
1680
1690
1700

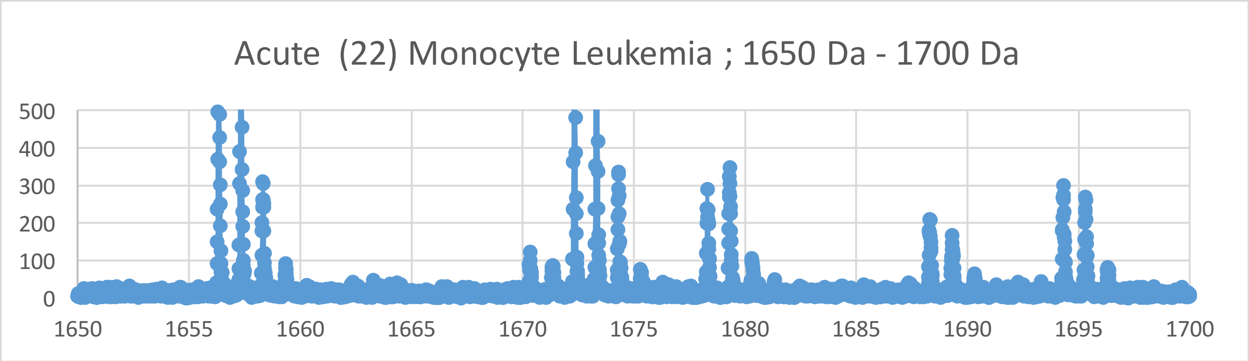
Acute (22) Monocyte Leukemia ; 1650 Da - 1700 Da
500
400
300
200
100
0
1650
1655
1660
1665
1670
1675
1680
1685
1690
1695
1700

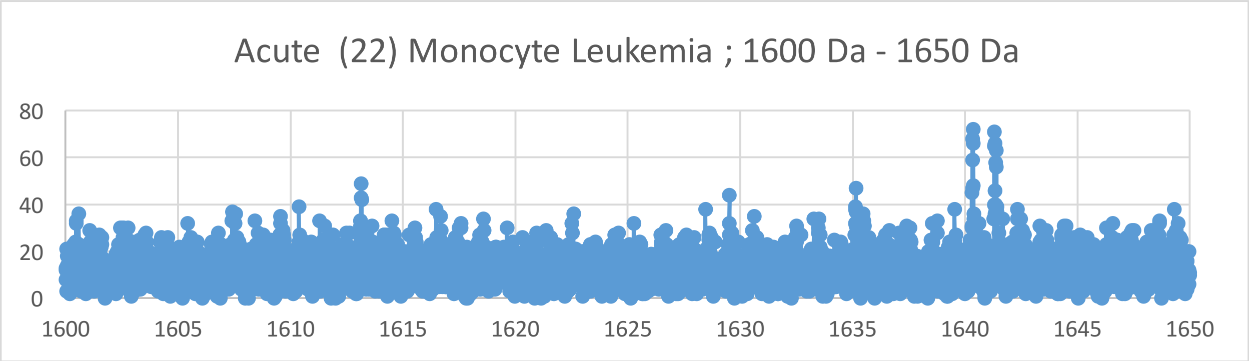
Acute (22) Monocyte Leukemia ; 1600 Da - 1650 Da
80
60
40
20
0
1600
1605
1610
1615
1620
1625
1630
1635
1640
1645
1650

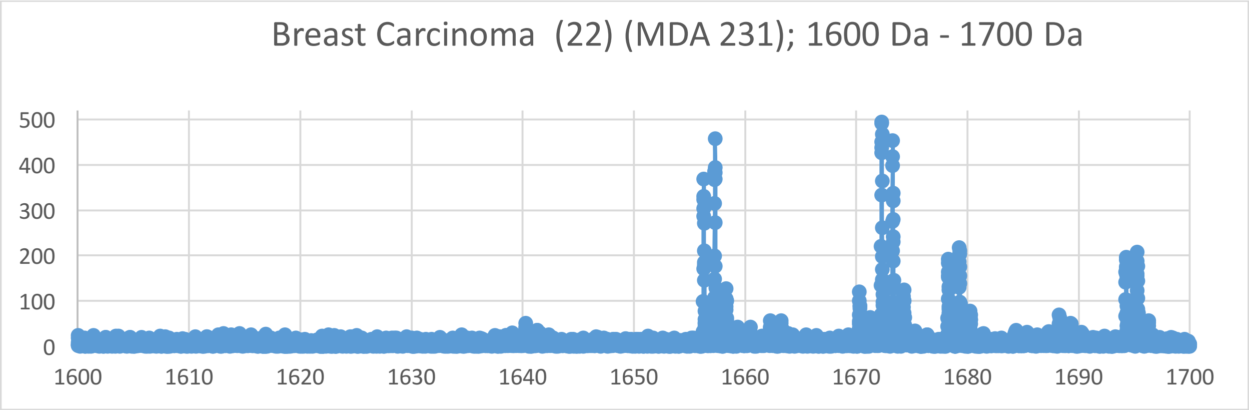
Breast Carcinoma (22) (MDA 231); 1600 Da - 1700 Da
500
400
300
200
100
0
1600 1610 1620 1630 1640 1650 1660 1670 1680 1690 1700

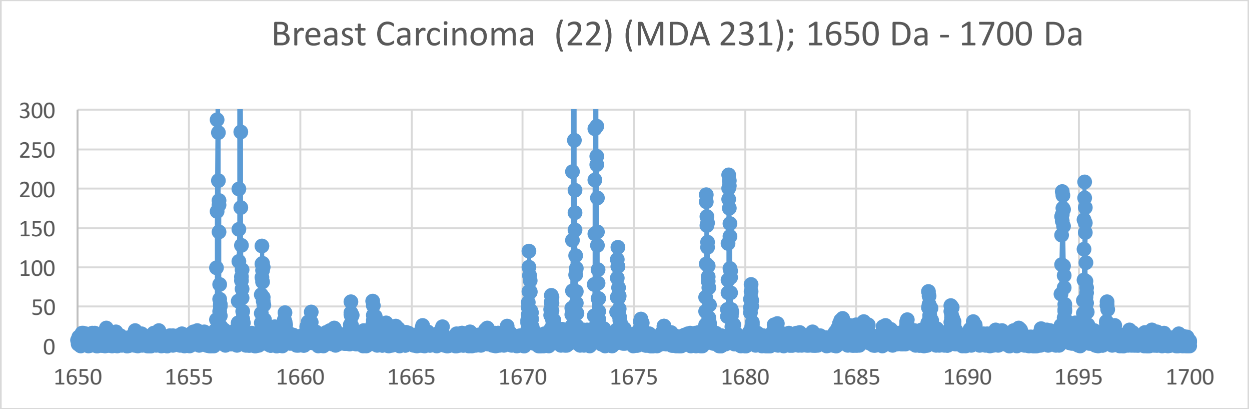
Breast Carcinoma (22) (MDA 231); 1650 Da - 1700 Da
300
250
200
150
100
50
0
1650 1655 1660 1665 1670 1675 1680 1685 1690 1695 1700

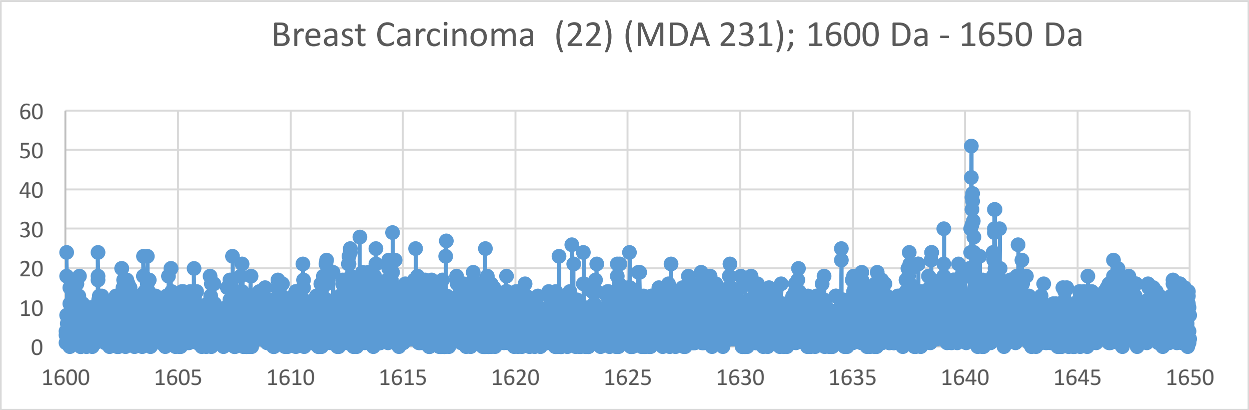
Breast Carcinoma (22) (MDA 231); 1600 Da - 1650 Da
60
50
40
30
20
10
0
1600 1605 1610 1615 1620 1625 1630 1635 1640 1645 1650

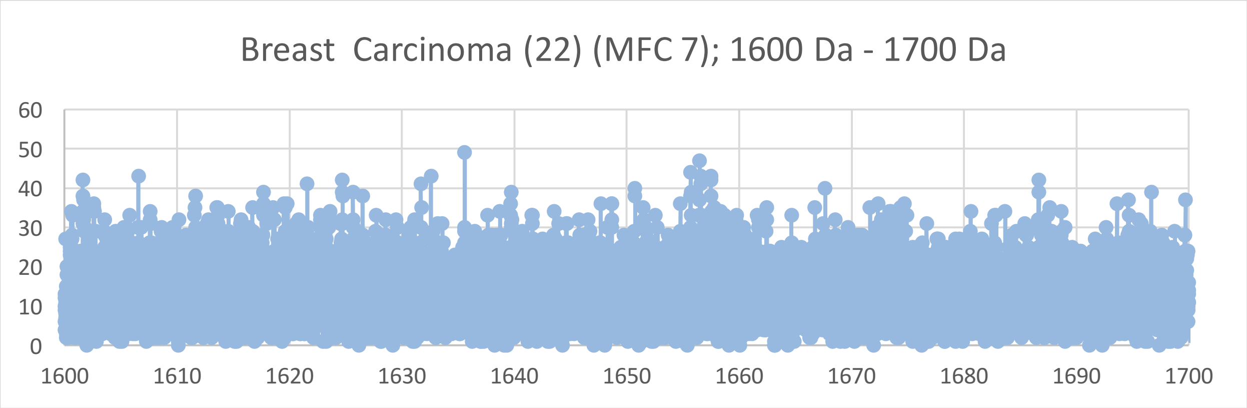
Breast Carcinoma (22) (MFC 7); 1600 Da - 1700 Da
60
50
40
30
20
10
0
1600
1610
1620
1630
1640
1650
1660
1670
1680
1690
1700

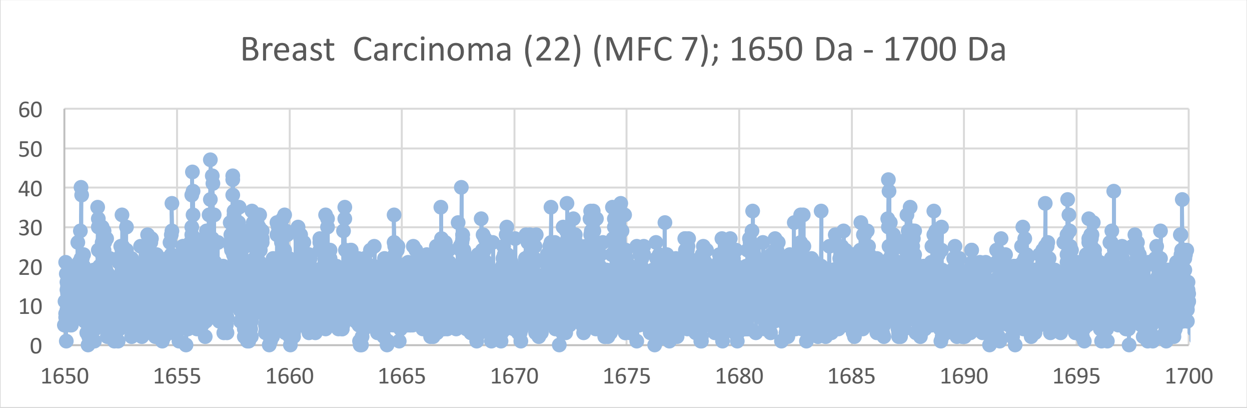
Breast Carcinoma (22) (MFC 7); 1650 Da - 1700 Da
60
50
40
30
20
10
0
1650
1655
1660
1665
1670
1675
1680
1685
1690
1695
1700

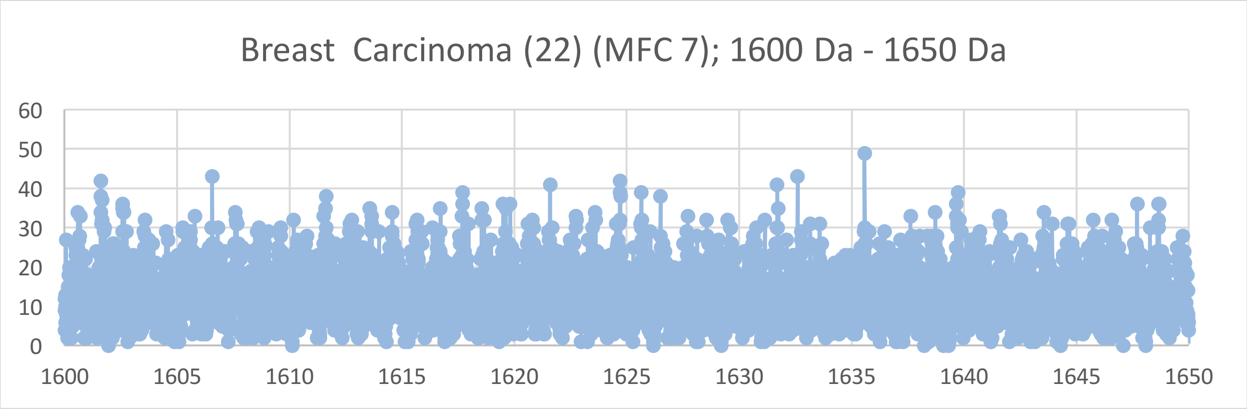
Breast Carcinoma (22) (MFC 7); 1600 Da - 1650 Da
60
50
40
30
20
10
0
1600
1605
1610
1615
1620
1625
1630
1635
1640
1645
1650

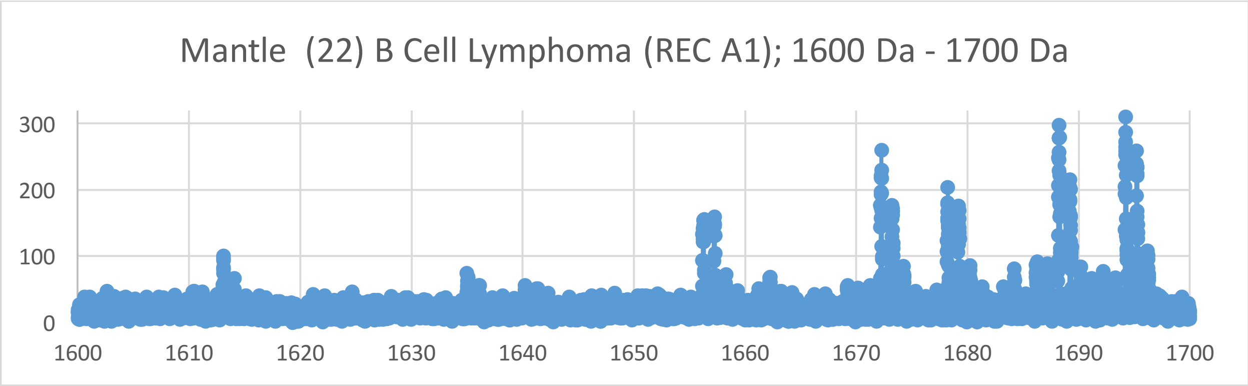
Mantle (22) B Cell Lymphoma (REC A1); 1600 Da - 1700 Da
300
200
100
0
1600 1610 1620 1630 1640 1650 1660 1670 1680 1690 1700

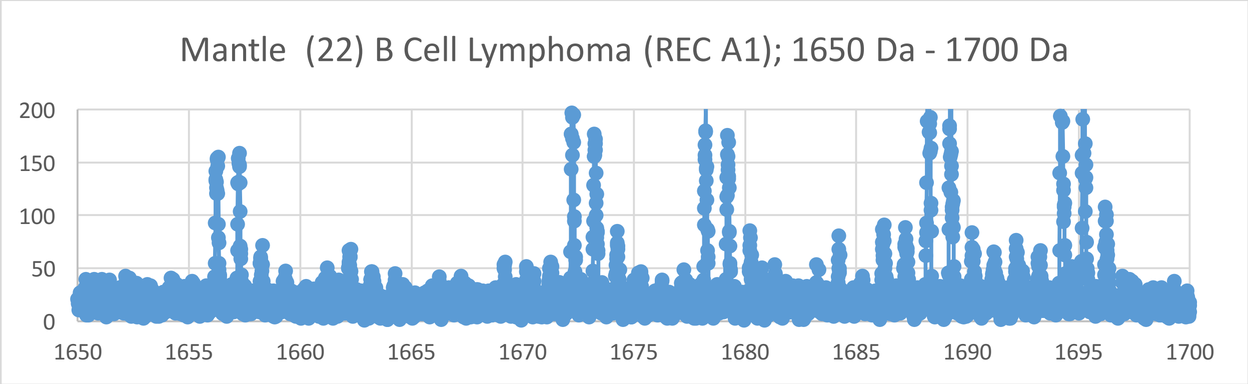
Mantle (22) B Cell Lymphoma (REC A1); 1650 Da - 1700 Da
200
150
100
50
0
1650 1655 1660 1665 1670 1675 1680 1685 1690 1695 1700

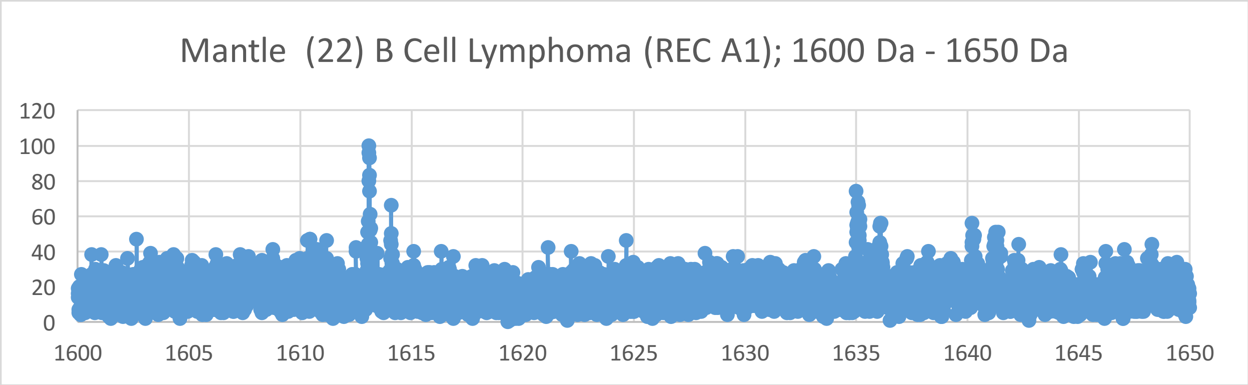
Mantle (22) B Cell Lymphoma (REC A1); 1600 Da - 1650 Da
120
100
80
60
40
20
0
1600 1605 1610 1615 1620 1625 1630 1635 1640 1645 1650

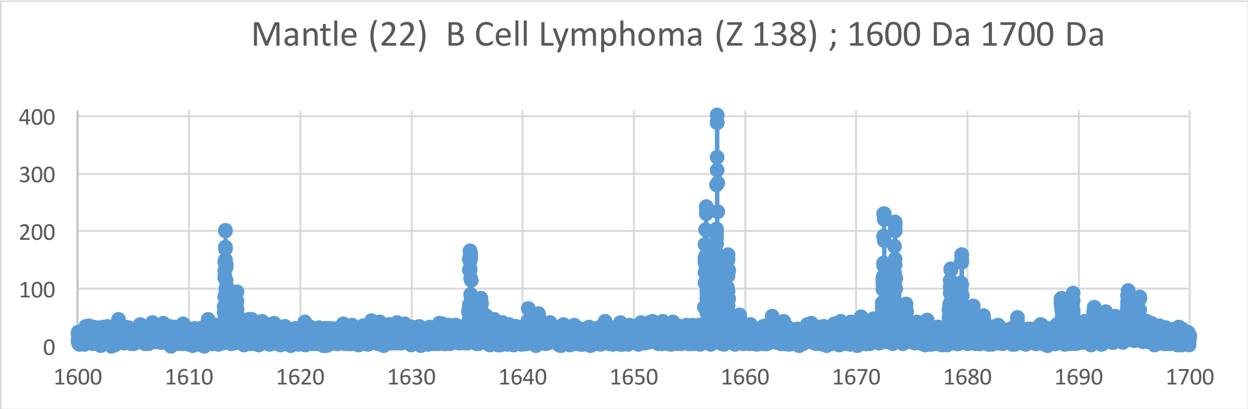
Mantle (22) B Cell Lymphoma (Z 138) ; 1600 Da 1700 Da
400
300
200
100
0
1600 1610 1620 1630 1640 1650 1660 1670 1680 1690 1700

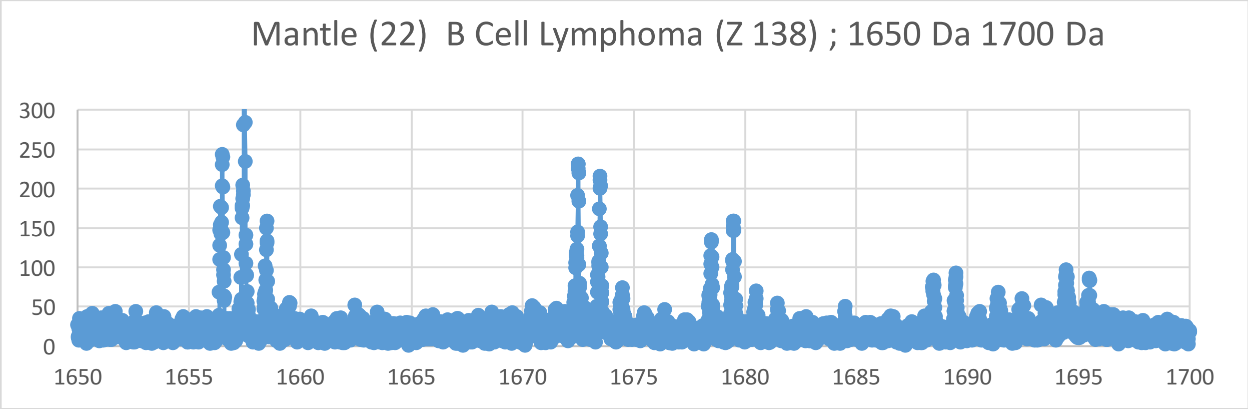
Mantle (22) B Cell Lymphoma (Z 138) ; 1650 Da 1700 Da
300
250
200
150
100
50
0
1650 1655 1660 1665 1670 1675 1680 1685 1690 1695 1700

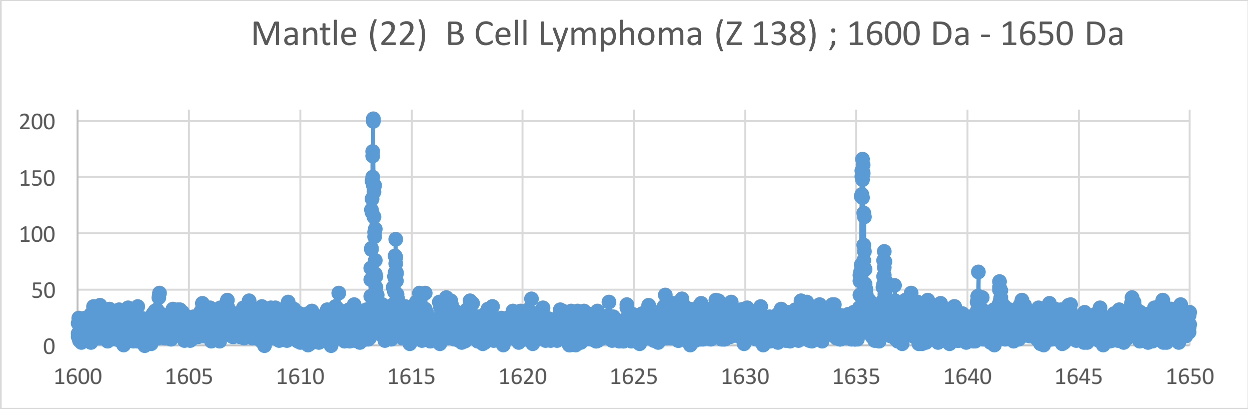
Mantle (22) B Cell Lymphoma (Z 138) ; 1600 Da - 1650 Da
200
150
100
50
0
1600 1605 1610 1615 1620 1625 1630 1635 1640 1645 1650

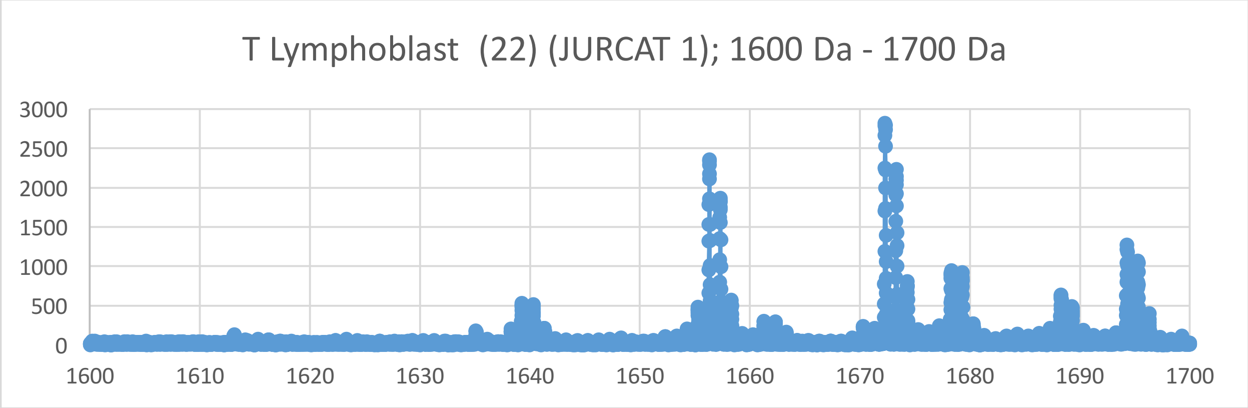
T Lymphoblast (22) (JURCAT 1); 1600 Da - 1700 Da
3000
2500
2000
1500
1000
500
0
1600
1610
1620
1630
1640
1650
1660
1670
1680
1690
1700

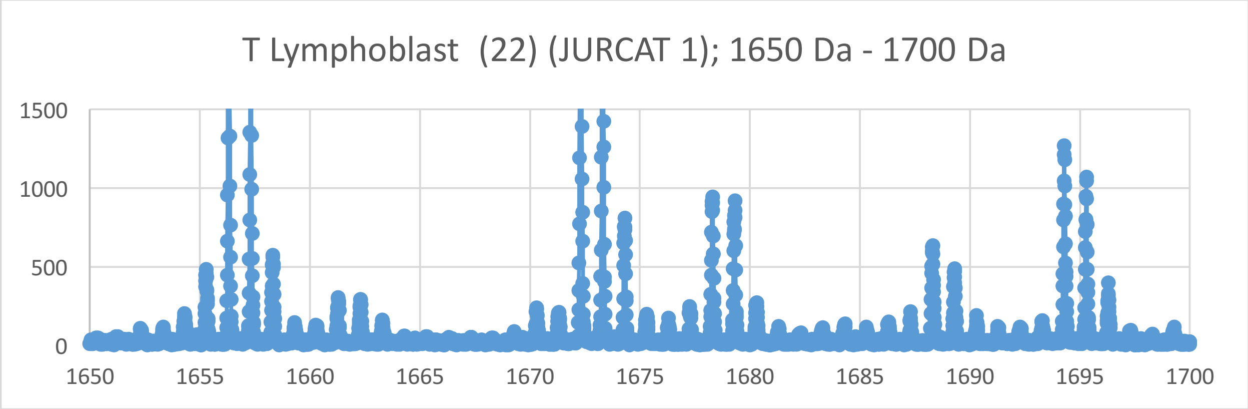
T Lymphoblast (22) (JURCAT 1); 1650 Da - 1700 Da
1500
1000
500
0
1650
1655
1660
1665
1670
1675
1680
1685
1690
1695
1700

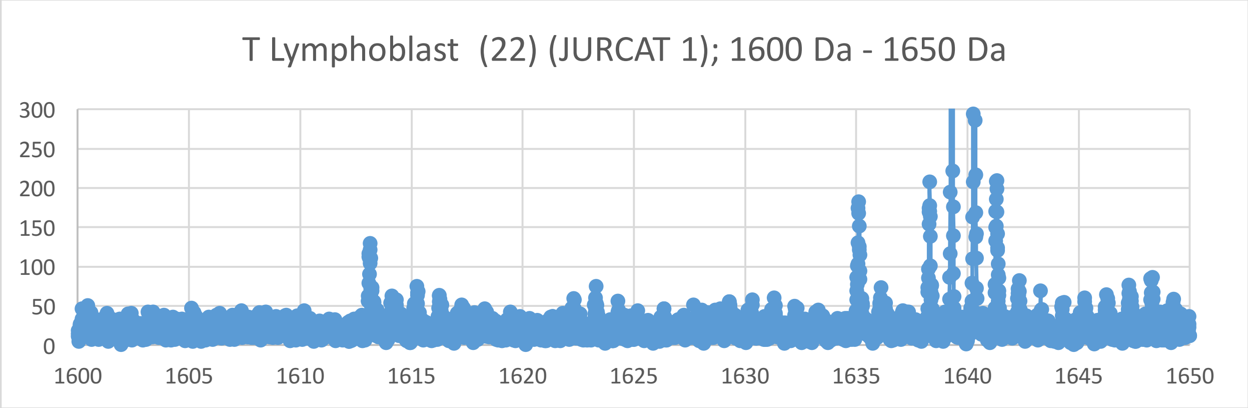
T Lymphoblast (22) (JURCAT 1); 1600 Da - 1650 Da
300
250
200
150
100
50
0
1600
1605
1610
1615
1620
1625
1630
1635
1640
1645
1650

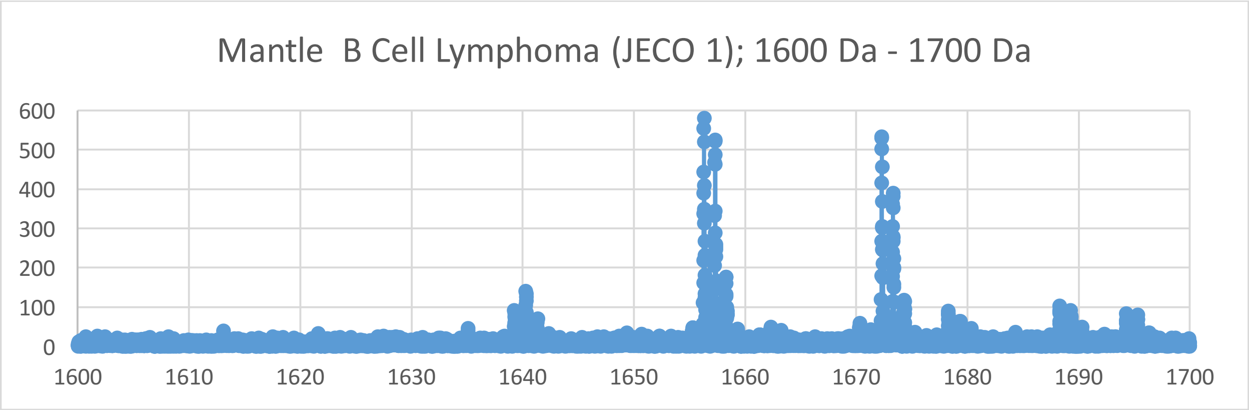
Mantle B Cell Lymphoma (JECO 1); 1600 Da - 1700 Da
600
500
400
300
200
100
0
1600
1610
1620
1630
1640
1650
1660
1670
1680
1690
1700

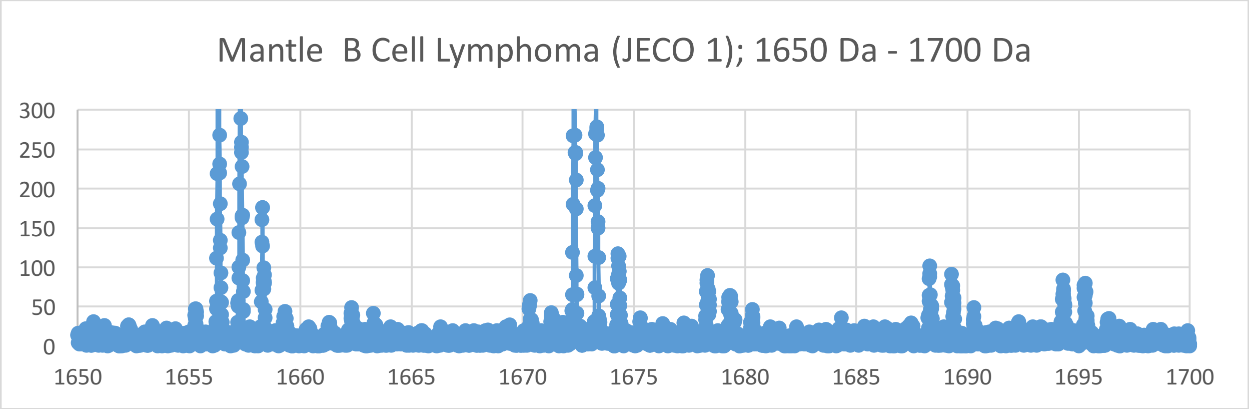
Mantle B Cell Lymphoma (JECO 1); 1650 Da - 1700 Da
300
250
200
150
100
50
0
1650
1655
1660
1665
1670
1675
1680
1685
1690
1695
1700

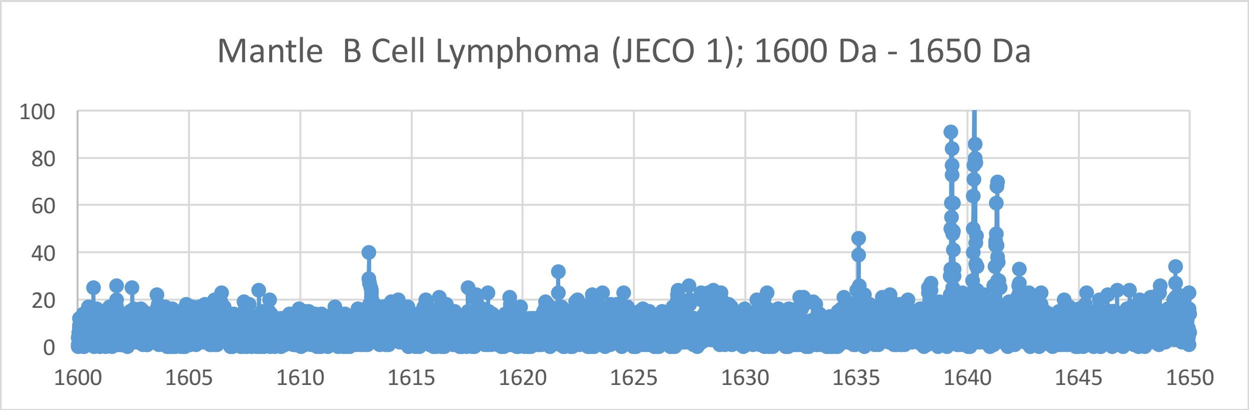
Mantle B Cell Lymphoma (JECO 1); 1600 Da - 1650 Da
100
80
60
40
20
0
1600
1605
1610
1615
1620
1625
1630
1635
1640
1645
1650

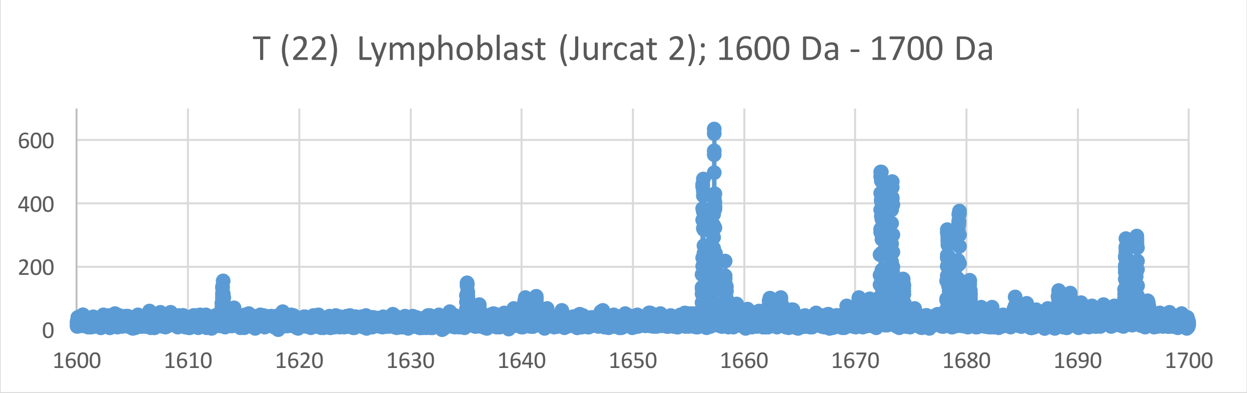
T (22) Lymphoblast (Jurcat 2); 1600 Da - 1700 Da
0
200
400
600
1600
1610
1620
1630
1640
1650
1660
1670
1680
1690
1700

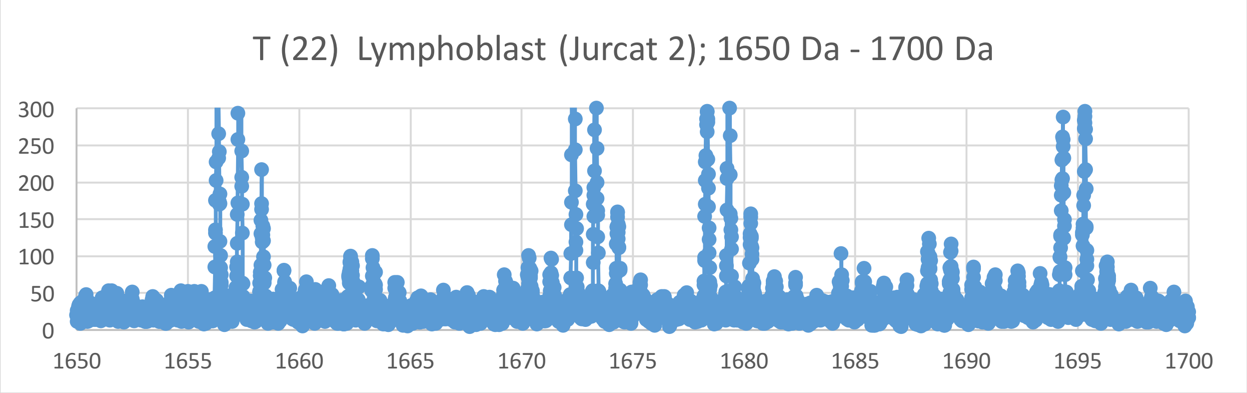
T (22) Lymphoblast (Jurcat 2); 1650 Da - 1700 Da
0
50
100
150
200
250
300
1650
1655
1660
1665
1670
1675
1680
1685
1690
1695
1700

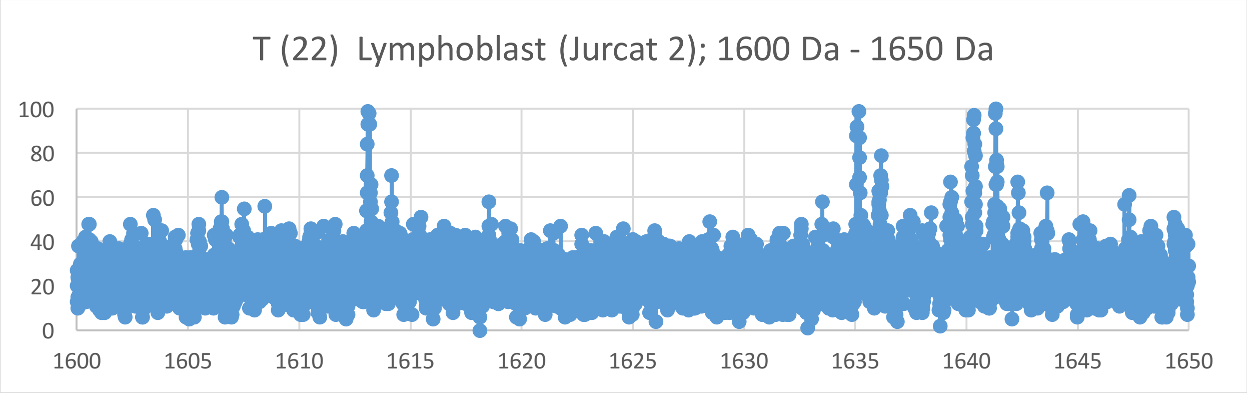
T (22) Lymphoblast (Jurcat 2); 1600 Da - 1650 Da
0
20
40
60
80
100
1600
1605
1610
1615
1620
1625
1630
1635
1640
1645
1650

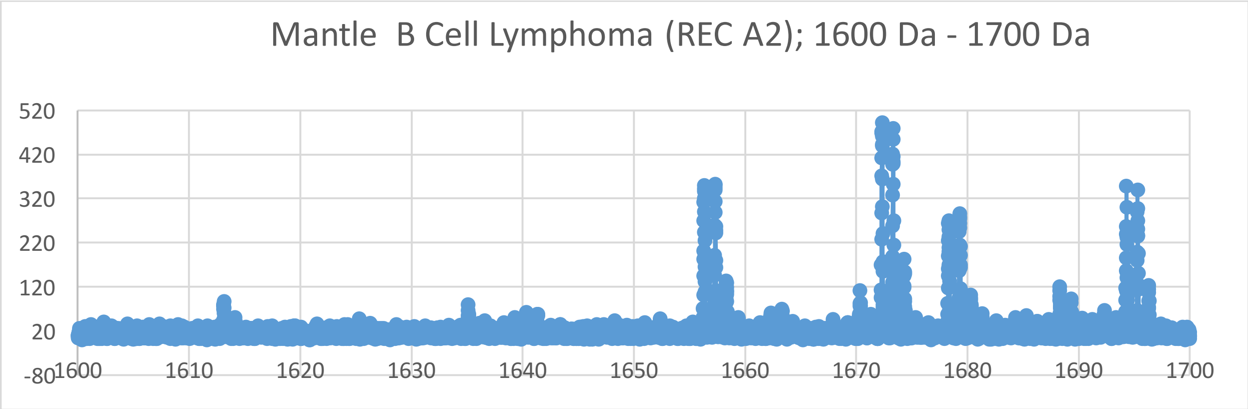
Mantle B Cell Lymphoma (REC A2); 1600 Da - 1700 Da
520
420
320
220
120
20
-80
1600
1610
1620
1630
1640
1650
1660
1670
1680
1690
1700

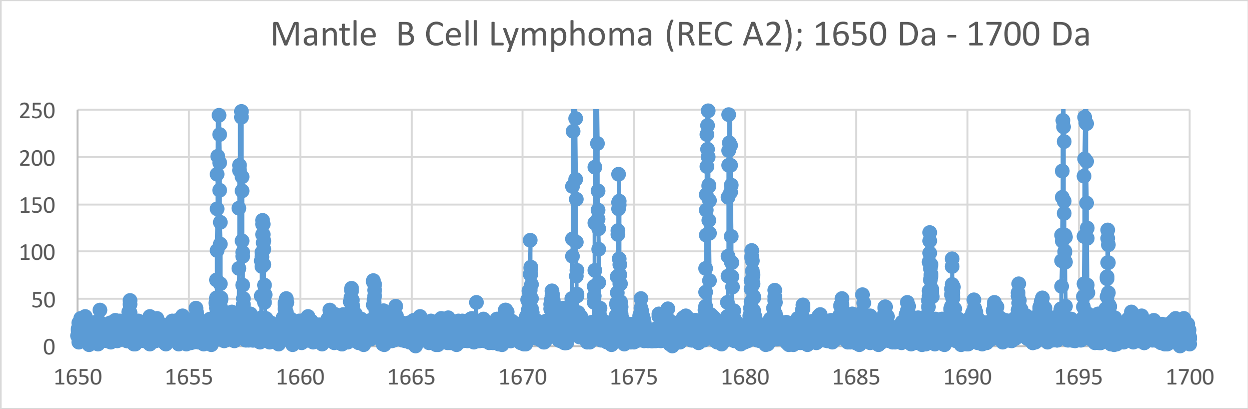
Mantle B Cell Lymphoma (REC A2); 1650 Da - 1700 Da
250
200
150
100
50
0
1650
1655
1660
1665
1670
1675
1680
1685
1690
1695
1700

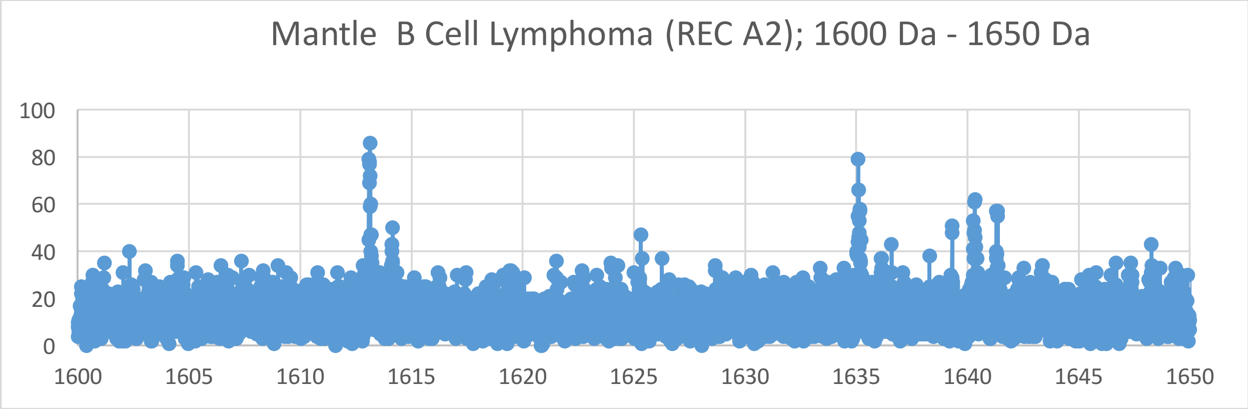
Mantle B Cell Lymphoma (REC A2); 1600 Da - 1650 Da
100
80
60
40
20
0
1600
1605
1610
1615
1620
1625
1630
1635
1640
1645
1650

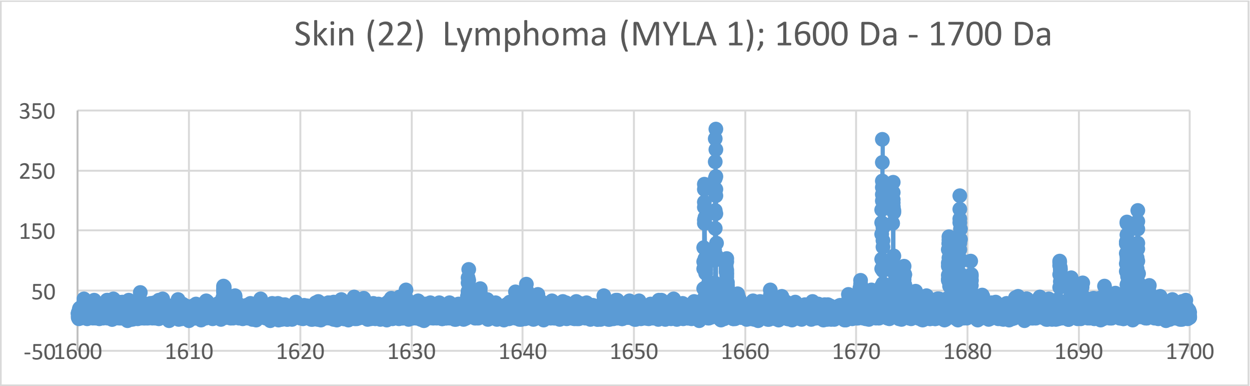
Skin (22)  Lymphoma (MYLA 1); 1600 Da - 1700 Da
350
250
150
50
-50
1600
1610
1620
1630
1640
1650
1660
1670
1680
1690
1700

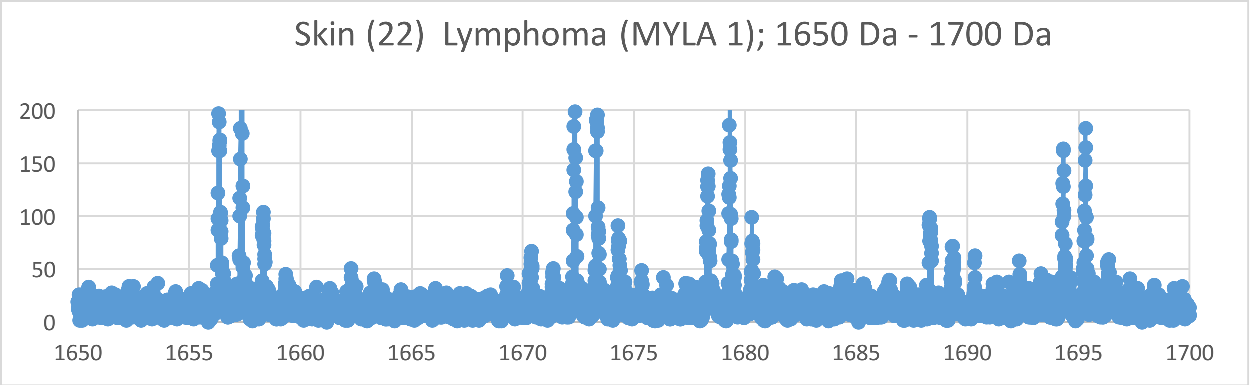
Skin (22)  Lymphoma (MYLA 1); 1650 Da - 1700 Da
200
150
100
50
0
1650
1655
1660
1665
1670
1675
1680
1685
1690
1695
1700

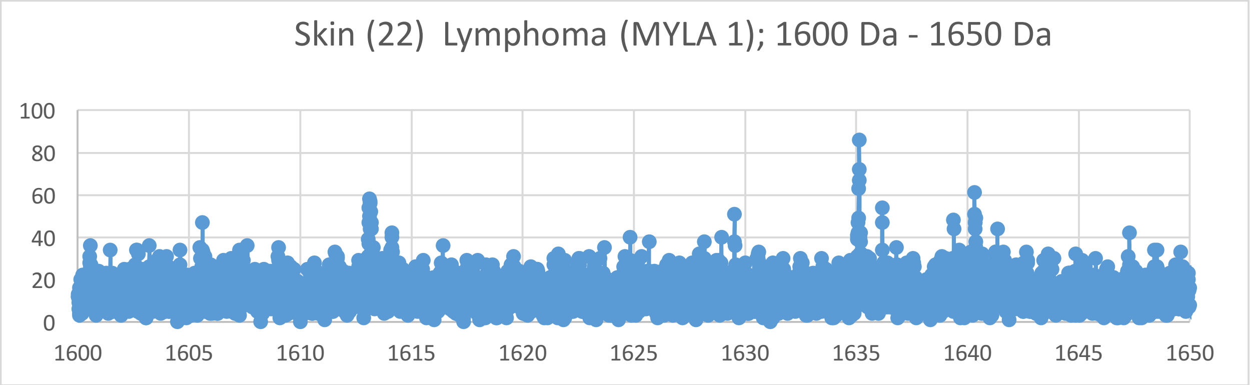
Skin (22)  Lymphoma (MYLA 1); 1600 Da - 1650 Da
100
80
60
40
20
0
1600
1605
1610
1615
1620
1625
1630
1635
1640
1645
1650

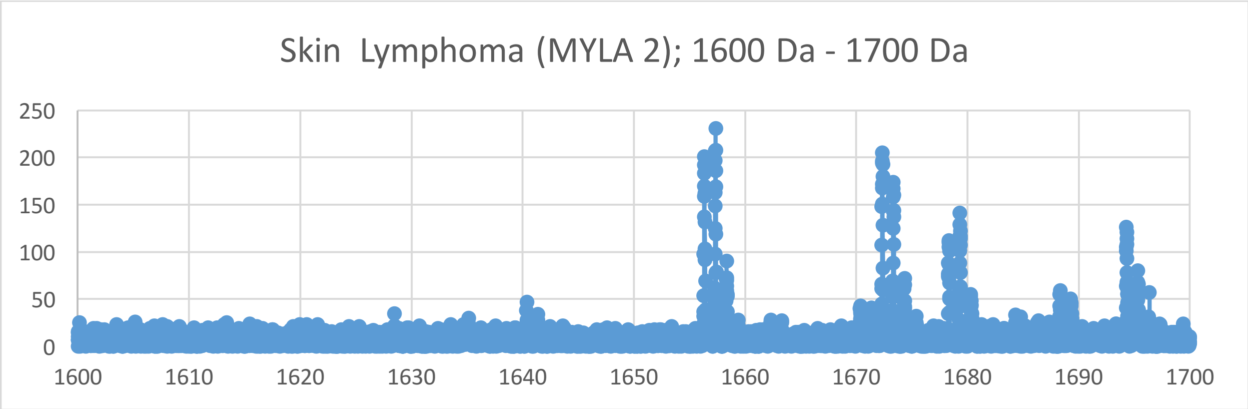
Skin Lymphoma (MYLA 2); 1600 Da - 1700 Da
250
200
150
100
50
0
1600
1610
1620
1630
1640
1650
1660
1670
1680
1690
1700

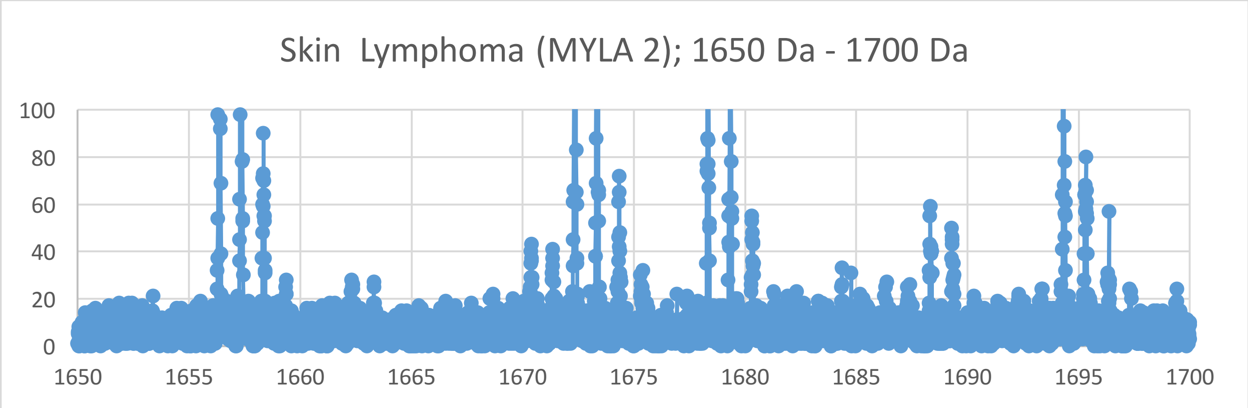
Skin Lymphoma (MYLA 2); 1650 Da - 1700 Da
100
80
60
40
20
0
1650
1655
1660
1665
1670
1675
1680
1685
1690
1695
1700

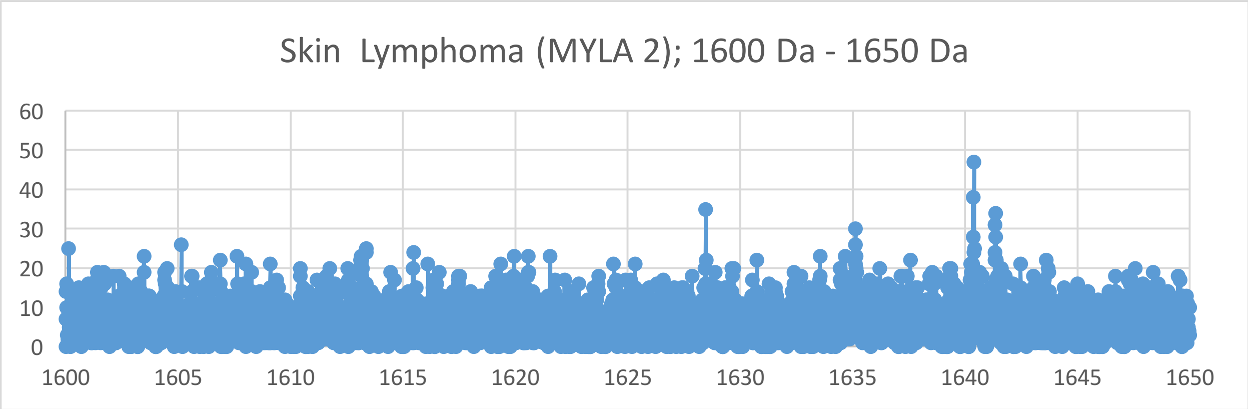
Skin Lymphoma (MYLA 2); 1600 Da - 1650 Da
60
50
40
30
20
10
0
1600
1605
1610
1615
1620
1625
1630
1635
1640
1645
1650

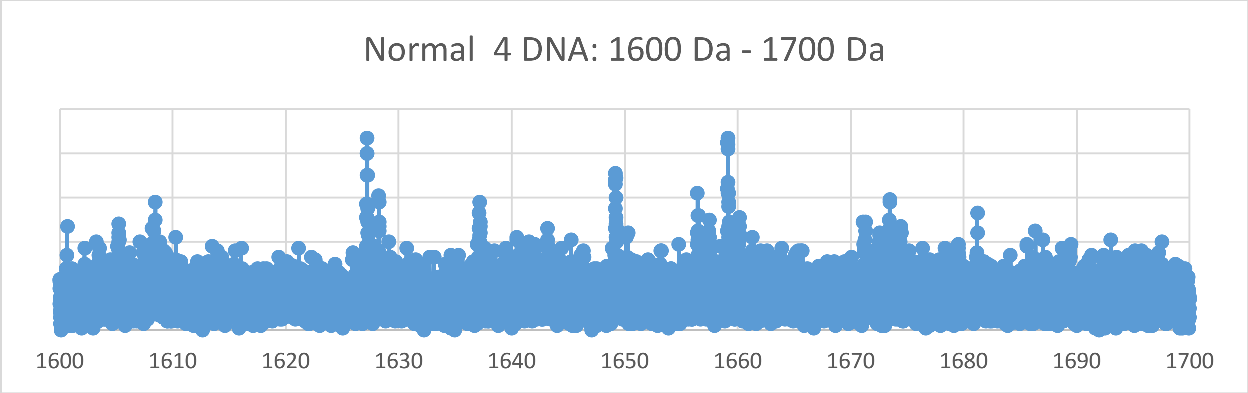
Normal 4 DNA: 1600 Da - 1700 Da
1600
1610
1620
1630
1640
1650
1660
1670
1680
1690
1700

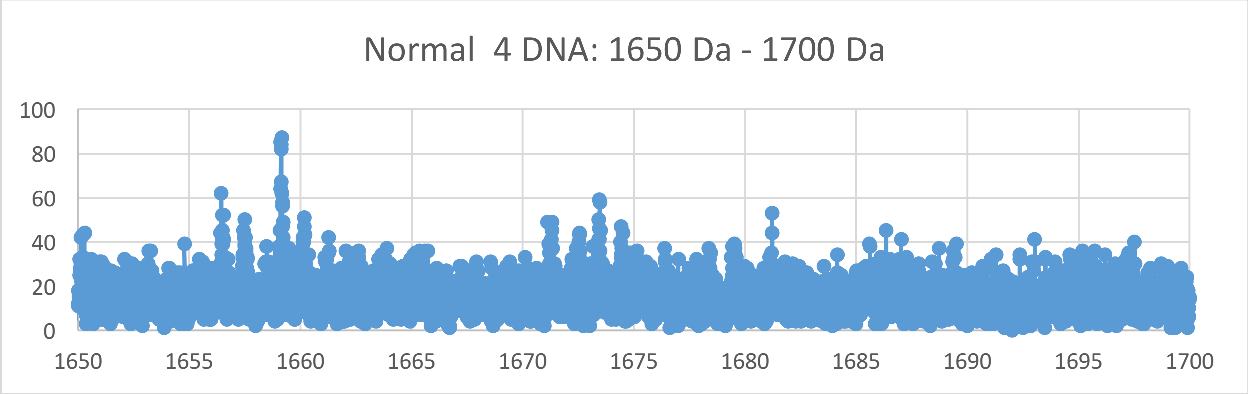
Normal 4 DNA: 1650 Da - 1700 Da
100
80
60
40
20
0
1650
1655
1660
1665
1670
1675
1680
1685
1690
1695
1700

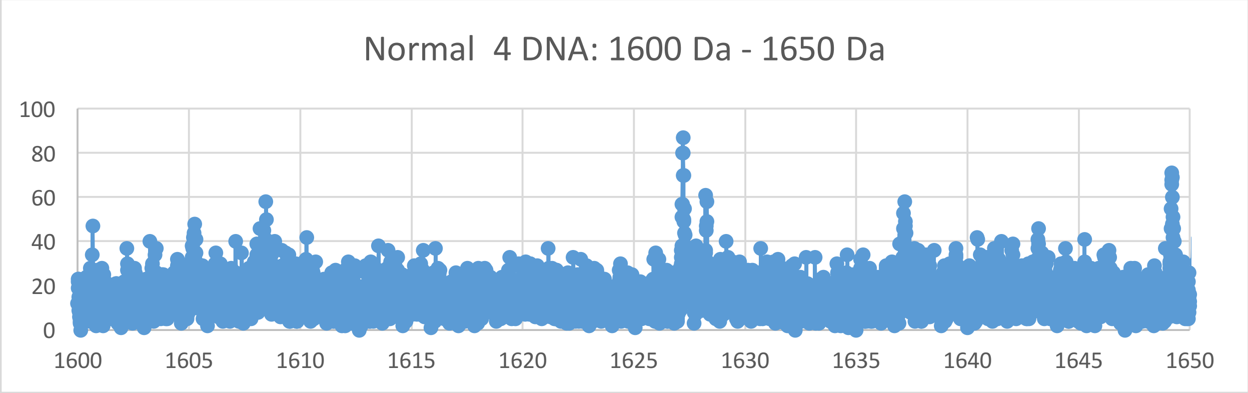
Normal 4 DNA: 1600 Da - 1650 Da
100
80
60
40
20
0
1600
1605
1610
1615
1620
1625
1630
1635
1640
1645
1650

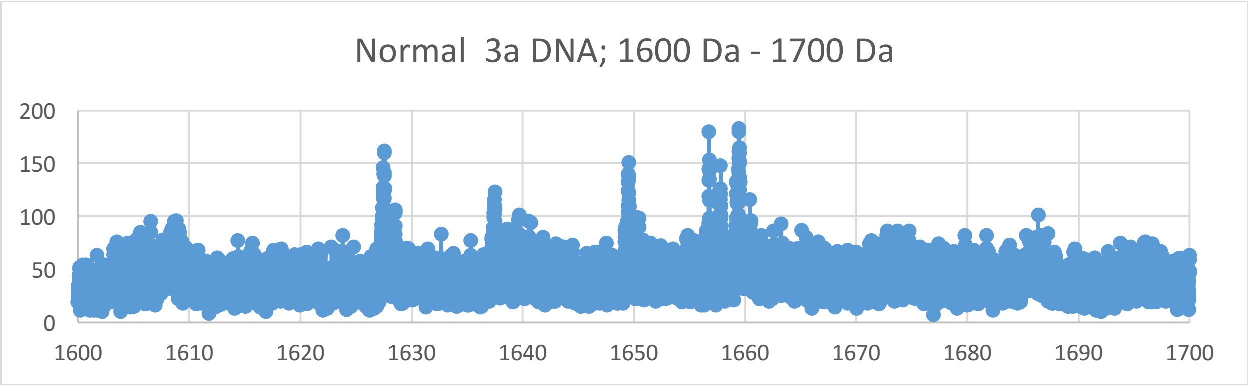

Normal 3a DNA; 1600 Da - 1700 Da
200
150
100
50
0
1600
1610
1620
1630
1640
1650
1660
1670
1680
1690
1700


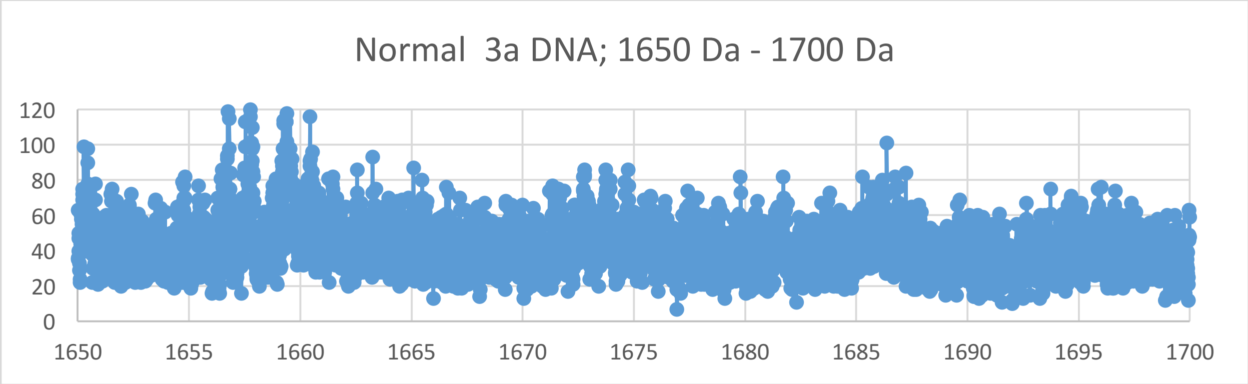

Normal 3a DNA; 1650 Da - 1700 Da
120
100
80
60
40
20
0
1650
1655
1660
1665
1670
1675
1680
1685
1690
1695
1700


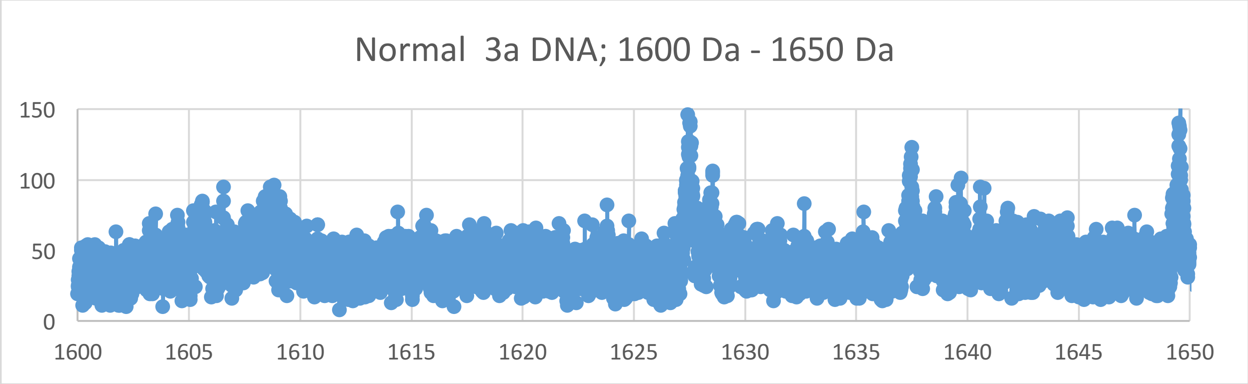

Normal 3a DNA; 1600 Da - 1650 Da
150
100
50
0
1600
1605
1610
1615
1620
1625
1630
1635
1640
1645
1650

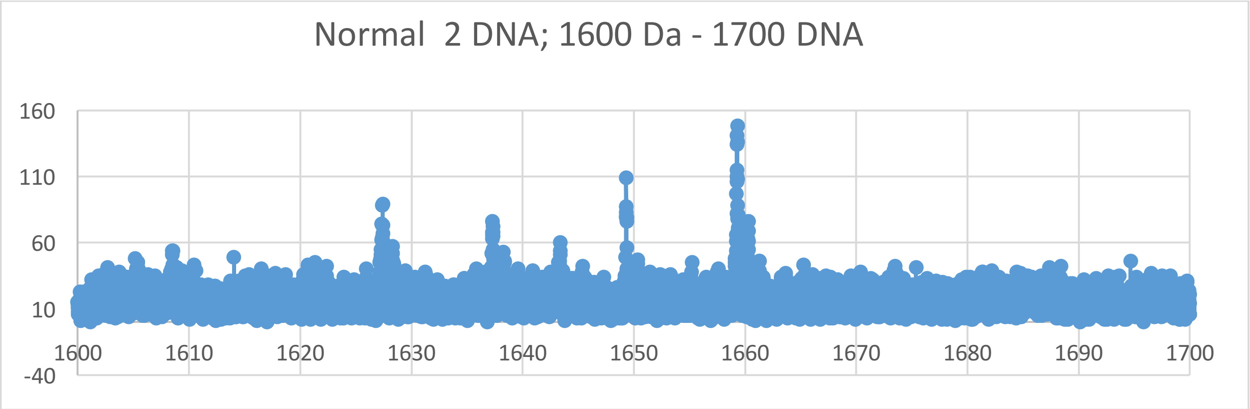
Normal 2 DNA; 1600 Da - 1700 DNA
160
110
60
10
-40
1600
1610
1620
1630
1640
1650
1660
1670
1680
1690
1700

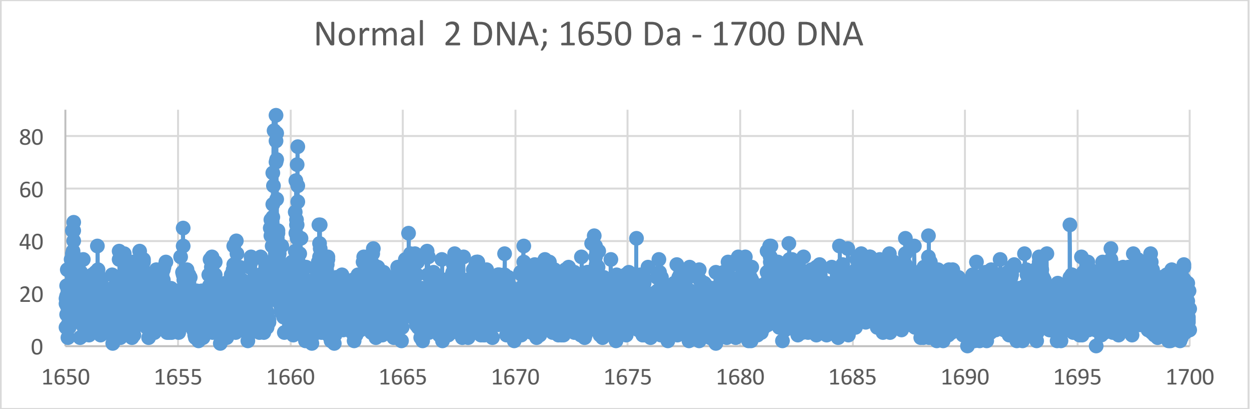
Normal 2 DNA; 1650 Da - 1700 DNA
80
60
40
20
0
1650
1655
1660
1665
1670
1675
1680
1685
1690
1695
1700

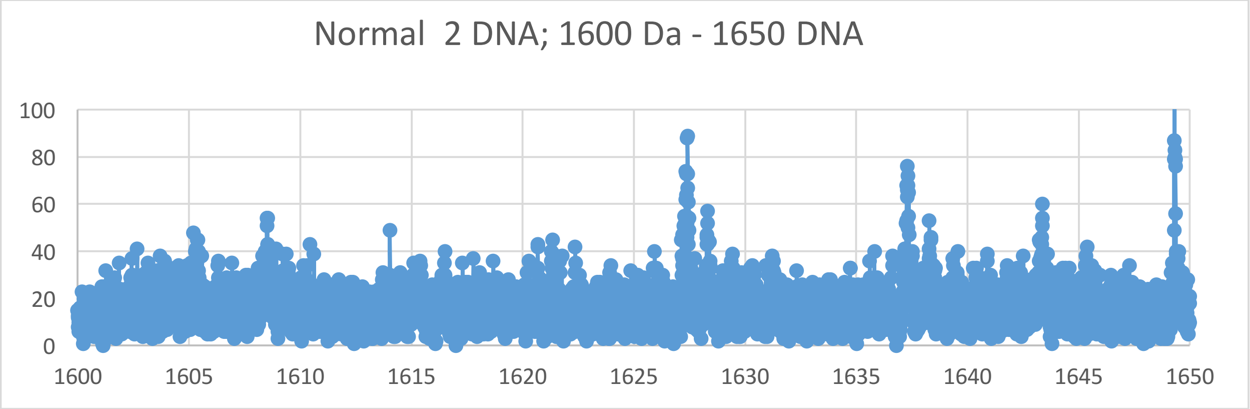
Normal 2 DNA; 1600 Da - 1650 DNA
100
80
60
40
20
0
1600
1605
1610
1615
1620
1625
1630
1635
1640
1645
1650

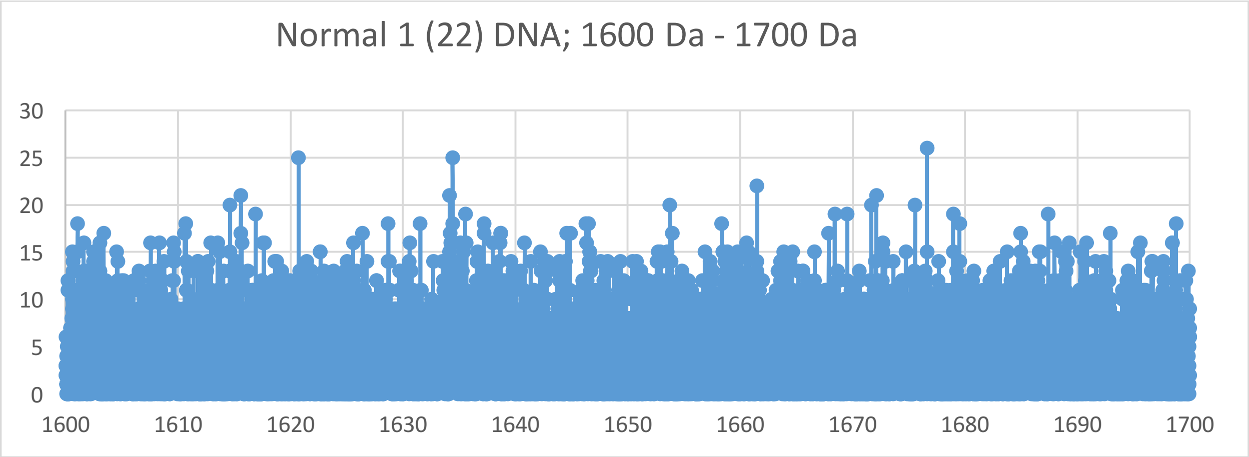

Normal 1 (22) DNA; 1600 Da - 1700 Da
30
25
20
15
10
5
0
1600
1610
1620
1630
1640
1650
1660
1670
1680
1690
1700


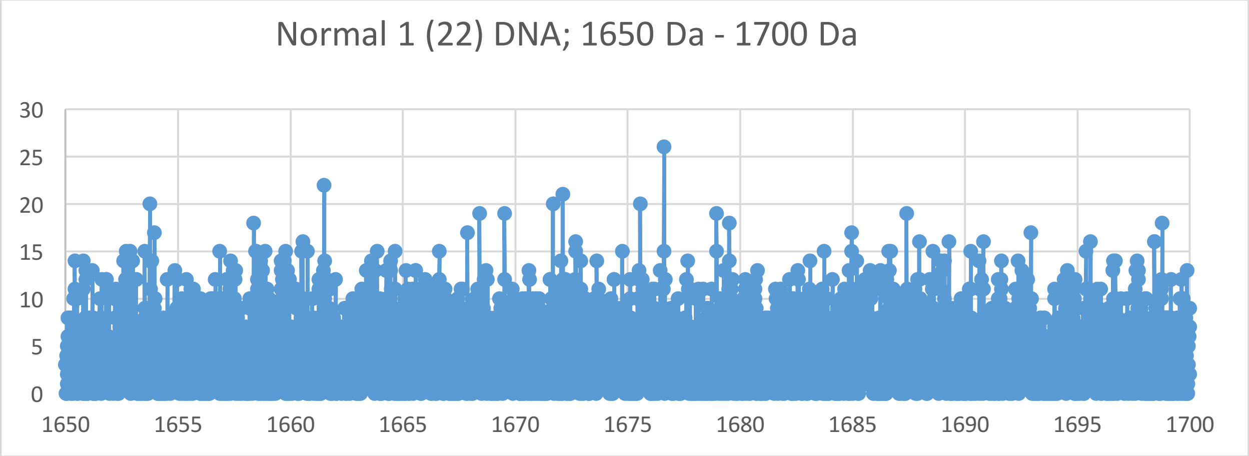

Normal 1 (22) DNA; 1650 Da - 1700 Da
30
25
20
15
10
5
0
1650
1655
1660
1665
1670
1675
1680
1685
1690
1695
1700


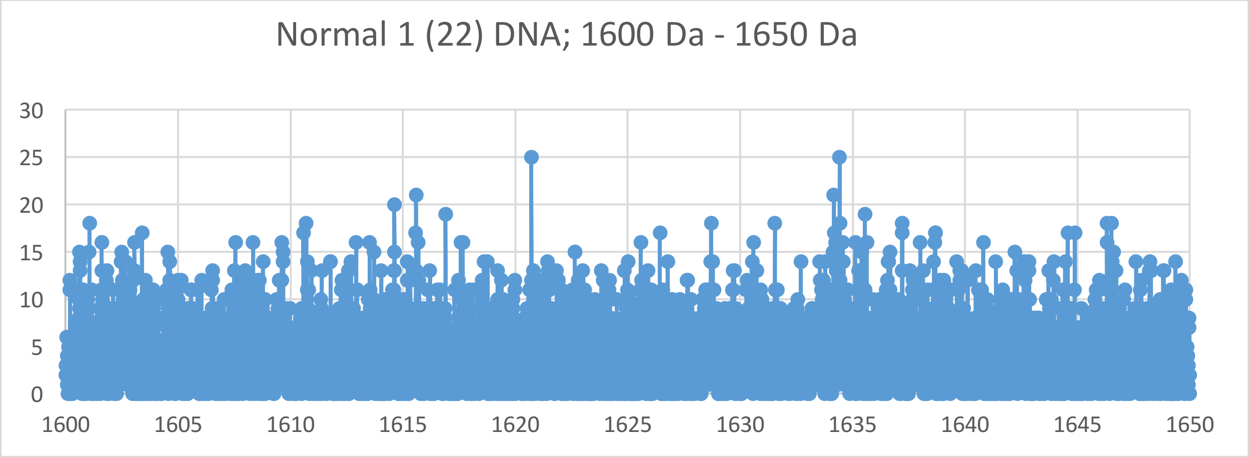

Normal 1 (22) DNA; 1600 Da - 1650 Da
30
25
20
15
10
5
0
1600
1605
1610
1615
1620
1625
1630
1635
1640
1645
1650
