## Supplementary Materials V for "Oncogenesis and Aging by Isotopic Functionalizations of the Proteins and Nucleic Acids"

V Supplementary Materials –Mass Spectra from 1400 Da -1600 Da

Results for Supplementary 5

So for the tetranucleotides from $(phos)_7$ to $(phos)_6$, the telomeric and nontelomeric peaks for cancer increase in number and the peaks for normal decrease in number for more stable normal tetranucleotides and less stable cancerous tetranucleotides with these dephosphorylations. The masses range from 1400 Da to 1600 Da. See Supplementary Figures V for graphs. These results and observations point specific differences between cancerous and normal oligonucleotides chemically and isotopically. It seems that below some number of phosphates, the telomeres and nontelomers behave in same way toward cancer and normal. So this suggest normal may be favored and distinguished from cancer by high phosphorylations. The mating telomeric peaks for normal increase with these dephosphorylations of the tetranucleotides. These dephosphorylations from $(phos)_7$ to $(phos)_6$ caused the resulting more intense and stable cancer and normal mixed tetranucleotides to be more nontelomeric and telomeric and negligibly isotopically enriched; but the mating telomeres are more slightly intense in both normal tetranucleotides and negligibly enriched isotopically. The mating telomeres are not as stabilized with cancer, mating telomeric tetranucleotides are negligibly isotopic enriched by these dephosphorylations. These dephosphorylations from $(phos)_7$ to $(phos)_6$ also caused the nontelomeric tetranucleotides to be more from intense in cancer DNA and more isotopically enriched. It seems this dephosphorylations are related to the isotopic enrichments and depletions; but the telomeres are more intense in cancer and less isotopically enriched whereas the intensity of the mating telomeres are less intense in cancer altered and the isotopic enrichments are greater. The telomeric tetranucleotides of cancer tended to also be less enriched so it seems the cancer stabilities of the telomeric tetranucleotides are related to there negligible isotopic enrichments for these dephosphorylations and purely driven by dephosphorylations. Enrichments weaken the nontelomeric normal tetranucleotides while negligibly strengthening cancerous telomeric tetranucleotides and weakening mating cancerous telomeric tetranucleotides. The more intense normal nontelomeric tetranucleotides were also slightly less isotopically enriched. These dephosphorylations from $(phos)_7$ to $(phos)_6$ slightly increased the isotopic enrichments of the cancer relative to the normal for the nontelomeric tetranucleotides. But the mating telomeric tetranucleotides were slightly enriched from these dephosphorylations for less enrichments and stronger more stable normal tetranucleotides for mating telomeres. Such seems compatible for replications and transcriptions as stable telomeres seem compatible to unstable mating telomeres for the unravelings and constructions of DNA mating telomeres onto the building templates of the telomeres. These dephosphorylations from $(phos)_7$ to $(phos)_6$ caused methylations to cause more normal intensities and stabilities but negligibly less isotopic enrichments for telomeres, mating telomeres and nontelomeres and (slightly for nontelomers) with large effects on mating telomeric tetranucleotides as the intensities of normal increased with negligible isotopic enrichments. But the methylations of the mating telomeres increased normal stabilities and with negligible isotopic enrichments for similar effects on mating telomeres compared to telomeres. This dephosphorylations from $(phos)_7$ to $(phos)_6$ caused acetylations to caused negligible changes relative in normal and cancer stabilities for telomeres and similar negligible changes in mixed cancer and normal stabilities of mating telomeres; the mating telomere are more cancer stable and more isotopically enriched by the acetylations. Thereby this dephosphorylations $(phos)_7$ to $(phos)_6$ and there varying consequences with respect to other dephosphorylations determines patterns of dephosphorylations and isotopic enrichment patterns in normal DNA, which mutate and change for different patterns in DNA of cells that become cancerous. These dephosphorylations $(phos)_7$ to $(phos)_6$ induced isotopic dynamics from varying methylations and acetylations in cancer DNA relative to normal DNA couple to other isotopically induced varied phosphorylations, methylations and acetylations throughout the DNA for emerging order from nanodisorder.

So for the tetranucleotides from $(phos)_6$ to $(phos)_5$, the telomeric, mating telomeric and nontelomeic peaks for cancer increase in number and the peaks for normal decrease in number for more stable normal tetranucleotides and less stable cancerous tetranucleotides with these dephosphorylations. The masses range from 1300 Da to 1550 Da. See Supplementary Figures V for graphs. These results and observations point specific differences between cancerous and normal oligonucleotides chemically and isotopically. It seems that below some number of phosphates, the telomeres, mating telomeres and nontelomers behave in same way toward cancer and normal. So this suggests normal may be favored and distinguished from cancer by high phosphorylations. The mating telomeric peaks for normal decrease with these dephosphorylations of the tetranucleotides. These dephosphorylations from $(phos)_6$ to $(phos)_5$ caused the resulting more intense and stable cancer and normal mixed tetranucleotides to be more nontelomeric and telomeric and more isotopically enriched; but the mating telomeres are more intense in both normal and cancer tetranucleotides and enriched isotopically. The mating telomeric are stabilized with cancer, mating telomeric tetranucleotides are isotopically enriched by these dephosphorylations. These dephosphorylations from $(phos)_6$ to $(phos)_5$ also caused the nontelomeric tetranucleotides to be more intense in cancer DNA and more isotopically enriched. It seems these dephosphorylations are related to the isotopic enrichments and depletions; but the telomeres are more intense in cancer and less isotopically enriched whereas the intensities of the mating telomeres are less intense in normal altered and the isotopic enrichments are negligible. The telomeric tetranucleotides of cancer tended to also be more enriched so it seems the cancer stabilities of the telomeric tetranucleotides are related to there more isotopic enrichments for this dephosphorylations and purely driven by dephosphorylations and enrichments. Enrichments strengthen the nontelomeric normal tetranucleotides while slightly weakening cancerous telomeric and mating telomeric teranucleotides. The more intense normal nontelomeric tetranucleotides were also isotopically enriched. This dephosphorylations from $(phos)_6$ to $(phos)_5$ increased the isotopic enrichments of the cancer and normal for the mating telomeric and telomeric tetranucleotides. But the mating telomeric tetranucleotides were enriched from these dephosphorylations for more enrichments and stronger more stable cancer tetranucleotides for mating telomeres. Such seems compatible for replications and transcriptions as stable telomeres seem compatible to unstable mating telomeres for the unravelings and constructions of DNA mating telomeres onto the building templates of the telomeres. These dephosphorylations from $(phos)_6$ to $(phos)_5$ caused methylations to cause more normal intensities and stabilities but slightly more isotopic enrichments for nontelomeres. But the methylations of the telomeres and mating telomeres slightly increased cancer stabilities and dramatically more isotopic enrichments for mating telomeres and telomeres. These dephosphorylations from $(phos)_6$ to $(phos)_5$ caused acetylations to cause less cancer stabilities and more normal stabilities for telomeres, mating telomeres and nontelomeres with more isotopic enrichments relative to methylations for telomeres and nontelomeres; the mating telomeres are more cancer stable and more isotopically enriched by the acetylations. Thereby these dephosphorylations $(phos)_6$ to $(phos)_5$ and these varying consequences with respect to other dephosphrylations determine patterns of dephosphorylations and isotopic enrichment patterns in normal DNA which mutate and change for different patterns in DNA of cells that become cancerous. These dephosphorylations $(phos)_6$ to $(phos)_5$ induced isotopic dynamics from varying methylations and acetylations in cancer DNA relative to normal DNA couple to other isotopically induced varied phosphorylations, methylations and acetylations throughout the DNA for emerging order from nanodisorder.

This mass range has tetranucleotides overlapping the trinucleotides in similar masses. So for the trinucleotides at $(phos)_9$, the telomeric peaks for normal strongly increase in number and the peaks for cancer decreases in number for more stable normal trinucleotides and less stable cancer trinucleotides with the dephosphorylations. But the mating telomeres and nontelomeres have decrease in numbers and stabilities for normal trinucleotides while the cancer trinucleotides are greater in peaks and stabilities. But as decrease the phosphate by 1 unit, the trinucleotides change so that the cancer trinucleotides fragment and break from the DNA with more number and stabilities for the telomeres, mating telomeres and nontelomeres from $(phos)_9$ to $(phos)_8$. The masses range from 1310 Da to 1550 Da. See Supplementary Figures 5 for graphs. It seems that below some number of phosphates, the telomeres, mating telomeres and nontelomers behave in same way toward cancer and normal. So these suggest normal may be favored and distinguished from cancer by high (or changing) saturated phosphorylations. The telomeric peaks for cancer increase with these dephosphorylations of the trinucleotides. The dephosphorylations from $(phos)_9$ to $(phos)_8$ caused the resulting more intense and stable cancer trinucleotides to be for mating telomeric, telomeric and nontelomeric trinucleotides and more isotopically enriched for telomeres, mating telomeres and nontelomeres. It is important to consider that there are more intense peaks for the trinucleotides relative to the tetranucleotides for the saturated phosphate units. The dephosphorylations from $(phos)_9$ to $(phos)_8$ increases isotopic enrichments of heavy isotopes of $^{13}C$ and $^{17}O$. The dephosphorylations from $(phos)_9$ to $(phos)_8$ also caused the telomeric, mating telomeric and nontelomeric trinucleotides to be more intense in cancer DNA and more isotopically enriched. It seems the dephosphorylations are related to the isotopic enrichments and depletions; but the telomeres are more intense in cancer and more isotopically enriched whereas the intensities of the mating telomeres and nontelomeres are less intense in cancer altered and the isotopic enrichments are more. The telomeric, mating telomeric and nontelomeric trianucleotides of cancer tended to also be more enriched so it seems the cancer stabilities of the telomeric, mating telomeric and nontelomeric trinucleotides are related to their more isotopic enrichments for the dephosphorylations from $(phos)_9$ to $(phos)_8$ and purely driven by dephosphorylations and enrichments. Enrichments strengthen the nontelomeric, telomeric and mating telomeric cancer trinucleotides while strongly weakening normal telomeric and nontelomeric trinucleotides. The more intense cancer nontelomeric, telomeric and nontelomeric trinucleotides were also more isotopically enriched. The dephosphorylations from $(phos)_9$ to $(phos)_8$ increased the isotopic enrichments of the cancer for the nontelomeric, mating telomeric and telomeric trinucleotides. But the nontelomeric trinucleotides were more enriched from the dephosphorylations for more enrichments and stronger more stable cancer trinucleotides for nontelomeres. Such seems compatible for replications and transcriptions as stable telomeres seem compatible to unstable mating telomeres for the unravelings and constructions of DNA mating telomeres onto the building templates of the telomeres. The dephosphorylations from $(phos)_9$ to $(phos)_8$ caused methylations to cause slightly more normal intensities and stabilities but slightly less isotopic enrichments for telomeres and nontelomeres. But the methylations of the mating telomeres increased cancer stabilities and greater isotopic enrichments for mating telomere. The dephosphorylations from $(phos)_9$ to $(phos)_8$ caused acetylations to cause more normal stabilities for nontelomeres and more cancer stabilities for telomeres and slightly more normal stabilities for the mating telomeres with more enrichments relative to methylations for telomeres and nontelomeres.

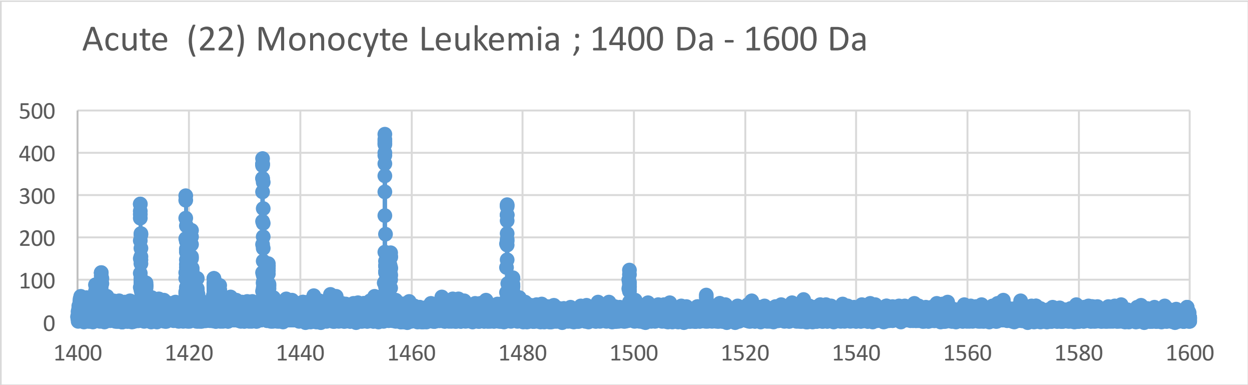
Acute (22) Monocyte Leukemia ; 1400 Da - 1600 Da
500
400
300
200
100
0
1400 1420 1440 1460 1480 1500 1520 1540 1560 1580 1600

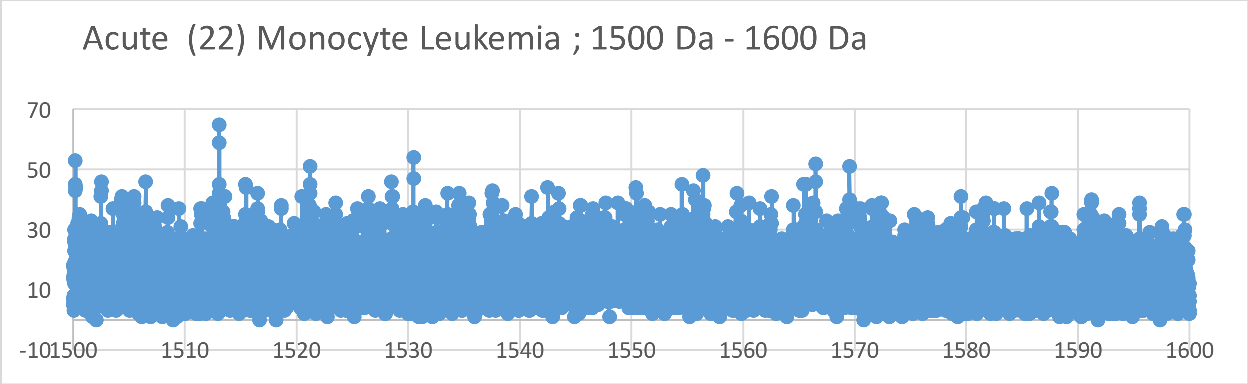
Acute (22) Monocyte Leukemia ; 1500 Da - 1600 Da
70
50
30
10
-10
1500 1510 1520 1530 1540 1550 1560 1570 1580 1590 1600

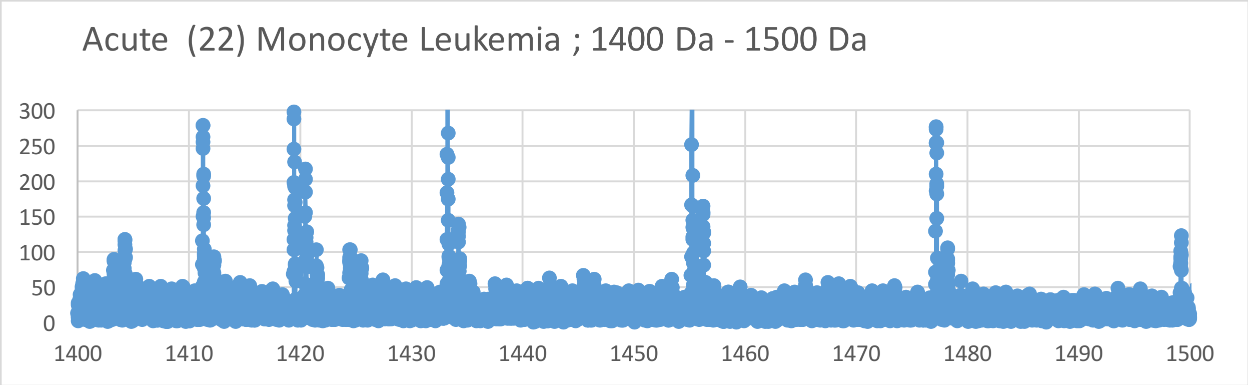
Acute (22) Monocyte Leukemia ; 1400 Da - 1500 Da
300
250
200
150
100
50
0
1400 1410 1420 1430 1440 1450 1460 1470 1480 1490 1500

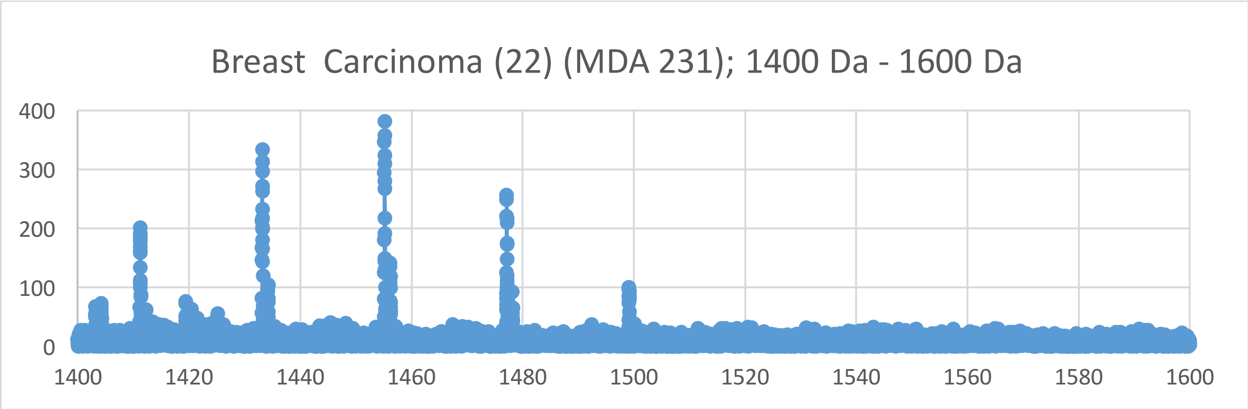

Breast  Carcinoma (22) (MDA 231); 1400 Da - 1600 Da
400
300
200
100
0
1400
1420
1440
1460
1480
1500
1520
1540
1560
1580
1600


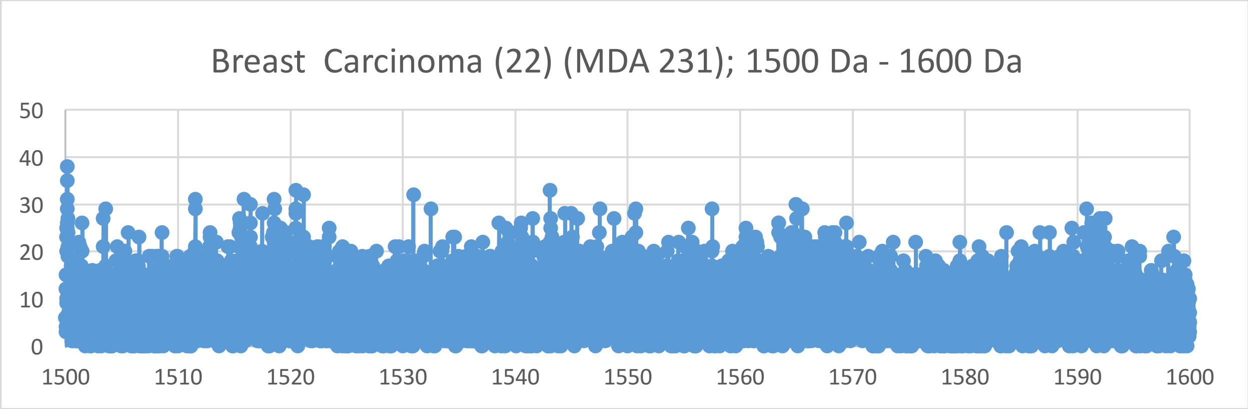

Breast  Carcinoma (22) (MDA 231); 1500 Da - 1600 Da
50
40
30
20
10
0
1500
1510
1520
1530
1540
1550
1560
1570
1580
1590
1600


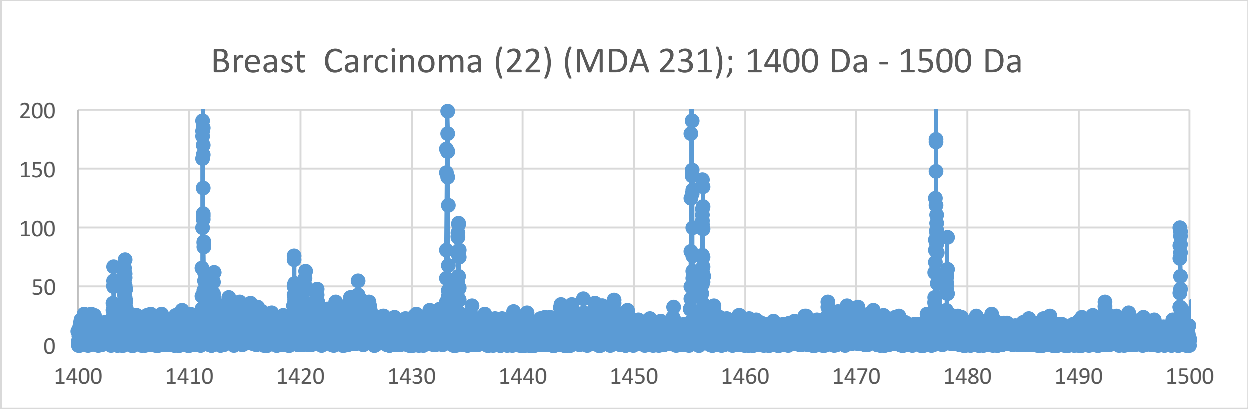

Breast  Carcinoma (22) (MDA 231); 1400 Da - 1500 Da
200
150
100
50
0
1400
1410
1420
1430
1440
1450
1460
1470
1480
1490
1500

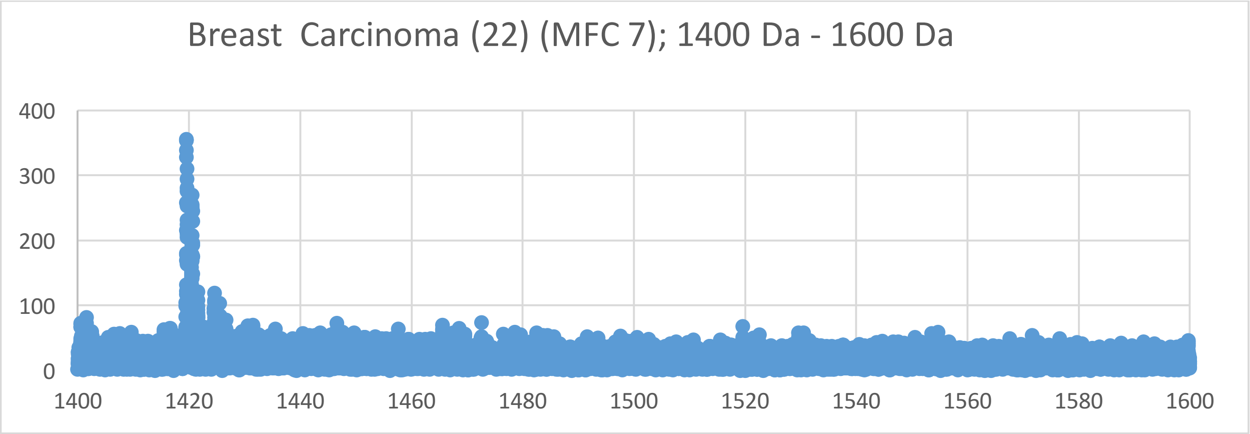
Breast Carcinoma (22) (MFC 7); 1400 Da - 1600 Da
400
300
200
100
0
1400
1420
1440
1460
1480
1500
1520
1540
1560
1580
1600

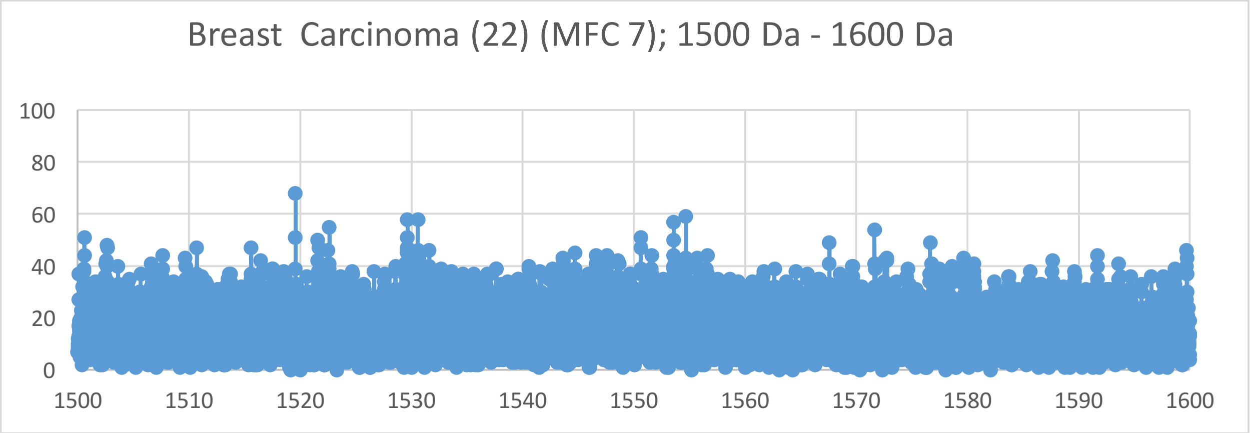
Breast Carcinoma (22) (MFC 7); 1500 Da - 1600 Da
100
80
60
40
20
0
1500
1510
1520
1530
1540
1550
1560
1570
1580
1590
1600

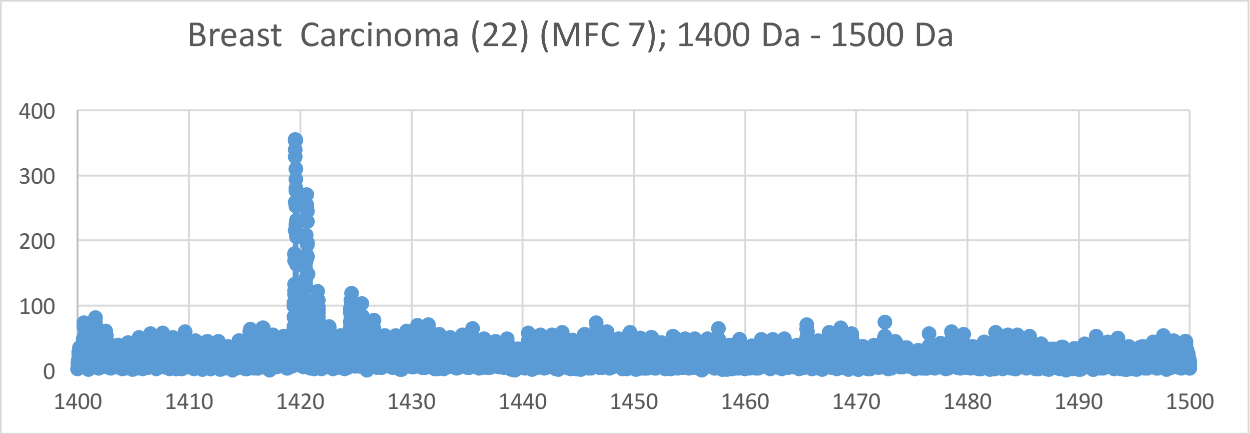
Breast Carcinoma (22) (MFC 7); 1400 Da - 1500 Da
400
300
200
100
0
1400
1410
1420
1430
1440
1450
1460
1470
1480
1490
1500

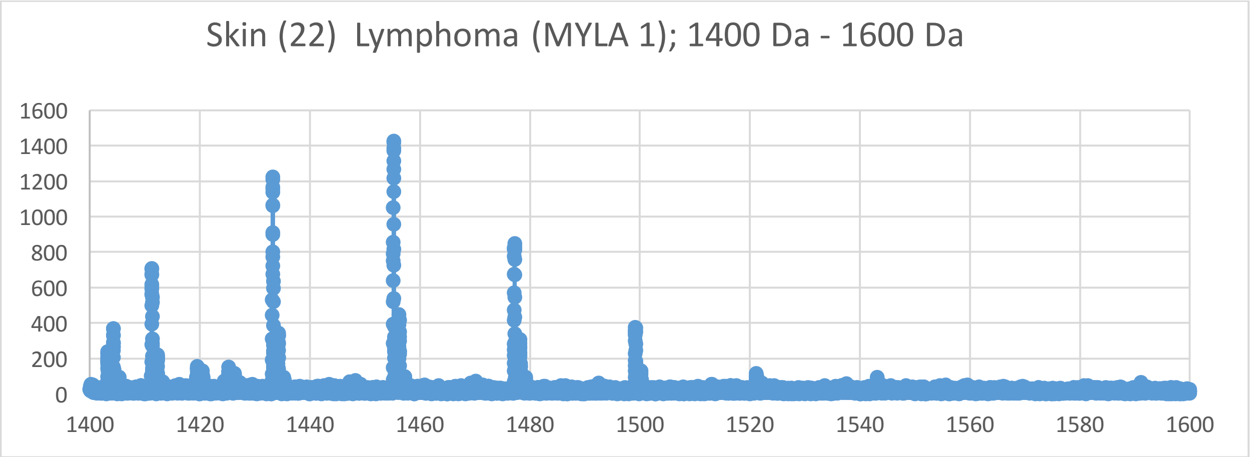
Skin (22) Lymphoma (MYLA 1); 1400 Da - 1600 Da
0
200
400
600
800
1000
1200
1400
1600
1400
1420
1440
1460
1480
1500
1520
1540
1560
1580
1600

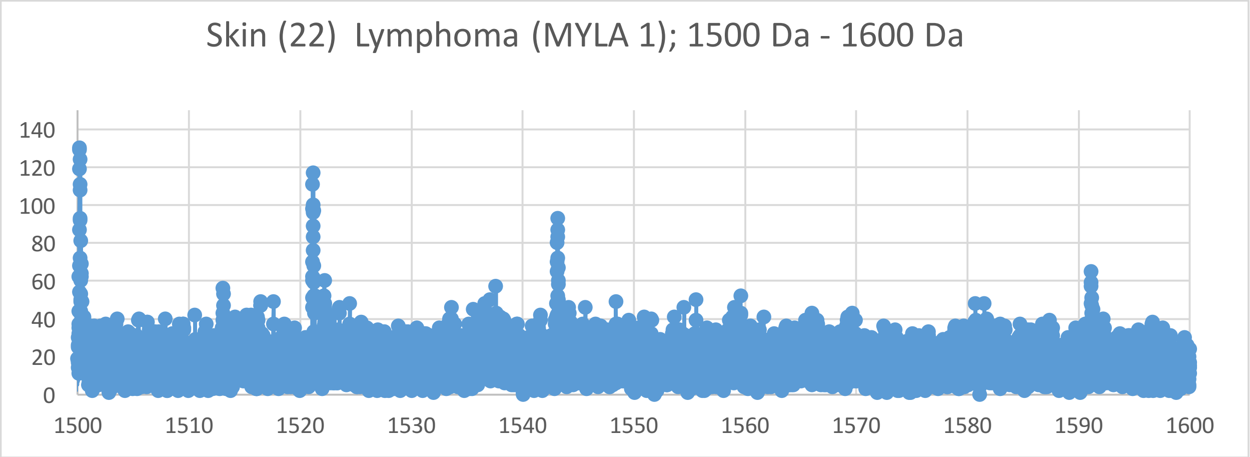
Skin (22) Lymphoma (MYLA 1); 1500 Da - 1600 Da
0
20
40
60
80
100
120
140
1500
1510
1520
1530
1540
1550
1560
1570
1580
1590
1600

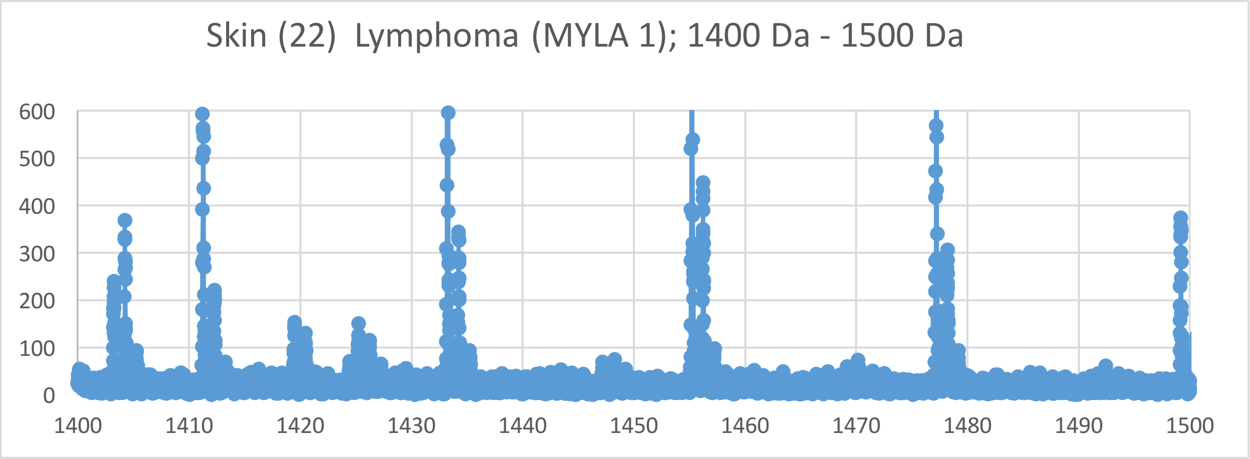
Skin (22) Lymphoma (MYLA 1); 1400 Da - 1500 Da
0
100
200
300
400
500
600
1400
1410
1420
1430
1440
1450
1460
1470
1480
1490
1500

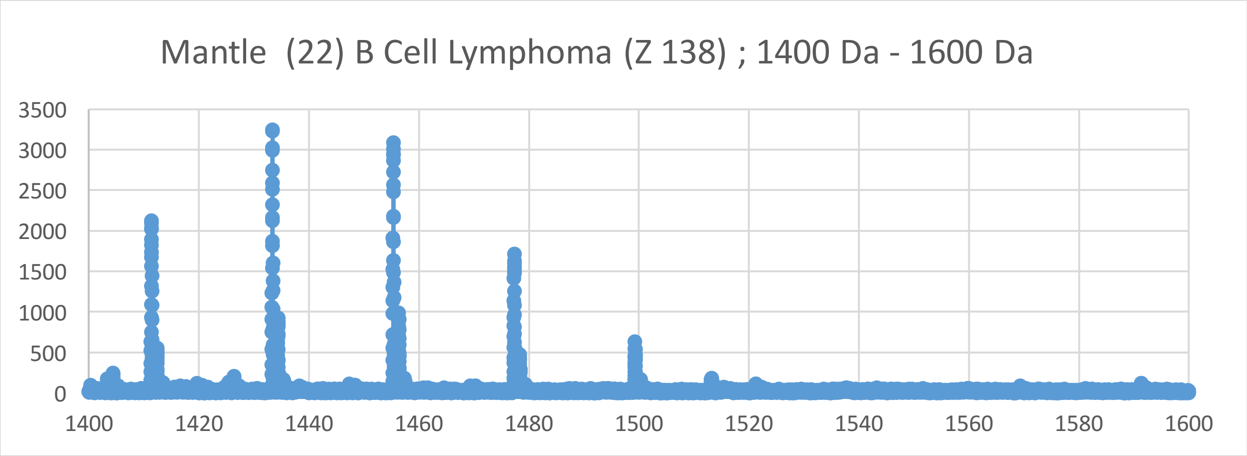
Mantle (22) B Cell Lymphoma (Z 138) ; 1400 Da - 1600 Da
3500
3000
2500
2000
1500
1000
500
0
1400
1420
1440
1460
1480
1500
1520
1540
1560
1580
1600

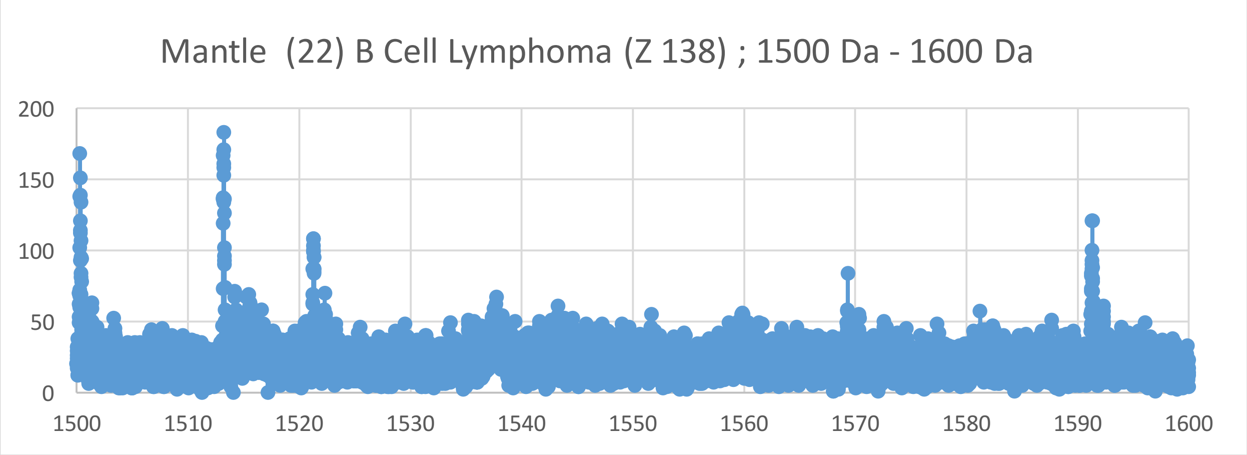
Mantle (22) B Cell Lymphoma (Z 138) ; 1500 Da - 1600 Da
200
150
100
50
0
1500
1510
1520
1530
1540
1550
1560
1570
1580
1590
1600

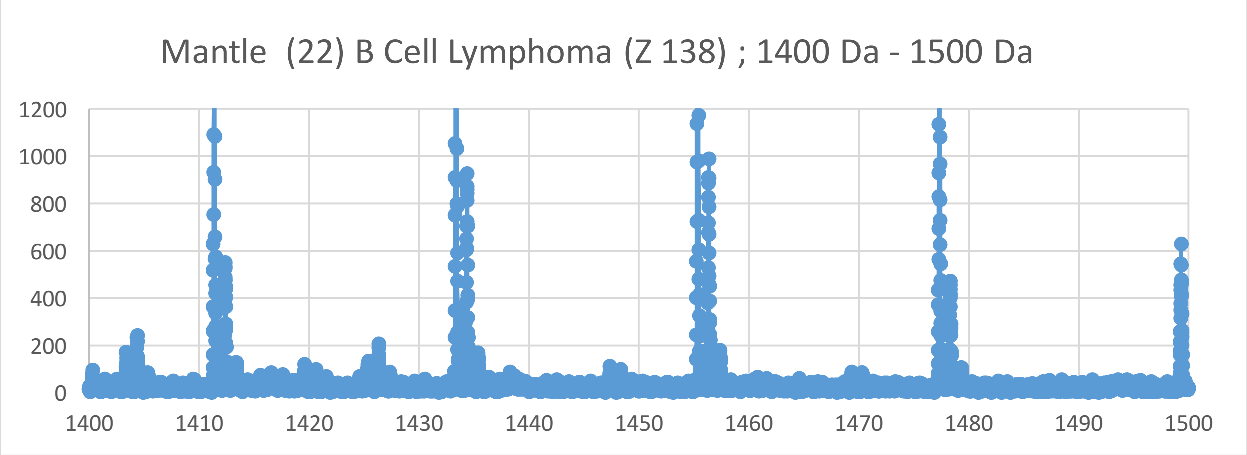
Mantle (22) B Cell Lymphoma (Z 138) ; 1400 Da - 1500 Da
1200
1000
800
600
400
200
0
1400
1410
1420
1430
1440
1450
1460
1470
1480
1490
1500

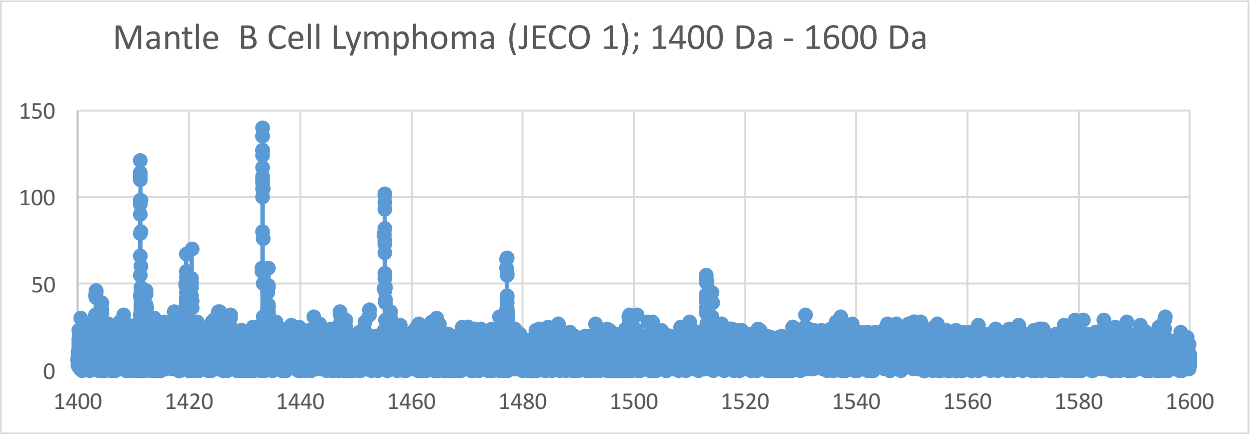
Mantle B Cell Lymphoma (JECO 1); 1400 Da - 1600 Da
150
100
50
0
1400
1420
1440
1460
1480
1500
1520
1540
1560
1580
1600

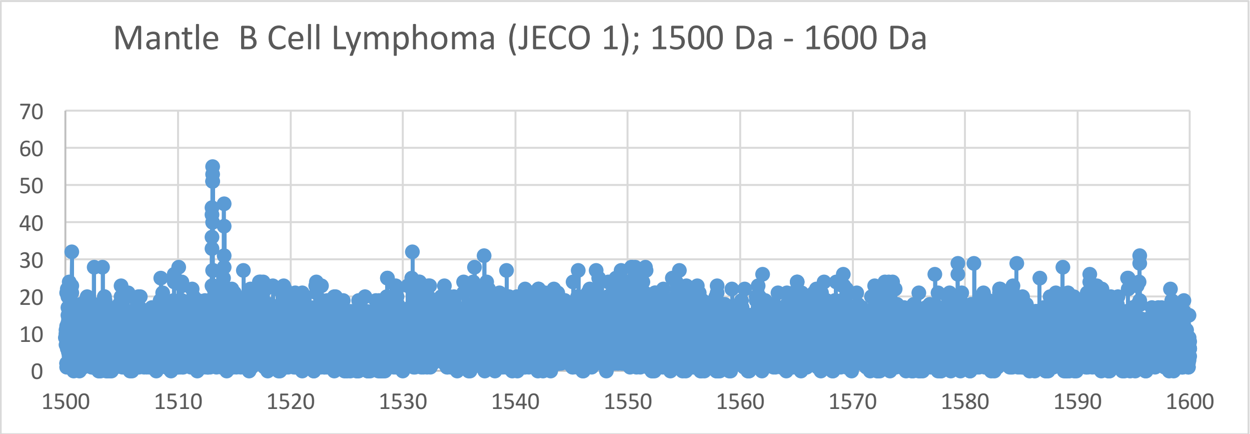
Mantle B Cell Lymphoma (JECO 1); 1500 Da - 1600 Da
70
60
50
40
30
20
10
0
1500
1510
1520
1530
1540
1550
1560
1570
1580
1590
1600

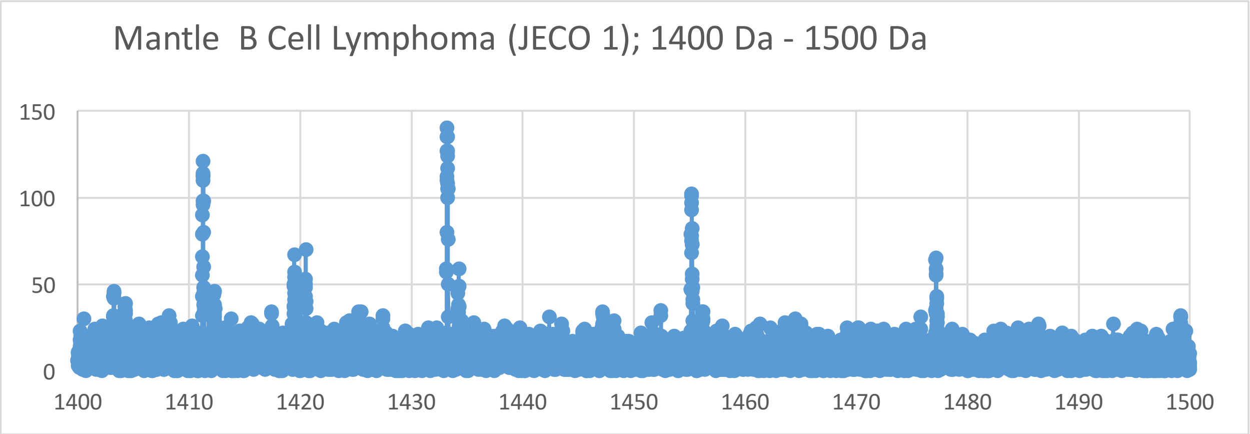
Mantle B Cell Lymphoma (JECO 1); 1400 Da - 1500 Da
150
100
50
0
1400
1410
1420
1430
1440
1450
1460
1470
1480
1490
1500

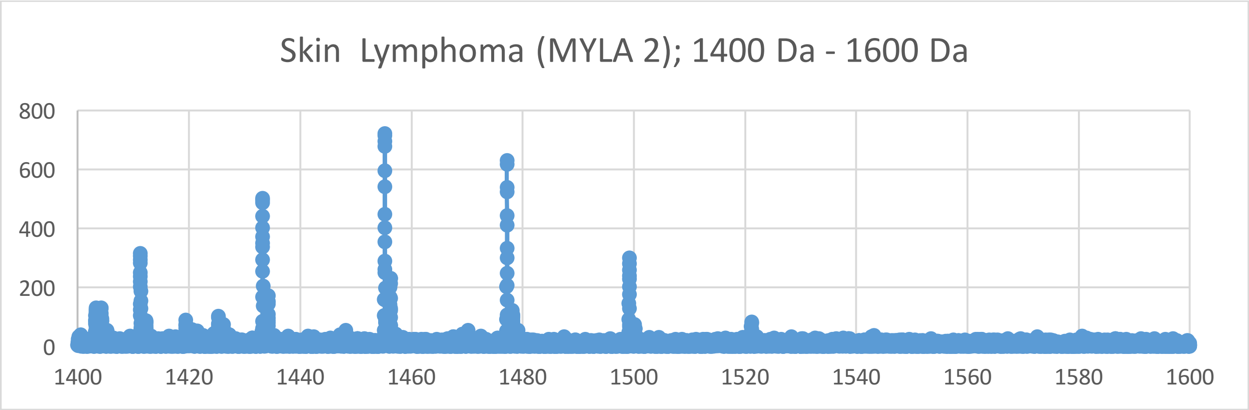
Skin Lymphoma (MYLA 2); 1400 Da - 1600 Da
800
600
400
200
0
1400
1420
1440
1460
1480
1500
1520
1540
1560
1580
1600

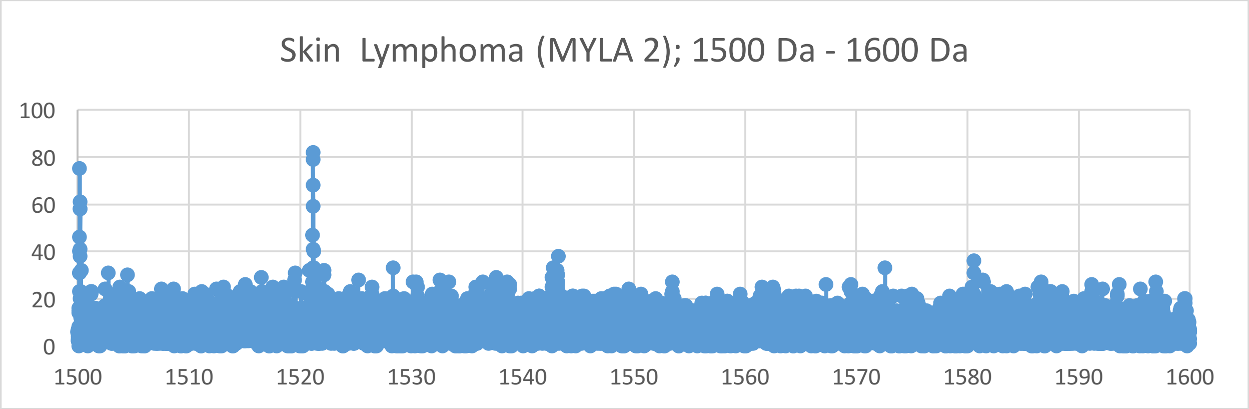
Skin Lymphoma (MYLA 2); 1500 Da - 1600 Da
100
80
60
40
20
0
1500
1510
1520
1530
1540
1550
1560
1570
1580
1590
1600

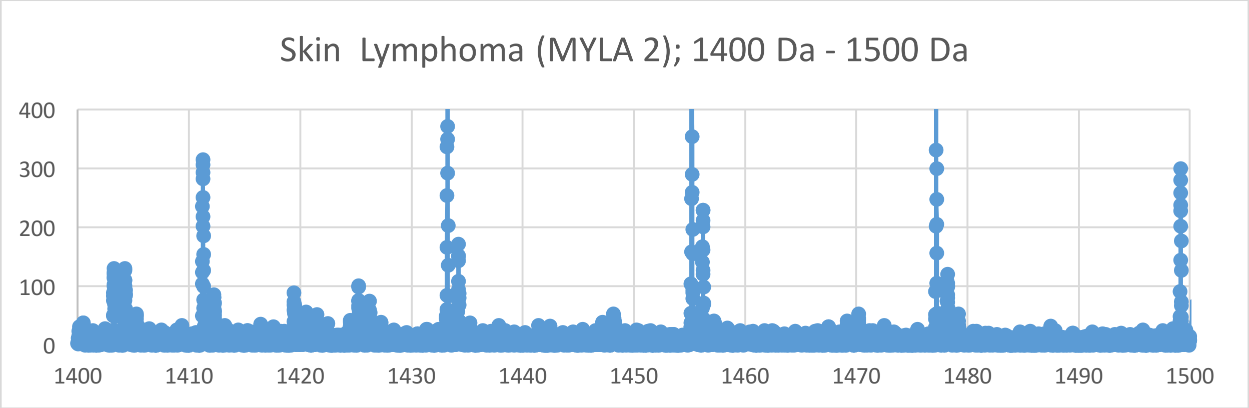
Skin Lymphoma (MYLA 2); 1400 Da - 1500 Da
400
300
200
100
0
1400
1410
1420
1430
1440
1450
1460
1470
1480
1490
1500

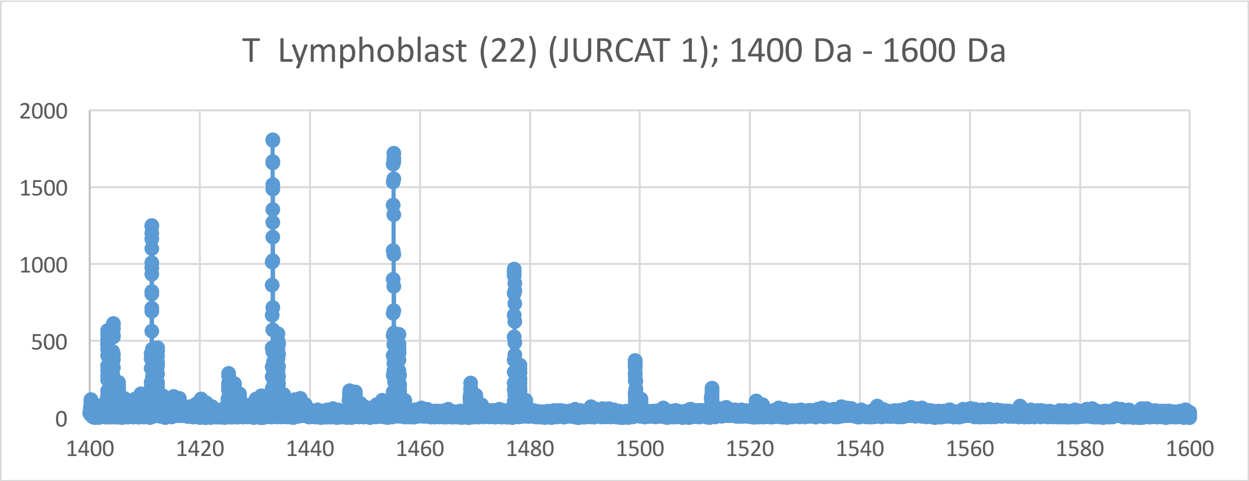
T Lymphoblast (22) (JURCAT 1); 1400 Da - 1600 Da
2000
1500
1000
500
0
1400
1420
1440
1460
1480
1500
1520
1540
1560
1580
1600

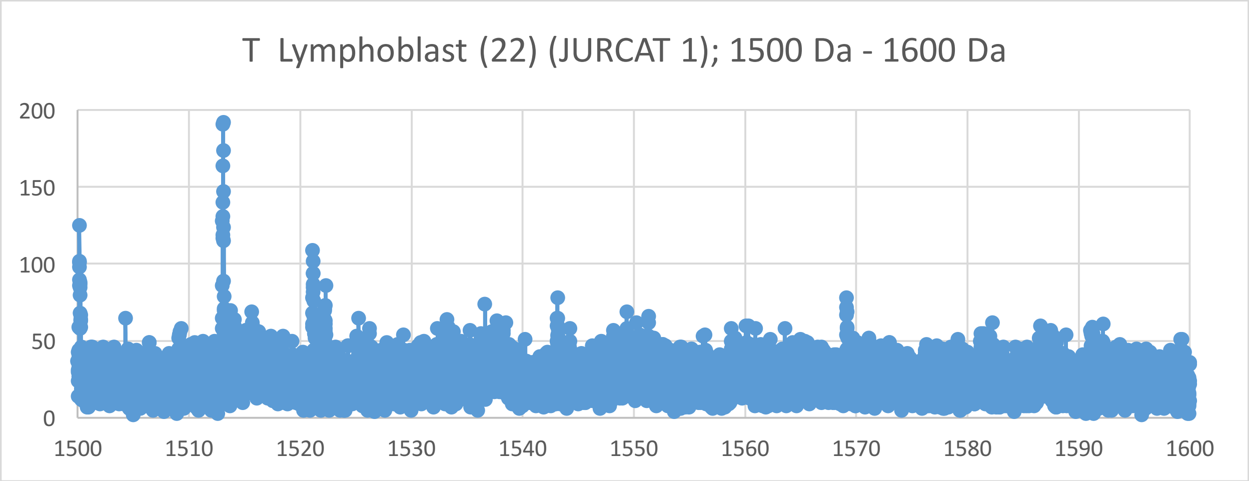
T Lymphoblast (22) (JURCAT 1); 1500 Da - 1600 Da
200
150
100
50
0
1500
1510
1520
1530
1540
1550
1560
1570
1580
1590
1600

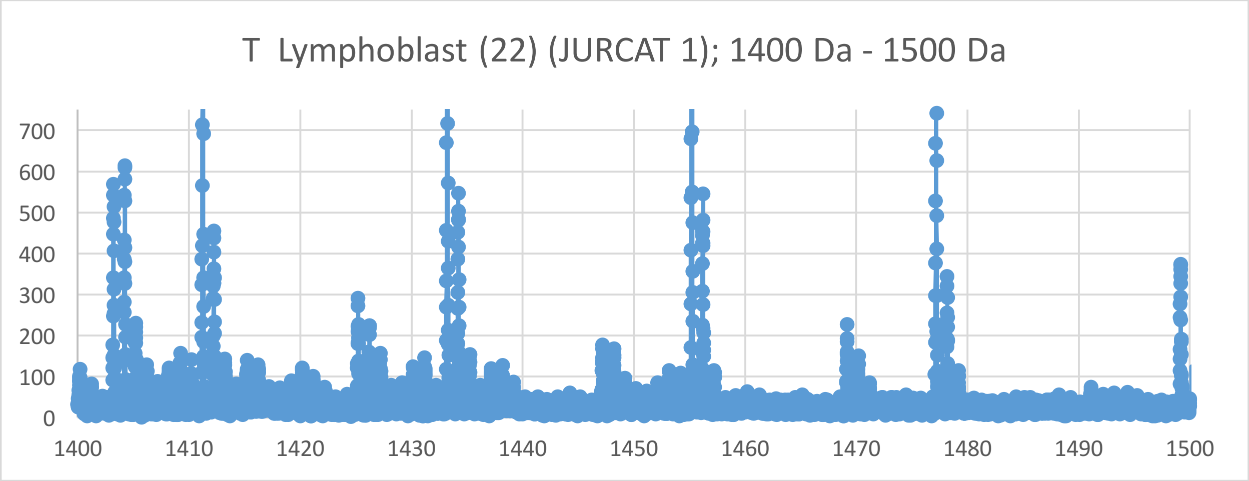
T Lymphoblast (22) (JURCAT 1); 1400 Da - 1500 Da
700
600
500
400
300
200
100
0
1400
1410
1420
1430
1440
1450
1460
1470
1480
1490
1500

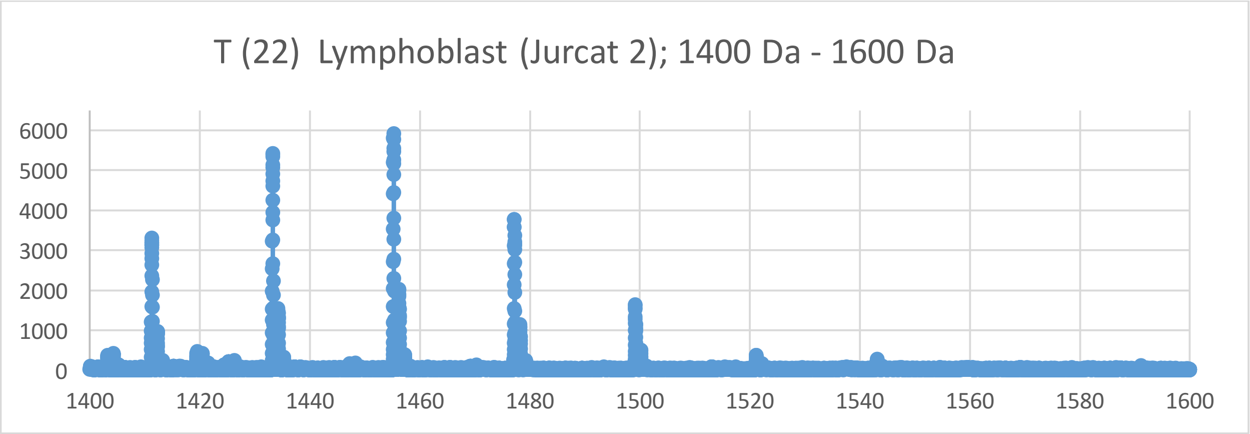
T (22) Lymphoblast (Jurcat 2); 1400 Da - 1600 Da
6000
5000
4000
3000
2000
1000
0
1400
1420
1440
1460
1480
1500
1520
1540
1560
1580
1600

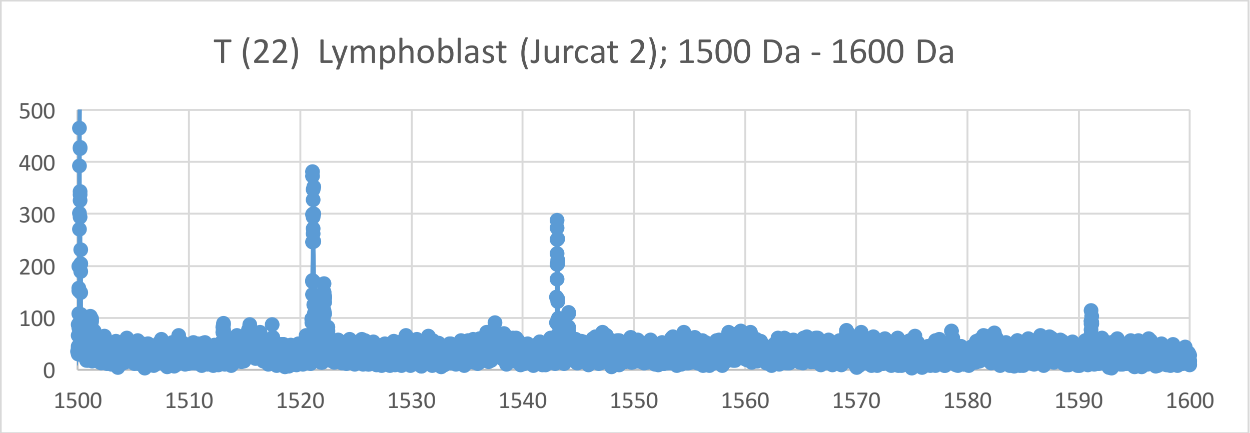
T (22) Lymphoblast (Jurcat 2); 1500 Da - 1600 Da
500
400
300
200
100
0
1500
1510
1520
1530
1540
1550
1560
1570
1580
1590
1600

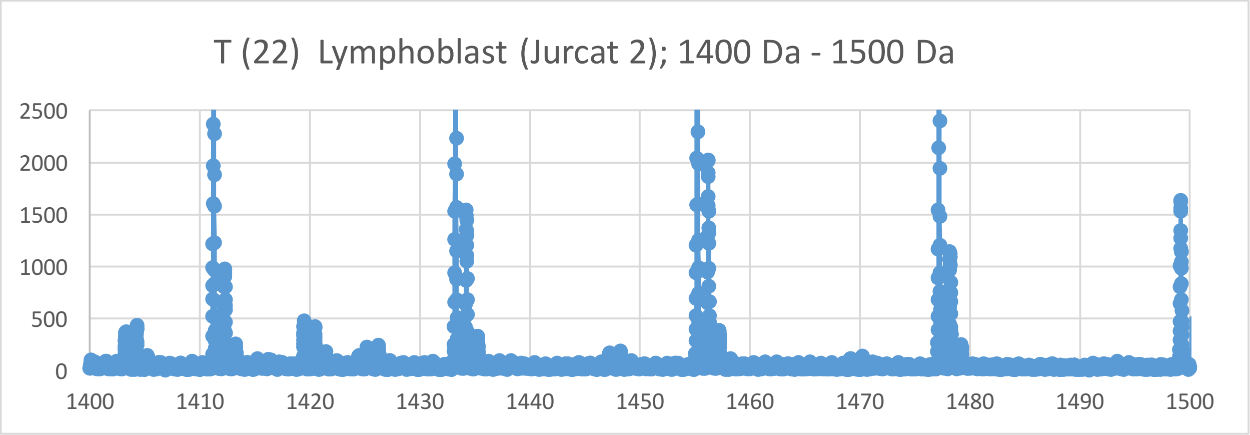
T (22) Lymphoblast (Jurcat 2); 1400 Da - 1500 Da
2500
2000
1500
1000
500
0
1400
1410
1420
1430
1440
1450
1460
1470
1480
1490
1500

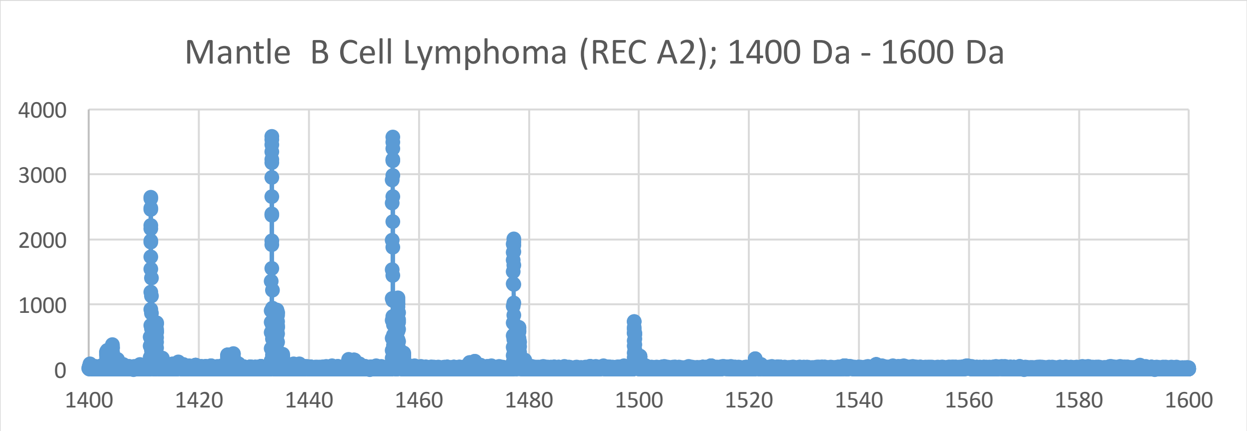
Mantle B Cell Lymphoma (REC A2); 1400 Da - 1600 Da
4000
3000
2000
1000
0
1400
1420
1440
1460
1480
1500
1520
1540
1560
1580
1600

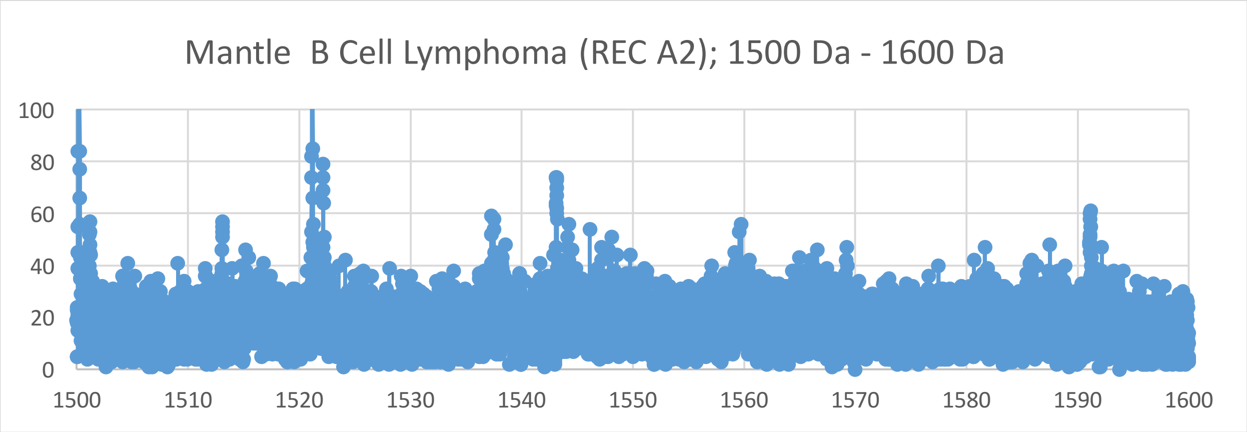
Mantle B Cell Lymphoma (REC A2); 1500 Da - 1600 Da
100
80
60
40
20
0
1500
1510
1520
1530
1540
1550
1560
1570
1580
1590
1600

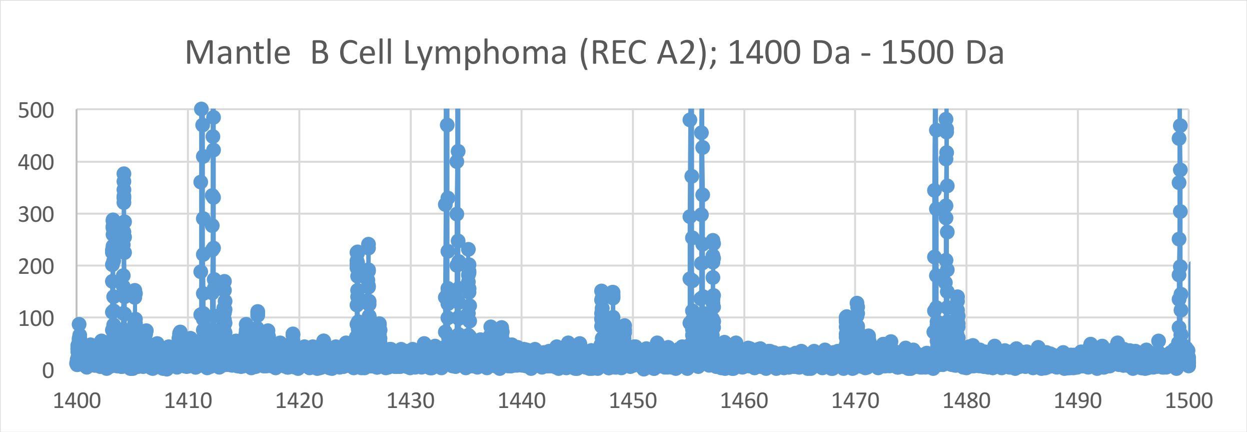
Mantle B Cell Lymphoma (REC A2); 1400 Da - 1500 Da
500
400
300
200
100
0
1400
1410
1420
1430
1440
1450
1460
1470
1480
1490
1500

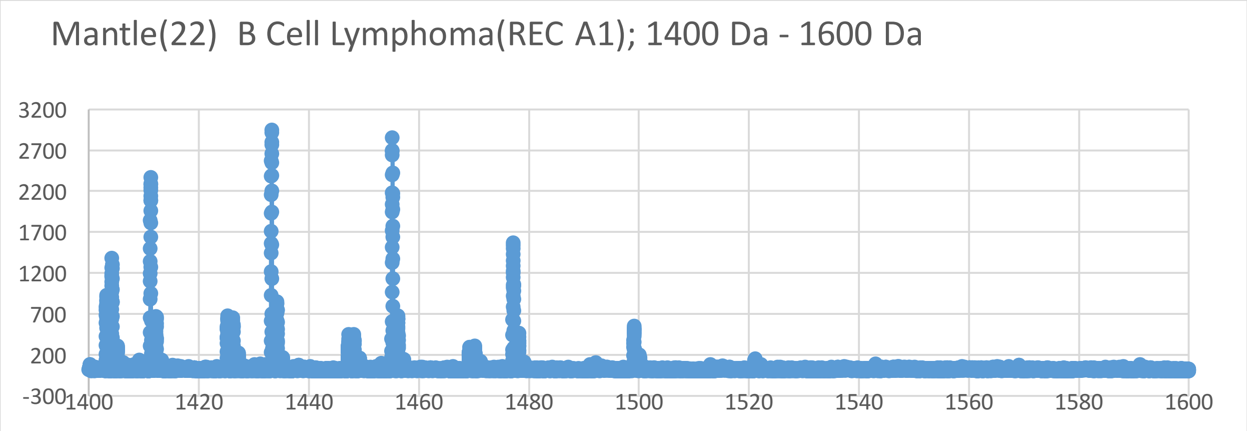
Mantle(22) B Cell Lymphoma(REC A1); 1400 Da - 1600 Da
3200
2700
2200
1700
1200
700
200
-300
1400
1420
1440
1460
1480
1500
1520
1540
1560
1580
1600

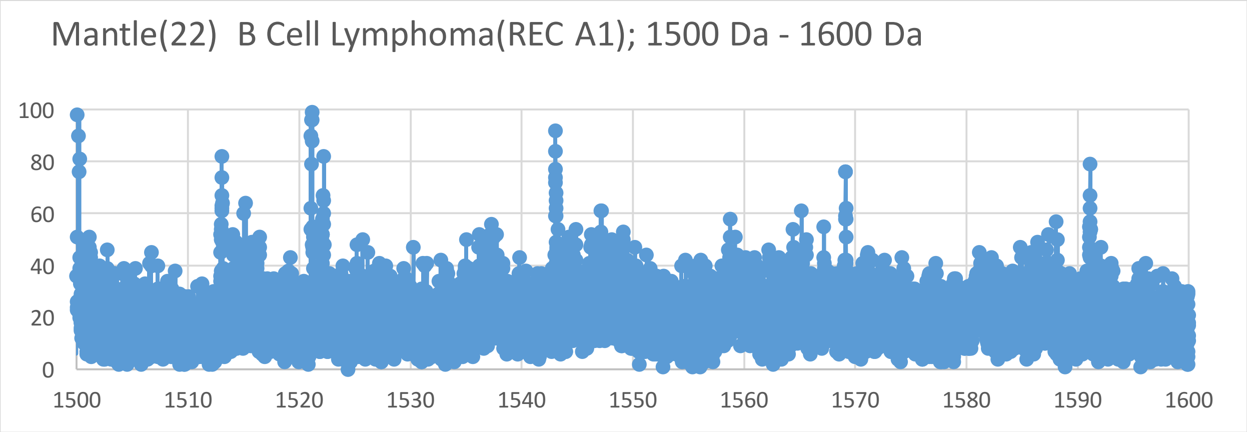
Mantle(22) B Cell Lymphoma(REC A1); 1500 Da - 1600 Da
100
80
60
40
20
0
1500
1510
1520
1530
1540
1550
1560
1570
1580
1590
1600

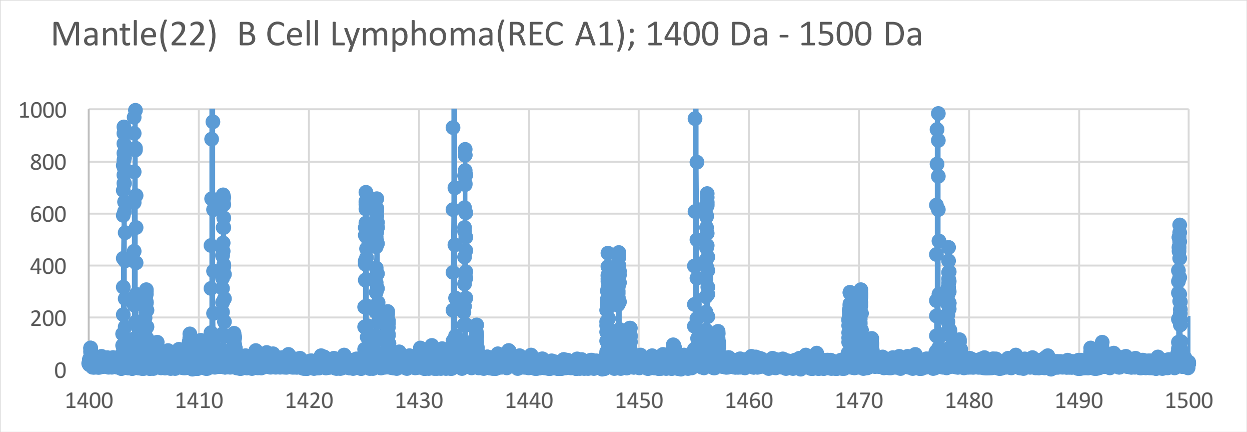
Mantle(22) B Cell Lymphoma(REC A1); 1400 Da - 1500 Da
1000
800
600
400
200
0
1400
1410
1420
1430
1440
1450
1460
1470
1480
1490
1500

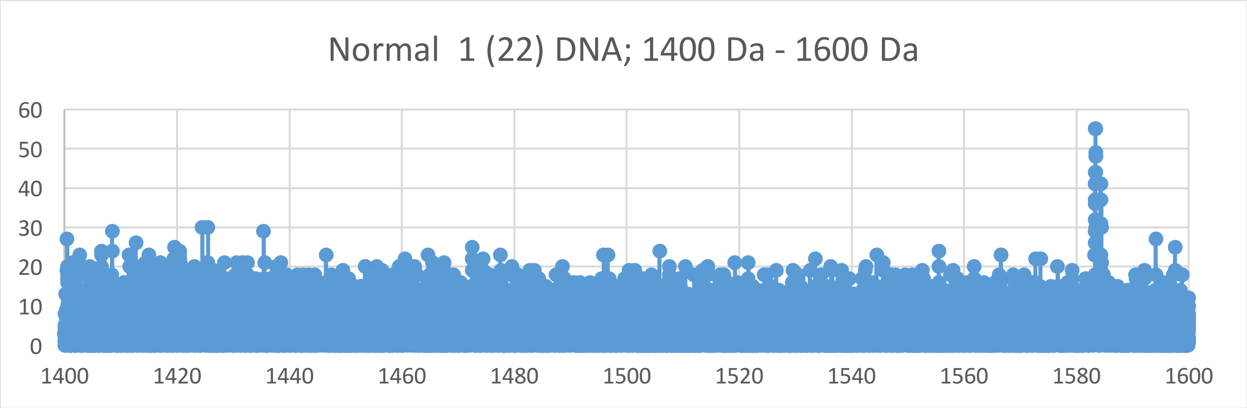
Normal 1 (22) DNA; 1400 Da - 1600 Da
60
50
40
30
20
10
0
1400
1420
1440
1460
1480
1500
1520
1540
1560
1580
1600

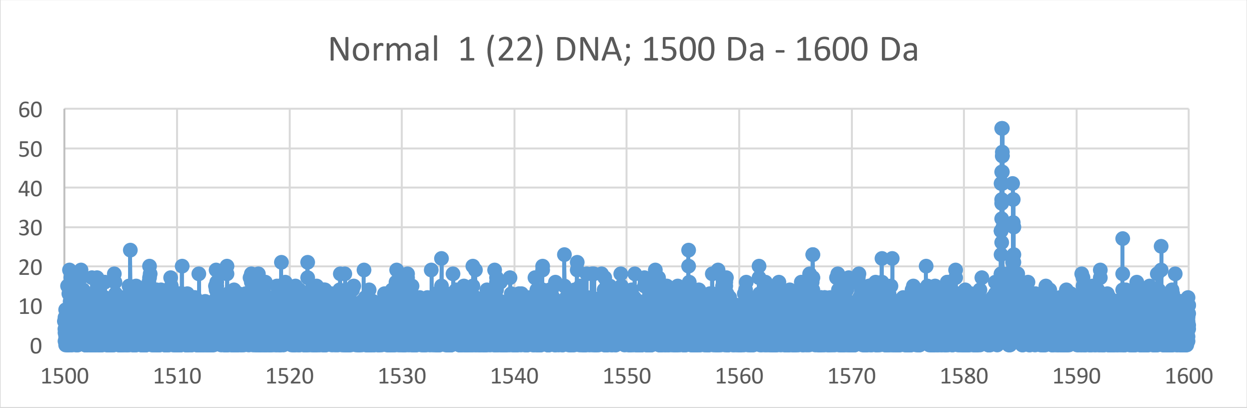
Normal 1 (22) DNA; 1500 Da - 1600 Da
60
50
40
30
20
10
0
1500
1510
1520
1530
1540
1550
1560
1570
1580
1590
1600

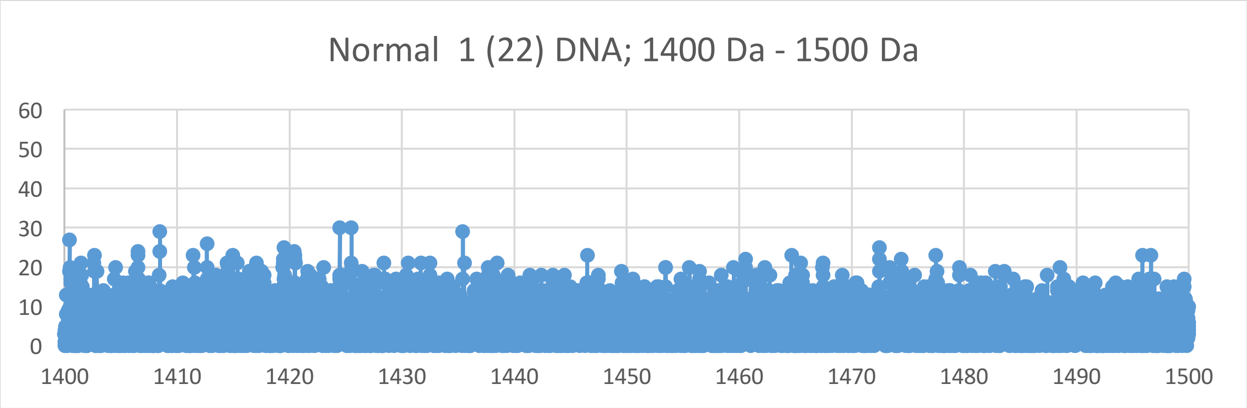
Normal 1 (22) DNA; 1400 Da - 1500 Da
60
50
40
30
20
10
0
1400
1410
1420
1430
1440
1450
1460
1470
1480
1490
1500

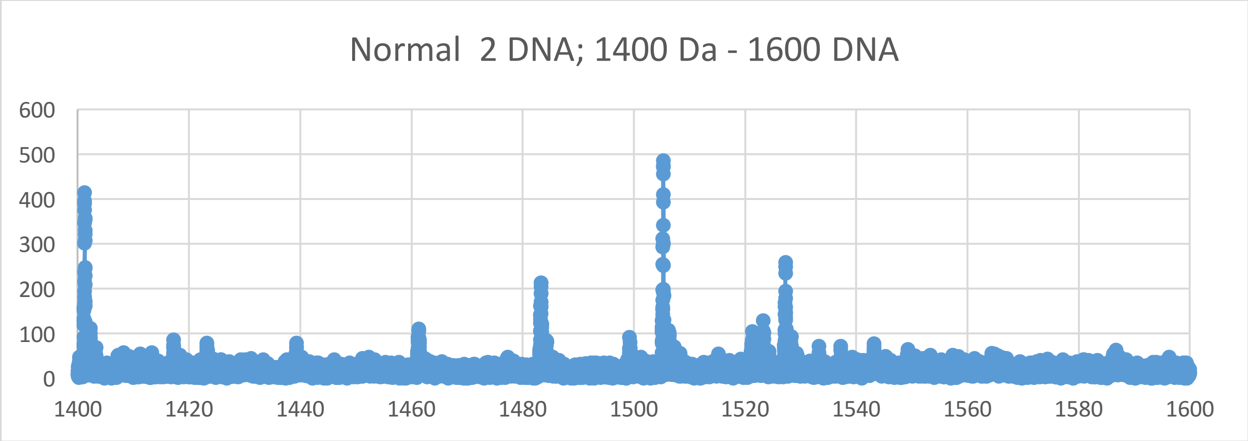
Normal 2 DNA; 1400 Da - 1600 DNA
600
500
400
300
200
100
0
1400
1420
1440
1460
1480
1500
1520
1540
1560
1580
1600

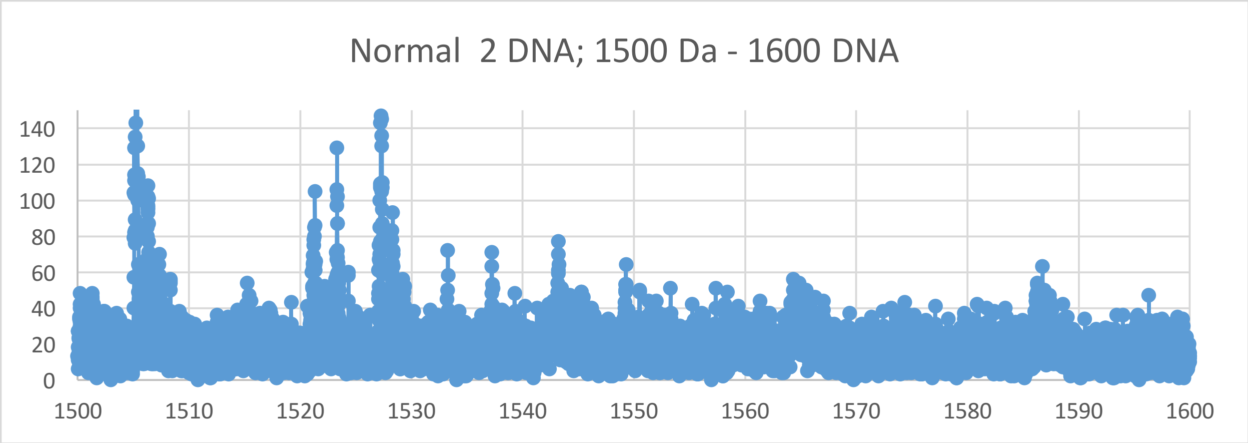
Normal 2 DNA; 1500 Da - 1600 DNA
140
120
100
80
60
40
20
0
1500
1510
1520
1530
1540
1550
1560
1570
1580
1590
1600

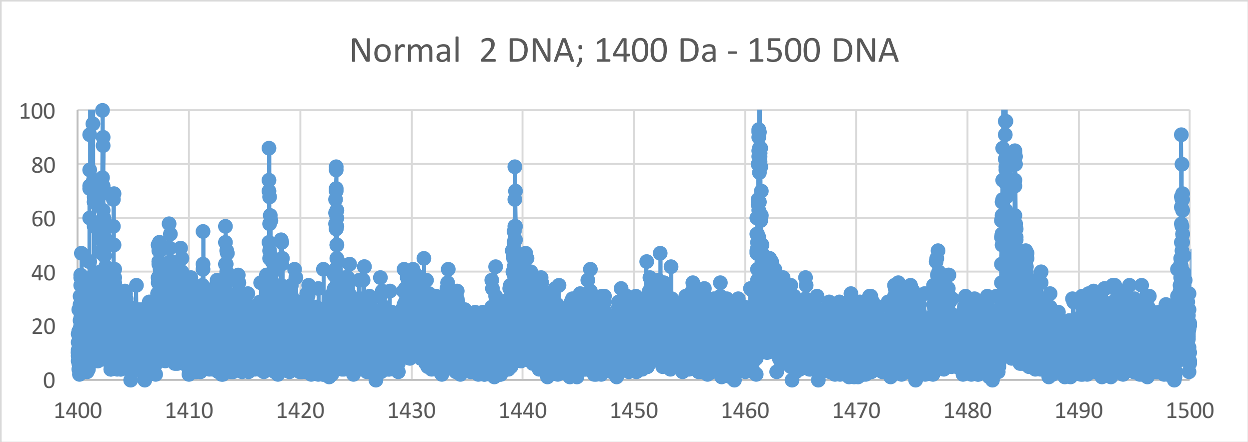
Normal 2 DNA; 1400 Da - 1500 DNA
100
80
60
40
20
0
1400
1410
1420
1430
1440
1450
1460
1470
1480
1490
1500

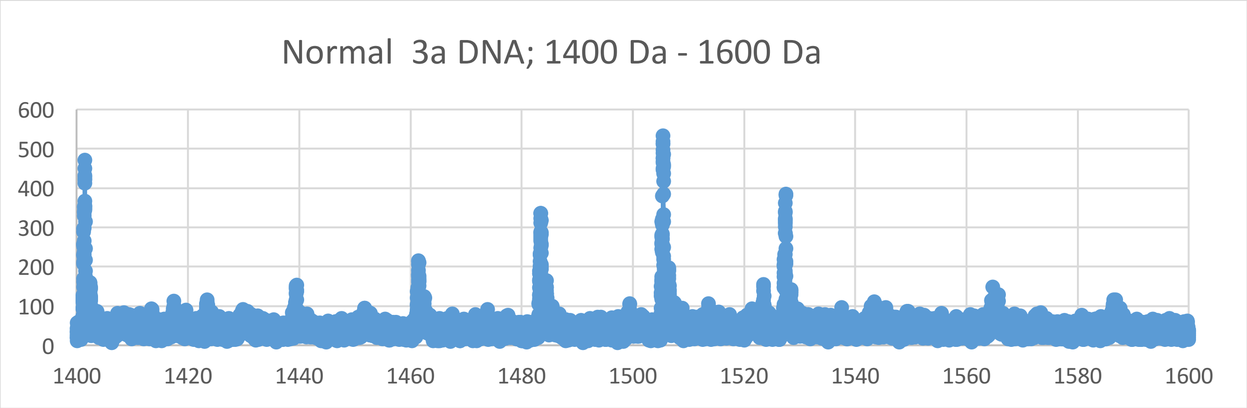
Normal 3a DNA; 1400 Da - 1600 Da
600
500
400
300
200
100
0
1400 1420 1440 1460 1480 1500 1520 1540 1560 1580 1600

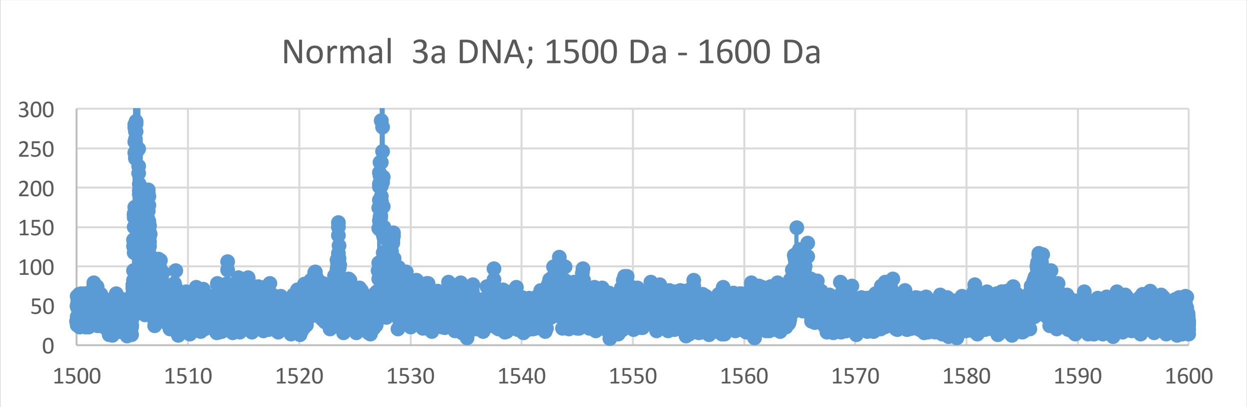
Normal 3a DNA; 1500 Da - 1600 Da
300
250
200
150
100
50
0
1500 1510 1520 1530 1540 1550 1560 1570 1580 1590 1600

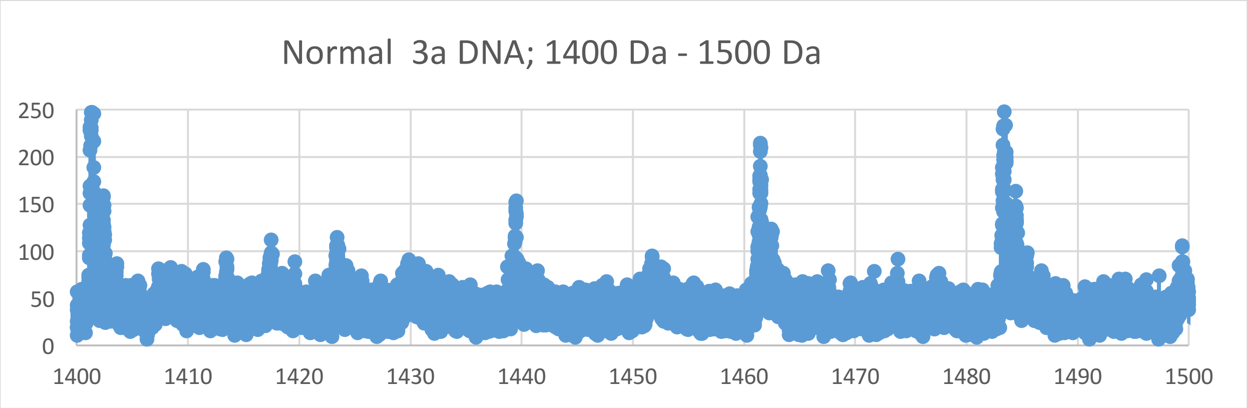
Normal 3a DNA; 1400 Da - 1500 Da
250
200
150
100
50
0
1400 1410 1420 1430 1440 1450 1460 1470 1480 1490 1500

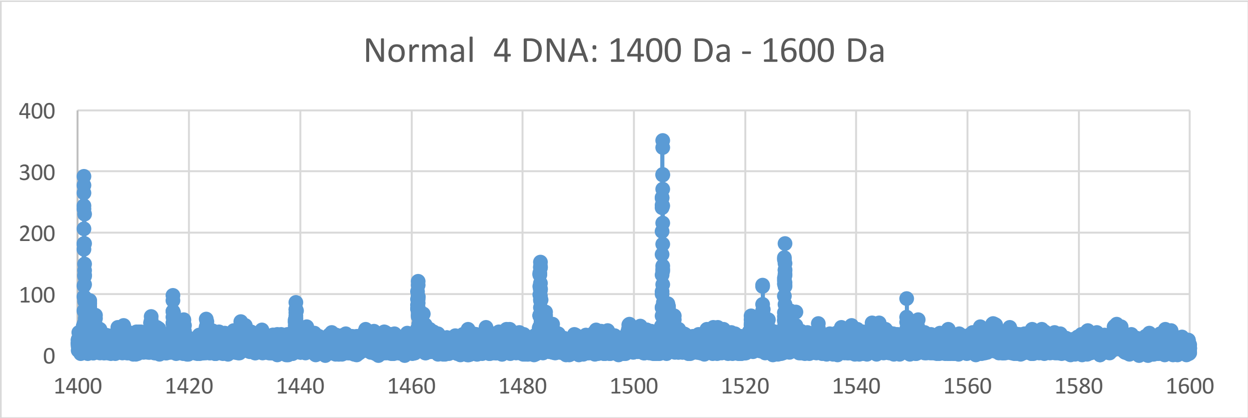

Normal 4 DNA: 1400 Da - 1600 Da
400
300
200
100
0
1400
1420
1440
1460
1480
1500
1520
1540
1560
1580
1600


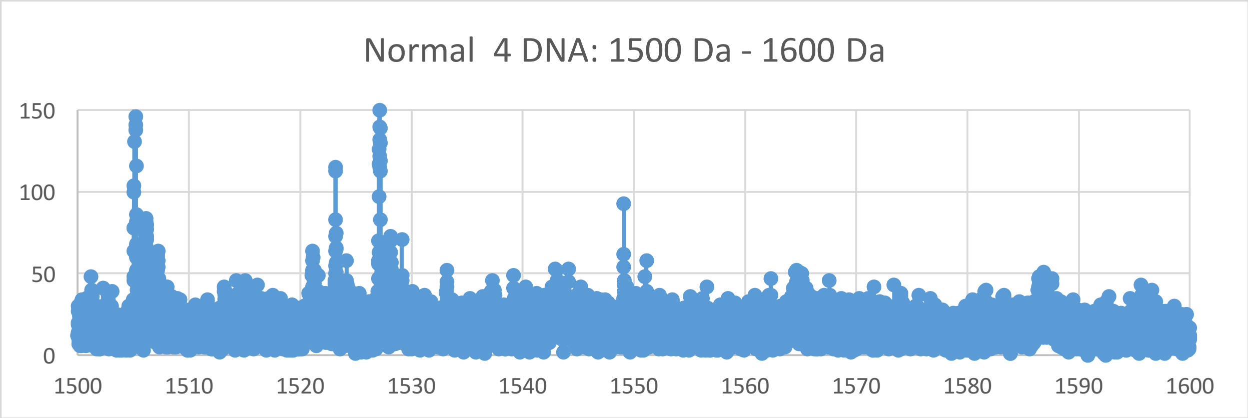

Normal 4 DNA: 1500 Da - 1600 Da
150
100
50
0
1500
1510
1520
1530
1540
1550
1560
1570
1580
1590
1600


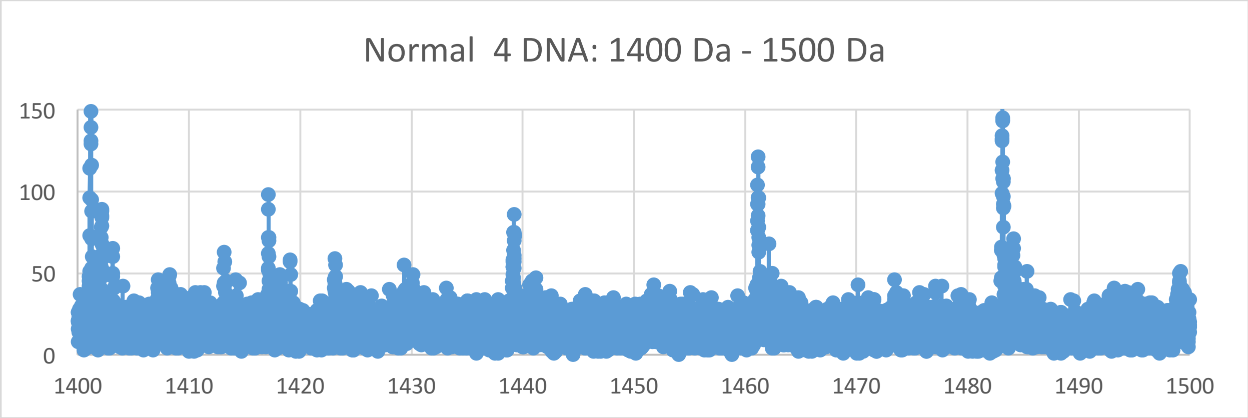

Normal 4 DNA: 1400 Da - 1500 Da
150
100
50
0
1400
1410
1420
1430
1440
1450
1460
1470
1480
1490
1500
