## Supplementary Materials VI for "Oncogenesis and Aging by Isotopic Functionalizations of the Proteins and Nucleic Acids"

VI Supplementary Materials – New Mass Spectra from 1300 Da -1400 Da

Results for Supplementary 6

So for the tetranucleotides from $(phos)_5$ to $(phos)_4$, the telomeric and nontelomeric peaks for cancer strongly increase in number and the peaks for normal strongly decreases in number for more stable cancer tetranucleotides and less stable normal tetranucleotides with these dephosphorylations. The masses range from 1350 Da to 1500 Da. See Supplementary Figures XI for graphs. These results and observations point specific differences between cancerous and normal oligonucleotides chemically and isotopically. It seems that below some number of phosphates, the telomeres, mating telomeres and nontelomers behave in same way toward cancer and normal. So this suggests normal may be favored and distinguished from cancer by high phosphorylations. The mating telomeric peaks for normal increase with these dephosphorylations of the tetranucleotides. These dephosphorylations from $(phos)_5$ to $(phos)_4$ caused the resulting more intense and stable cancer tetranucleotides to be more nontelomeric and telomeric and more isotopically enriched for telomeres and less for nontelomeres; but the mating telomeres are more intense in normal tetranucleotides and enriched isotopically. The mating telomeres are not as stabilized with cancer, mating telomeric tetranucleotides are isotopically enriched by these dephosphorylations. These dephosphorylations from $(phos)_5$ to $(phos)_4$ also caused the nontelomeric tetranucleotides to be more intense in cancer DNA and more isotopically enriched. It seems these dephosphorylations are related to the isotopic enrichments and depletions; but the telomeres are more intense in cancer and more isotopically enriched whereas the intensities of the mating telomeres are less intense in cancer and the isotopic enrichments are more. The telomeric tetranucleotides of cancer tended to also be more enriched so it seems the cancer stabilities of the telomeric tetranucleotides are related to their more isotopic enrichments for this dephosphorylations and purely driven by dephosphorylations and enrichments. Enrichments weaken the nontelomeric normal tetranucleotides while slightly weakening normal telomeres and strengthening the normal mating telomeric teranucleotides. The more intense cancer nontelomeric tetranucleotides were also slightly less isotopically enriched. These dephosphorylations from $(phos)_5$ to $(phos)_4$ increased the isotopic enrichments of the cancer for the nontelomeric and telomeric tetranucleotides. The mating telomeric tetranucleotides were more enriched from these dephosphorylations for more enrichments and stronger more stable normal tetranucleotides for mating telomeres. Such seems compatible for replications and transcriptions as stable telomeres seem compatible to unstable mating telomeres for the unravelings and constructions of DNA mating telomeres onto the building templates of the telomeres. These dephosphorylations from $(phos)_5$ to $(phos)_4$ caused methylations to cause slightly more normal intensities and stabilities but slightly less isotopic enrichments for telomeres and nontelomeres. But the methylations of the mating telomeres increased cancer stabilities and less isotopic enrichments for mating telomeres. This dephosphorylations from $(phos)_6$ to $(phos)_5$ caused acetylations to cause slightly more normal stabilities and slightly less cancer stabilities for telomeres and nontelomeres with more enrichments relative to methylations for telomeres and nontelomeres; the mating telomeres are slightly more cancer stable and less isotopically enriched by the acetylations. Thereby these dephosphorylations $(phos)_5$ to $(phos)_4$ and their varying consequences with respect to other dephosphorylations determine patterns of dephosphorylations and isotopic enrichment patterns in normal DNA which mutate and change for different patterns in DNA of cells that become cancerous. These dephosphorylations from $(phos)_5$ to $(phos)_4$ induced isotopic dynamics from of varying methylations and acetylations in cancer DNA relative to normal DNA couple to other isotopically induced varied phosphorylations, methylations and acetylations throughout the DNA for emerging order from nanodisorder.

In this mass range the tetranucleotides overlap with the trinucleotides so these masses are also in theory involving trinucleotides. So for the trinucleotides at $(phos)_8$, the telomeric peaks for cancer strongly increase in number and the peaks for normal decreases in number for more stable cancer trinucleotides and less stable normal trinucleotides with the $(phos)_8$. But the mating telomeres and nontelomeres have increase in numbers and stabilities for cancer trinucleotides while the normal trinucleotides are less in peaks and stabilities with $(phos)_8$. But as decrease the phosphate by 1 unit, the trinucleotides change so that the normal trinucleotides more fragment and break from the DNA with more numbers and stabilities for the nontelomeres, with slightly more fragmenting and stabilities of the normal from DNA for telomeres and mating telomeres from $(phos)_8$ to $(phos)_7$ . The masses range from 1324 Da to 1468 Da. See Supplementary Figures VI for graphs. It seems that below some number of phosphates, the telomeres, mating telomeres and nontelomers behave in same way toward cancer and normal. So this suggest normal may be favored and distinguished from cancer by changing saturated phosphorylations. The telomeric peaks for normal increase with this dephosphorylations of the trinucleotides from $(phos)_8$ to $(phos)_7$. The dephosphorylations from $(phos)_8$ to $(phos)_7$ caused the resulting more intense and stable normal trinucleotides to be more mating telomeric, telomeric and nontelomeric trinucleotides and less isotopic enrichments for telomeres, mating telomeres and nontelomeres. It is important to consider that there are more intense peaks for the trinucleotides relative to the tetranucleotides for the saturated phosphate units. For the trinucleotides, the intensities of normal increased and the intensities of cancer fragments decreased from $(phos)_8$ to $(phos)_7$. The dephosphorylations from $(phos)_8$ to $(phos)_7$ decreases isotopic enrichments of heavy isotopes of $^{13}C$ and $^{17}O$. The dephosphorylations from $(phos)_8$ to $(phos)_7$ also caused the telomeric, mating telomeric and nontelomeric trinucleotides to be more intense in normal DNA and less isotopically enriched. It seems the dephosphorylations are related to the isotopic enrichments and depletions; but the telomeres and nontelomeres are more intense in normal and less isotopically enriched whereas the intensities of the mating telomeres are more intense in normal and the less isotopic enriched. The telomeric, mating telomeric and nontelomeric trinucleotides of normal tended to also be less enriched so it seems the cancer stabilities of the telomeric, mating telomeric and nontelomeric trinucleotides are related to their less isotopic enrichments for the dephosphorylations from $(phos)_8$ to $(phos)_7$ and purely driven by dephosphorylations and enrichments. Enrichments strengthen the nontelomeric cancer trinucleotides while strongly weakening normal telomeric, mating telomeric and nontelomeric trinucleotides. The more intense normal nontelomeric, telomeric and nontelomeric trinucleotides were also less isotopically enriched. The dephosphorylations from $(phos)_8$ to $(phos)_7$ decreased the isotopic enrichments of the normal for the nontelomeric, mating telomeric and telomeric trinucleotides.

But the nontelomeric trinucleotides were less enriched from the dephosphorylations for less enrichments and stronger more stable normal trinucleotides for nontelomeres. Such seems compatible for replications and transcriptions as stable telomeres seem compatible to unstable mating telomeres for the unravelings and constructions of DNA mating telomeres onto the building templates of the telomeres. The dephosphorylations from $(phos)_8$ to $(phos)_7$ caused methylations to cause more cancer intensities and stabilities but more isotopic enrichments for telomeres and nontelomeres. But the methylations of the mating telomeres increased cancer stabilities and slightly more isotopic enrichments for mating telomeres. The methylations of the nontelomeres increased cancer intensities and stabilities with more enrichments. It seems from this data that during biological processes the dephosphorylations and stabilizations of normal or vice versa dephosphorylations and stabilizations of cancer are or can be countered by methylations and destabilizations of the normal for cancer or vice versa dephosphorylations and destabilizations of the cancer by methylations as this research discovers coupled phosphorylations and methylations or coupled dephosphorylations and methylations with involved greater enrichments for normal biological functions and explaining cancer! The dephosphorylations from $(phos)_8$ to $(phos)_7$ caused acetylations to cause more cancer stabilities for telomeres and nontelomeres and more cancer stabilities for telomeres and nontelomeres with more enrichments for telomeres and more enrichments for nontelomeres. The acetylations from $(phos)_8$ to $(phos)_7$ caused more normal stabilities and intensities for mating telomeres with more enrichments. This contrast in mating telomeres and telomere from the acetylations may reflect the developments of cancer as the telomeres fragment but the mating telomeres more holds together for damaged DNA for cancer.

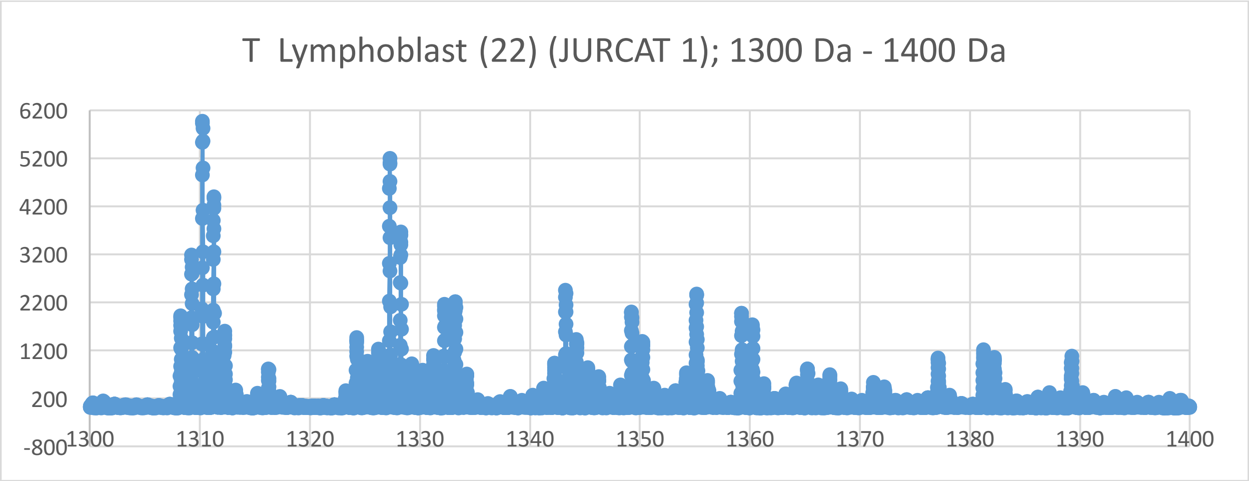
T Lymphoblast (22) (JURCAT 1); 1300 Da - 1400 Da
6200
5200
4200
3200
2200
1200
200
-800
1300
1310
1320
1330
1340
1350
1360
1370
1380
1390
1400

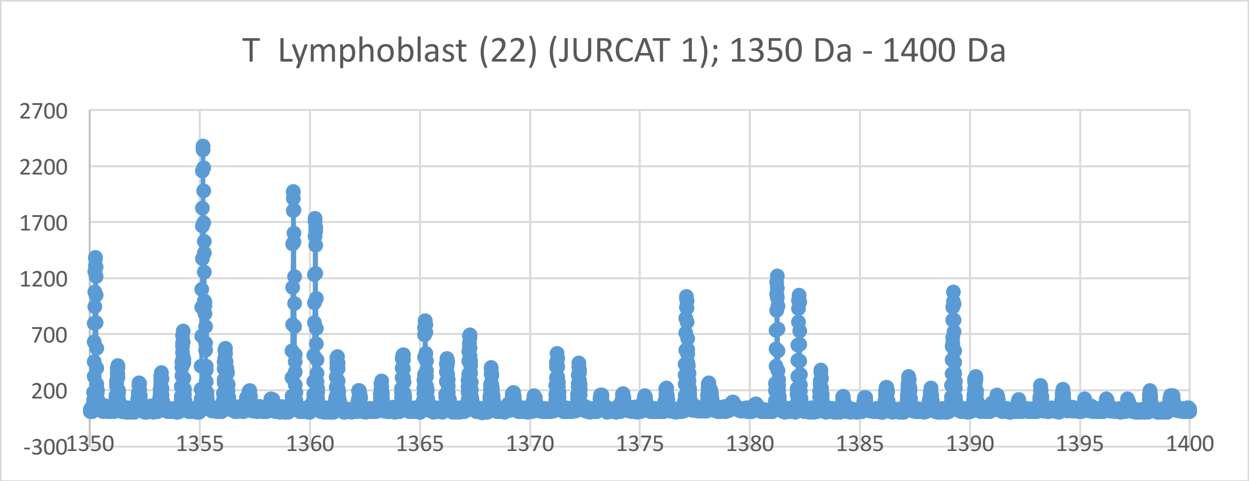
T Lymphoblast (22) (JURCAT 1); 1350 Da - 1400 Da
2700
2200
1700
1200
700
200
-300
1350
1355
1360
1365
1370
1375
1380
1385
1390
1395
1400

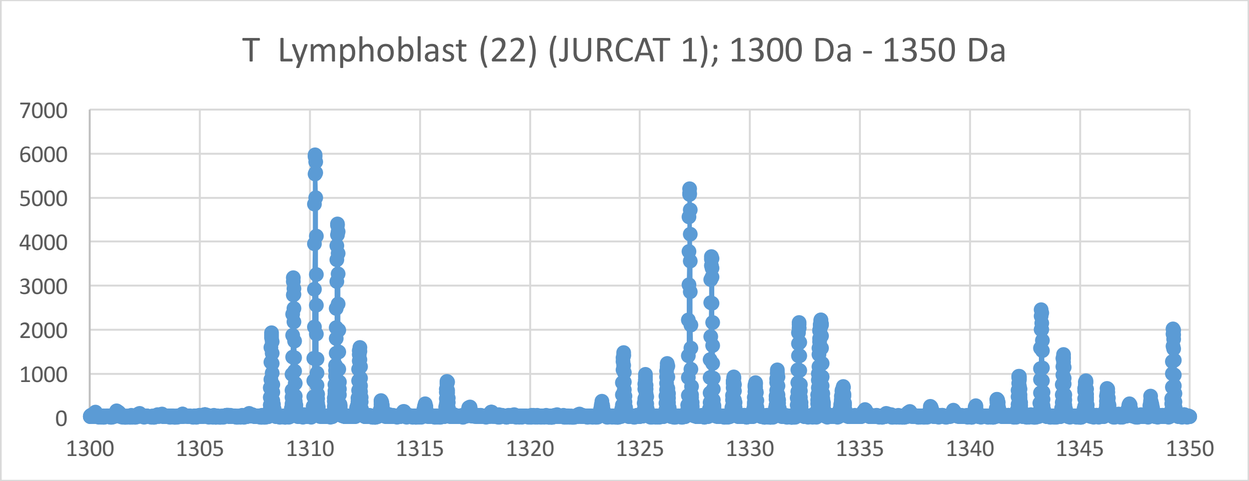
T Lymphoblast (22) (JURCAT 1); 1300 Da - 1350 Da
7000
6000
5000
4000
3000
2000
1000
0
1300
1305
1310
1315
1320
1325
1330
1335
1340
1345
1350

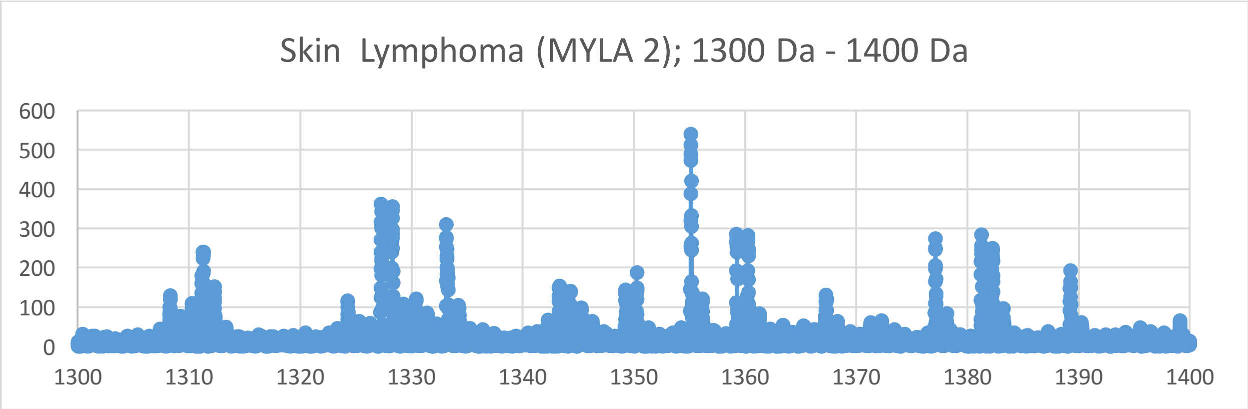
Skin Lymphoma (MYLA 2); 1300 Da - 1400 Da
600
500
400
300
200
100
0
1300
1310
1320
1330
1340
1350
1360
1370
1380
1390
1400

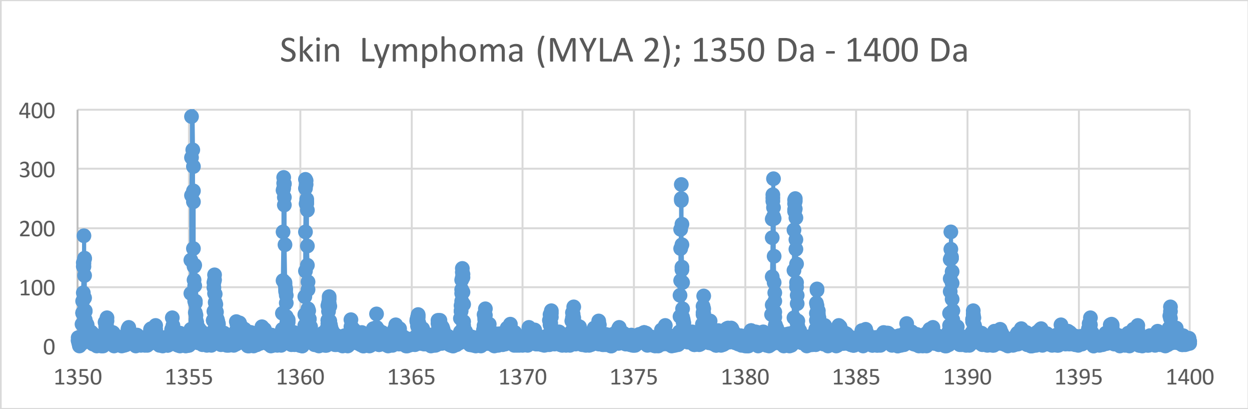
Skin Lymphoma (MYLA 2); 1350 Da - 1400 Da
400
300
200
100
0
1350
1355
1360
1365
1370
1375
1380
1385
1390
1395
1400

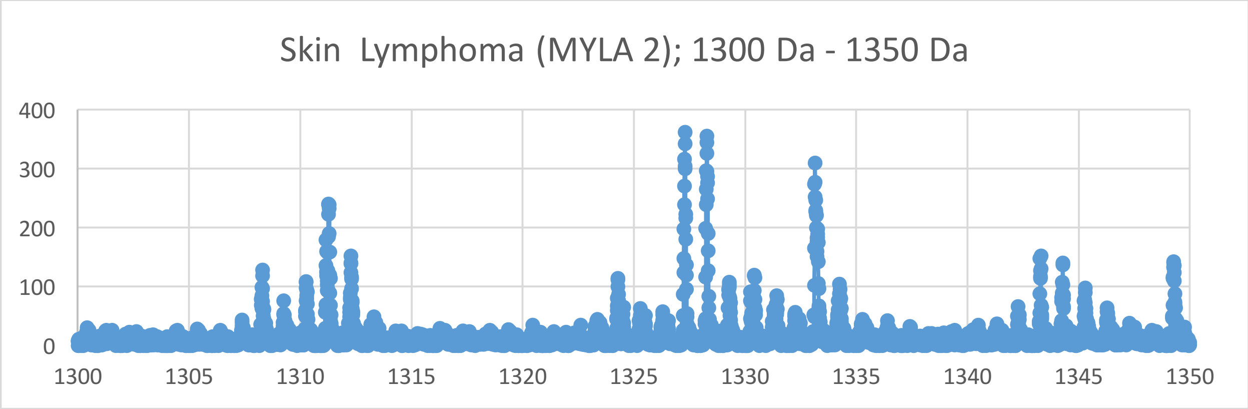
Skin Lymphoma (MYLA 2); 1300 Da - 1350 Da
400
300
200
100
0
1300
1305
1310
1315
1320
1325
1330
1335
1340
1345
1350

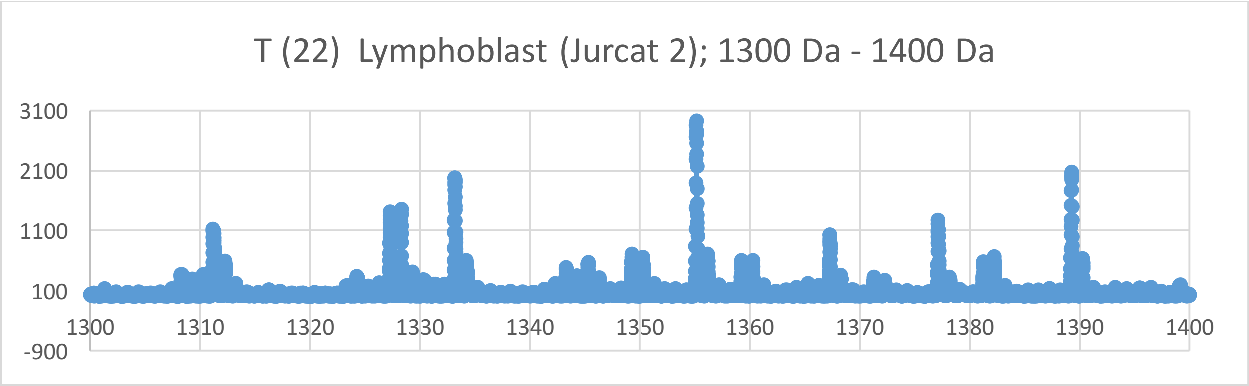
T (22) Lymphoblast (Jurcat 2); 1300 Da - 1400 Da
3100
2100
1100
100
-900
1300
1310
1320
1330
1340
1350
1360
1370
1380
1390
1400

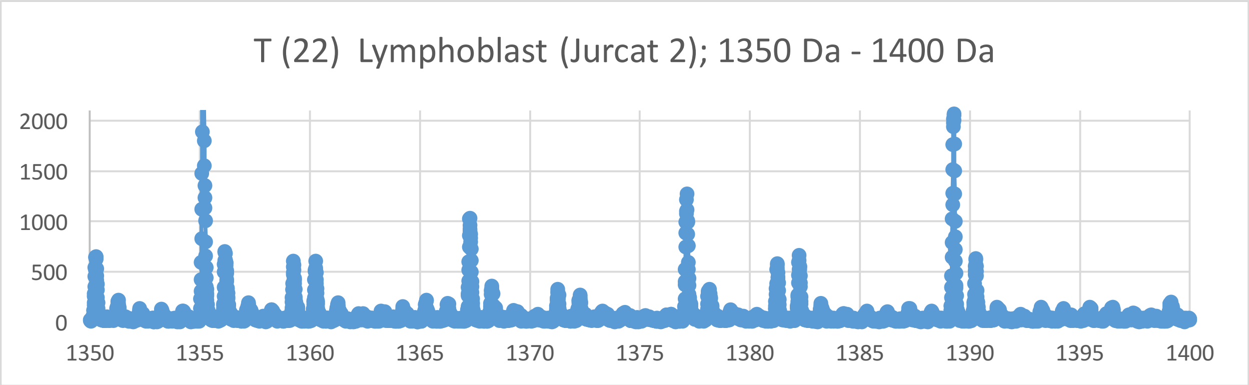
T (22) Lymphoblast (Jurcat 2); 1350 Da - 1400 Da
2000
1500
1000
500
0
1350
1355
1360
1365
1370
1375
1380
1385
1390
1395
1400

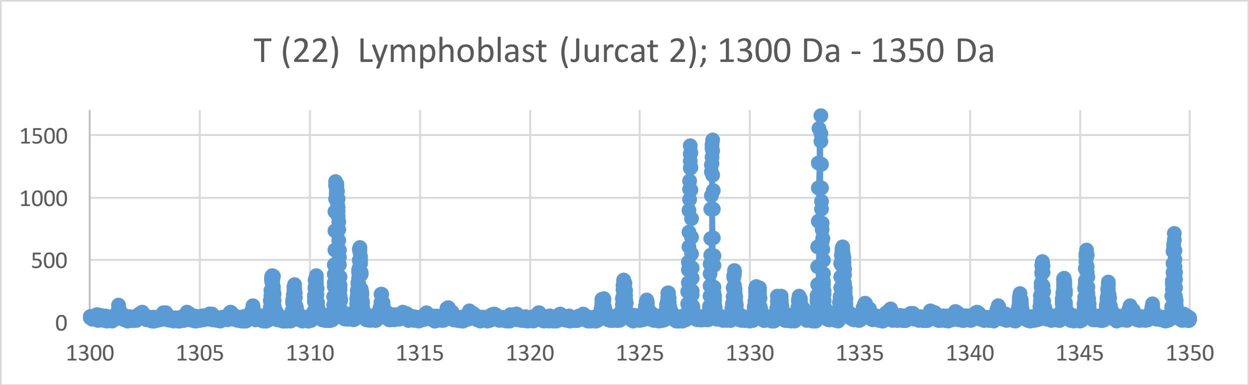
T (22) Lymphoblast (Jurcat 2); 1300 Da - 1350 Da
1500
1000
500
0
1300
1305
1310
1315
1320
1325
1330
1335
1340
1345
1350

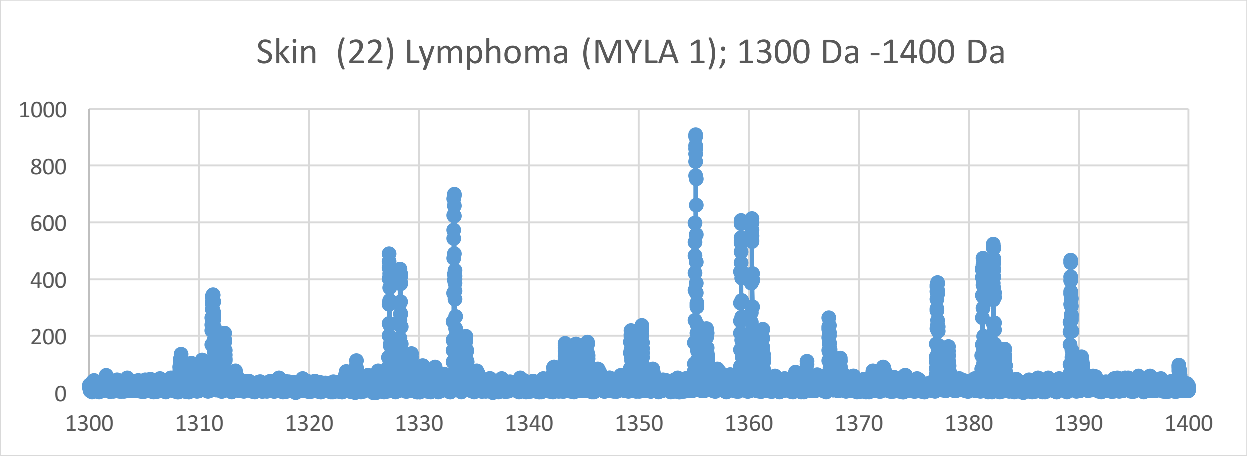
Skin (22) Lymphoma (MYLA 1); 1300 Da -1400 Da
1000
800
600
400
200
0
1300
1310
1320
1330
1340
1350
1360
1370
1380
1390
1400

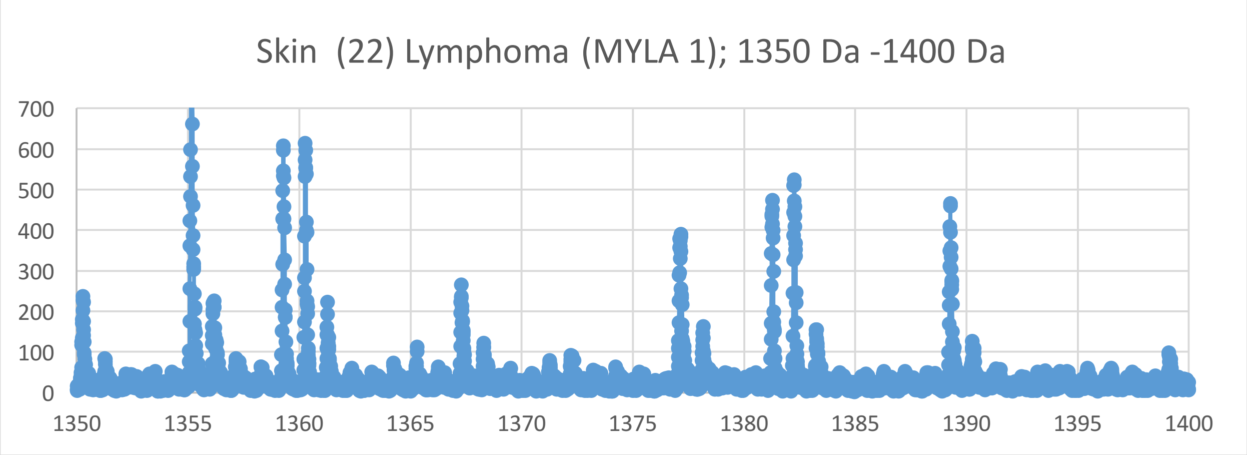
Skin (22) Lymphoma (MYLA 1); 1350 Da -1400 Da
700
600
500
400
300
200
100
0
1350
1355
1360
1365
1370
1375
1380
1385
1390
1395
1400

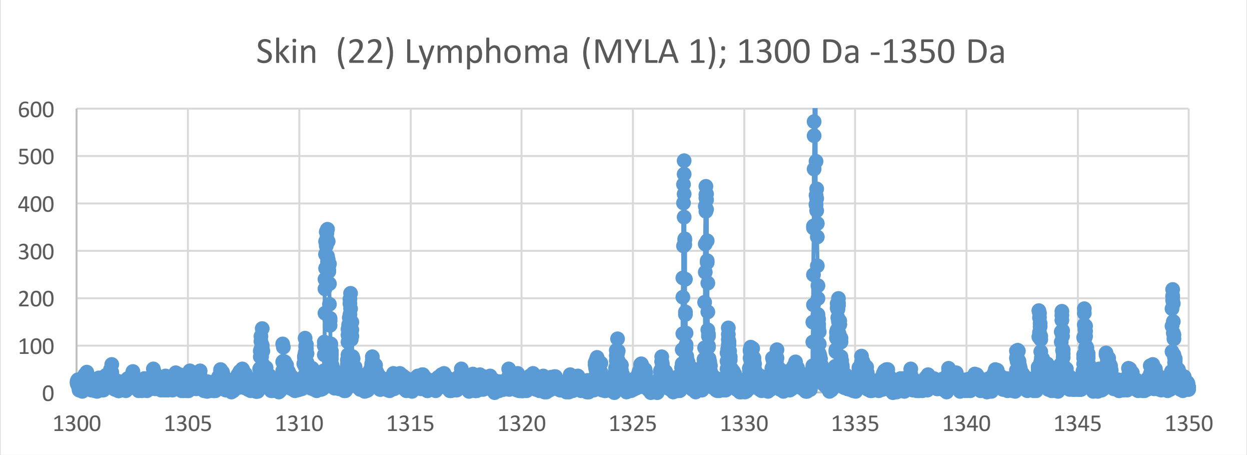
Skin (22) Lymphoma (MYLA 1); 1300 Da -1350 Da
600
500
400
300
200
100
0
1300
1305
1310
1315
1320
1325
1330
1335
1340
1345
1350

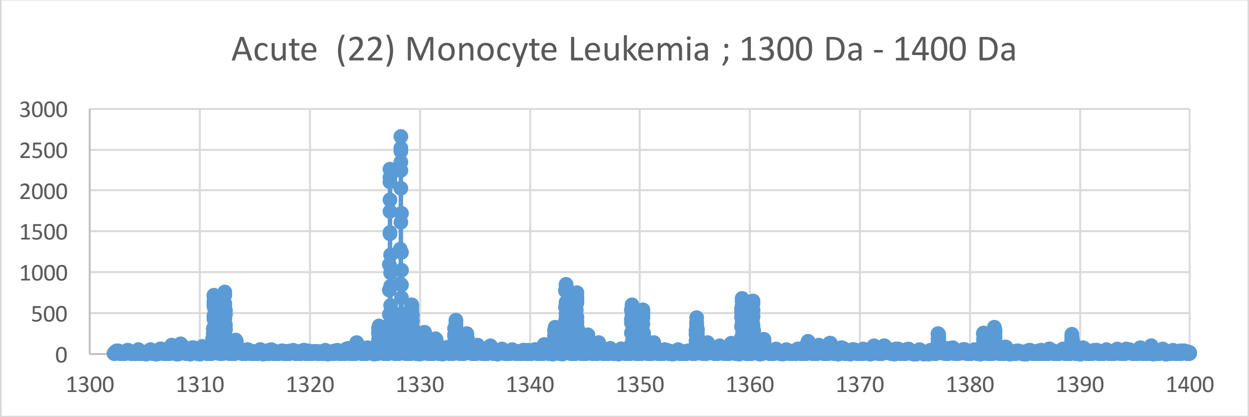
Acute (22) Monocyte Leukemia ; 1300 Da - 1400 Da
3000
2500
2000
1500
1000
500
0
1300
1310
1320
1330
1340
1350
1360
1370
1380
1390
1400

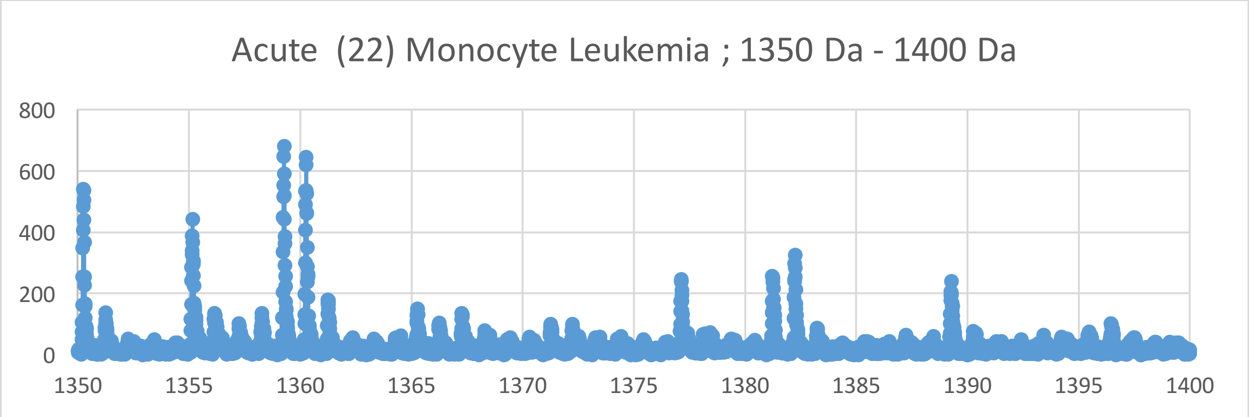
Acute (22) Monocyte Leukemia ; 1350 Da - 1400 Da
800
600
400
200
0
1350
1355
1360
1365
1370
1375
1380
1385
1390
1395
1400

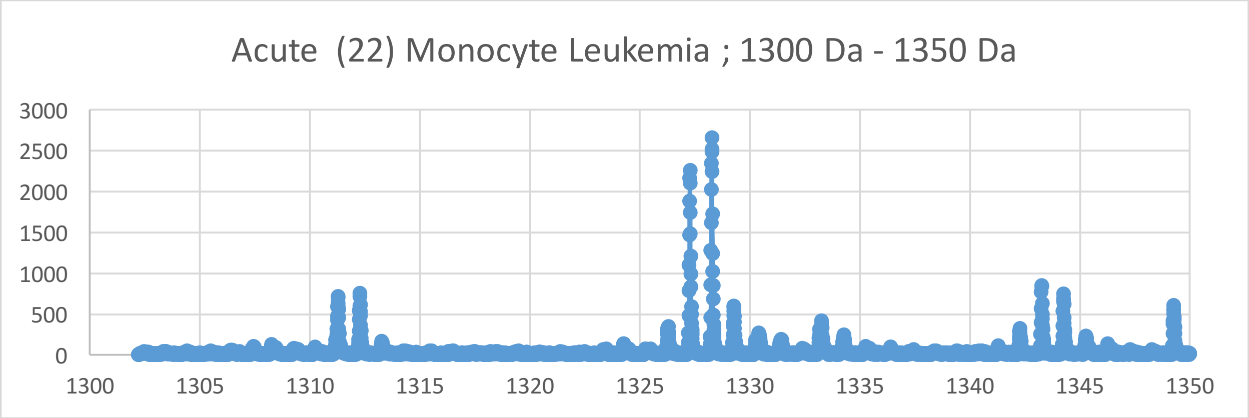
Acute (22) Monocyte Leukemia ; 1300 Da - 1350 Da
3000
2500
2000
1500
1000
500
0
1300
1305
1310
1315
1320
1325
1330
1335
1340
1345
1350

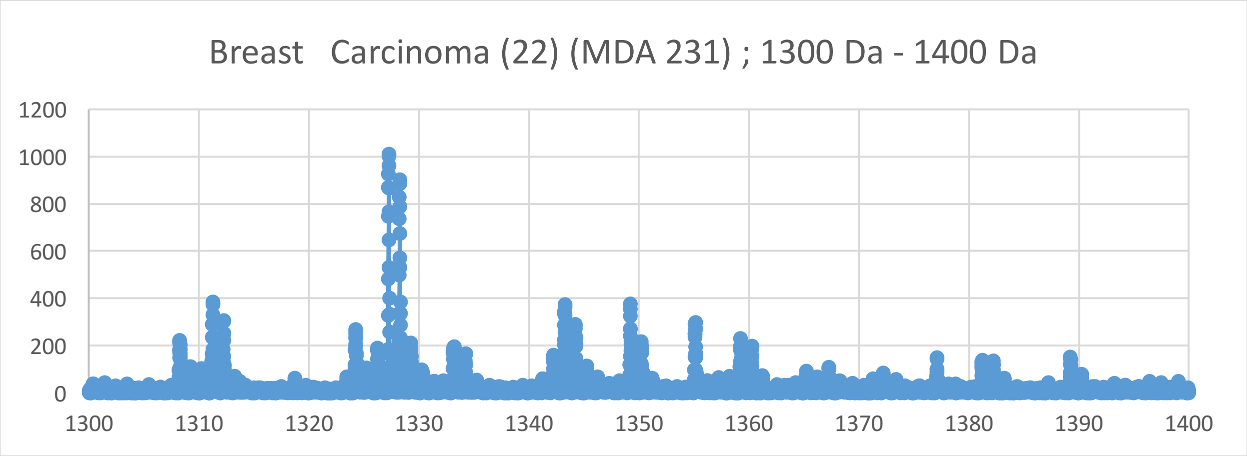
Breast Carcinoma (22) (MDA 231) ; 1300 Da - 1400 Da
1200
1000
800
600
400
200
0
1300
1310
1320
1330
1340
1350
1360
1370
1380
1390
1400

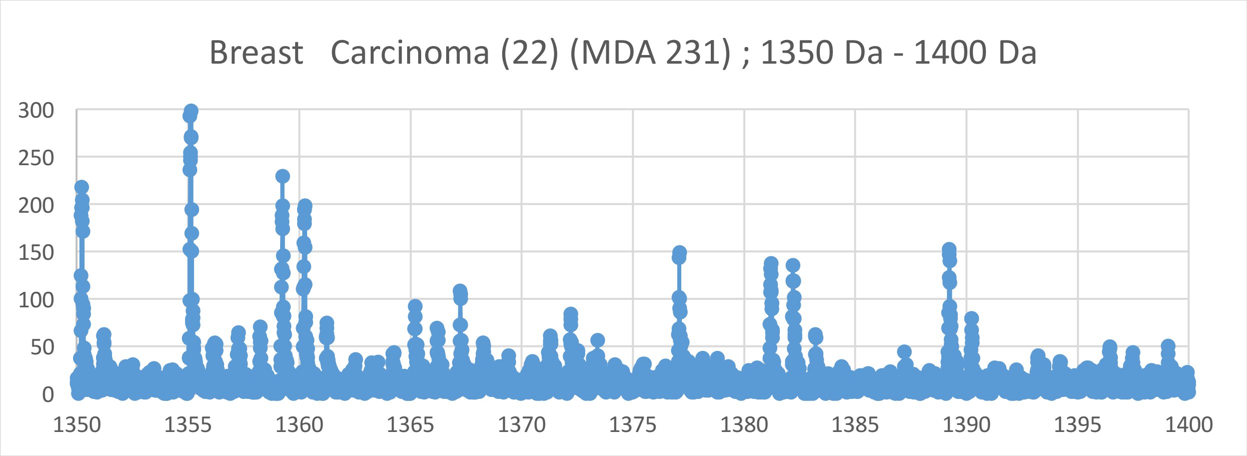
Breast Carcinoma (22) (MDA 231) ; 1350 Da - 1400 Da
300
250
200
150
100
50
0
1350
1355
1360
1365
1370
1375
1380
1385
1390
1395
1400

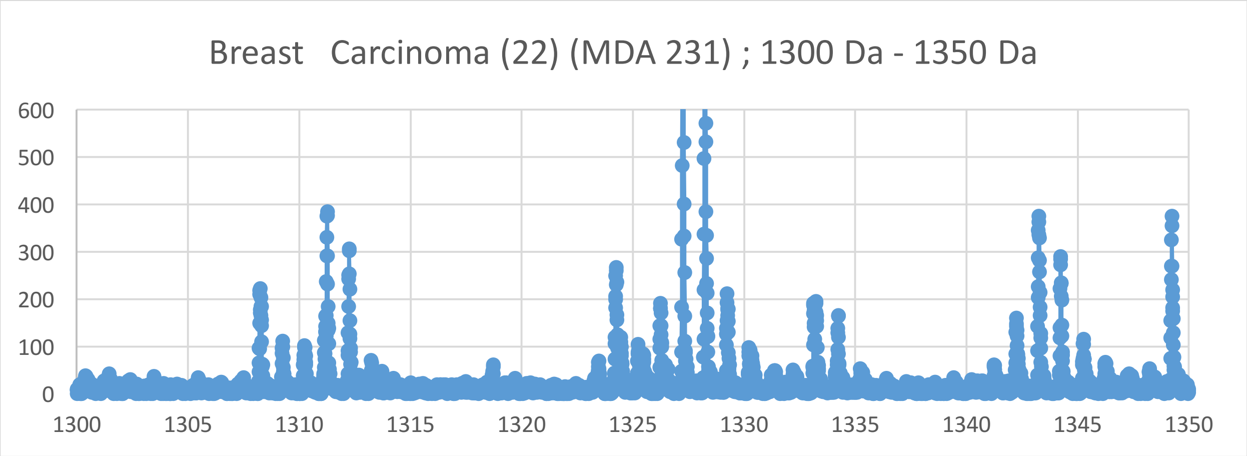
Breast Carcinoma (22) (MDA 231) ; 1300 Da - 1350 Da
600
500
400
300
200
100
0
1300
1305
1310
1315
1320
1325
1330
1335
1340
1345
1350

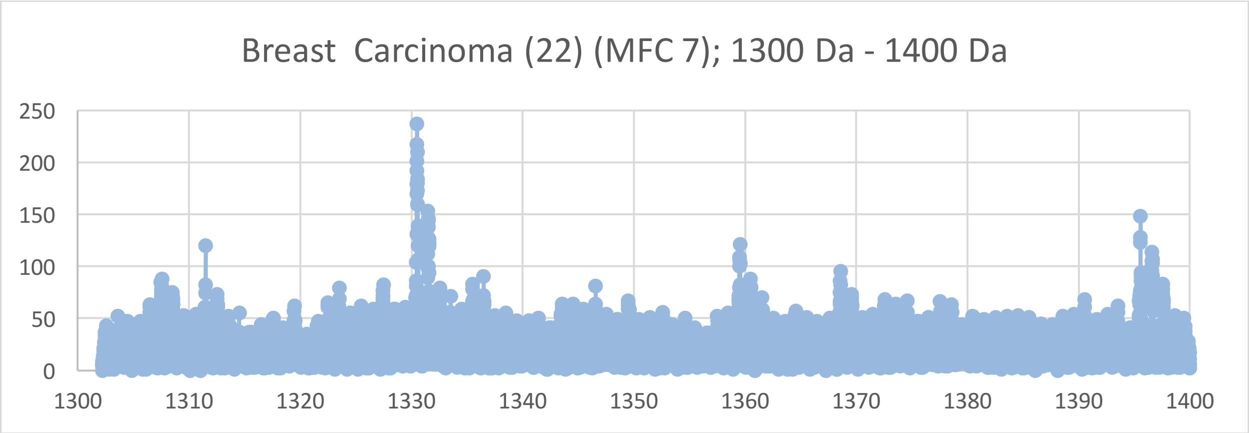

Breast Carcinoma (22) (MFC 7); 1300 Da - 1400 Da
250
200
150
100
50
0
1300
1310
1320
1330
1340
1350
1360
1370
1380
1390
1400


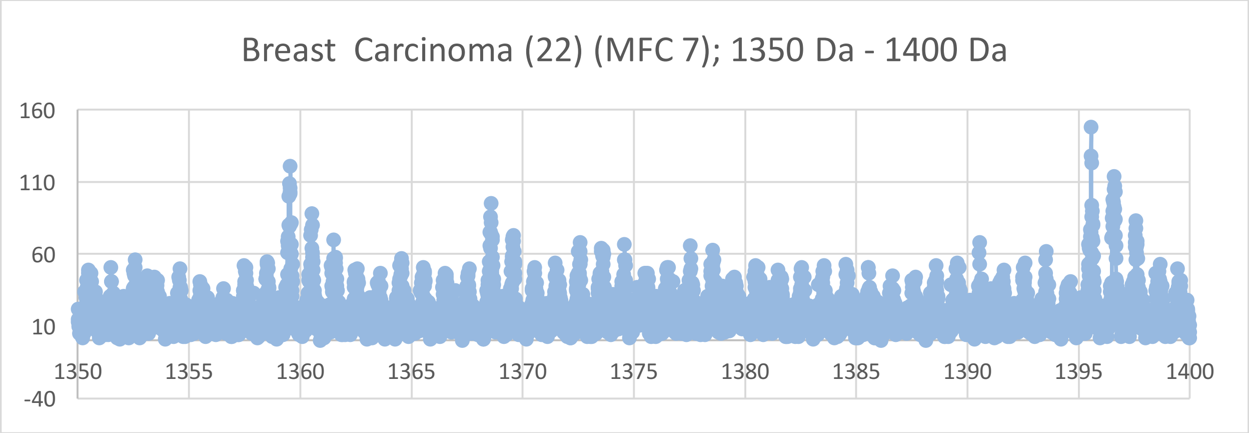

Breast Carcinoma (22) (MFC 7); 1350 Da - 1400 Da
160
110
60
10
-40
1350
1355
1360
1365
1370
1375
1380
1385
1390
1395
1400


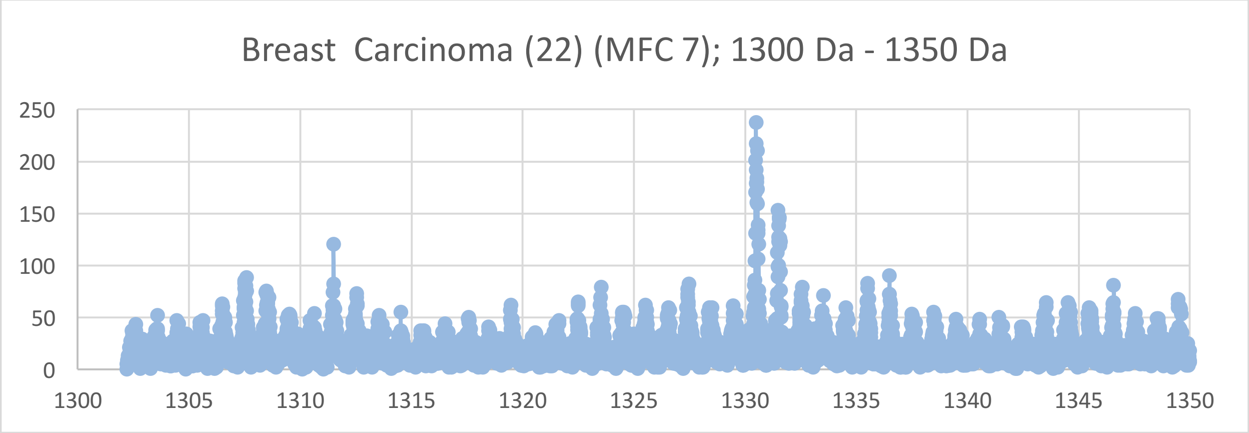

Breast Carcinoma (22) (MFC 7); 1300 Da - 1350 Da
250
200
150
100
50
0
1300
1305
1310
1315
1320
1325
1330
1335
1340
1345
1350

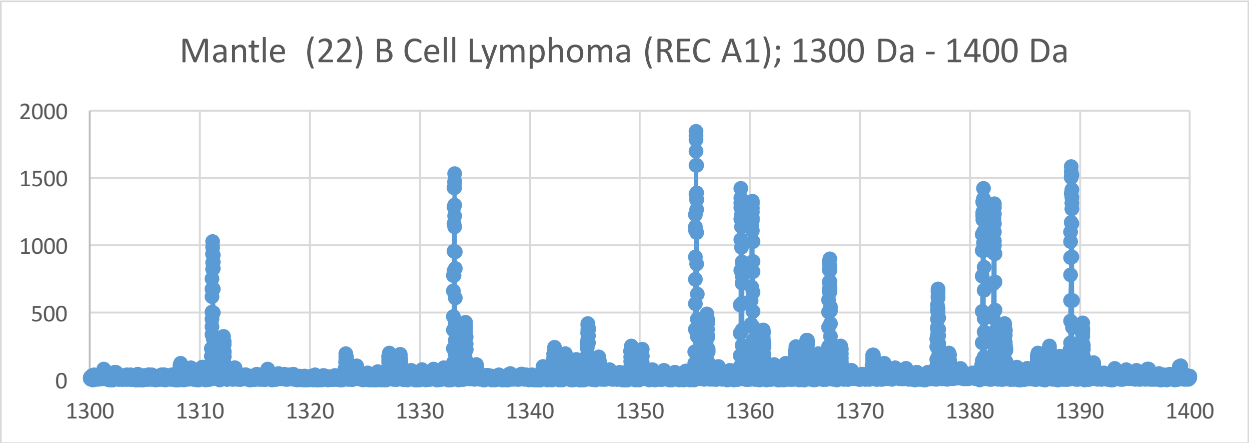
Mantle (22) B Cell Lymphoma (REC A1); 1300 Da - 1400 Da
2000
1500
1000
500
0
1300
1310
1320
1330
1340
1350
1360
1370
1380
1390
1400

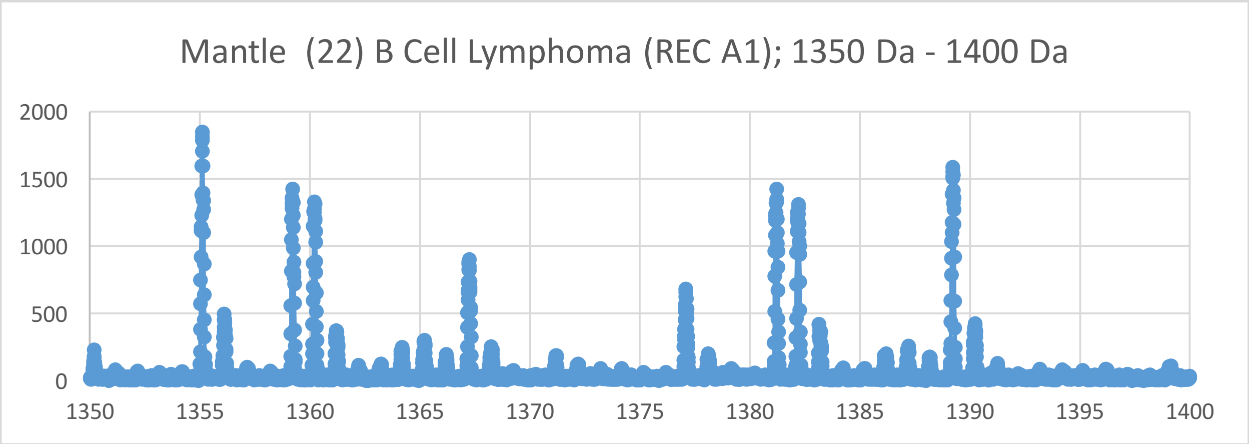
Mantle (22) B Cell Lymphoma (REC A1); 1350 Da - 1400 Da
2000
1500
1000
500
0
1350
1355
1360
1365
1370
1375
1380
1385
1390
1395
1400

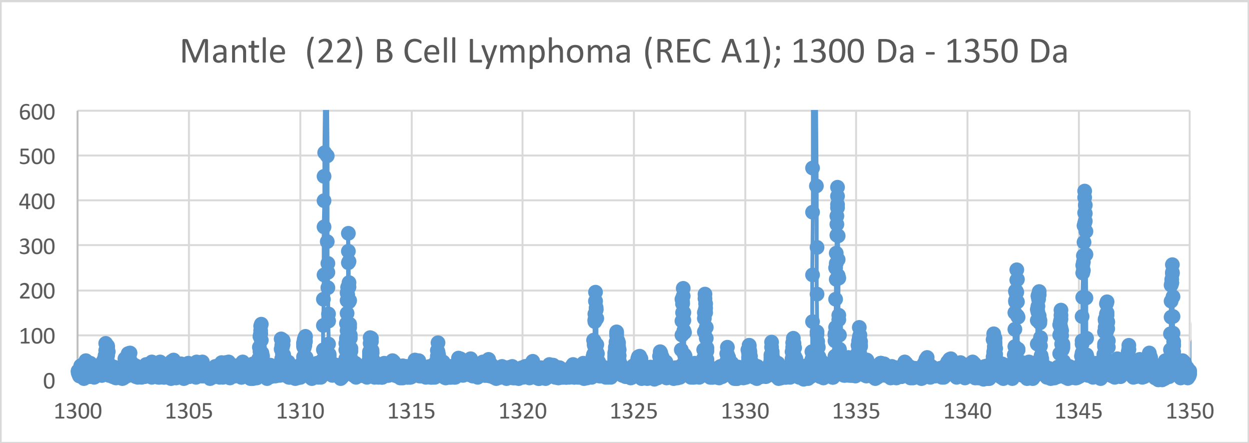
Mantle (22) B Cell Lymphoma (REC A1); 1300 Da - 1350 Da
600
500
400
300
200
100
0
1300
1305
1310
1315
1320
1325
1330
1335
1340
1345
1350

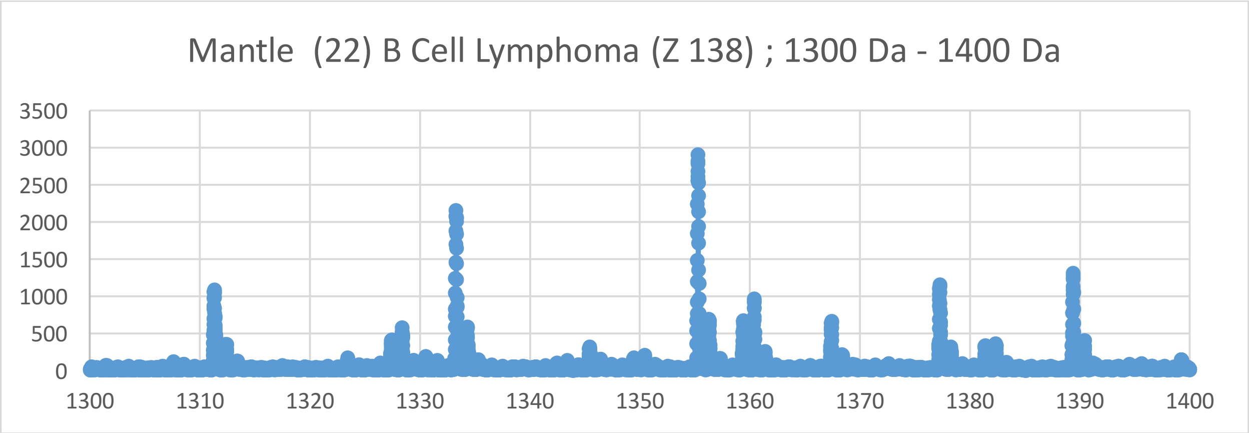
Mantle (22) B Cell Lymphoma (Z 138) ; 1300 Da - 1400 Da
3500
3000
2500
2000
1500
1000
500
0
1300
1310
1320
1330
1340
1350
1360
1370
1380
1390
1400

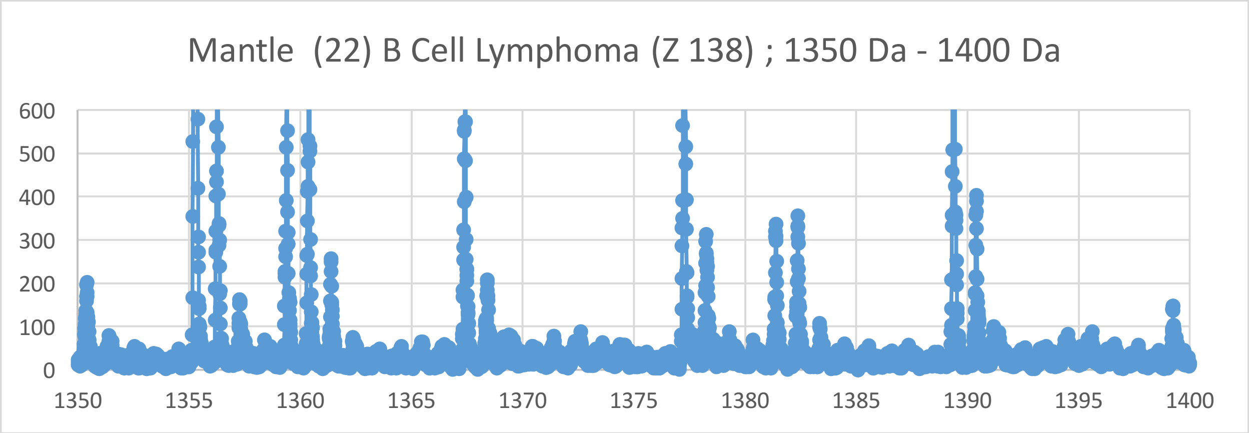
Mantle (22) B Cell Lymphoma (Z 138) ; 1350 Da - 1400 Da
600
500
400
300
200
100
0
1350
1355
1360
1365
1370
1375
1380
1385
1390
1395
1400

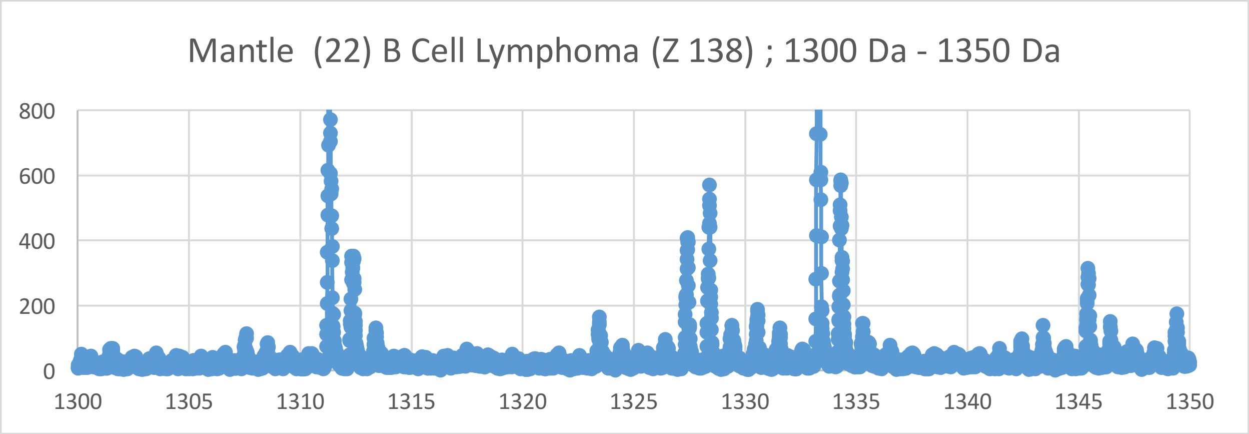
Mantle (22) B Cell Lymphoma (Z 138) ; 1300 Da - 1350 Da
800
600
400
200
0
1300
1305
1310
1315
1320
1325
1330
1335
1340
1345
1350

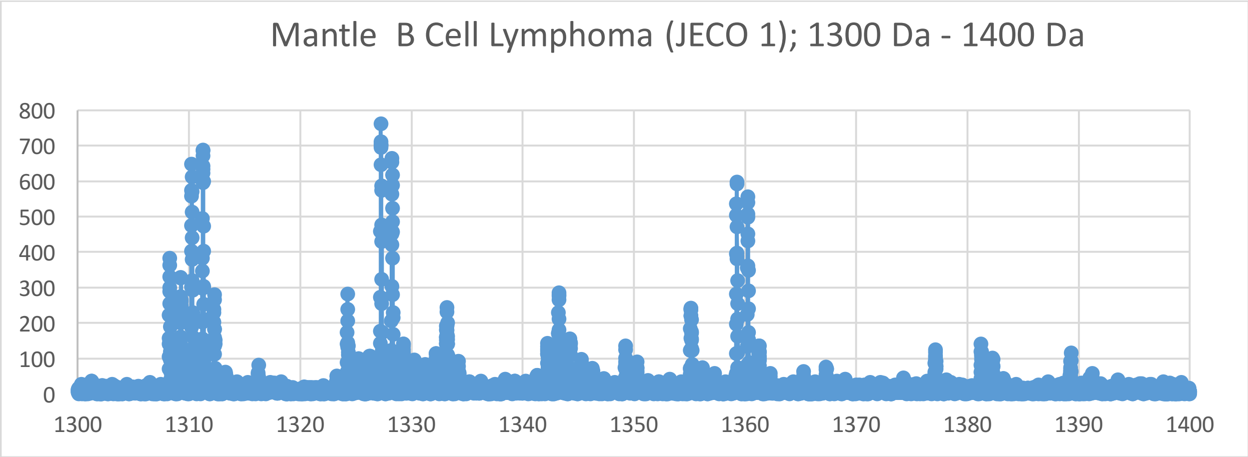
Mantle B Cell Lymphoma (JECO 1); 1300 Da - 1400 Da
800
700
600
500
400
300
200
100
0
1300
1310
1320
1330
1340
1350
1360
1370
1380
1390
1400

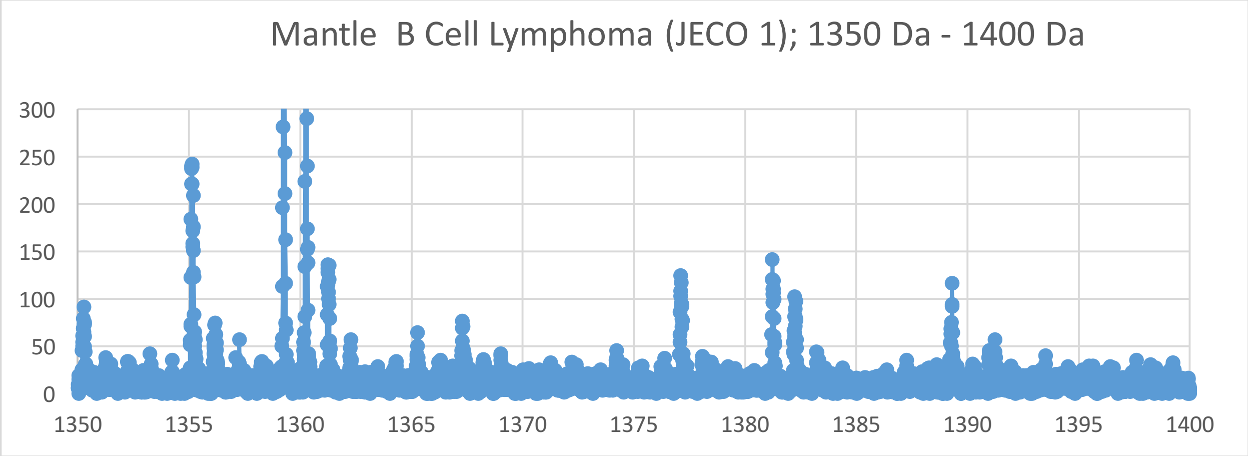
Mantle B Cell Lymphoma (JECO 1); 1350 Da - 1400 Da
300
250
200
150
100
50
0
1350
1355
1360
1365
1370
1375
1380
1385
1390
1395
1400

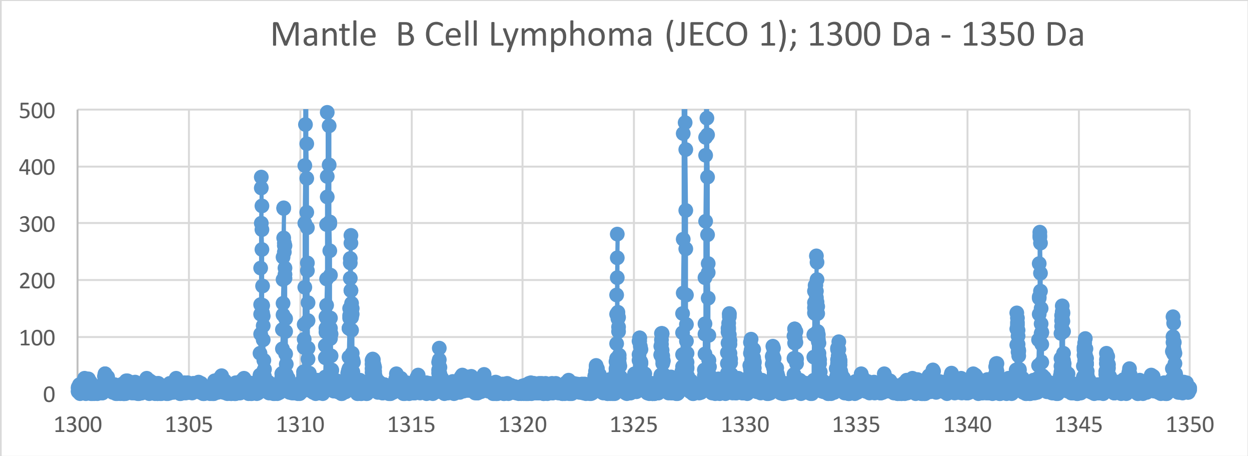
Mantle B Cell Lymphoma (JECO 1); 1300 Da - 1350 Da
500
400
300
200
100
0
1300
1305
1310
1315
1320
1325
1330
1335
1340
1345
1350

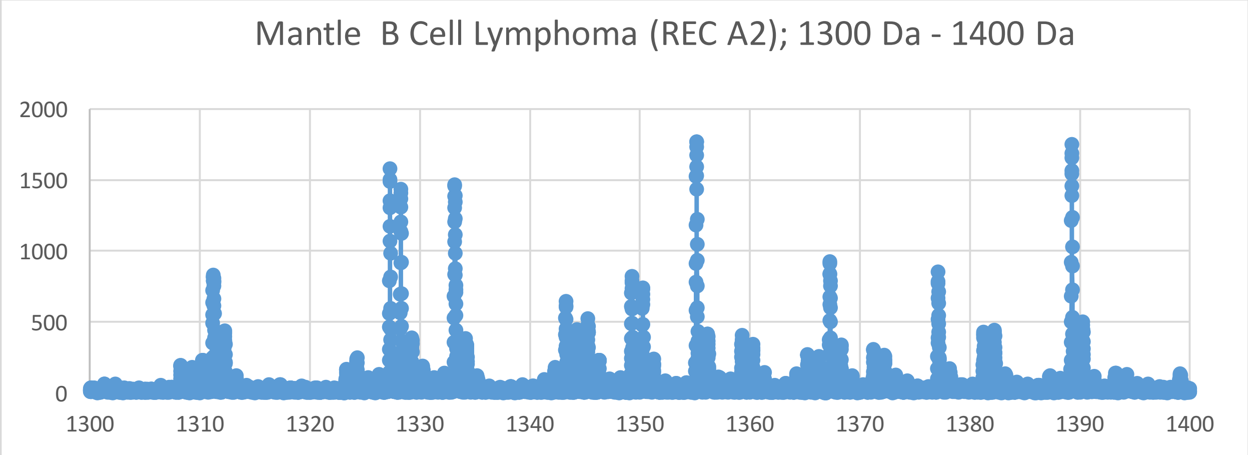
Mantle B Cell Lymphoma (REC A2); 1300 Da - 1400 Da
2000
1500
1000
500
0
1300
1310
1320
1330
1340
1350
1360
1370
1380
1390
1400

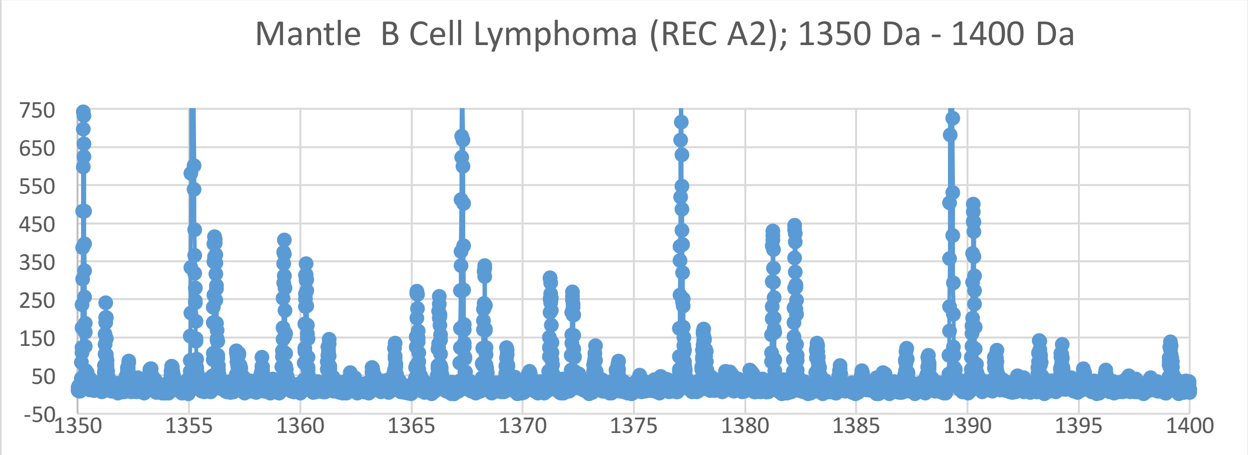
Mantle B Cell Lymphoma (REC A2); 1350 Da - 1400 Da
750
650
550
450
350
250
150
50
-50
1350
1355
1360
1365
1370
1375
1380
1385
1390
1395
1400

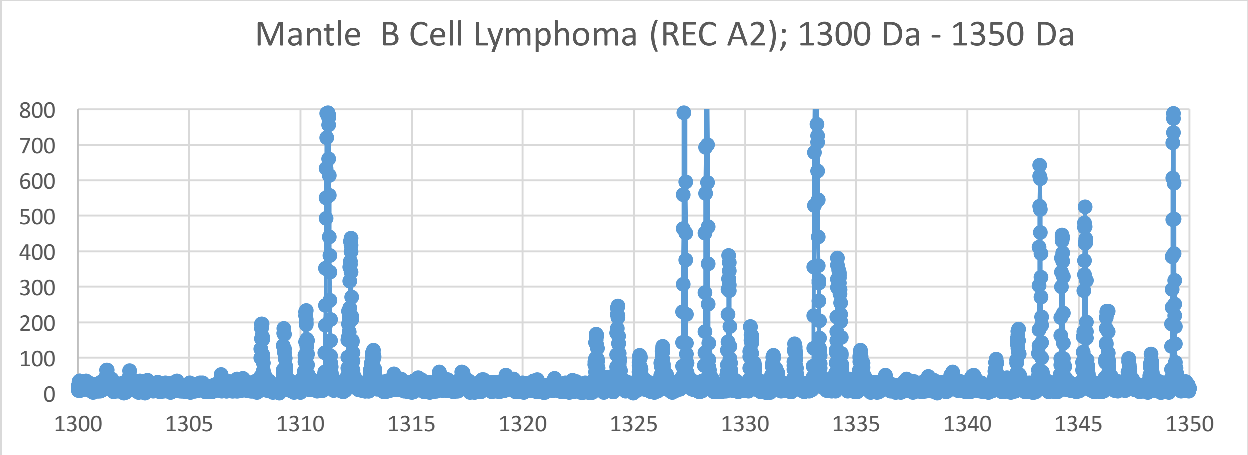
Mantle B Cell Lymphoma (REC A2); 1300 Da - 1350 Da
800
700
600
500
400
300
200
100
0
1300
1305
1310
1315
1320
1325
1330
1335
1340
1345
1350

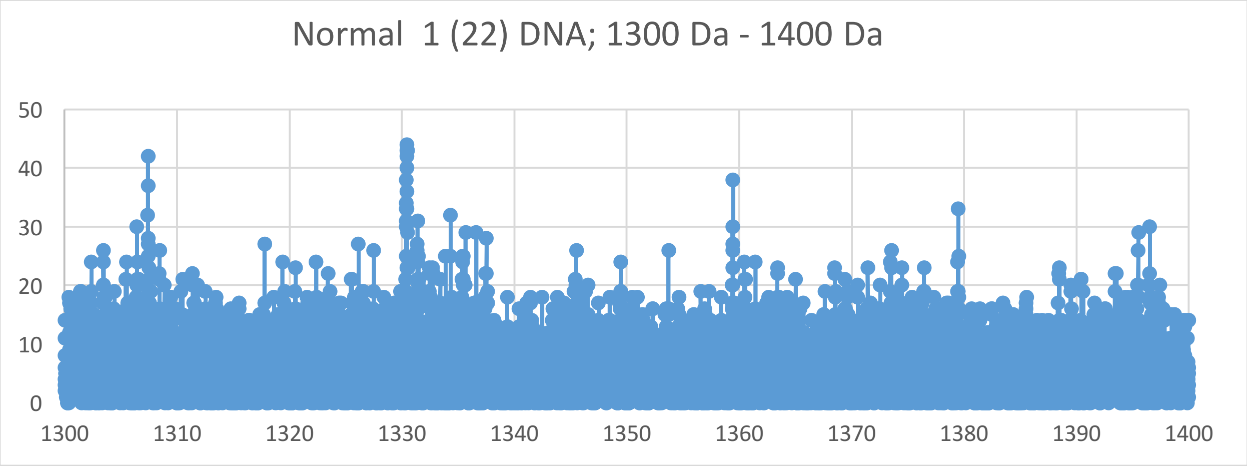
Normal 1 (22) DNA; 1300 Da - 1400 Da
50
40
30
20
10
0
1300
1310
1320
1330
1340
1350
1360
1370
1380
1390
1400

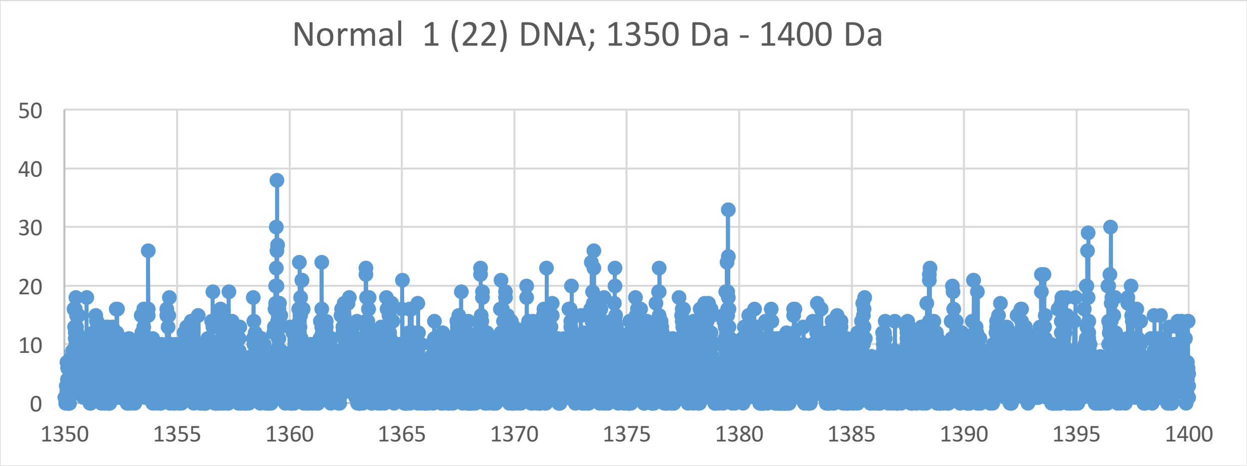
Normal 1 (22) DNA; 1350 Da - 1400 Da
50
40
30
20
10
0
1350
1355
1360
1365
1370
1375
1380
1385
1390
1395
1400

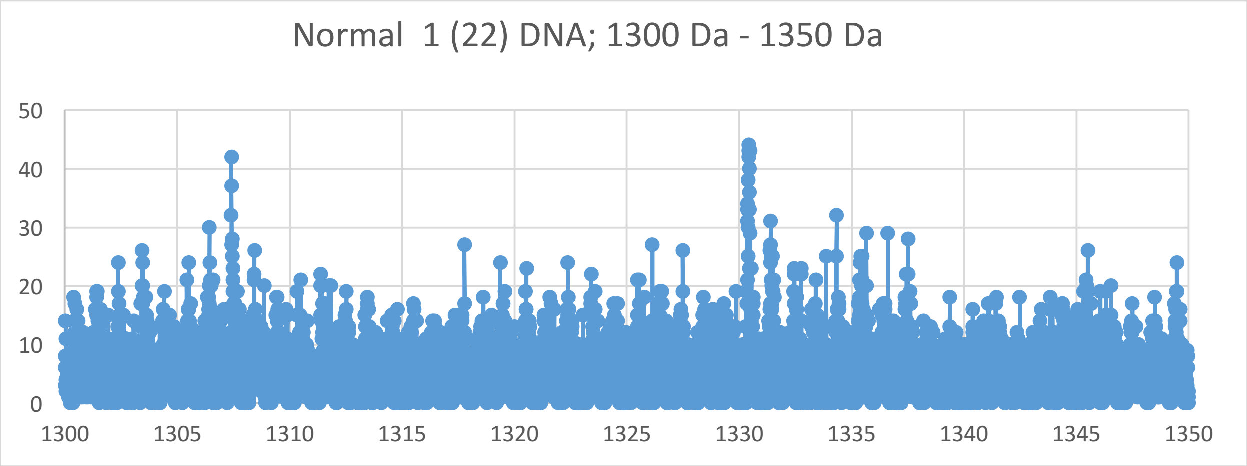
Normal 1 (22) DNA; 1300 Da - 1350 Da
50
40
30
20
10
0
1300
1305
1310
1315
1320
1325
1330
1335
1340
1345
1350

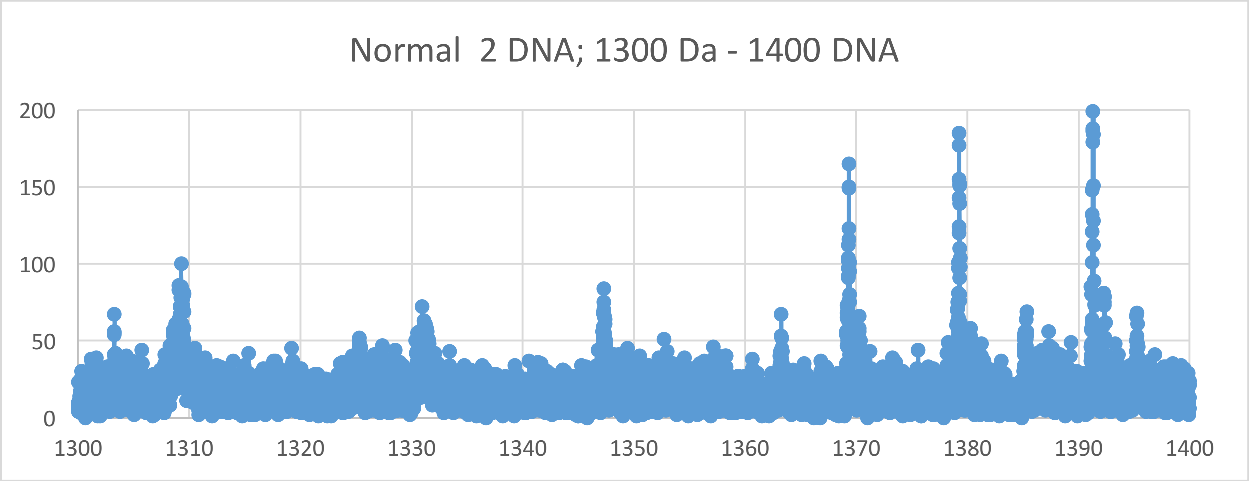
Normal 2 DNA; 1300 Da - 1400 DNA
200
150
100
50
0
1300
1310
1320
1330
1340
1350
1360
1370
1380
1390
1400

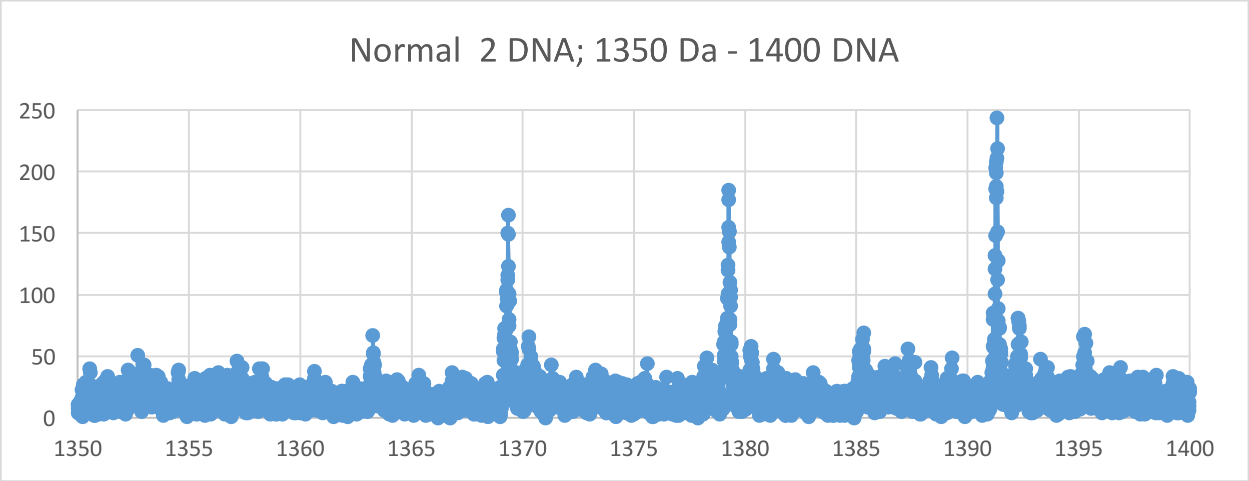
Normal 2 DNA; 1350 Da - 1400 DNA
250
200
150
100
50
0
1350
1355
1360
1365
1370
1375
1380
1385
1390
1395
1400

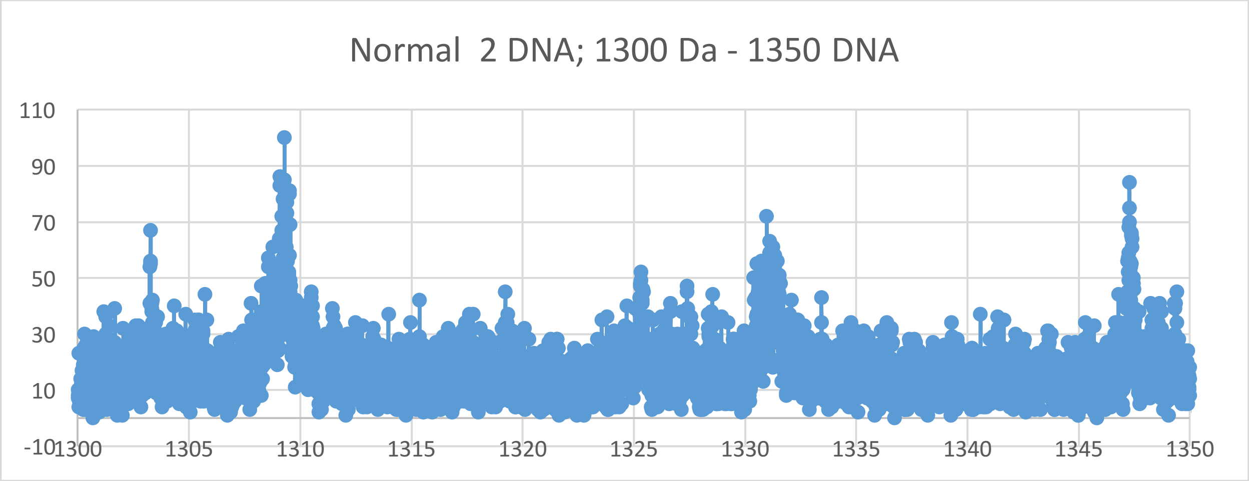
Normal 2 DNA; 1300 Da - 1350 DNA
110
90
70
50
30
10
-10
1300
1305
1310
1315
1320
1325
1330
1335
1340
1345
1350

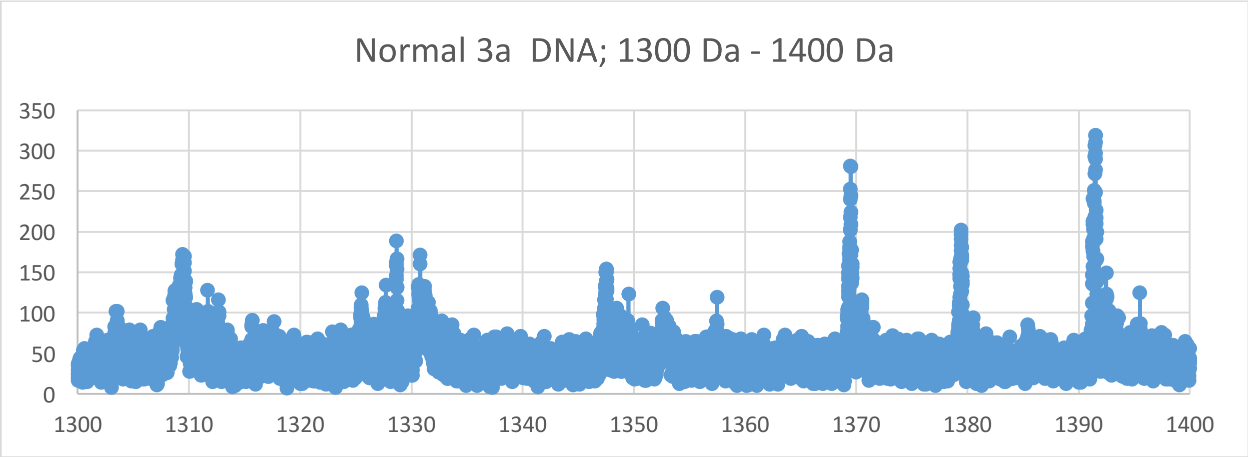
Normal 3a  DNA; 1300 Da - 1400 Da
350
300
250
200
150
100
50
0
1300
1310
1320
1330
1340
1350
1360
1370
1380
1390
1400

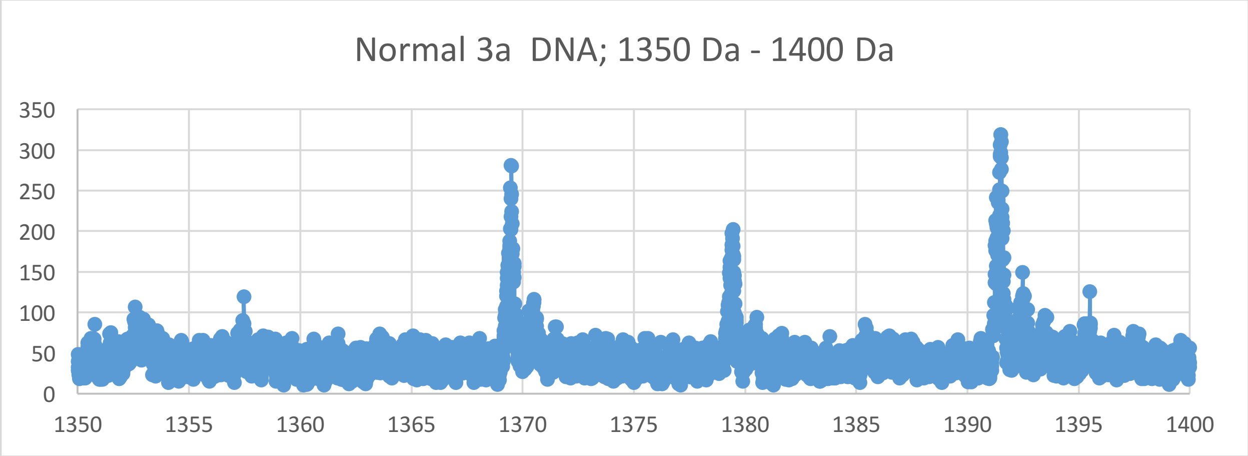
Normal 3a  DNA; 1350 Da - 1400 Da
350
300
250
200
150
100
50
0
1350
1355
1360
1365
1370
1375
1380
1385
1390
1395
1400

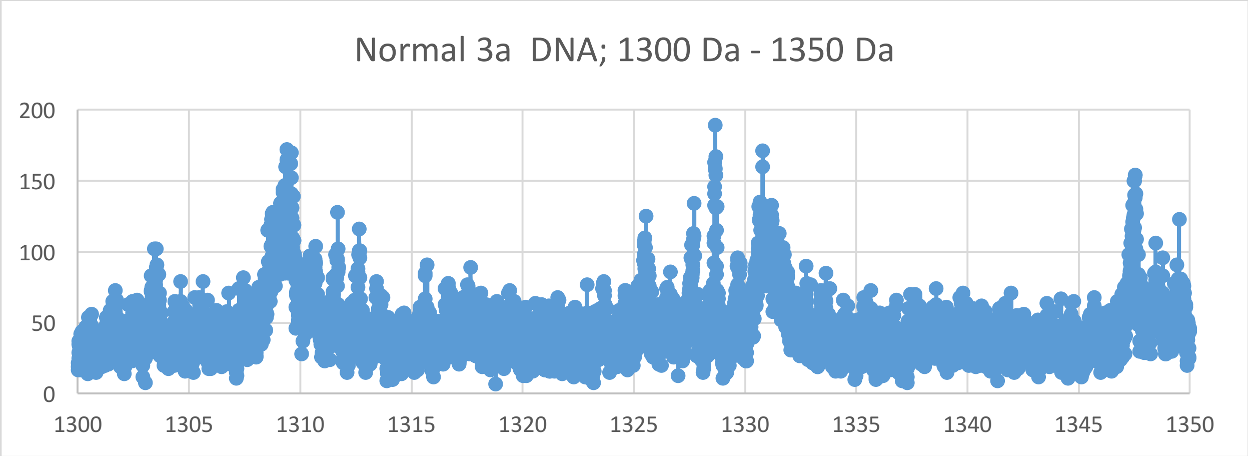
Normal 3a  DNA; 1300 Da - 1350 Da
200
150
100
50
0
1300
1305
1310
1315
1320
1325
1330
1335
1340
1345
1350

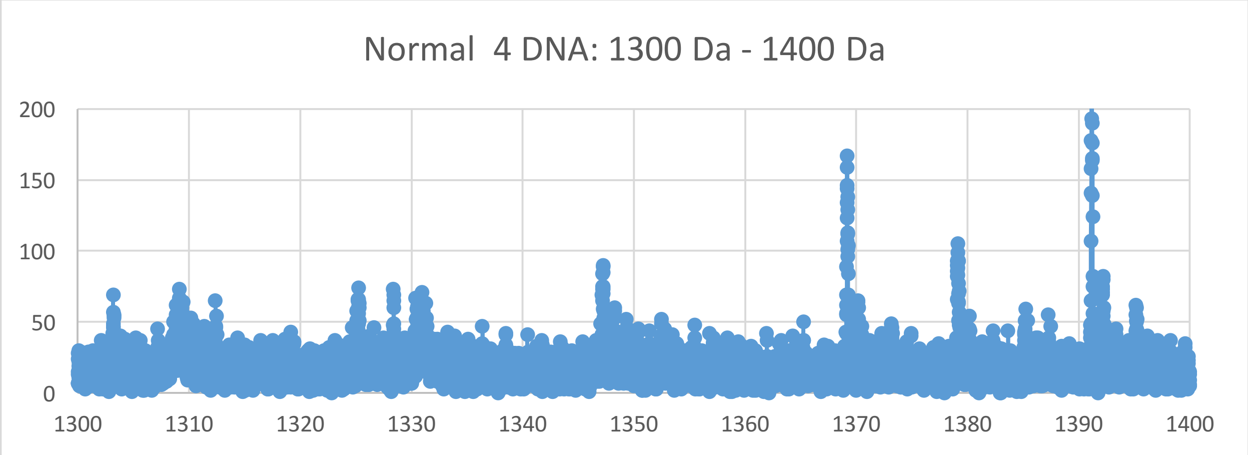
Normal 4 DNA: 1300 Da - 1400 Da
0
50
100
150
200
1300
1310
1320
1330
1340
1350
1360
1370
1380
1390
1400

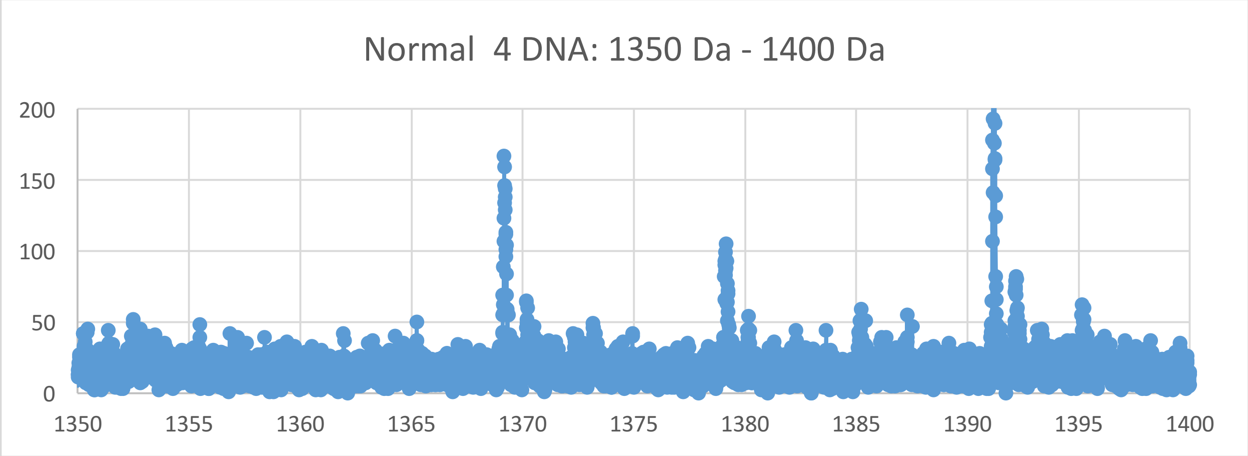
Normal 4 DNA: 1350 Da - 1400 Da
0
50
100
150
200
1350
1355
1360
1365
1370
1375
1380
1385
1390
1395
1400

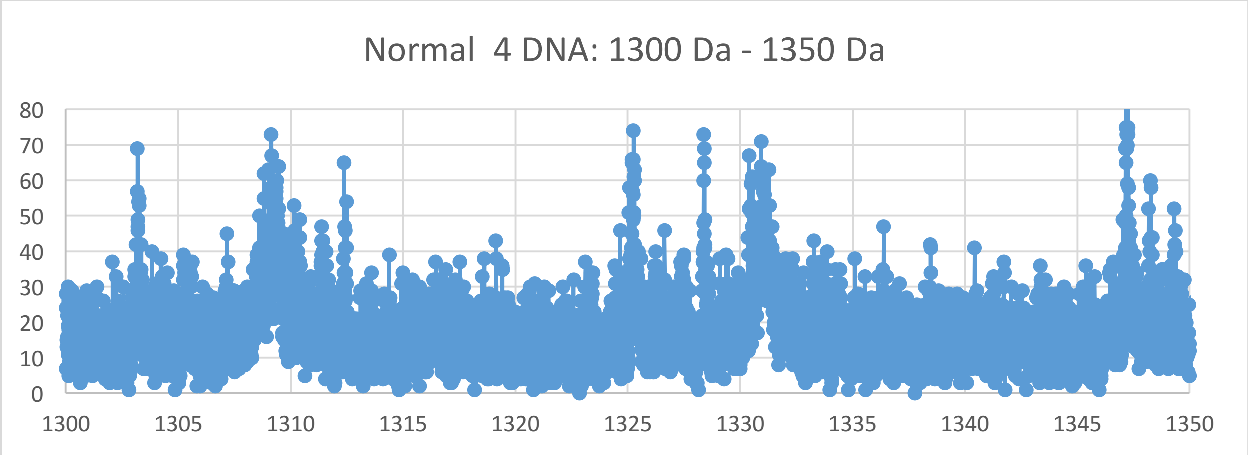
Normal 4 DNA: 1300 Da - 1350 Da
0
10
20
30
40
50
60
70
80
1300
1305
1310
1315
1320
1325
1330
1335
1340
1345
1350
