## Supplementary Materials VII for "Oncogenesis and Aging by Isotopic Functionalizations of the Proteins and Nucleic Acids"

VII Supplementary Materials – New Mass Spectra from 1200 Da -1300 Da

Results for Supplementary 7

So for the trinucleotides at $(phos)_7$, the telomeric peaks for cancer strongly decrease in intensities and number and the peaks for normal increases in number and intensity for less stable cancer trinucleotides and more stable normal trinucleotides with the $(phos)_7$. But the mating telomeres and nontelomeres have decrease in number and stabilities for cancer trinucleotides while the normal trinucleotides are slightly more in peaks and stabilities with $(phos)_7$. But as decrease the phosphate by 1 unit, the trinucleotides change so that the normal trinucleotides more fragment and break from the DNA with more number and stabilities for the telomeres, mating telomeres and nontelomeres, with more fragmenting and stabilities of the normal from DNA for telomeres and mating telomeres from $(phos)_7$ to $(phos)_6$ . The masses range from 1200 Da to 1300 Da See Supplementary Figures 7 for graphs. It seems that below some number of phosphates, the telomeres, mating telomeres and nontelomers behave in same way toward cancer and normal. So this suggests normal may be favored and distinguished from cancer by increasing or changing saturated phosphorylations. The telomeric peaks for normal increase with these dephosphorylations of the trinucleotides. The dephosphorylations from $(phos)_7$ to $(phos)_6$ caused the resulting more intense and stable normal trinucleotides to be more mating telomeric, telomeric and nontelomeric trinucleotides with more isotopic enrichments for telomeres and nontelomeres. It is important to consider that there are more intense peaks for the trinucleotides relative to the tetranucleotides for the saturated phosphate units. For the trinucleotides, the intensities of normal increased and the intensities of cancer fragments decreased from $(phos)_7$ to $(phos)_6$ . The dephosphorylations from $(phos)_7$ to $(phos)_6$ increase isotopic enrichments of heavy isotopes of $^{13}C$ and $^{17}O$ for mating telomeres. The dephosphorylations from $(phos)_7$ to $(phos)_6$ also caused the telomeric and nontelomeric trinucleotides to be more intense in normal DNA and more isotopically enriched from $(phos)_7$ to $(phos)_6$. It seems the dephosphorylations are related to the isotopic enrichments and depletions; but the telomeres and nontelomeres are more intense in normal and more isotopically enriched whereas the intensities of the mating telomeres are negligibly changed in cancer and the more isotopic enriched from $(phos)_7$ to $(phos)_6$. The telomeric, mating telomeric and nontelomeric trinucleotides of normal tended to also be more enriched so it seems the normal stabilities of the telomeric and nontelomeric trinucleotides and the cancer stabilities of the mating telomeres are related to their more isotopic enrichments for the dephosphorylations from $(phos)_7$ to $(phos)_6$ and and not purely driven by dephosphorylations and enrichments. Enrichments strengthen the telomeric and nontelomeric normal trinucleotides while strongly weakening normal mating telomeric trinucleotides. The more intense normal nontelomeric and telomeric trinucleotides were also more isotopically enriched. The dephosphorylations from $(phos)_7$ to $(phos)_6$ strongly increased the isotopic enrichments of the normal for the nontelomeric and telomeric trinucleotides and cancer for mating telomeres had strongly enrichments of nonprimordials. But the nontelomeric trinucleotides were even more enriched from the dephosphorylations for greater enrichments and stronger more stable cancer trinucleotides for nontelomeres. Such seems compatible for replications and transcriptions as stable telomeres seem compatible to unstable mating telomers for the unravelings and constructions of DNA mating telomeres onto the building templates of the telomeres. The dephosphorylations from $(phos)_7$ to $(phos)_6$ caused methylations to cause negligible changes of cancer intensities, isotopic enrichments and stabilities for telomeres and nontelomeres. But the methylations of the mating telomeres slightly increased cancer stabilities and decreased isotopic enrichments for mating telomeres. Such loss of differing behavior of telomeres and nontelomeres with greater dephosphorylations points to malfunctions of DNA by highly dephosphorylated nucleotides and plays a role in cancer. The methylations of the nontelomeres does not slightly increased cancer intensities and stabilities with much less enrichments from $(phos)_7$ to $(phos)_6$. It seems from these data that during biological processes the dephosphorylations and stabilizations of normal or vice versa dephosphorylations and stabilizations of cancer are or can be countered by methylations and destabilizations of the normal for cancer or vice versa dephosphorylations and destabilizations of the cancer by methylations as this research discovers coupled phosphorylations and methylations or coupled dephosphorylations and methylations with involved greater enrichments for normal biological functions and explaining cancer! It seems that if nucleotides lose to many phosphates in regulating methylations and actylations that the differences among genes and in particular telomeric genes and nontelomeric genes are loss and the activity of cancer arises! The dephosphorylations from $(phos)_7$ to $(phos)_6$ caused acetylations to cause slightly more cancer stabiliies for telomeres and nontelomeres and more normal instabilities for telomeres and nontelomeres with slightly less enrichments for telomeres, mating telomeres and nontelomeres. The acetylations from $(phos)_7$ to $(phos)_6$ caused more cancer stabilities and intensities for mating telomeres with much less enrichments. This contrast in mating telomeres and telomeres from the acetylations may reflect the developments of cancer as the telomeres fragment but the mating telomeres more holds together for damaged DNA for cancer.

So for the trinucleotides at $(phos)_6$, the telomeric and nontelomeric peaks for mix of cancer and normal strongly increase in intensities and number and the peaks for some cancer and normal decrease in number and intensities for more mix of stabilities of normal and cancer trinucleotides and a few less stable cancer and normal trinucleotides with the $(phos)_6$. But the mating telomeres have increase in number and stabilities for cancer trinucleotides while the normal trinucleotides are mostly less in peaks and stabilities with $(phos)_6$. But as decrease the phosphate by 1 unit, the trinucleotides change so that the normal trinucleotides more fragment and break from the DNA with more number and stabilities for the telomeres and nontelomeres, with slightly more fragmenting and stabilities of the normal from DNA for telomeres from $(phos)_6$ to $(phos)_5$. The masses range from 1169 Da to 1312 Da. See Supplementary Figures VII for graphs. It seems that below some number of phosphates, the telomeres, mating telomeres and nontelomers behave in same way toward cancer and normal. So this suggest normal may be favored and distinguished from cancer by changing phosphorylations. The telomeric peaks for normal increase in numbers with these dephosphorylations of the trinucleotides. The dephosphorylations from $(phos)_6$ to $(phos)_5$ caused the resulting more intense and stable normal trinucleotides to be slightly more for mating telomeric trinucleotides, more normal stabilities for telomeric and nontelomeric trinucleotides and less isotopically enrichments for telomeres, but more isotopic enrichments for mating telomeres and nontelomeres. It is important to consider that there are more intense peaks for the trinucleotides relative to the tetranucleotides for the saturated phosphate units. For the trinucleotides the intensities of both normal and cancer fragments decreased from $(phos)_6$ to $(phos)_5$. The dephosphorylations from $(phos)_6$ to $(phos)_5$ decreases isotopic enrichments of heavy isotopes of $^{13}C$ and $^{17}O$ for telomeric trinucleotides but not for mating telomeric and nontelomeric trinucleotides. The dephosphorylations from $(phos)_6$ to $(phos)_5$ also caused the telomeric and nontelomeric trinucleotides to be more intense in normal DNA and less isotopically enriched, but mating telomeres change to be more intense in both normal and cancer with more isotopic enrichments for mixed trinucleotides by dephosphorylations from $(phos)_6$ to $(phos)_5$. It seems the dephosphorylations are related to the isotopic enrichments and depletions; but the telomeres and nontelomeres are slightly more numerous in normal and less isotopically enriched whereas the frequencies of the mating telomeres are mixed more numerous in cancer and normal trinucleotides and the more isotopic enriched from $(phos)_6$ to $(phos)_5$. In general it seems with less phosphorylations there is less isotopic enrichments and the difference between telomeric, nontelomeric and mating telomeric binding and stabilities diminish to induce cancer by such chemical mutations! The telomeric, mating telomeric and nontelomeric trinucleotides of normal tended to also be less enriched with heavy $^{13}C$ and $^{17}O$ and more enriched with light $^{15}N$; so it seems the normal stabilities of the telomeric, mating telomeric and nontelomeric trinucleotides and the cancer stabilities of the mating telomeres are related to their less isotopic enrichments with $^{13}C$ and $^{17}O$ but more enrichments with $^{15}N$ for the dephosphorylations from $(phos)_6$ to $(phos)_5$ and not purely driven by dephosphorylations but both dephosphorylations and enrichments. Enrichments strengthen the telomeric and nontelomeric normal trinucleotides while strongly weakening normal mating telomeric trinucleotides. The more intense normal nontelomeric and telomeric trinucleotides were also less isotopically enriched with heavy $^{13}C$ and $^{17}O$ but more isotopically enriched with $^{15}N$. The dephosphorylations from $(phos)_6$ to $(phos)_5$ strongly increased the $^{15}N$ isotopic enrichments of the normal for the nontelomeric and telomeric trinucleotides and cancer and normal for mating telomeres had strongly enrichments of $^{13}C$, $^{17}O$ and $^{15}N$ nonprimordials. But the mating telomeres and nontelomeric trinucleotides were even more enriched from the dephosphorylations for greater enrichments and stronger more stable cancer and normal trinucleotides for mating telomeres and nontelomeres. Such seems compatible for replications and transcriptions as stable telomeres seem compatible to unstable mating telomeres and nontelomeres for the unravelings and constructions of DNA mating telomeres onto the building templates of the telomeres. The dephosphorylations from $(phos)_6$ to $(phos)_5$ caused methylations to cause slightly increase cancer number of peaks but increase the intensities of normal peaks and stabilities with for telomeres and nontelomeres with more isotopic enrichments for telomeres and and more isotopic enrichments for mating telomeres and nontelomeres. But the methylations of the mating telomeres increased normal stabilities and slightly increased isotopic enrichments for mating telomeres. Such loss of differing behavior of telomeres and nontelomers with greater dephosphorylations points to malfunctions of DNA by highly dephosphorylated nucleotides and plays a role in cancer. The methylations of the telomeres does slightly increased cancer intensities and stabilities with less enrichment from $(phos)_6$ to $(phos)_5$ relative to the nontelomeres. It seems from these data that during biological processes the dephosphorylations and stabilizations of normal or vice versa dephosphorylations and stabilizations of cancer are or can be countered by methylations and destabilizations of the normal for cancer or vice versa dephosphorylations and destabilizations of the cancer by methylations as this research discovers coupled phosphorylations and methylations or coupled dephosphorylations and methylations with involved greater enrichments for normal biological functions and explaining cancer! It seems that if nucleotides lose to many phosphates in regulating methylations and acteylations that the differences among genes and in particular telomeric genes and nontelomeric genes are lost and the activity of cancer arises! The dephosphorylations from $(phos)_6$ to $(phos)_5$ caused acetylation to cause greatly more cancer stabilities for telomeres and slightly more cancer stabilities for nontelomeres with more enrichments for telomeres and nontelomeres. The acetylations from $(phos)_6$ to $(phos)_5$ caused more cancer stabilities and intensities for mating telomeres with much less enrichments of light isotopes but more enrichments of heavy isotopes. This contrast in mating telomeres and telomeres from the acetylations may reflect the developments of cancer as the telomeres fragment but the mating telomeres more holds together for damaged DNA for cancer.

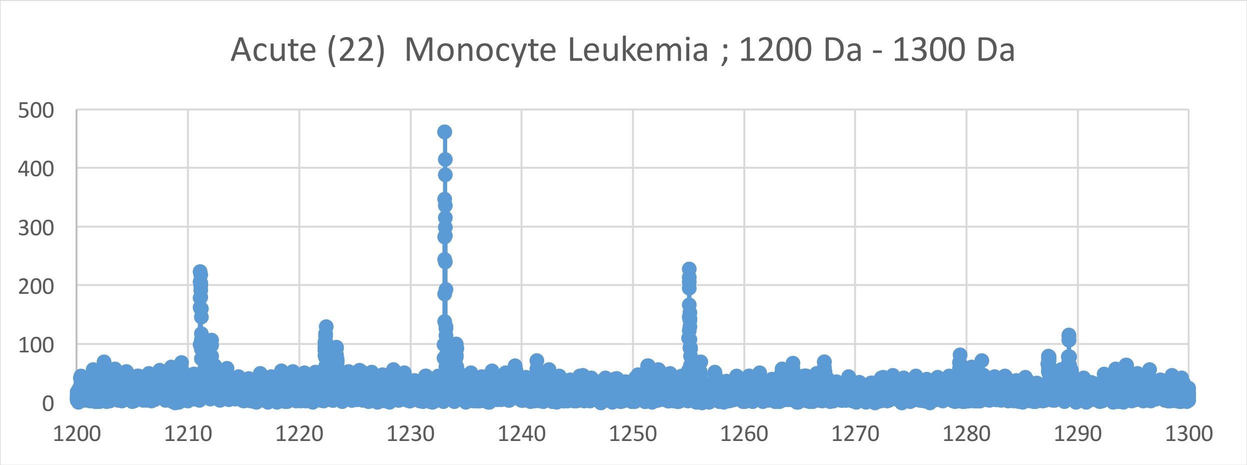

Acute (22) Monocyte Leukemia ; 1200 Da - 1300 Da
500
400
300
200
100
0
1200
1210
1220
1230
1240
1250
1260
1270
1280
1290
1300


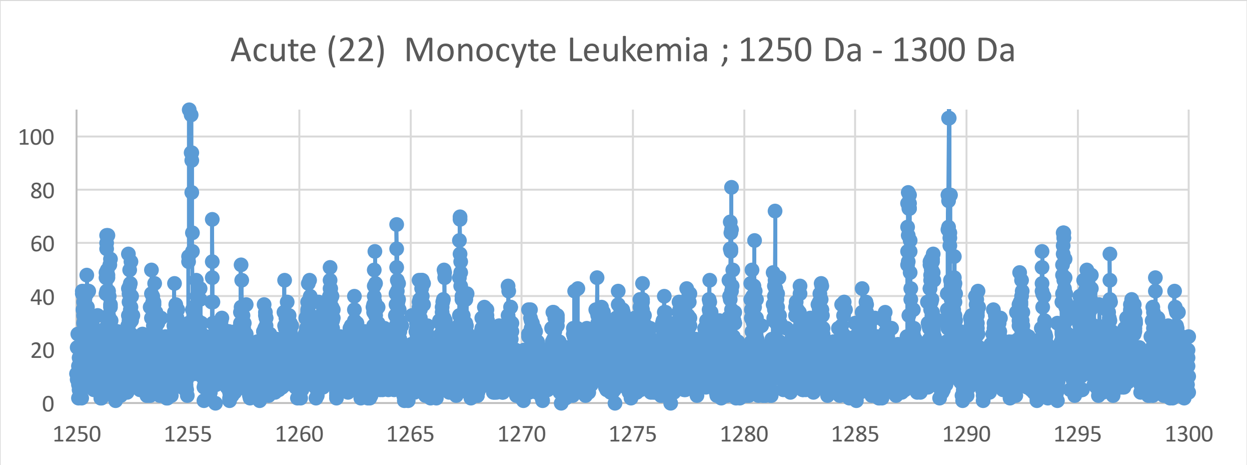

Acute (22) Monocyte Leukemia ; 1250 Da - 1300 Da
100
80
60
40
20
0
1250
1255
1260
1265
1270
1275
1280
1285
1290
1295
1300


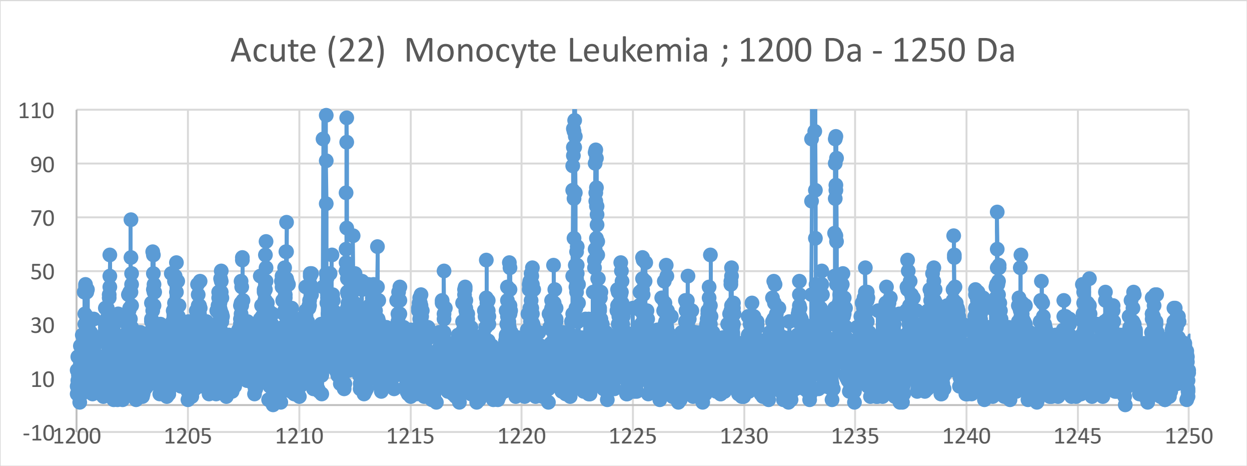

Acute (22) Monocyte Leukemia ; 1200 Da - 1250 Da
110
90
70
50
30
10
-10
1200
1205
1210
1215
1220
1225
1230
1235
1240
1245
1250

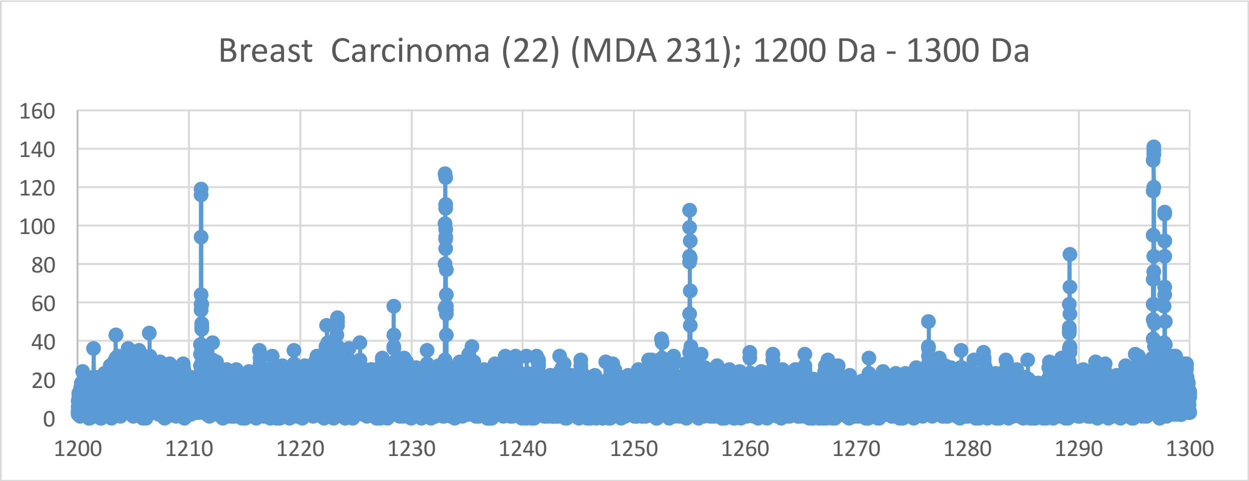
Breast Carcinoma (22) (MDA 231); 1200 Da - 1300 Da
0
20
40
60
80
100
120
140
160
1200
1210
1220
1230
1240
1250
1260
1270
1280
1290
1300

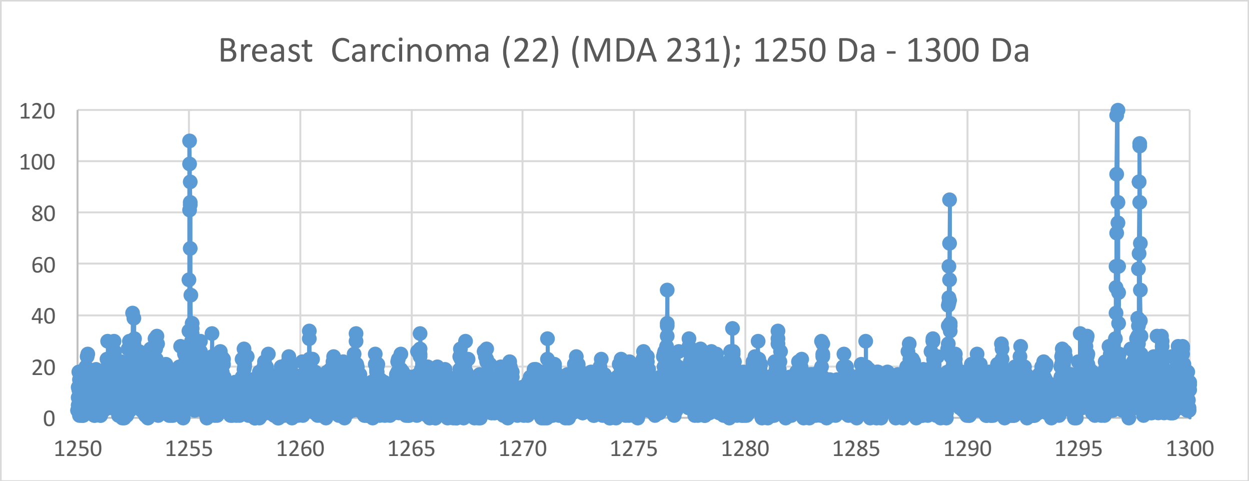
Breast Carcinoma (22) (MDA 231); 1250 Da - 1300 Da
0
20
40
60
80
100
120
1250
1255
1260
1265
1270
1275
1280
1285
1290
1295
1300

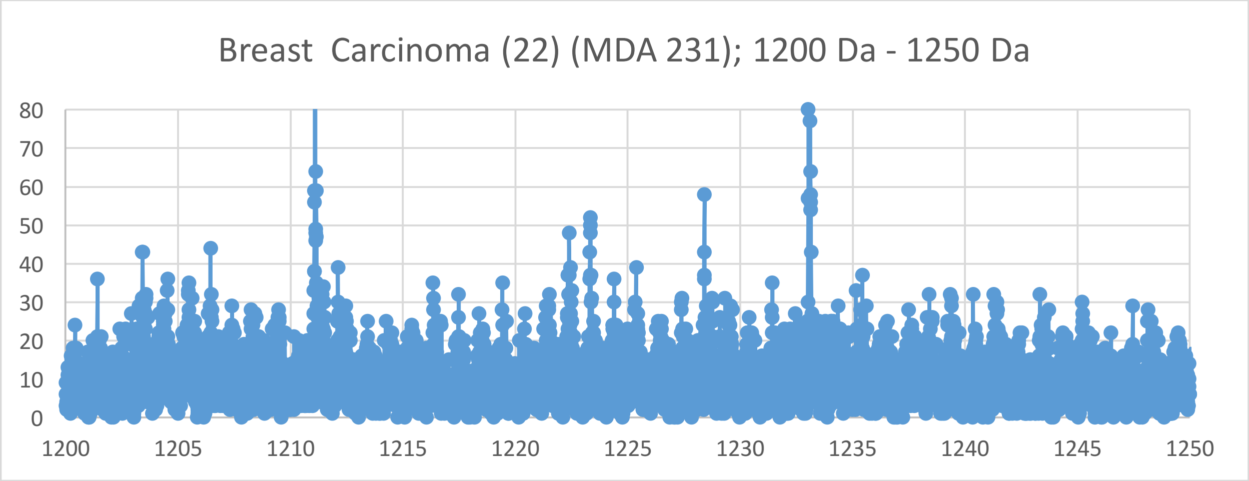
Breast Carcinoma (22) (MDA 231); 1200 Da - 1250 Da
0
10
20
30
40
50
60
70
80
1200
1205
1210
1215
1220
1225
1230
1235
1240
1245
1250

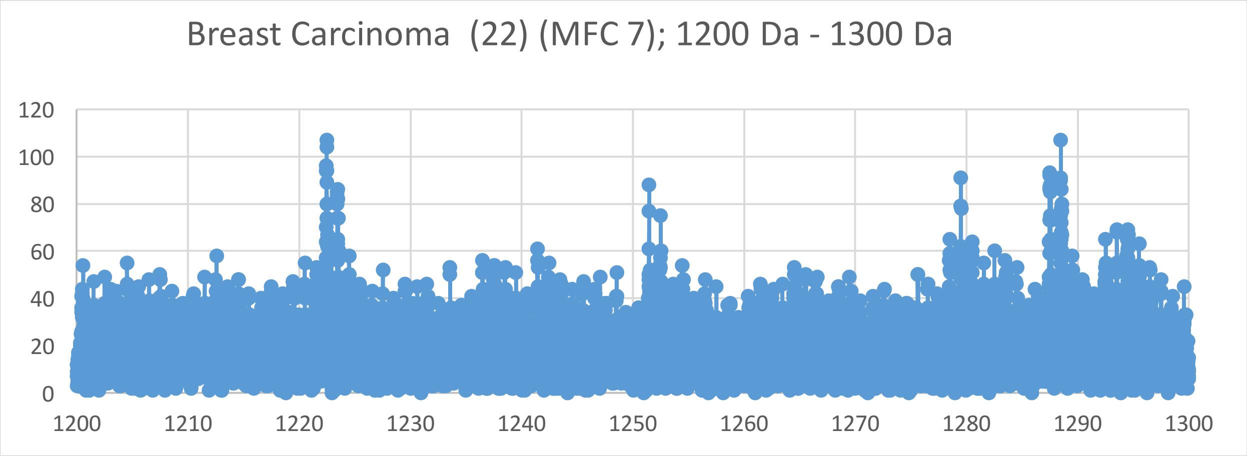

Breast Carcinoma (22) (MFC 7); 1200 Da - 1300 Da
120
100
80
60
40
20
0
1200
1210
1220
1230
1240
1250
1260
1270
1280
1290
1300


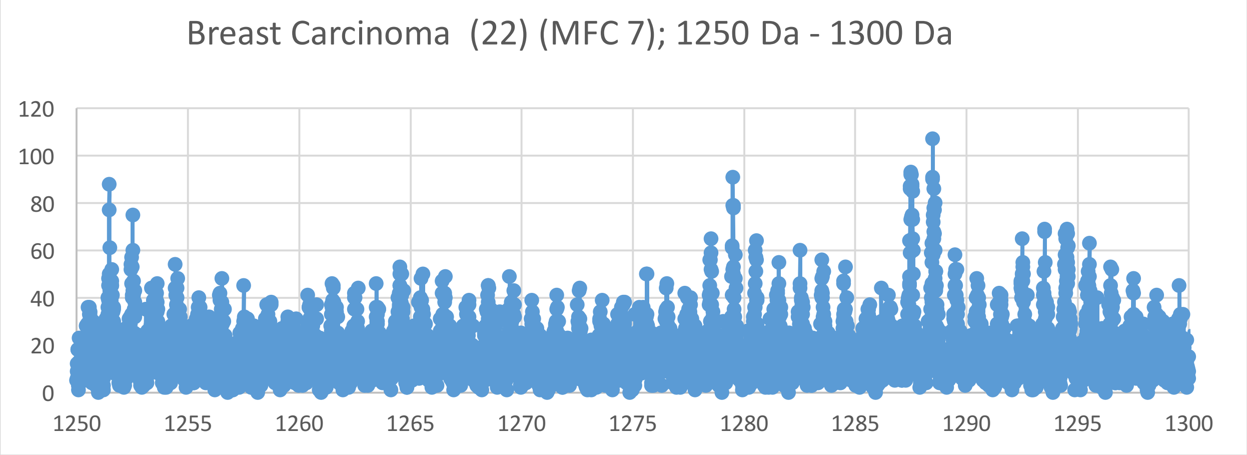

Breast Carcinoma (22) (MFC 7); 1250 Da - 1300 Da
120
100
80
60
40
20
0
1250
1255
1260
1265
1270
1275
1280
1285
1290
1295
1300


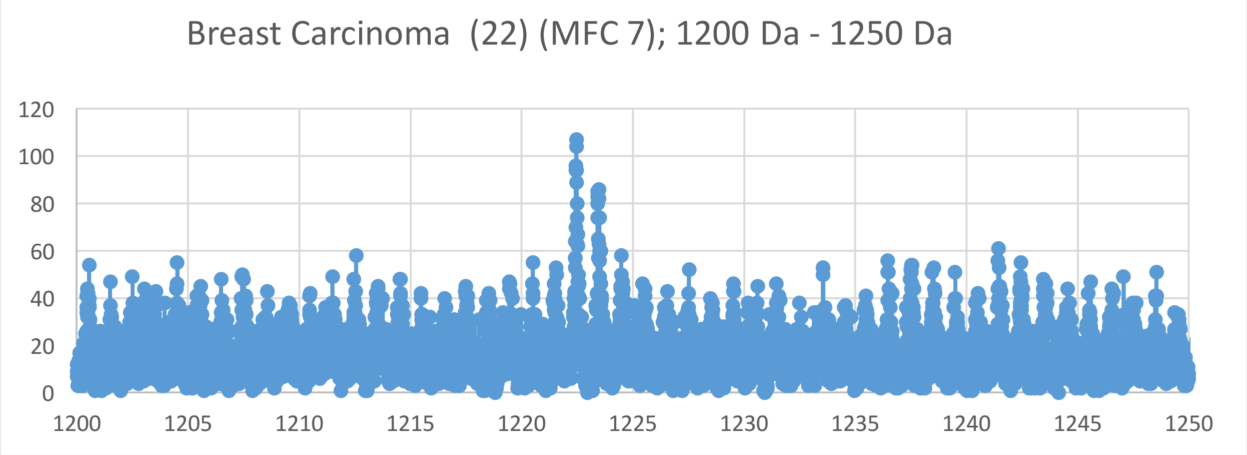

Breast Carcinoma (22) (MFC 7); 1200 Da - 1250 Da
120
100
80
60
40
20
0
1200
1205
1210
1215
1220
1225
1230
1235
1240
1245
1250

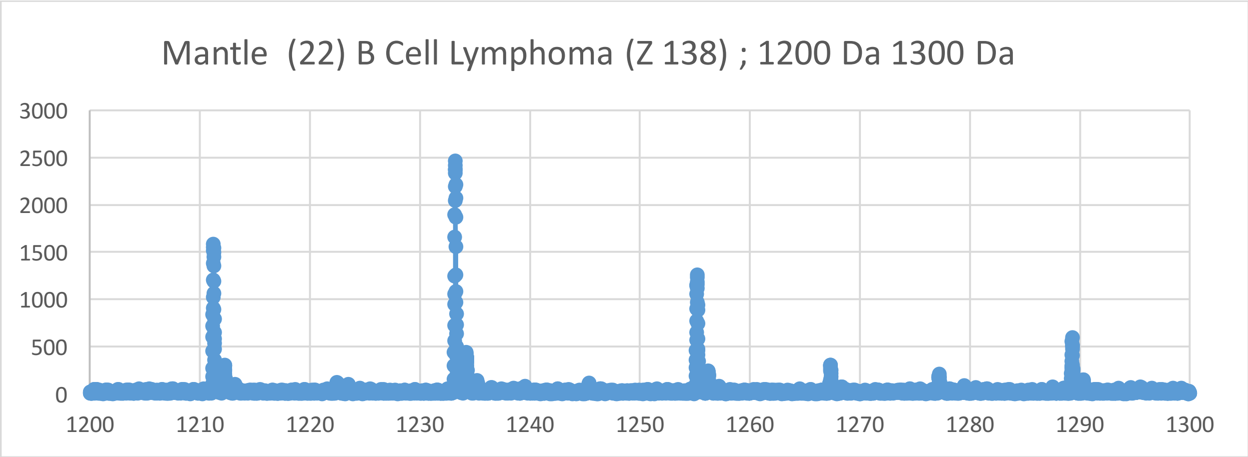
Mantle (22) B Cell Lymphoma (Z 138) ; 1200 Da 1300 Da
3000
2500
2000
1500
1000
500
0
1200
1210
1220
1230
1240
1250
1260
1270
1280
1290
1300

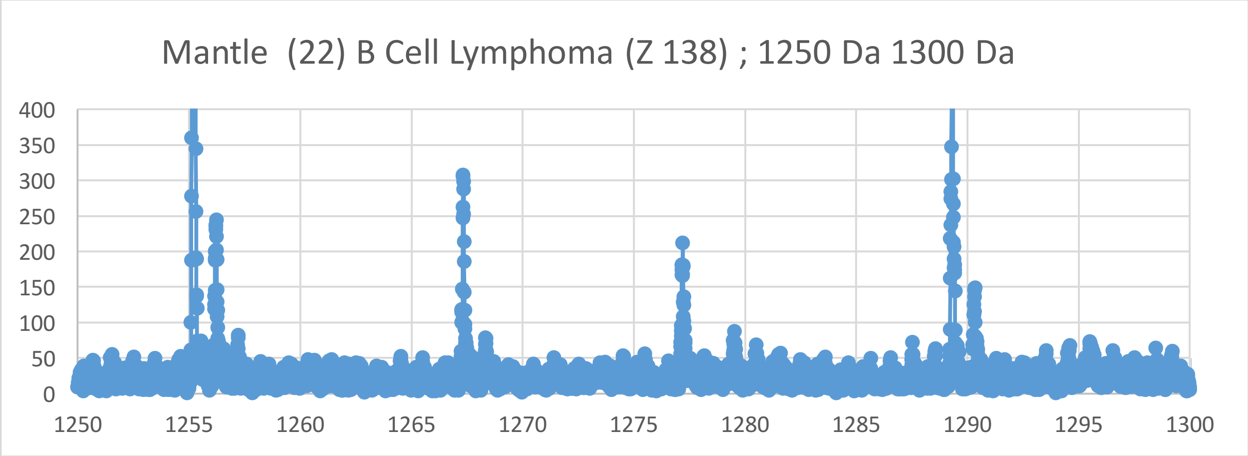
Mantle (22) B Cell Lymphoma (Z 138) ; 1250 Da 1300 Da
400
350
300
250
200
150
100
50
0
1250
1255
1260
1265
1270
1275
1280
1285
1290
1295
1300

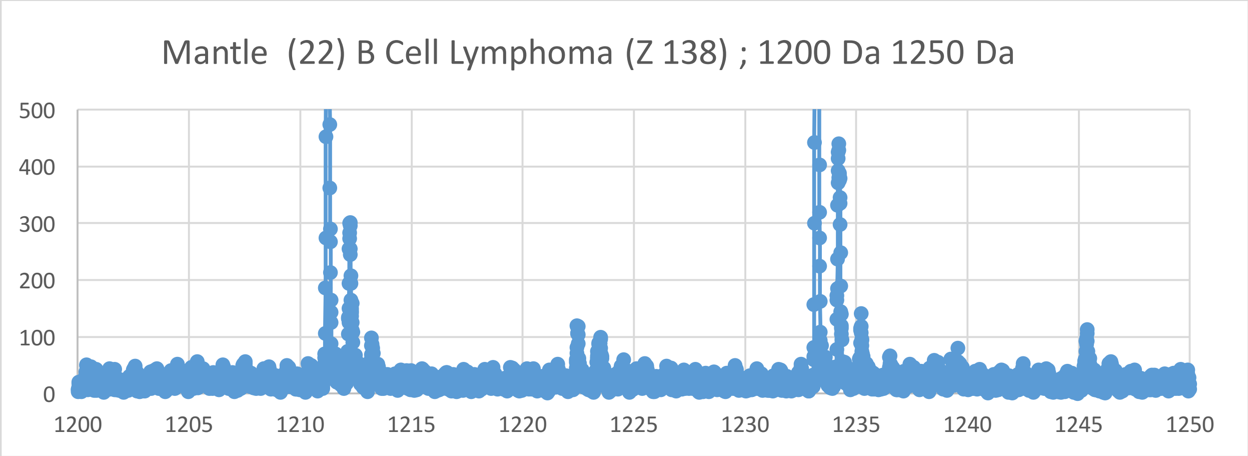
Mantle (22) B Cell Lymphoma (Z 138) ; 1200 Da 1250 Da
500
400
300
200
100
0
1200
1205
1210
1215
1220
1225
1230
1235
1240
1245
1250

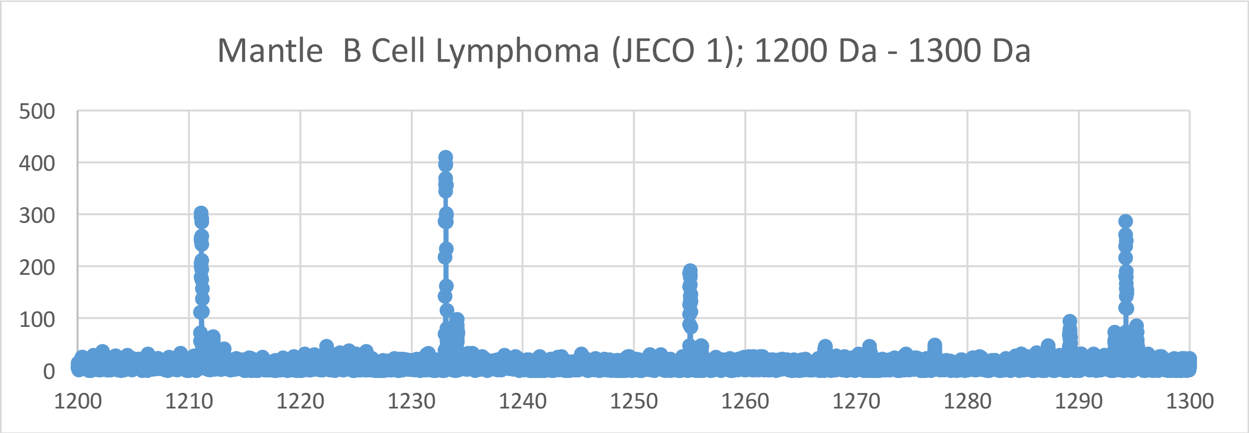
Mantle B Cell Lymphoma (JECO 1); 1200 Da - 1300 Da
500
400
300
200
100
0
1200
1210
1220
1230
1240
1250
1260
1270
1280
1290
1300

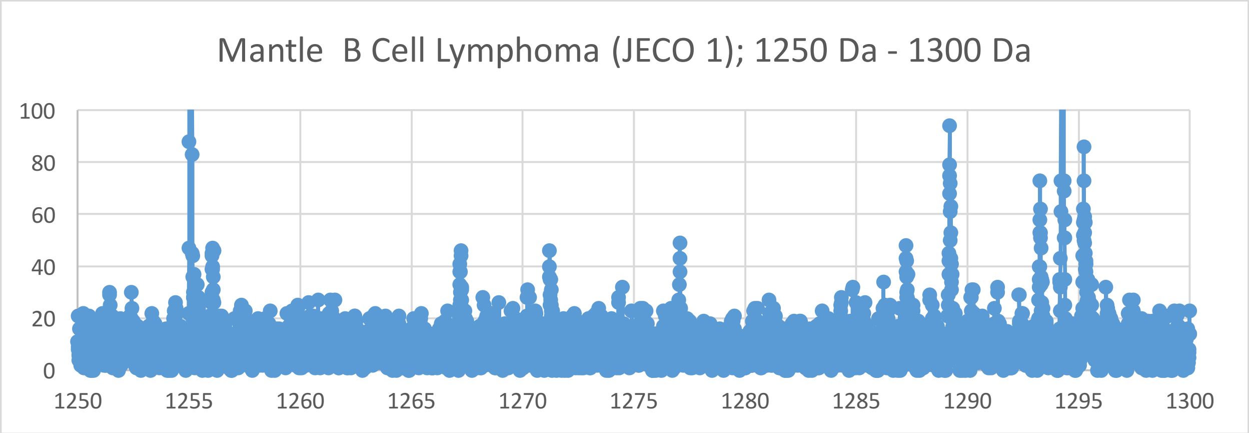
Mantle B Cell Lymphoma (JECO 1); 1250 Da - 1300 Da
100
80
60
40
20
0
1250
1255
1260
1265
1270
1275
1280
1285
1290
1295
1300

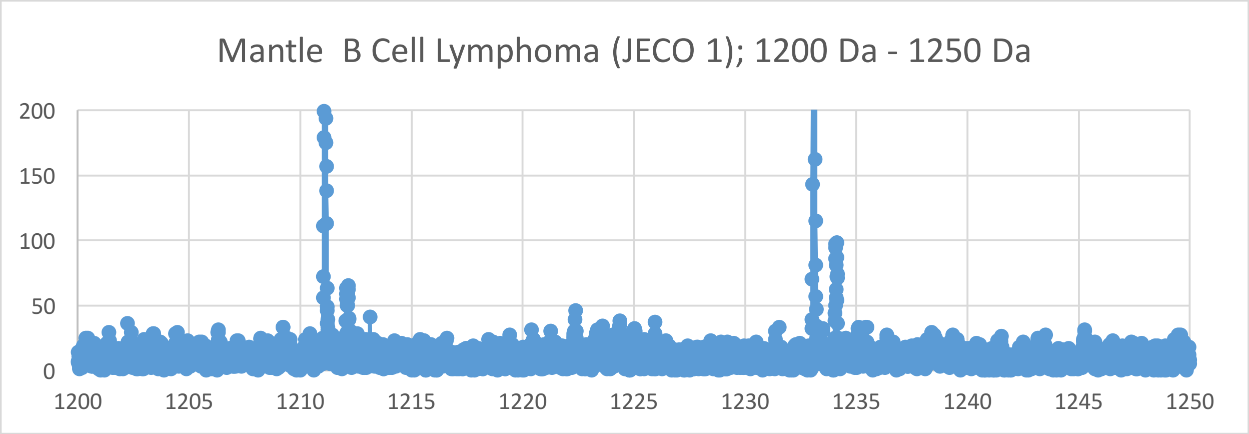
Mantle B Cell Lymphoma (JECO 1); 1200 Da - 1250 Da
200
150
100
50
0
1200
1205
1210
1215
1220
1225
1230
1235
1240
1245
1250

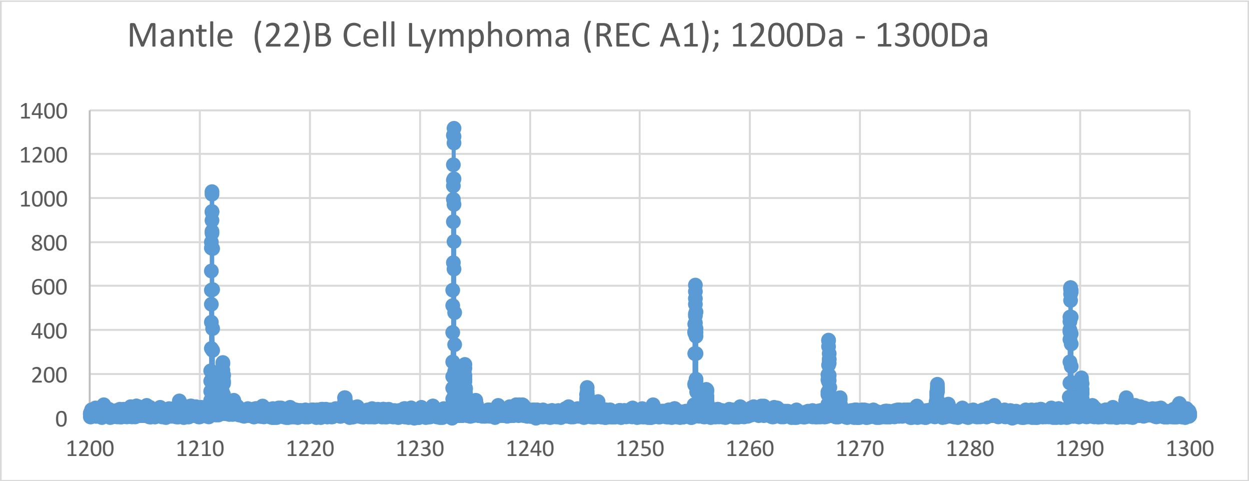
Mantle (22)B Cell Lymphoma (REC A1); 1200Da - 1300Da
1400
1200
1000
800
600
400
200
0
1200
1210
1220
1230
1240
1250
1260
1270
1280
1290
1300

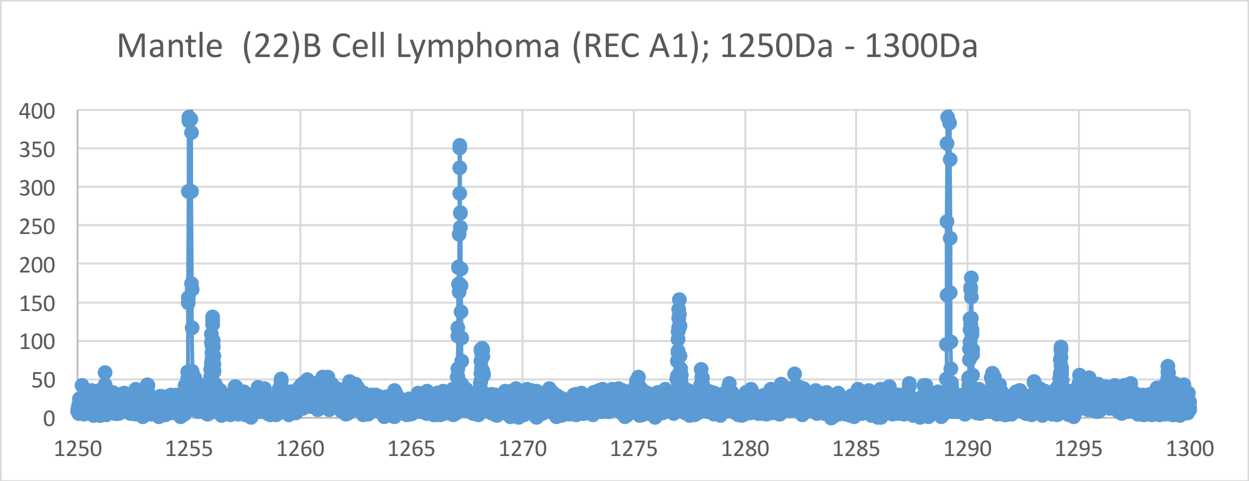
Mantle (22)B Cell Lymphoma (REC A1); 1250Da - 1300Da
400
350
300
250
200
150
100
50
0
1250
1255
1260
1265
1270
1275
1280
1285
1290
1295
1300

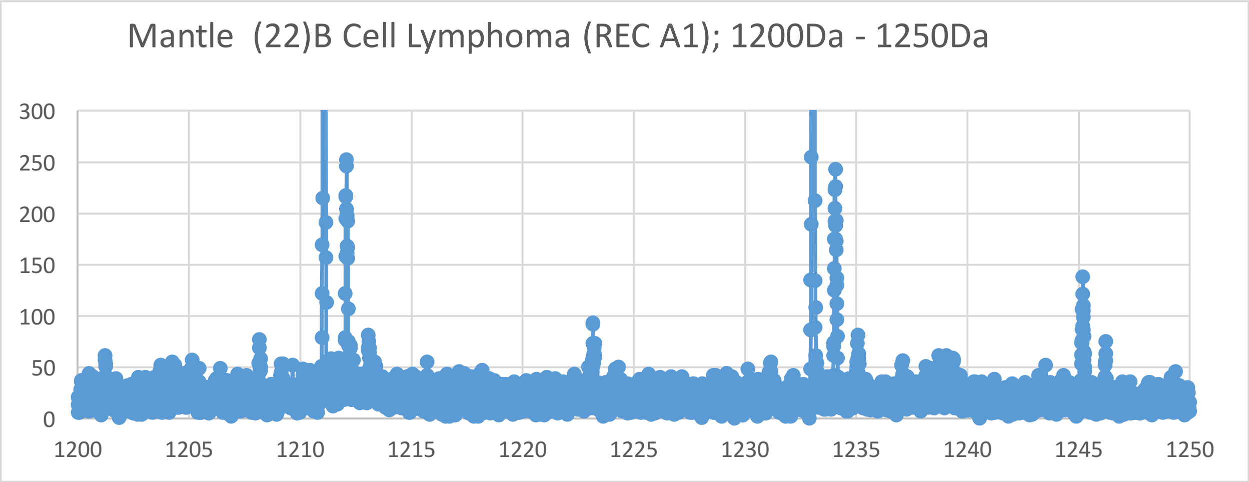
Mantle (22)B Cell Lymphoma (REC A1); 1200Da - 1250Da
300
250
200
150
100
50
0
1200
1205
1210
1215
1220
1225
1230
1235
1240
1245
1250

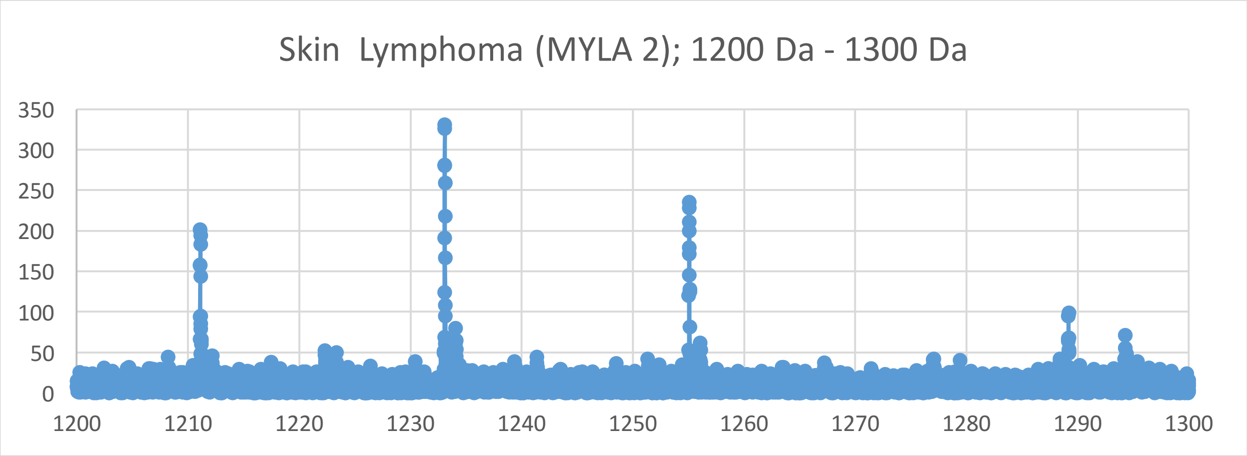
Skin Lymphoma (MYLA 2); 1200 Da - 1300 Da
350
300
250
200
150
100
50
0
1200
1210
1220
1230
1240
1250
1260
1270
1280
1290
1300

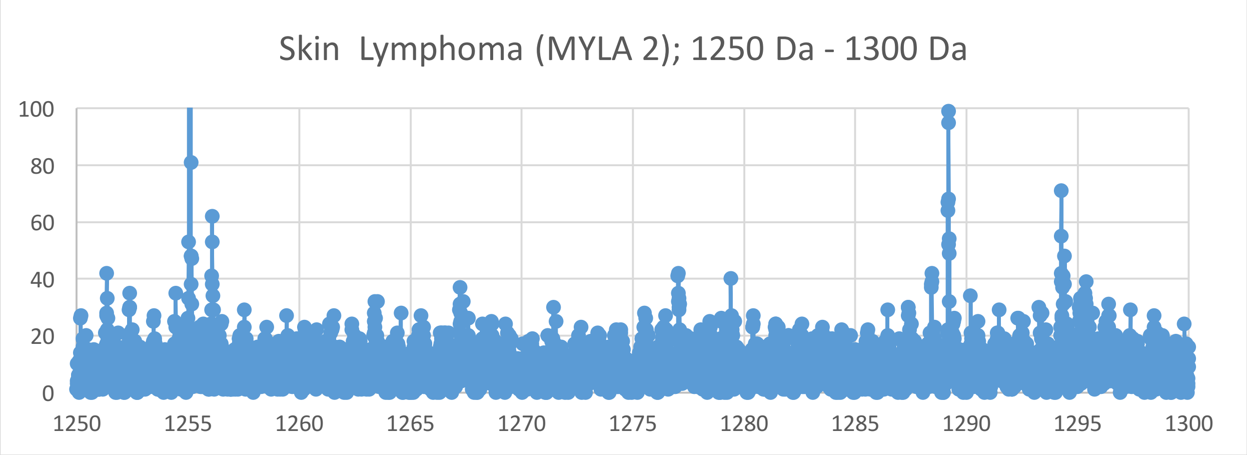
Skin Lymphoma (MYLA 2); 1250 Da - 1300 Da
100
80
60
40
20
0
1250
1255
1260
1265
1270
1275
1280
1285
1290
1295
1300

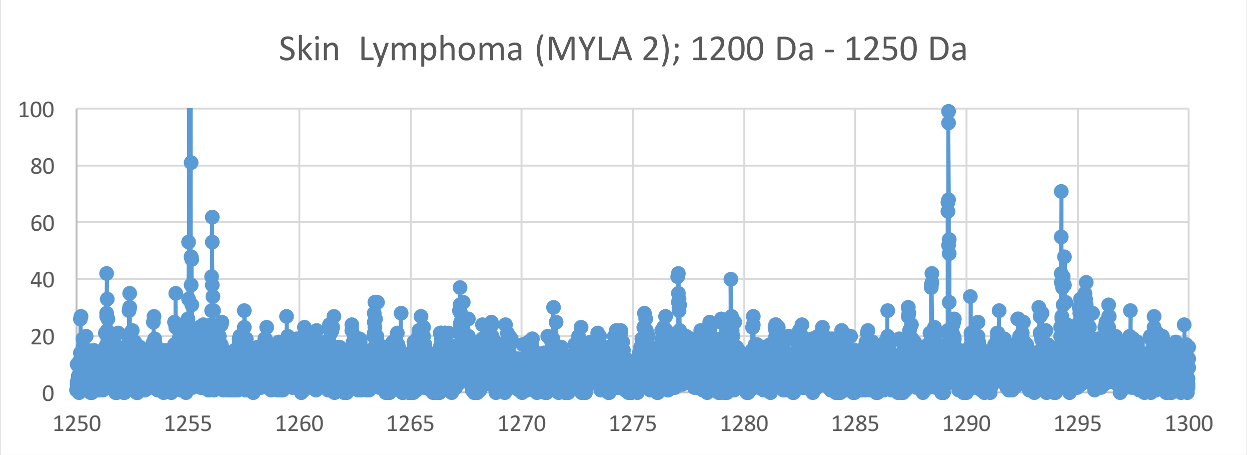
Skin Lymphoma (MYLA 2); 1200 Da - 1250 Da
100
80
60
40
20
0
1250
1255
1260
1265
1270
1275
1280
1285
1290
1295
1300

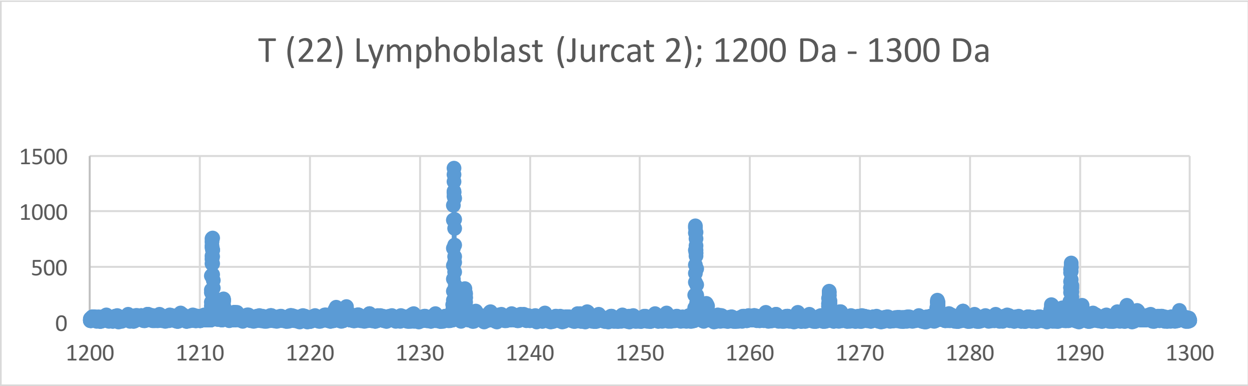
T (22) Lymphoblast (Jurcat 2); 1200 Da - 1300 Da
1500
1000
500
0
1200
1210
1220
1230
1240
1250
1260
1270
1280
1290
1300

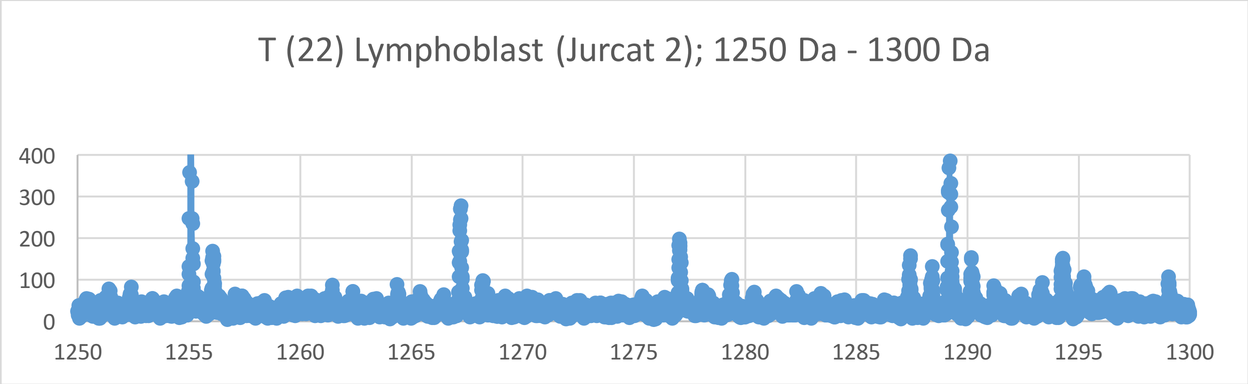
T (22) Lymphoblast (Jurcat 2); 1250 Da - 1300 Da
400
300
200
100
0
1250
1255
1260
1265
1270
1275
1280
1285
1290
1295
1300

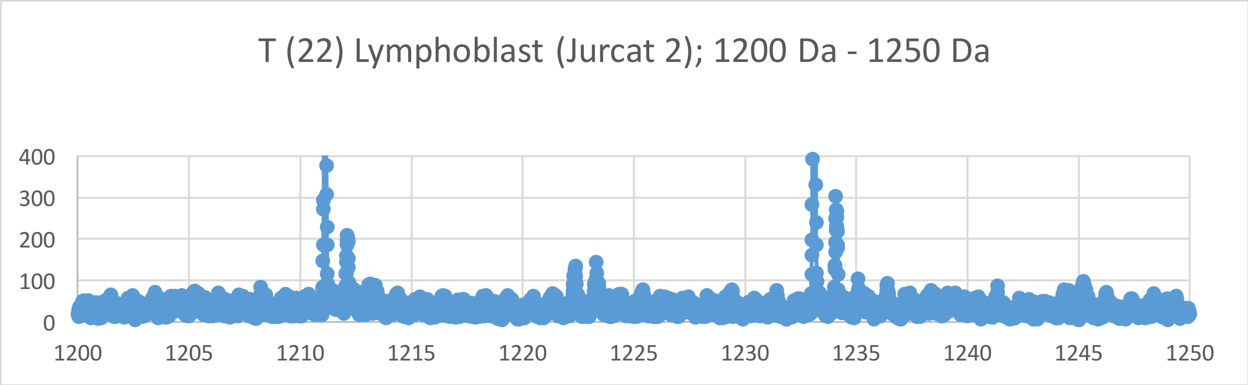
T (22) Lymphoblast (Jurcat 2); 1200 Da - 1250 Da
400
300
200
100
0
1200
1205
1210
1215
1220
1225
1230
1235
1240
1245
1250

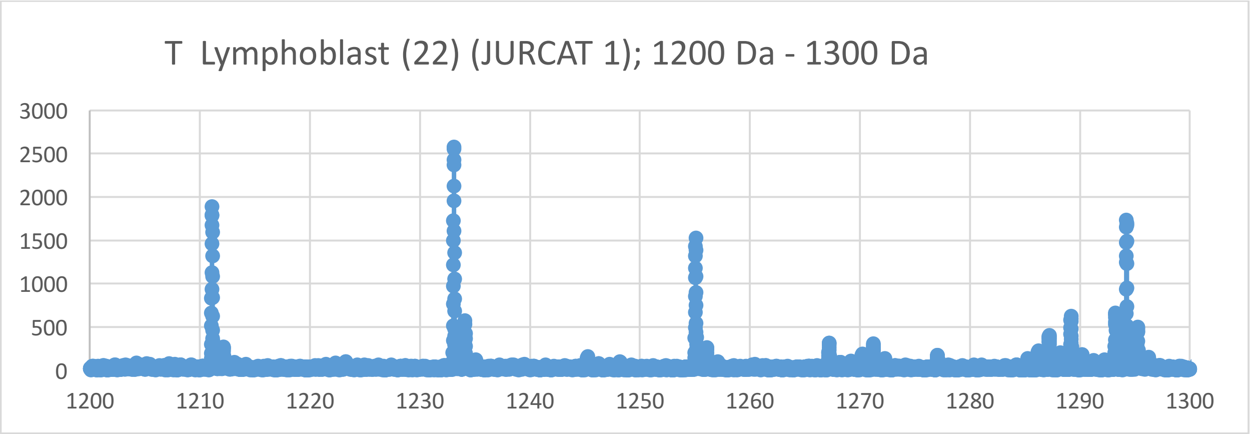
T Lymphoblast (22) (JURCAT 1); 1200 Da - 1300 Da
3000
2500
2000
1500
1000
500
0
1200
1210
1220
1230
1240
1250
1260
1270
1280
1290
1300

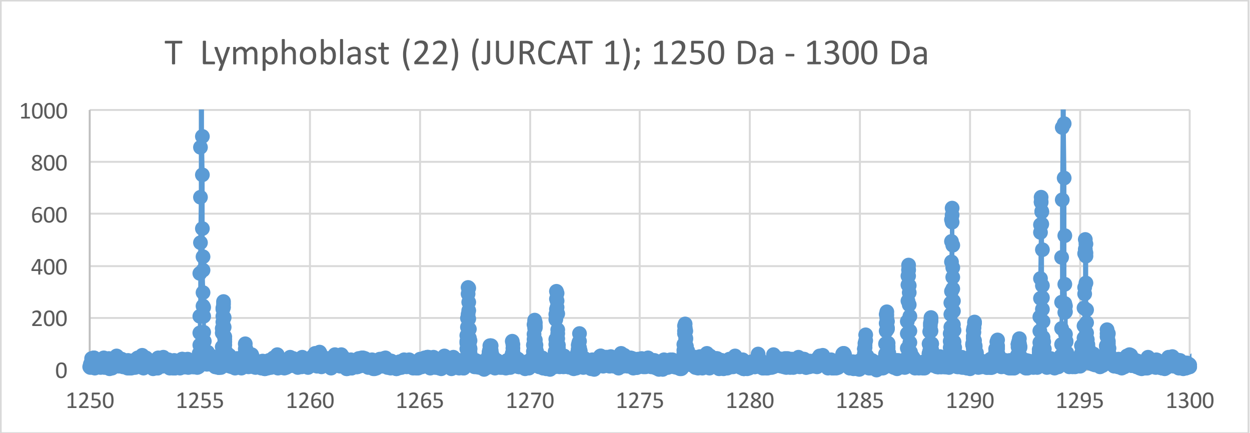
T Lymphoblast (22) (JURCAT 1); 1250 Da - 1300 Da
1000
800
600
400
200
0
1250
1255
1260
1265
1270
1275
1280
1285
1290
1295
1300

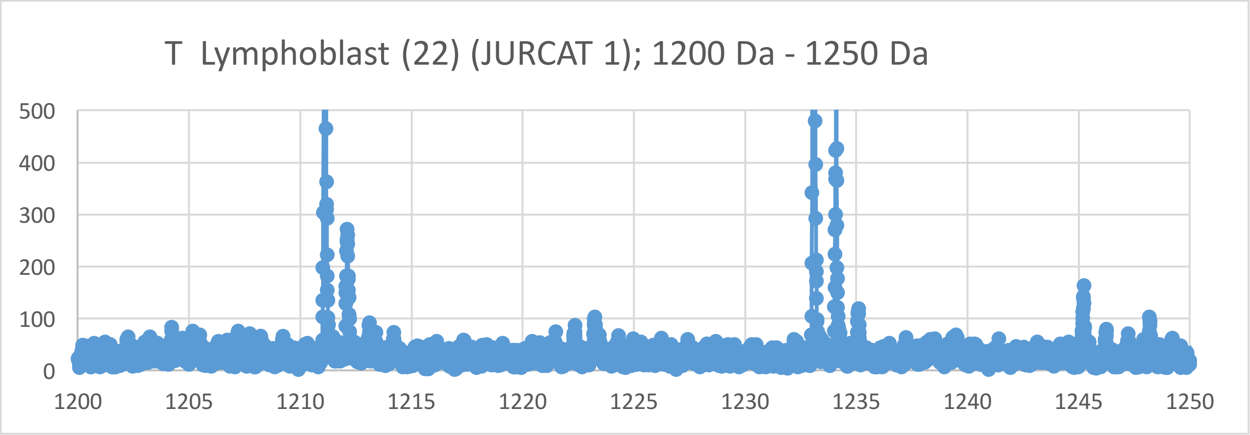
T Lymphoblast (22) (JURCAT 1); 1200 Da - 1250 Da
500
400
300
200
100
0
1200
1205
1210
1215
1220
1225
1230
1235
1240
1245
1250

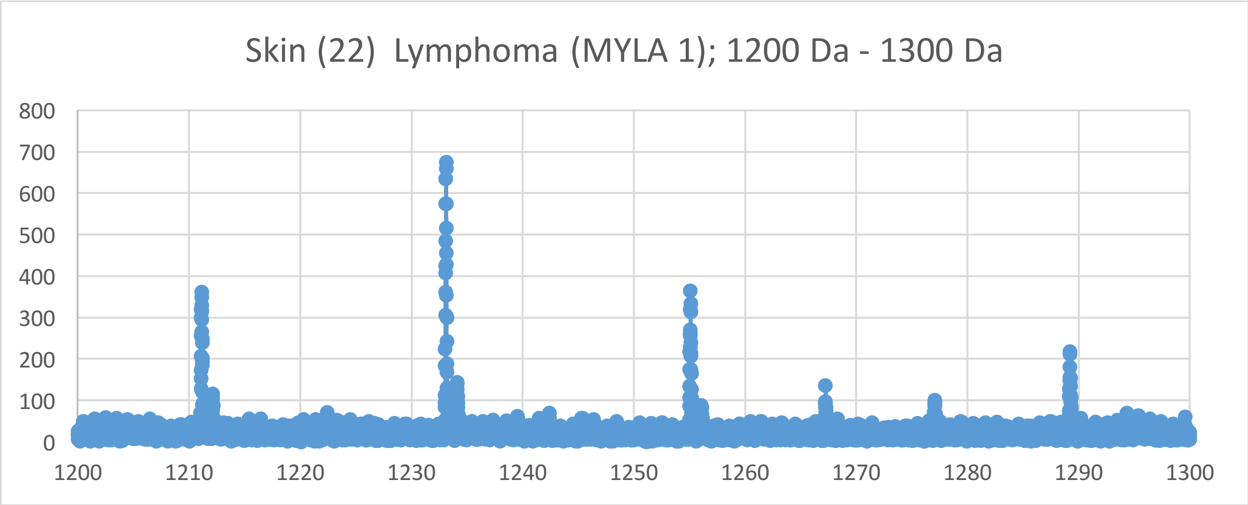
Skin (22) Lymphoma (MYLA 1); 1200 Da - 1300 Da
800
700
600
500
400
300
200
100
0
1200
1210
1220
1230
1240
1250
1260
1270
1280
1290
1300

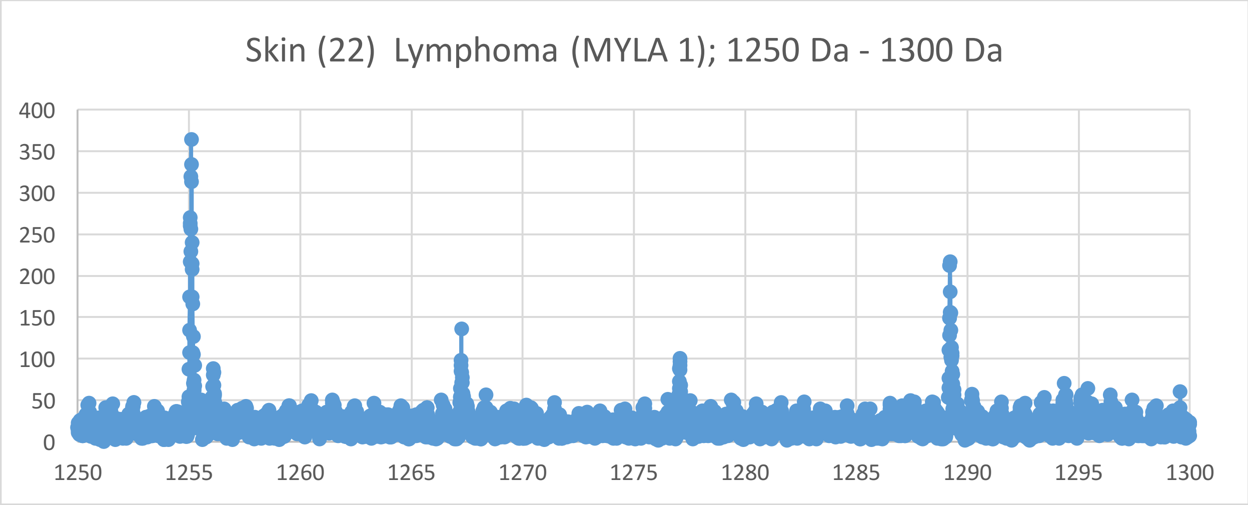
Skin (22) Lymphoma (MYLA 1); 1250 Da - 1300 Da
400
350
300
250
200
150
100
50
0
1250
1255
1260
1265
1270
1275
1280
1285
1290
1295
1300

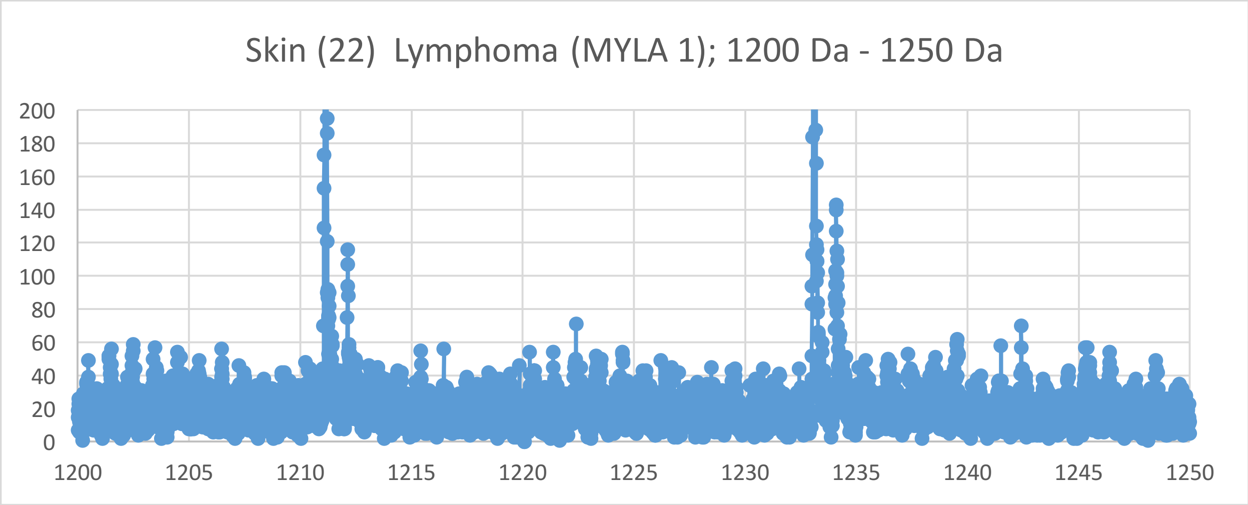
Skin (22) Lymphoma (MYLA 1); 1200 Da - 1250 Da
200
180
160
140
120
100
80
60
40
20
0
1200
1205
1210
1215
1220
1225
1230
1235
1240
1245
1250

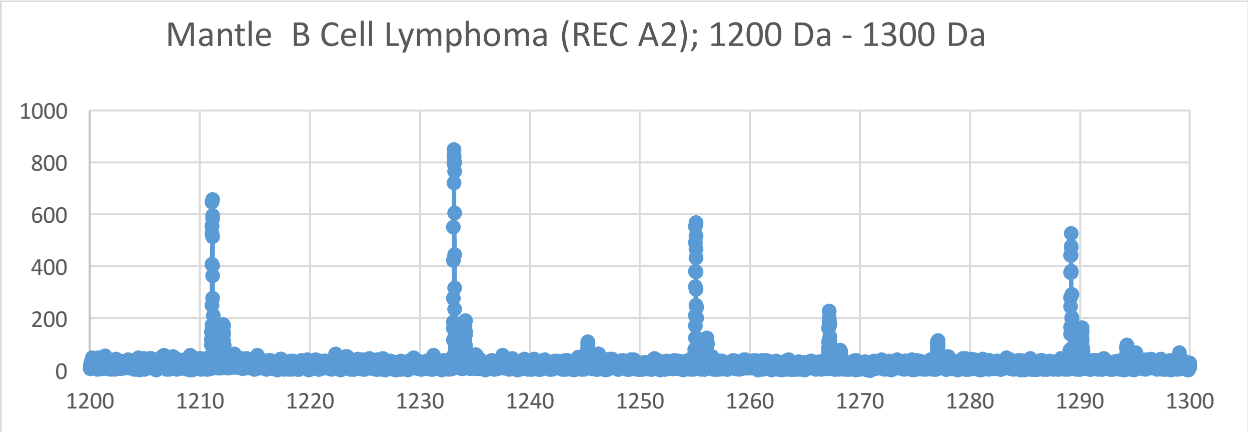
Mantle B Cell Lymphoma (REC A2); 1200 Da - 1300 Da
1000
800
600
400
200
0
1200
1210
1220
1230
1240
1250
1260
1270
1280
1290
1300

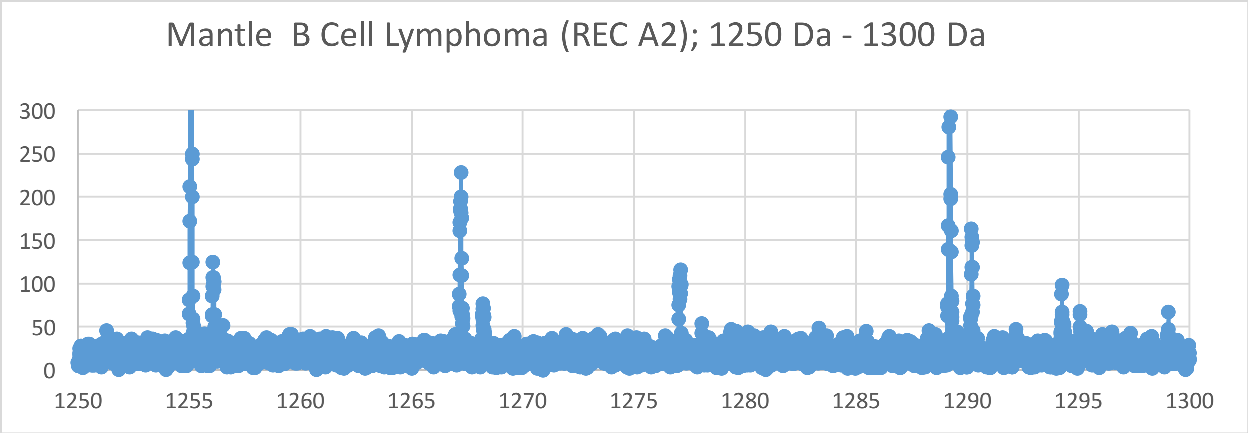
Mantle B Cell Lymphoma (REC A2); 1250 Da - 1300 Da
300
250
200
150
100
50
0
1250
1255
1260
1265
1270
1275
1280
1285
1290
1295
1300

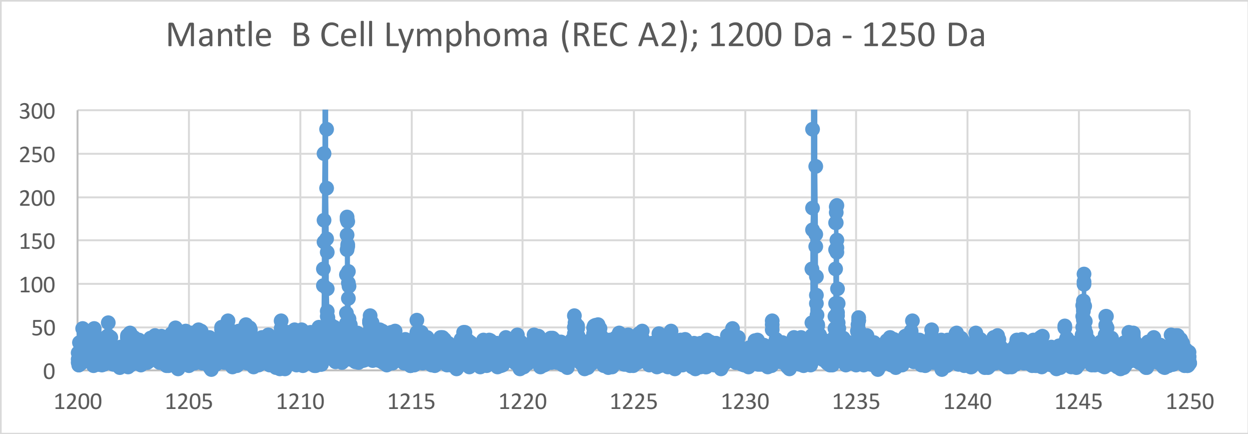
Mantle B Cell Lymphoma (REC A2); 1200 Da - 1250 Da
300
250
200
150
100
50
0
1200
1205
1210
1215
1220
1225
1230
1235
1240
1245
1250

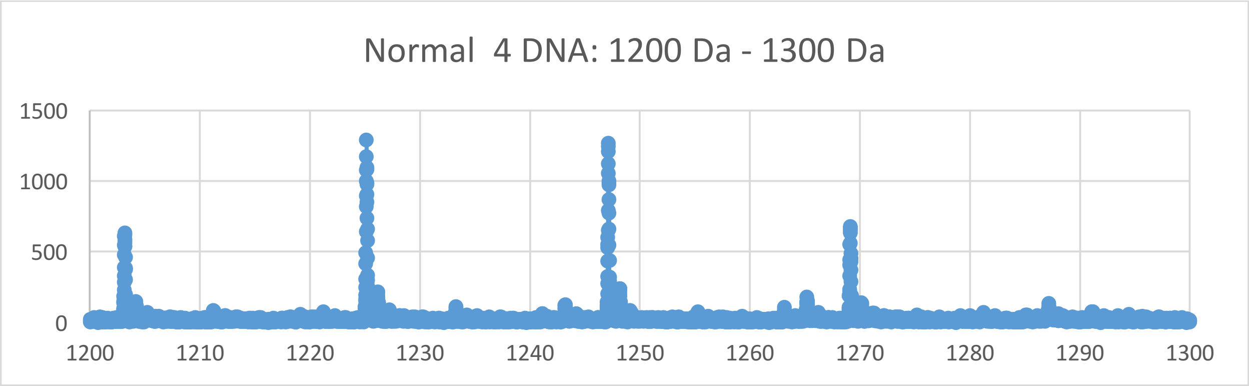
Normal 4 DNA: 1200 Da - 1300 Da
1500
1000
500
0
1200
1210
1220
1230
1240
1250
1260
1270
1280
1290
1300

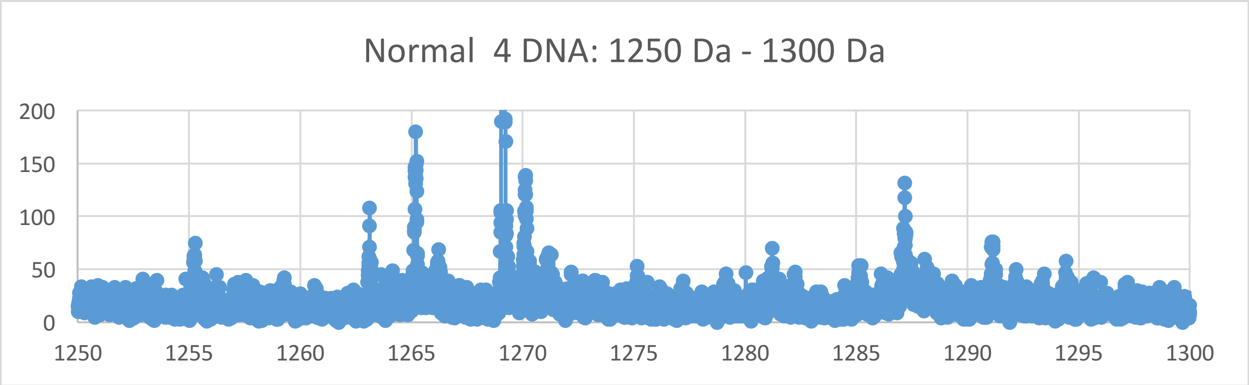
Normal 4 DNA: 1250 Da - 1300 Da
200
150
100
50
0
1250
1255
1260
1265
1270
1275
1280
1285
1290
1295
1300

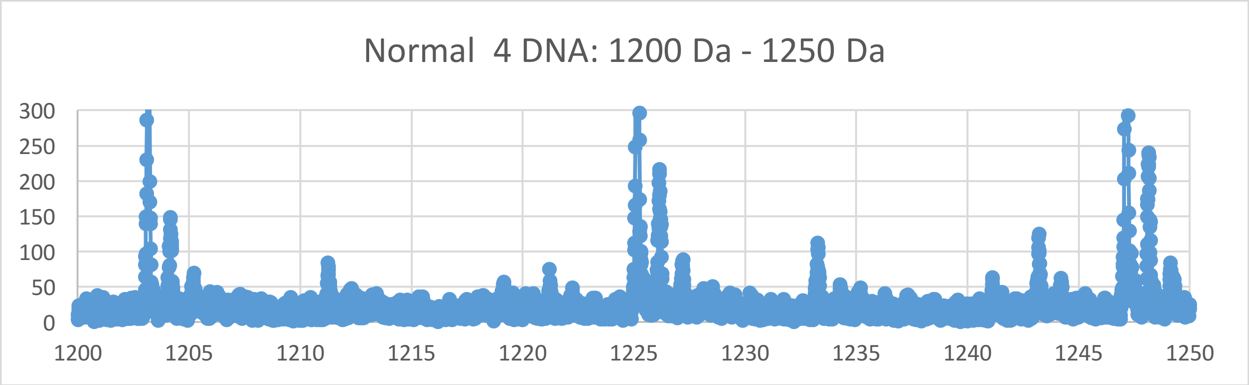
Normal 4 DNA: 1200 Da - 1250 Da
300
250
200
150
100
50
0
1200
1205
1210
1215
1220
1225
1230
1235
1240
1245
1250

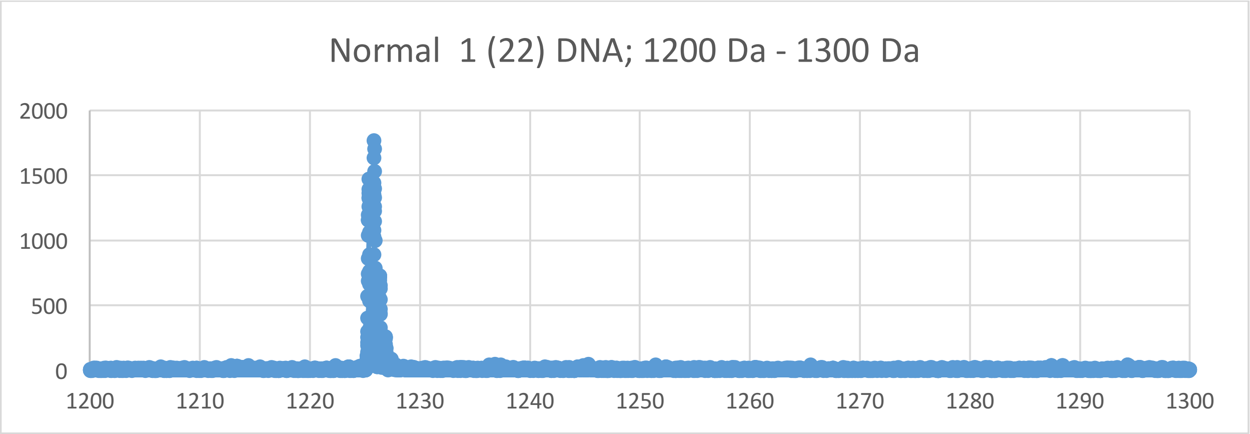
Normal 1 (22) DNA; 1200 Da - 1300 Da
2000
1500
1000
500
0
1200
1210
1220
1230
1240
1250
1260
1270
1280
1290
1300

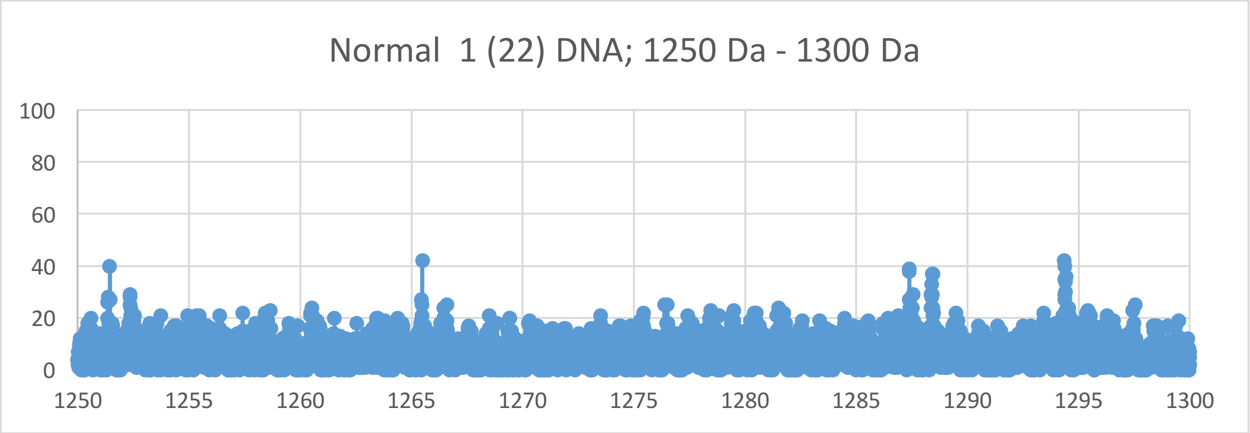
Normal 1 (22) DNA; 1250 Da - 1300 Da
100
80
60
40
20
0
1250
1255
1260
1265
1270
1275
1280
1285
1290
1295
1300

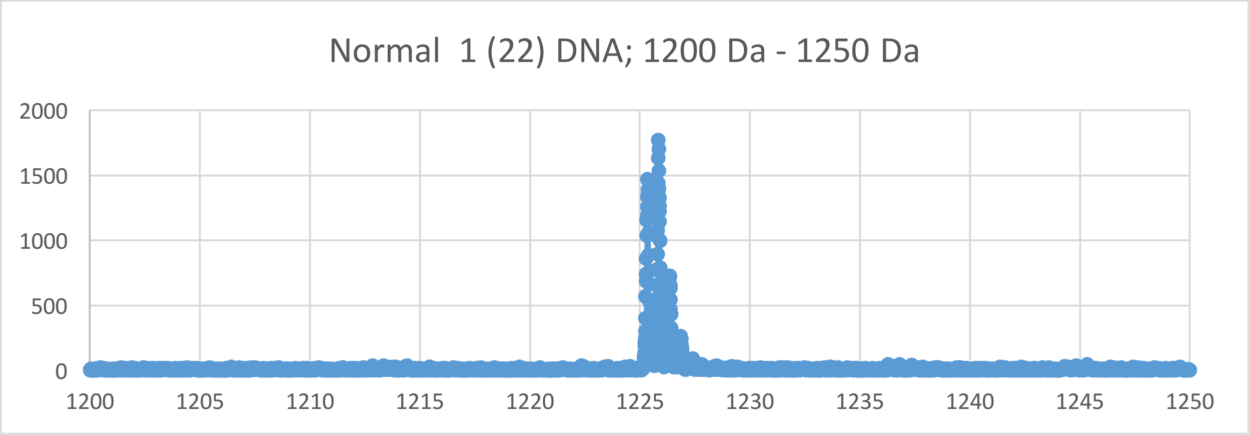
Normal 1 (22) DNA; 1200 Da - 1250 Da
2000
1500
1000
500
0
1200
1205
1210
1215
1220
1225
1230
1235
1240
1245
1250

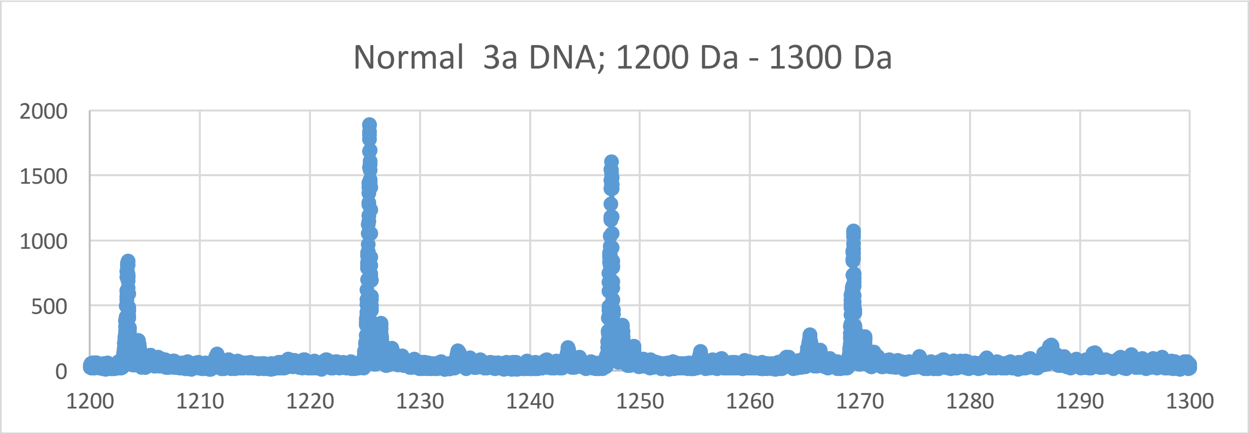
Normal 3a DNA; 1200 Da - 1300 Da
2000
1500
1000
500
0
1200
1210
1220
1230
1240
1250
1260
1270
1280
1290
1300

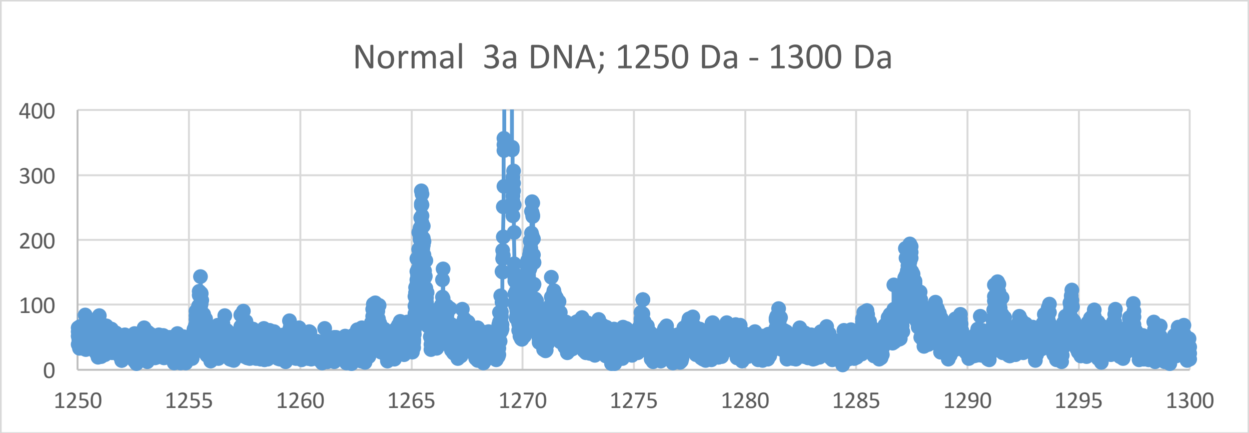
Normal 3a DNA; 1250 Da - 1300 Da
400
300
200
100
0
1250
1255
1260
1265
1270
1275
1280
1285
1290
1295
1300

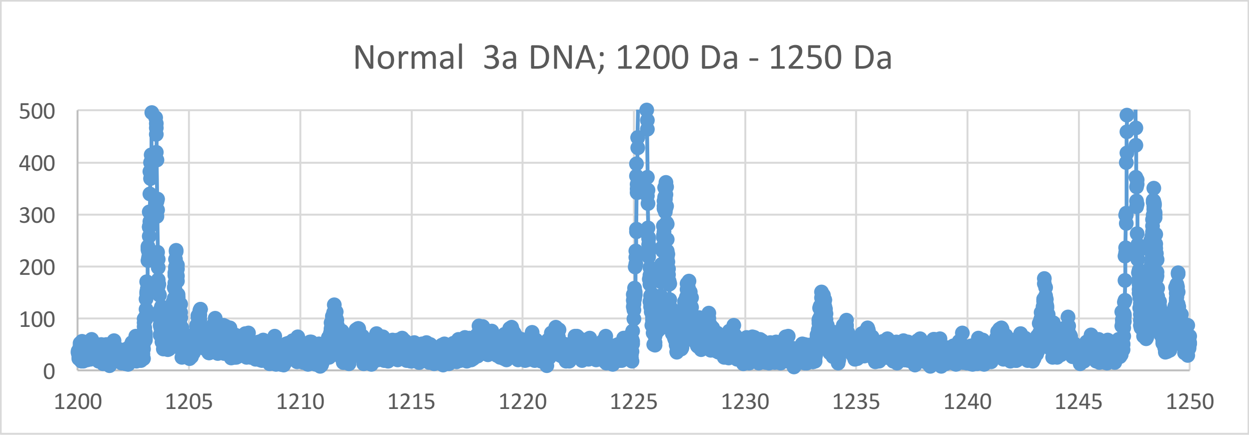
Normal 3a DNA; 1200 Da - 1250 Da
500
400
300
200
100
0
1200
1205
1210
1215
1220
1225
1230
1235
1240
1245
1250

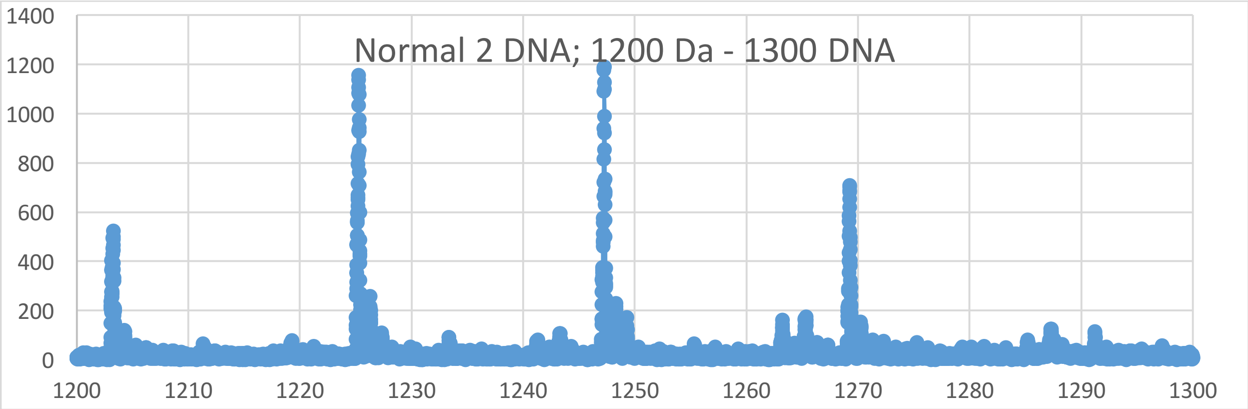
Normal 2 DNA; 1200 Da - 1300 DNA
0
200
400
600
800
1000
1200
1400
1200
1210
1220
1230
1240
1250
1260
1270
1280
1290
1300

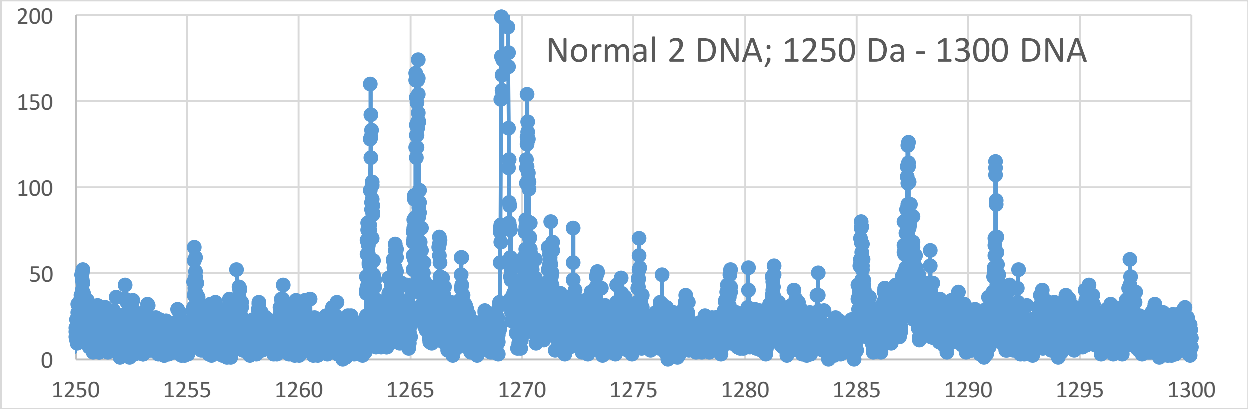
Normal 2 DNA; 1250 Da - 1300 DNA
0
50
100
150
200
1250
1255
1260
1265
1270
1275
1280
1285
1290
1295
1300

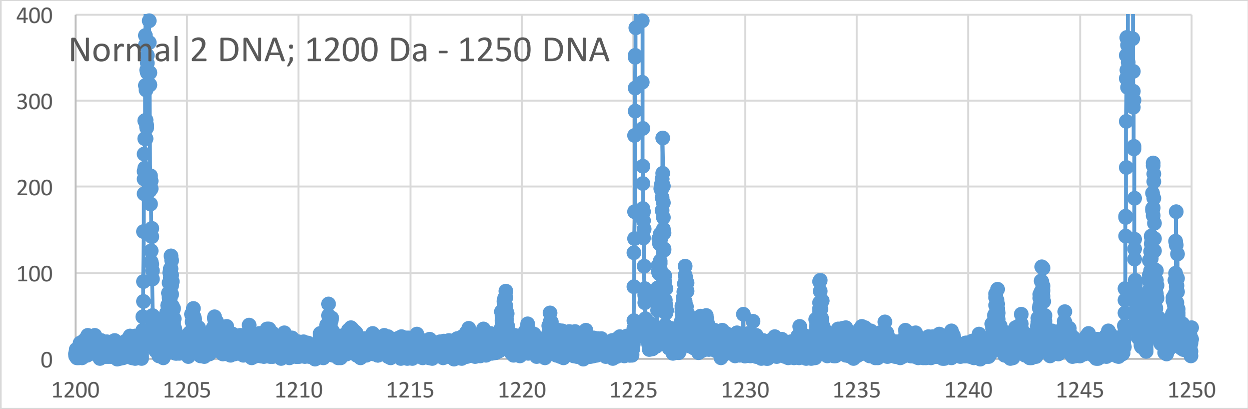
Normal 2 DNA; 1200 Da - 1250 DNA
0
100
200
300
400
1200
1205
1210
1215
1220
1225
1230
1235
1240
1245
1250
