## Supplementary Materials VIII for "Oncogenesis and Aging by Isotopic Functionalizations of the Proteins and Nucleic Acids"

VIII Supplementary Materials – New Mass Spectra from 1000 Da -1200 Da

Results for Supplementary 8

So for the trinucleotides at $(phos)_5$, the telomeric and nontelomeric peaks for normal strongly increase in intensities and number and the peaks for cancer decrease in number and intensities for stabilities of normal telomeric and nontelomeric trinucleotides. But the mating telomeres have increase in numbers, intensities and stabilities for cancer and normal trinucleotides at $(phos)_5$. But as decrease the phosphate by 1 unit, the trinucleotides change so that the normal trinucleotides less fragment and break from the DNA with less number and stabilities for the telomeres and nontelomeres, with less fragmentings and stabilities of the normal from DNA for telomeres and nontelomeres from $(phos)_5$ to $(phos)_4$. The masses range from 1000 Da to 1200 Da. See Supplementary Figures VIII for graphs. It seems that below some number of phosphates, the telomeres, mating telomeres and nontelomers behave in same way toward cancer and normal. So this suggests normal may be favored and distinguished from cancer by changing phosphorylations. The telomeric peaks for normal decrease in number with these dephosphorylations of the trinucleotides. The dephosphorylations from $(phos)_5$ to $(phos)_4$ caused the resulting less intense and stable normal trinucleotides to be less for telomeric and nontelomeric trinucleotides with negligible change in isotopic enrichments and the dephosphorylations caused even more cancer stabilities for mating telomeric trinucleotides and greater isotopically enrichments for telomeres. It is important to consider that there are more intense peaks for the trinucleotides relative to the tetranucleotides for the saturated phosphate units. For the trinucleotides the intensities of both normal and cancer fragments increased from $(phos)_5$ to $(phos)_4$. The dephosphorylations from $(phos)_5$ to $(phos)_4$ increases isotopic enrichments of light isotopes of $^{15}N$ for telomeric and nontelomeric trinucleotides but not for mating telomeric trinucleotides. The dephosphorylations from $(phos)_5$ to $(phos)_4$ also caused the telomeric and nontelomeric trinucleotides to be more intense in cancer DNA and more isotopically enriched, but mating telomeres change to be less intense in the normal and less isotopically enriched from $(phos)_5$ to $(phos)_4$. It seems the dephosphorylations are related to the isotopic enrichments and depletions; but the telomeres and nontelomeres are slightly more numerous in cancer and more isotopically enriched in light isotopes, whereas the frequencies of the mating telomeres is more numerous in cancer and the less isotopic enriched from $(phos)_5$ to $(phos)_4$. In general, it seems with less phosphorylations there are changes in isotopic enrichments and the difference between telomeric, nontelomeric and mating telomeric oligonucleotide bindings / stabilities with diminishment to induce cancer by such chemical mutations! The telomeric, mating telomeric and nontelomeric trinucleotides of normal tended to also be less enriched with heavy $^{13}C$ and $^{17}O$ and more enriched with light $^{15}N$ so it seems the cancer stabilities of the telomeric, mating telomeric and nontelomeric trinucleotides are related to their less isotopic enrichments with $^{13}C$ and $^{17}O$ but more enrichments with $^{15}N$ for the dephosphorylations from $(phos)_5$ to $(phos)_4$ and not purely driven by dephosphorylations but both dephosphorylations and enrichments. Enrichments strengthen the cancerous telomeric and nontelomeric normal trinucleotides while strongly weakening normal mating telomeric trinucleotides. The more intense cancerous nontelomeric and telomeric trinucleotides were also less isotopically enriched with heavy $^{13}C$ and $^{17}O$ but more isotopically enriched with $^{15}N$. The dephosphorylations from $(phos)_5$ to $(phos)_4$ strongly increased the $^{15}N$ isotopic enrichments of the cancer for the nontelomeric and telomeric trinucleotides and cancer for mating telomeres had weaker enrichments of $^{13}C$, $^{17}O$ nonprimordials. But the mating telomeric trinucleotides were even more enriched from the dephosphorylations for greater enrichments and stronger more stable cancer trinucleotides for mating telomeres. Such seems compatible for replications and transcriptions as stable telomeres seem compatible to unstable mating telomeres and nontelomeres for the unravelings and constructions of DNA mating telomeres onto the building templates of the telomeres. The dephosphorylations from $(phos)_5$ to $(phos)_4$ caused methylations to cause slightly increase normal intensities and numbers of peaks telomeres with negligible changes in isotopic enrichments. The methylations caused negligible change in intensities and stabilities of nontelomeric trinucleotides with negligible change in enrichments and the methylations caused increased intensities of both normal and cancer for mating telomeric trinucleotides with increase in enrichments. Such loss of differing behavior of telomeres and nontelomers with greater dephosphorylations points to malfunctions of DNA by highly dephosphorylated nucleotides and plays a role in cancer. The methylations of the telomeres does slightly increased normal intensities and stabilities with less enrichments from $(phos)_5$ to $(phos)_4$ relative to the nontelomeres. It seems from this data that during biological processes the dephosphorylations and stabilizations of normal or vice versa dephosphorylations and stabilizations of cancer are or can be countered by methylations and destabilizations of the normal for cancer or vice versa dephosphorylations and destabilizations of the cancer by methylations as this research discovers coupled phosphorylations and methylations or coupled dephosphorylations and methylations with involved greater enrichments for normal biological functions and explaining cancer! It seems that if nucleotides loose too many phosphates in regulating methylations and acteylations that the differences among genes and in particular telomere genes and nontelomere genes are lost and the activity of cancer arises! The dephosphorylations from $(phos)_5$ to $(phos)_4$ caused acetylations to cause increased normal stabilities for telomeres with increased enrichments for telomeres and nontelomeres. The acetylations from $(phos)_5$ to $(phos)_4$ caused cancer stabilities and intensities for mating telomeres with negligible change in enrichments of light and heavy isotopes. This contrast in mating telomeres and telomeres from the acetylations may reflect the developments of cancer as the telomeres fragment but the mating telomeres more holds together for damaged DNA for cancer.

So for the trinucleotides at $(phos)_4$, the telomeric (and mating telomeric) peaks for cancer strongly increase in intensities and number for trinucleotides. But the nontelomeres have increase in numbers and stabilities for normal trinucleotides while the cancer trinucleotides are mostly less in peaks and stabilities for normal trinucleotides with $(phos)_4$. But as decrease the phosphate by 1 unit, the trinucleotides change so that the cancer trinucleotides more fragment and break from the DNA with more intensities and stabilities for the telomeres and mating telomeres, with slightly more fragmenting and stabilities of the cancer from DNA for telomeres and mating telomeres from $(phos)_4$ to $(phos)_3$. The masses range from 1000 Da to 1200 Da. See Supplementary Figures 8 for graphs. It seems that below some number of phosphates, the telomeres, mating telomeres and nontelomers behave in same way toward cancer and normal. So this suggest normal may be favored and distinguished from cancer by changing phosphorylations. The telomeric and mating telomeric peaks for cancer increase in number and intensities with these dephosphorylations of the trinucleotides. The dephosphorylations from $(phos)_4$ to $(phos)_3$ caused the resulting more intense and stable cancer trinucleotides to be more for telomeric and mating telomeric trinucleotides with more isotopic enrichments of heavy $^{13}C$ and $^{17}O$ and the dephosphorylations caused even more normal stabilities for nontelomeric trinucleotides and greater isotopic enrichments of $^{13}C$ and $^{17}O$ for nontelomeres. It is important to consider that there are more intense peaks for the trinucleotides relative to the tetranucleotides for the saturated phosphate units and for some reason for large dephosphorylations the intensities increase from some intermediate drops for medium phosphorylations. For the trinucleotides, the intensities of both normal and cancer fragments increased from $(phos)_4$ to $(phos)_{3.}$ The dephosphorylations from $(phos)_4$ to $(phos)_3$ increase isotopic enrichments of light isotopes of $^{15}N$ for nontelomeric trinucleotides but not for mating telomeric trinucleotides. The dephosphorylations from $(phos)_4$ to $(phos)_3$ also caused the telomeric and mating telomeric trinucleotides to be more intense in cancer DNA and more isotopically enriched, but mating telomeres tend to enrich in heavier isotopes but cancer telomeres tend to enrich in light isotopes from $(phos)_5$ to $(phos)_4$. This is unusual as cancer fragments are usually enriched in heavy isotopes of $^{13}C$ and $^{17}O$ but for trinucleotide fragments here the cancer is enriched in $^{15}N$ lighter isotopes and this is consistent with less clumping in oligonucleotides of fewer nucleotides so the $^{14}N$ and $^{15}N$ play larger role in dynamics of smaller oligonucleotides. It seems the dephosphorylations are related to the isotopic enrichments and depletions; but the telomeres and mating telomeres are slightly more numerous in cancer and more isotopically enriched in light isotopes (for telomeres) and heavy (for mating telomers), whereas the frequencies of the nontelomeres are more numerous in normal and the more isotopically enriched in $^{13}C$ and $^{17}O$ from $(phos)_4$ to $(phos)_3$. In general, it seems with less phosphorylations there is change in isotopic enrichments and the difference between telomeric, nontelomeric and mating telomeric binding / stability diminishes to induce cancer by such chemical mutations! The mating telomeric trinucleotides of cancer tended to also be less enriched with heavy $^{13}C$ and $^{17}O$ and more enriched with light $^{15}N$ so it seems the cancer stabilities of the mating telomeric trinucleotides are related to their less isotopic enrichments with $^{13}C$ and $^{17}O$ but more enrichments with $^{15}N$ for the dephosphorylations from $(phos)_4$ to $(phos)_3$ and not purely driven by dephosphorylations but both dephosphorylations and enrichments. Enrichments strengthen the cancerous telomeric and mating telomeric trinucleotides while strongly weakening cancer non- telomeric trinucleotides. The more intense cancerous telomeric trinucleotides were also less isotopically enriched with heavy $^{13}C$ and $^{17}O$ but more isotopically enriched with $^{15}N$. The dephosphorylations from $(phos)_4$ to $(phos)_3$ strongly increased the $^{15}N$ isotopic enrichments of the cancer for the telomeric trinucleotides and cancer for mating telomeres had greater enrichments of $^{13}C$, $^{17}O$ nonprimordials. The cancer and normal peaks for nontelomeric nucleotides were observed enriched with heavier isotopes. But the nontelomeric trinucleotides were even more enriched from the dephosphorylations for greater enrichments with heavier isotopes and stronger more stable cancer trinucleotides than for telomeres. Such seems compatible for replications and transcriptions as stable telomeres seem compatible to unstable mating telomeres and nontelomeres for the unravelings and constructions of DNA mating telomeres onto the building templates of the telomeres. The dephosphorylations from $(phos)_4$ to $(phos)_3$ caused methylations to cause increase cancer intensities and numbers of peaks telomeres with changes in isotopic enrichments. The methylations caused increase in normal intensities and stabilities of mating telomeric trinucleotides with reductions in enrichments and the methylations caused increased intensities of cancer for nontelomeric trinucleotides with reductions in enrichments. Such loss of differing behaviors of telomeres and nontelomers with greater dephosphorylations points to malfunctions of DNA by highly dephosphorylated nucleotides and plays a role in cancer. The methylations of the telomeres does slightly increased cancer intensities and stabilities with negligible changes in enrichments from $(phos)_4$ to $(phos)_3$ relative to the nontelomeres. It seems from this data that during biological processes the dephosphorylations and stabilizations of normal or vice versa dephosphorylations and stabilizations of cancer are or can be countered by methylations and destabilizations of the normal for cancer or vice versa dephosphorylations and destabilizations of the cancer by methylations as this research discovers coupled phosphorylations and methylations or coupled dephosphorylations and methylations with involved greater enrichments for normal biological functions and explaining cancer! It seems that if nucleotides lose too many phosphates in regulating methylations and acteylations that the differences among genes and in particular telomeric genes and nontelomeric genes is lost and the activity of cancer arises! The dephosphorylations from $(phos)_4$ to $(phos)_3$ caused acetylations to cause decrease in cancer stabilities for telomeres with decreased enrichments of heavier isotopes for telomeres. The acetylations caused increase stabilities, intensities and number of cancer peaks for mating telomeres with increase light isotopic enrichments. The acetylations from $(phos)_4$ to $(phos)_3$ caused slight normal stabilities and intensities for nontelomeres with decreased enrichments of heavy isotopes but enrichments of light isotopes. This contrast in mating telomeres and telomeres from the acetylations may reflect the developments of cancer as the telomeres fragment but the mating telomeres more holds together for damaged DNA for cancer.

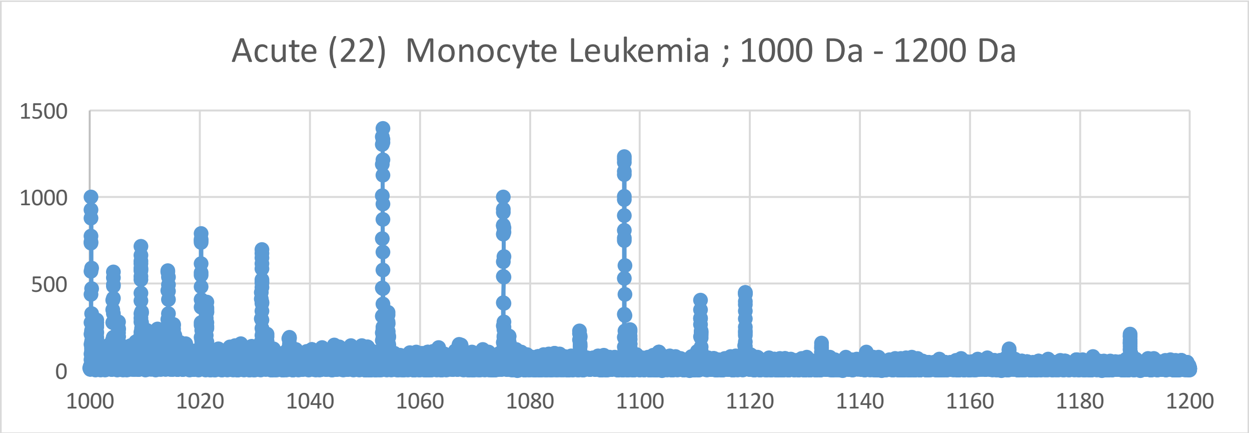
Acute (22) Monocyte Leukemia ; 1000 Da - 1200 Da
1500
1000
500
0
1000
1020
1040
1060
1080
1100
1120
1140
1160
1180
1200

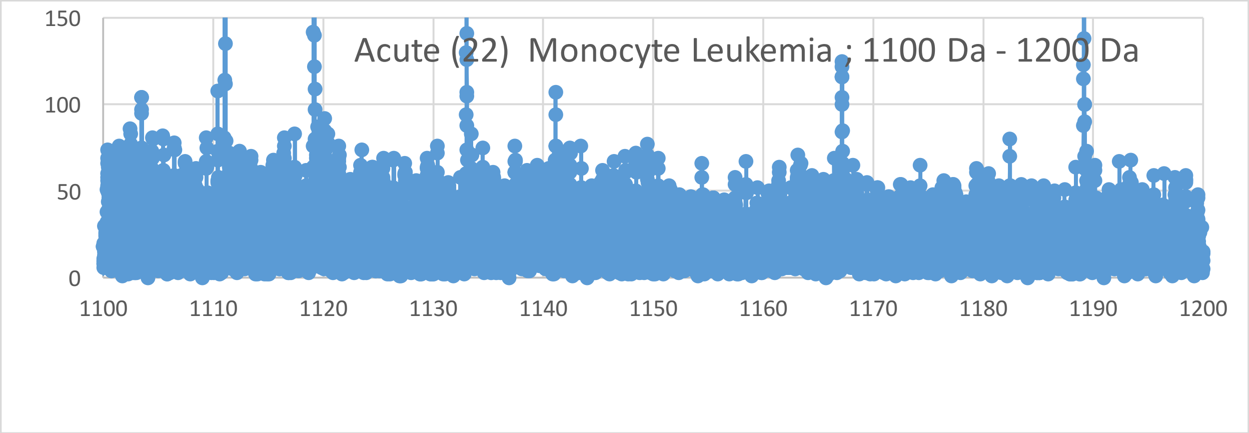
Acute (22) Monocyte Leukemia ; 1100 Da - 1200 Da
150
100
50
0
1100
1110
1120
1130
1140
1150
1160
1170
1180
1190
1200

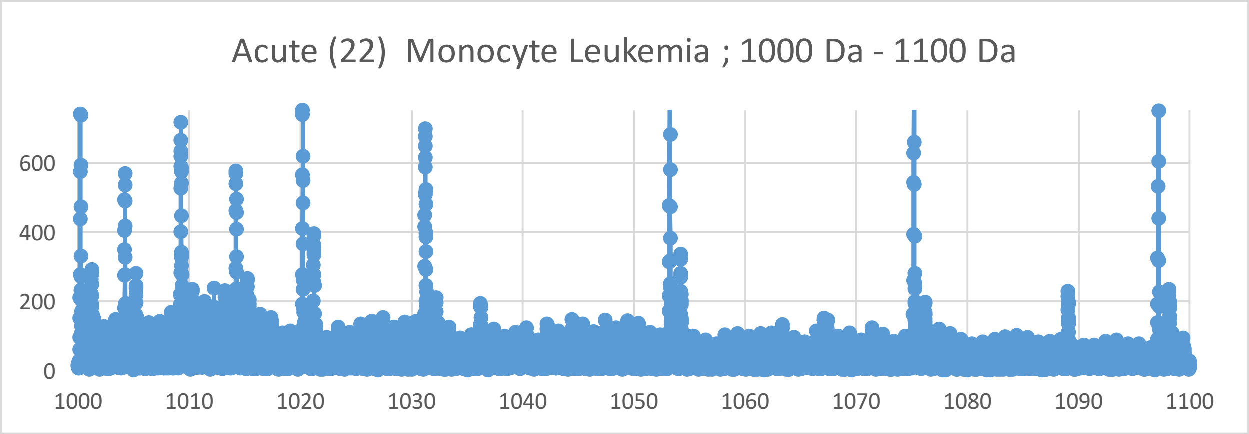
Acute (22) Monocyte Leukemia ; 1000 Da - 1100 Da
600
400
200
0
1000
1010
1020
1030
1040
1050
1060
1070
1080
1090
1100

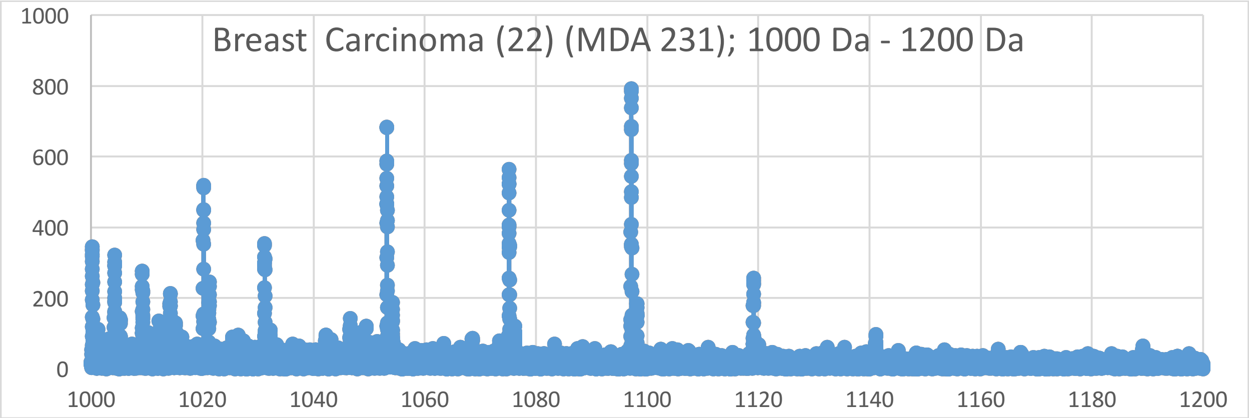
Breast Carcinoma (22) (MDA 231); 1000 Da - 1200 Da
1000
800
600
400
200
0
1000
1020
1040
1060
1080
1100
1120
1140
1160
1180
1200

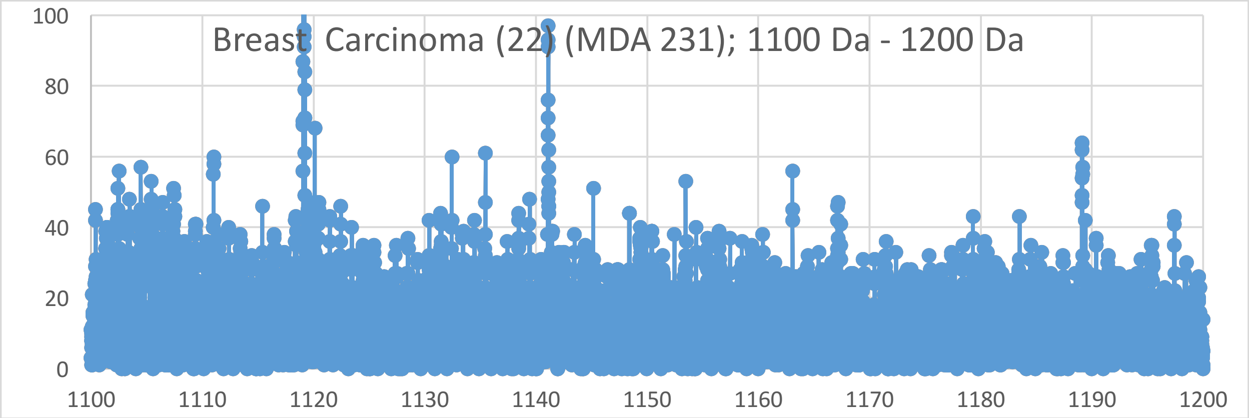
Breast Carcinoma (22) (MDA 231); 1100 Da - 1200 Da
100
80
60
40
20
0
1100
1110
1120
1130
1140
1150
1160
1170
1180
1190
1200

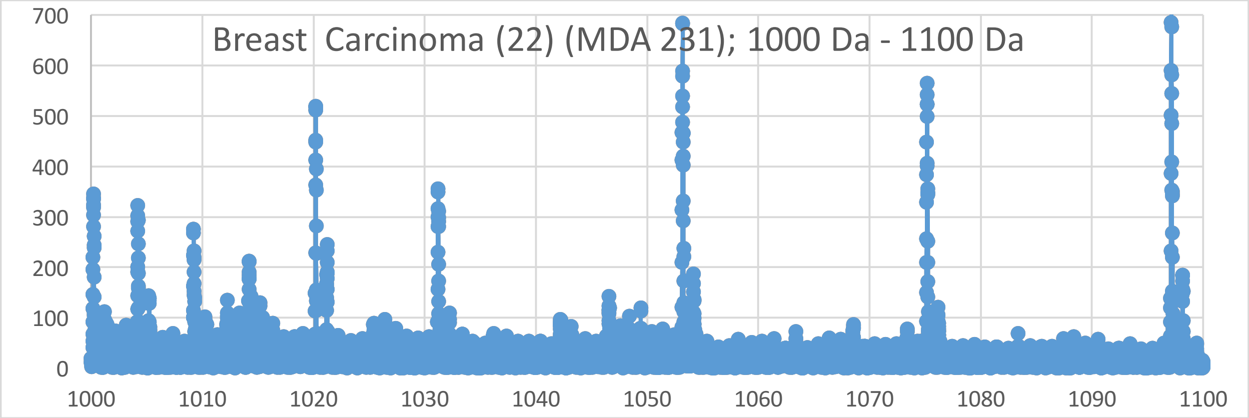
Breast Carcinoma (22) (MDA 231); 1000 Da - 1100 Da
700
600
500
400
300
200
100
0
1000
1010
1020
1030
1040
1050
1060
1070
1080
1090
1100

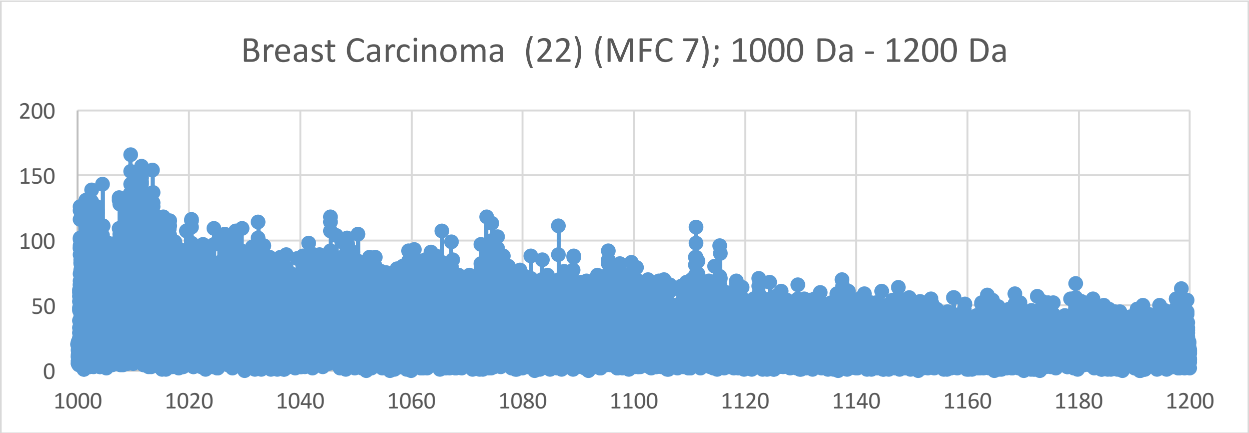

Breast Carcinoma (22) (MFC 7); 1000 Da - 1200 Da
0
50
100
150
200
1000
1020
1040
1060
1080
1100
1120
1140
1160
1180
1200


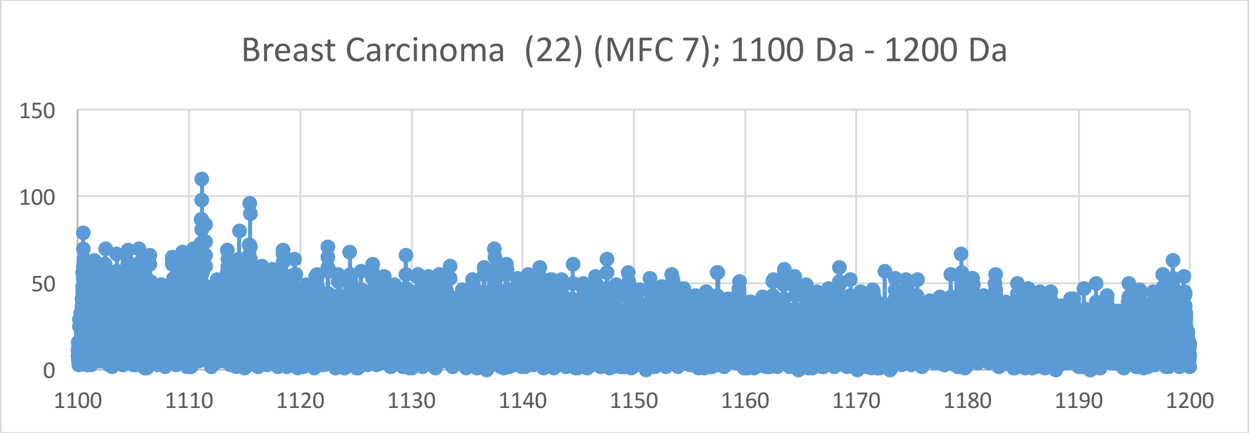

Breast Carcinoma (22) (MFC 7); 1100 Da - 1200 Da
0
50
100
150
1100
1110
1120
1130
1140
1150
1160
1170
1180
1190
1200


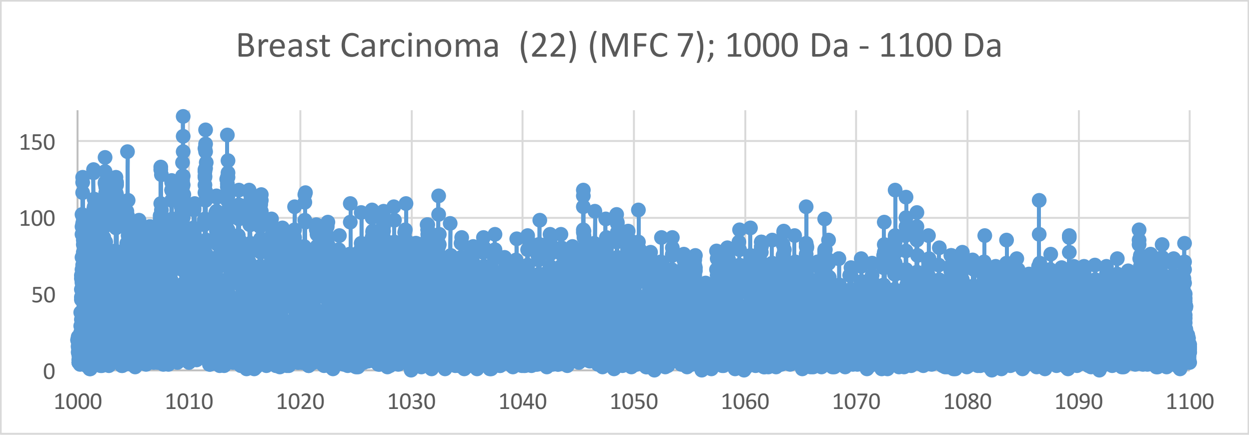

Breast Carcinoma (22) (MFC 7); 1000 Da - 1100 Da
0
50
100
150
1000
1010
1020
1030
1040
1050
1060
1070
1080
1090
1100

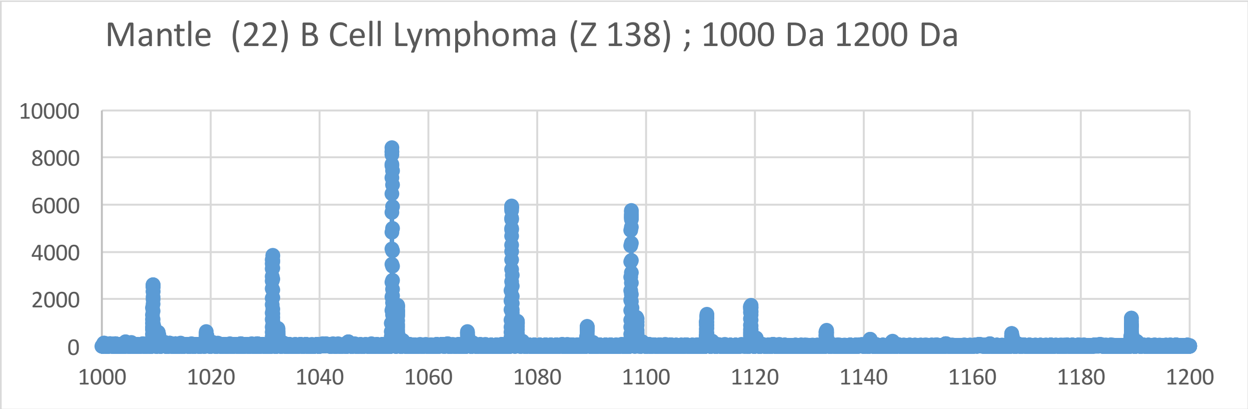
Mantle (22) B Cell Lymphoma (Z 138) ; 1000 Da 1200 Da
10000
8000
6000
4000
2000
0
1000
1020
1040
1060
1080
1100
1120
1140
1160
1180
1200

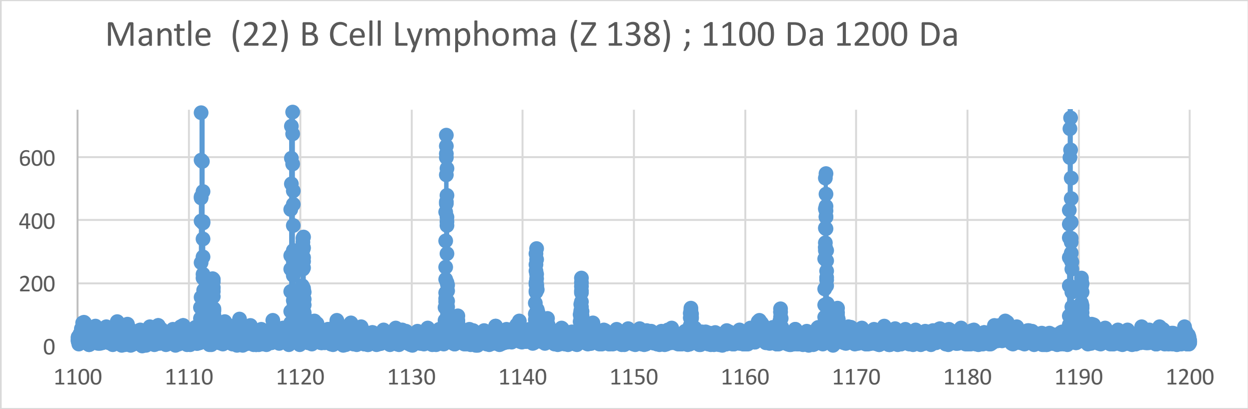
Mantle (22) B Cell Lymphoma (Z 138) ; 1100 Da 1200 Da
600
400
200
0
1100
1110
1120
1130
1140
1150
1160
1170
1180
1190
1200

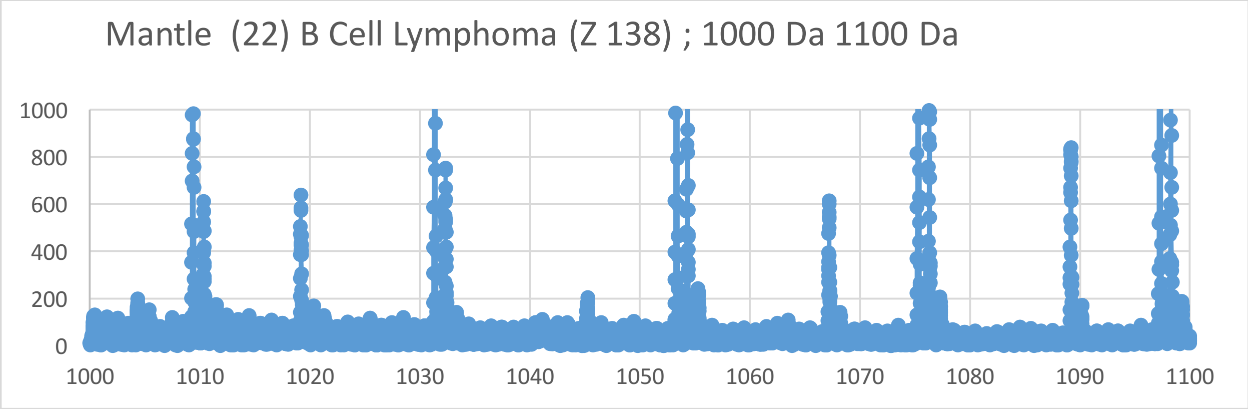
Mantle (22) B Cell Lymphoma (Z 138) ; 1000 Da 1100 Da
1000
800
600
400
200
0
1000
1010
1020
1030
1040
1050
1060
1070
1080
1090
1100

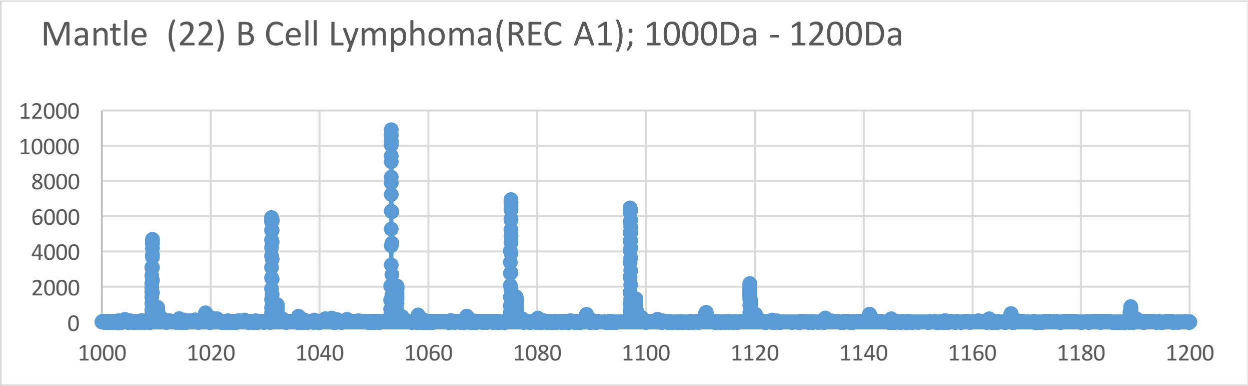
Mantle (22) B Cell Lymphoma(REC A1); 1000Da - 1200Da
12000
10000
8000
6000
4000
2000
0
1000
1020
1040
1060
1080
1100
1120
1140
1160
1180
1200

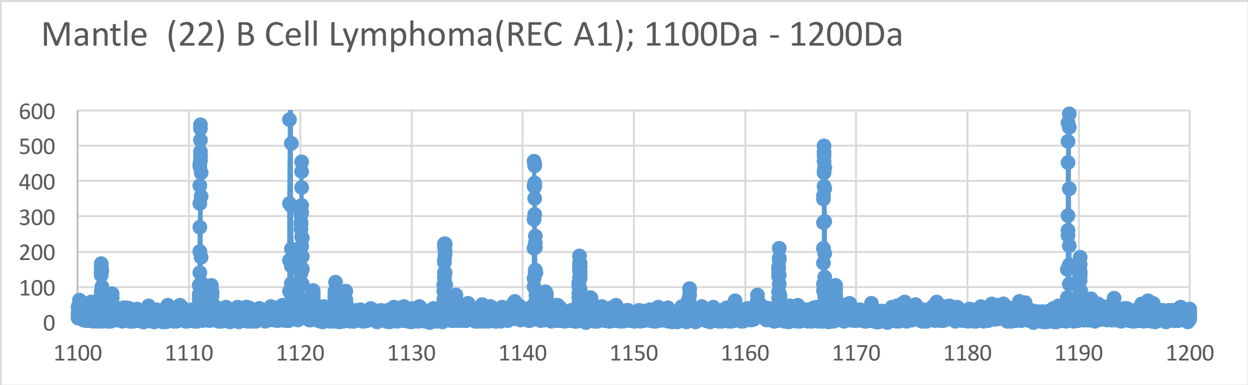
Mantle (22) B Cell Lymphoma(REC A1); 1100Da - 1200Da
600
500
400
300
200
100
0
1100
1110
1120
1130
1140
1150
1160
1170
1180
1190
1200

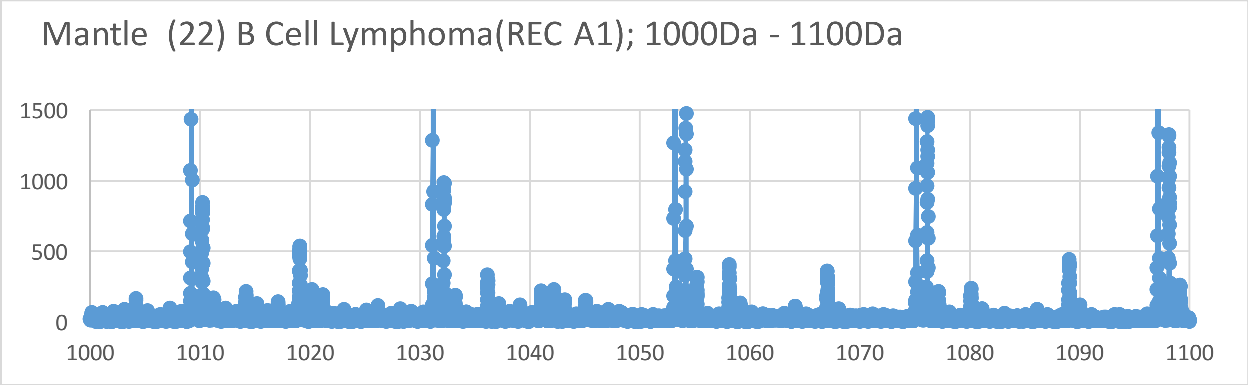
Mantle (22) B Cell Lymphoma(REC A1); 1000Da - 1100Da
1500
1000
500
0
1000
1010
1020
1030
1040
1050
1060
1070
1080
1090
1100

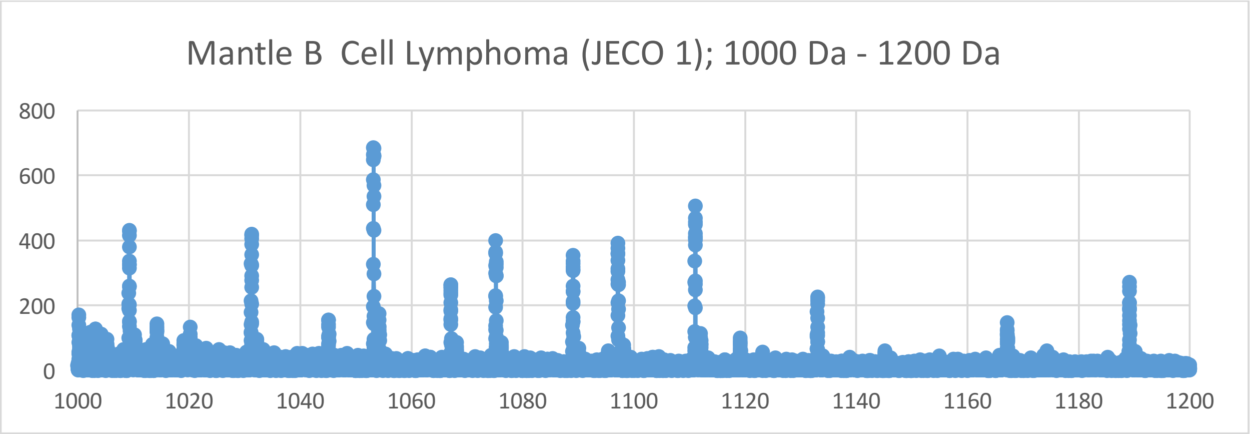
Mantle B Cell Lymphoma (JECO 1); 1000 Da - 1200 Da
800
600
400
200
0
1000
1020
1040
1060
1080
1100
1120
1140
1160
1180
1200

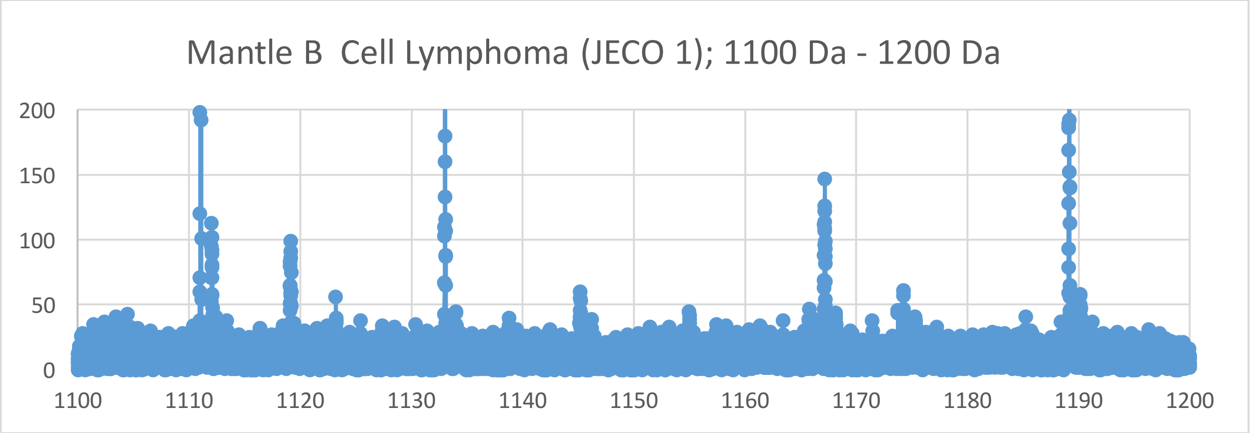
Mantle B Cell Lymphoma (JECO 1); 1100 Da - 1200 Da
200
150
100
50
0
1100
1110
1120
1130
1140
1150
1160
1170
1180
1190
1200

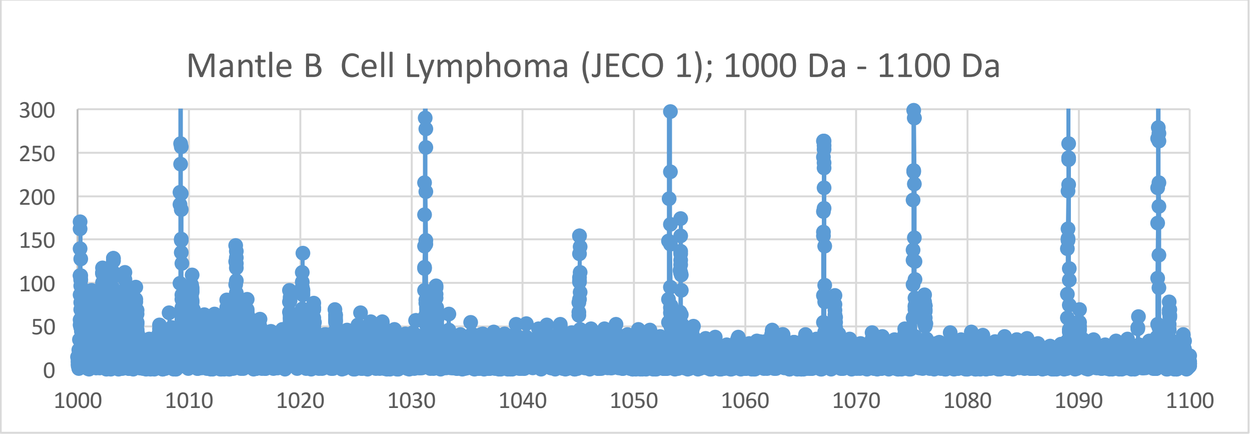
Mantle B Cell Lymphoma (JECO 1); 1000 Da - 1100 Da
300
250
200
150
100
50
0
1000
1010
1020
1030
1040
1050
1060
1070
1080
1090
1100

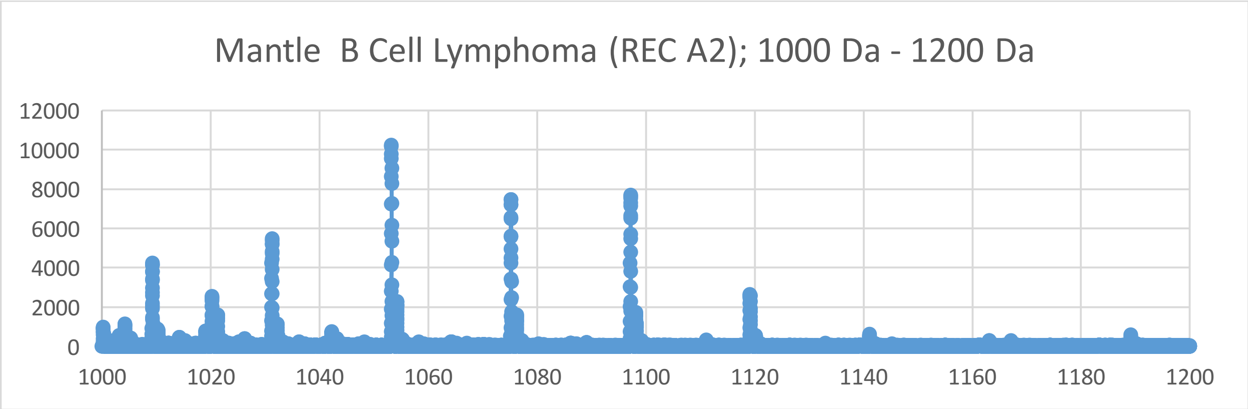
Mantle B Cell Lymphoma (REC A2); 1000 Da - 1200 Da
12000
10000
8000
6000
4000
2000
0
1000
1020
1040
1060
1080
1100
1120
1140
1160
1180
1200

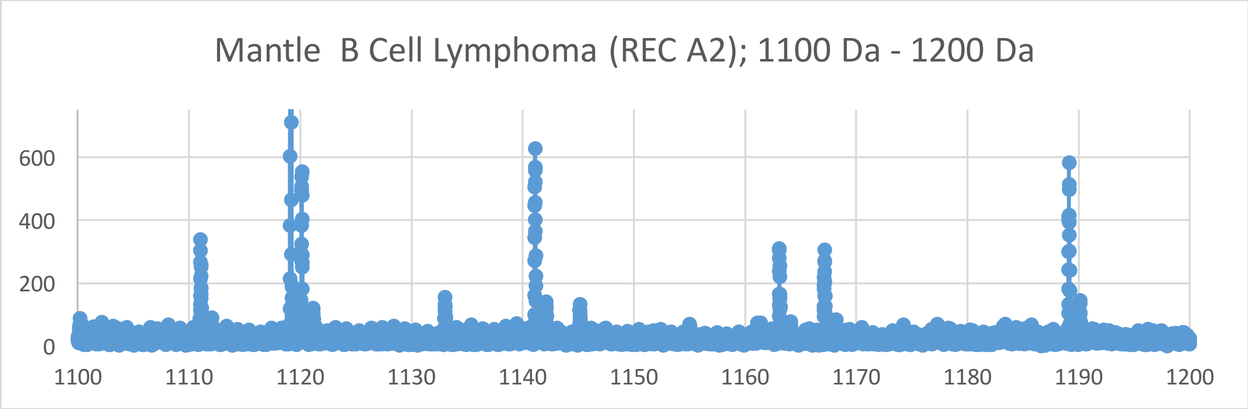
Mantle B Cell Lymphoma (REC A2); 1100 Da - 1200 Da
600
400
200
0
1100
1110
1120
1130
1140
1150
1160
1170
1180
1190
1200

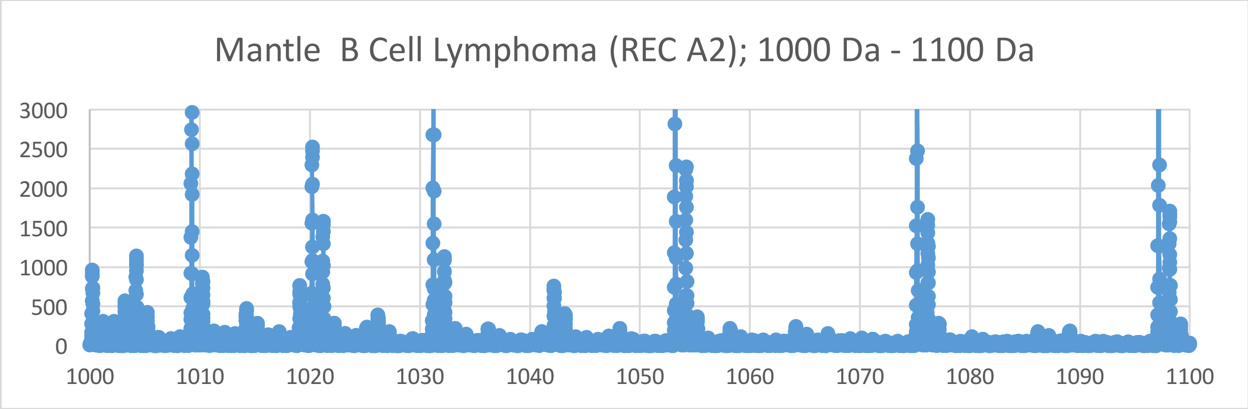
Mantle B Cell Lymphoma (REC A2); 1000 Da - 1100 Da
3000
2500
2000
1500
1000
500
0
1000
1010
1020
1030
1040
1050
1060
1070
1080
1090
1100

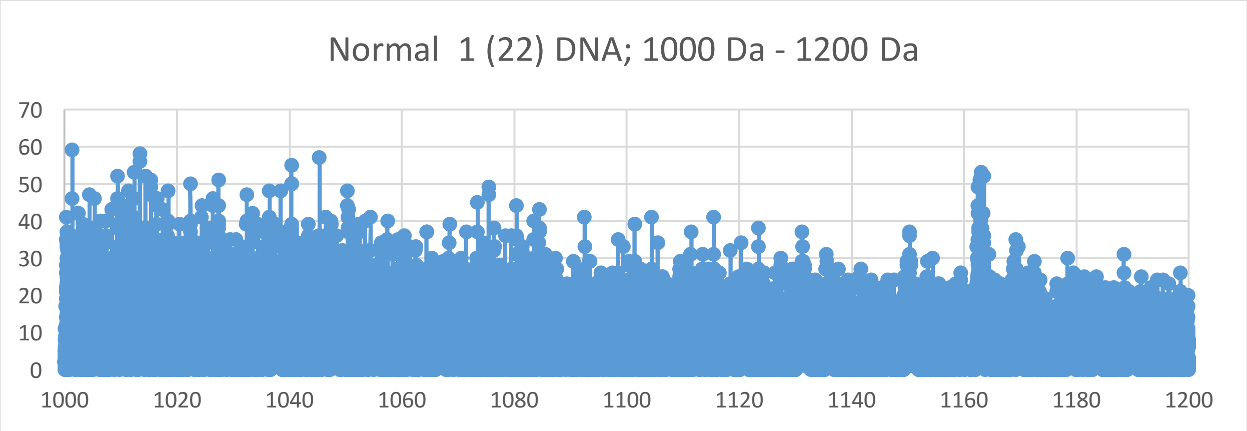
Normal 1 (22) DNA; 1000 Da - 1200 Da
0
10
20
30
40
50
60
70
1000
1020
1040
1060
1080
1100
1120
1140
1160
1180
1200

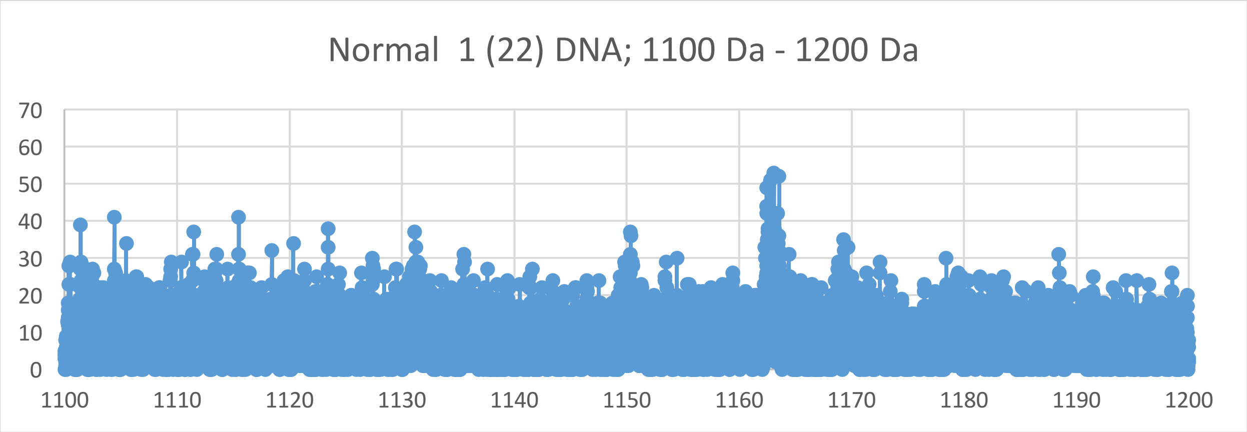
Normal 1 (22) DNA; 1100 Da - 1200 Da
0
10
20
30
40
50
60
70
1100
1110
1120
1130
1140
1150
1160
1170
1180
1190
1200

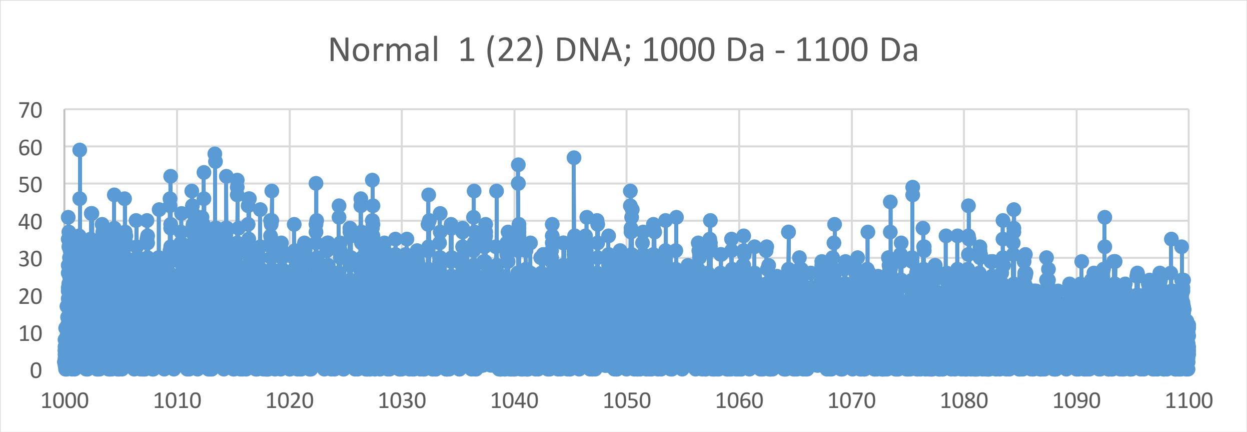
Normal 1 (22) DNA; 1000 Da - 1100 Da
0
10
20
30
40
50
60
70
1000
1010
1020
1030
1040
1050
1060
1070
1080
1090
1100

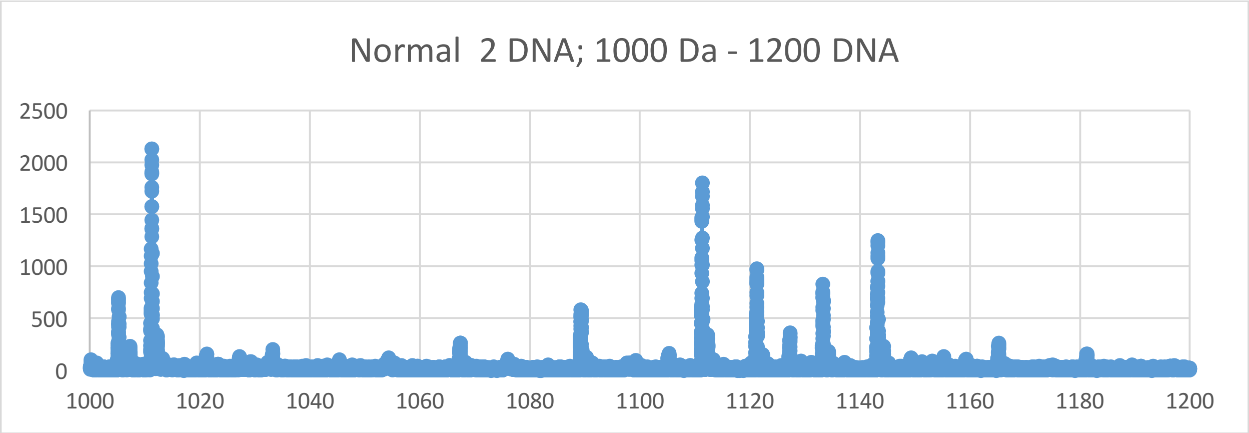
Normal 2 DNA; 1000 Da - 1200 DNA
2500
2000
1500
1000
500
0
1000
1020
1040
1060
1080
1100
1120
1140
1160
1180
1200

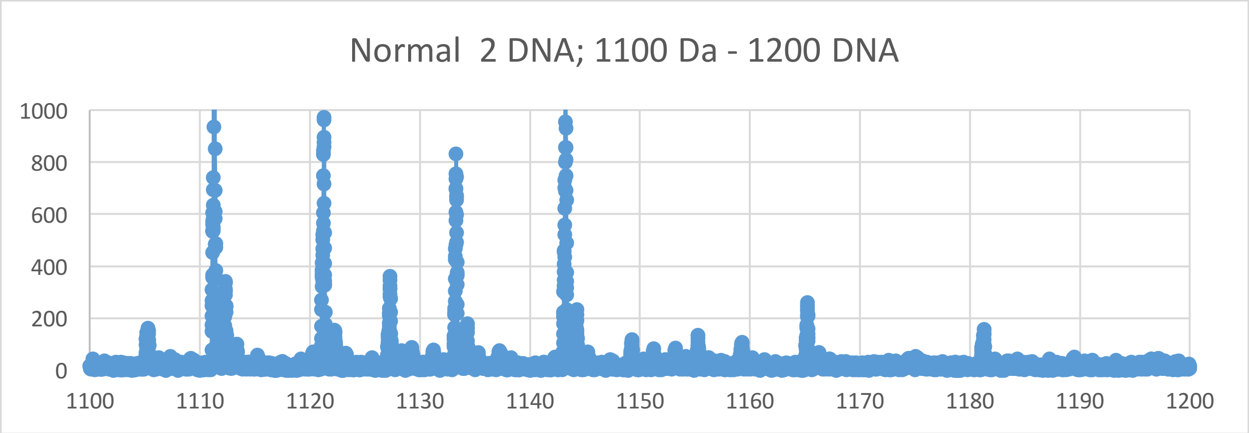
Normal 2 DNA; 1100 Da - 1200 DNA
1000
800
600
400
200
0
1100
1110
1120
1130
1140
1150
1160
1170
1180
1190
1200

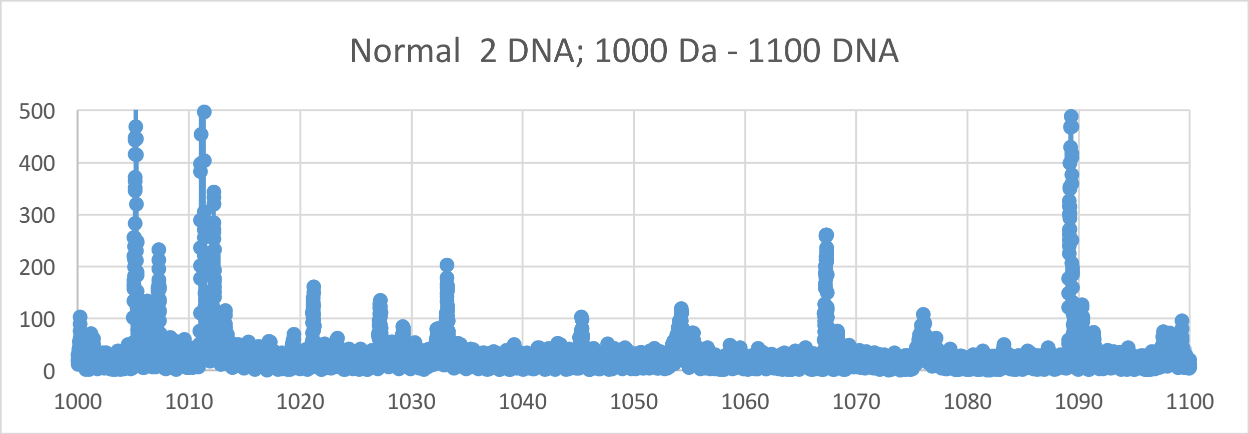
Normal 2 DNA; 1000 Da - 1100 DNA
500
400
300
200
100
0
1000
1010
1020
1030
1040
1050
1060
1070
1080
1090
1100

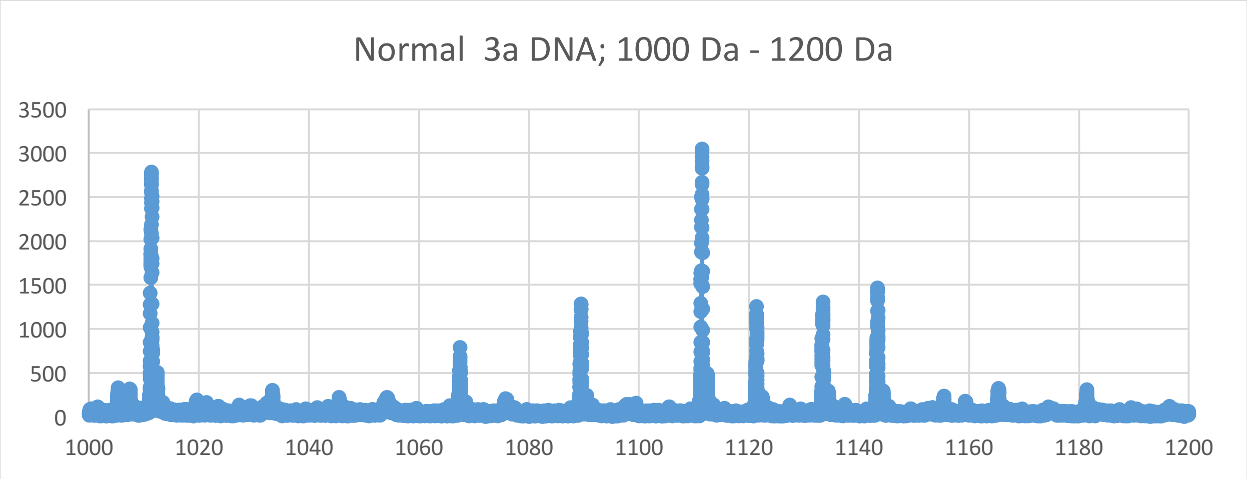

Normal 3a DNA; 1000 Da - 1200 Da
3500
3000
2500
2000
1500
1000
500
0
1000
1020
1040
1060
1080
1100
1120
1140
1160
1180
1200


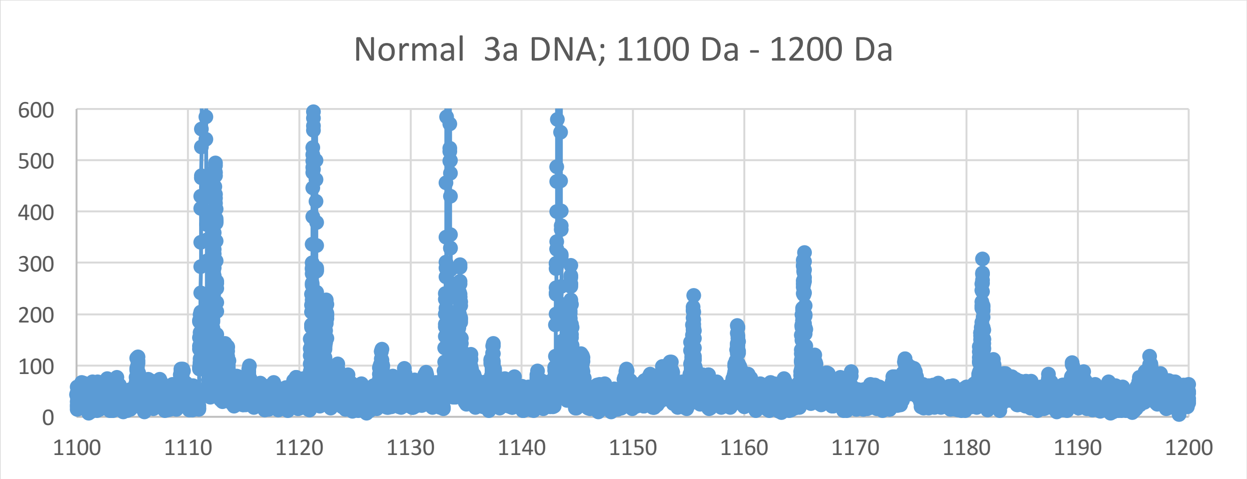

Normal 3a DNA; 1100 Da - 1200 Da
600
500
400
300
200
100
0
1100
1110
1120
1130
1140
1150
1160
1170
1180
1190
1200


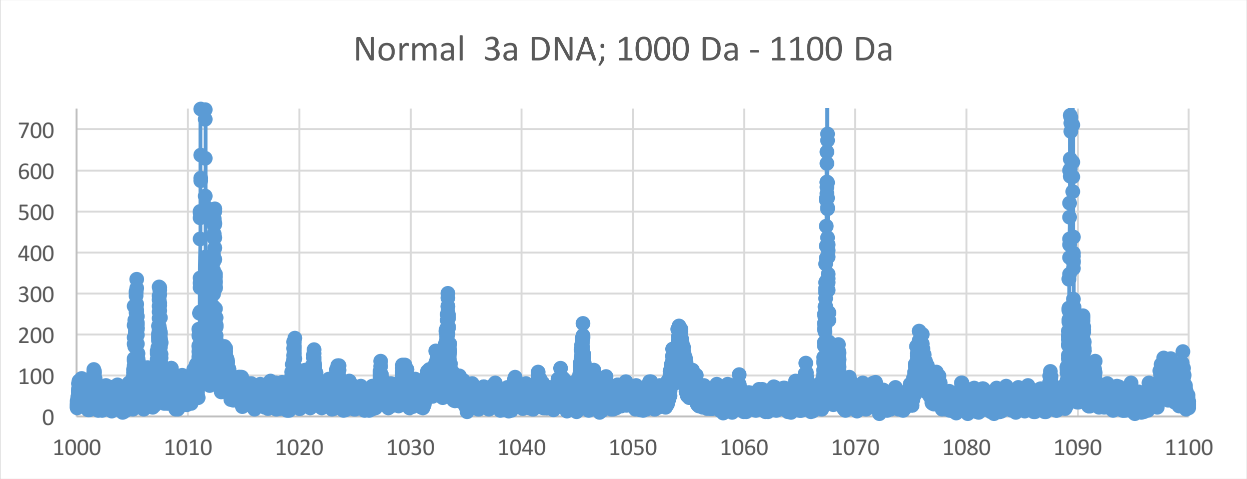

Normal 3a DNA; 1000 Da - 1100 Da
700
600
500
400
300
200
100
0
1000
1010
1020
1030
1040
1050
1060
1070
1080
1090
1100

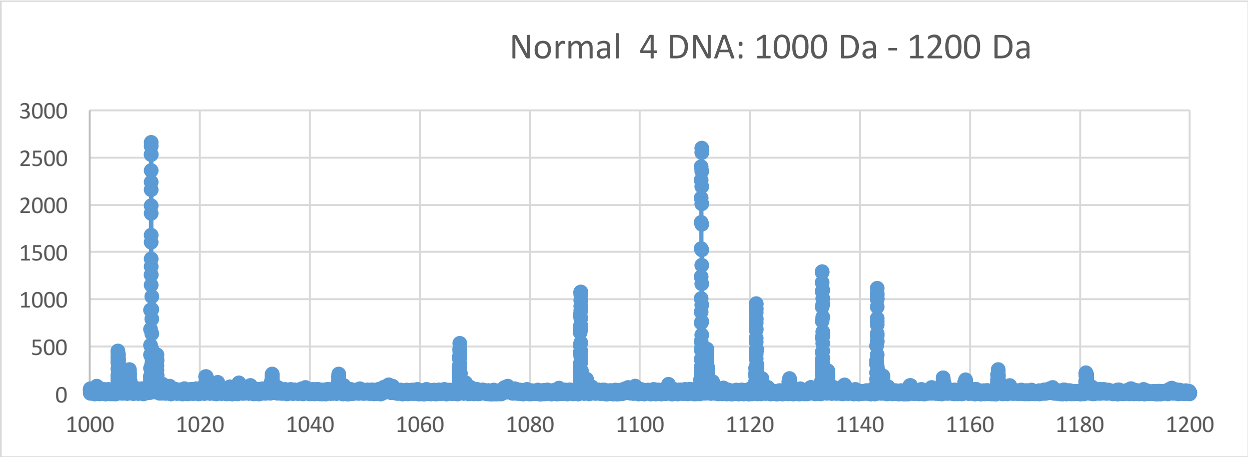
Normal 4 DNA: 1000 Da - 1200 Da
3000
2500
2000
1500
1000
500
0
1000
1020
1040
1060
1080
1100
1120
1140
1160
1180
1200

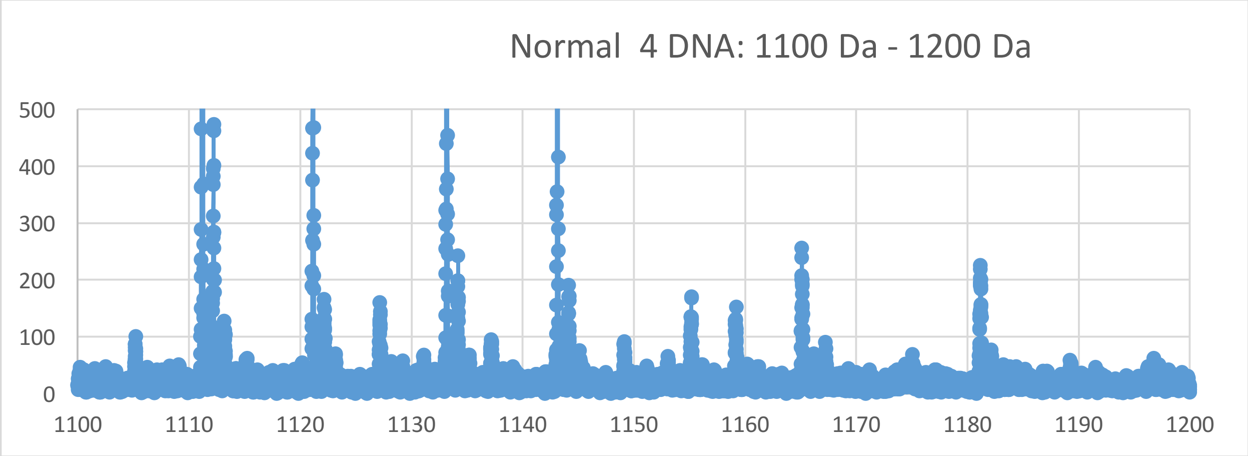
Normal 4 DNA: 1100 Da - 1200 Da
500
400
300
200
100
0
1100
1110
1120
1130
1140
1150
1160
1170
1180
1190
1200

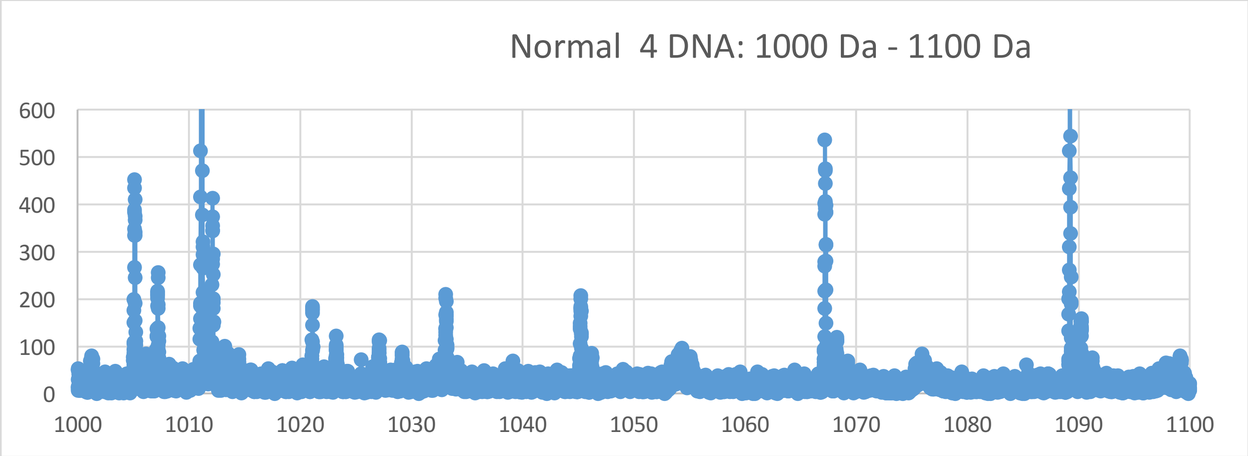
Normal 4 DNA: 1000 Da - 1100 Da
600
500
400
300
200
100
0
1000
1010
1020
1030
1040
1050
1060
1070
1080
1090
1100

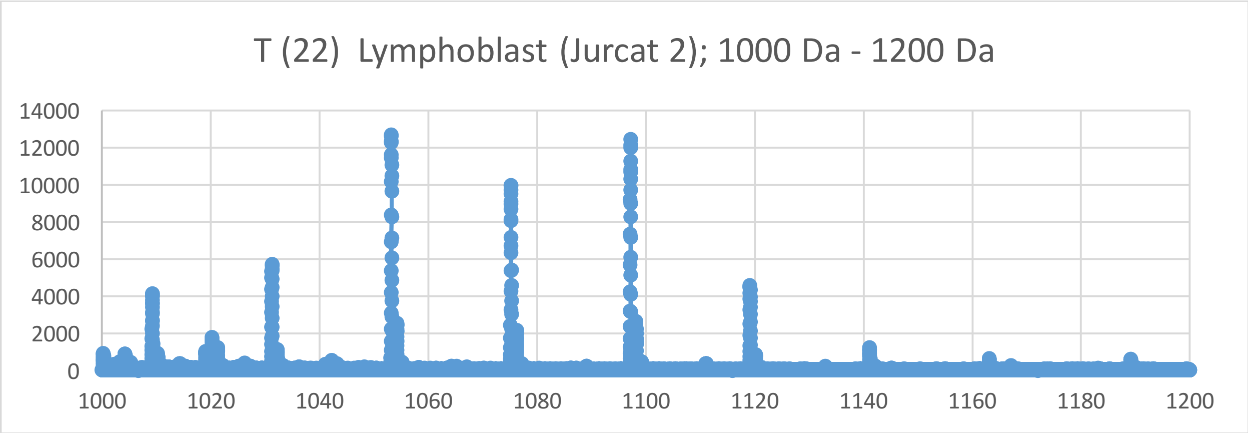
T (22) Lymphoblast (Jurcat 2); 1000 Da - 1200 Da
14000
12000
10000
8000
6000
4000
2000
0
1000
1020
1040
1060
1080
1100
1120
1140
1160
1180
1200

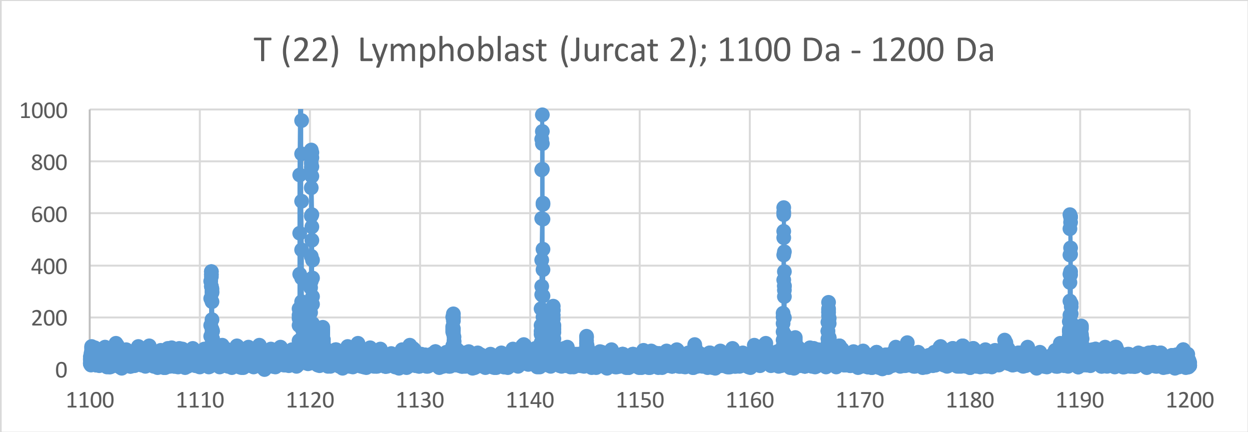
T (22) Lymphoblast (Jurcat 2); 1100 Da - 1200 Da
1000
800
600
400
200
0
1100
1110
1120
1130
1140
1150
1160
1170
1180
1190
1200

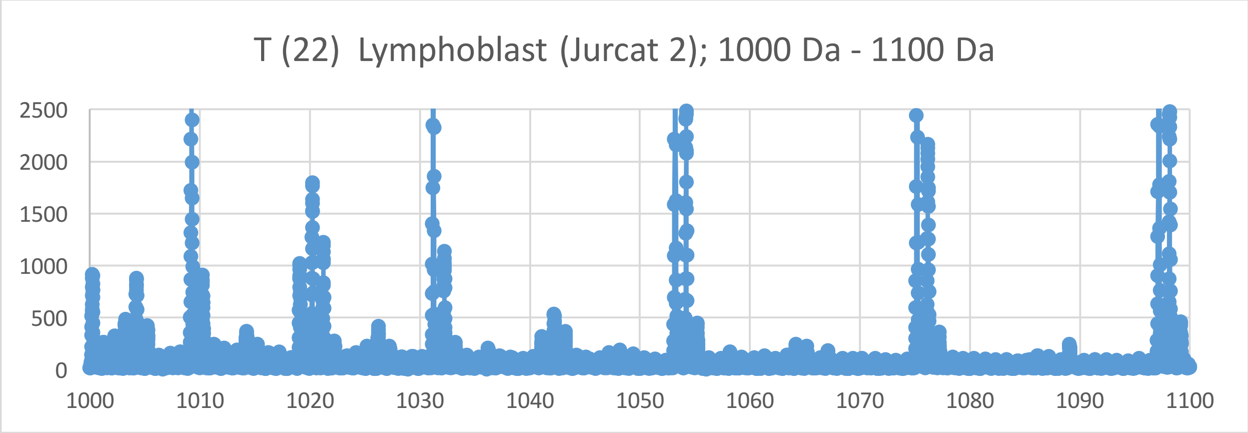
T (22) Lymphoblast (Jurcat 2); 1000 Da - 1100 Da
2500
2000
1500
1000
500
0
1000
1010
1020
1030
1040
1050
1060
1070
1080
1090
1100

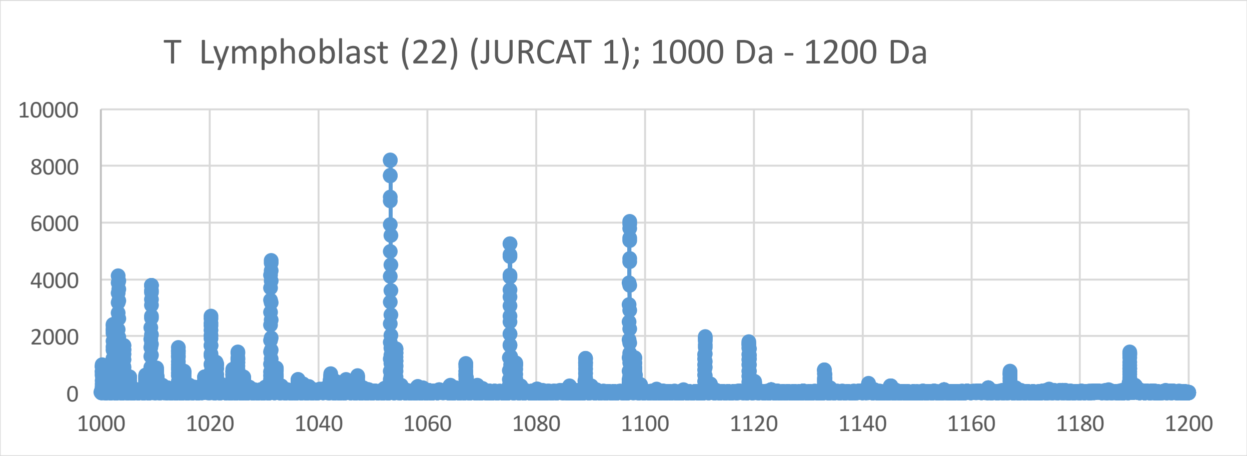
T Lymphoblast (22) (JURCAT 1); 1000 Da - 1200 Da
10000
8000
6000
4000
2000
0
1000
1020
1040
1060
1080
1100
1120
1140
1160
1180
1200

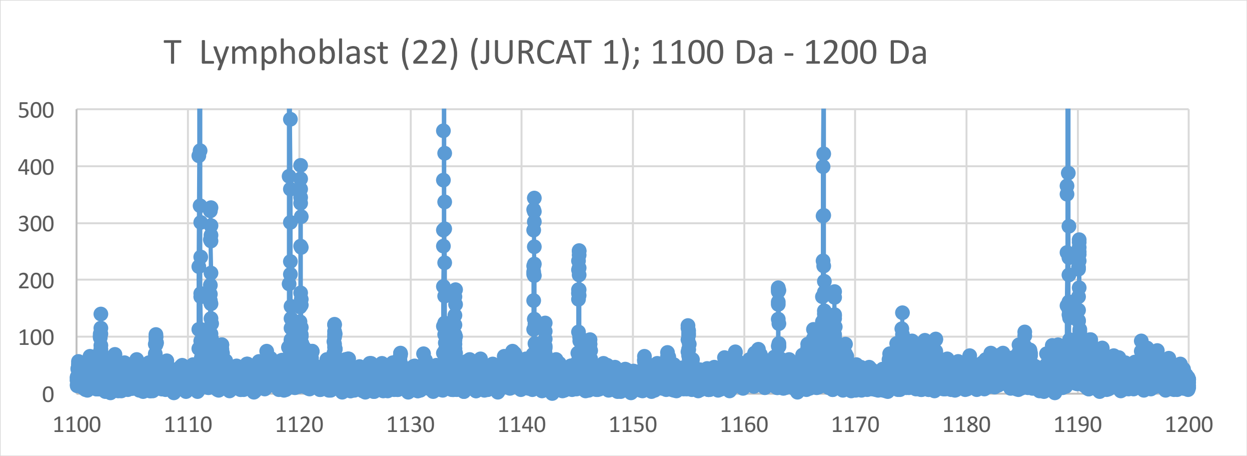
T Lymphoblast (22) (JURCAT 1); 1100 Da - 1200 Da
500
400
300
200
100
0
1100
1110
1120
1130
1140
1150
1160
1170
1180
1190
1200

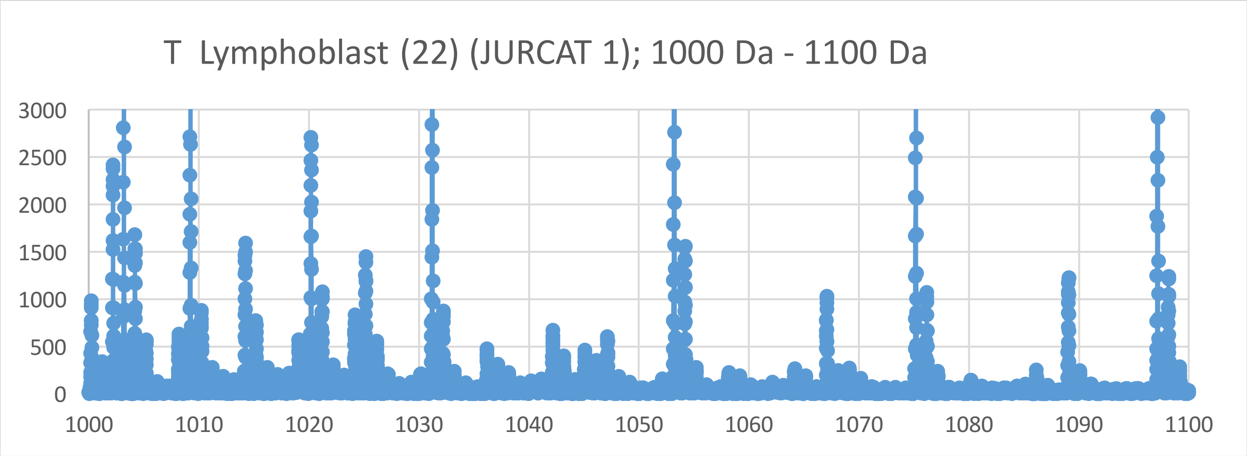
T Lymphoblast (22) (JURCAT 1); 1000 Da - 1100 Da
3000
2500
2000
1500
1000
500
0
1000
1010
1020
1030
1040
1050
1060
1070
1080
1090
1100

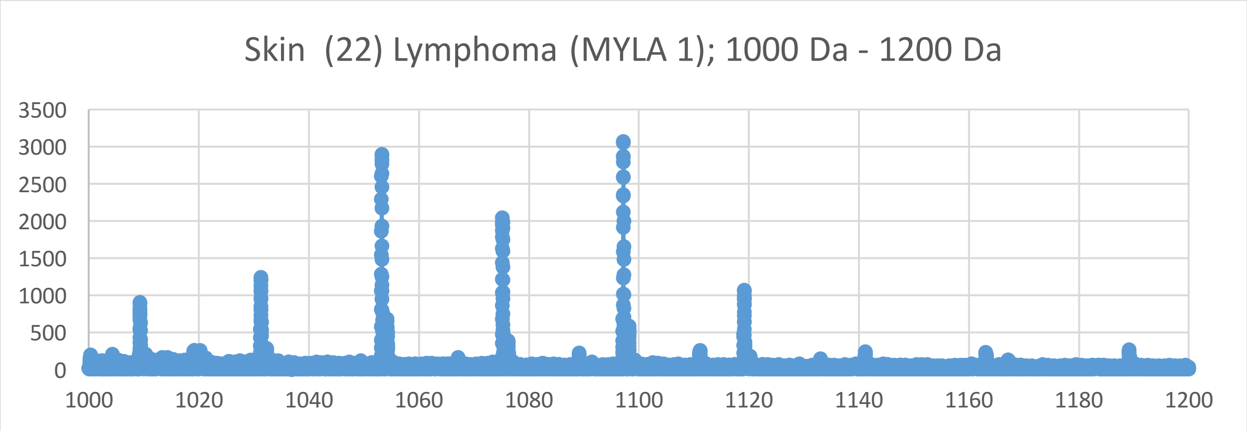
Skin (22) Lymphoma (MYLA 1); 1000 Da - 1200 Da
3500
3000
2500
2000
1500
1000
500
0
1000
1020
1040
1060
1080
1100
1120
1140
1160
1180
1200

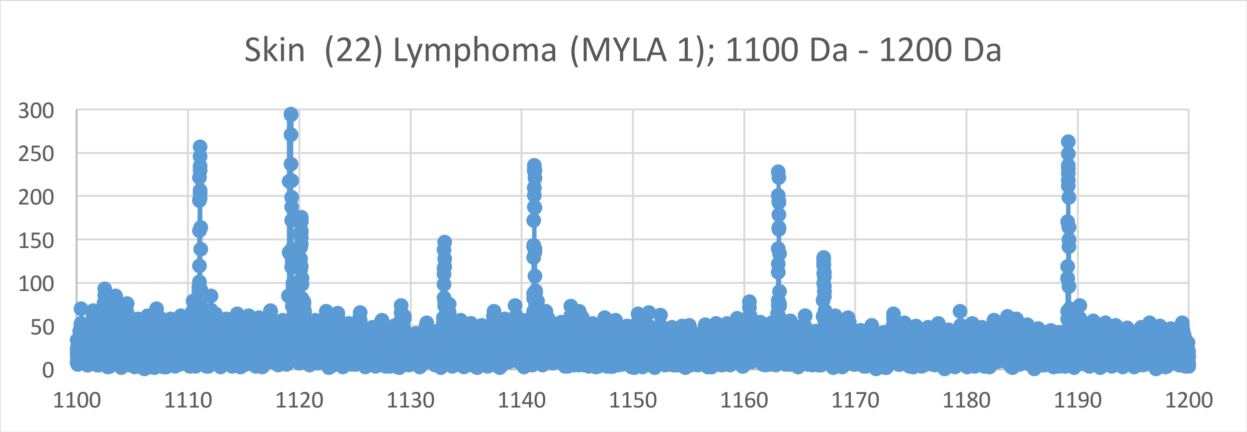
Skin (22) Lymphoma (MYLA 1); 1100 Da - 1200 Da
300
250
200
150
100
50
0
1100
1110
1120
1130
1140
1150
1160
1170
1180
1190
1200

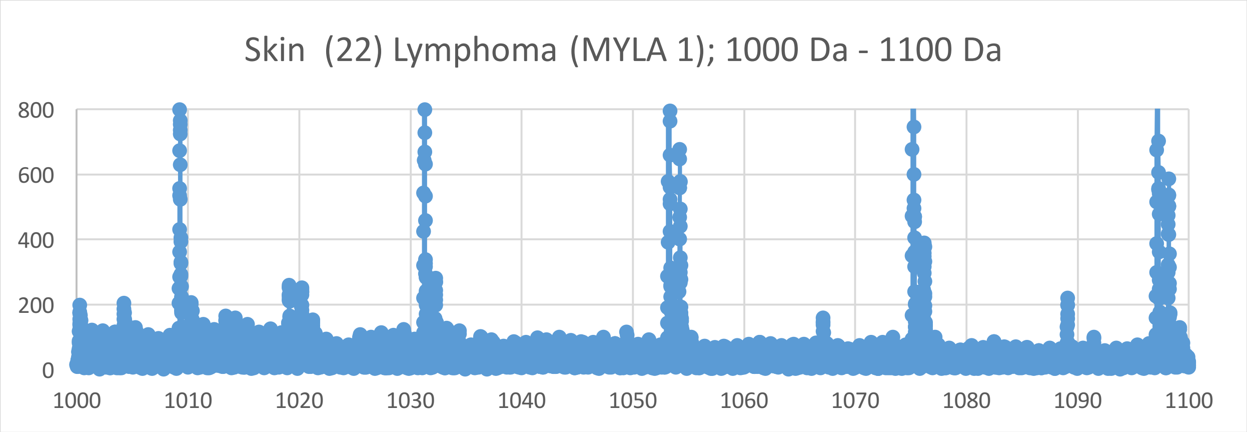
Skin (22) Lymphoma (MYLA 1); 1000 Da - 1100 Da
800
600
400
200
0
1000
1010
1020
1030
1040
1050
1060
1070
1080
1090
1100

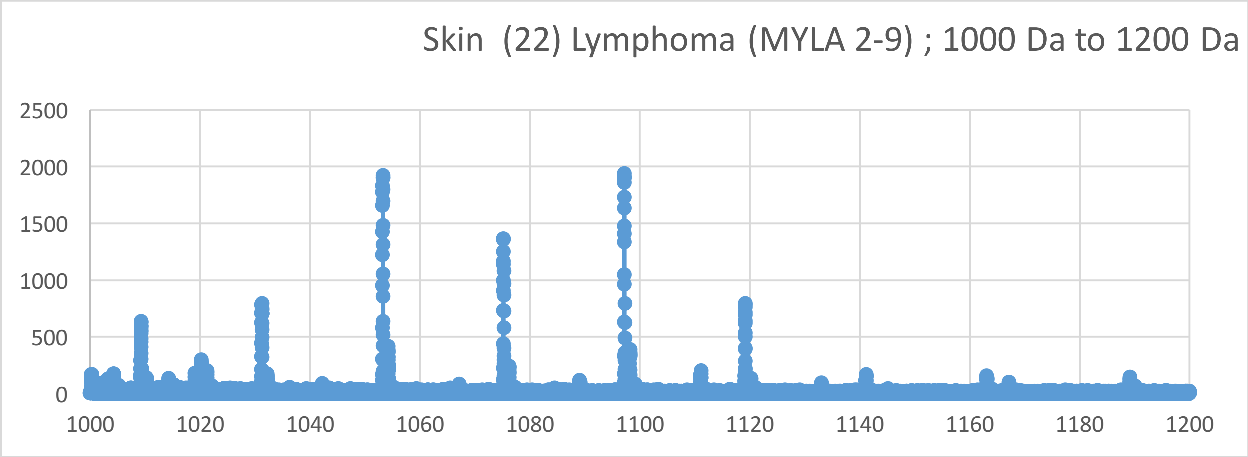
Skin (22) Lymphoma (MYLA 2-9) ; 1000 Da to 1200 Da
2500
2000
1500
1000
500
0
1000 1020 1040 1060 1080 1100 1120 1140 1160 1180 1200

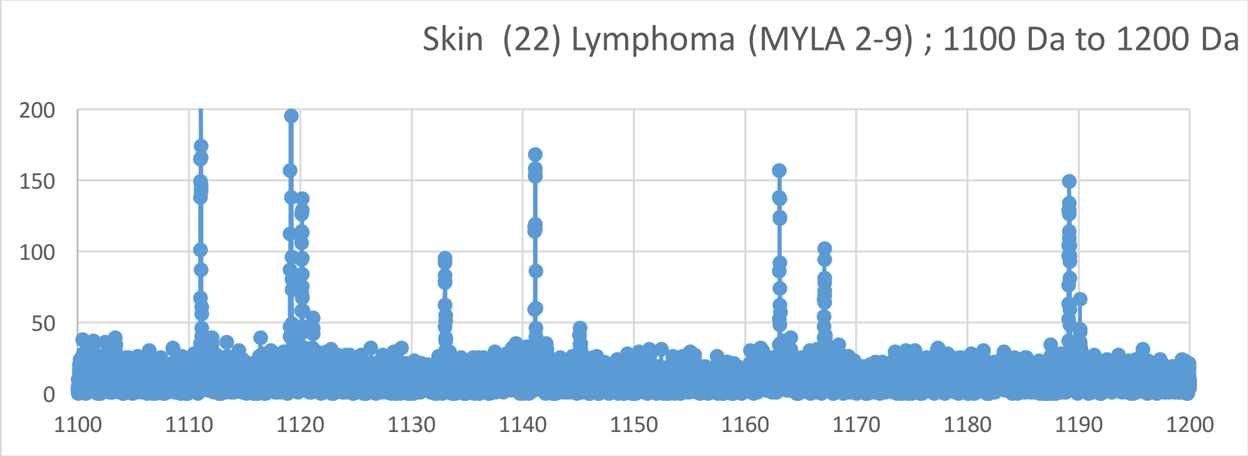
Skin (22) Lymphoma (MYLA 2-9) ; 1100 Da to 1200 Da
200
150
100
50
0
1100 1110 1120 1130 1140 1150 1160 1170 1180 1190 1200

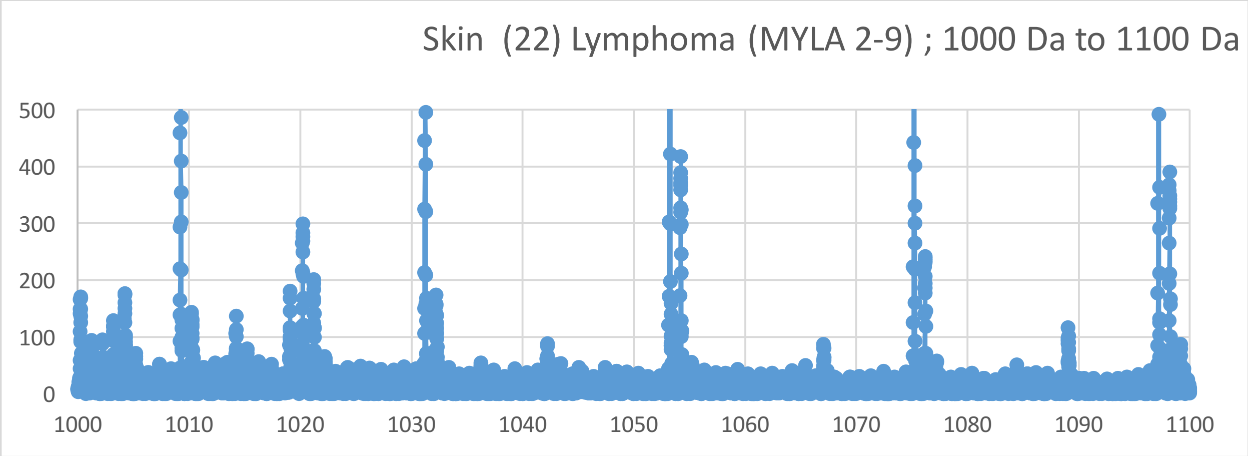
Skin (22) Lymphoma (MYLA 2-9) ; 1000 Da to 1100 Da
500
400
300
200
100
0
1000 1010 1020 1030 1040 1050 1060 1070 1080 1090 1100

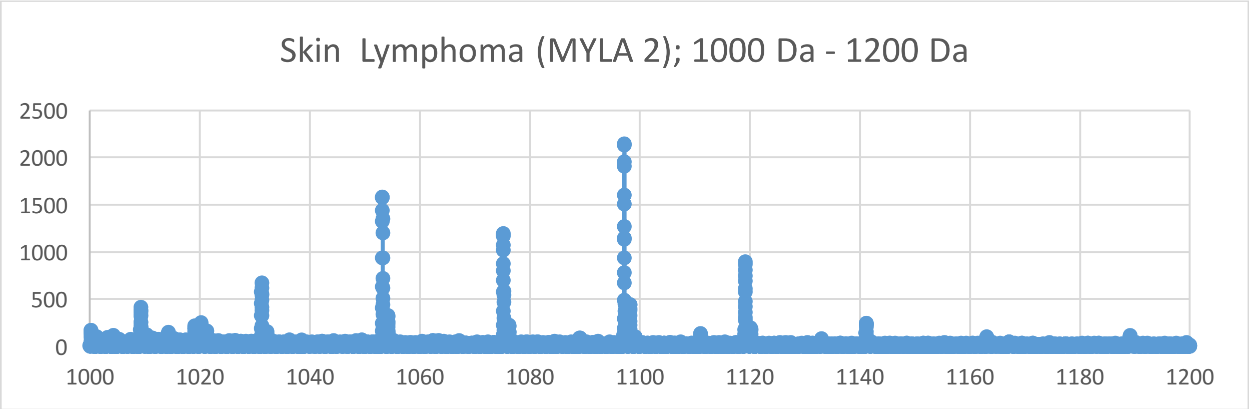
Skin Lymphoma (MYLA 2); 1000 Da - 1200 Da
2500
2000
1500
1000
500
0
1000
1020
1040
1060
1080
1100
1120
1140
1160
1180
1200

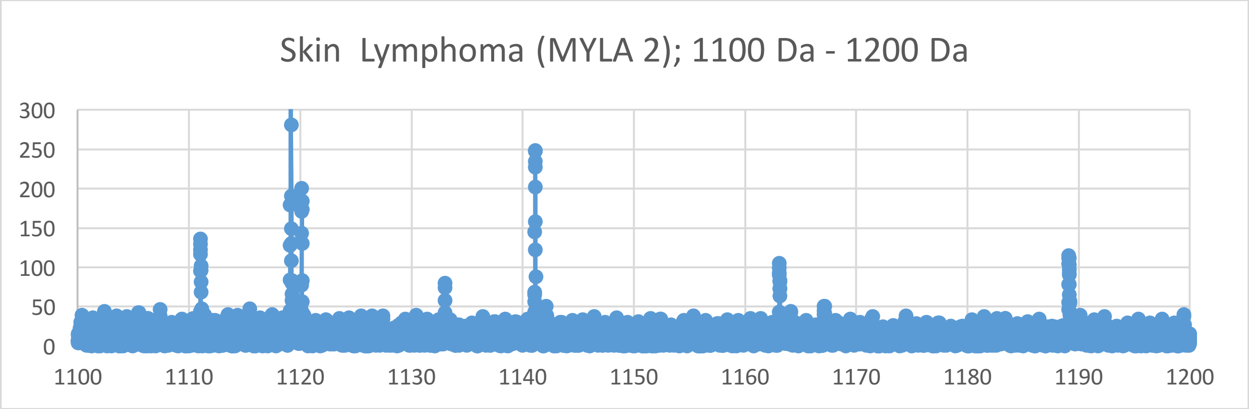
Skin Lymphoma (MYLA 2); 1100 Da - 1200 Da
300
250
200
150
100
50
0
1100
1110
1120
1130
1140
1150
1160
1170
1180
1190
1200

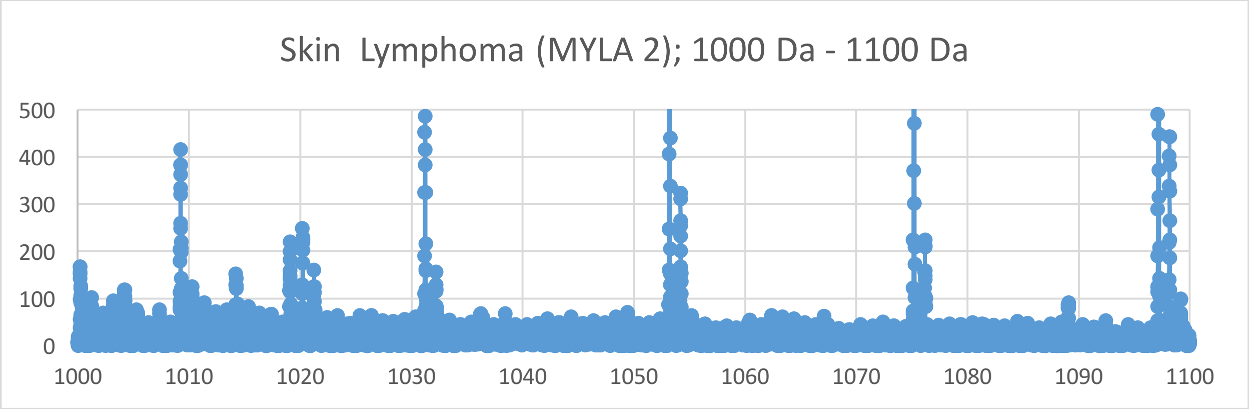
Skin Lymphoma (MYLA 2); 1000 Da - 1100 Da
500
400
300
200
100
0
1000
1010
1020
1030
1040
1050
1060
1070
1080
1090
1100
