## Supplementary Materials IX for "Oncogenesis and Aging by Isotopic Functionalizations of the Proteins and Nucleic Acids"

IX Supplementary Materials – Mass Spectra from 800 Da -1000 Da

Results for Supplementary 9

So for the trinucleotides at $(phos)_3$, the telomeric and mating telomeric peaks for cancer dramatically increase in intensities and number and the peaks for normal nucleotides decrease in number for stabilities of cancer telomeric trinucleotides. But the nontelomeres have increased in number and stabilities for cancer trinucleotides while the normal trinucleotides are mostly less in peaks and stabilities with $(phos)_3$. But as decrease the phosphate by 1 unit, the trinucleotides change so that the cancer trinucleotides more fragment and break from the DNA with negligible increase in intensities and stabilities for the telomeres and mating telomeres, with about the same fragmenting and stabilities of the cancer from DNA for telomeres and mating telomeres from $(phos)_3$ to $(phos)_2$. The masses range from 800 Da to 1000 Da. See Supplementary Materials IX for graphs. It seems that below some number of phosphate number the telomeres, mating telomeres and nontelomeres behave in same way toward cancer and normal. So this suggests normal may be favored and distinguished from cancer by changing phosphorylations. The telomeric and mating telomeric peaks for cancer negligibly increase in number and intensities with these dephosphorylations of the trinucleotides. The dephosphorylations from $(phos)_3$ to $(phos)_2$ caused the resulting negligible increase in intensities and stabilities of cancer trinucleotides to be more for telomeric and mating telomeric trinucleotides with less isotopic enrichments of heavy $^{13}C$ and $^{17}O$ and the dephosphorylations caused even negligibly more normal stabilities for nontelomeric trinucleotides and less isotopically enrichments of $^{13}C$ and $^{17}O$ for nontelomeres. It is important to consider that there are more intense peaks for the trinucleotides relative to the tetranucleotides for the saturated phosphate units and for some reason for large dephosphorylations, the intensities increase from some intermediate drops for medium phosphorylations. For the trinucleotides the intensities of both normal and cancer fragments slightly increased from $(phos)_3$ to $(phos)_2$. The dephosphorylations from $(phos)_3$ to $(phos)_2$ decrease isotopic enrichments of light isotopes of $^{15}N$ for telomeric and mating telimeric trinucleotides. The dephosphorylations from $(phos)_3$ to $(phos)_2$ also caused the telomeric, nontelomeric and mating telomeric trinucleotides to be negligibly more intense in cancer DNA and less isotopically enriched from $(phos)_3$ to $(phos)_2$. It seems the dephosphorylations are related to the isotopic enrichments and depletions; but the telomeres and mating telomeres are slightly more numerous in cancer and less isotopically enriched isotopes for telomeres, mating telomeres and nontelomeres from $(phos)_3$ to $(phos)_2$. In general, it seems with less phosphorylations there are changes in isotopic enrichments and the differences between telomeric, nontelomeric and mating telomeric oligonucleotide binding / stability diminish to induce cancer by such chemical mutations! The mating telomeric, telomeric and nontelomeric trinucleotides of cancer tended to also be less enriched so it seems the cancer stabilities of the mating telomeric, telomeric and nontelomeric trinucleotides are related to their less isotopic enrichments from $(phos)_3$ to $(phos)_2$ and not purely driven by dephosphorylations but both dephosphorylations and enrichments. Enrichments strengthen the cancerous telomeric, nontelomeric and mating telomeric trinucleotides while strongly weakening normal telomeric, matic telomeric and non-telomeric trinucleotides. The more intense cancerous telomeric, mating telomeric and nontelomeric trinucleotides were also less isotopically enriched. Such seems compatible for replications and transcriptions as stable telomeres seem compatible to unstable mating telomeres and nontelomeres for the unravelings and constructions of DNA mating telomeres onto the building templates of the telomeres. The dephosphorylations from $(phos)_3$ to $(phos)_2$ caused methylations to cause increase normal intensities and numbers of peaks telomeres and nontelomeres with increased isotopic enrichments. The methylations caused slight increase in cancer intensities and stabilities of mating telomeric trinucleotides with slight increased enrichments. Such loss of differing behaviors of telomeres and nontelomeres with greater dephosphorylations point to malfunctions of DNA by highly dephosphorylated nucleotides and plays a role in cancer. The methylations of the telomeres does slightly increased cancer intensities and stabilities with negligible changes in enrichments from $(phos)_3$ to $(phos)_2$ relative to the nontelomeres. It seems from these data that during biological processes the dephosphorylations and stabilizations of normal or vice versa dephosphorylations and stabilizations of cancer are or can be countered by methylations and destabilizations of the normal for cancer or vice versa dephosphorylations and destabilizations of the cancer by methylations as this research discovers coupled phosphorylations and methylations or coupled dephosphorylations and methylations with involved greater enrichments for normal biological functions and explaining cancer! It seems that if nucleotides lose too many phosphates in regulating methylations and acetylations that the differences among genes and in particular telomeric genes and nontelomeric genes are lost and the activity of cancer arises! The dephosphorylations from $(phos)_3$ to $(phos)_2$ caused acetylations to cause decrease in cancer stabilities for telomeres with decreased enrichments of heavier isotopes for telomeres . The acetylations caused increase stabilities, intensities and number of cancer peaks for telomeres with less isotopic enrichments. The mating telomeres slightly increase stabilities of normal with acetylations with negligible change in isotopic enrichments; where as the nontelomeres undergo negligible changes in intensities of cancer and normal trinucleotides with increased isotopic enrichments. This contrast in mating telomeres and telomeres from the acetylations may reflect the development of cancer as the telomeres fragment but the mating telomeres more holds together for damaged DNA for cancer.

So for the dinucleotides at $(phos)_6$, the telomeric peaks for cancer strongly increase in number and the peaks for normal decreases in number for more stable cancer dinucleotides and less stable normal dinucleotides with the $(phos)_6$. But the mating telomeres and nontelomeres have increase in number and stabilities for normal dinucleotides while the cancer dinucleotides are less in peaks and stabilities with $(phos)_6$. But as decrease the phosphate by 1 unit, the dinucleotides change so that the normal dinucleotides more fragment and break and are stable from the DNA with more number and stabilities for the telomeres and nontelomeres, with slightly more fragmenting and stabilities of the normal from DNA for telomeres and nontelomeres from $(phos)_6$ to $(phos)_5$. The masses range from 800 Da to 1000 Da. See Supplementary Figures IX for graphs. It seems that below some number of phosphates, the telomeres, mating telomeres and nontelomers behave in same way within cancer and normal DNA. So this suggests normal may be favored and distinguished from cancer by changing saturated phosphorylations. The telomeric peaks for normal increase with these dephosphorylations of the dinucleotides. The dephosphorylations from $(phos)_6$ to $(phos)_5$

caused the resulting more intense and stable normal dinucleotides to be more telomeric and nontelomeric dinucleotides and more isotopic enrichments for telomeres and nontelomeres. It is important to consider that there are more intense peaks for the dinucleotides relative to the trinucleotides and tetranucleotides for the saturated phosphate units. For the dinucleotides the intensities of normal increased and the intensities of cancer fragments decreased from $(phos)_6$ to $(phos)_{5.}$ The dephosphorylations of mating telomeric dinucleotides from $(phos)_6$ to $(phos)_5$ decrease normal stabilities and isotopic enrichments of heavy isotopes of $^{13}C$ and $^{17}O$. The dephosphorylations from $(phos)_6$ to $(phos)_5$ also caused the telomeric and nontelomeric dinucleotides to be more intense in normal DNA and less isotopically enriched. It seems the dephosphorylations are related to the isotopic enrichments and depletions; but the telomeres and nontelomeres are slightly more intense in normal and less isotopically enriched whereas the intensities of the mating telomeres are more intense in cancer and the less isotopic enriched. The telomeric and nontelomeric dinucleotides of normal tended to also be less enriched so it seems the cancer stabilities of the telomeric and nontelomeric dinucleotides are related to their less isotopic enrichments for the dephosphorylations from $(phos)_6$ to $(phos)_5$ and purely driven by dephosphorylations and enrichments. Enrichments weaken the nontelomeric cancer dinucleotides while strongly strengthening normal telomeric dinucleotides. The more intense cancer nontelomeric dinucleotides were also less isotopically enriched, but more intense cancerous telomeric dinucleotides were less enriched isotopically. The dephosphorylations from $(phos)_6$ to $(phos)_5$ decreased the isotopic enrichments of the normal for the nontelomeric and cancer mating telomeric dinucleotides. But the nontelomeric dinucleotides were less enriched from the dephosphorylations for less enrichments and stronger more stable normal dinucleotides for nontelomeres. Such seems compatible for replications and transcriptions as stable telomeres seem compatible to unstable mating telomeres for the unravelings and constructions of DNA mating telomeres onto the building templates of the telomeres. The dephosphorylations from $(phos)_6$ to $(phos)_5$ caused methylations to cause slightly less cancer intensities and stabilities and less isotopic enrichments for nontelomeres. But the methylations of the telomeres and mating telomeres caused negligible alterations cancer stabilities and isotopic enrichments, but the mating telomeres had slightly more isotopic enrichments. The methylations of the nontelomeres slightly increased cancer intensities and stabilities with slightly negligible enrichments. It seems from these data that during biological processes the dephosphorylations and stabilizations of normal or vice versa dephosphorylations and stabilizations of cancer are or can be countered by methylations and destabilizations of the normal for cancer or vice versa dephosphorylations and destabilizations of the cancer by methylations as this research discovers coupled phosphorylations and methylations or coupled dephosphorylations and methylations with involved greater enrichments for normal biological function and explaining cancer! The dephosphorylations from $(phos)_6$ to $(phos)_5$ caused acetylations to cause more normal stabilities for mating telomeres and nontelomeres with strong isotopic enrichments and negligible cancer stabilities for telomeres with strong enrichments. The acetylations from $(phos)_6$ to $(phos)_5$ caused more normal stabilities and intensities for mating telomeres and nontelomeres with strongly more enrichments. This contrast in mating telomeres and telomeres with telomeres from the acetylations may reflect the developments of cancer as the telomeres fragment but the mating telomeres and nontelomeres more hold together for damaged DNA for cancer.

So for the dinucleotides at $(phos)_5$, the telomeric peaks for normal strongly increase in number and the peaks for cancer decreases in number for more stable normal dinucleotides and less stable cancer dinucleotides with the $(phos)_5$. But the mating telomeres have increase in number and stability for mixed cancer dinucleotides while the mixed cancer and normal dinucleotides are less in peaks and stabilities with $(phos)_5$. The nontelomeres favor cancer at $(phos)_5$. But as decrease the phosphate by 1 unit, the dinucleotides change so that the cancer dinucleotides more fragment and break and are stable from the DNA with more numbers and stabilities for the telomeres, mating telomeres and nontelomeres $(phos)_5$ to $(phos)_4$. The masses range from 800 Da to 1000 Da. See Supplementary Figures IX for graphs. It seems that within the saturation range of some number of phosphates, the telomeres, mating telomeres and nontelomers behave in different way within cancer and normal DNA. So this suggest cancer may be favored and distinguished from normal by changing saturated phosphorylations from $(phos)_5$ to $(phos)_4$. The telomeric, mating telomeric and nontelomeric peaks for cancer increase with this dephosphorylations of the dinucleotides. The dephosphorylations from $(phos)_5$ to $(phos)_4$ caused the resulting more intense and stable cancer dinucleotides to be more telomeric, mating telomeric and nontelomeric dinucleotides and strongly more isotopic enrichments for telomeres, mating telomeres and nontelomere. It is important to consider that there are more intense peaks for the dinucleotides relative to the trinucleotides and tetranucleotides for the saturated phosphate units. For the dinucleotides the intensities of cancer increased and the intensities of normal fragments decreased from $(phos)_5$ to $(phos)_4$. The dephosphorylations of telomeric, mating telomeric and nontelomeric dinucleotides from $(phos)_5$ to $(phos)_4$ strongly increases isotopic enrichments of heavy isotopes of $^{13}C$ and $^{17}O$. This may be the greatest data demonstrating the isotopic cause for mutations and phosphate driven isotopic enrichments and phosphate and isotopic enrichments for genesis of cancer! The dephosphorylations from $(phos)_5$ to $(phos)_4$ also caused the telomeric, mating telomeric and nontelomeric dinucleotides to be more intense in cancer DNA and greater isotopically enriched. It seems the dephosphorylations are related to the isotopic enrichments and depletions; but the telomeres, mating telomeres and nontelomeres are hugely more intense in cancer and much more isotopically enriched. The telomeric, mating telomeric and nontelomeric dinucleotides of cancer tended to also be more isotopically enriched so it seems the cancer stabilities of the telomeric, mating telomeric and nontelomeric dinucleotides are related to their greater isotopic enrichments from the dephosphorylations from $(phos)_5$ to $(phos)_4$ and the mutations purely driven by dephosphorylations and enrichments. Enrichments weaken the nontelomeric normal dinucleotides while strongly strengthening cancerous telomeric dinucleotides. The more intense cancer telomeric, mating telomeric and nontelomeric dinucleotides were also more isotopically enriched, but more intense cancerous telomeric, mating telomeric and nontelomeric dinucleotides were more enriched isotopically. The dephosphorylations from $(phos)_5$ to $(phos)_4$ increased the isotopic enrichments of the cancer for the telomeric, nontelomeric and mating telomeric dinucleotides. Such seems compatible for replications and transcriptions as stable telomeres seem compatible to unstable mating telomeres for the unravelings and constructions of DNA mating telomeres onto the building templates of the telomeres. But in this case of all normal telomeric, mating telomeric and nontelomeric dinucleotides are enriched and unstable then the replications, transcriptions and translations cannot occur normally. The dephosphorylations from $(phos)_5$ to $(phos)_4$ caused methylations to cause negligibly more cancer intensities and stabilities but lowered isotopic enrichments for nontelomeres. But the methylations of the nontelomeres and mating telomeres caused negligible alterations cancer stabilities and isotopic enrichments. The methylations of the nontelomeres slightly increased cancer intensities and stabilities with slightly negligible enrichments. It seems from these data that during biological processes the dephosphorylations and stabilizations of normal or vice versa dephosphorylations and stabilizations of cancer are or can be countered by methylations (and isotopic enrichments) and destabilizations of the normal for cancer or vice versa dephosphorylations and destabilizations of the cancer by methylations as this research discovers coupled phosphorylations and methylations (and isotopic enrichments) or coupled dephosphorylations and methylations (and isotopic enrichments) with involved greater enrichments for normal biological function and explaining cancer! The dephosphorylations from $(phos)_5$ to $(phos)_4$ caused acetylations to cause negligible change in stabilities of cancer for mating telomeres and nontelomeres with strong isotopic enrichments and negligible change in intensities of cancer for telomeres with strong enrichments. The acetylations from $(phos)_5$ to $(phos)_4$ caused less normal stabilities and intensities for telomeres and mating telomeres with strongly more enrichments. This contrast in mating telomeres and telomeres with telomeres from the acetylations may reflect the developments of cancer as the telomeres fragment but the mating telomeres and nontelomeres more hold together for damaged DNA for cancer.

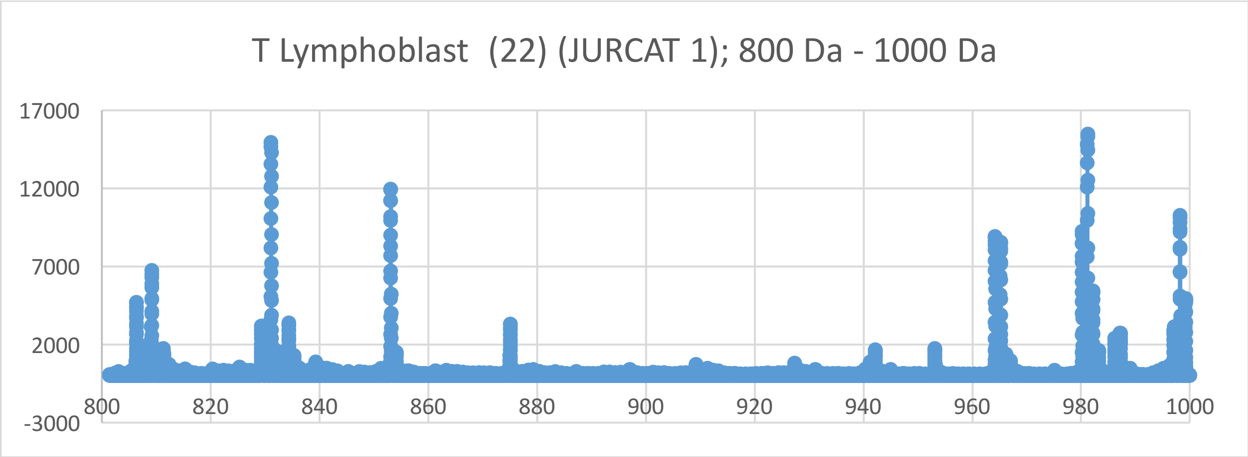
T Lymphoblast (22) (JURCAT 1); 800 Da - 1000 Da
17000
12000
7000
2000
-3000
800
820
840
860
880
900
920
940
960
980
1000

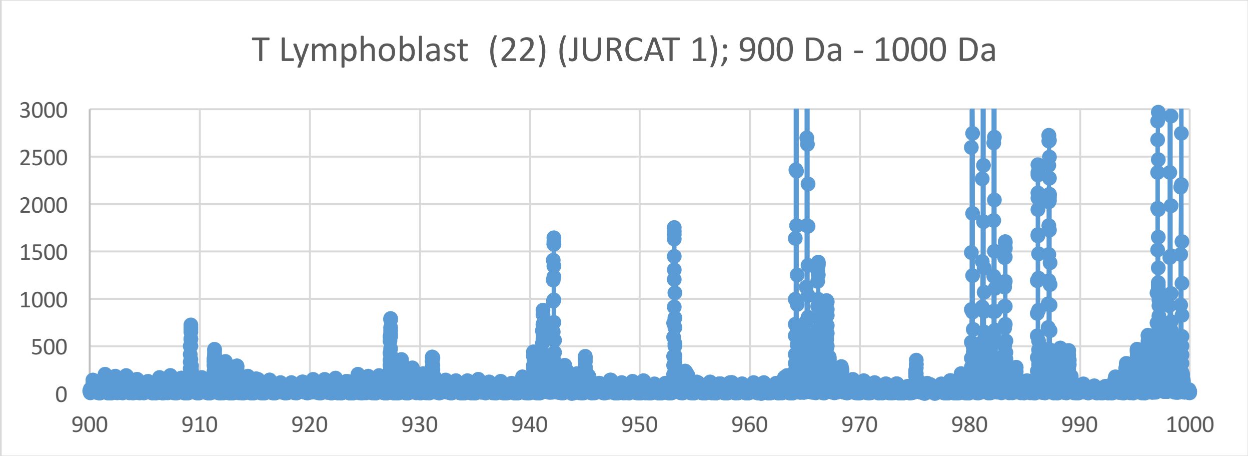
T Lymphoblast (22) (JURCAT 1); 900 Da - 1000 Da
3000
2500
2000
1500
1000
500
0
900
910
920
930
940
950
960
970
980
990
1000

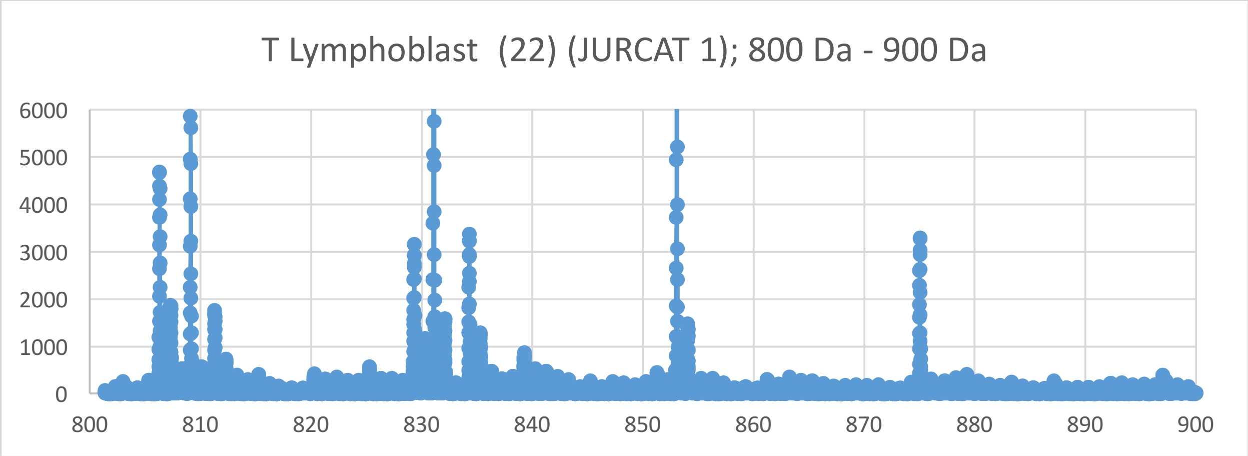
T Lymphoblast (22) (JURCAT 1); 800 Da - 900 Da
6000
5000
4000
3000
2000
1000
0
800
810
820
830
840
850
860
870
880
890
900

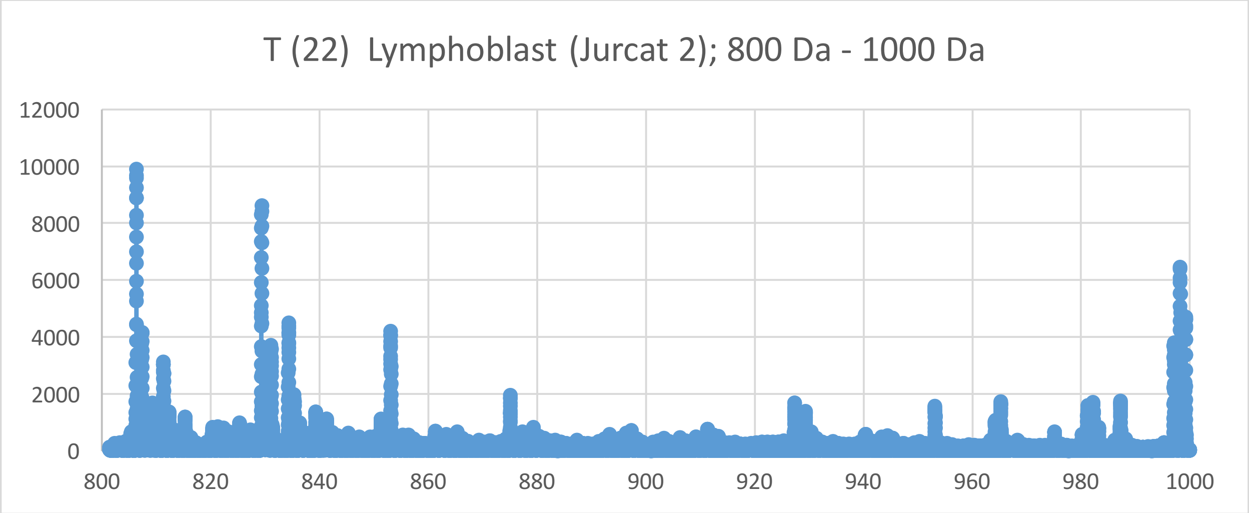
T (22) Lymphoblast (Jurcat 2); 800 Da - 1000 Da
12000
10000
8000
6000
4000
2000
0
800
820
840
860
880
900
920
940
960
980
1000

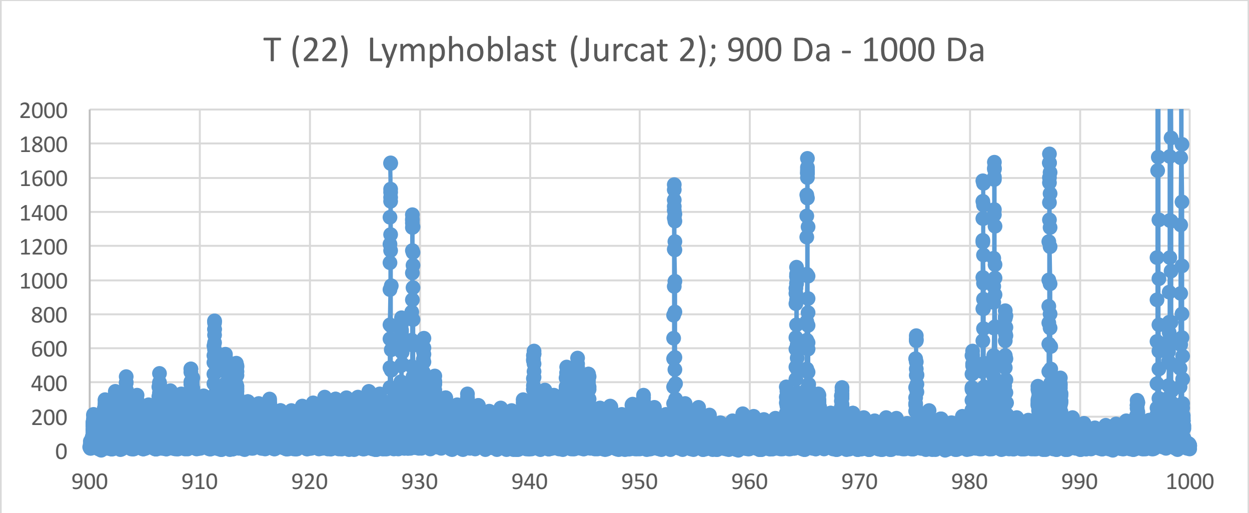
T (22) Lymphoblast (Jurcat 2); 900 Da - 1000 Da
2000
1800
1600
1400
1200
1000
800
600
400
200
0
900
910
920
930
940
950
960
970
980
990
1000

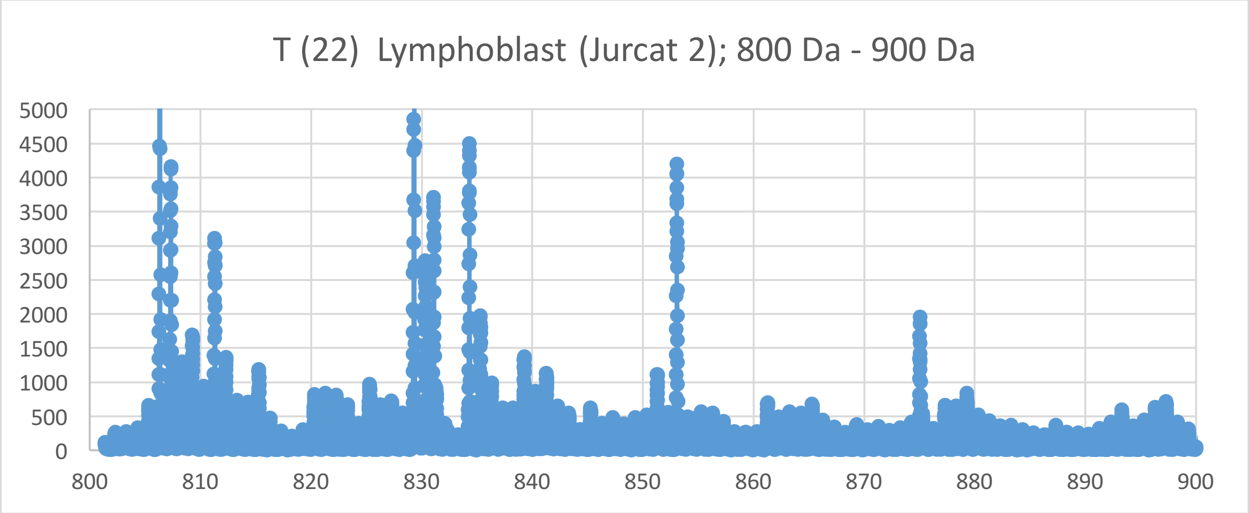
T (22) Lymphoblast (Jurcat 2); 800 Da - 900 Da
5000
4500
4000
3500
3000
2500
2000
1500
1000
500
0
800
810
820
830
840
850
860
870
880
890
900

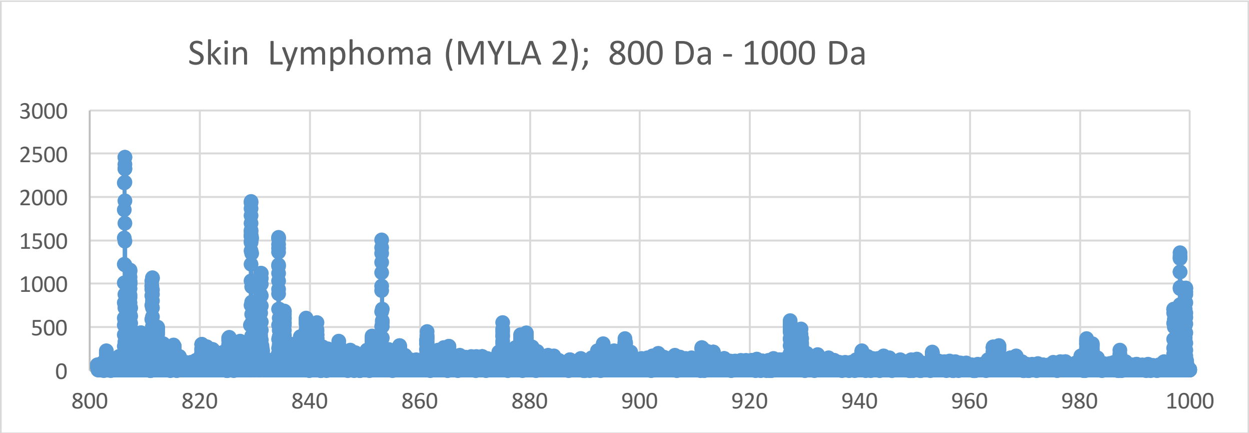
Skin Lymphoma (MYLA 2); 800 Da - 1000 Da
3000
2500
2000
1500
1000
500
0
800
820
840
860
880
900
920
940
960
980
1000

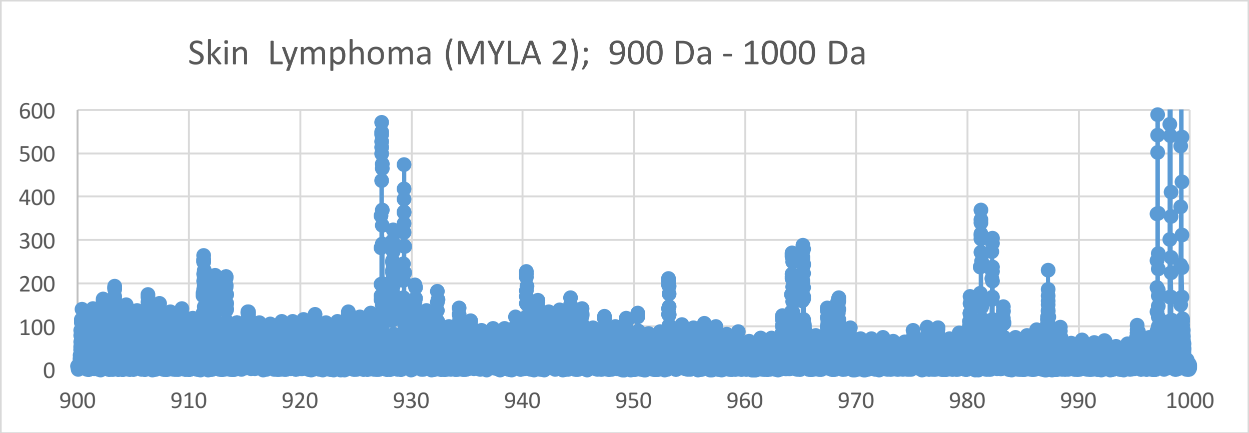
Skin Lymphoma (MYLA 2); 900 Da - 1000 Da
600
500
400
300
200
100
0
900
910
920
930
940
950
960
970
980
990
1000

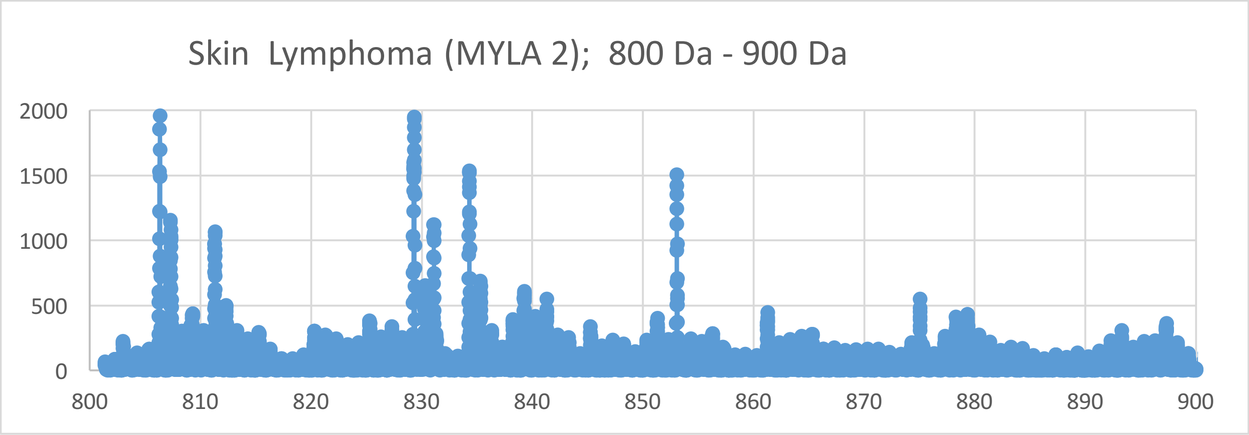
Skin Lymphoma (MYLA 2); 800 Da - 900 Da
2000
1500
1000
500
0
800
810
820
830
840
850
860
870
880
890
900

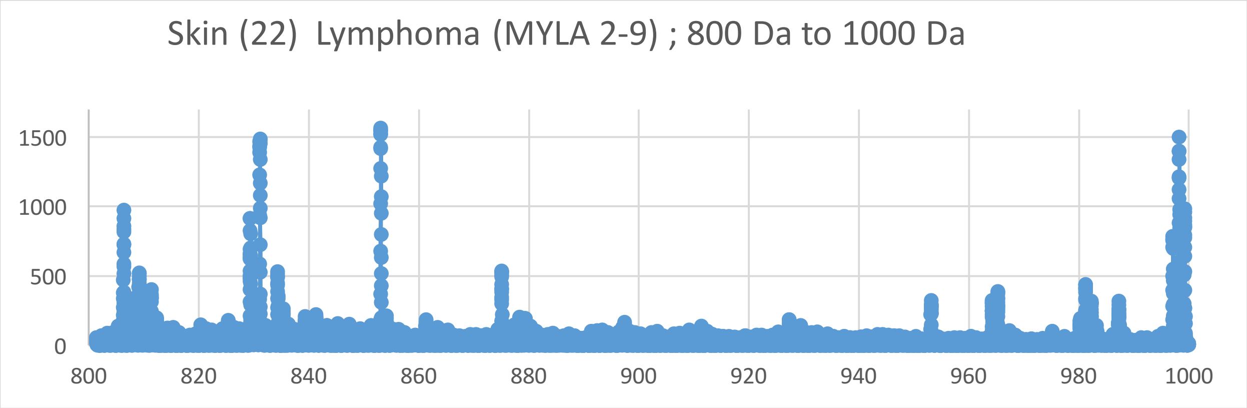
Skin (22) Lymphoma (MYLA 2-9) ; 800 Da to 1000 Da
1500
1000
500
0
800
820
840
860
880
900
920
940
960
980
1000

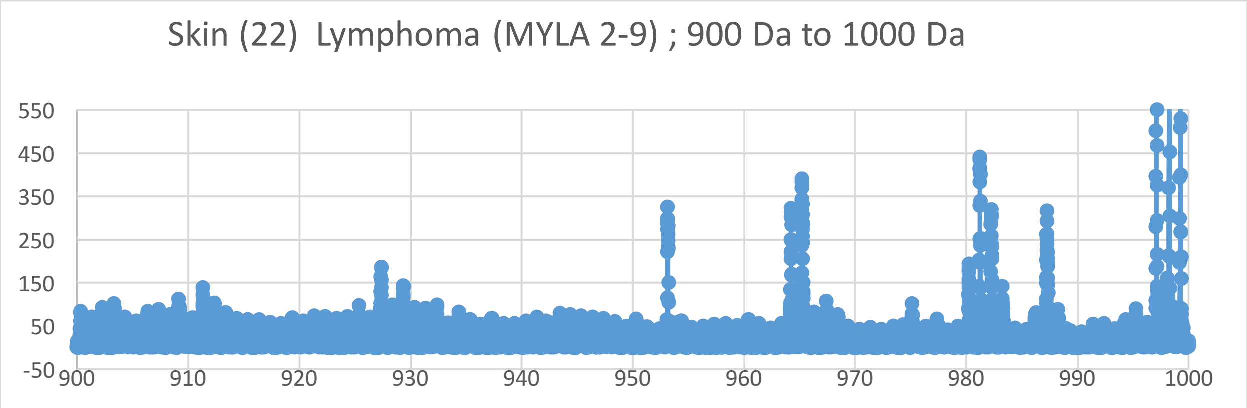
Skin (22) Lymphoma (MYLA 2-9) ; 900 Da to 1000 Da
550
450
350
250
150
50
-50
900
910
920
930
940
950
960
970
980
990
1000

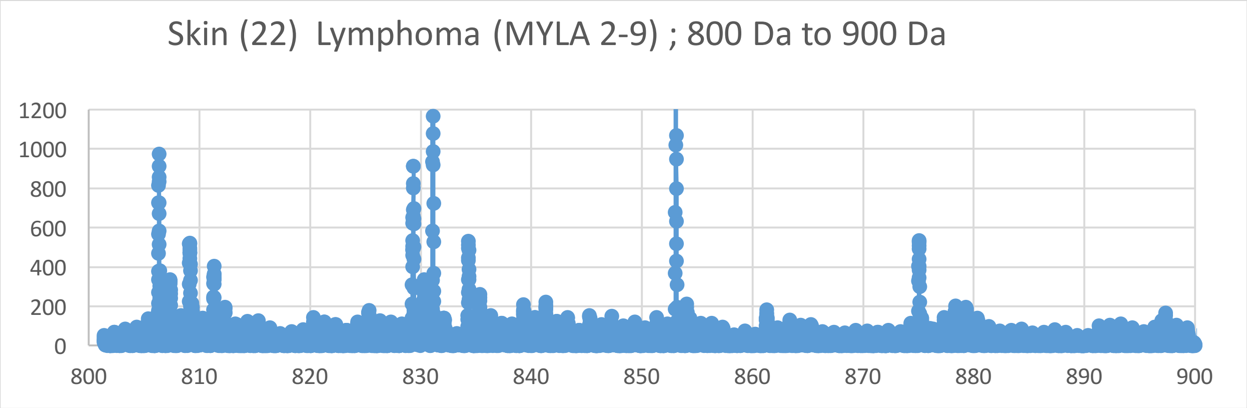
Skin (22) Lymphoma (MYLA 2-9) ; 800 Da to 900 Da
1200
1000
800
600
400
200
0
800
810
820
830
840
850
860
870
880
890
900

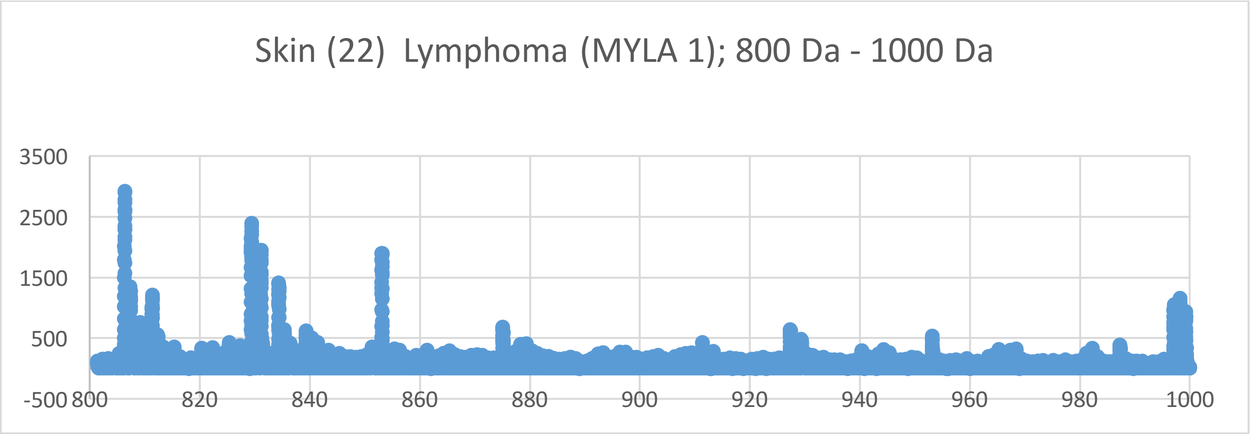
Skin (22) Lymphoma (MYLA 1); 800 Da - 1000 Da
3500
2500
1500
500
-500
800
820
840
860
880
900
920
940
960
980
1000

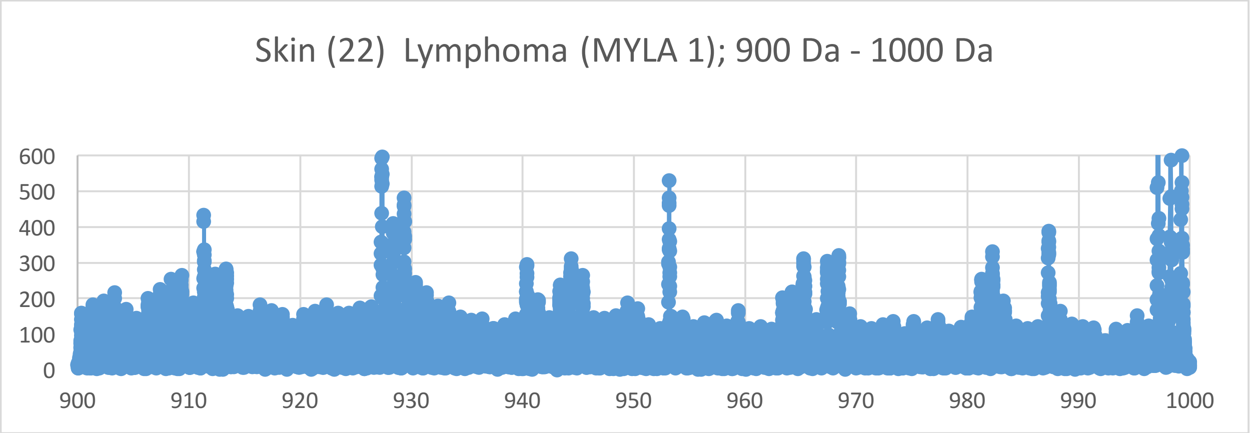
Skin (22) Lymphoma (MYLA 1); 900 Da - 1000 Da
600
500
400
300
200
100
0
900
910
920
930
940
950
960
970
980
990
1000

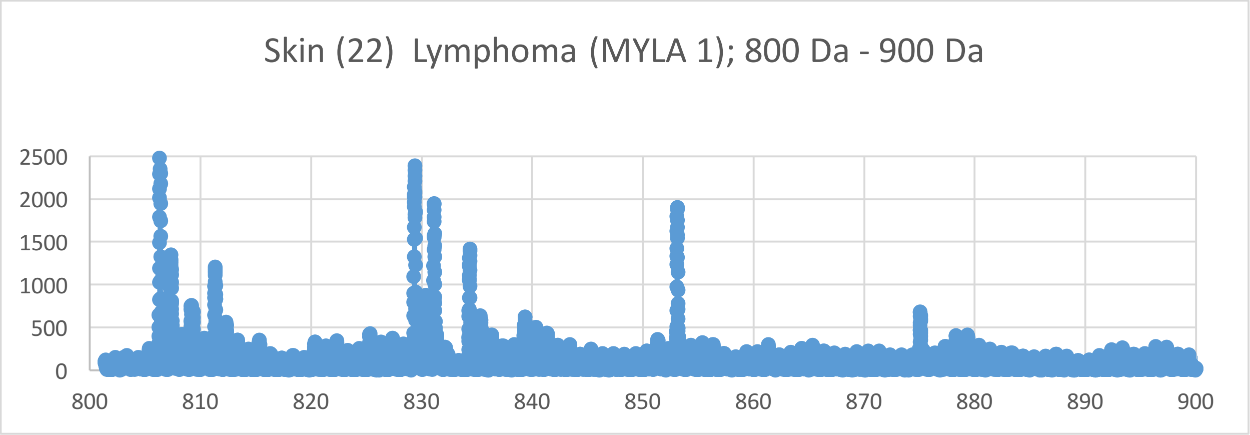
Skin (22) Lymphoma (MYLA 1); 800 Da - 900 Da
2500
2000
1500
1000
500
0
800
810
820
830
840
850
860
870
880
890
900

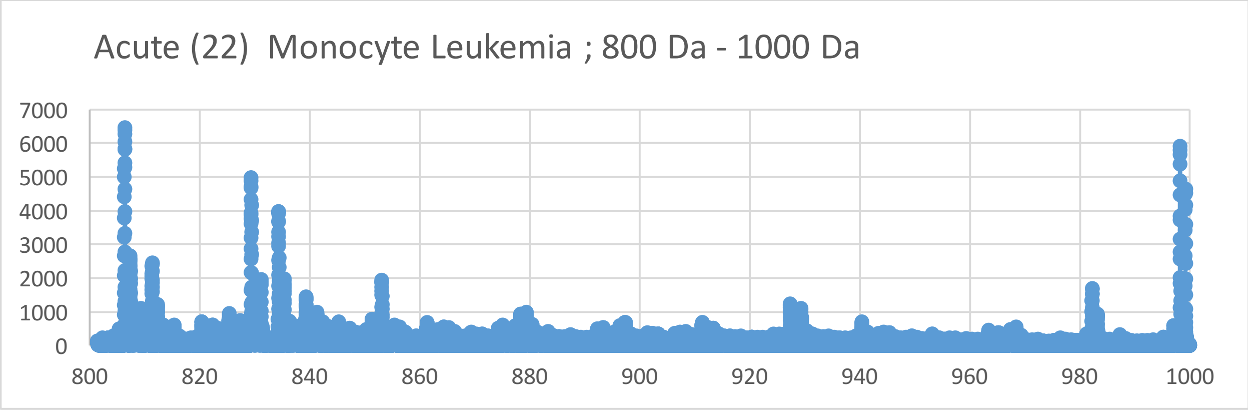
Acute (22) Monocyte Leukemia ; 800 Da - 1000 Da
7000
6000
5000
4000
3000
2000
1000
0
800
820
840
860
880
900
920
940
960
980
1000

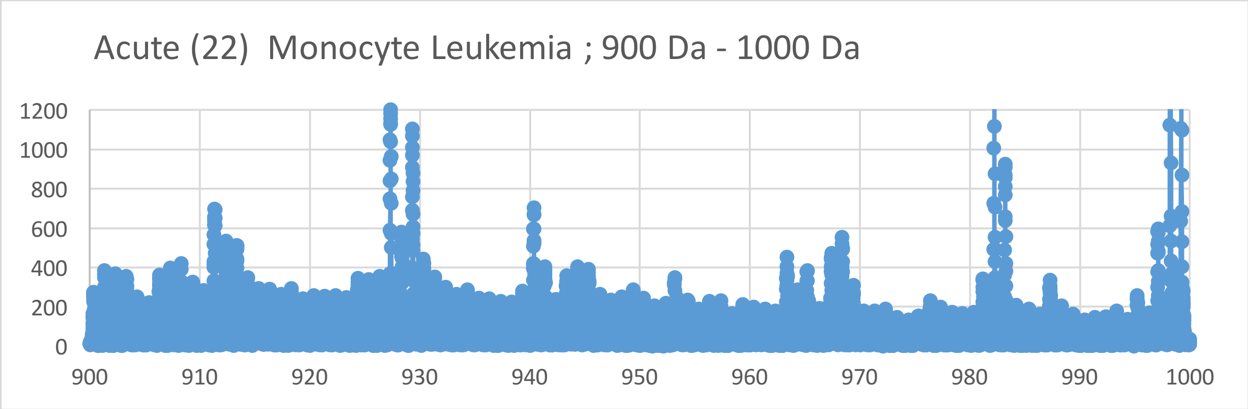
Acute (22) Monocyte Leukemia ; 900 Da - 1000 Da
1200
1000
800
600
400
200
0
900
910
920
930
940
950
960
970
980
990
1000

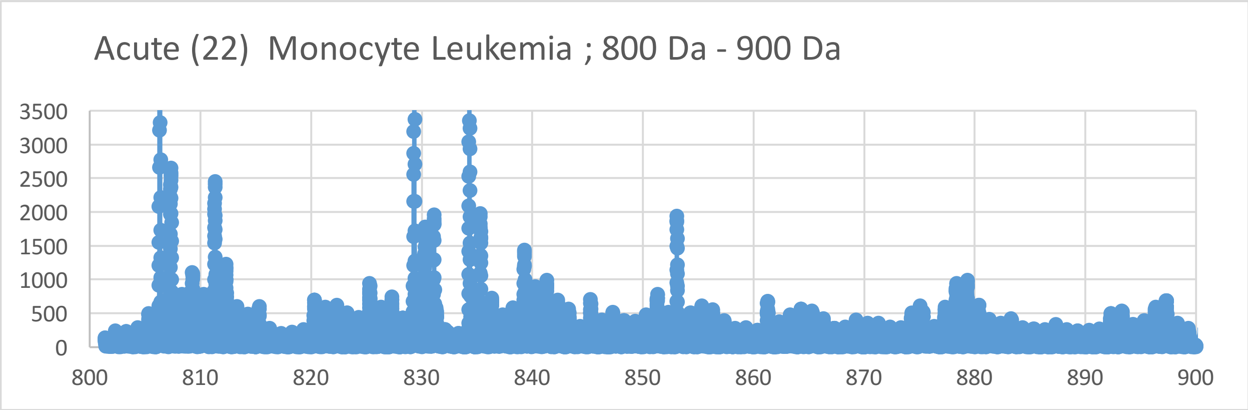
Acute (22) Monocyte Leukemia ; 800 Da - 900 Da
3500
3000
2500
2000
1500
1000
500
0
800
810
820
830
840
850
860
870
880
890
900

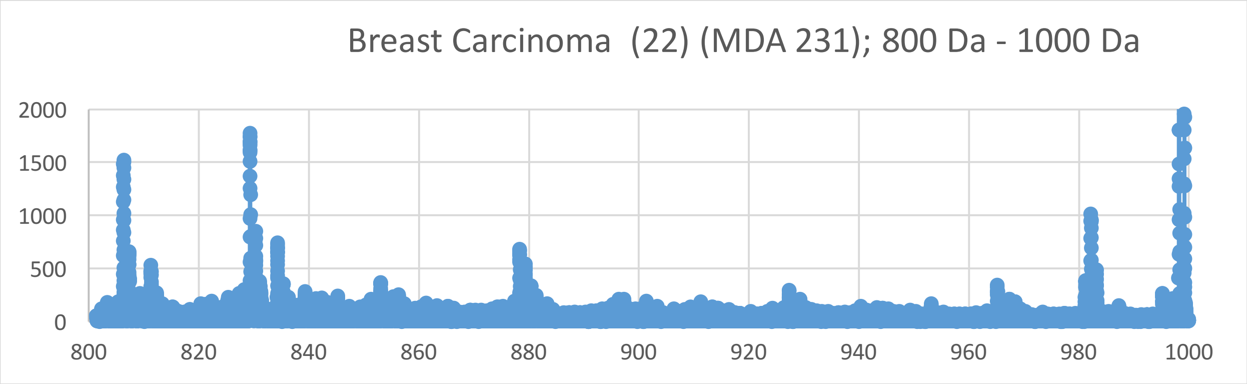
Breast Carcinoma (22) (MDA 231); 800 Da - 1000 Da
2000
1500
1000
500
0
800
820
840
860
880
900
920
940
960
980
1000

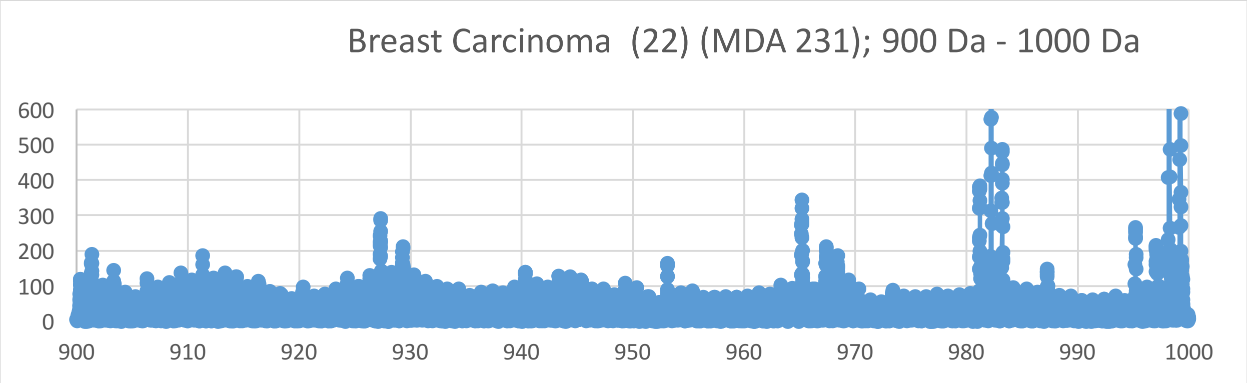
Breast Carcinoma (22) (MDA 231); 900 Da - 1000 Da
600
500
400
300
200
100
0
900
910
920
930
940
950
960
970
980
990
1000

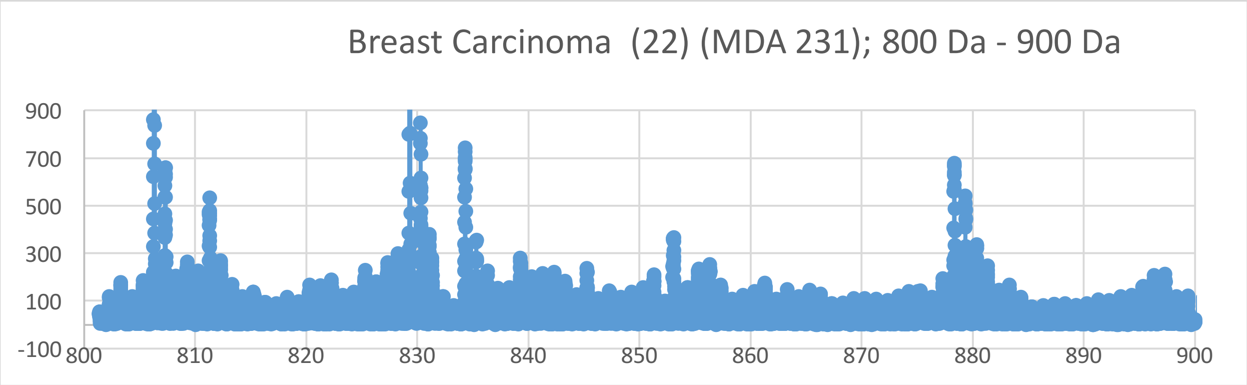
Breast Carcinoma (22) (MDA 231); 800 Da - 900 Da
900
700
500
300
100
-100
800
810
820
830
840
850
860
870
880
890
900

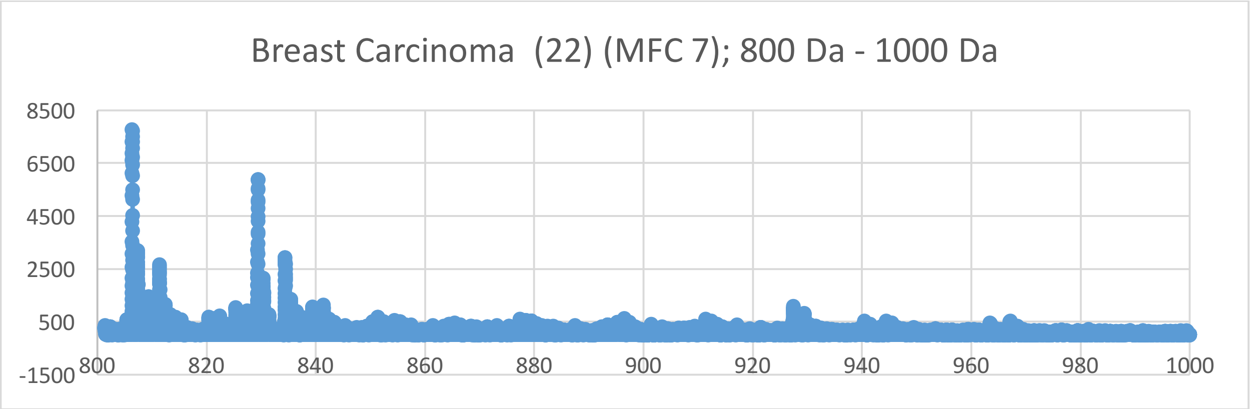
Breast Carcinoma (22) (MFC 7); 800 Da - 1000 Da
8500
6500
4500
2500
500
-1500
800
820
840
860
880
900
920
940
960
980
1000

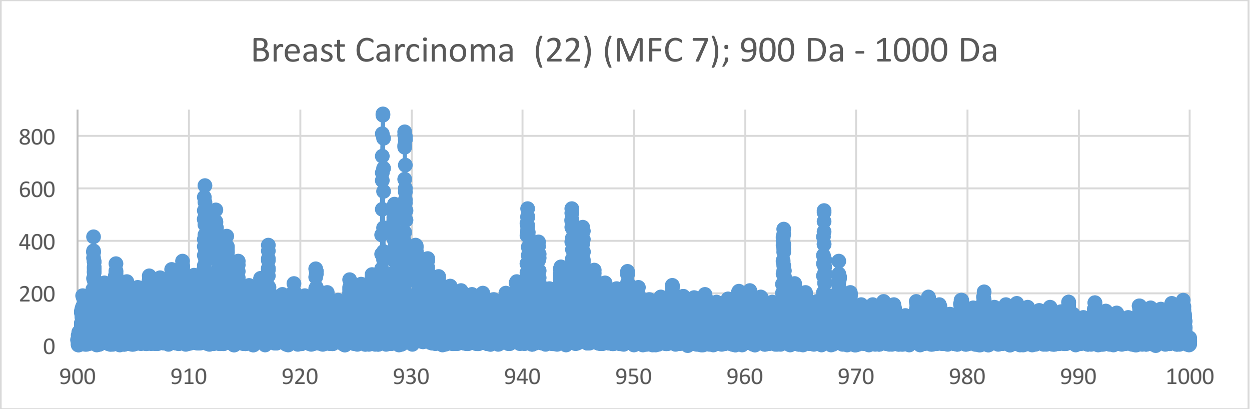
Breast Carcinoma (22) (MFC 7); 900 Da - 1000 Da
800
600
400
200
0
900
910
920
930
940
950
960
970
980
990
1000

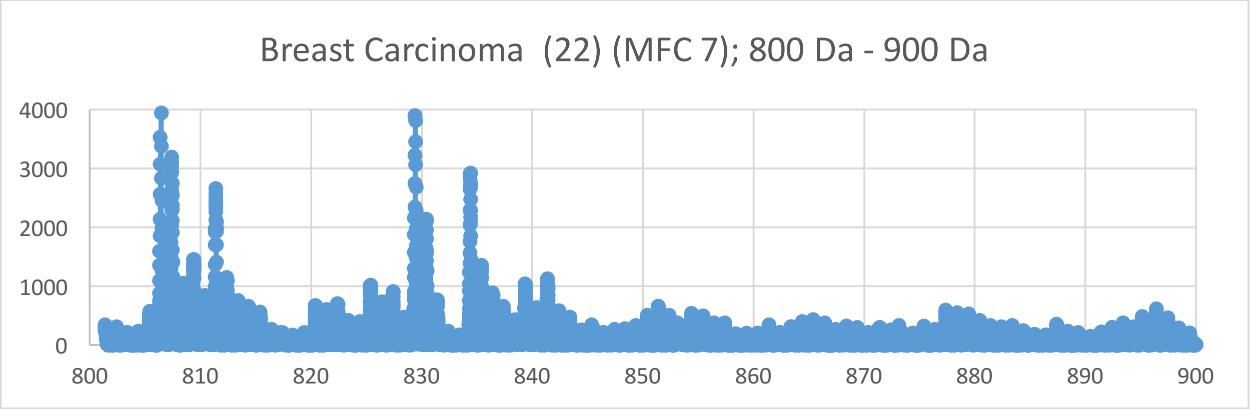
Breast Carcinoma (22) (MFC 7); 800 Da - 900 Da
4000
3000
2000
1000
0
800
810
820
830
840
850
860
870
880
890
900

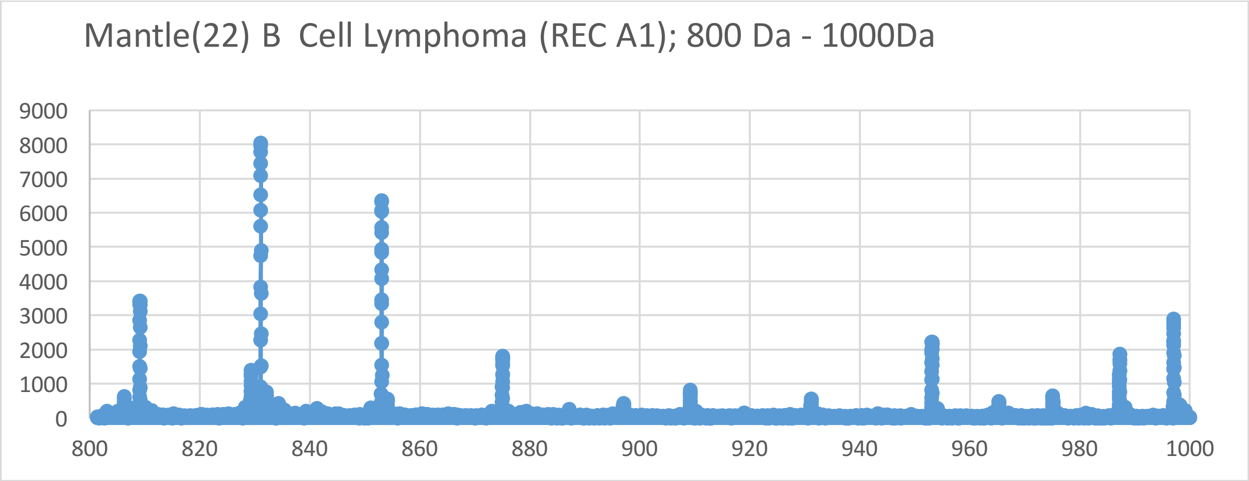
Mantle(22) B Cell Lymphoma (REC A1); 800 Da - 1000Da
9000
8000
7000
6000
5000
4000
3000
2000
1000
0
800
820
840
860
880
900
920
940
960
980
1000

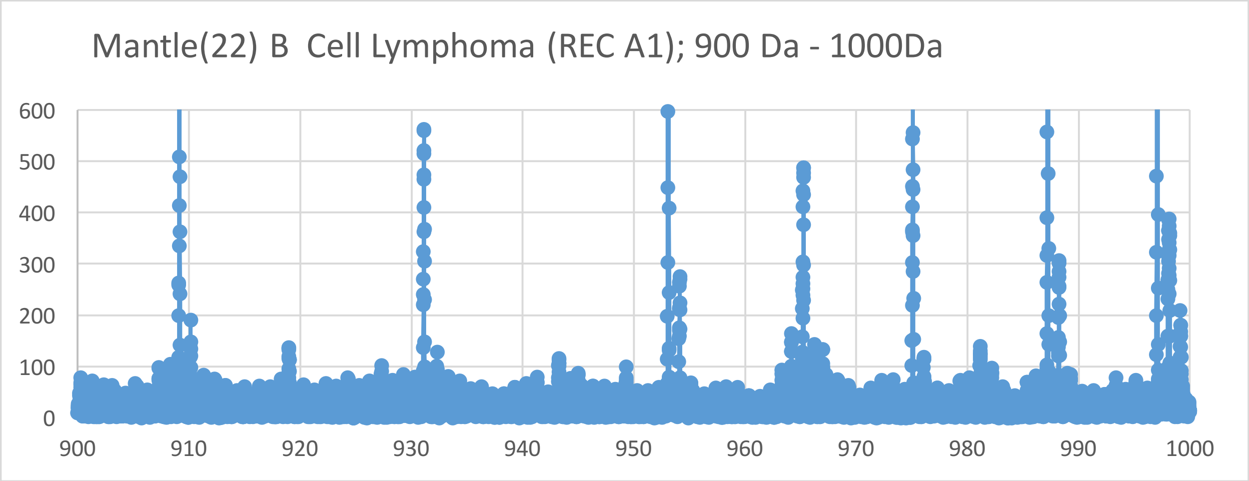
Mantle(22) B Cell Lymphoma (REC A1); 900 Da - 1000Da
600
500
400
300
200
100
0
900
910
920
930
940
950
960
970
980
990
1000

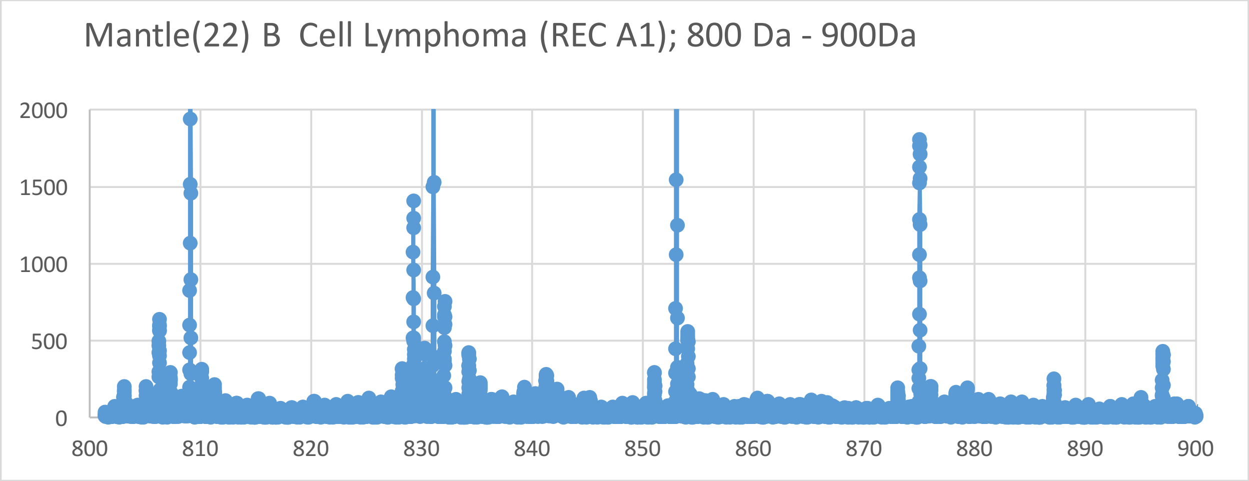
Mantle(22) B Cell Lymphoma (REC A1); 800 Da - 900Da
2000
1500
1000
500
0
800
810
820
830
840
850
860
870
880
890
900

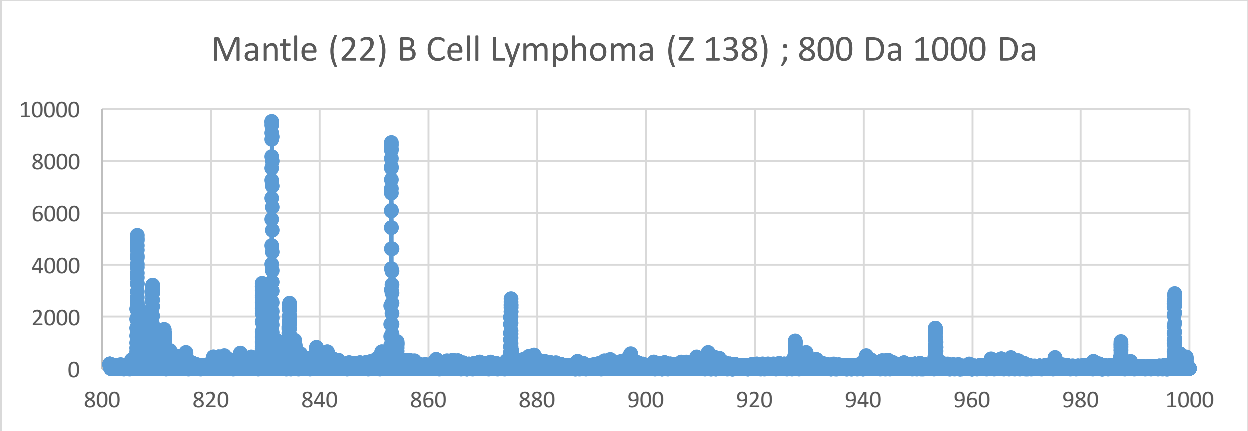
Mantle (22) B Cell Lymphoma (Z 138) ; 800 Da 1000 Da
10000
8000
6000
4000
2000
0
800
820
840
860
880
900
920
940
960
980
1000

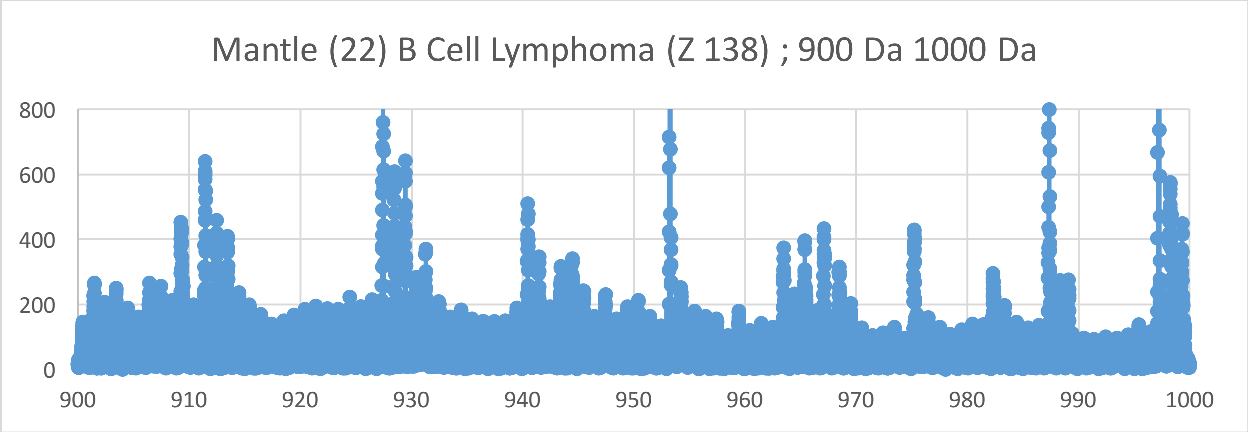
Mantle (22) B Cell Lymphoma (Z 138) ; 900 Da 1000 Da
800
600
400
200
0
900
910
920
930
940
950
960
970
980
990
1000

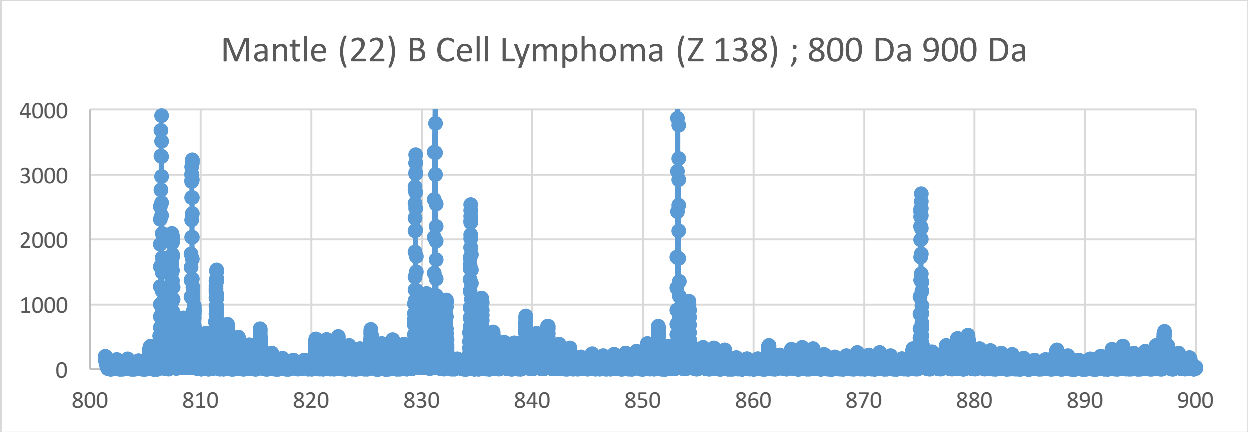
Mantle (22) B Cell Lymphoma (Z 138) ; 800 Da 900 Da
4000
3000
2000
1000
0
800
810
820
830
840
850
860
870
880
890
900

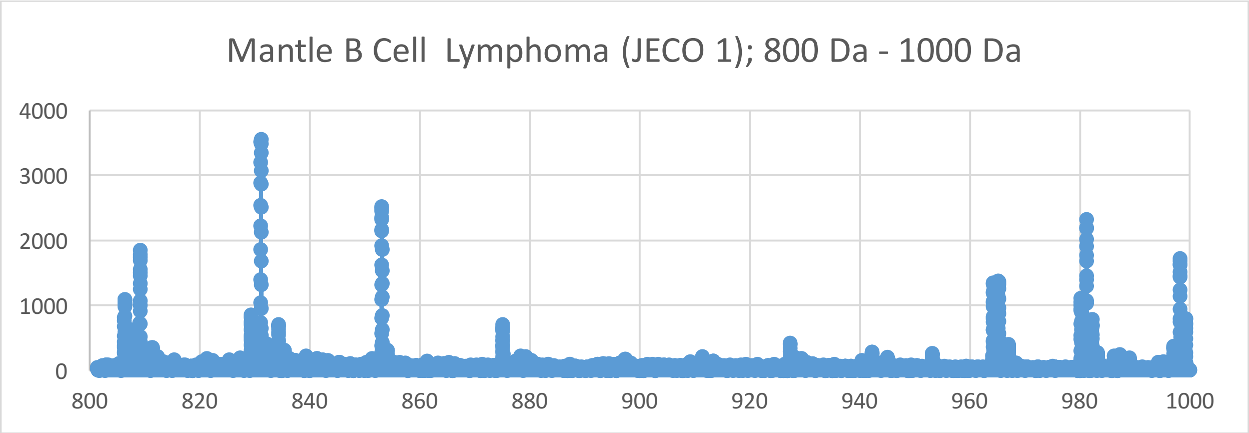
Mantle B Cell Lymphoma (JECO 1); 800 Da - 1000 Da
4000
3000
2000
1000
0
800
820
840
860
880
900
920
940
960
980
1000

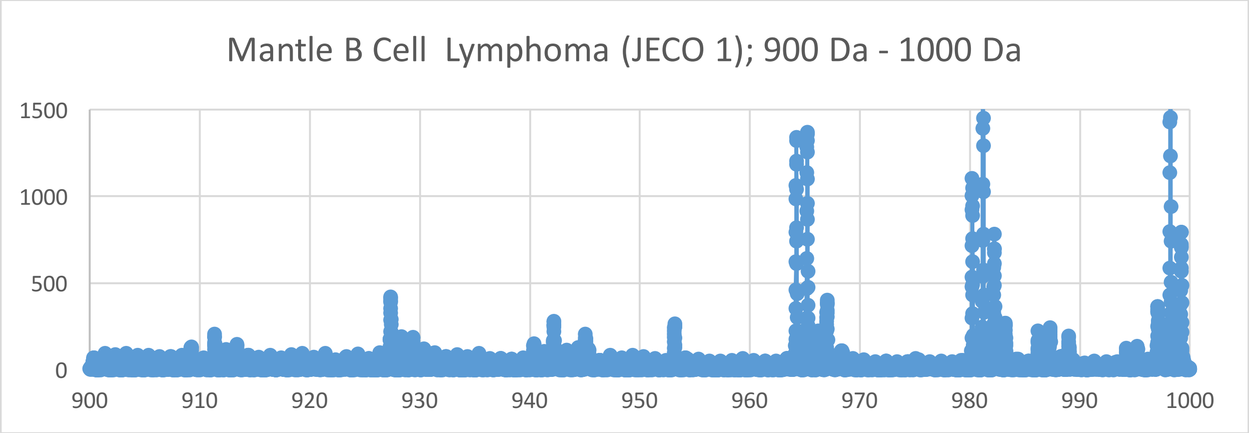
Mantle B Cell Lymphoma (JECO 1); 900 Da - 1000 Da
1500
1000
500
0
900
910
920
930
940
950
960
970
980
990
1000

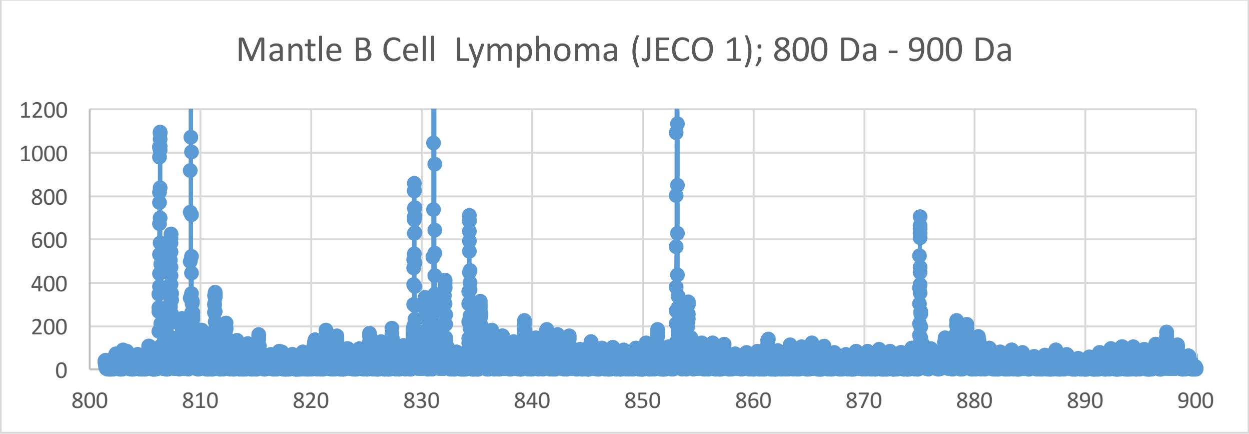
Mantle B Cell Lymphoma (JECO 1); 800 Da - 900 Da
1200
1000
800
600
400
200
0
800
810
820
830
840
850
860
870
880
890
900

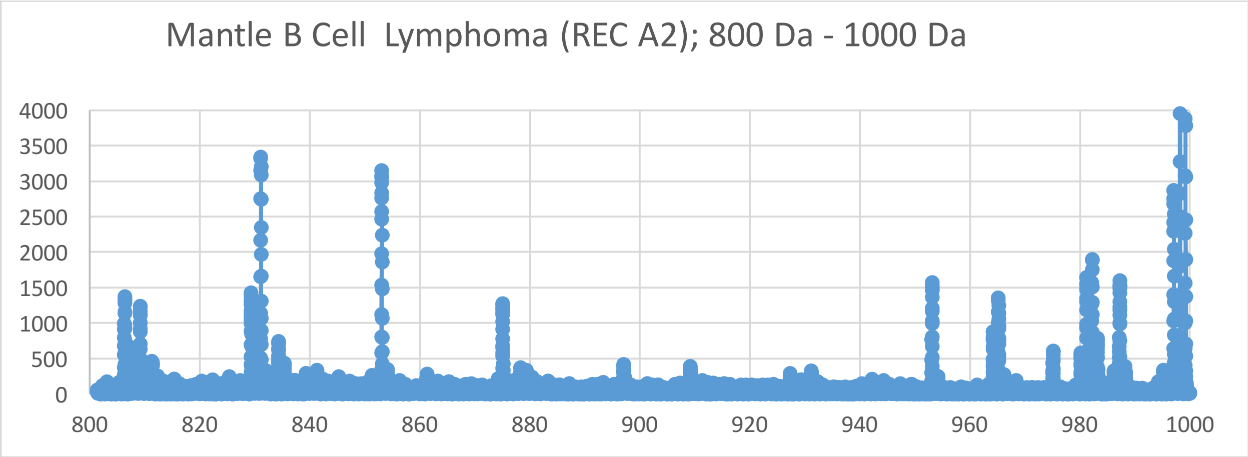
Mantle B Cell  Lymphoma (REC A2); 800 Da - 1000 Da
4000
3500
3000
2500
2000
1500
1000
500
0
800
820
840
860
880
900
920
940
960
980
1000

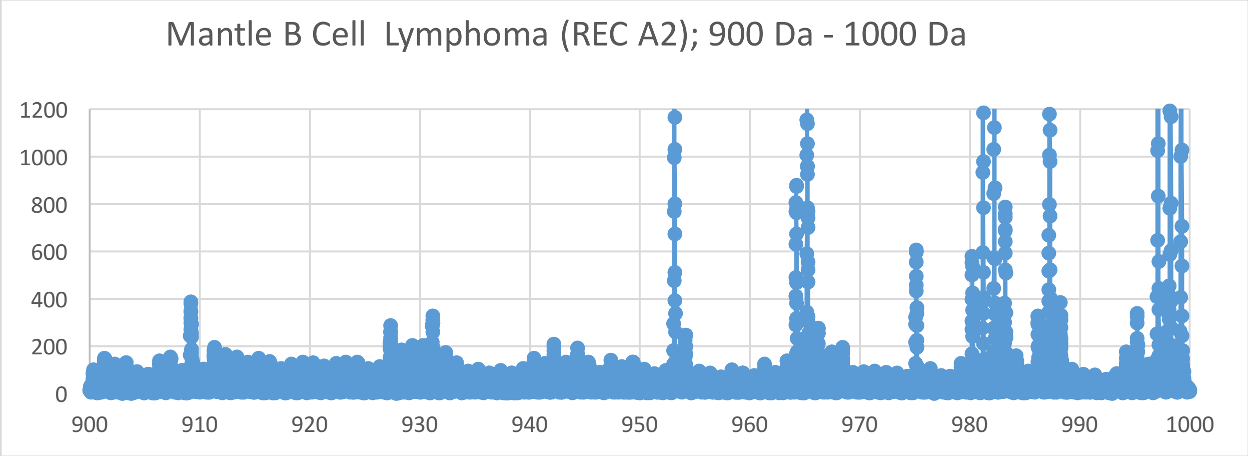
Mantle B Cell  Lymphoma (REC A2); 900 Da - 1000 Da
1200
1000
800
600
400
200
0
900
910
920
930
940
950
960
970
980
990
1000

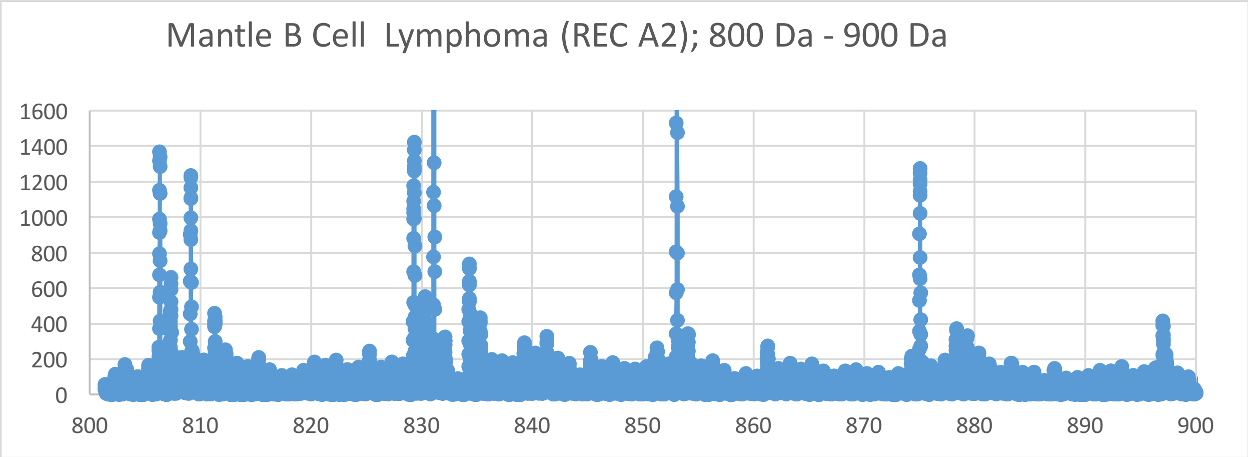
Mantle B Cell  Lymphoma (REC A2); 800 Da - 900 Da
1600
1400
1200
1000
800
600
400
200
0
800
810
820
830
840
850
860
870
880
890
900

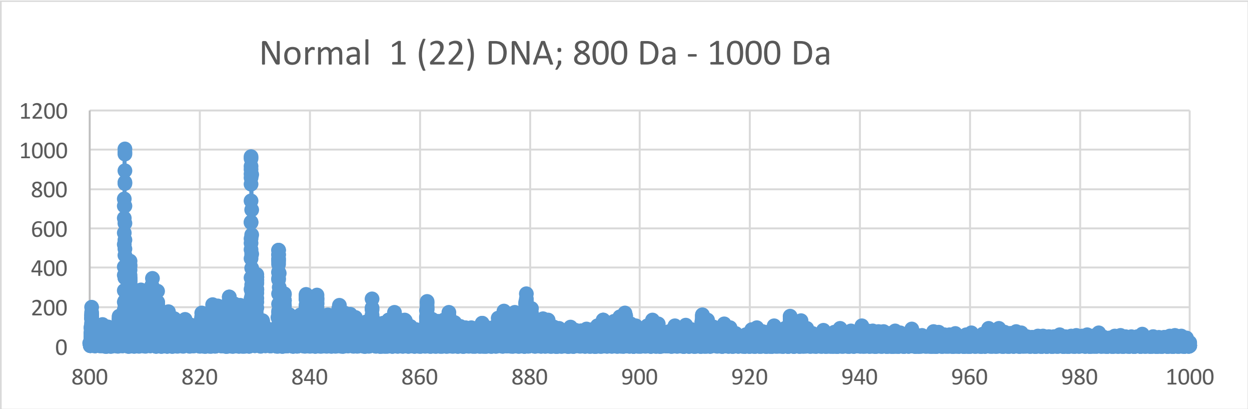
Normal 1 (22) DNA; 800 Da - 1000 Da
1200
1000
800
600
400
200
0
800
820
840
860
880
900
920
940
960
980
1000

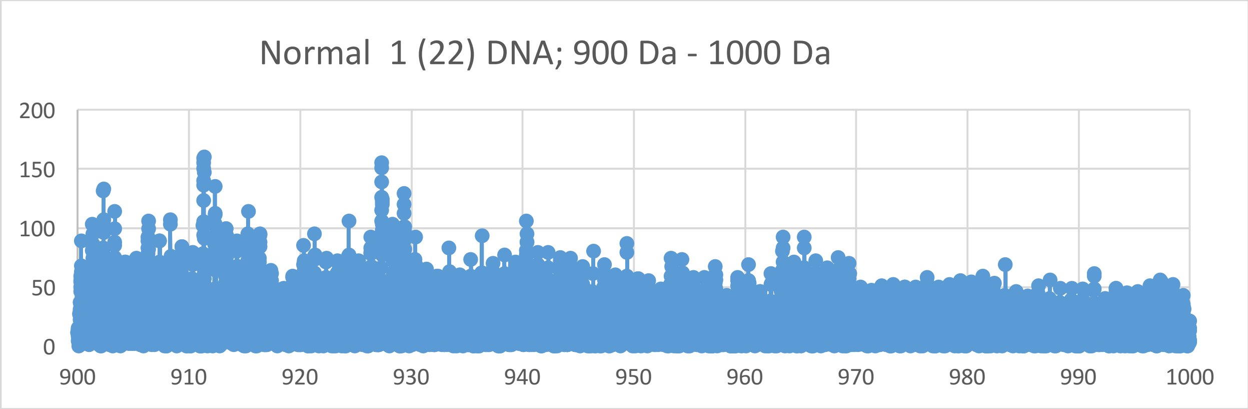
Normal 1 (22) DNA; 900 Da - 1000 Da
200
150
100
50
0
900
910
920
930
940
950
960
970
980
990
1000

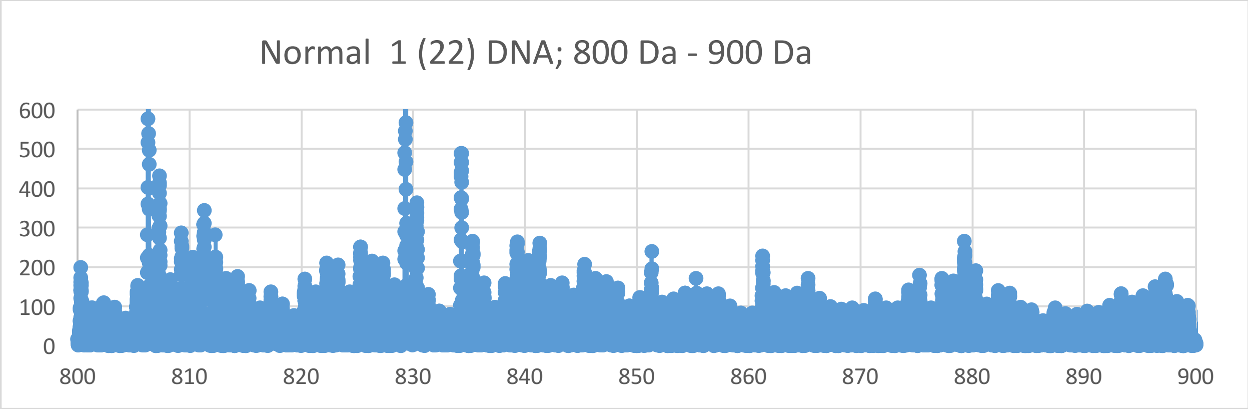
Normal 1 (22) DNA; 800 Da - 900 Da
600
500
400
300
200
100
0
800
810
820
830
840
850
860
870
880
890
900

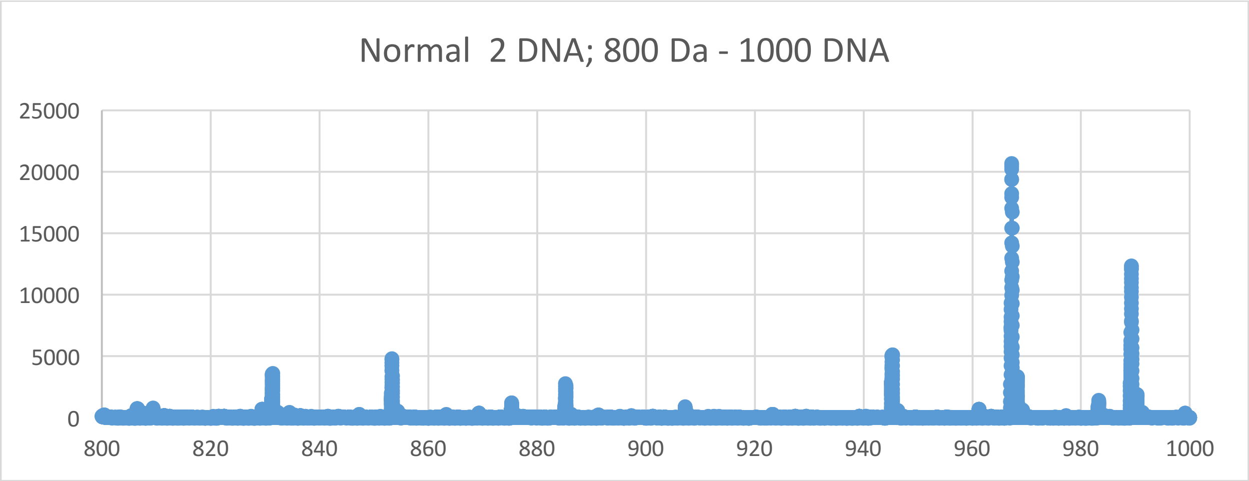
Normal  2 DNA; 800 Da - 1000 DNA
25000
20000
15000
10000
5000
0
800
820
840
860
880
900
920
940
960
980
1000

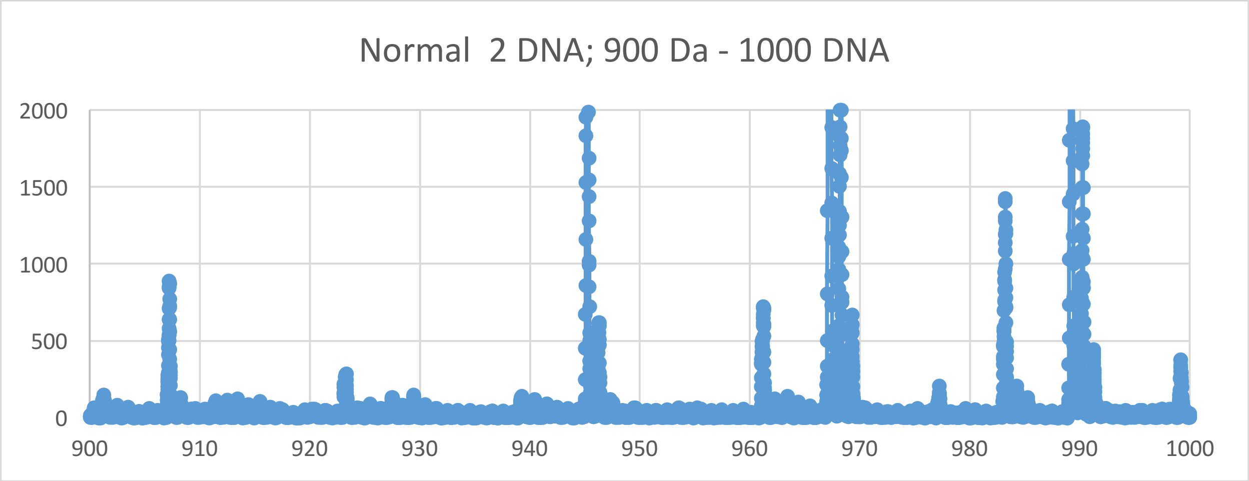
Normal  2 DNA; 900 Da - 1000 DNA
2000
1500
1000
500
0
900
910
920
930
940
950
960
970
980
990
1000

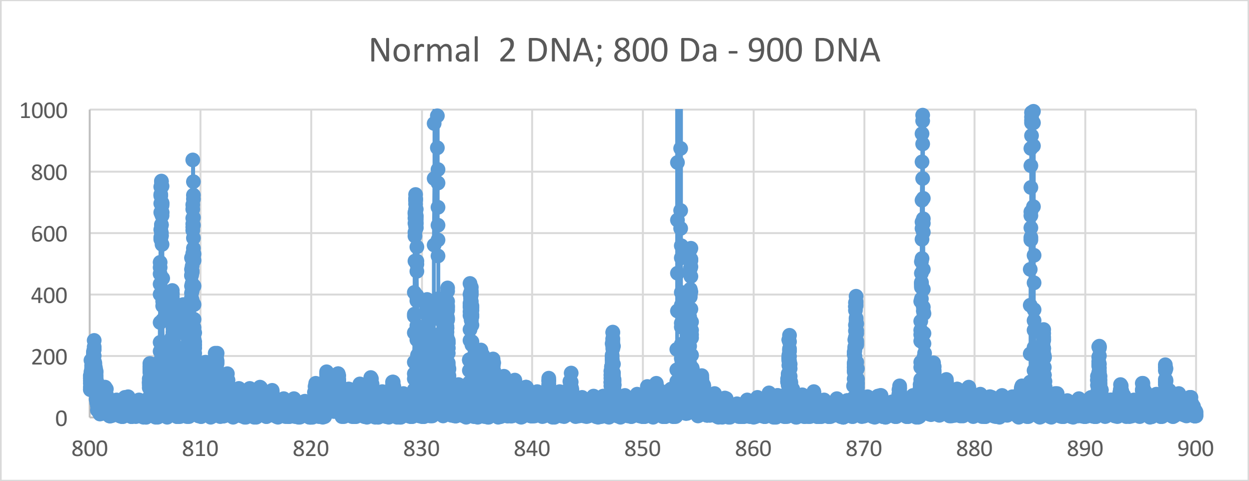
Normal  2 DNA; 800 Da - 900 DNA
1000
800
600
400
200
0
800
810
820
830
840
850
860
870
880
890
900

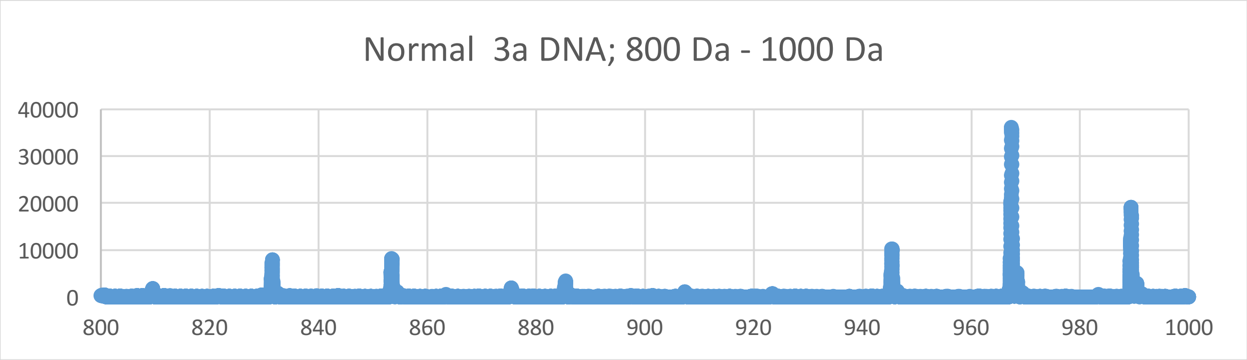
Normal 3a DNA; 800 Da - 1000 Da
40000
30000
20000
10000
0
800
820
840
860
880
900
920
940
960
980
1000

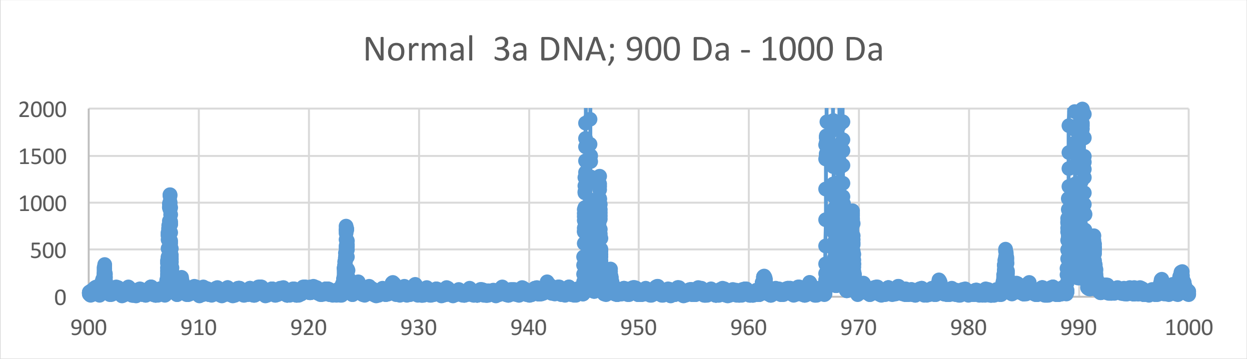
Normal 3a DNA; 900 Da - 1000 Da
2000
1500
1000
500
0
900
910
920
930
940
950
960
970
980
990
1000

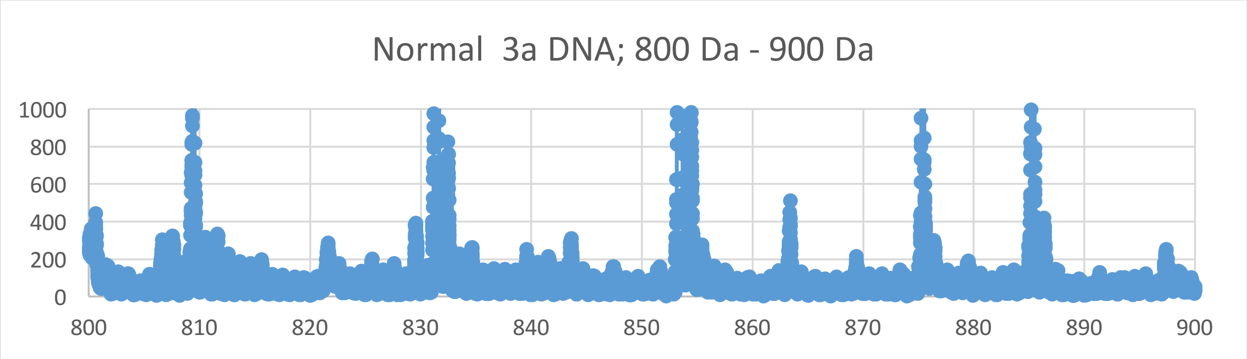
Normal 3a DNA; 800 Da - 900 Da
1000
800
600
400
200
0
800
810
820
830
840
850
860
870
880
890
900

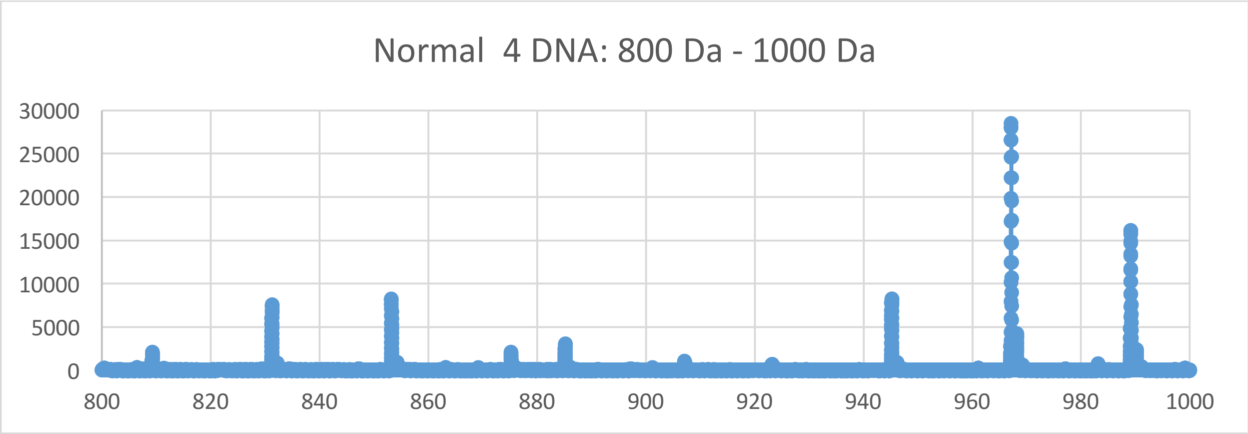
Normal 4 DNA: 800 Da - 1000 Da
30000
25000
20000
15000
10000
5000
0
800
820
840
860
880
900
920
940
960
980
1000

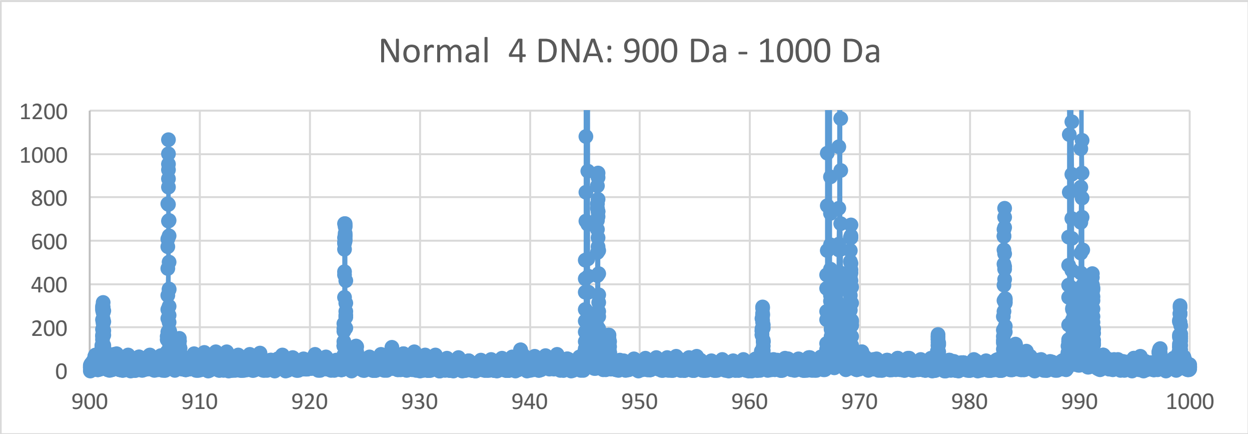
Normal 4 DNA: 900 Da - 1000 Da
1200
1000
800
600
400
200
0
900
910
920
930
940
950
960
970
980
990
1000

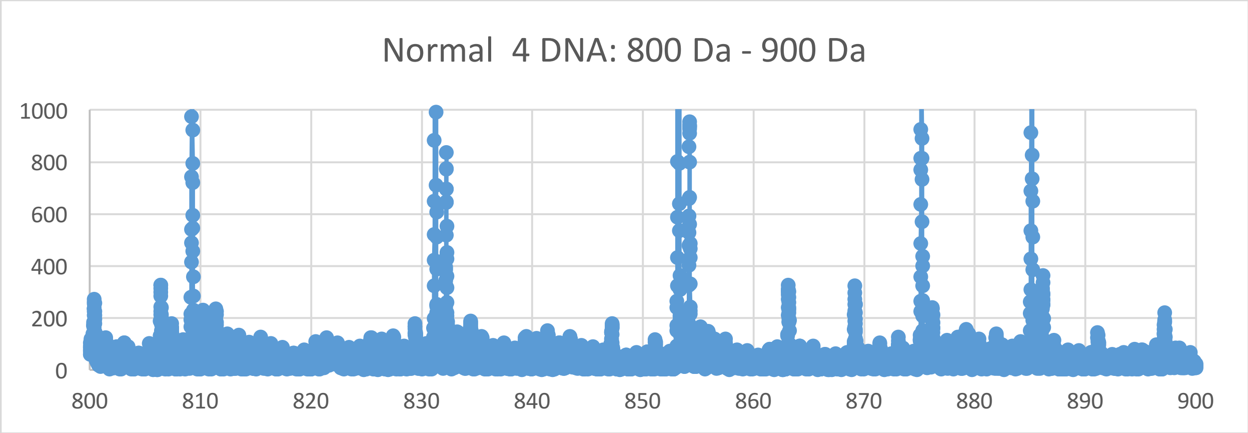
Normal 4 DNA: 800 Da - 900 Da
1000
800
600
400
200
0
800
810
820
830
840
850
860
870
880
890
900
