## Supplementary Materials X for "Oncogenesis and Aging by Isotopic Functionalizations of the Proteins and Nucleic Acids"

X Supplementary Materials – New Mass Spectra from 600 Da -800 Da

Results for Supplementary 10

So for the dinucleotides at $(phos)_4$, the telomeric, mating telomeric and nontelomeric peaks for cancer dramatically change in number and the peaks for normal dinucleotides dramatically change in number for more stable cancer dinucleotides and less stable normal dinucleotides with the $(phos)_4$ from the $(phos)_5$. The telomeres, mating telomeres and nontelomeres favor cancer at $(phos)_4$. But as decrease the phosphate by 1 unit, the dinucleotides change so that the cancer dinucleotides more fragment and break and are stable from the DNA with more number and stabilities for the telomeres, mating telomeres and nontelomeres from $(phos)_4$ to $(phos)_3$. The masses range from 885 Da to 705 Da. See Supplementary Figures X for graphs. It seems that below the saturation range of some number of phosphates, the telomeres, mating telomeres and nontelomeres behave in similar way within cancer and normal DNA. So this suggests cancer may be favored and distinguished from normal by changing saturated phosphorylations from $(phos)_4$ to $(phos)_3$. The telomeric, mating telomeric and nontelomere peaks for cancer slightly increase with these dephosphorylations of the dinucleotides. The dephosphorylations from $(phos)_4$ to $(phos)_3$ caused the resulting more intense and stable cancer dinucleotides to be more telomeric, mating telomeric and nontelomeric dinucleotides and slightly lesser isotopically enrichments for telomeres, mating telomeres and nontelomeres. It is important to consider that there are more intense peaks for the dinucleotides relative to the trinucleotides and tetranucleotides for the saturated phosphate units. For the dinucleotides the intensities of cancer increased and the intensities of normal fragments decreased from $(phos)_4$ to $(phos)_{3.}$ The dephosphorylations of telomeric, mating telomeric and nontelomeric dinucleotides from $(phos)_4$ to $(phos)_3$ slightly decreases isotopic enrichments of heavy isotopes of $^{15}N$, $^{13}C$ and $^{17}O$. The dephosphorylations from $(phos)_4$ to $(phos)_3$ also caused the telomeric, mating telomeric and nontelomeric dinucleotides to be more intense in cancer DNA and slightly less isotopically enriched. It seems the dephosphorylations are related to the isotopic enrichments and depletions; but the telomeres, mating telomeres and nontelomeres are hugely more intense in cancer and slightly less isotopically enriched. The telomeric, mating telomeric and nontelomeric dinucleotides of cancer tended to also be slightly less isotopically enriched so it seems the cancer stabilities of the telomeric, mating telomeric and nontelomeric dinucleotides are related to their slightly lesser isotopic enrichments from the dephosphorylations from $(phos)_4$ to $(phos)_3$ and the mutations purely driven by dephosphorylations and enrichments. Enrichments weaken the mating telomeric normal dinucleotides while strongly strenthening cancerous telomeric dinucleotides. The more intense cancer telomeric, mating telomeric and nontelomeric dinucleotides were also slightly less isotopically enriched, but more intense cancerous telomeric, mating telomeric and nontelomeric dinucleotides were less enriched isotopically. The dephosphorylations from $(phos)_4$ to $(phos)_3$ increased the isotopic enrichments of the cancer for the telomeric, nontelomeric and mating telomeric dinucleotides. Such seems compatible for replications and transcriptions as stable telomeres seem compatible to unstable mating telomeres for the unravelings and constructions of DNA mating telomeres onto the building templates of the telomeres. But in this case if all normal telomeric, mating telomeric and nontelomeric dinucleotides are enriched and unstable then the replications, transcriptions and translations cannot occur normally. The dephosphorylations from $(phos)_4$ to $(phos)_3$ caused methylations to cause slightly less cancer intensities and stabilities but lowered isotopic enrichments for telomeres and mating telomeres. But the methylations of the nontelomeres caused negligible alterations cancer stabilities and isotopic enrichments. The methylations of the nontelomeres slightly decreased cancer intensities and stabilities with negligible enrichments. It seems from these data that during biological processes the dephosphorylations and stabilizations of normal or vice versa dephosphorylations and stabilizations of cancer are or can be countered by methylations (and isotopic enrichments) and destabilizations of the normal for cancer or vice versa dephosphorylations and destabilizations of the cancer by methylations as this research discovers coupled phosphorylations and methylations (and isotopic enrichments) or coupled dephosphorylations and methylations (and isotopic enrichments) with involved greater enrichments for normal biological functions and explaining cancer! The dephosphorylations from $(phos)_4$ to $(phos)_3$ caused acetylations to cause more cancer stabilities for telomeres, mating telomeres and nontelomeres with strong isotopic enrichments and strong cancer stabilities for telomeres with strong enrichments of heavy isotopes. The acetylations from $(phos)_4$ to $(phos)_3$ caused more cancer stabilities and intensities for nontelomeres with strongly more enrichments. This contrast in mating telomeres and telomeres with telomeres from the acetylations may reflect the developments of cancer as the telomeres fragment but the mating telomeres and nontelomeres more hold together for damaged DNA for cancer.

So for the dinucleotides at $(phos)_3$, the telomeric, mating telomeric and nontelomeric peaks for cancer negligibly change in number and the peaks for normal strongly change in number for more similar stable cancer dinucleotides and less stable normal dinucleotides with the $(phos)_3$ as was for $(phos)_4$. The telomeres, mating telomeres and nontelomeres favor cancer at $(phos)_3$. But as decrease the phosphate by 1 unit, the dinucleotides change so that the normal dinucleotides more fragment and break and are stable from the DNA with more number and stabilities for the telomeres and mating telomeres $(phos)_3$ to $(phos)_2$. The masses range from 600 Da to 800 Da. See Supplementary Figures X for graphs. It seems that below the saturation range of some number of phosphates, the telomeres, mating telomeres and nontelomeres behave in different ways within cancer and normal DNA. So this suggests normal may be favored and distinguished from cancer by changing saturated phosphorylations from $(phos)_3$ to $(phos)_2$. The mating telomeric peaks for normal increase with these dephosphorylations of the dinucleotides with more isotopic enrichments. The dephosphorylations from $(phos)_3$ to $(phos)_2$ caused the resulting more intense and stable normal dinucleotides to be more telomeric and more isotopically enriched for telomeres. The nontelomeric dinucleotides had change in intensities and isotopic enrichments for the dephosphorylations from $(phos)_3$ to $(phos)_2$ . It is important to consider that there are more intense peaks for the dinucleotides relative to the trinucleotides and tetranucleotides for the saturated phosphate units. For the dinucleotides, the intensities of normal increased and the intensities of cancer fragments decreased from $(phos)_3$ to $(phos)_{2.}$ The dephosphorylations of telomeric and mating telomeric dinucleotides from $(phos)_3$ to $(phos)_2$ increases isotopic enrichments of heavy isotopes of $^{15}N$, $^{13}C$ and $^{17}O$. The dephosphorylations from $(phos)_3$ to $(phos)_2$ also caused the telomeric dinucleotides to be more intense in normal DNA and more isotopically enriched. It seems the dephosphorylations are related to the isotopic enrichments and depletions; but the telomeres, mating telomeres are hugely more intense in normal and more isotopically enriched. The nontelomeric dinucleotides of cancer tended to also be more isotopically enriched so it seems the cancer stabilities of the nontelomeric dinucleotides are related to their greater isotopic enrichments from the dephosphorylations from $(phos)_3$ to $(phos)_2$ and the mutations purely driven by dephosphorylations and enrichments. Enrichments weaken the nontelomeric cancer dinucleotides while strongly strengthening normal, mating telomeric and telomeric dinucleotides. The more intense cancer telomeric and mating telomeric dinucleotides were also slightly more isotopically enriched, but more intense cancerous nontelomeric dinucleotides were more enriched isotopically. The dephosphorylations from $(phos)_3$ to $(phos)_2$ decreased the isotopic enrichments of the cancer for the telomeric and nontelomeric dinucleotides. Such seems compatible for replications and transcriptions as stable telomeres seem compatible to unstable mating telomeres for the unravelings and constructions of DNA mating telomeres onto the building templates of the telomeres. But in this case if all normal telomeric, mating telomeric and nontelomeric dinucleotides are enriched and unstable then the replications, transcriptions and translations cannot occur normally. The dephosphorylations from $(phos)_3$ to $(phos)_2$ caused methylations to cause negligible alterations of normal intensities and stabilities with negligible isotopic enrichments for telomeres, nontelomeres and mating telomeres. It seems for many dephosphorylations that the loss of too many phosphates diminish the effect of methylations for causing or preventing cancer. It seems from this data that during biological processes the dephosphorylations and stabilizations of normal or vice versa dephosphorylations and stabilizations of cancer are or can be countered by methylations (and isotopic enrichments) and destabilizations of the normal for cancer or vice versa dephosphorylations and destabilizations of the cancer by methylations as this research discovers coupled phosphorylations and methylations (and isotopic enrichments) or coupled dephosphorylations and methylations (and isotopic enrichments) with involved greater enrichments for normal biological functions and explaining cancer! The dephosphorylations from $(phos)_3$ to $(phos)_2$ caused acetylations to cause more cancer stabilities for mating telomeres and nontelomeres with strong isotopic enrichments of light $^{15}N$ and slightly more cancer stabilities for telomeres with enrichments of light isotopes. The acetylations from $(phos)_3$ to $(phos)_2$ caused more cancer stabilities and intensities for mating telomeres, telomeres and nontelomeres with strongly more enrichments. This contrast in mating telomeres and telomeres with telomeres from the acetylations may reflect the developments of cancer as the telomeres fragment but the mating telomeres and nontelomeres more hold together for damaged DNA for cancer.

So for the dinucleotides at $(phos)_2$, the telomeric and mating telomeric peaks for cancer decrease in number and the peaks for cancer for more similar stable normal dinucleotides and similar less stable cancer dinucleotides with the $(phos)_2$ as was for $(phos)_3$. The nontelomeres favors normal at $(phos)_2$. But as decrease the phosphate by 1 unit, the dinucleotides change so that the cancer dinucleotides more fragment and break and are stable from the DNA with more number and stabilities for the telomeres, mating telomeres and nontelomeres $(phos)_2$ to

$(phos)_1$ . The masses range from 550 Da to 732 Da. See Supplementary Figures X for graphs. It seems that below the saturation range of some number of phosphates, the telomeres, mating telomeres and nontelomeres behave in similiar way within cancer and normal DNA. So this suggest cancer may be favored and distinguished from normal by decreasing saturated phosphorylations from $(phos)_2$ to $(phos)_1$. The telomeric, mating telomeric and nontelomeric peaks for cancer increase with these dephosphorylations of the dinucleotides. The dephosphorylations from $(phos)_2$ to $(phos)_1$ caused the resulting more intense and stable cancer dinucleotides to be more telomeric, mating telomeric and nontelomeric dinucleotides and less isotopically enrichments for telomeres, mating telomeres and nontelomeres. It is important to consider that there are more intense peaks for the dinucleotides relative to the trinucleotides and tetranucleotides for the saturated phosphate units. For the dinucleotides, the intensities of cancer increased and the intensities of normal fragments decreased from $(phos)_2$ to $(phos)_{1.}$ The dephosphorylations of telomeric, mating telomeric and nontelomeric dinucleotides from $(phos)_2$ to $(phos)_1$ decrease isotopic enrichments of heavy isotopes of $^{15}N$, $^{13}C$ and $^{17}O$. The dephosphorylations from $(phos)_2$ to $(phos)_1$ also caused the telomeric, mating telomeric and nontelomeric dinucleotides to be more intense in cancer DNA and less isotopically enriched. It seems that the dephosphorylations not only diminish effects of methylations but also diminish the ability to enrich or deplete isotopes. It seems the dephosphorylations are related to the isotopic enrichments and depletions; but the telomeres, mating telomeres and nontelomeres are hugely more intense in cancer and less isotopically enriched. The telomeric, mating telomeric and nontelomeric dinucleotides of cancer tended to also be less isotopically enriched so it seems the cancer stabilities of the telomeric, mating telomeric and nontelomeric dinucleotides are related to their lesser isotopic enrichments from the dephosphorylations from $(phos)_2$ to $(phos)_1$ and the mutations purely driven by dephosphorylations and enrichments. Enrichments weaken the nontelomeric, mating telomeric and telomeric cancerous dinucleotides while strongly strengthening normal telomeric, mating telomeric and nontelomeric dinucleotides. The more intense cancer telomeric, mating telomeric and nontelomeric dinucleotides were also slightly less isotopically enriched, so more intense cancerous telomeric, mating telomeric and nontelomeric dinucleotides were less enriched isotopically. The dephosphorylations from $(phos)_2$ to $(phos)_1$ decreased the isotopic enrichments of the cancer for the telomeric, nontelomeric and mating telomeric dinucleotides. Such seems compatible for replications and transcriptions as stable telomeres seem compatible to unstable mating telomeres for the unravelings and constructions of DNA mating telomeres onto the building templates of the telomeres. But in this case if all normal telomeric, mating telomeric and nontelomeric dinucleotides are enriched and unstable then the replications, transcriptions and translations cannot as well occur normally. The dephosphorylations from $(phos)_2$ to $(phos)_1$ caused methylations to greatly cause cancer intensities and stabilities but stronger isotopic enrichments for telomeres, nontelomeres and mating telomeres. It seems from these data that during biological processes the dephosphorylations and stabilizations of normal or vice versa dephosphorylations and stabilizations of cancer are or can be countered by methylations (and isotopic enrichments) and destabilizations of the normal for cancer or vice versa dephosphorylations and destabilizations of the cancer by methylations as this research discovers coupled phosphorylations and methylations (and isotopic enrichments) or coupled dephosphorylations and methylations (and isotopic enrichments) with involved greater enrichments for normal biological functions and explaining cancer! But these differing effects of methylations on cancer and normal DNA are lossed as the dephosphorylations become severe. The phosphorylations are involved with the methylations and the methylations modulates of cancer and normal DNA stabilities. The dephosphorylations from $(phos)_2$ to $(phos)_1$ caused acetylations to cause more normal stabilities for telomeres and mating telomeres with strong isotopic enrichments and negligible cancer stabilities for telomeres with slightly less enrichments of heavy isotopes. The acetylations from $(phos)_2$ to $(phos)_1$ caused more dramatic increase in cancer isotopic enrichments, stabilities and intensities for nontelomeres. This contrast in mating telomeres and telomeres with telomeres from the acetylations may reflect the developments of cancer as the telomeres fragment but the mating telomeres and nontelomeres more hold together for damaged DNA for cancer.

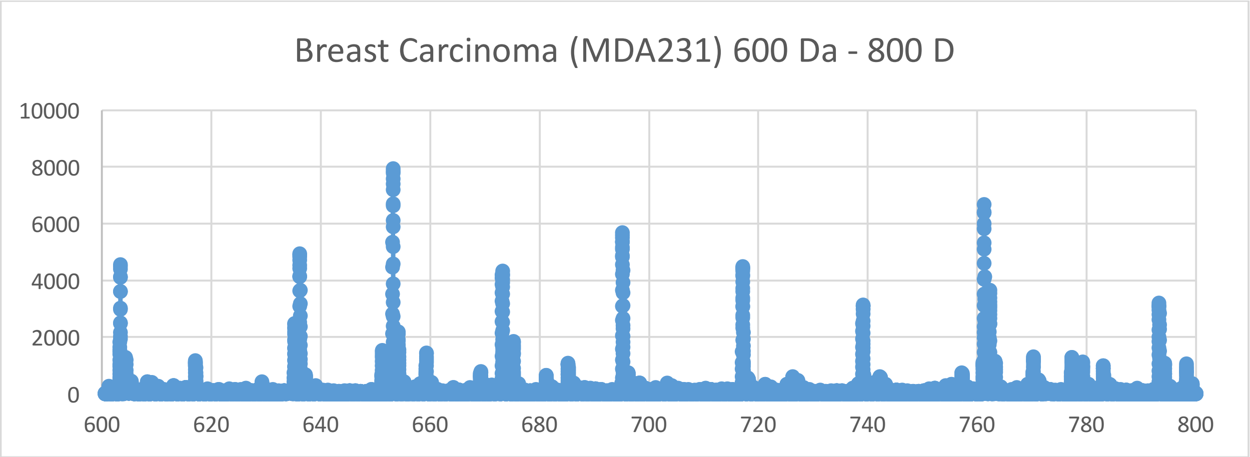
Breast Carcinoma (MDA231) 600 Da - 800 D
10000
8000
6000
4000
2000
0
600
620
640
660
680
700
720
740
760
780
800

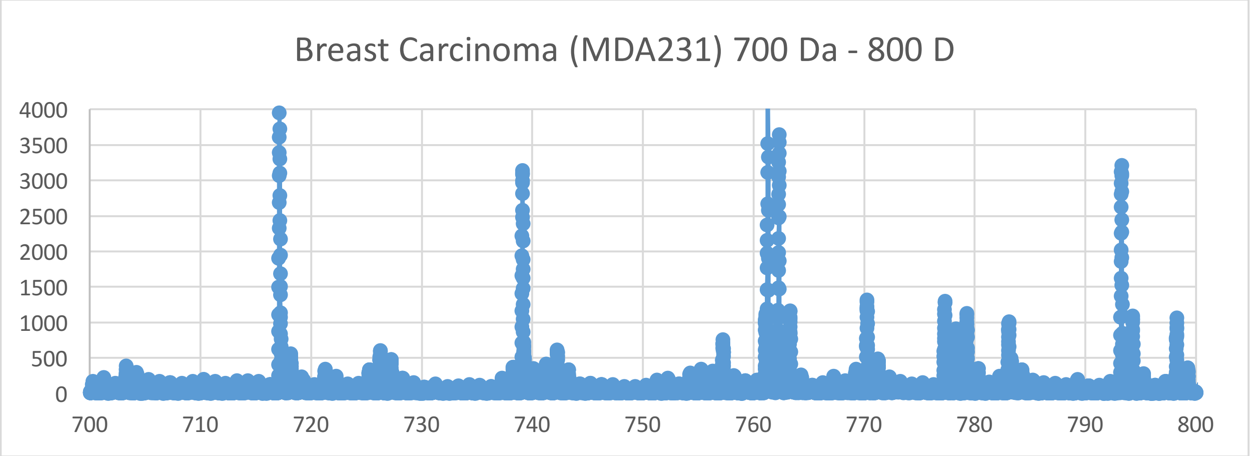
Breast Carcinoma (MDA231) 700 Da - 800 D
4000
3500
3000
2500
2000
1500
1000
500
0
700
710
720
730
740
750
760
770
780
790
800

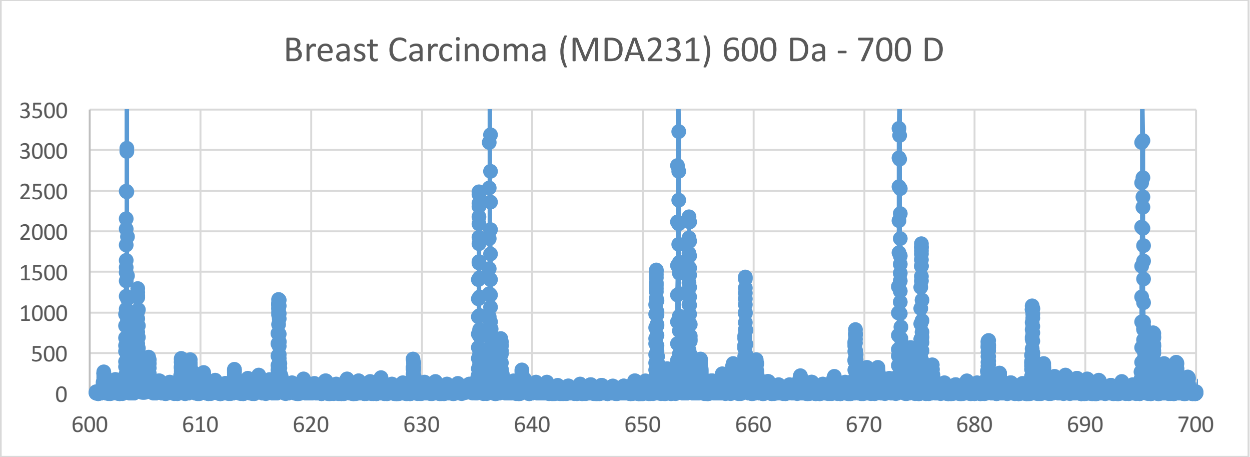
Breast Carcinoma (MDA231) 600 Da - 700 D
3500
3000
2500
2000
1500
1000
500
0
600
610
620
630
640
650
660
670
680
690
700

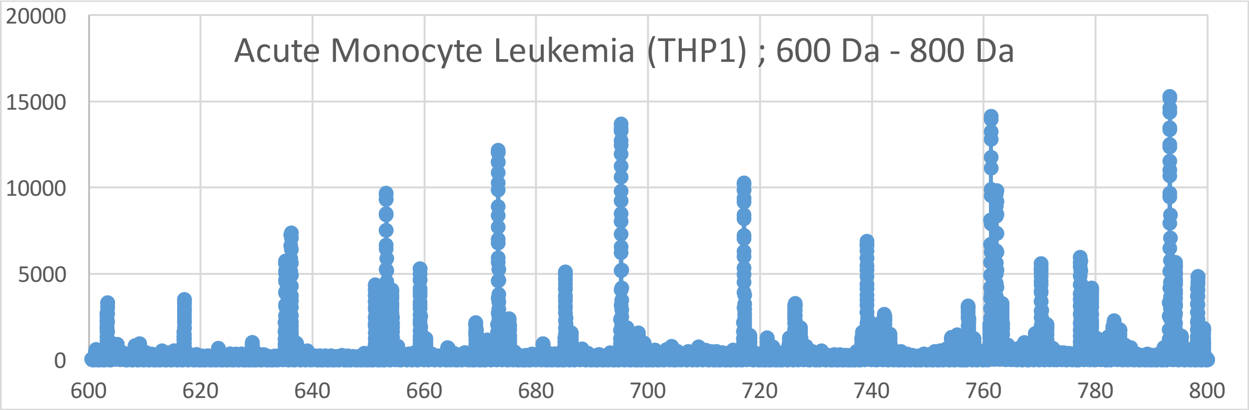

Acute Monocyte Leukemia (THP1) ; 600 Da - 800 Da
20000
15000
10000
5000
0
600
620
640
660
680
700
720
740
760
780
800


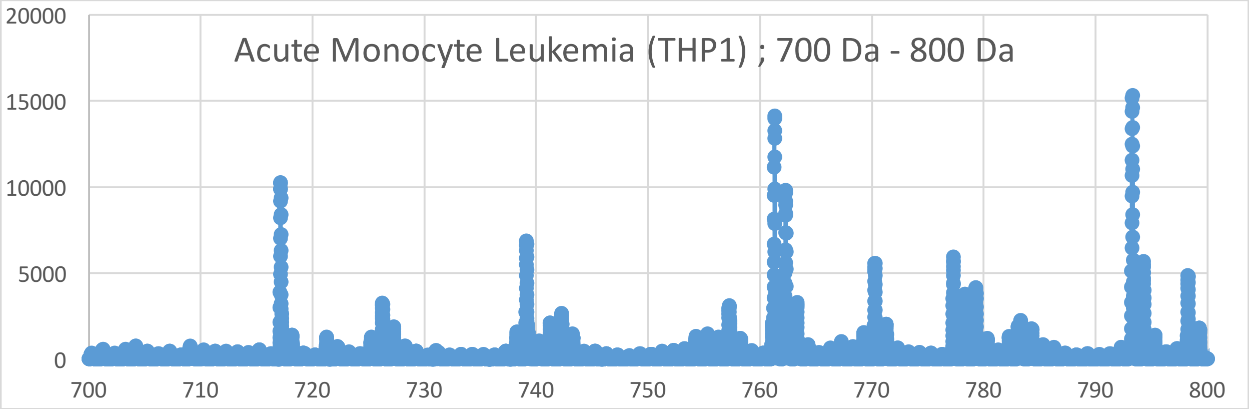

Acute Monocyte Leukemia (THP1) ; 700 Da - 800 Da
20000
15000
10000
5000
0
700
710
720
730
740
750
760
770
780
790
800


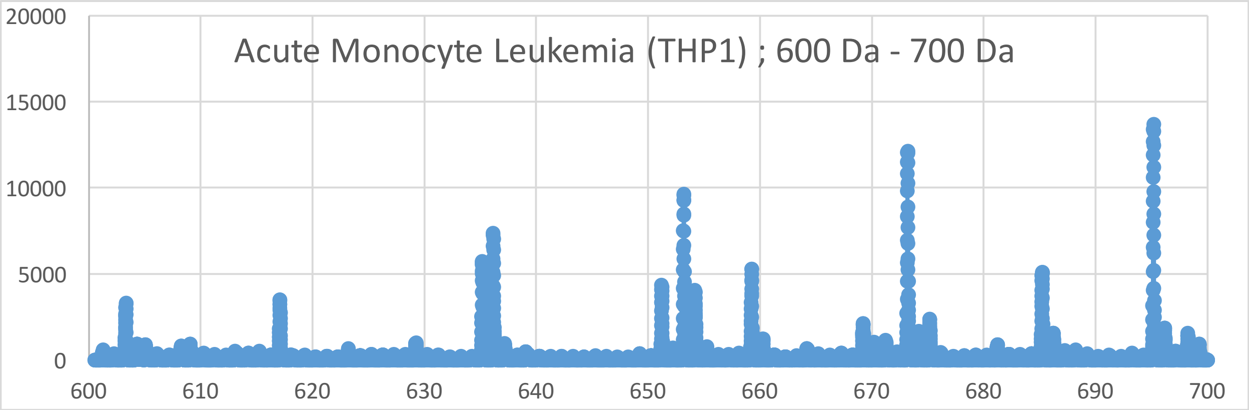

Acute Monocyte Leukemia (THP1) ; 600 Da - 700 Da
20000
15000
10000
5000
0
600
610
620
630
640
650
660
670
680
690
700

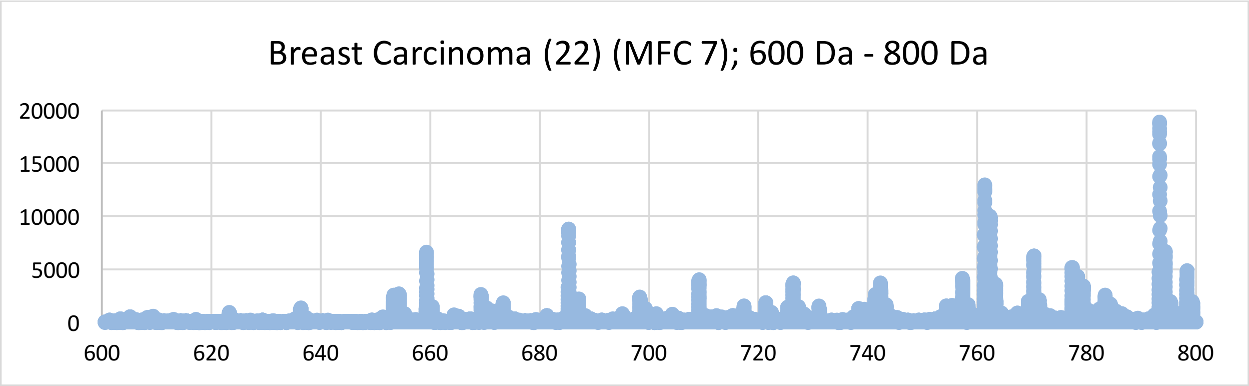
Breast Carcinoma (22) (MFC 7); 600 Da - 800 Da
20000
15000
10000
5000
0
600
620
640
660
680
700
720
740
760
780
800

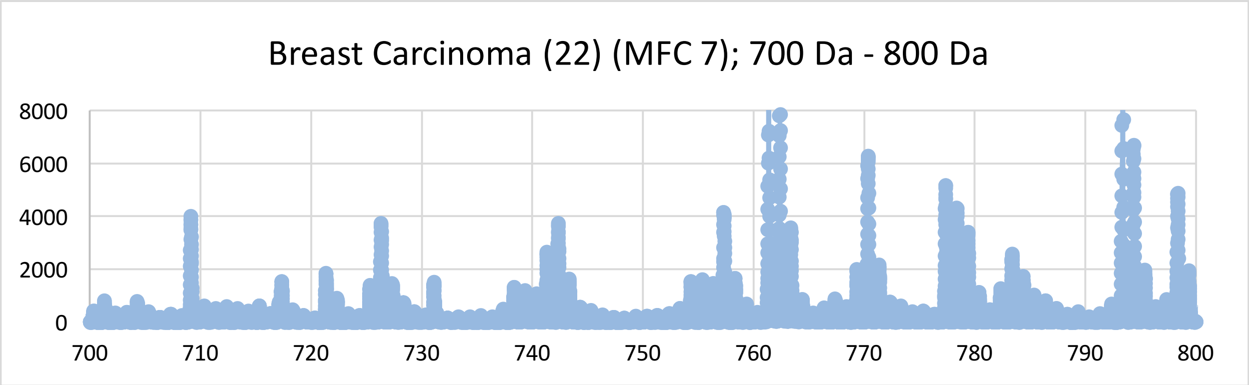
Breast Carcinoma (22) (MFC 7); 700 Da - 800 Da
8000
6000
4000
2000
0
700
710
720
730
740
750
760
770
780
790
800

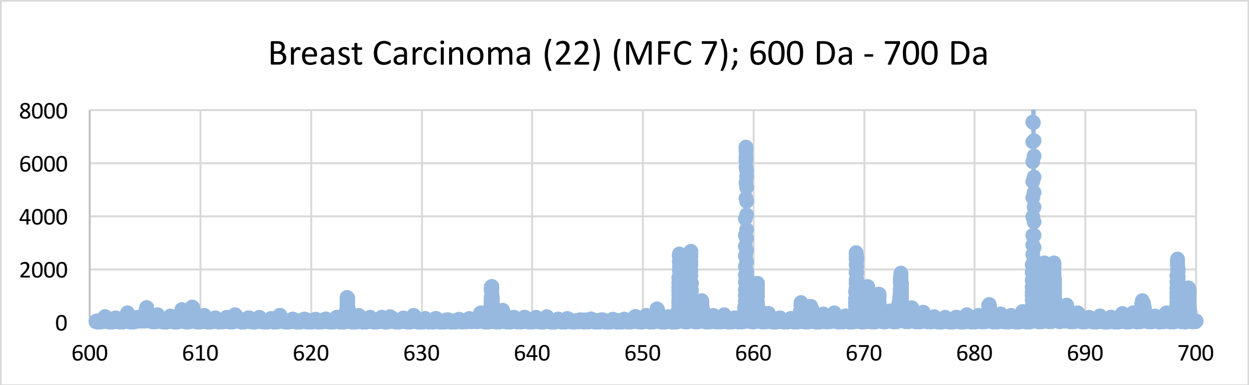
Breast Carcinoma (22) (MFC 7); 600 Da - 700 Da
8000
6000
4000
2000
0
600
610
620
630
640
650
660
670
680
690
700

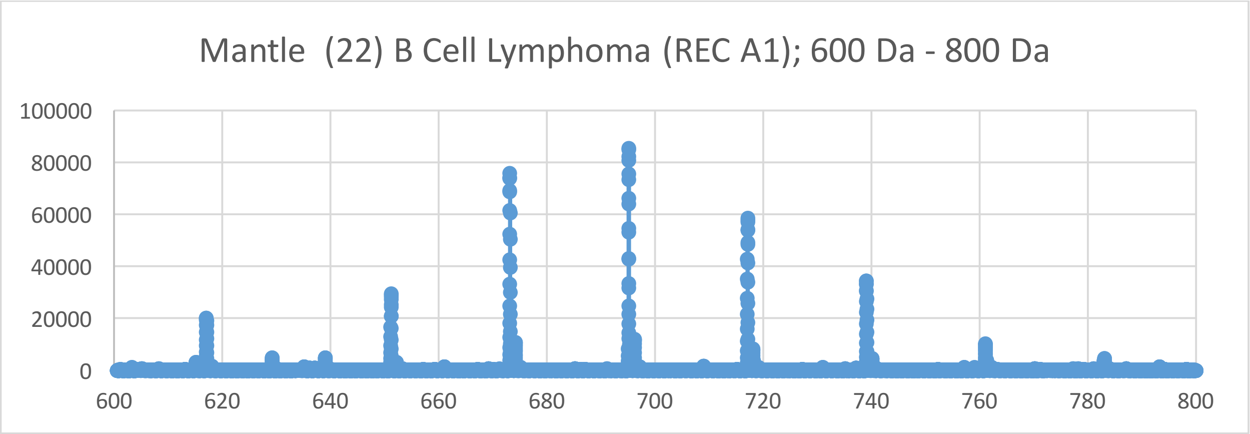

Mantle (22) B Cell Lymphoma (REC A1); 600 Da - 800 Da
100000
80000
60000
40000
20000
0
600
620
640
660
680
700
720
740
760
780
800


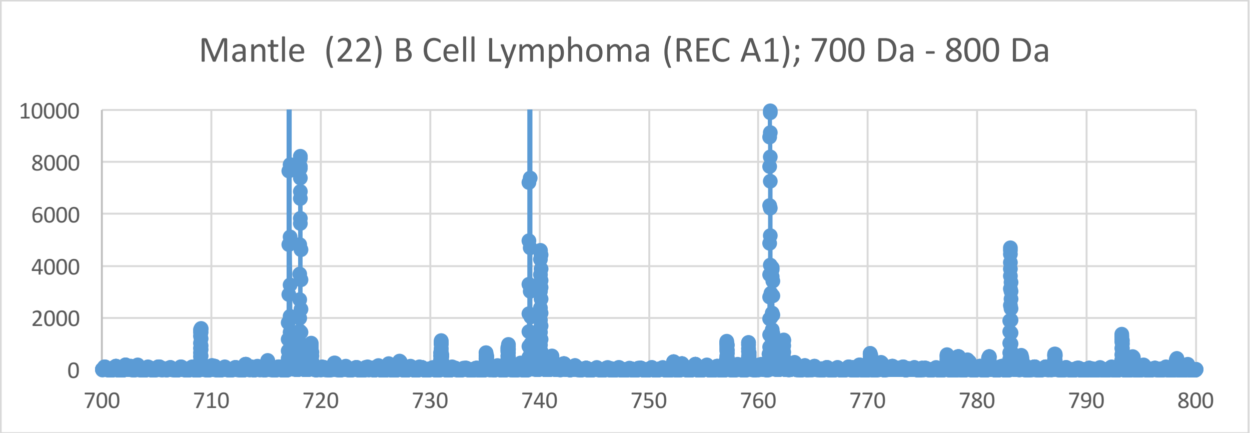

Mantle (22) B Cell Lymphoma (REC A1); 700 Da - 800 Da
10000
8000
6000
4000
2000
0
700
710
720
730
740
750
760
770
780
790
800


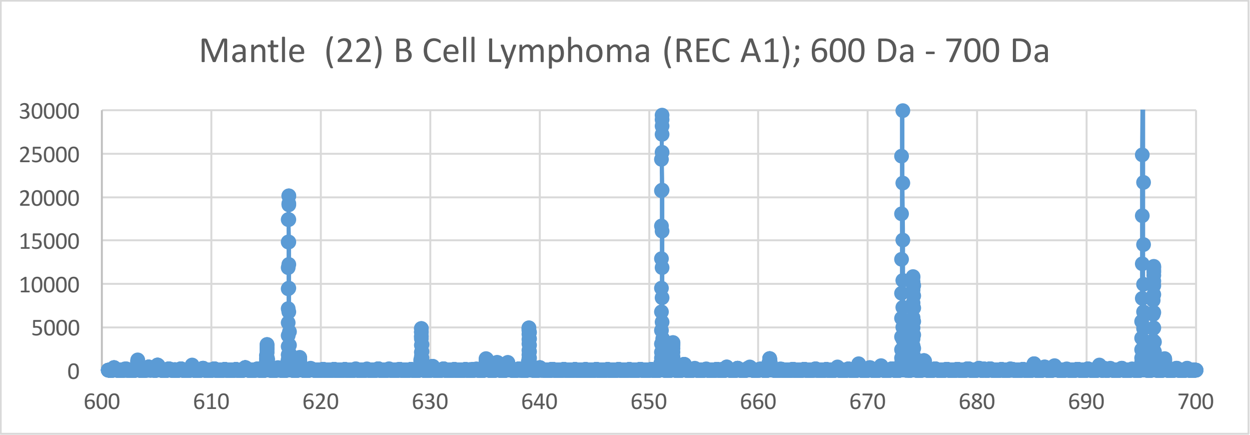

Mantle (22) B Cell Lymphoma (REC A1); 600 Da - 700 Da
30000
25000
20000
15000
10000
5000
0
600
610
620
630
640
650
660
670
680
690
700

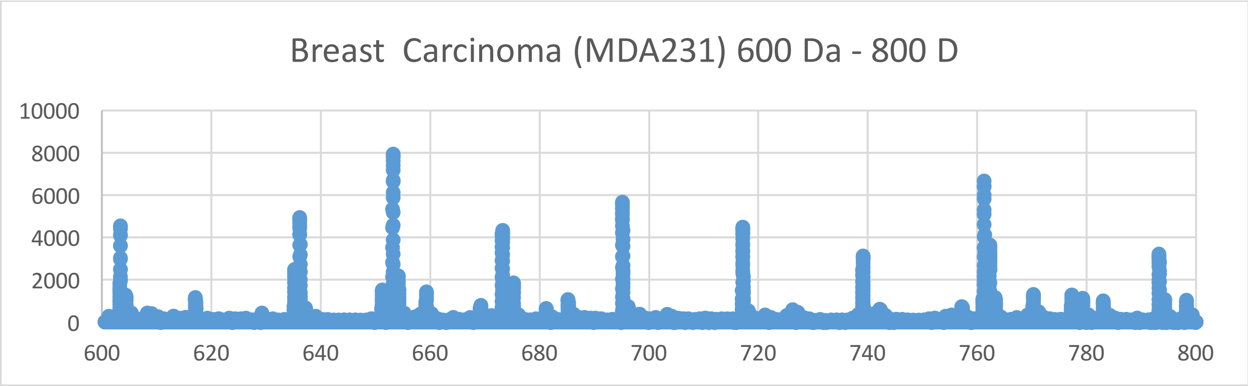
Breast  Carcinoma (MDA231) 600 Da - 800 D
10000
8000
6000
4000
2000
0
600
620
640
660
680
700
720
740
760
780
800

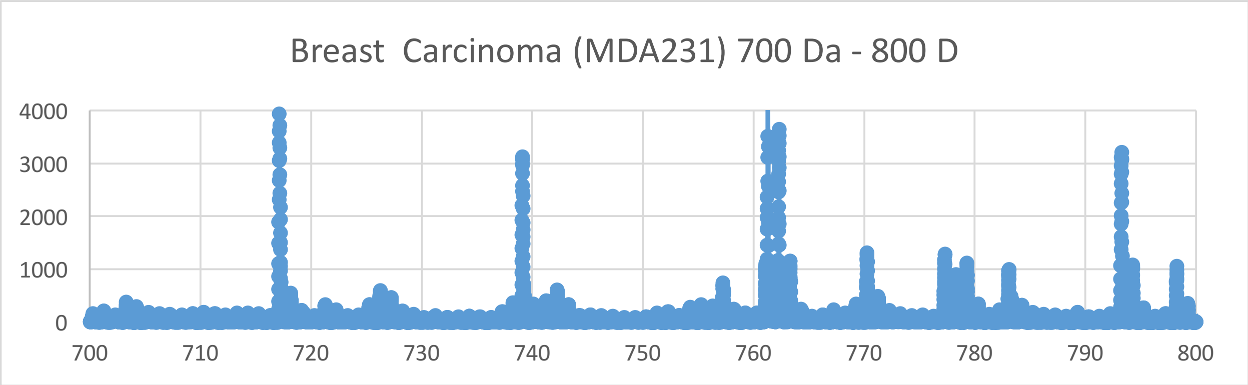
Breast  Carcinoma (MDA231) 700 Da - 800 D
4000
3000
2000
1000
0
700
710
720
730
740
750
760
770
780
790
800

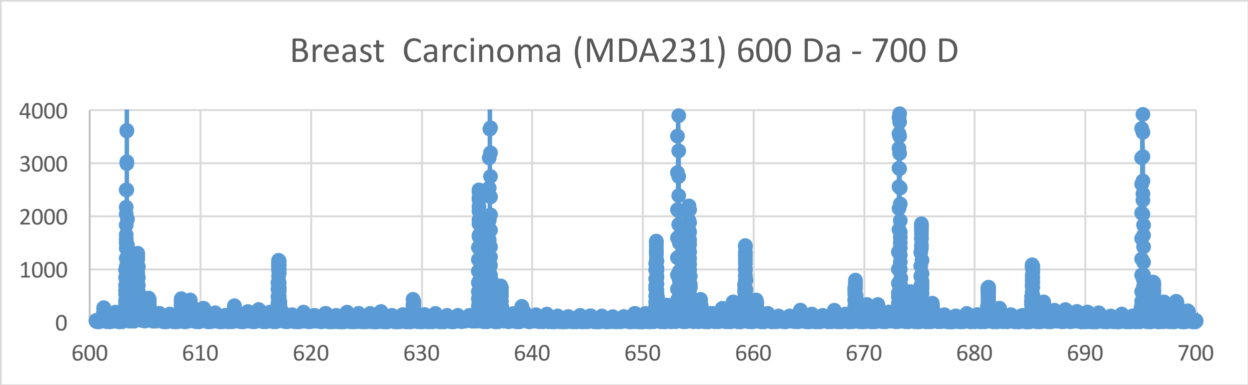
Breast  Carcinoma (MDA231) 600 Da - 700 D
4000
3000
2000
1000
0
600
610
620
630
640
650
660
670
680
690
700

Mantle (2) BCell Lymphoma (JECO 1) ; 600 Da - 800 Da

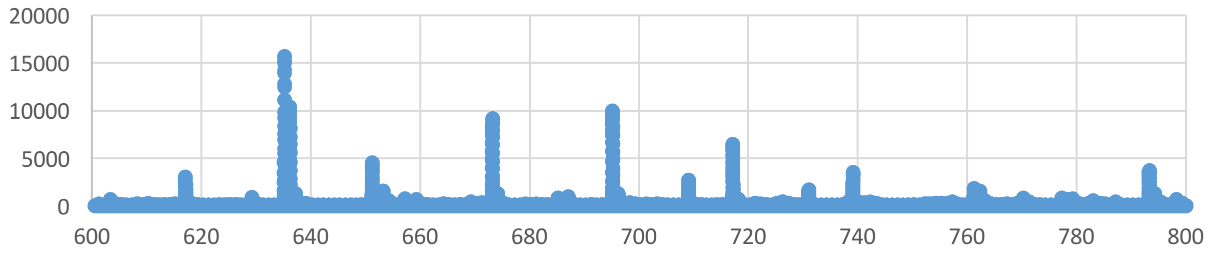


Mantle (2) BCell Lymphoma (JECO 1) ; 700 Da - 800 Da

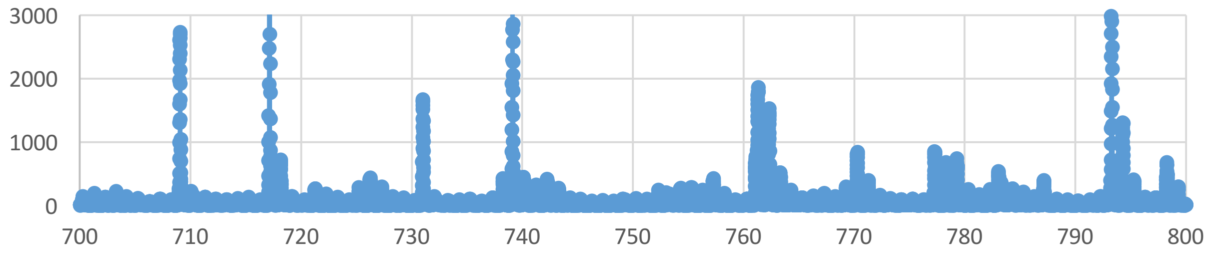


Mantle (2) BCell Lymphoma (JECO 1) ; 600 Da - 700 Da

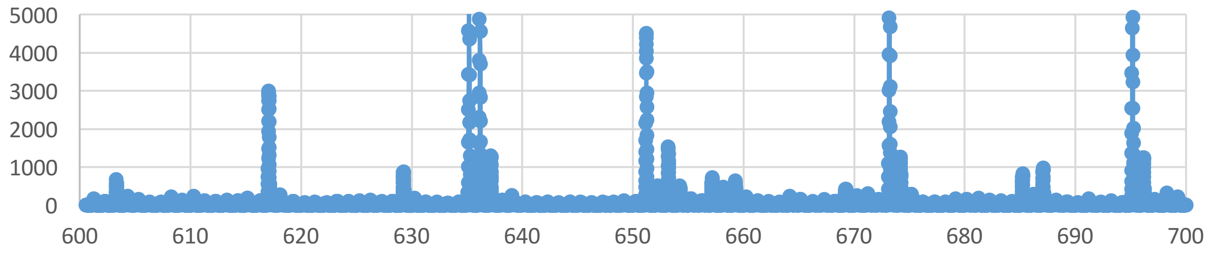

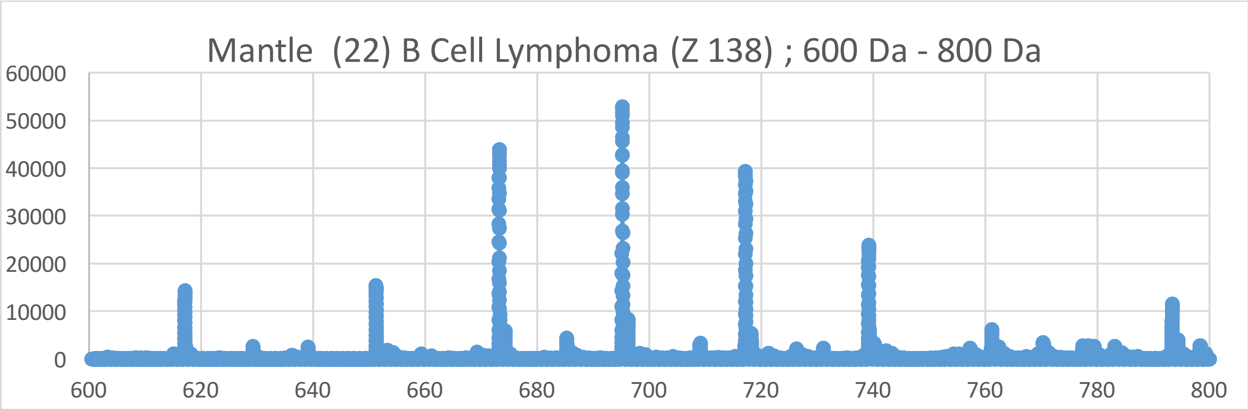
Mantle (22) B Cell Lymphoma (Z 138) ; 600 Da - 800 Da
60000
50000
40000
30000
20000
10000
0
600
620
640
660
680
700
720
740
760
780
800

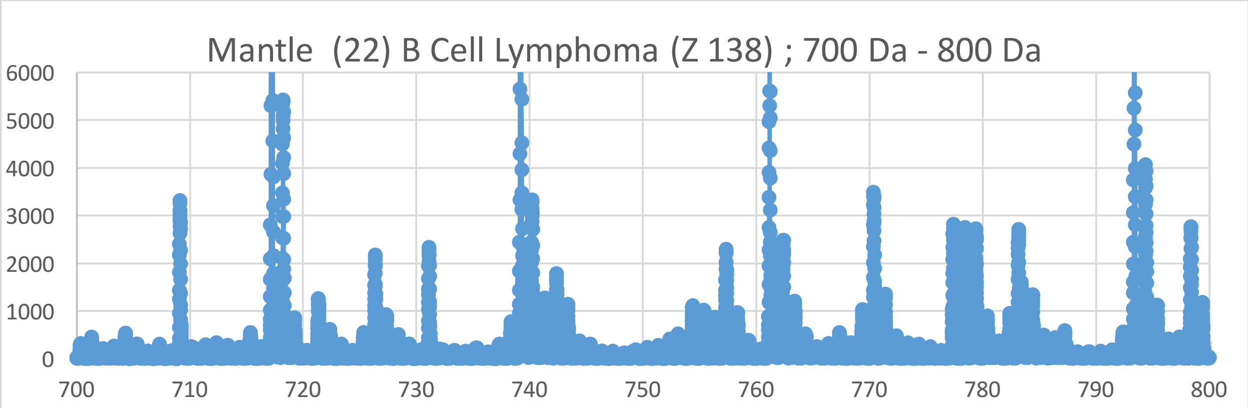
Mantle (22) B Cell Lymphoma (Z 138) ; 700 Da - 800 Da
6000
5000
4000
3000
2000
1000
0
700
710
720
730
740
750
760
770
780
790
800

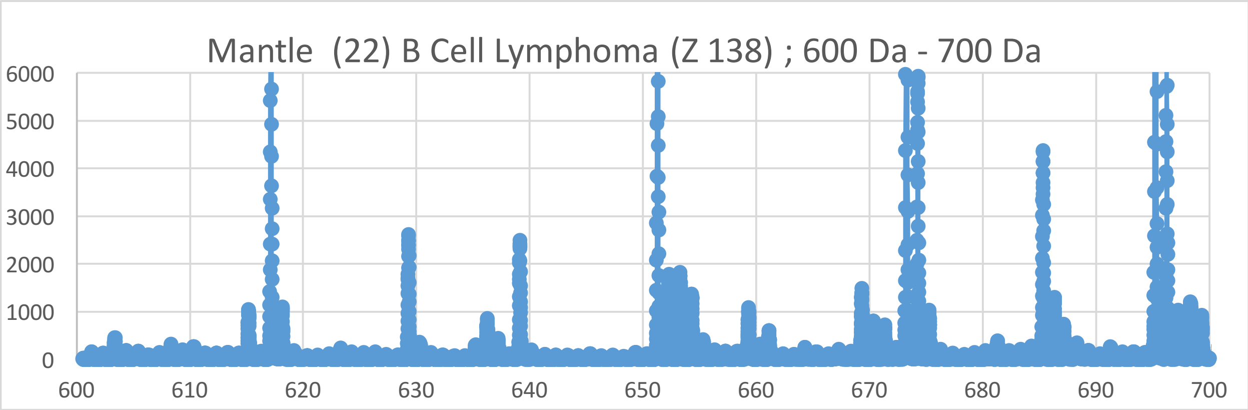
Mantle (22) B Cell Lymphoma (Z 138) ; 600 Da - 700 Da
6000
5000
4000
3000
2000
1000
0
600
610
620
630
640
650
660
670
680
690
700

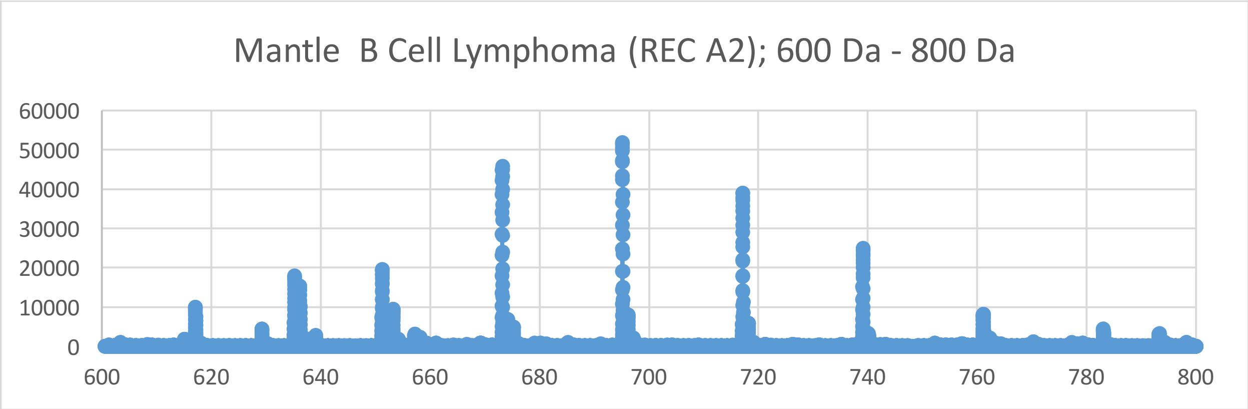
Mantle B Cell Lymphoma (REC A2); 600 Da - 800 Da
60000
50000
40000
30000
20000
10000
0
600
620
640
660
680
700
720
740
760
780
800

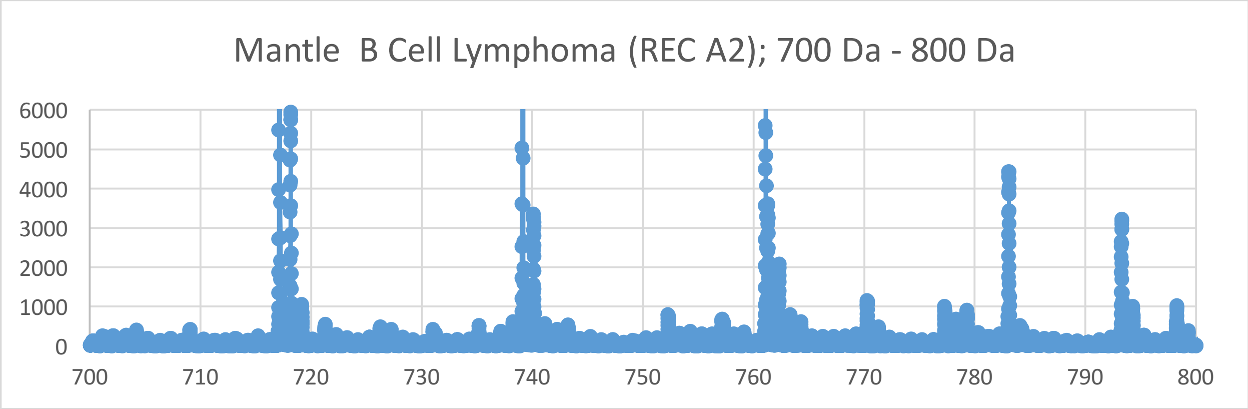
Mantle B Cell Lymphoma (REC A2); 700 Da - 800 Da
6000
5000
4000
3000
2000
1000
0
700
710
720
730
740
750
760
770
780
790
800

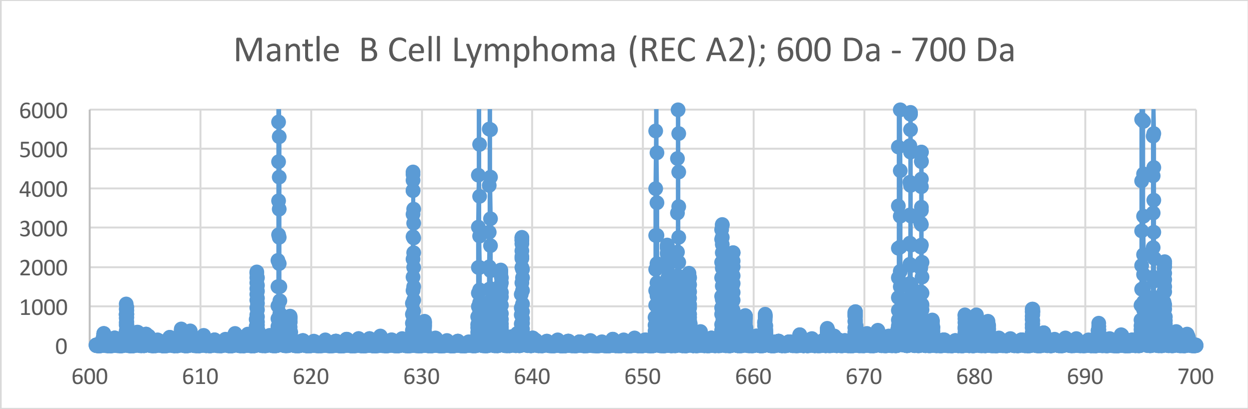
Mantle B Cell Lymphoma (REC A2); 600 Da - 700 Da
6000
5000
4000
3000
2000
1000
0
600
610
620
630
640
650
660
670
680
690
700

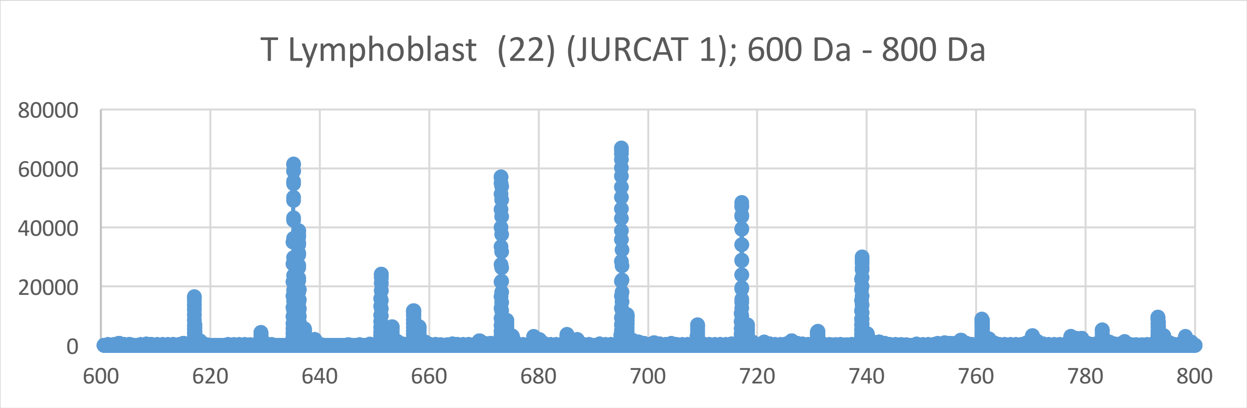
T Lymphoblast (22) (JURCAT 1); 600 Da - 800 Da
80000
60000
40000
20000
0
600
620
640
660
680
700
720
740
760
780
800

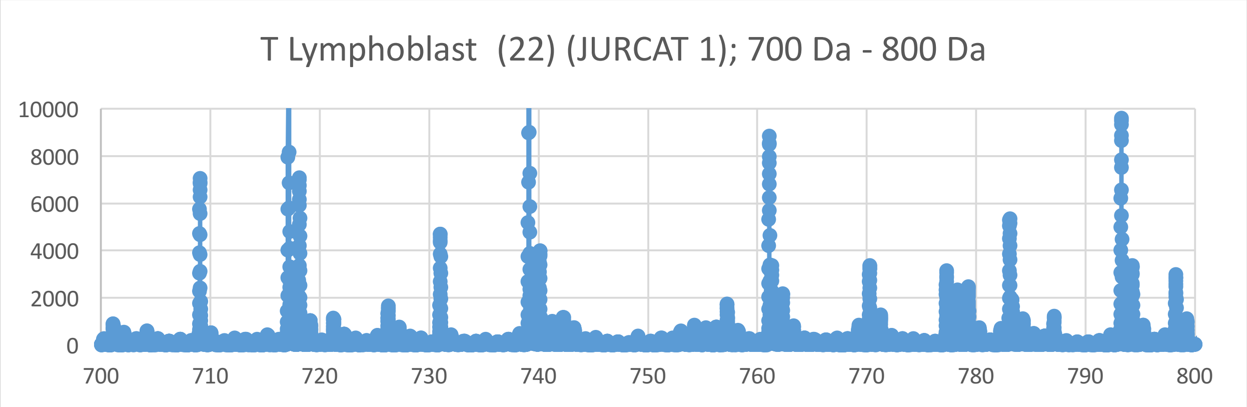
T Lymphoblast (22) (JURCAT 1); 700 Da - 800 Da
10000
8000
6000
4000
2000
0
700
710
720
730
740
750
760
770
780
790
800

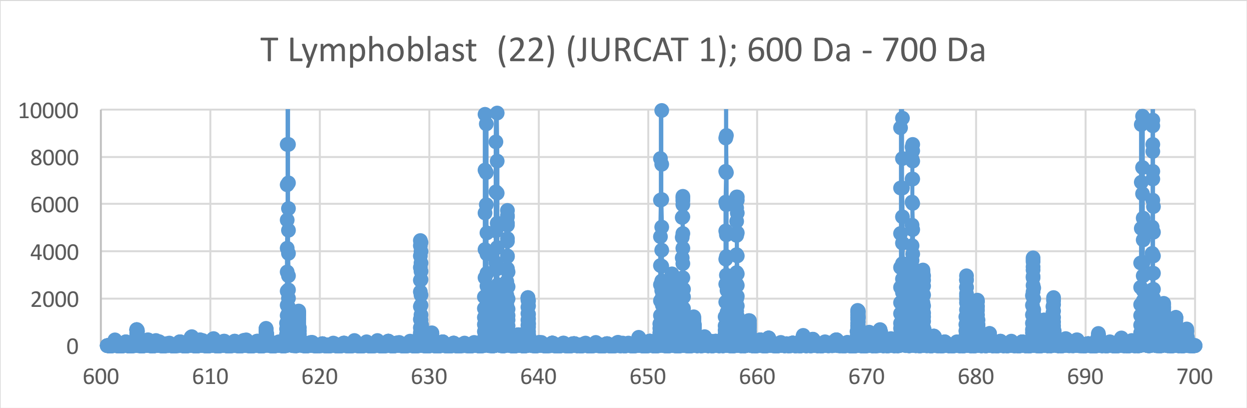
T Lymphoblast (22) (JURCAT 1); 600 Da - 700 Da
10000
8000
6000
4000
2000
0
600
610
620
630
640
650
660
670
680
690
700

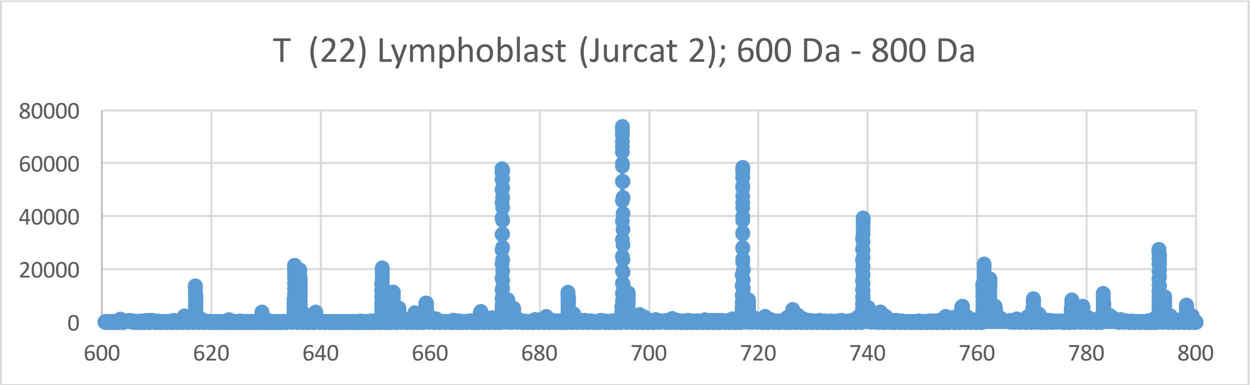
T (22) Lymphoblast (Jurcat 2); 600 Da - 800 Da
80000
60000
40000
20000
0
600
620
640
660
680
700
720
740
760
780
800

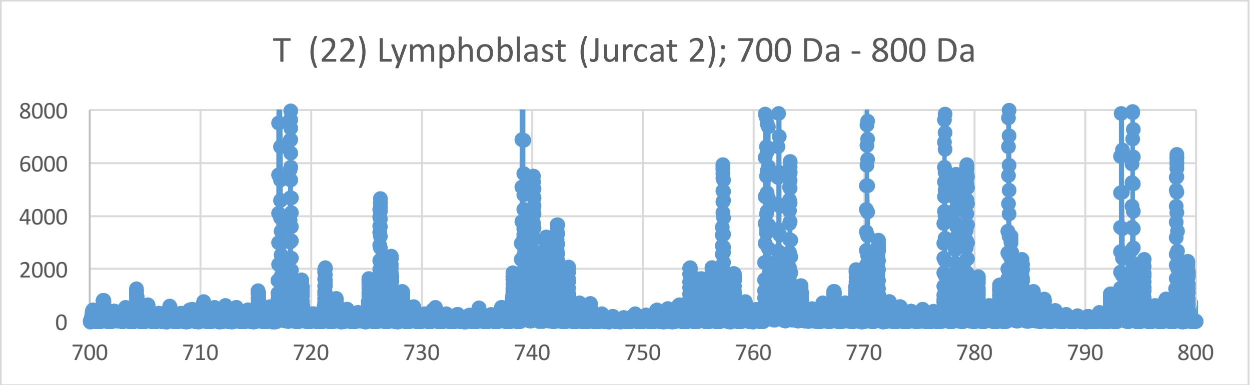
T (22) Lymphoblast (Jurcat 2); 700 Da - 800 Da
8000
6000
4000
2000
0
700
710
720
730
740
750
760
770
780
790
800

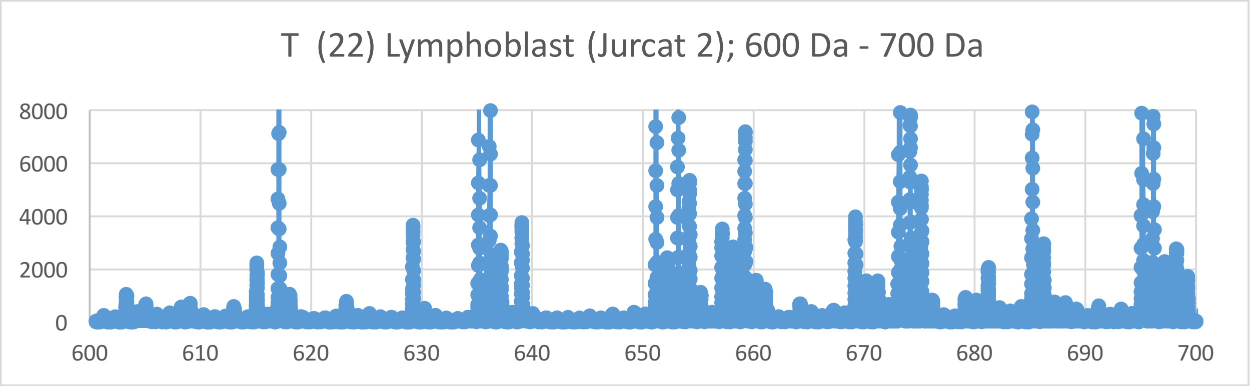
T (22) Lymphoblast (Jurcat 2); 600 Da - 700 Da
8000
6000
4000
2000
0
600
610
620
630
640
650
660
670
680
690
700

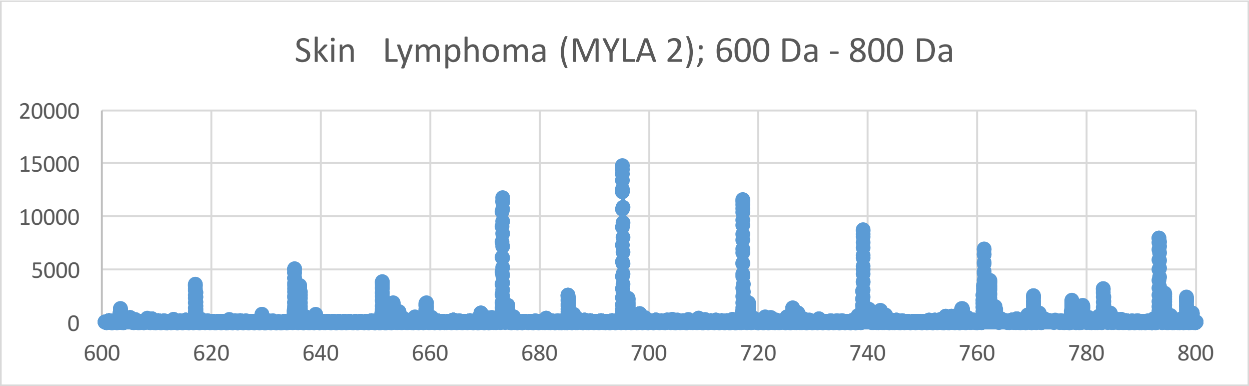
Skin Lymphoma (MYLA 2); 600 Da - 800 Da
20000
15000
10000
5000
0
600
620
640
660
680
700
720
740
760
780
800

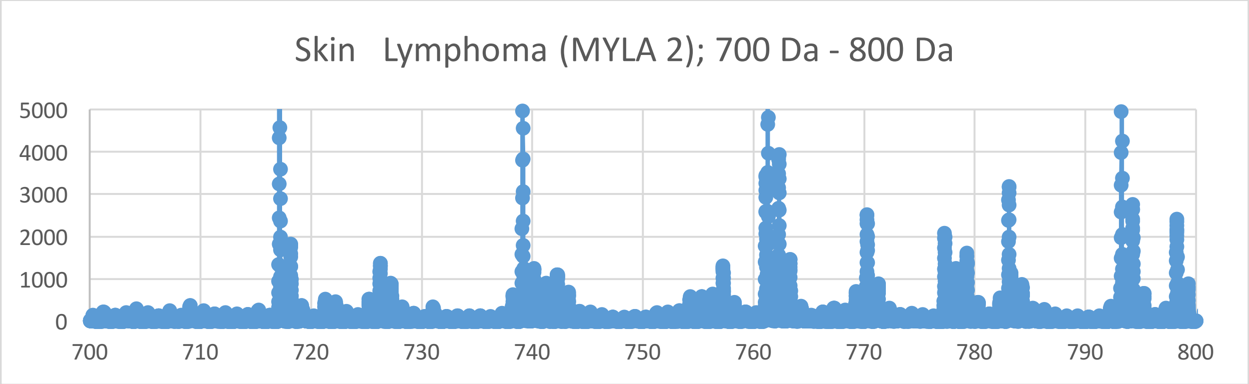
Skin Lymphoma (MYLA 2); 700 Da - 800 Da
5000
4000
3000
2000
1000
0
700
710
720
730
740
750
760
770
780
790
800

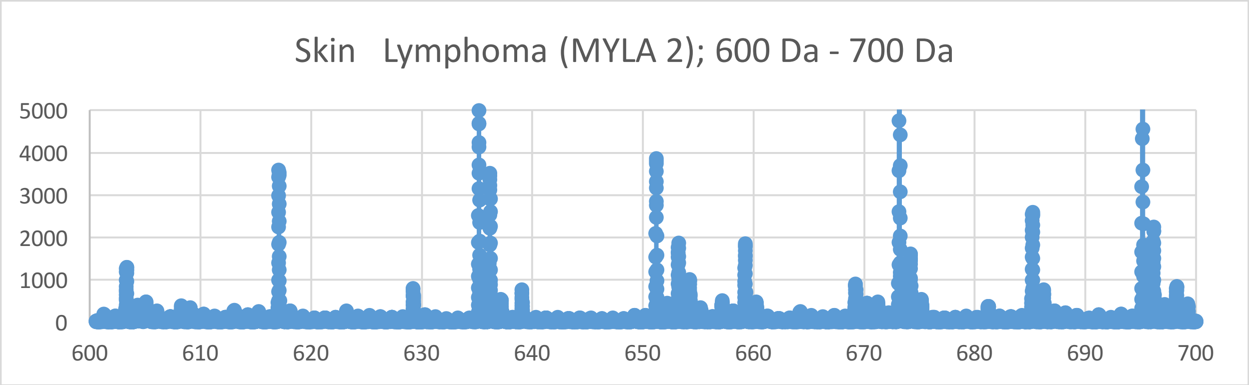
Skin Lymphoma (MYLA 2); 600 Da - 700 Da
5000
4000
3000
2000
1000
0
600
610
620
630
640
650
660
670
680
690
700

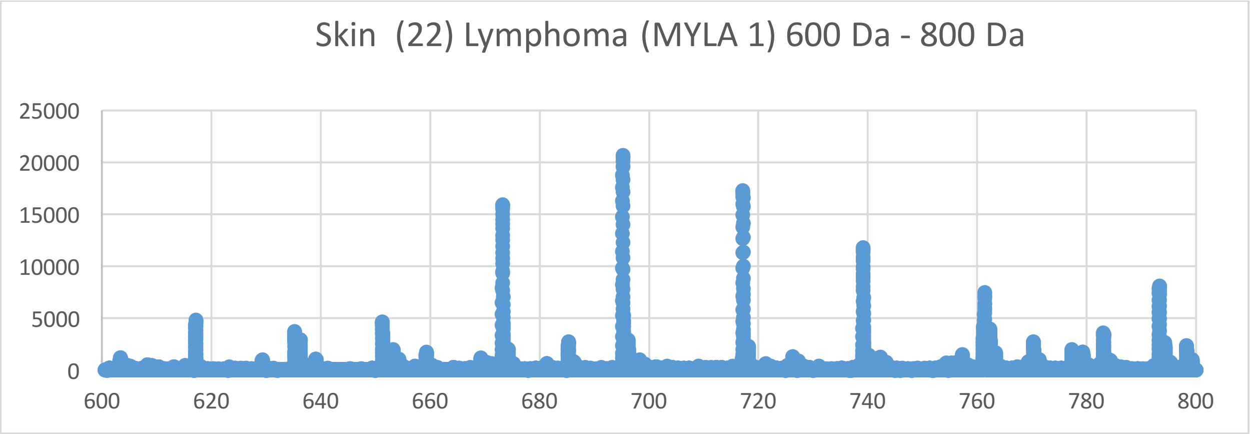
Skin (22) Lymphoma (MYLA 1) 600 Da - 800 Da
25000
20000
15000
10000
5000
0
600
620
640
660
680
700
720
740
760
780
800

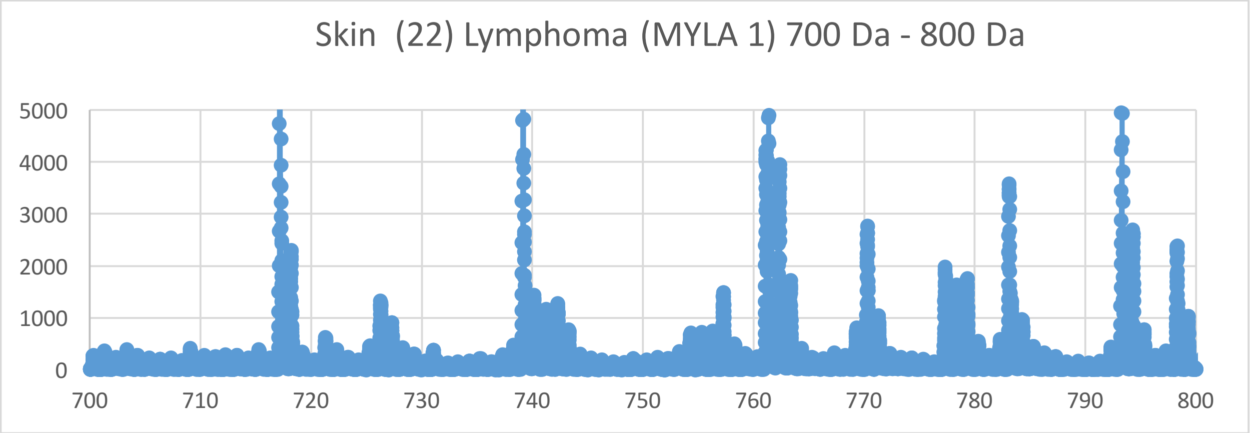
Skin (22) Lymphoma (MYLA 1) 700 Da - 800 Da
5000
4000
3000
2000
1000
0
700
710
720
730
740
750
760
770
780
790
800

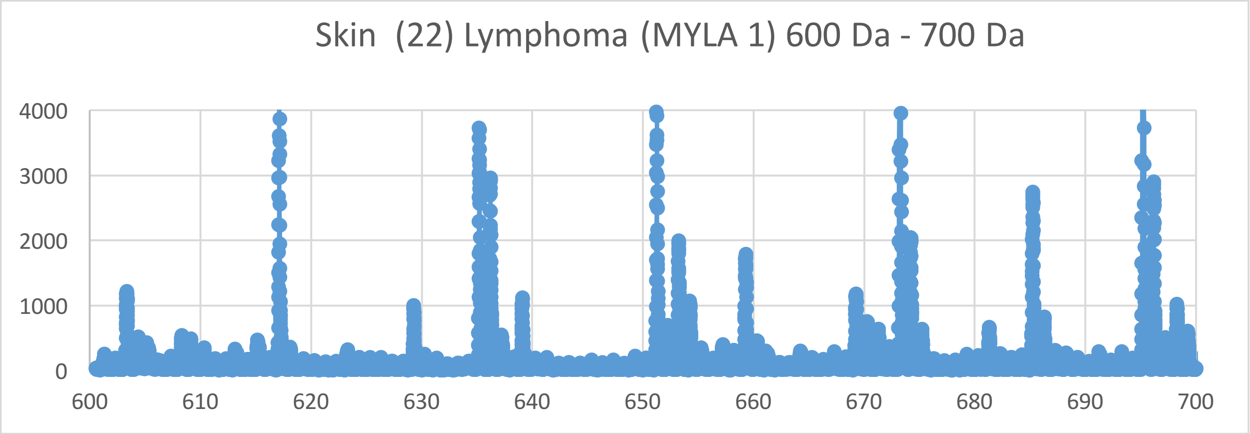
Skin (22) Lymphoma (MYLA 1) 600 Da - 700 Da
4000
3000
2000
1000
0
600
610
620
630
640
650
660
670
680
690
700

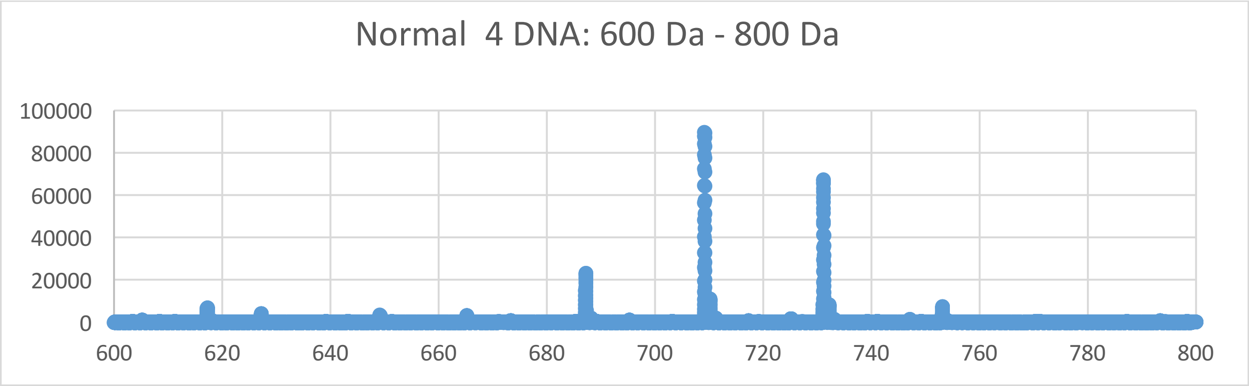
Normal 4 DNA: 600 Da - 800 Da
100000
80000
60000
40000
20000
0
600
620
640
660
680
700
720
740
760
780
800

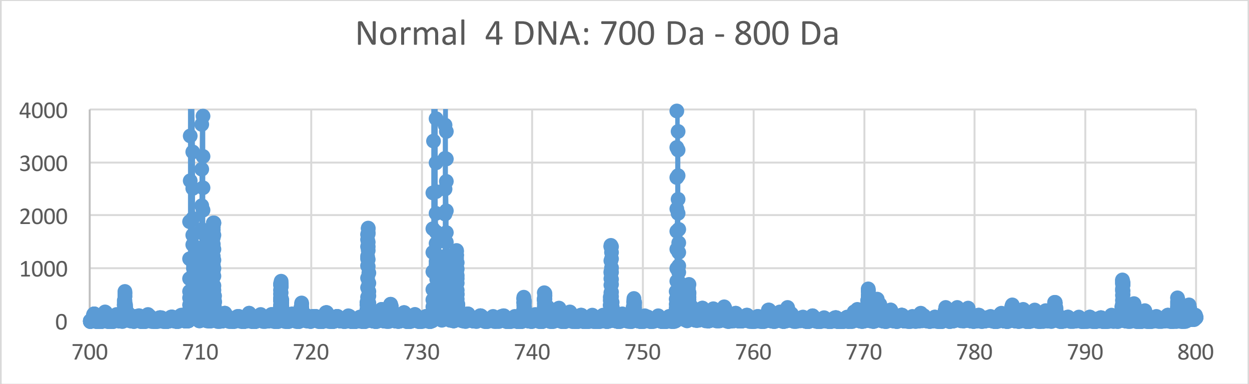
Normal 4 DNA: 700 Da - 800 Da
4000
3000
2000
1000
0
700
710
720
730
740
750
760
770
780
790
800

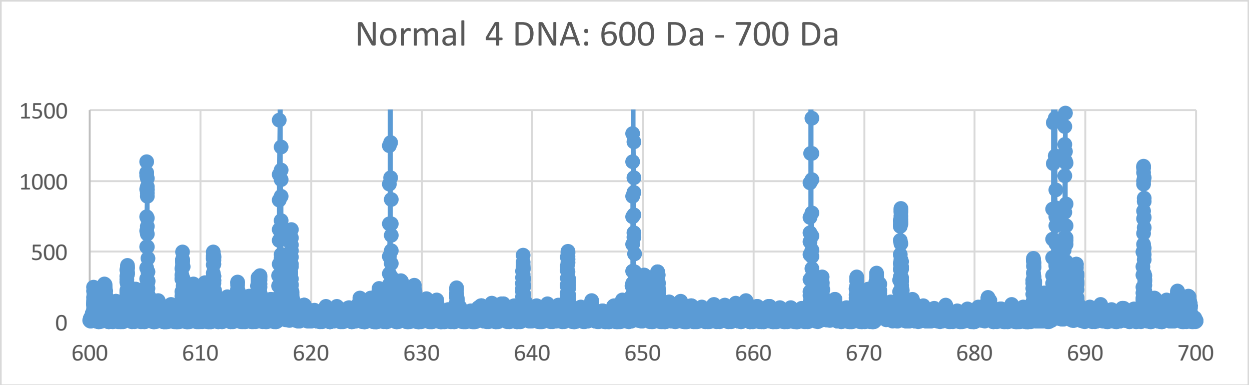
Normal 4 DNA: 600 Da - 700 Da
1500
1000
500
0
600
610
620
630
640
650
660
670
680
690
700

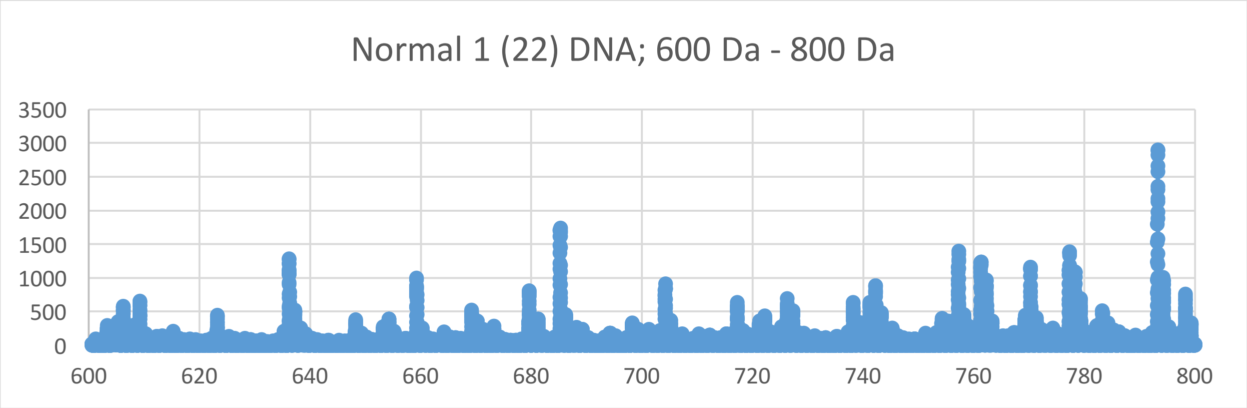

Normal 1 (22) DNA; 600 Da - 800 Da
3500
3000
2500
2000
1500
1000
500
0
600
620
640
660
680
700
720
740
760
780
800


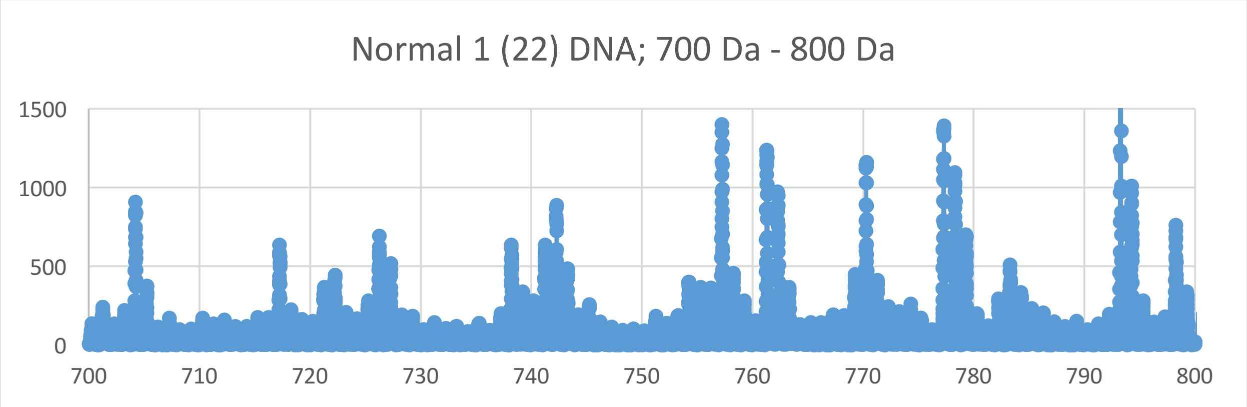

Normal 1 (22) DNA; 700 Da - 800 Da
1500
1000
500
0
700
710
720
730
740
750
760
770
780
790
800


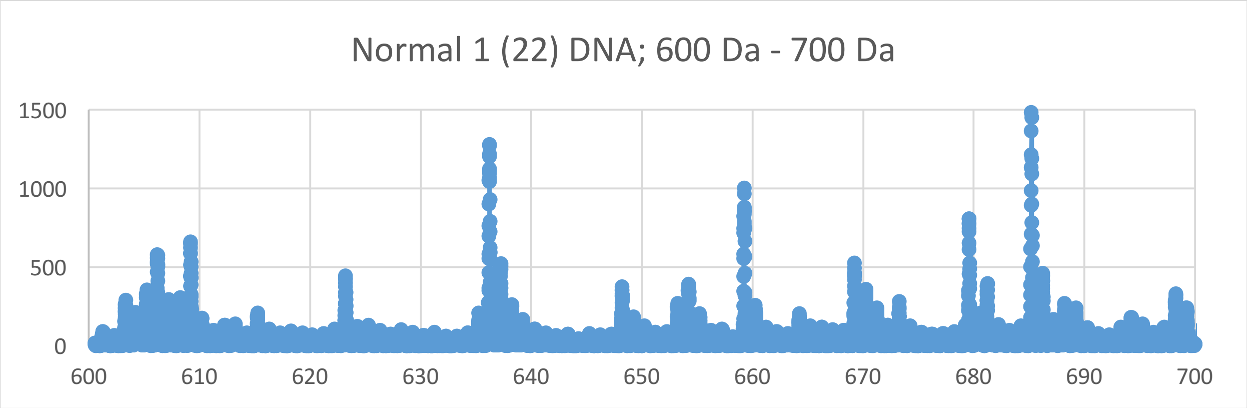

Normal 1 (22) DNA; 600 Da - 700 Da
1500
1000
500
0
600
610
620
630
640
650
660
670
680
690
700

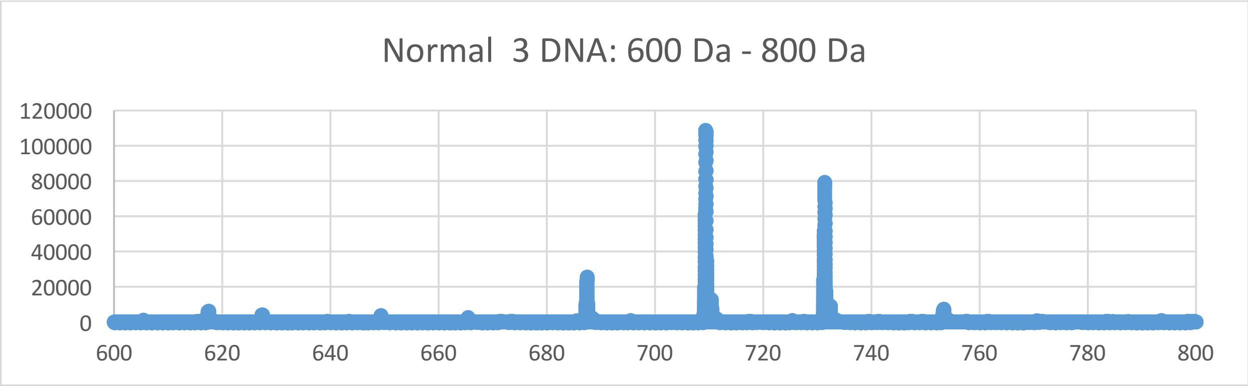
Normal 3 DNA: 600 Da - 800 Da
120000
100000
80000
60000
40000
20000
0
600
620
640
660
680
700
720
740
760
780
800

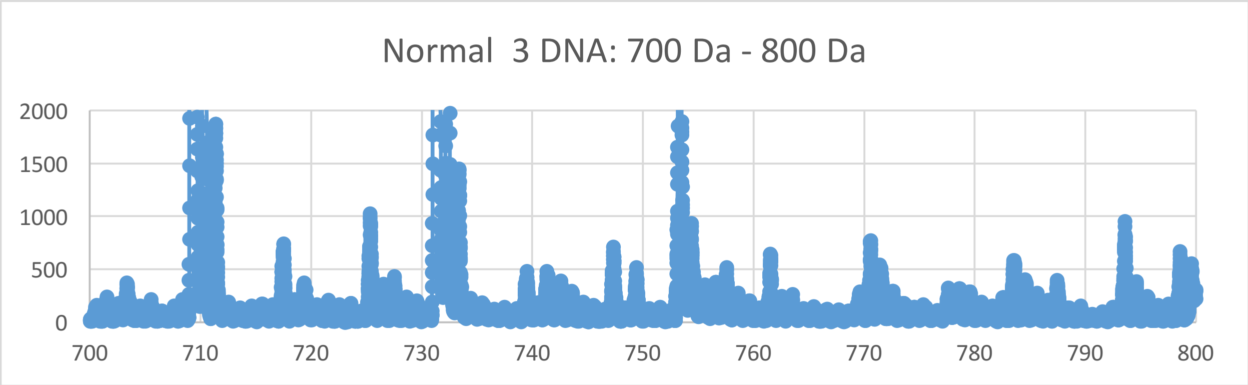
Normal 3 DNA: 700 Da - 800 Da
2000
1500
1000
500
0
700
710
720
730
740
750
760
770
780
790
800

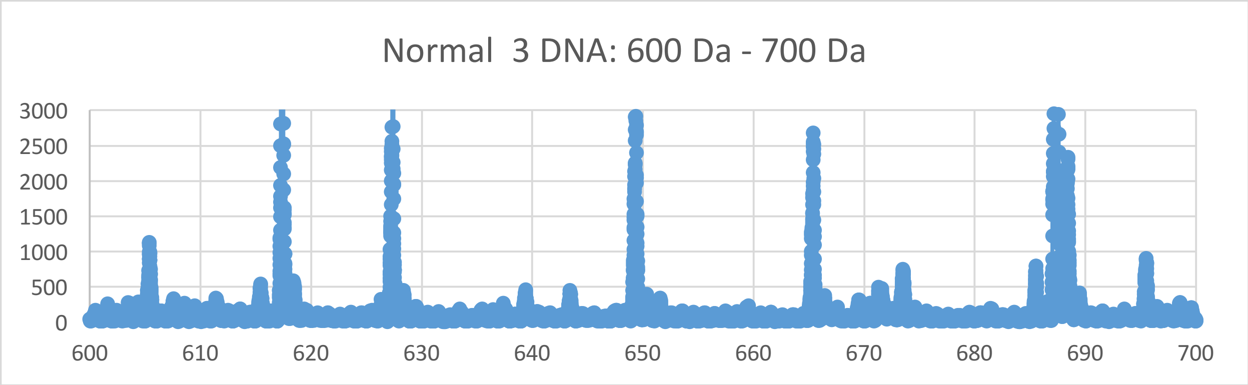
Normal 3 DNA: 600 Da - 700 Da
3000
2500
2000
1500
1000
500
0
600
610
620
630
640
650
660
670
680
690
700

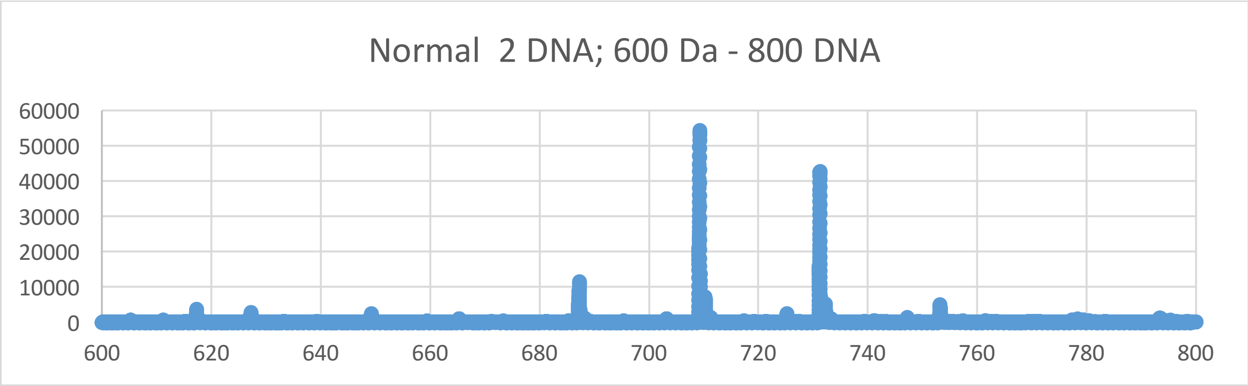
Normal 2 DNA; 600 Da - 800 DNA
60000
50000
40000
30000
20000
10000
0
600
620
640
660
680
700
720
740
760
780
800

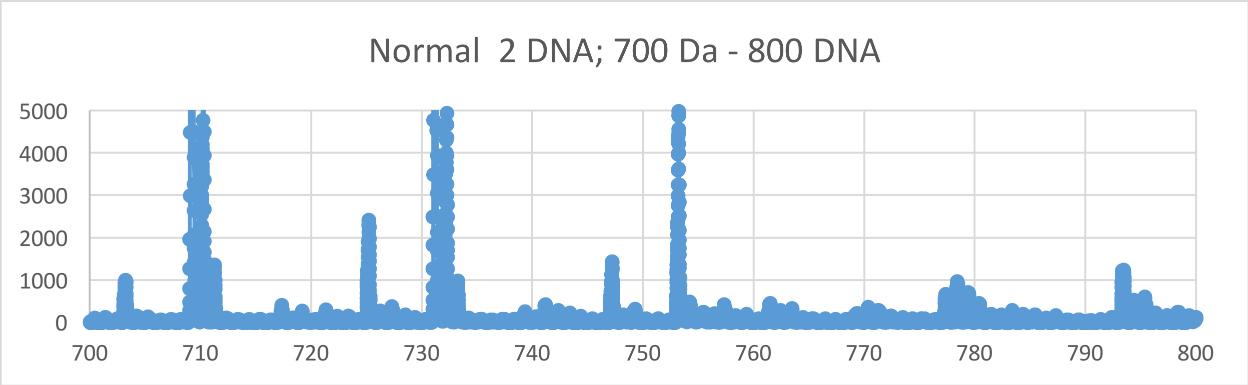
Normal 2 DNA; 700 Da - 800 DNA
5000
4000
3000
2000
1000
0
700
710
720
730
740
750
760
770
780
790
800

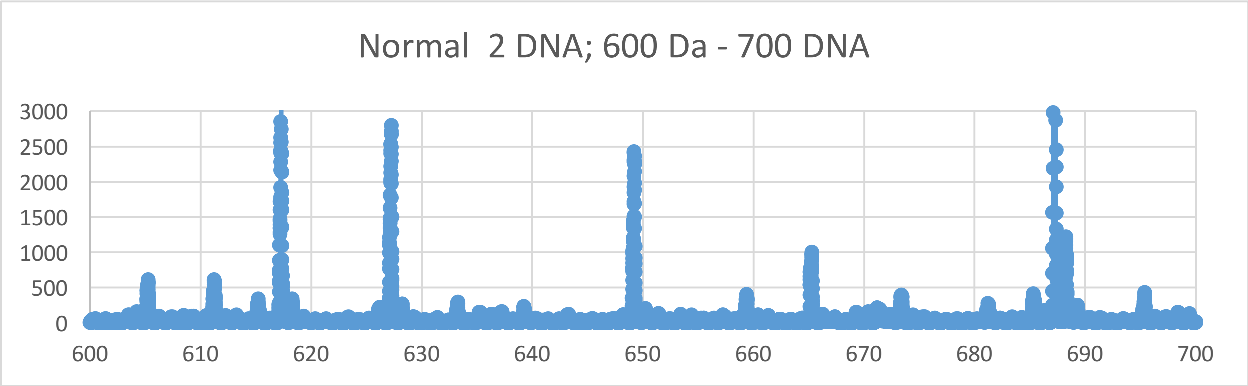
Normal 2 DNA; 600 Da - 700 DNA
3000
2500
2000
1500
1000
500
0
600
610
620
630
640
650
660
670
680
690
700
