## Supplementary Materials XI for "Oncogenesis and Aging by Isotopic Functionalizations of the Proteins and Nucleic Acids"

XI Supplementary Materials – Mass Spectra from 400 Da -600 Da

Results for Supplementary 11

So for the mononucleotides at $(phos)_3$, the A, T, G and C peaks for normal dramatically change in peak intensities and increase in numbers with the $(phos)_3$ to $(phos)_2$ dephosphorylations. The mononucleotides dramatically change fragmentation patterns for cancer at $(phos)_2$ from strong stable normal pieces at $(phos)_3$. But as decrease the phosphate by 1 unit, the mononucleotides change so that the normal mononucleotides more fragment and break with stabilities from the DNA with more numbers and stabilities for the A, T, C and G for $(phos)_3$ to $(phos)_2$. The masses range from 544 Da to 383 Da. The normals fragment more intense mononucleotides in this mass range. See Supplementary Figures XI for graphs. It seems that below the saturation range of some number of phosphates, the telomeres, mating telomeres and nontelomeres behave in similar way within cancer and normal DNA. So this suggests cancer may be favored and distinguished from normal by changing saturated phosphorylations from $(phos)_3$ to $(phos)_2$. The A, T, C and G mononucleotide peaks for normal strongly decrease with this dephosphorylations of the mononucleotides. The dephosphorylations from $(phos)_3$ to $(phos)_2$ caused the resulting dramatic, intense and stable normal mononucleotides to be more telomeric, mating telomeric and nontelomeric and cancerous and strongly isotopically enriched for telomeres, mating telomeres and nontelomeres. It is important to consider that there are more intense peaks for the mononucleotides relative to the dinucleotides, trinucleotides and tetranucleotides for the saturated phosphate units. For the mononucleotides, the intensities of normal strongly decreased and the intensities of cancer fragments slightly increased from $(phos)_3$ to $(phos)_{2.}$ The dephosphorylations of telomeric, mating telomeric and nontelomeric mononucleotides from $(phos)_3$ to $(phos)_2$ negligibly decreases isotopic enrichments of heavy isotopes of $^{15}N$, $^{13}C$ and $^{17}O$. The dephosphorylations from $(phos)_3$ to $(phos)_2$ also caused the telomeric, mating telomeric and nontelomeric mononucleotides to be more intense in cancer DNA and negligibly more isotopically enriched. It seems the dephosphorylations are related to the isotopic enrichments and depletions; but the telomeres are more intense in cancer and negligibly more isotopically enriched. The telomeric and mating telomeric mononucleotides of cancer tended to also be negligibly isotopically enriched; so it seems the cancer stabilities of the telomeric, mating telomeric and nontelomeric mononucleotides are related to their isotopic enrichments from the dephosphorylations from $(phos)_3$ to $(phos)_2$ and the mutations purely driven by dephosphorylations and enrichments. Enrichments weaken the mating normal mononucleotides while strengthening cancerous telomeric mononucleotides. The intense cancer telomeric, mating telomeric and nontelomeric mononucleotides were also more isotopically enriched. The dephosphorylations from $(phos)_3$ to $(phos)_2$ negligibly increased the isotopic enrichments of the cancer for the telomeric, nontelomeric and mating telomeric mononucleotides. Such seems compatible for replications and transcriptions as stable telomeres seem compatible to unstable mating telomeres for the unravelings and constructions of DNA mating telomeres onto the building templates of the telomeres. But in this case of all normal telomeric, mating telomeric and nontelomeric mononucleotides are negligibly enriched and unstable then the replications, transcriptions and translations cannot occur normally. The dephosphorylations from $(phos)_3$ to $(phos)_2$ caused demethylations to cause more cancer intensities and stabilities but lowered isotopic enrichments for G. But the demethylations of the G and A caused negligible alterations cancer stabilities and isotopic enrichments. Where isotopic changes occurred, the functionalizational effects were nonnegligible. The methylations of the C mononucleotides increased cancer intensities and stabilities with isotopic enrichments. It seems from these data that during biological processes the dephosphorylations and stabilizations of normal or vice versa dephosphorylations and stabilizations of cancer are or can be countered by methylations (and isotopic enrichments) and destabilizations of the normal for cancer or vice versa dephosphorylations and destabilizations of the cancer by methylations as this research discovers coupled phosphorylations and methylations (and isotopic enrichments) or coupled dephosphorylations and methylations (and isotopic enrichments) with involved greater enrichments for normal biological functions and explaining cancer! The dephosphorylations from $(phos)_3$ to $(phos)_2$ caused acetylations to cause less cancer stabilities for A and C with nonnegligible isotopic depletions and less cancer stabilities for G and T with nonnegligible depletions of heavy isotopes. The acetylations from $(phos)_3$ to $(phos)_2$ caused less cancer stabilities and intensities for A and C with more isotopic depletions. This contrast in A, T, C and G from the de-acetylations and dephosphorylations may reflect the developments of cancer as the mononucleotides fragment but the other mononucleotides more hold together for damaged DNA for cancer.

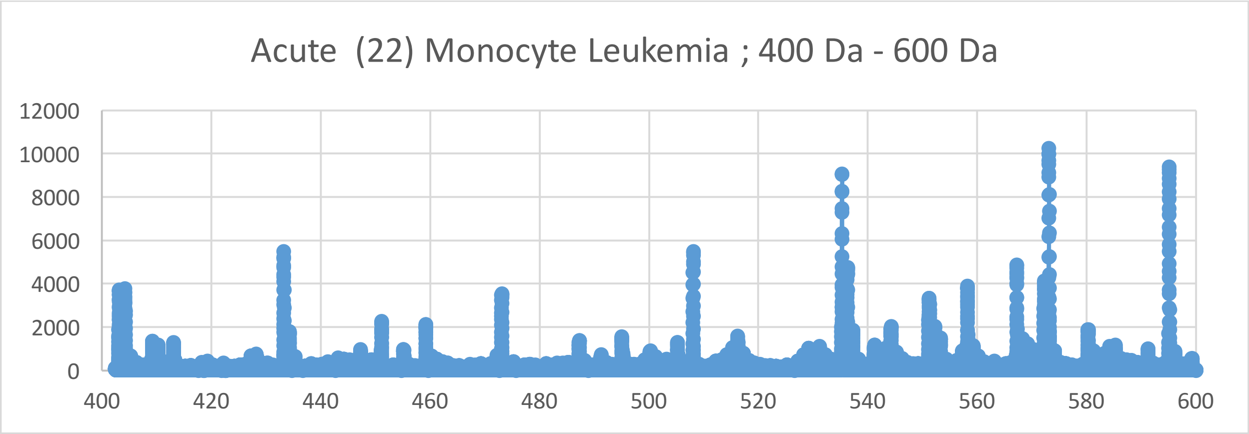
Acute (22) Monocyte Leukemia ; 400 Da - 600 Da
12000
10000
8000
6000
4000
2000
0
400
420
440
460
480
500
520
540
560
580
600

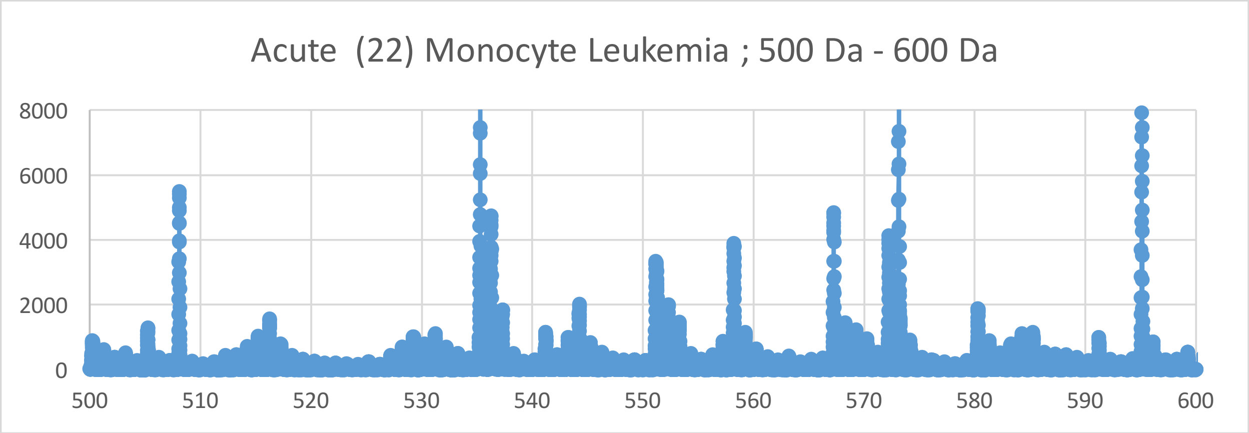
Acute (22) Monocyte Leukemia ; 500 Da - 600 Da
8000
6000
4000
2000
0
500
510
520
530
540
550
560
570
580
590
600

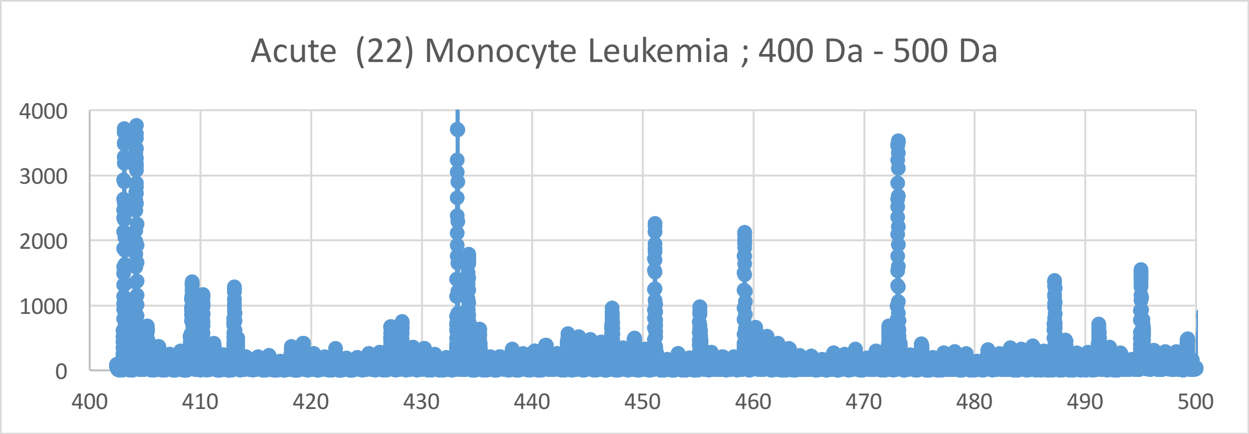
Acute (22) Monocyte Leukemia ; 400 Da - 500 Da
4000
3000
2000
1000
0
400
410
420
430
440
450
460
470
480
490
500

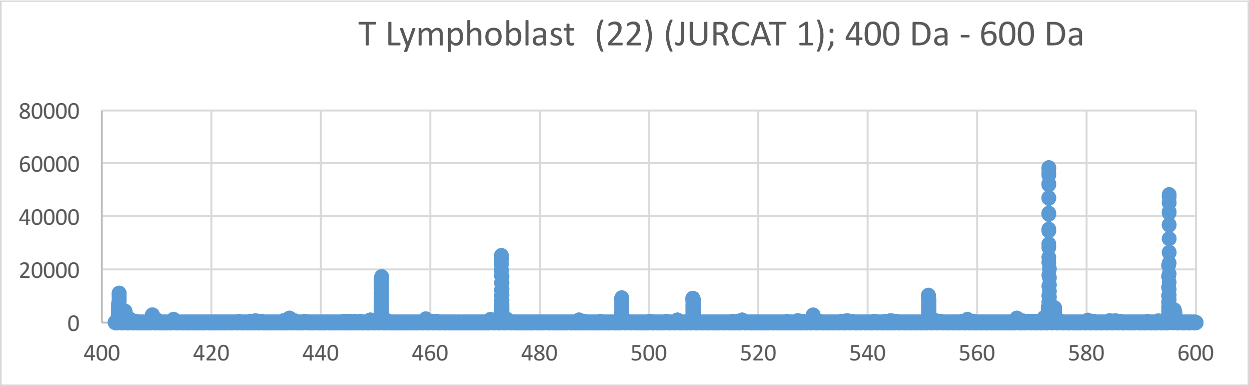
T Lymphoblast (22) (JURCAT 1); 400 Da - 600 Da
0
20000
40000
60000
80000
400 420 440 460 480 500 520 540 560 580 600

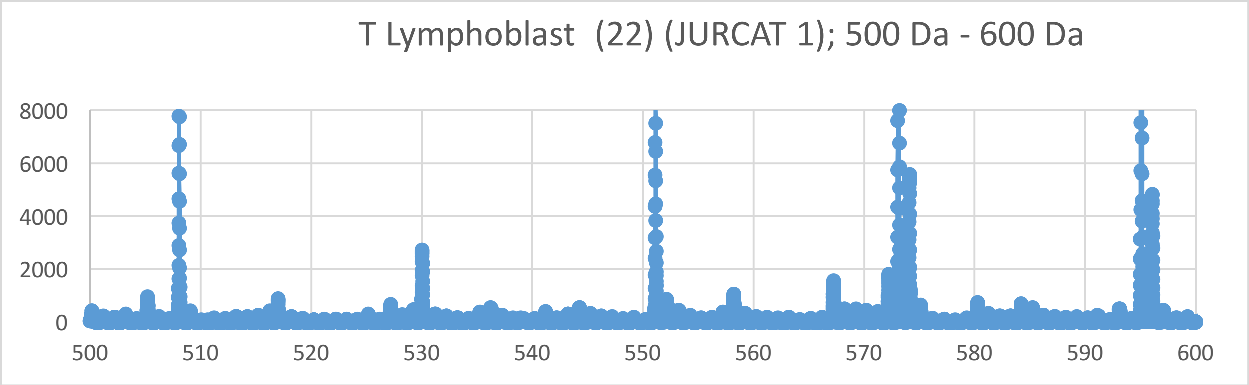
T Lymphoblast (22) (JURCAT 1); 500 Da - 600 Da
0
2000
4000
6000
8000
500 510 520 530 540 550 560 570 580 590 600

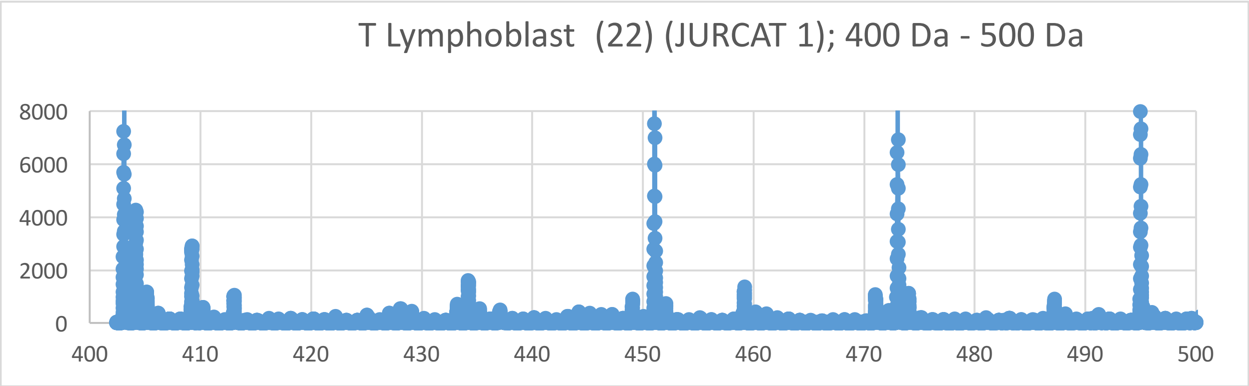
T Lymphoblast (22) (JURCAT 1); 400 Da - 500 Da
0
2000
4000
6000
8000
400 410 420 430 440 450 460 470 480 490 500

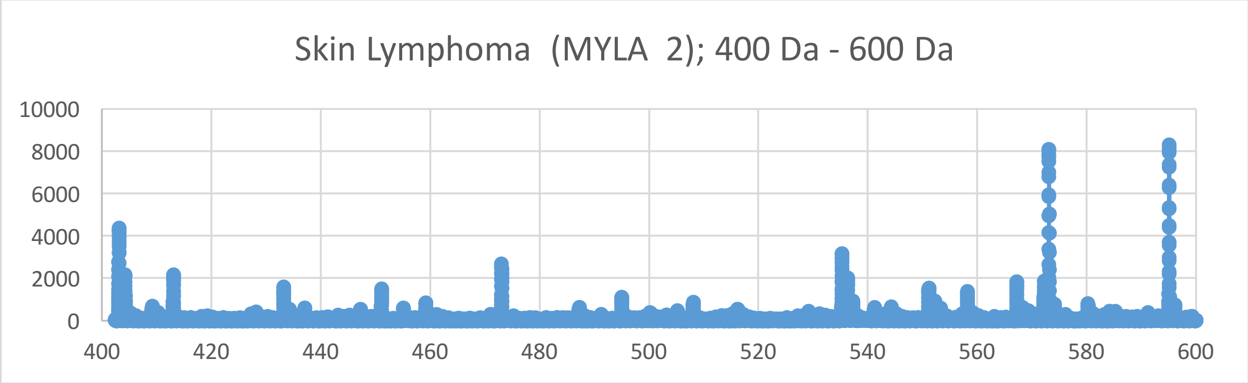
Skin Lymphoma (MYLA 2); 400 Da - 600 Da
10000
8000
6000
4000
2000
0
400
420
440
460
480
500
520
540
560
580
600

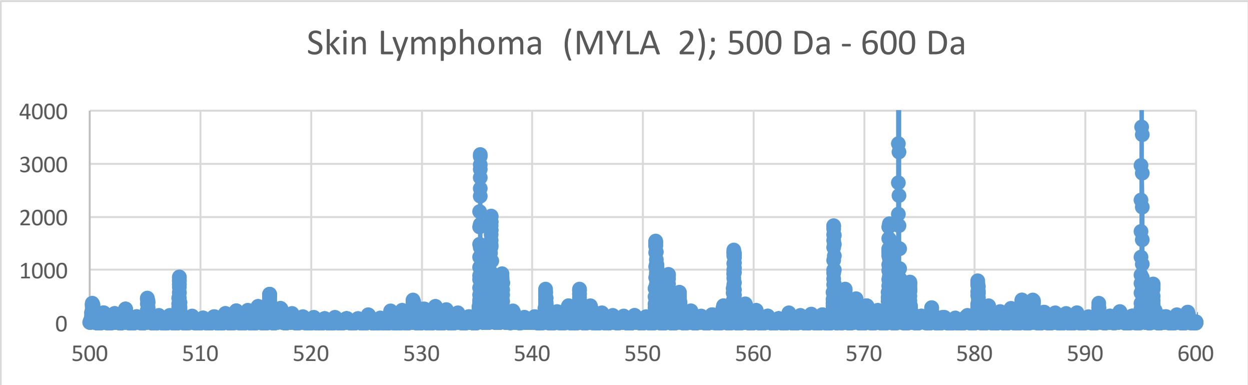
Skin Lymphoma (MYLA 2); 500 Da - 600 Da
4000
3000
2000
1000
0
500
510
520
530
540
550
560
570
580
590
600

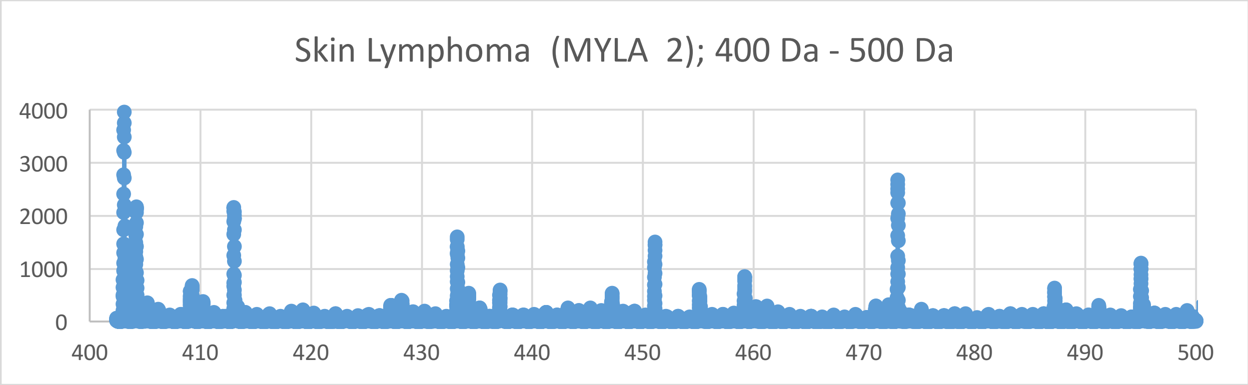
Skin Lymphoma (MYLA 2); 400 Da - 500 Da
4000
3000
2000
1000
0
400
410
420
430
440
450
460
470
480
490
500

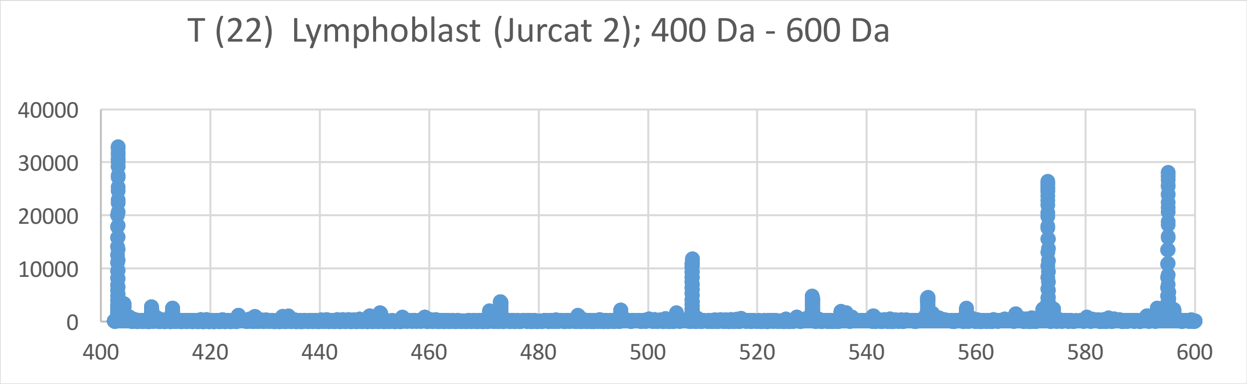

T (22) Lymphoblast (Jurcat 2); 400 Da - 600 Da
40000
30000
20000
10000
0
400
420
440
460
480
500
520
540
560
580
600


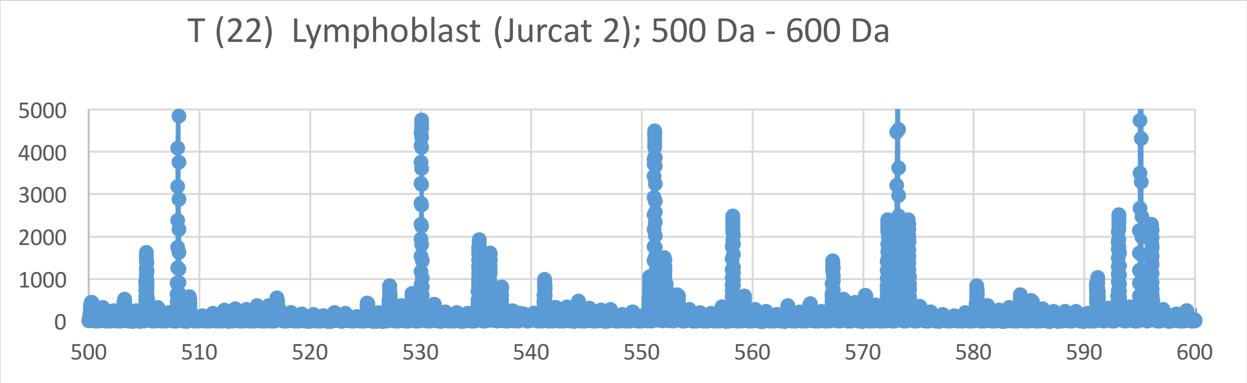

T (22) Lymphoblast (Jurcat 2); 500 Da - 600 Da
5000
4000
3000
2000
1000
0
500
510
520
530
540
550
560
570
580
590
600


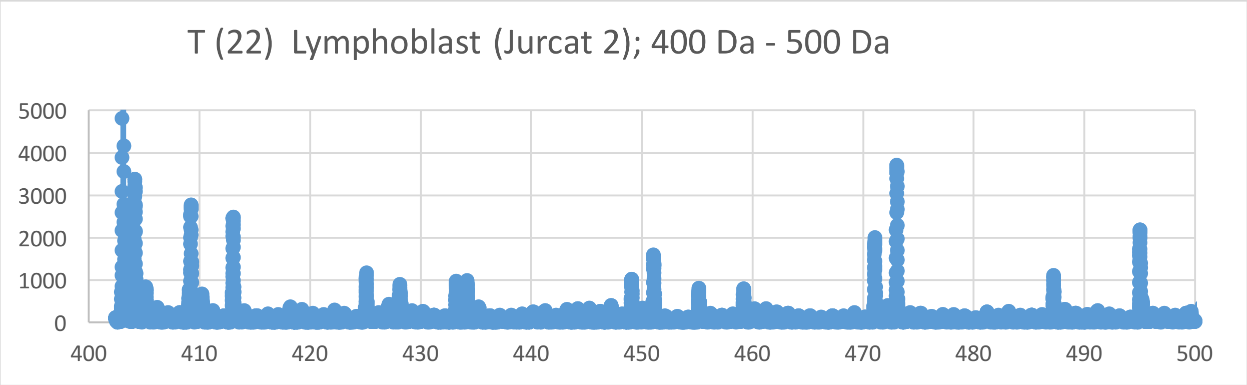

T (22) Lymphoblast (Jurcat 2); 400 Da - 500 Da
5000
4000
3000
2000
1000
0
400
410
420
430
440
450
460
470
480
490
500

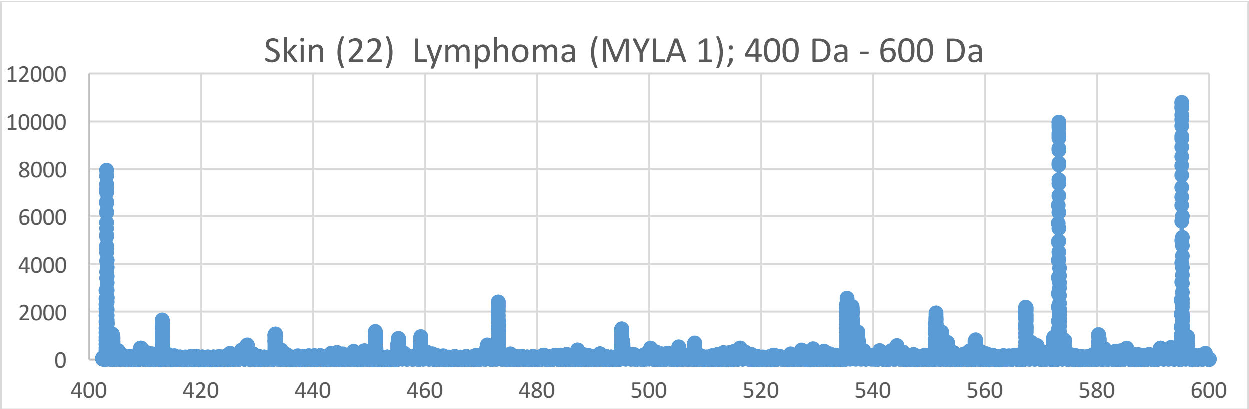
Skin (22) Lymphoma (MYLA 1); 400 Da - 600 Da
12000
10000
8000
6000
4000
2000
0
400
420
440
460
480
500
520
540
560
580
600

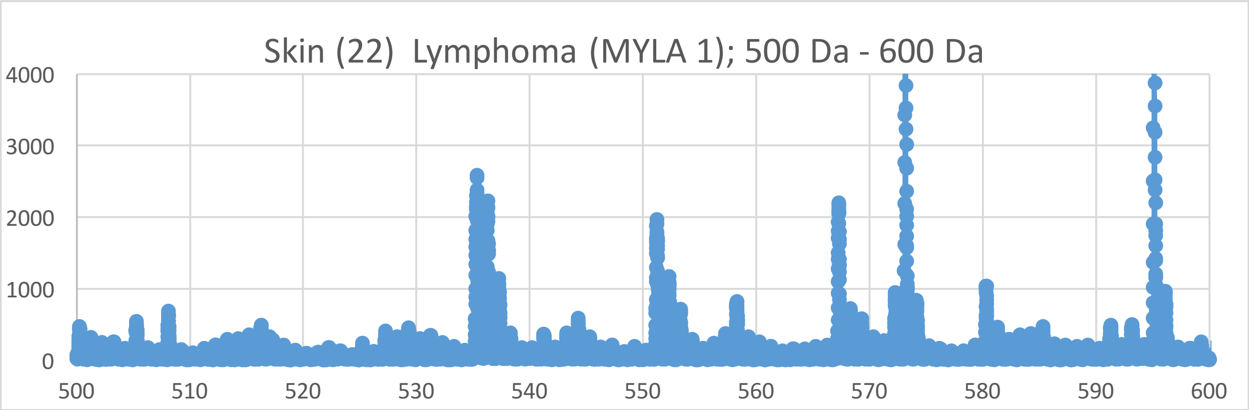
Skin (22) Lymphoma (MYLA 1); 500 Da - 600 Da
4000
3000
2000
1000
0
500
510
520
530
540
550
560
570
580
590
600

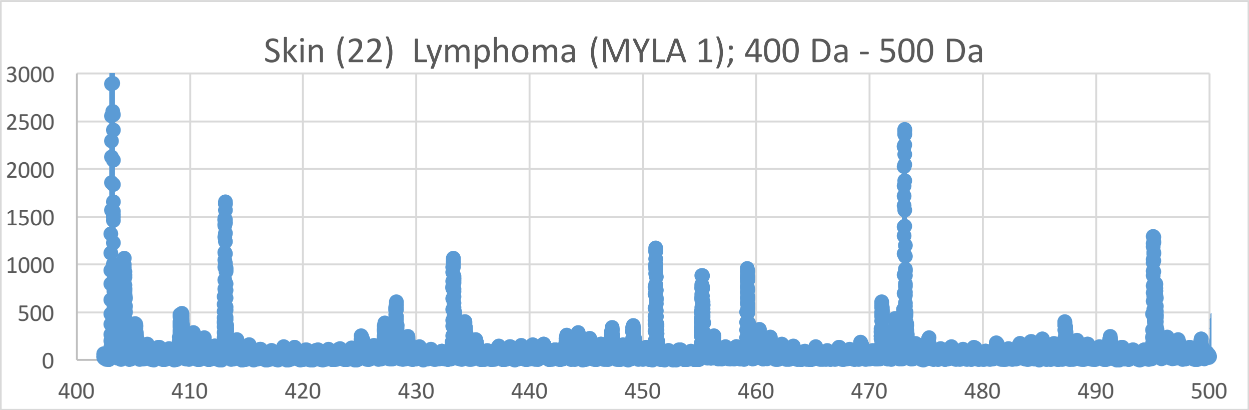
Skin (22) Lymphoma (MYLA 1); 400 Da - 500 Da
3000
2500
2000
1500
1000
500
0
400
410
420
430
440
450
460
470
480
490
500

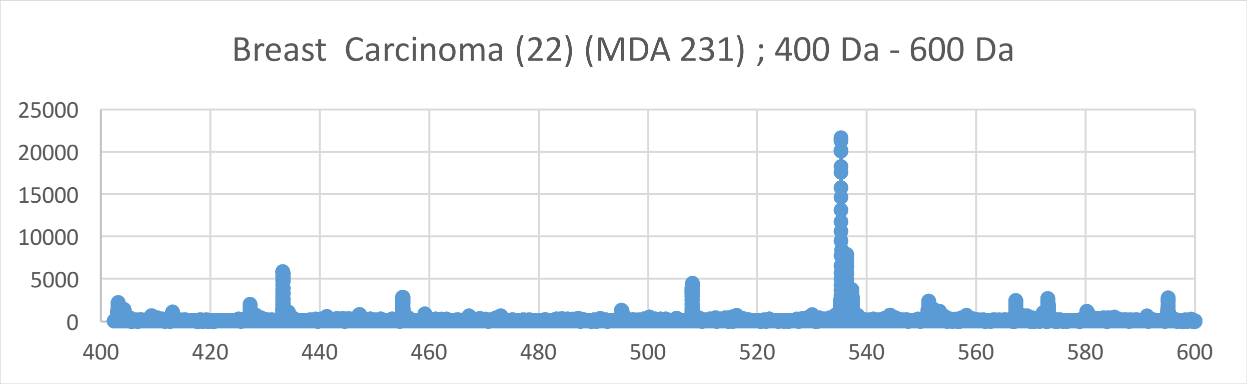

Breast  Carcinoma (22) (MDA 231) ; 400 Da - 600 Da
25000
20000
15000
10000
5000
0
400
420
440
460
480
500
520
540
560
580
600


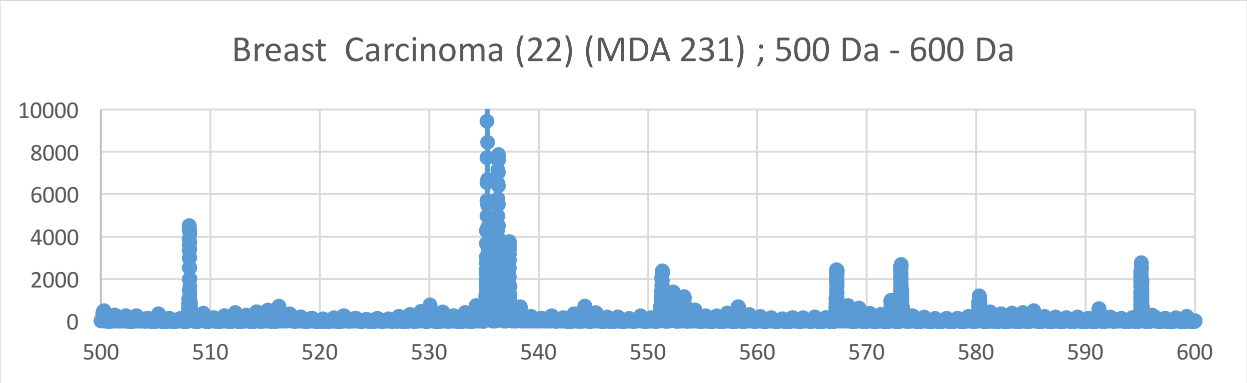

Breast  Carcinoma (22) (MDA 231) ; 500 Da - 600 Da
10000
8000
6000
4000
2000
0
500
510
520
530
540
550
560
570
580
590
600


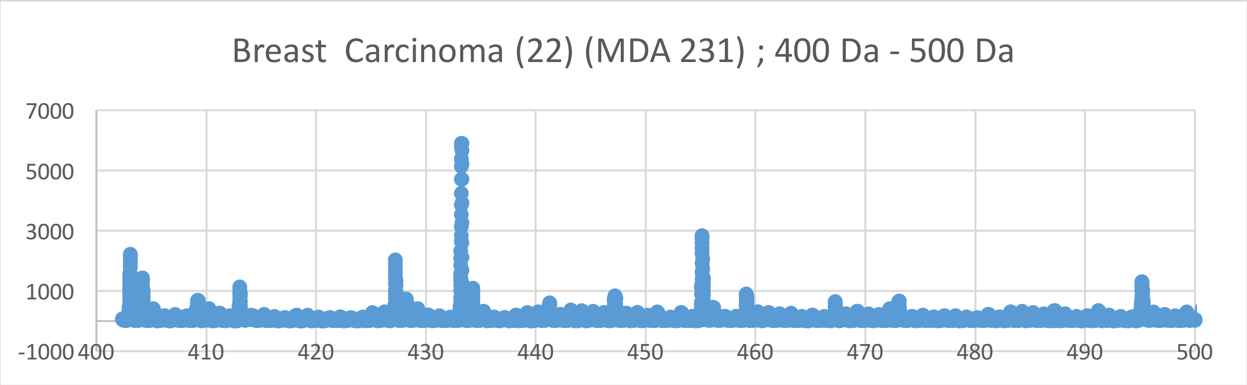

Breast  Carcinoma (22) (MDA 231) ; 400 Da - 500 Da
7000
5000
3000
1000
-1000
400
410
420
430
440
450
460
470
480
490
500

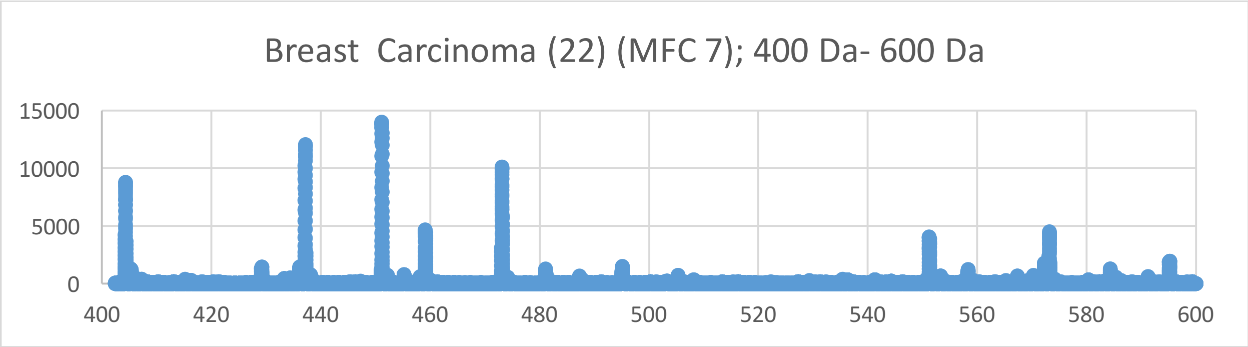
Breast  Carcinoma (22) (MFC 7); 400 Da- 600 Da
15000
10000
5000
0
400
420
440
460
480
500
520
540
560
580
600

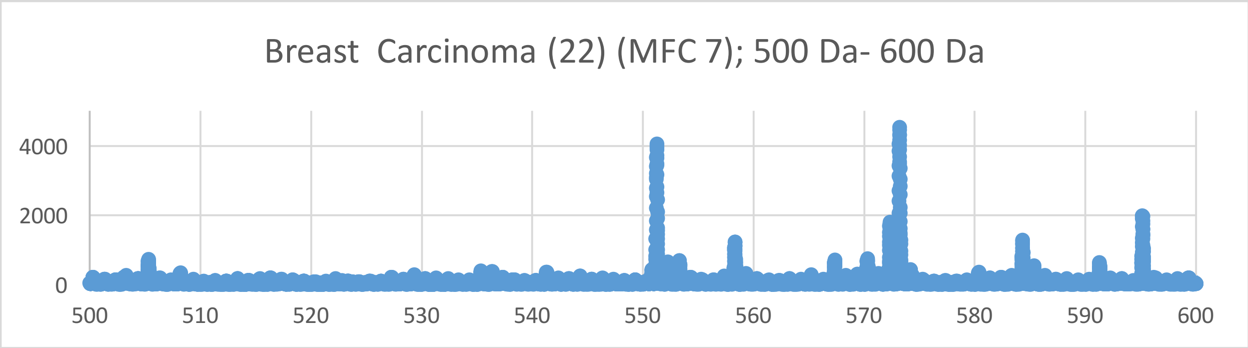
Breast  Carcinoma (22) (MFC 7); 500 Da- 600 Da
4000
2000
0
500
510
520
530
540
550
560
570
580
590
600

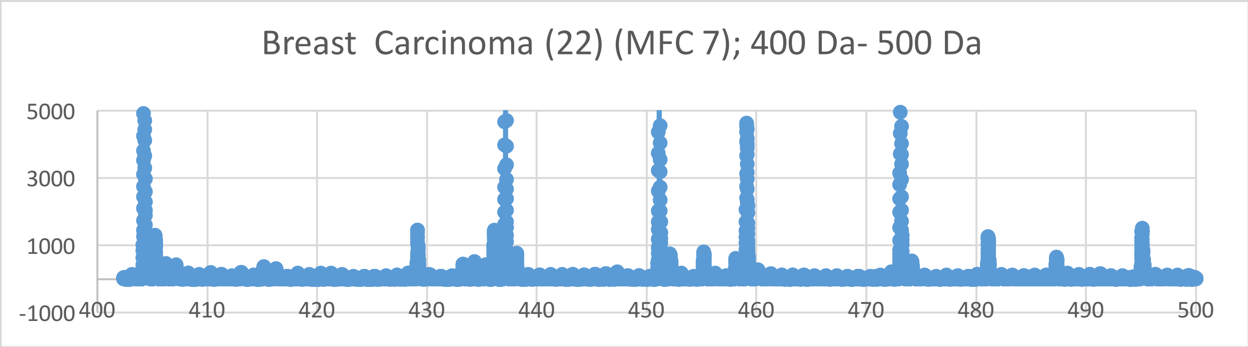
Breast  Carcinoma (22) (MFC 7); 400 Da- 500 Da
5000
3000
1000
-1000
400
410
420
430
440
450
460
470
480
490
500

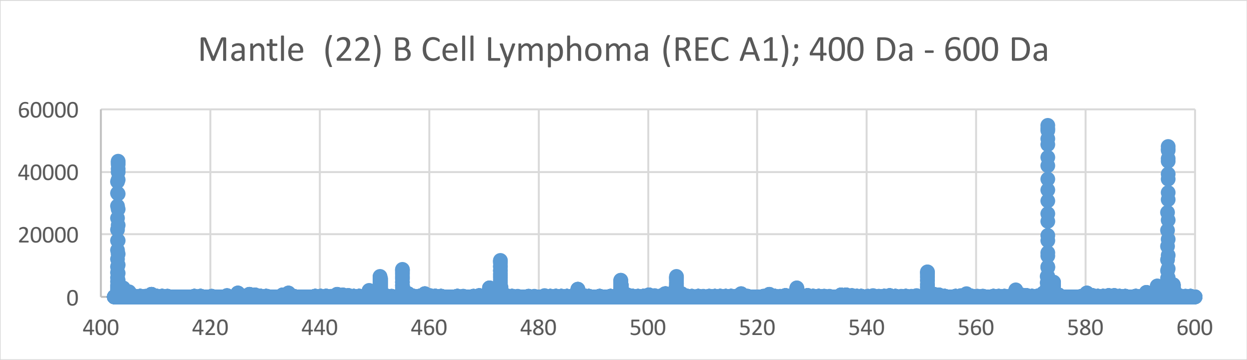
Mantle (22) B Cell Lymphoma (REC A1); 400 Da - 600 Da
60000
40000
20000
0
400
420
440
460
480
500
520
540
560
580
600

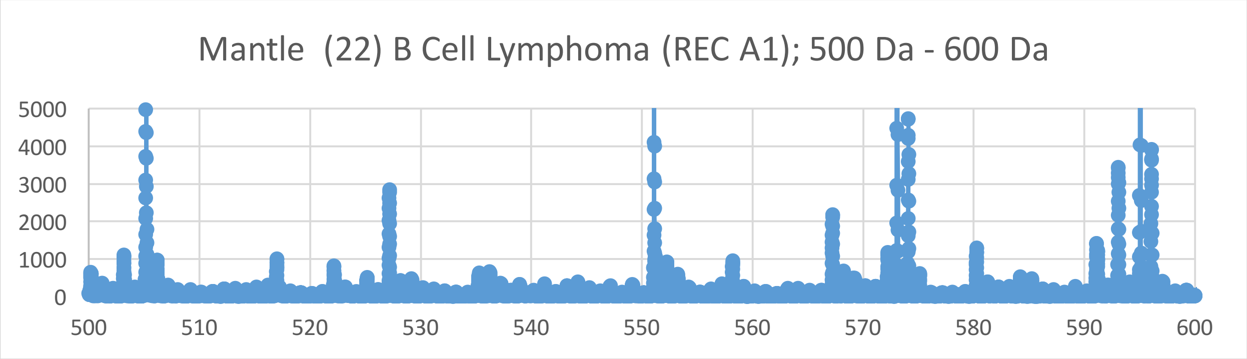
Mantle (22) B Cell Lymphoma (REC A1); 500 Da - 600 Da
5000
4000
3000
2000
1000
0
500
510
520
530
540
550
560
570
580
590
600

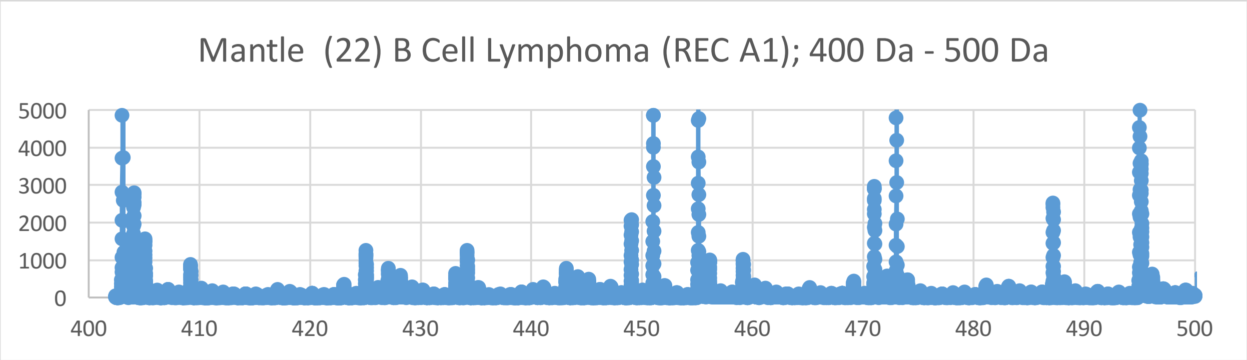
Mantle (22) B Cell Lymphoma (REC A1); 400 Da - 500 Da
5000
4000
3000
2000
1000
0
400
410
420
430
440
450
460
470
480
490
500

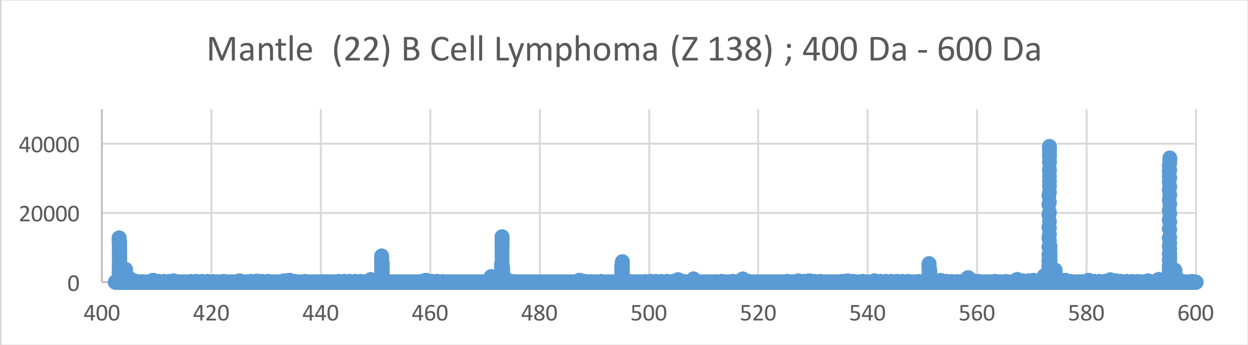
Mantle (22) B Cell Lymphoma (Z 138) ; 400 Da - 600 Da
40000
20000
0
400
420
440
460
480
500
520
540
560
580
600

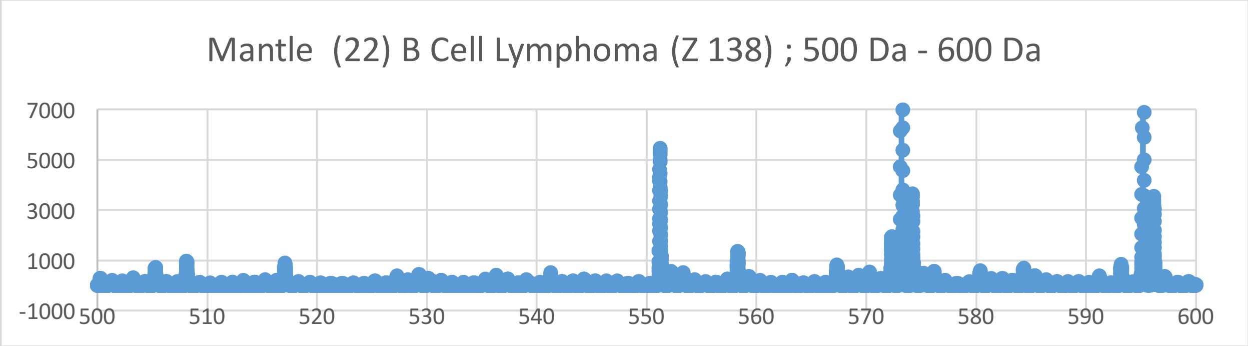
Mantle (22) B Cell Lymphoma (Z 138) ; 500 Da - 600 Da
7000
5000
3000
1000
-1000
500
510
520
530
540
550
560
570
580
590
600

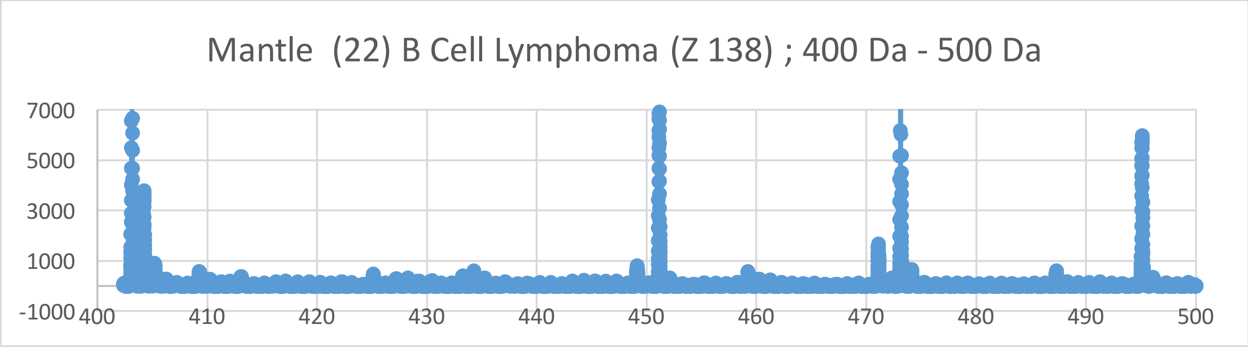
Mantle (22) B Cell Lymphoma (Z 138) ; 400 Da - 500 Da
7000
5000
3000
1000
-1000
400
410
420
430
440
450
460
470
480
490
500

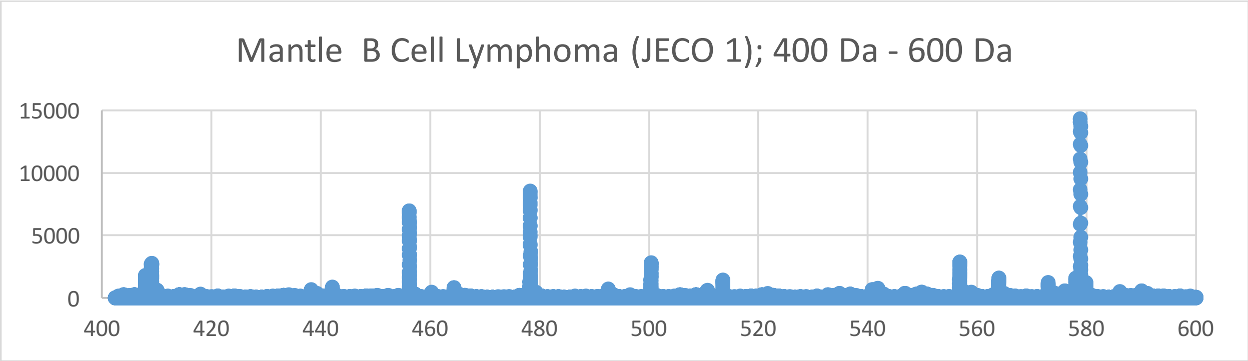
Mantle B Cell Lymphoma (JECO 1); 400 Da - 600 Da
15000
10000
5000
0
400
420
440
460
480
500
520
540
560
580
600

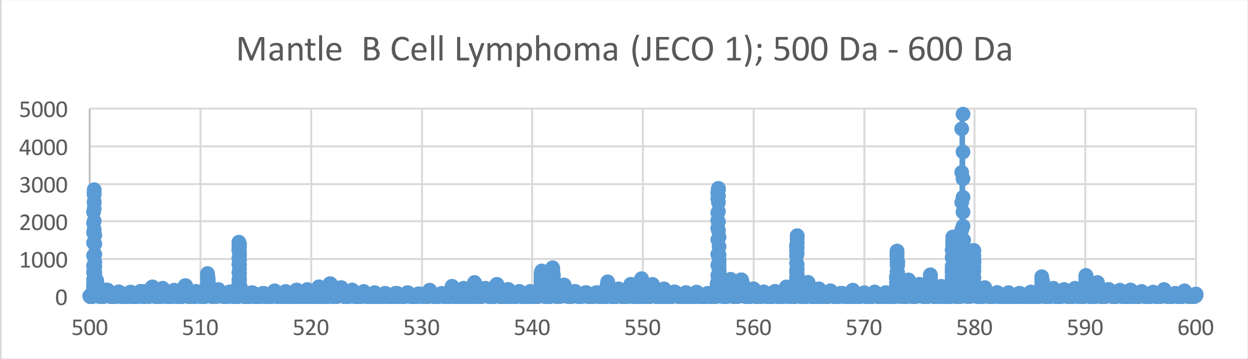
Mantle B Cell Lymphoma (JECO 1); 500 Da - 600 Da
5000
4000
3000
2000
1000
0
500
510
520
530
540
550
560
570
580
590
600

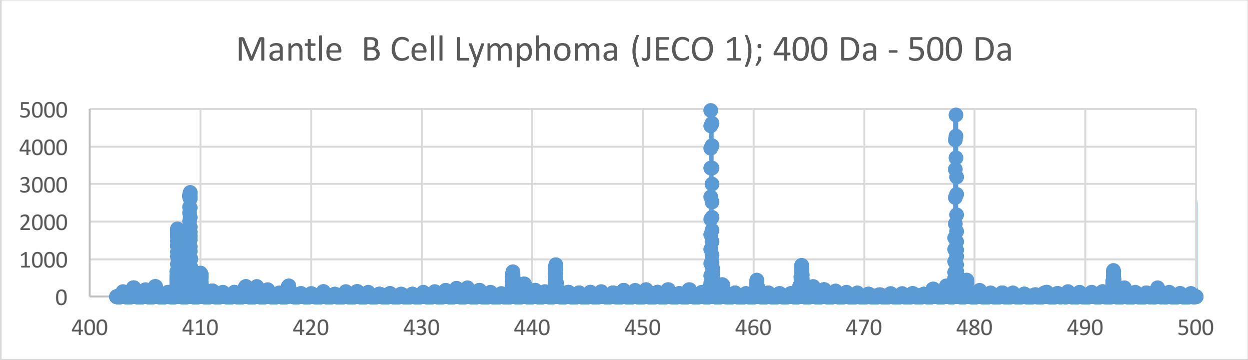
Mantle B Cell Lymphoma (JECO 1); 400 Da - 500 Da
5000
4000
3000
2000
1000
0
400
410
420
430
440
450
460
470
480
490
500

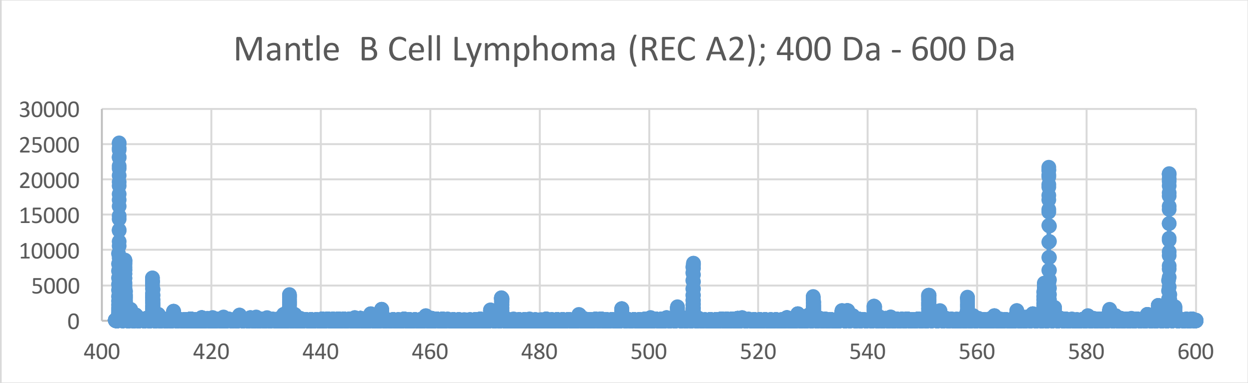
Mantle B Cell Lymphoma (REC A2); 400 Da - 600 Da
30000
25000
20000
15000
10000
5000
0
400 420 440 460 480 500 520 540 560 580 600

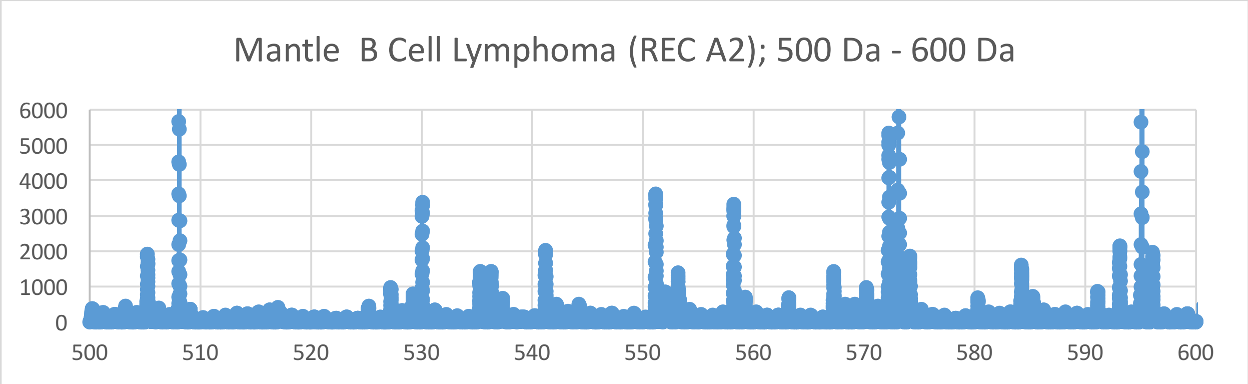
Mantle B Cell Lymphoma (REC A2); 500 Da - 600 Da
6000
5000
4000
3000
2000
1000
0
500 510 520 530 540 550 560 570 580 590 600

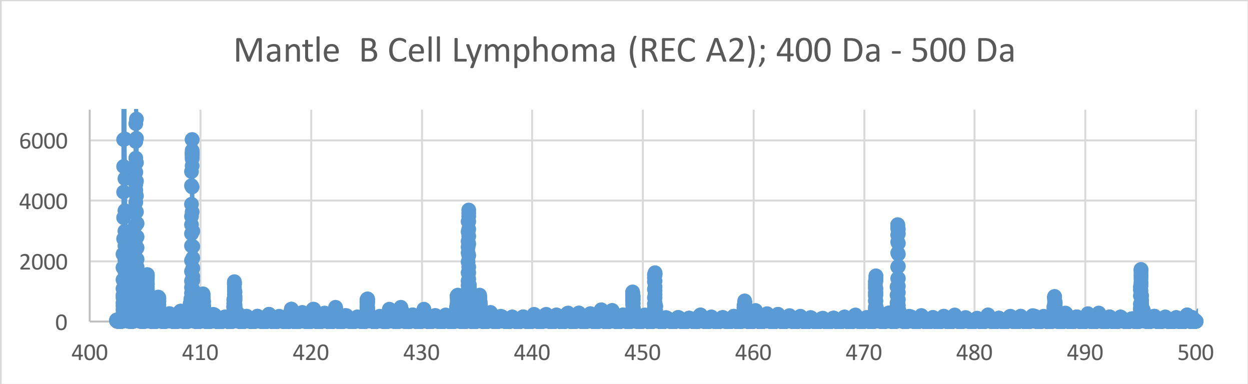
Mantle B Cell Lymphoma (REC A2); 400 Da - 500 Da
6000
4000
2000
0
400 410 420 430 440 450 460 470 480 490 500

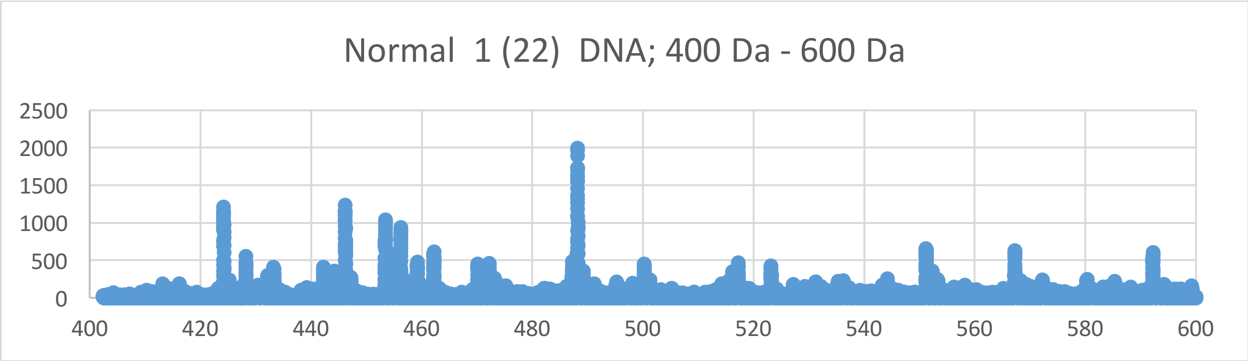
Normal 1 (22) DNA; 400 Da - 600 Da
2500
2000
1500
1000
500
0
400
420
440
460
480
500
520
540
560
580
600

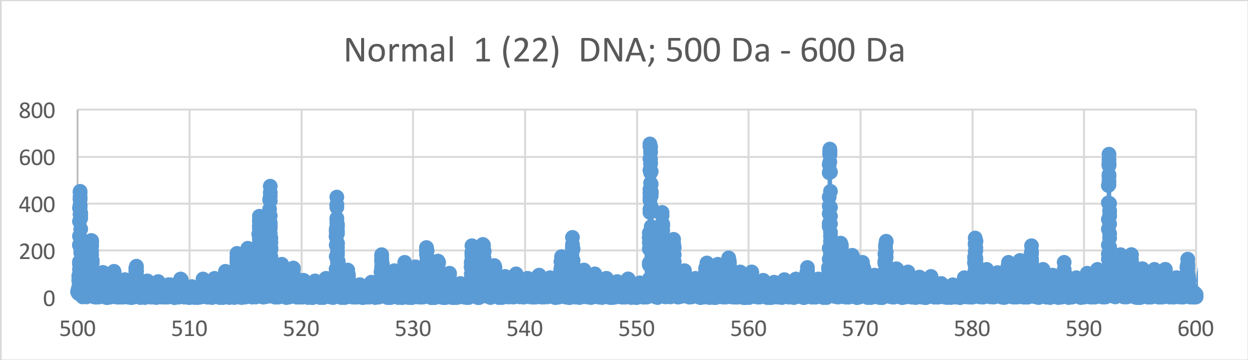
Normal 1 (22) DNA; 500 Da - 600 Da
800
600
400
200
0
500
510
520
530
540
550
560
570
580
590
600

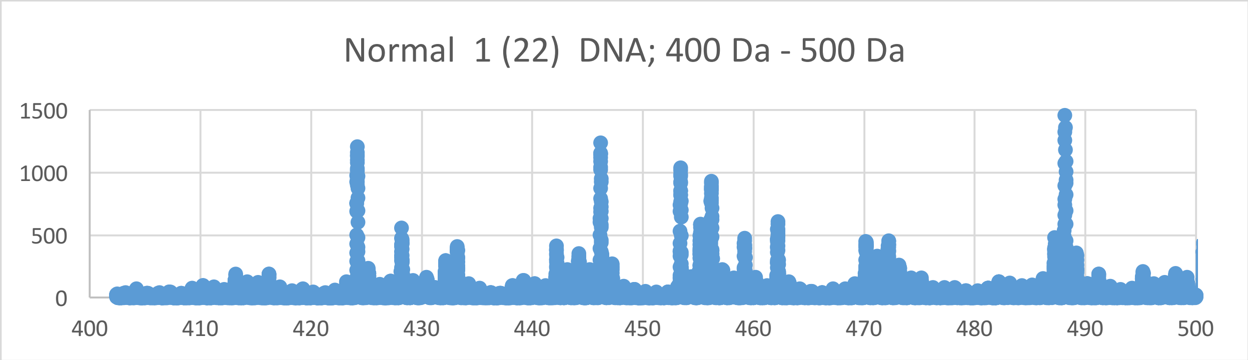
Normal 1 (22) DNA; 400 Da - 500 Da
1500
1000
500
0
400
410
420
430
440
450
460
470
480
490
500

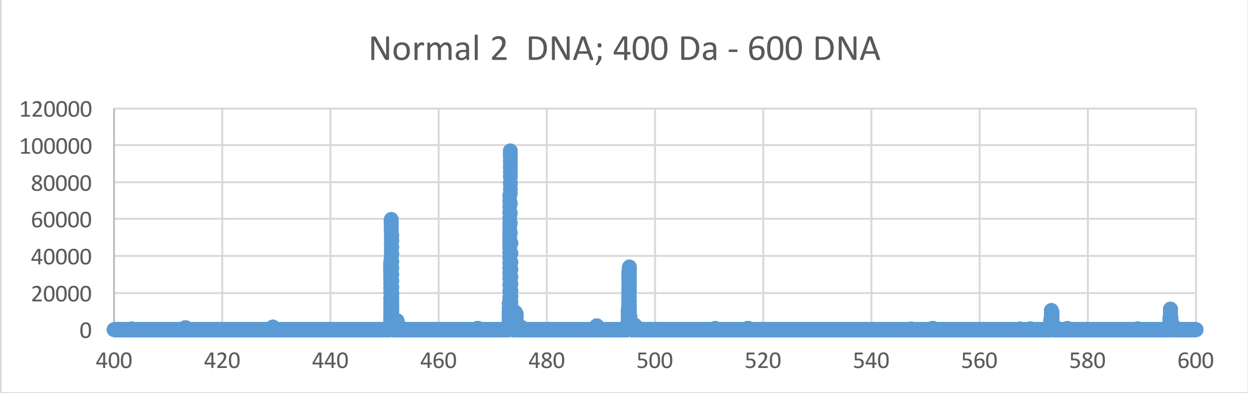
Normal 2 DNA; 400 Da - 600 DNA
120000
100000
80000
60000
40000
20000
0
400 420 440 460 480 500 520 540 560 580 600

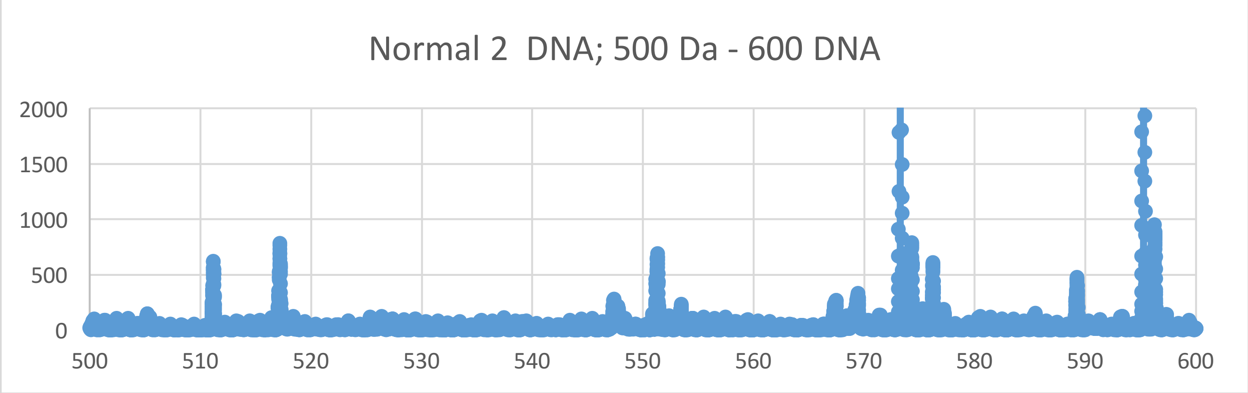
Normal 2 DNA; 500 Da - 600 DNA
2000
1500
1000
500
0
500 510 520 530 540 550 560 570 580 590 600

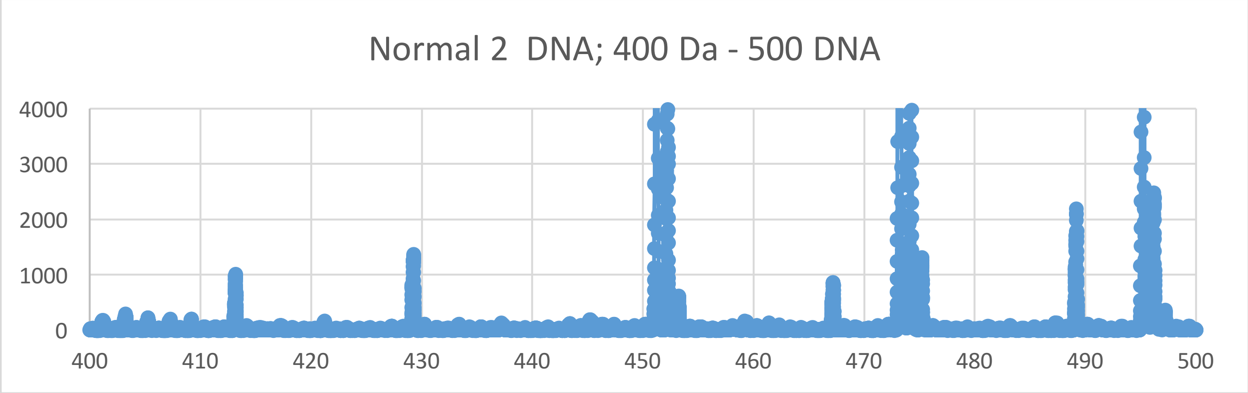
Normal 2 DNA; 400 Da - 500 DNA
4000
3000
2000
1000
0
400 410 420 430 440 450 460 470 480 490 500

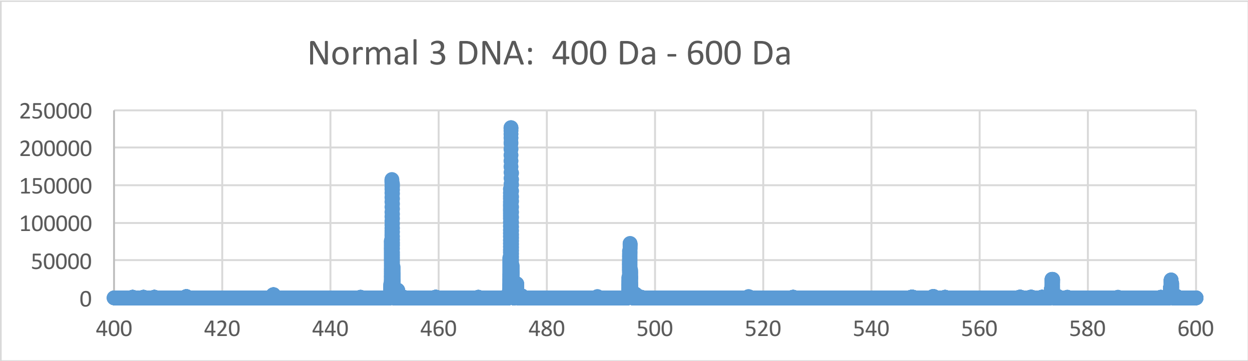
Normal 3 DNA: 400 Da - 600 Da
250000
200000
150000
100000
50000
0
400
420
440
460
480
500
520
540
560
580
600

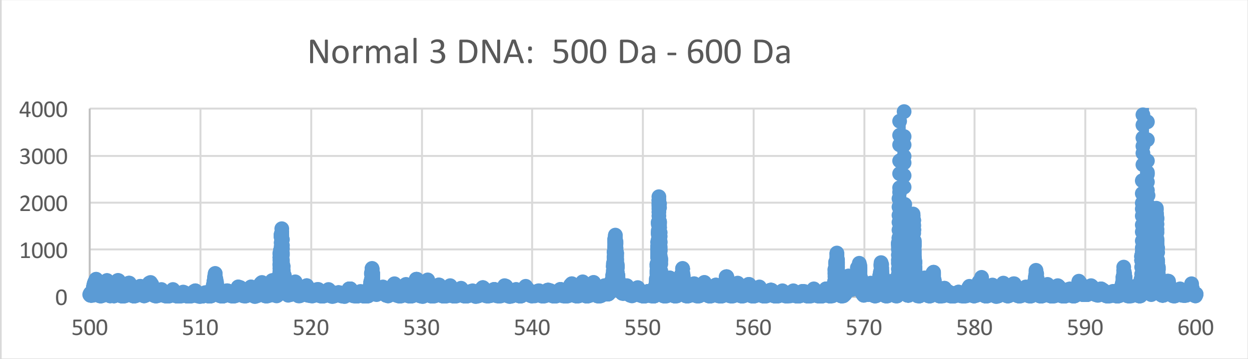
Normal 3 DNA: 500 Da - 600 Da
4000
3000
2000
1000
0
500
510
520
530
540
550
560
570
580
590
600

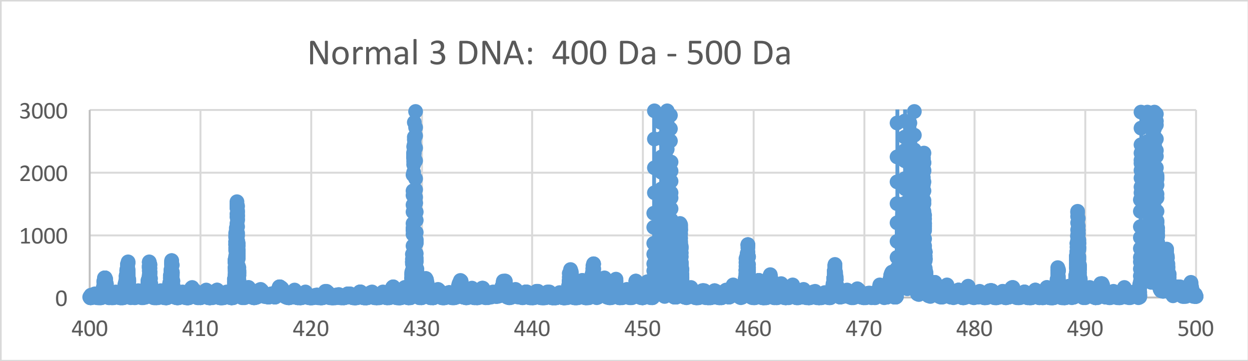
Normal 3 DNA: 400 Da - 500 Da
3000
2000
1000
0
400
410
420
430
440
450
460
470
480
490
500

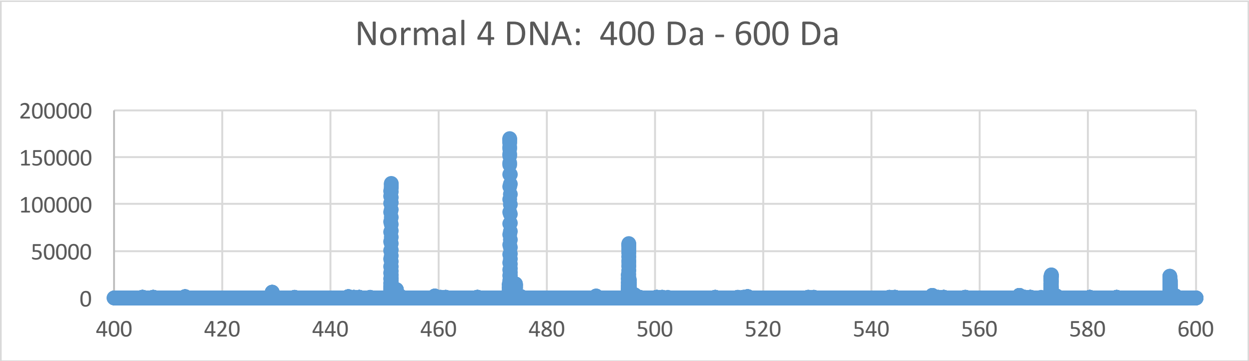
Normal 4 DNA: 400 Da - 600 Da
200000
150000
100000
50000
0
400 420 440 460 480 500 520 540 560 580 600

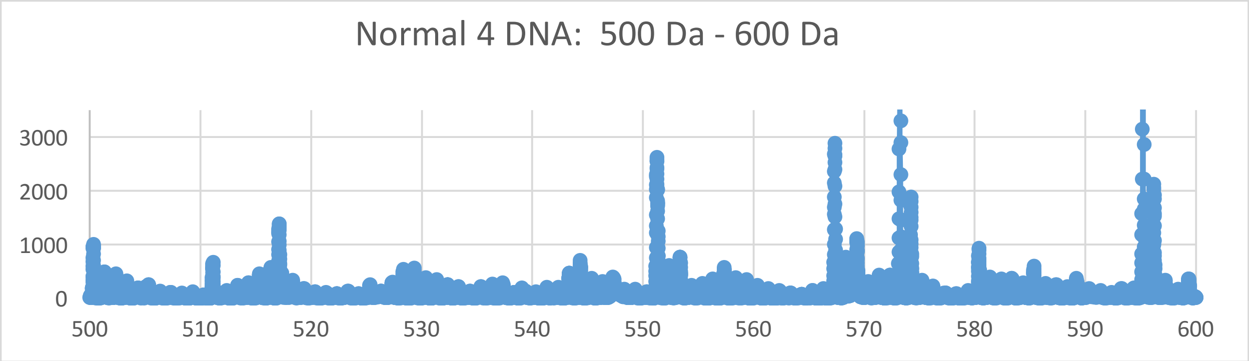
Normal 4 DNA: 500 Da - 600 Da
3000
2000
1000
0
500 510 520 530 540 550 560 570 580 590 600

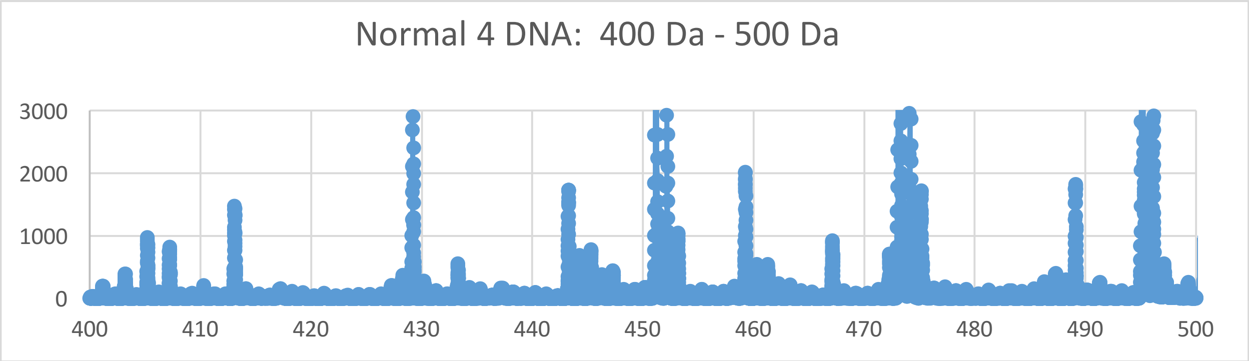
Normal 4 DNA: 400 Da - 500 Da
3000
2000
1000
0
400 410 420 430 440 450 460 470 480 490 500
