## Supplementary Materials XII for "Oncogenesis and Aging by Isotopic Functionalizations of the Proteins and Nucleic Acids"

XII Supplementary Materials – New Mass Spectra from 100 Da - 400 Da

Results for Supplementary 12

So for the mononucleotides at $(phos)_2$, the A, T, G and C peaks for cancer negligibly change in peak intensities and increase in numbers with the $(phos)_2$ to $(phos)_1$ dephosphorylations. The mononucleotides negligibly change fragmentation patterns for cancer at $(phos)_1$. But as decrease the phosphate by 1 unit, the mononucleotides change so that the cancer mononucleotides more fragment and break with stabilities from the DNA with more number and stabilities for the A, T, C and G for $(phos)_2$ to $(phos)_1$. The masses range from 469 Da to 304 Da. The normal fragment more intense mononucleotides in this mass range. See Supplementary Figures 12 for graphs. It seems that below the saturation range of some number of phosphates, the telomeres, mating telomeres and nontelomeres behave in similar way within cancer and normal DNA. So this suggests cancer may be favored and distinguished from normal by changing saturated phosphorylations from $(phos)_2$ to $(phos)_1$. The A, T, C and G mononucleotide peaks for cancer negligibly increase with these dephosphorylations of the mononucleotides. The dephosphorylations from $(phos)_2$ to $(phos)_1$ caused the resulting negligible intense and stable cancer mononucleotides to be more telomeric, mating telomeric and nontelomeric mononucleotides and strongly negligibly isotopically enriched for telomeres, mating telomeres and nontelomeres. It is important to consider that there are more intense peaks for the mononucleotides relative to the dinucleotides, trinucleotides and tetranucleotides for the saturated phosphate units. For the mononucleotides, the intensities of cancer negligibly increased and the intensities of normal fragments decreased from $(phos)_2$ to $(phos)_{1.}$ The dephosphorylations of telomeric, mating telomeric and nontelomeric mononucleotides from $(phos)_2$ to $(phos)_1$ negligibly increases isotopic enrichments of heavy isotopes of $^{15}N$, $^{13}C$ and $^{17}O$. The dephosphorylations from $(phos)_2$ to $(phos)_1$ also caused the telomeric, mating telomeric and nontelomeric mononucleotides to be negligibly more intense in cancer DNA and negligibly more isotopically enriched. It seems the dephosphorylations are related to the isotopic enrichments and depletions; but the telomeres, mating telomeres and nontelomeres are negligibly more intense in cancer and negligibly more isotopically enriched. The telomeric, mating telomeric and nontelomeric mononucleotides of cancer tended to also be negligibly isotopically enriched; so it seems the cancer stabilities of the telomeric, mating telomeric and nontelomeric mononucleotides are related to their negligibly isotopic enrichments from the dephosphorylations from $(phos)_2$ to $(phos)_1$ and the mutations purely driven by dephosphorylations and enrichments. Enrichments weaken the nontelomeric normal mononucleotides while negligibly strengthening cancerous telomeric mononucleotides. The negligibly intense cancer telomeric, mating telomeric and nontelomeric mononucleotides were also negligibly more isotopically enriched, but negligibly intense cancerous telomeric, mating telomeric and nontelomeric mononucleotides were negligibly enriched isotopically. The dephosphorylations from $(phos)_2$ to $(phos)_1$ negligibly increased the isotopic enrichments of the cancer for the telomeric, nontelomeric and mating telomeric mononucleotides. Such seems compatible for replications and transcriptions as stable telomeres seem compatible to unstable mating telomeres for the unravelings and constructions of DNA mating telomeres onto the building templates of the telomeres. But in this case of all normal telomeric, mating telomeric and nontelomeric mononucleotides are negligibly enriched and unstable then the replications, transcriptions and translations cannot occur normally. The dephosphorylations from $(phos)_2$ to $(phos)_1$ caused demethylations to cause negligibly more cancer intensities and stabilities but lowered isotopic enrichments for G. But the demethylations of the G and A caused negligible alterations cancer stabilities and isotopic enrichments. Where isotopic changes occurred, the functionalization effects were negligible. The methylations of the C mononucleotides negligibly decreased cancer intensities and stabilities with isotopic enrichments. It seems from this data that during biological processes the dephosphorylations and stabilizations of normal or vice versa dephosphorylations and stabilizations of cancer are or can be countered by methylations (and isotopic enrichments) and destabilizations of the normal for cancer or vice versa dephosphorylations and destabilizations of the cancer by methylations as this research discovers coupled phosphorylations and methylations (and isotopic enrichments) or coupled dephosphorylations and methylations (and isotopic enrichments) with involved greater enrichments for normal biological functions and explaining cancer! The dephosphorylations from $(phos)_2$ to $(phos)_1$ caused acetylations to cause less cancer stabilities for A and C with negligible isotopic depletions and negligible cancer stabilities for G and T with negligible depletions of heavy isotopes. The acetylations from $(phos)_2$ to $(phos)_1$ caused less cancer stabilites and intensities for A and C with negligibly more isotopic depletions. This contrast in A, T, C and G from the de-acetylations and dephosphorylations may reflect the developments of cancer as the mononucleotides fragment but the other mononucleotides more hold together for damaged DNA for cancer.

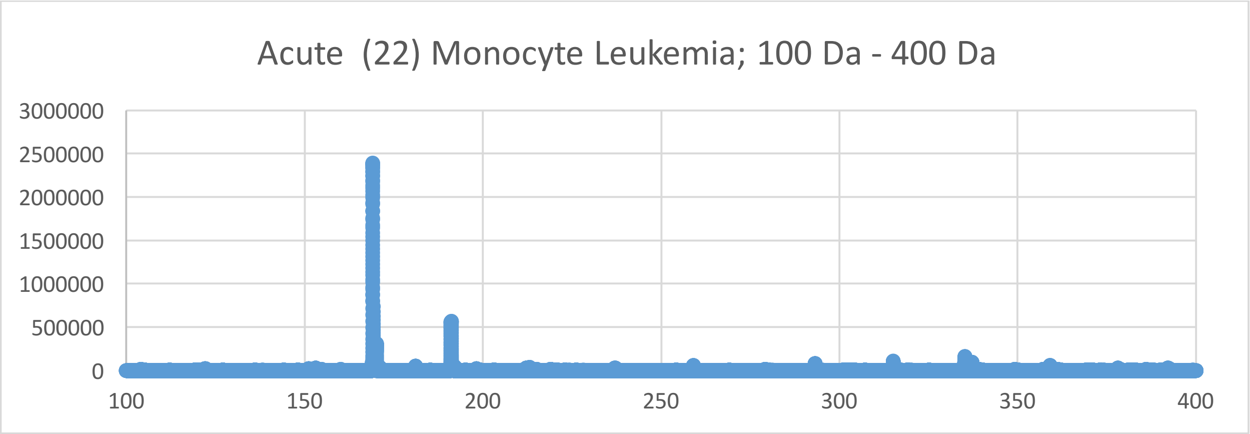
Acute (22) Monocyte Leukemia; 100 Da - 400 Da
3000000
2500000
2000000
1500000
1000000
500000
0
100
150
200
250
300
350
400

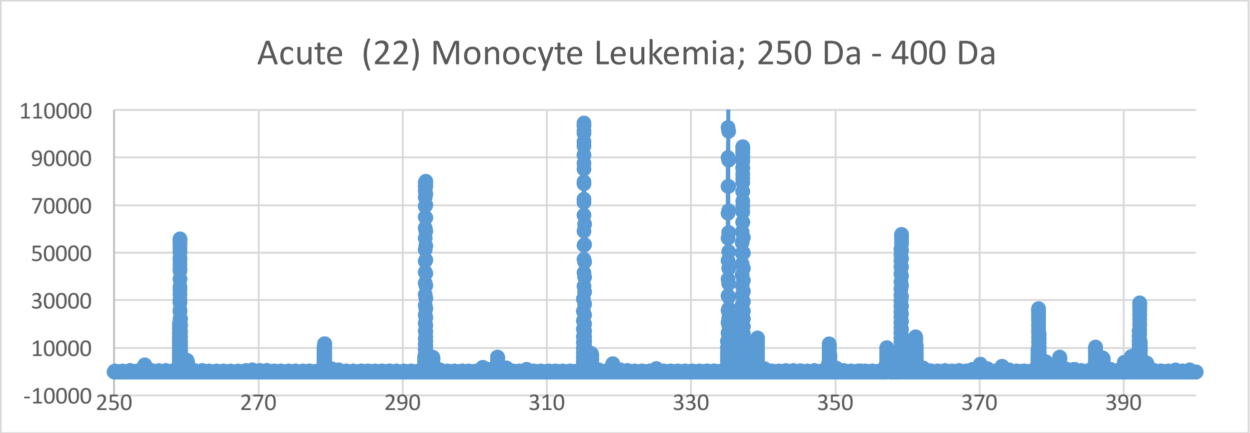
Acute (22) Monocyte Leukemia; 250 Da - 400 Da
110000
90000
70000
50000
30000
10000
-10000
250
270
290
310
330
350
370
390

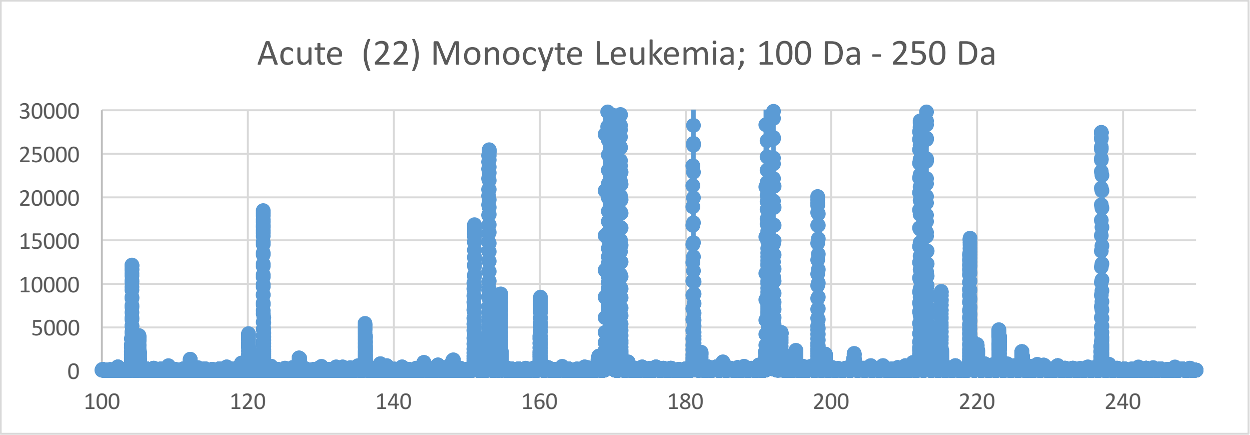
Acute (22) Monocyte Leukemia; 100 Da - 250 Da
30000
25000
20000
15000
10000
5000
0
100
120
140
160
180
200
220
240

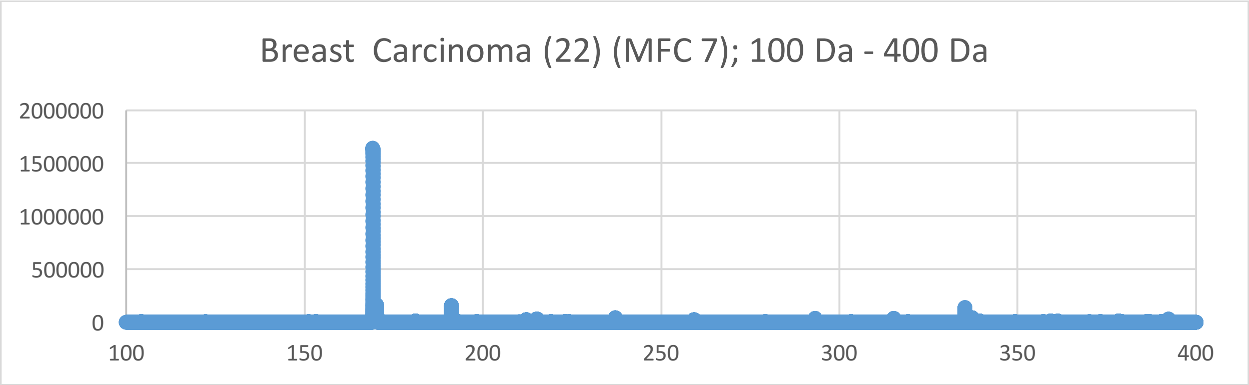
Breast Carcinoma (22) (MFC 7); 100 Da - 400 Da
2000000
1500000
1000000
500000
0
100
150
200
250
300
350
400

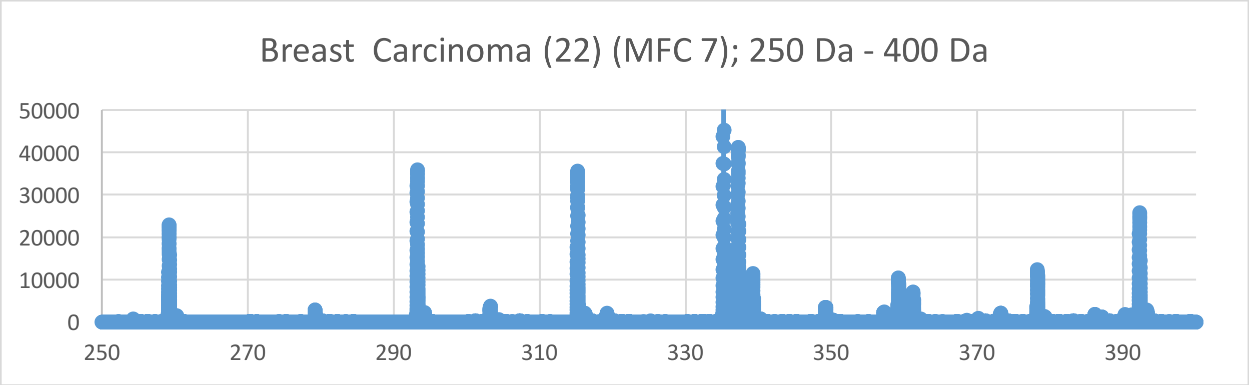
Breast Carcinoma (22) (MFC 7); 250 Da - 400 Da
50000
40000
30000
20000
10000
0
250
270
290
310
330
350
370
390

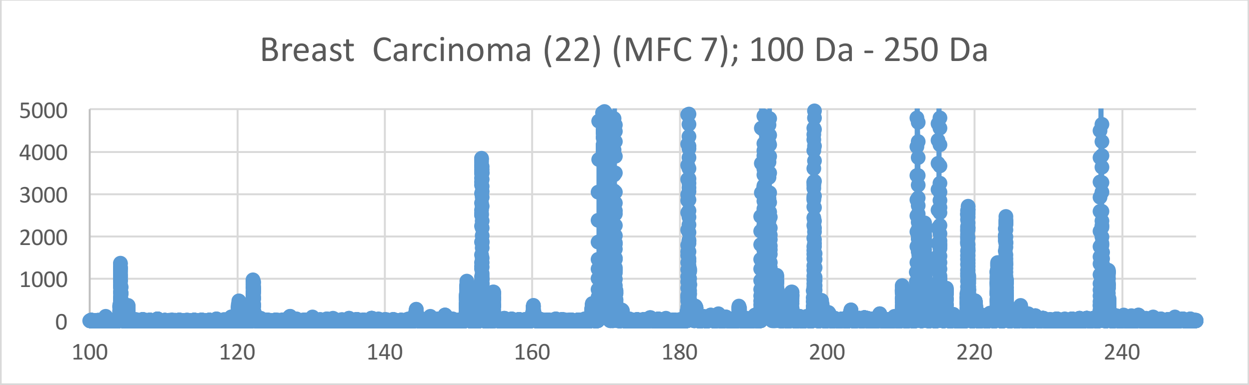
Breast Carcinoma (22) (MFC 7); 100 Da - 250 Da
5000
4000
3000
2000
1000
0
100
120
140
160
180
200
220
240

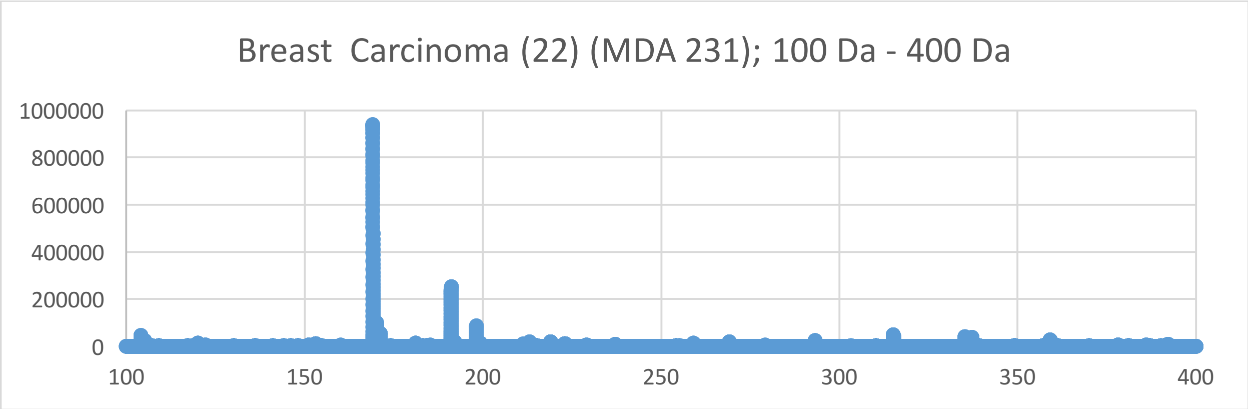
Breast Carcinoma (22) (MDA 231); 100 Da - 400 Da
1000000
800000
600000
400000
200000
0
100
150
200
250
300
350
400

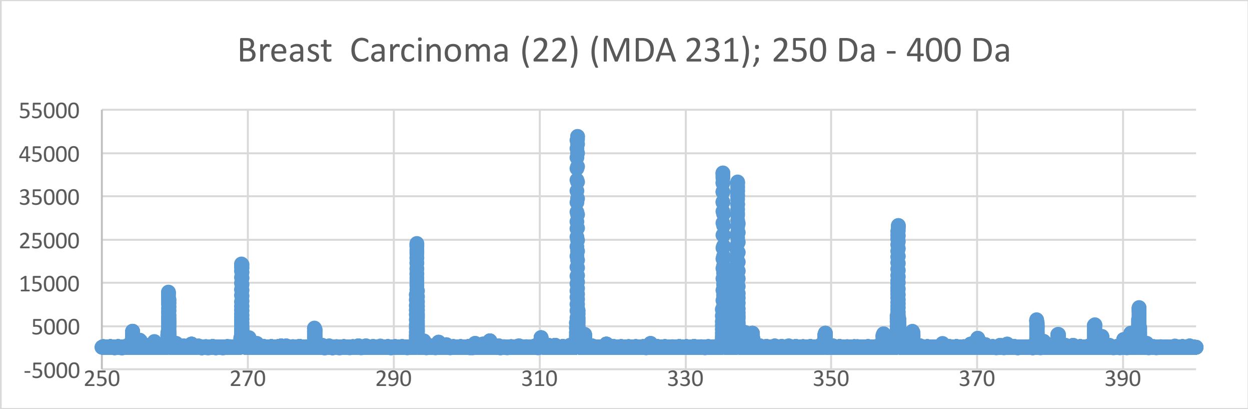
Breast Carcinoma (22) (MDA 231); 250 Da - 400 Da
55000
45000
35000
25000
15000
5000
-5000
250
270
290
310
330
350
370
390

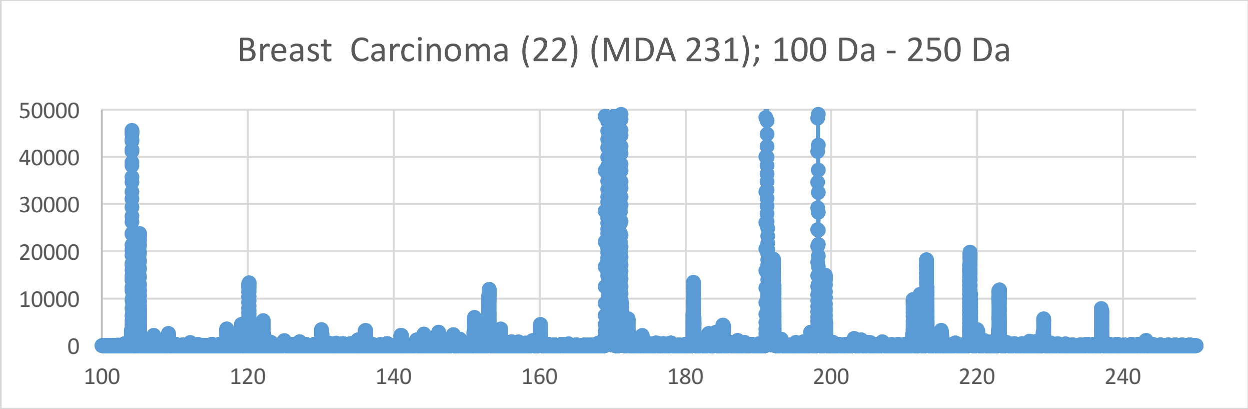
Breast Carcinoma (22) (MDA 231); 100 Da - 250 Da
50000
40000
30000
20000
10000
0
100
120
140
160
180
200
220
240

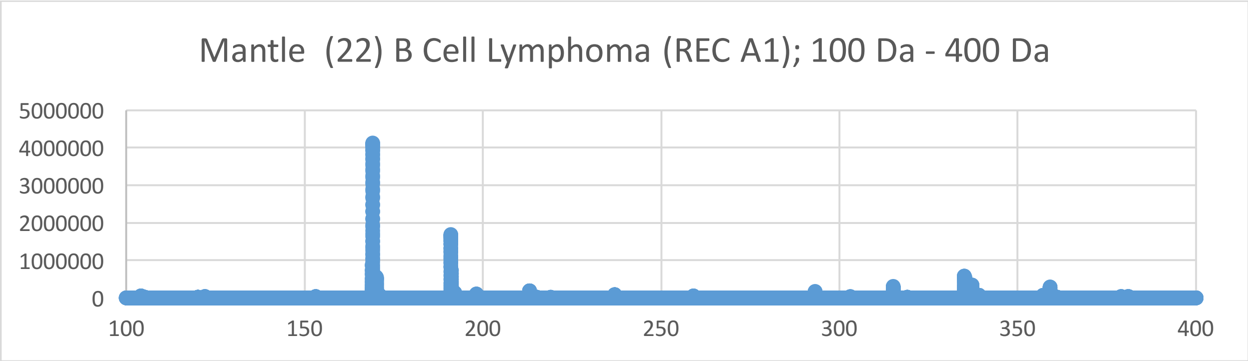
Mantle (22) B Cell Lymphoma (REC A1); 100 Da - 400 Da
5000000
4000000
3000000
2000000
1000000
0
100
150
200
250
300
350
400

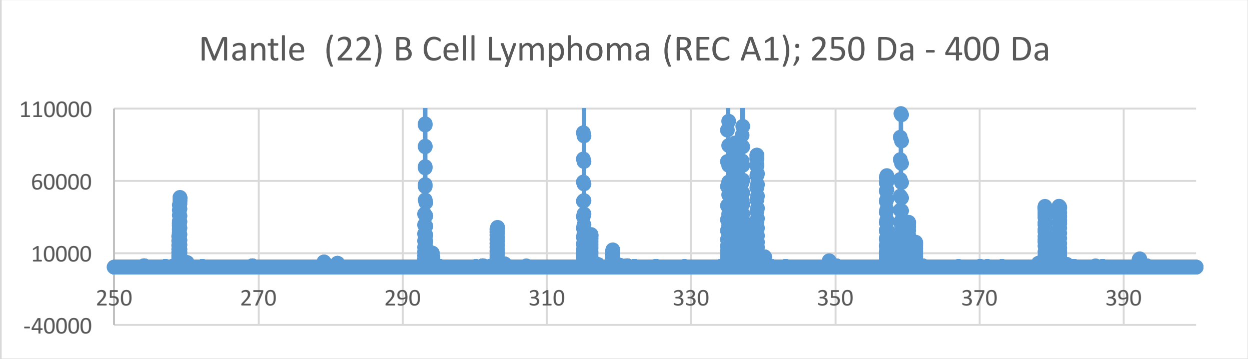
Mantle (22) B Cell Lymphoma (REC A1); 250 Da - 400 Da
110000
60000
10000
-40000
250
270
290
310
330
350
370
390

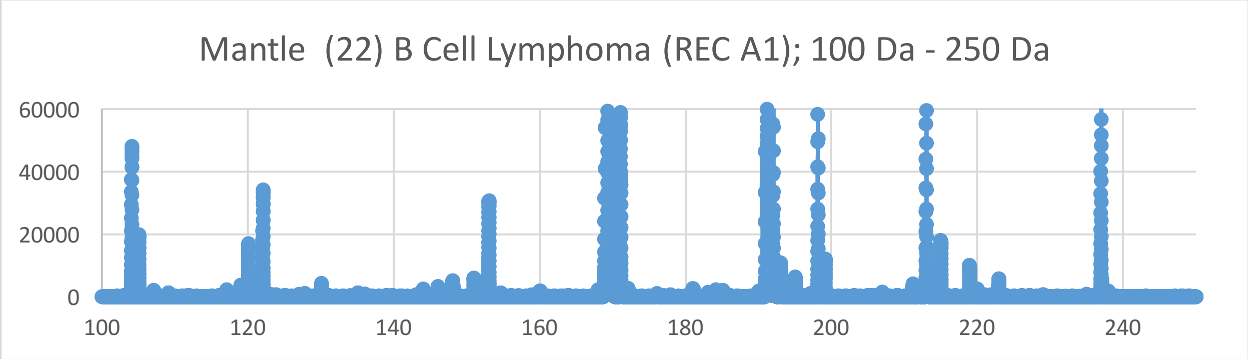
Mantle (22) B Cell Lymphoma (REC A1); 100 Da - 250 Da
60000
40000
20000
0
100
120
140
160
180
200
220
240

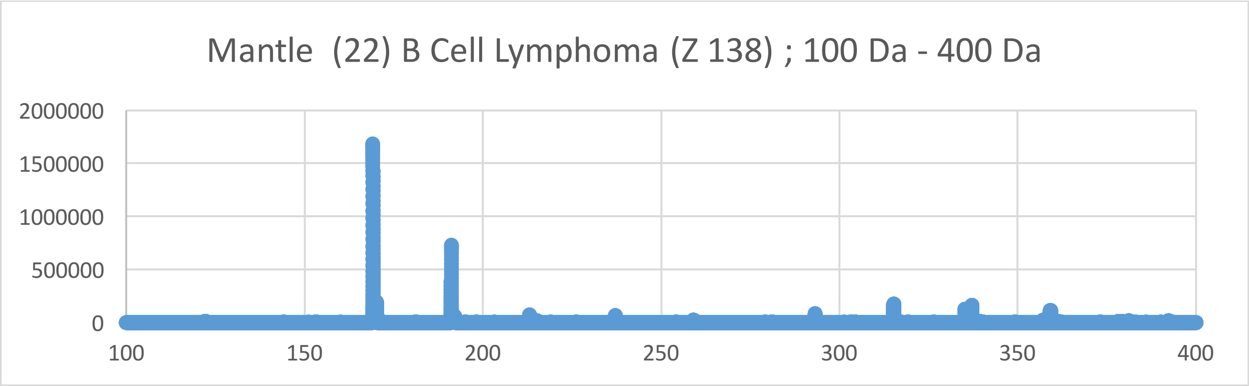
Mantle (22) B Cell Lymphoma (Z 138) ; 100 Da - 400 Da
2000000
1500000
1000000
500000
0
100
150
200
250
300
350
400

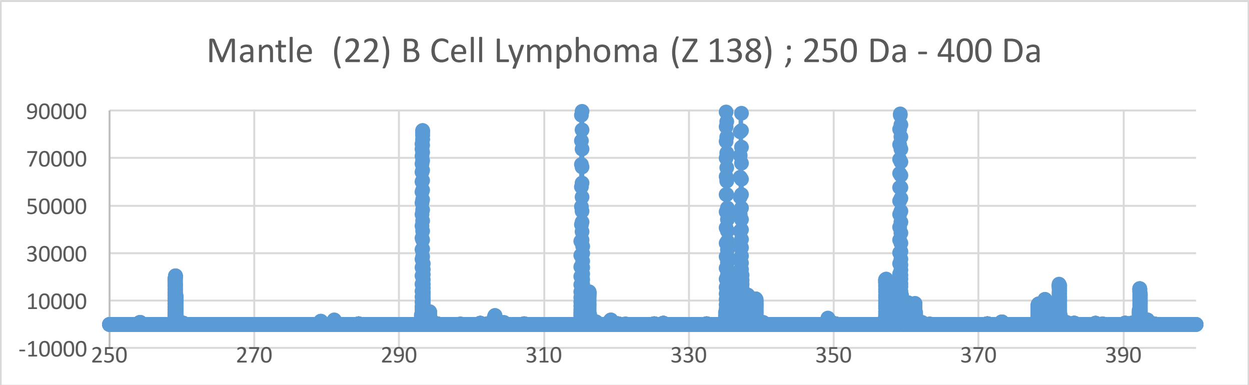
Mantle (22) B Cell Lymphoma (Z 138) ; 250 Da - 400 Da
90000
70000
50000
30000
10000
-10000
250
270
290
310
330
350
370
390

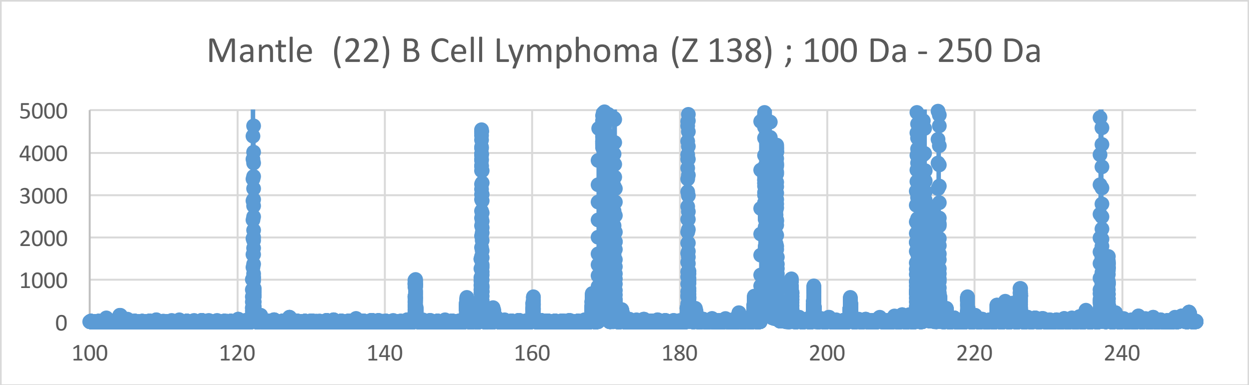
Mantle (22) B Cell Lymphoma (Z 138) ; 100 Da - 250 Da
5000
4000
3000
2000
1000
0
100
120
140
160
180
200
220
240

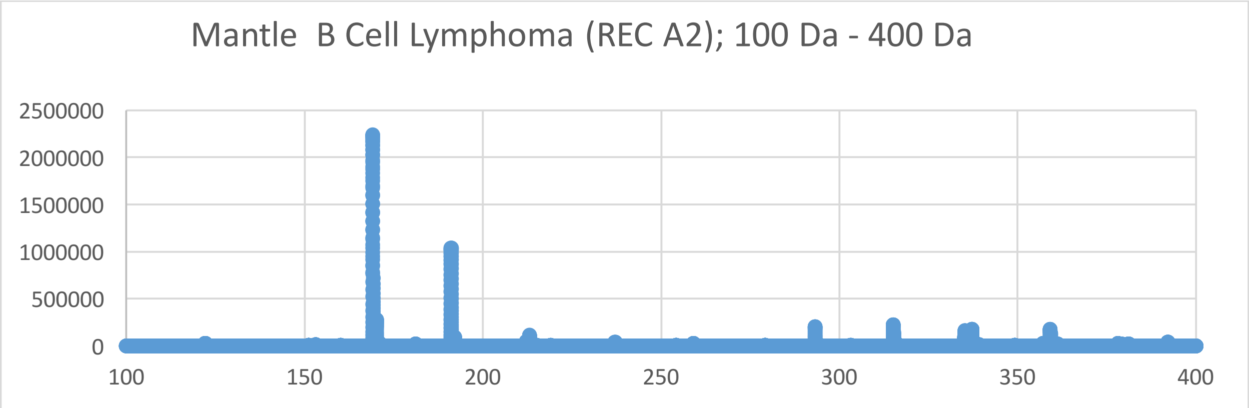
Mantle B Cell Lymphoma (REC A2); 100 Da - 400 Da
2500000
2000000
1500000
1000000
500000
0
100
150
200
250
300
350
400

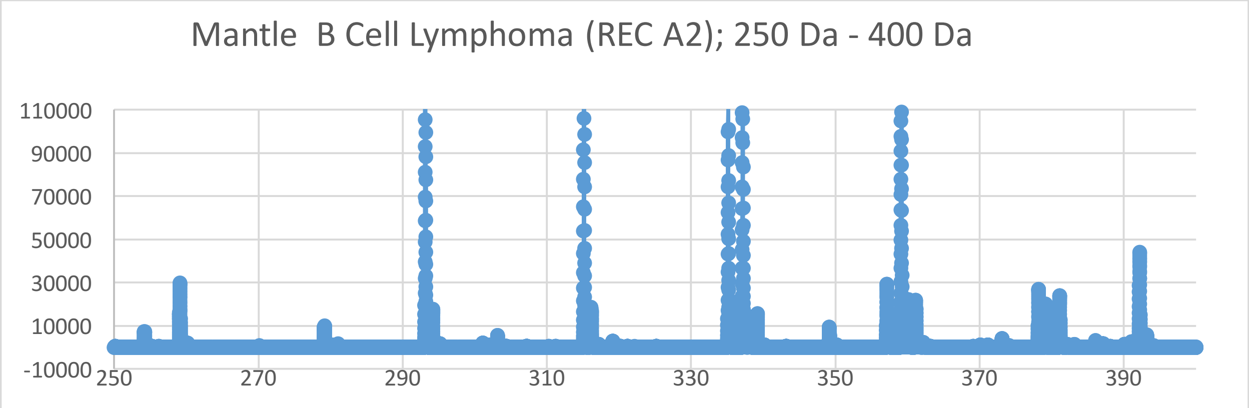
Mantle B Cell Lymphoma (REC A2); 250 Da - 400 Da
110000
90000
70000
50000
30000
10000
-10000
250
270
290
310
330
350
370
390

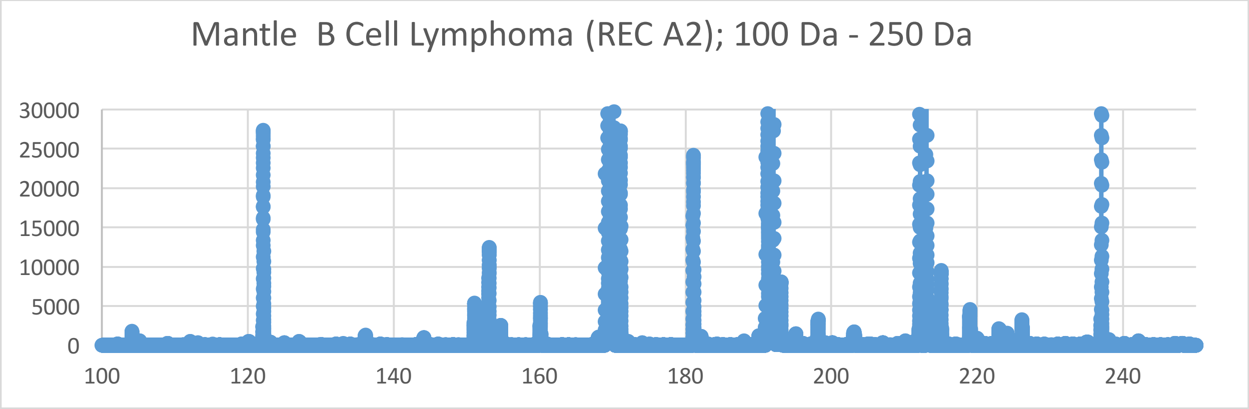
Mantle B Cell Lymphoma (REC A2); 100 Da - 250 Da
30000
25000
20000
15000
10000
5000
0
100
120
140
160
180
200
220
240

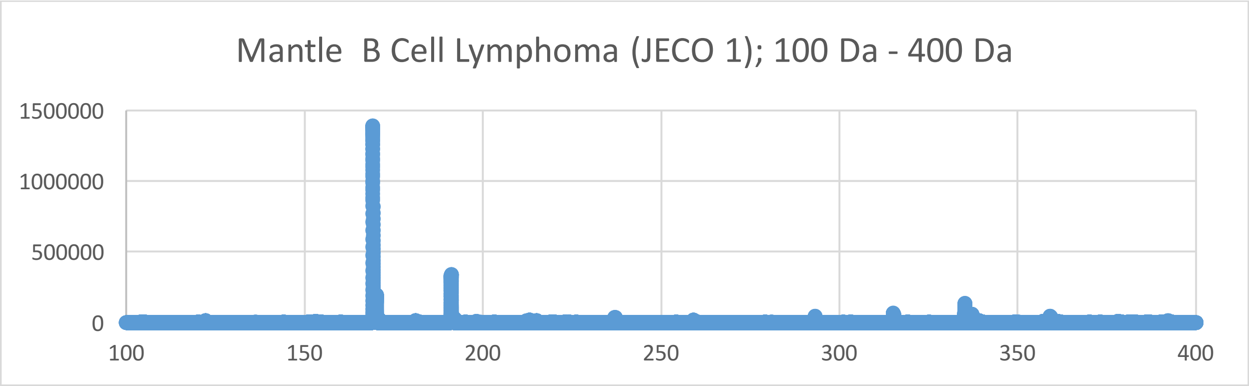
Mantle B Cell Lymphoma (JECO 1); 100 Da - 400 Da
1500000
1000000
500000
0
100
150
200
250
300
350
400

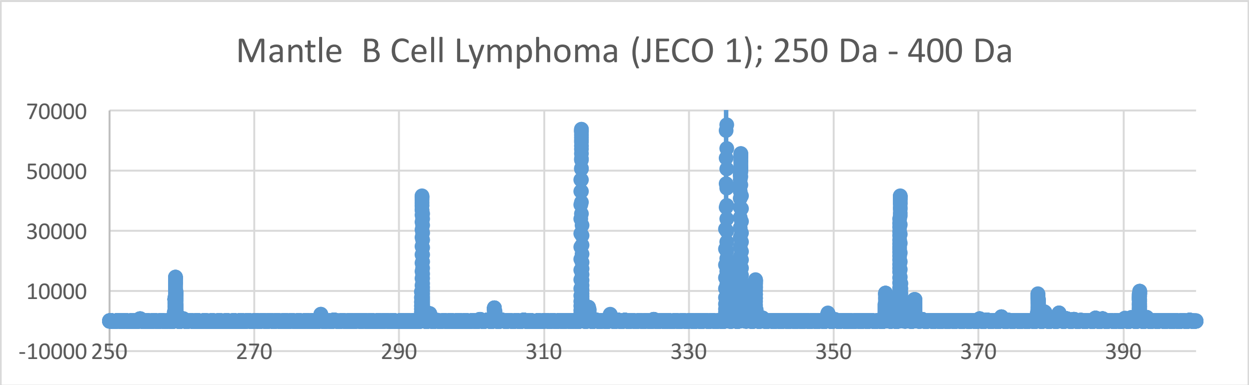
Mantle B Cell Lymphoma (JECO 1); 250 Da - 400 Da
70000
50000
30000
10000
-10000
250
270
290
310
330
350
370
390

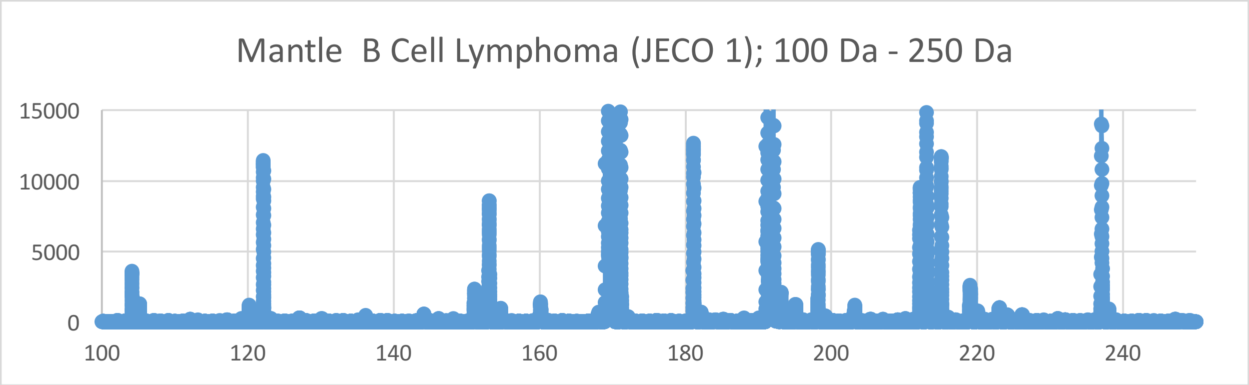
Mantle B Cell Lymphoma (JECO 1); 100 Da - 250 Da
15000
10000
5000
0
100
120
140
160
180
200
220
240

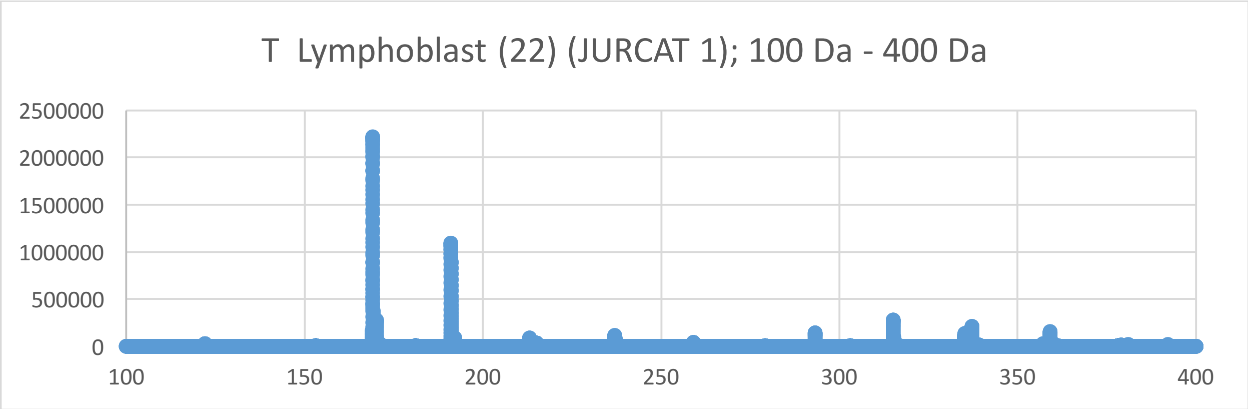
T Lymphoblast (22) (JURCAT 1); 100 Da - 400 Da
2500000
2000000
1500000
1000000
500000
0
100
150
200
250
300
350
400

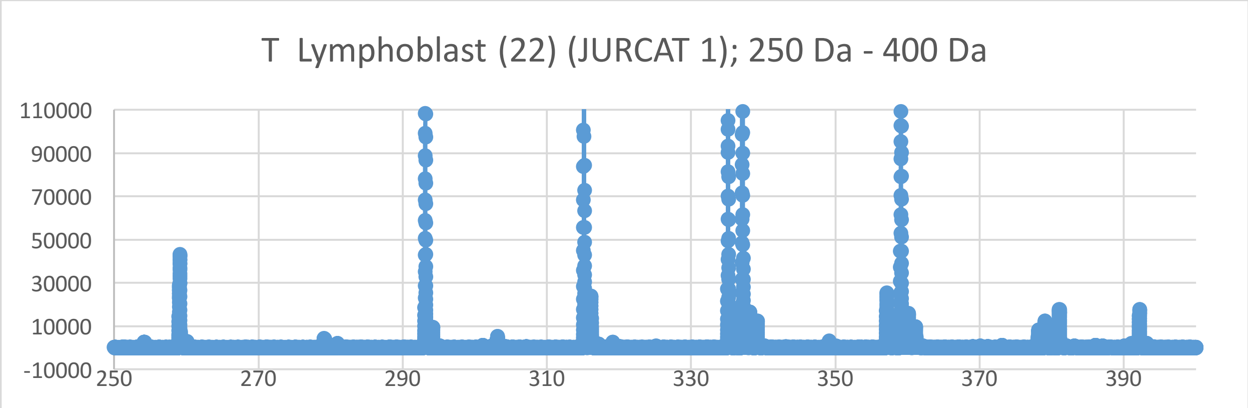
T Lymphoblast (22) (JURCAT 1); 250 Da - 400 Da
110000
90000
70000
50000
30000
10000
-10000
250
270
290
310
330
350
370
390

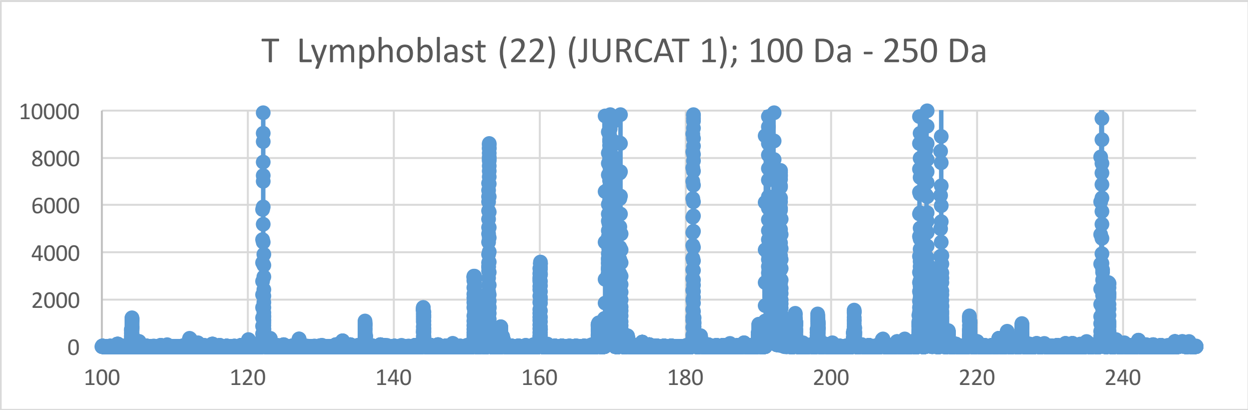
T Lymphoblast (22) (JURCAT 1); 100 Da - 250 Da
10000
8000
6000
4000
2000
0
100
120
140
160
180
200
220
240

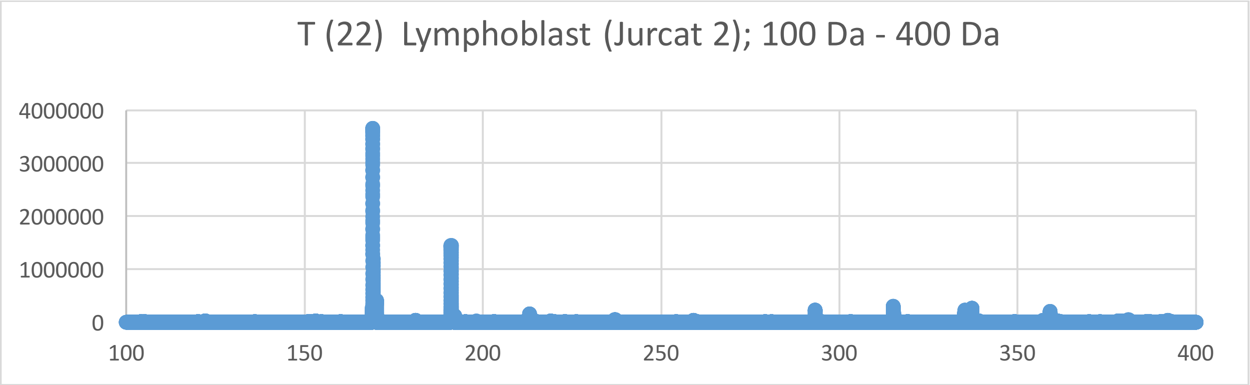
T (22) Lymphoblast (Jurcat 2); 100 Da - 400 Da
4000000
3000000
2000000
1000000
0
100
150
200
250
300
350
400
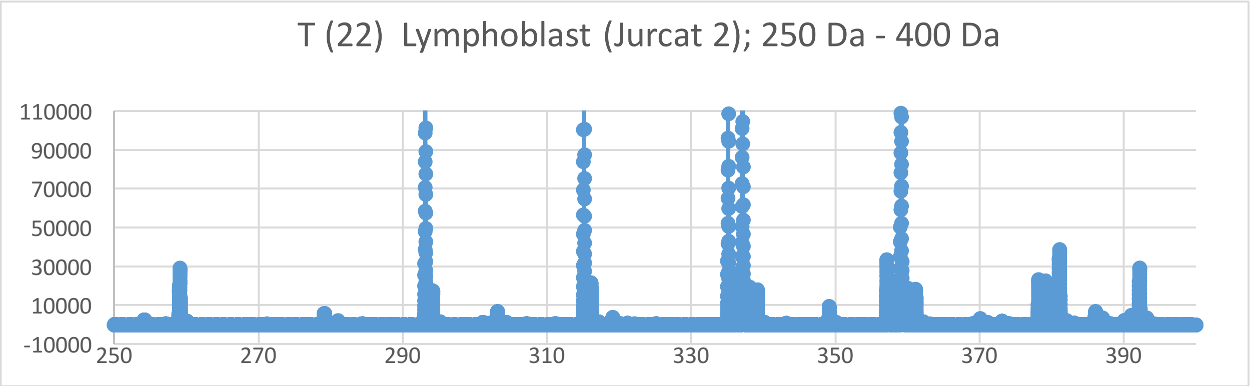
T (22) Lymphoblast (Jurcat 2); 250 Da - 400 Da
110000
90000
70000
50000
30000
10000
-10000
250
270
290
310
330
350
370
390
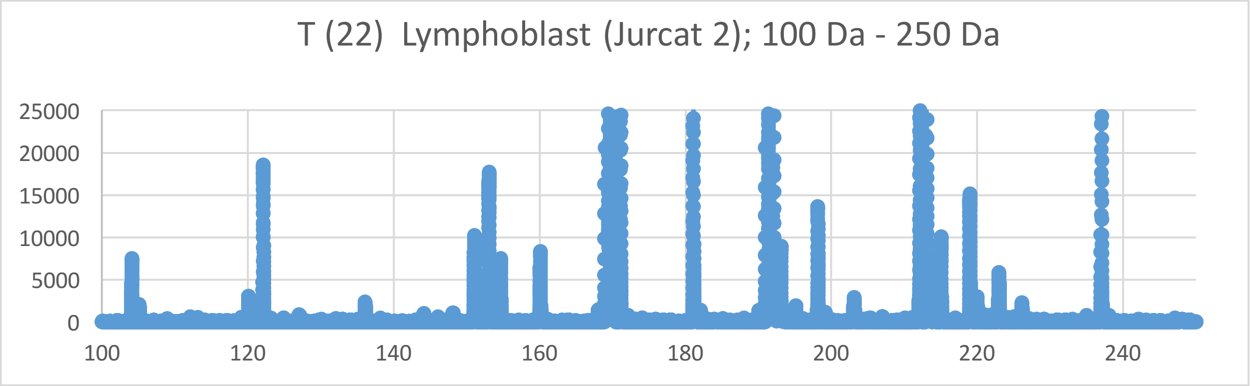
T (22) Lymphoblast (Jurcat 2); 100 Da - 250 Da
25000
20000
15000
10000
5000
0
100
120
140
160
180
200
220
240

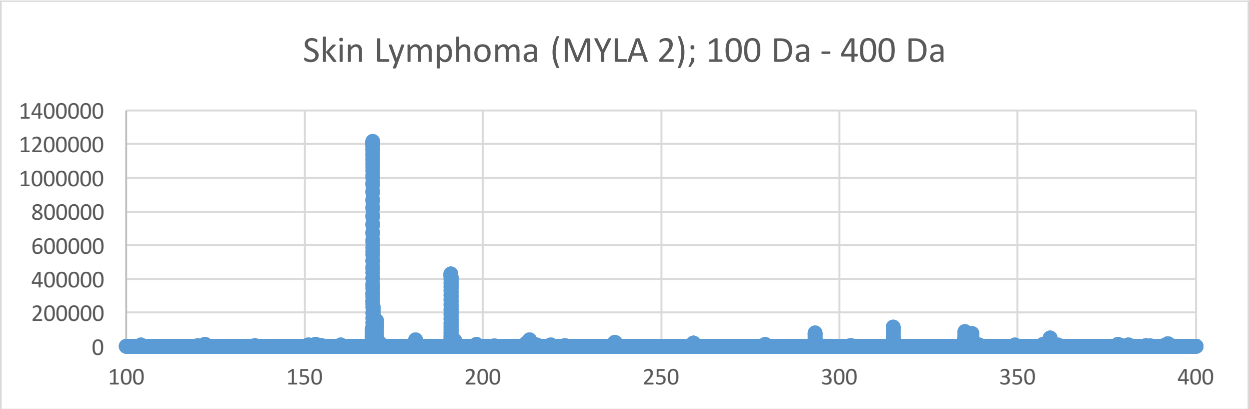
Skin Lymphoma (MYLA 2); 100 Da - 400 Da
1400000
1200000
1000000
800000
600000
400000
200000
0
100
150
200
250
300
350
400

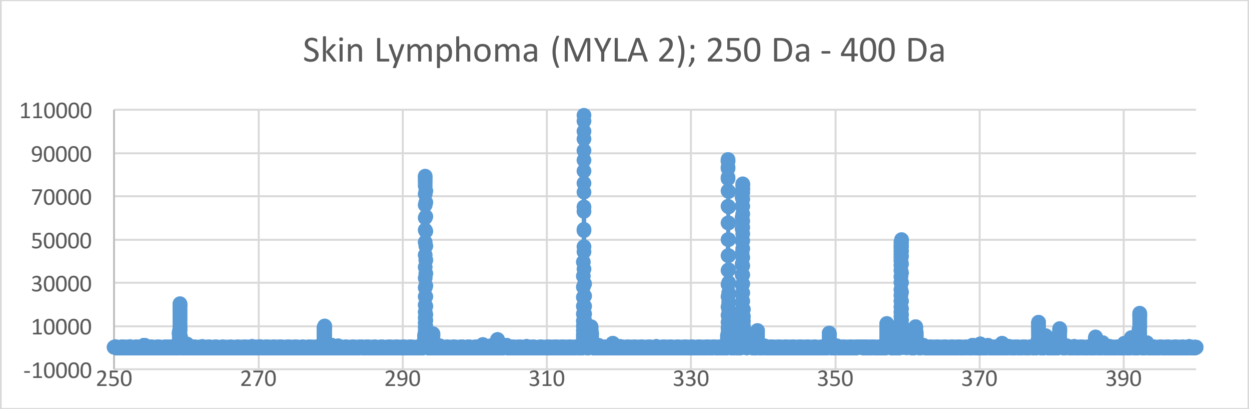
Skin Lymphoma (MYLA 2); 250 Da - 400 Da
110000
90000
70000
50000
30000
10000
-10000
250
270
290
310
330
350
370
390

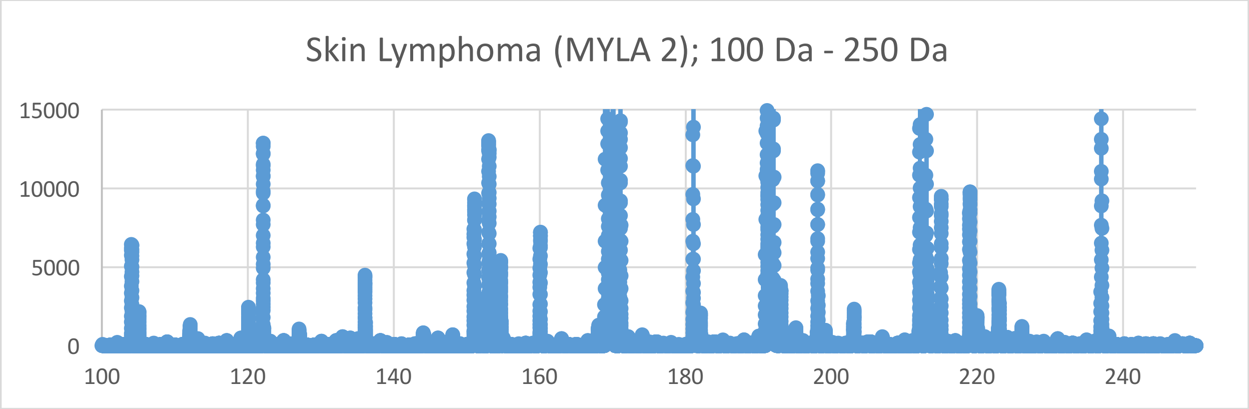
Skin Lymphoma (MYLA 2); 100 Da - 250 Da
15000
10000
5000
0
100
120
140
160
180
200
220
240

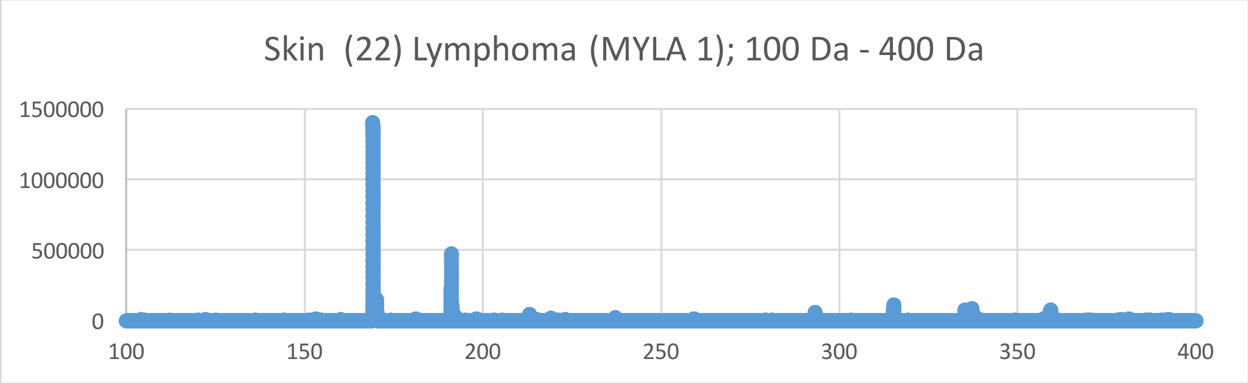
Skin (22) Lymphoma (MYLA 1); 100 Da - 400 Da
1500000
1000000
500000
0
100
150
200
250
300
350
400

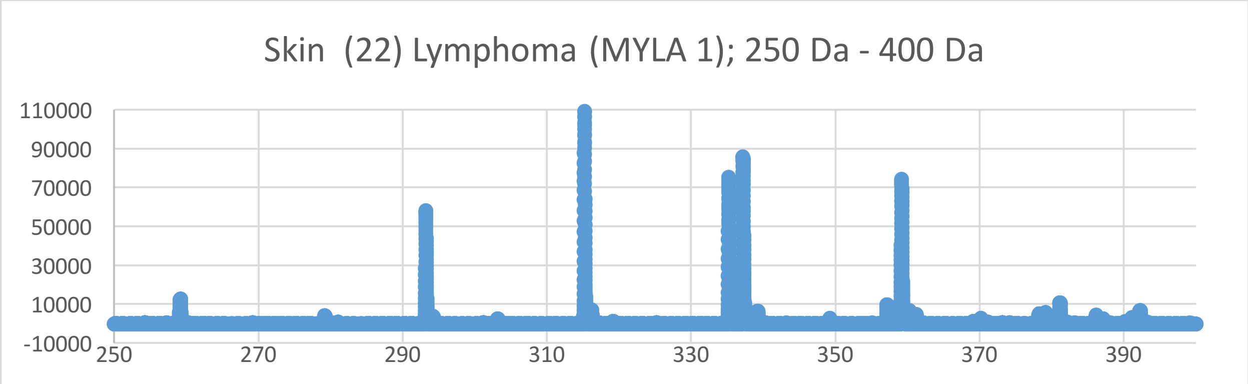
Skin (22) Lymphoma (MYLA 1); 250 Da - 400 Da
110000
90000
70000
50000
30000
10000
-10000
250
270
290
310
330
350
370
390

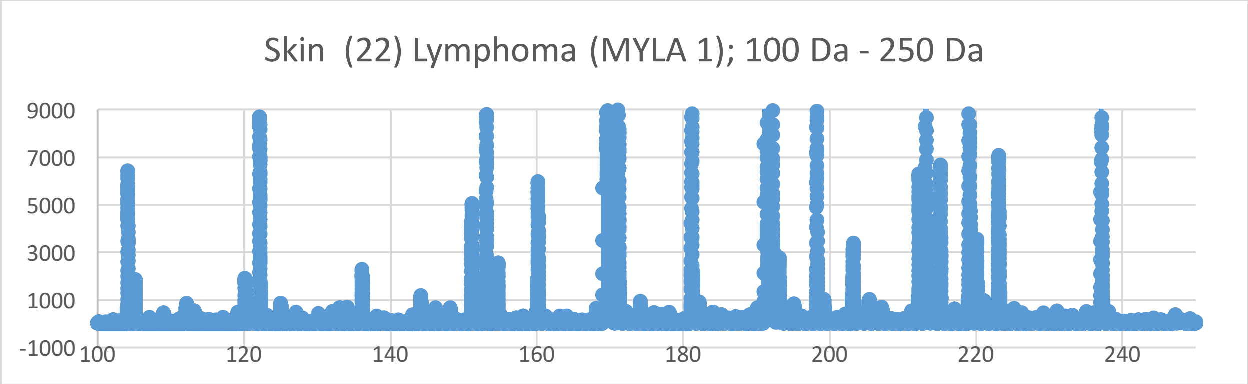
Skin (22) Lymphoma (MYLA 1); 100 Da - 250 Da
9000
7000
5000
3000
1000
-1000
100
120
140
160
180
200
220
240

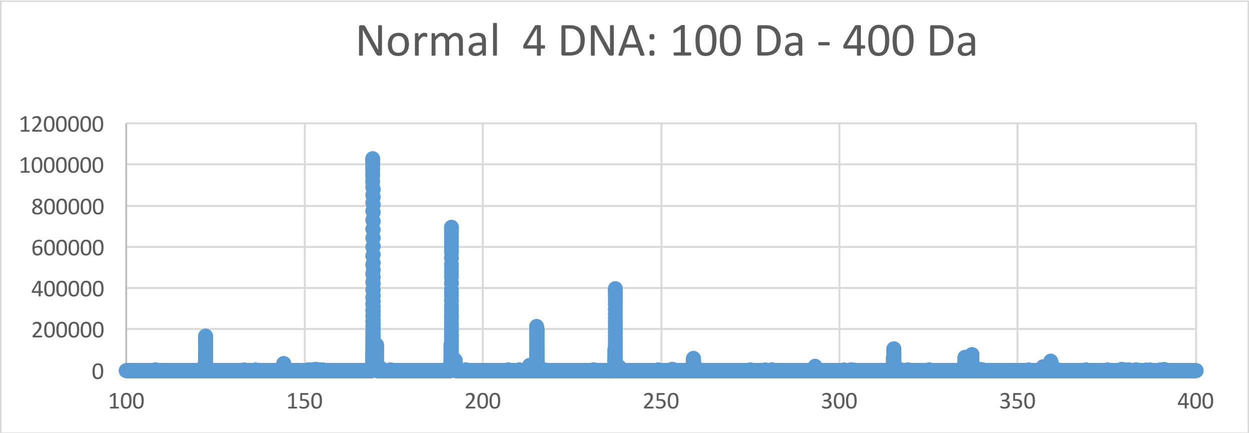
Normal 4 DNA: 100 Da - 400 Da
1200000
1000000
800000
600000
400000
200000
0
100
150
200
250
300
350
400

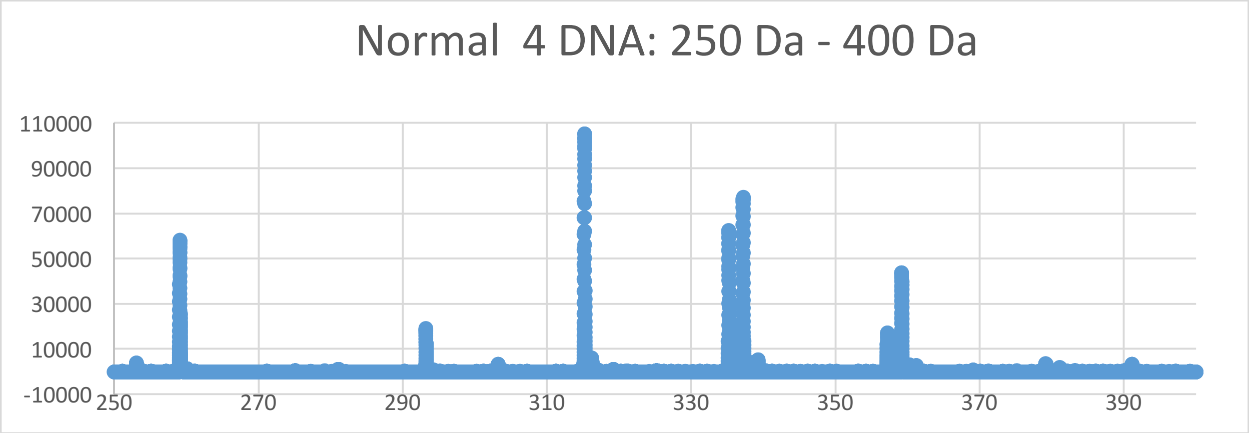
Normal 4 DNA: 250 Da - 400 Da
110000
90000
70000
50000
30000
10000
-10000
250
270
290
310
330
350
370
390

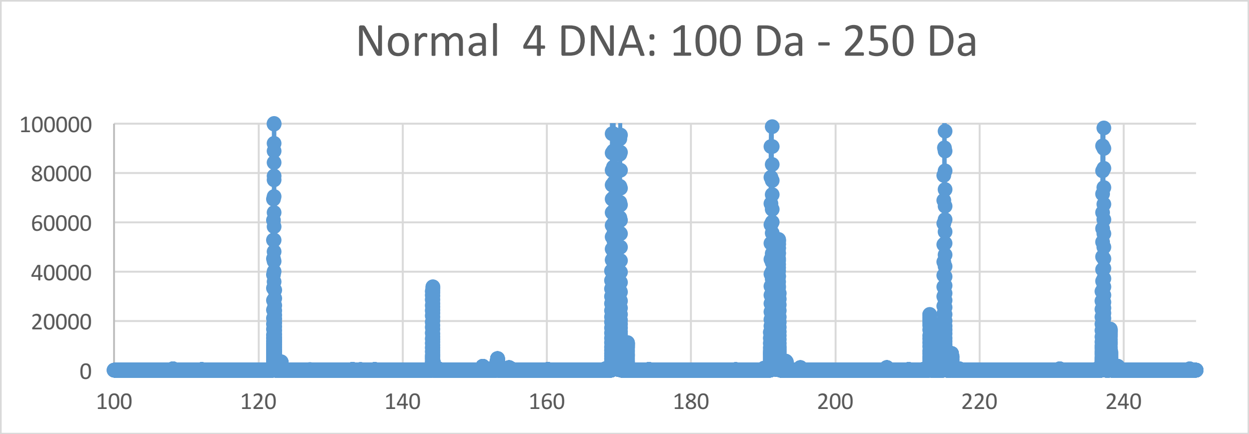
Normal 4 DNA: 100 Da - 250 Da
100000
80000
60000
40000
20000
0
100
120
140
160
180
200
220
240

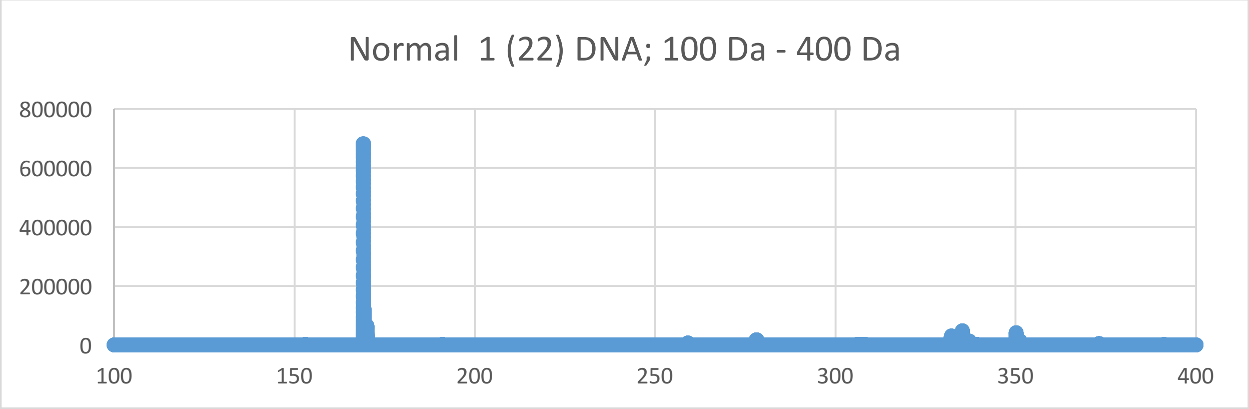
Normal 1 (22) DNA; 100 Da - 400 Da
800000
600000
400000
200000
0
100
150
200
250
300
350
400

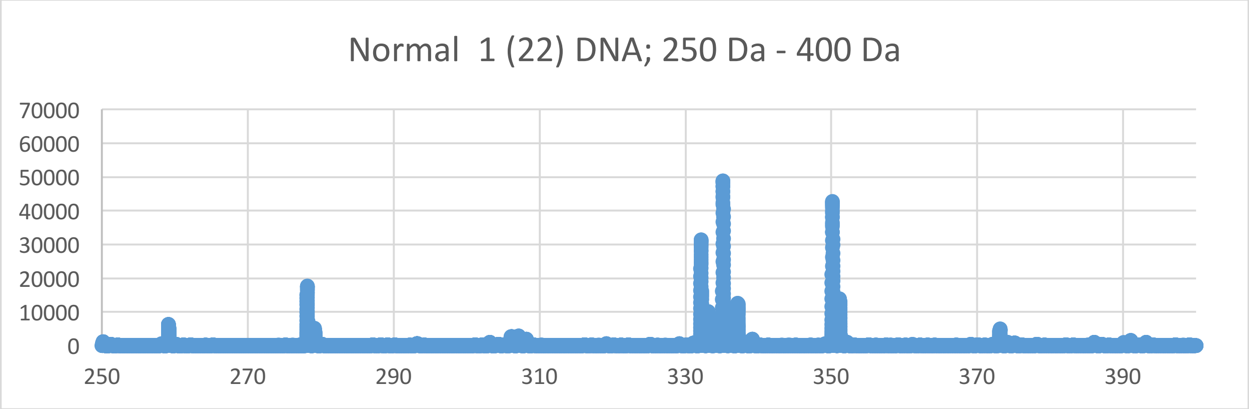
Normal 1 (22) DNA; 250 Da - 400 Da
70000
60000
50000
40000
30000
20000
10000
0
250
270
290
310
330
350
370
390

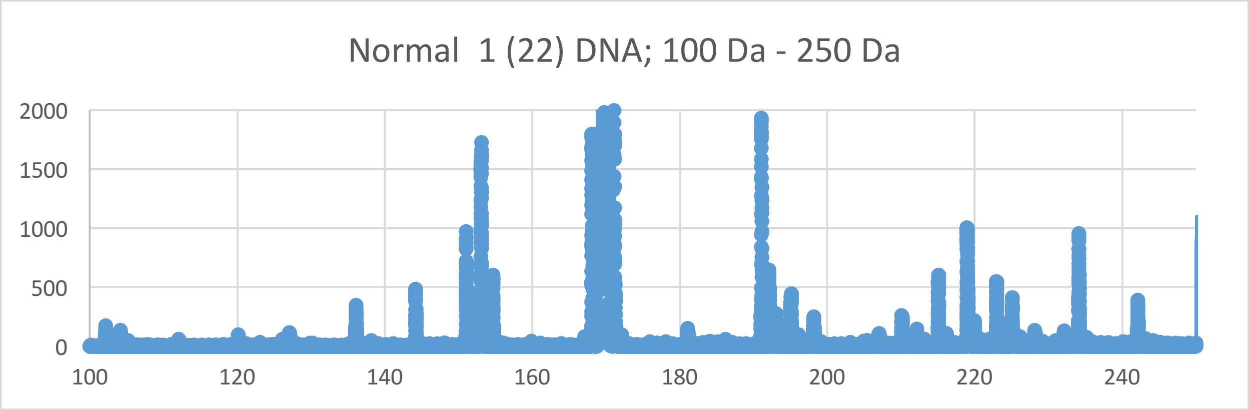
Normal 1 (22) DNA; 100 Da - 250 Da
2000
1500
1000
500
0
100
120
140
160
180
200
220
240

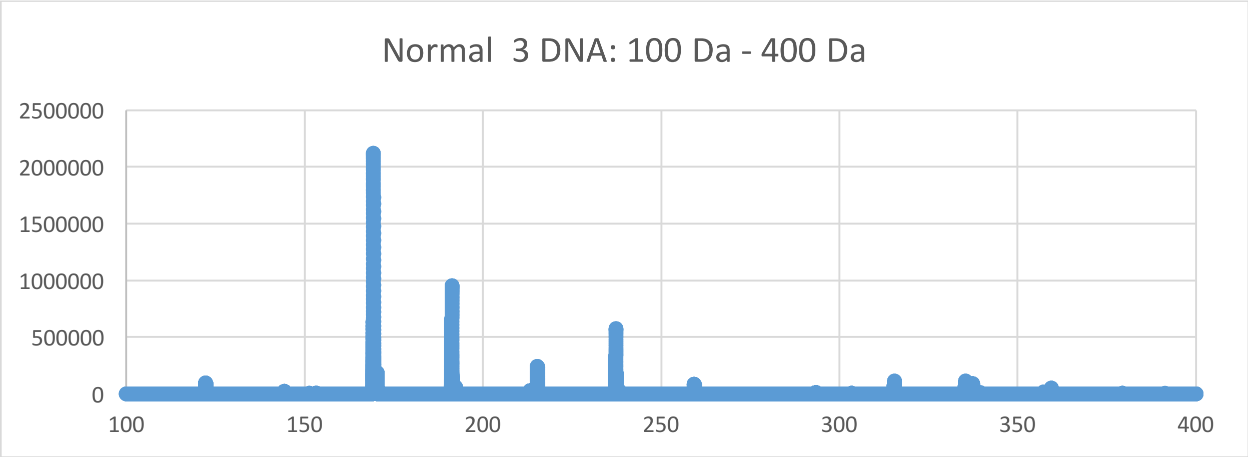
Normal 3 DNA: 100 Da - 400 Da
2500000
2000000
1500000
1000000
500000
0
100
150
200
250
300
350
400

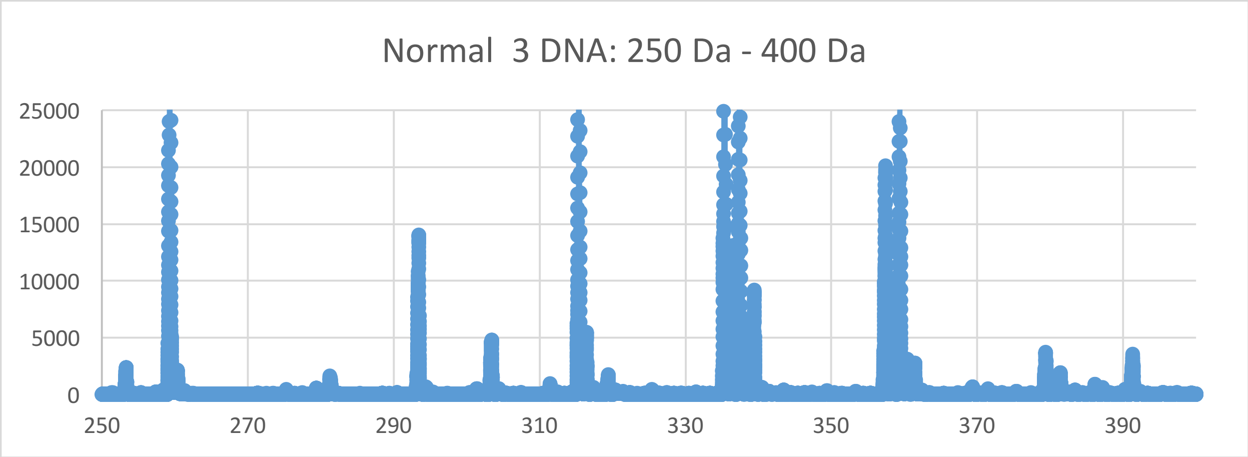
Normal 3 DNA: 250 Da - 400 Da
25000
20000
15000
10000
5000
0
250
270
290
310
330
350
370
390

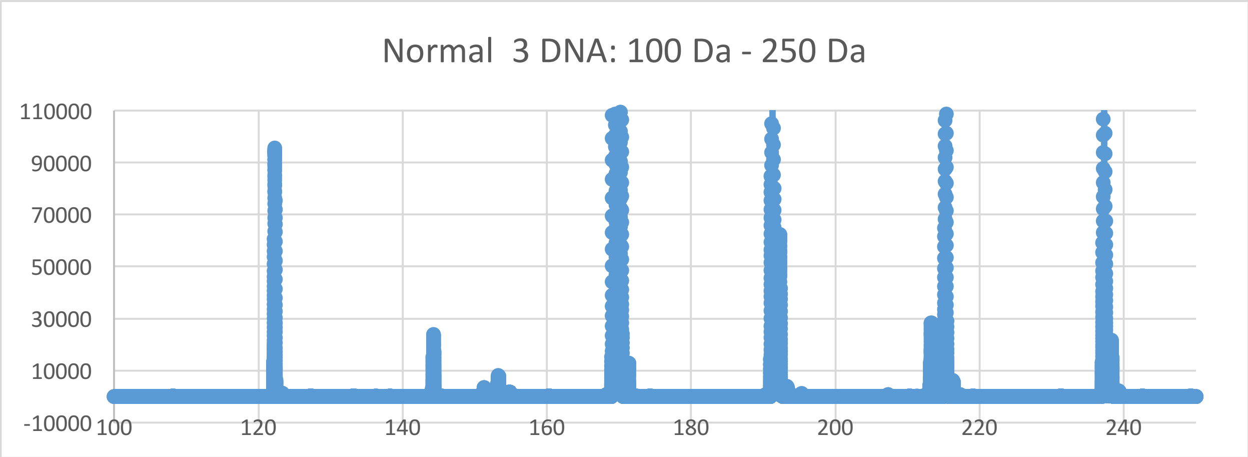
Normal 3 DNA: 100 Da - 250 Da
110000
90000
70000
50000
30000
10000
-10000
100
120
140
160
180
200
220
240

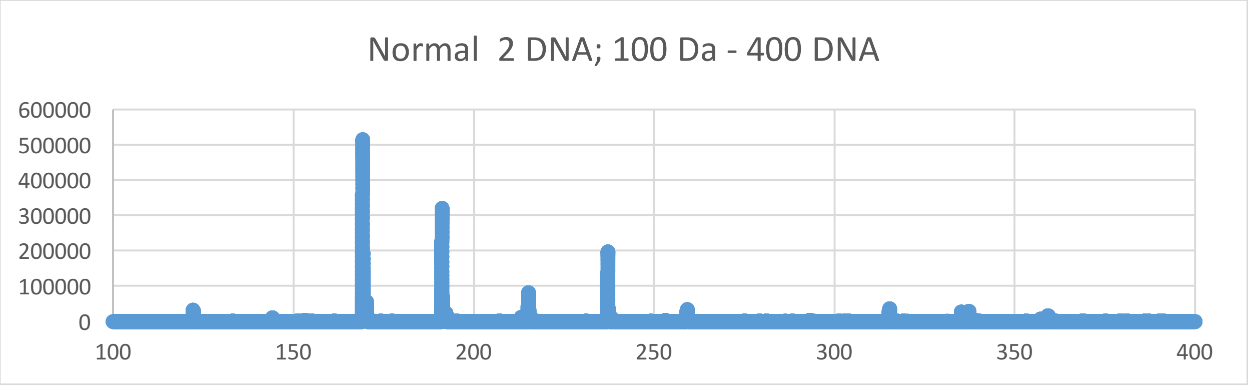
Normal 2 DNA; 100 Da - 400 DNA
600000
500000
400000
300000
200000
100000
0
100
150
200
250
300
350
400

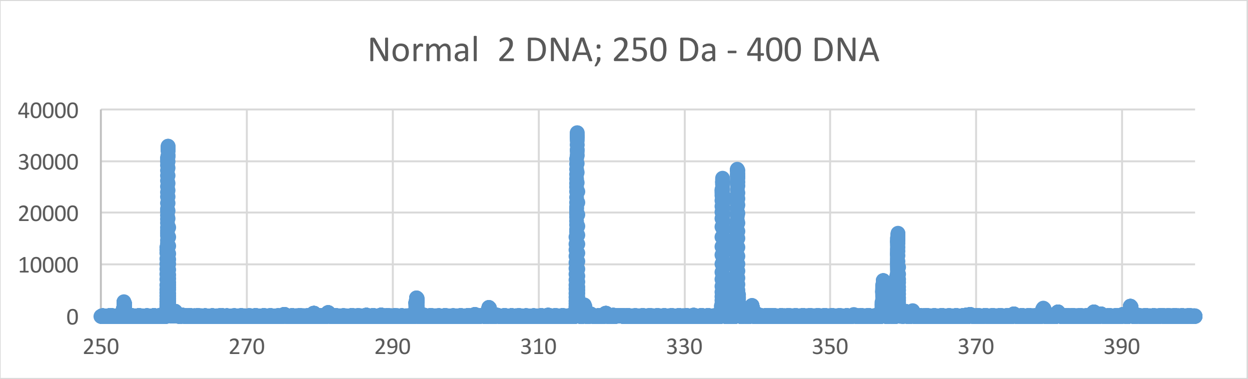
Normal 2 DNA; 250 Da - 400 DNA
40000
30000
20000
10000
0
250
270
290
310
330
350
370
390

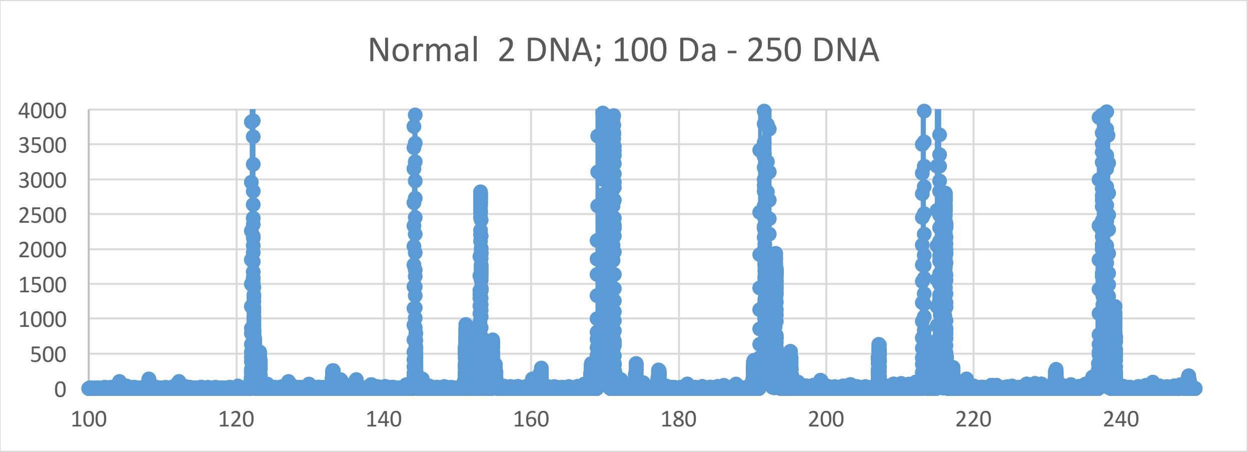
Normal 2 DNA; 100 Da - 250 DNA
4000
3500
3000
2500
2000
1500
1000
500
0
100
120
140
160
180
200
220
240
