## Supplementary Materials XIV for "Oncogenesis and Aging by Isotopic Functionalizations of the Proteins and Nucleic Acids"

XIV Supplementary Materials – Mass Spectra from 2000 Da - 2300 Da

Results for Supplementary 14

So for the pentanucleotides at $(phos)_{14}$, the A, T, G and C peaks for cancer decreases in peak intensities and decrease in numbers with the $(phos)_{14}$ to $(phos)_{13}$ dephosphorylations for telomeric oligonucleotides. The pentanucleotides dramatically change fragmentation patterns for cancer at $(phos)_{13}$. But as decrease the phosphate by 1 unit, the pentanucleotides change so that the mixed telomeric cancer and normal pentanucleotides more fragment and break with stabilities from the DNA with more numbers and stabilities for the A, T, C and G for $(phos)_{14}$ to $(phos)_{13}$ . The masses range from 2468 Da to 2190 Da. The cancer fragment more intense and stable pentanucleotides in this mass range. See Supplementary Materials XII for graphs. It seems that below the saturation range of some number of phosphates, the telomeres and mating telomeres behave in similar way within cancer and normal DNA. So this suggests cancer may be favored and distinguished from normal by changing saturated phosphorylations from $(phos)_{14}$ to $(phos)_{13}$. The A, T, C and G pentanucleotide peaks for cancer decrease with these dephosphorylations of the pentanucleotides. The dephosphorylations from $(phos)_{14}$ to $(phos)_{13}$ caused the resulting negligible changes in intensities and stabilitioes for cancer pentanucleotides for mating telomeric pentanucleotides and negligibly isotopically enrichments for mating telomeres. It is important to consider that there are less intense peaks for the pentanucleotides relative to the mononucleotides, dinucleotides, trinucleotides and tetranucleotides for the saturated phosphate units. For the pentanucleotides, the intensities of cancer negligibly increased and the intensities of normal fragments increased from $(phos)_{14}$ to $(phos)_{13.}$ The dephosphorylations of telomeric pentanucleotides from $(phos)_{14}$ to $(phos)_{13}$ slightly decreased isotopic enrichments of heavy isotopes of $^{15}N$, $^{13}C$ and $^{17}O$. The dephosphorylations from $(phos)_{14}$ to $(phos)_{13}$ also caused the mating telomeric pentanucleotides to be negligibly more intense in cancer DNA and more isotopically enriched. It seems the dephosphorylations are related to the isotopic enrichments and depletions; but the telomeres and mating telomeres are negligibly more intense in cancer and slightly more isotopically enriched. The telomeric and mating telomeric pentanucleotides of cancer tended to also be negligibly isotopically enriched so it seems the cancer stabilities of the telomeric and mating telomeric pentanucleotides are related to their negligibly isotopic enrichments from the dephosphorylations from $(phos)_{14}$ to $(phos)_{13}$ and the mutations purely driven by dephosphorylations and enrichments. Isotopic enrichments weaken the mating telomeric normal pentanucleotides while negligibly strengthening cancerous telomeric pentanucleotides. The negligibly intense cancer telomeric pentanucleotides were also slightly more isotopically enriched. The negligible intense cancer mating telomeric pentanucleotides were slightly more isotopically enriched by light isotopes. The dephosphorylations from $(phos)_{14}$ to $(phos)_{13}$ negligibly increased the isotopic enrichments of the cancer for the telomeric and mating telomeric pentanucleotides. Such seems compatible for replications and transcriptions as stable telomeres seem compatible to unstable mating telomeres for the unravelings and constructions of DNA mating telomeres onto the building templates of the telomeres. But in this case of all normal telomeric and mating telomeric pentanucleotides are negligibly enriched and unstable then the replications, transcriptions and translations cannot occur normally. The dephosphorylations from $(phos)_{14}$ to $(phos)_{13}$ caused demethylations to cause more cancer intensities and stabilities but greater isotopic enrichments for G containing pentanucleotides. But the demethylations of the T, C and/or A caused negligible alterations of cancer stabilities and slight increased isotopic enrichments for mating telomeres. Where isotopic changes occurred the functionalization effects were not negligible. The methylations of the C pentanucleotides negligibly decreased cancer intensities and stabilities with isotopic enrichments. It seems from these data that during biological processes the dephosphorylations and stabilizations of normal or vice versa dephosphorylations and stabilizations of cancer are or can be countered by methylations (and isotopic enrichments) and destabilizations of the normal for cancer or vice versa dephosphorylations and destabilizations of the cancer by methylations as this research discovers coupled phosphorylations and methylations (and isotopic enrichments) or coupled dephosphorylations and methylations (and isotopic enrichments) with involved greater enrichments for normal biological functions and explaining cancer! The dephosphorylations from $(phos)_{14}$ to $(phos)_{13}$ caused acetylations to cause more cancer stabilities for A and C with negligibly isotopic depletions and slight cancer stabilities for G and T with negligibly depletions of heavy isotopes for telomeric pentanucleotides. The acetylations from $(phos)_{14}$ to $(phos)_{13}$ caused less cancer stabilities and intensities for A and C with negligibly more isotopic depletions for the mating telomeric pentanucleotides. This contrast in A, T, C and G from the de-acetylations and dephosphorylations may reflect the developments of cancer as the pentanucleotides fragment but the other pentanucleotides more hold together for damaged DNA for cancer.

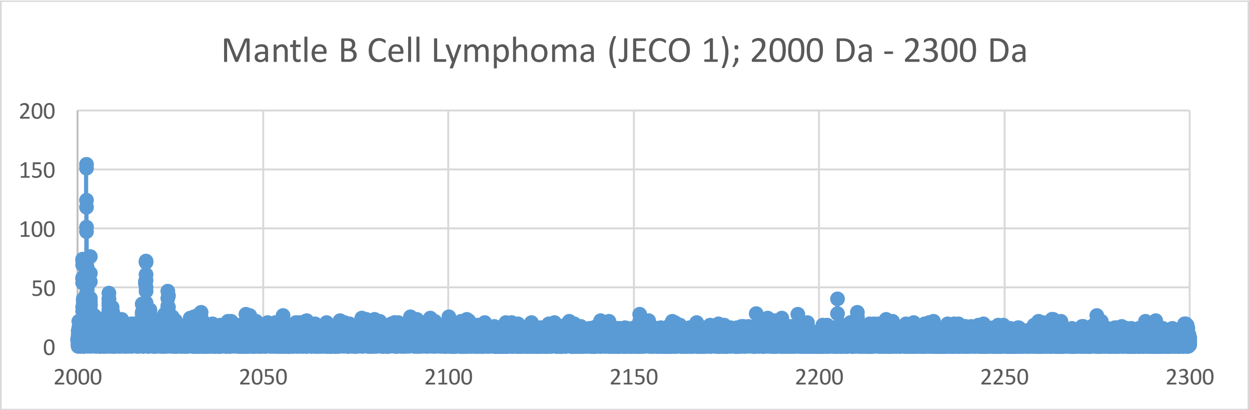
Mantle B Cell Lymphoma (JECO 1); 2000 Da - 2300 Da
200
150
100
50
0
2000
2050
2100
2150
2200
2250
2300

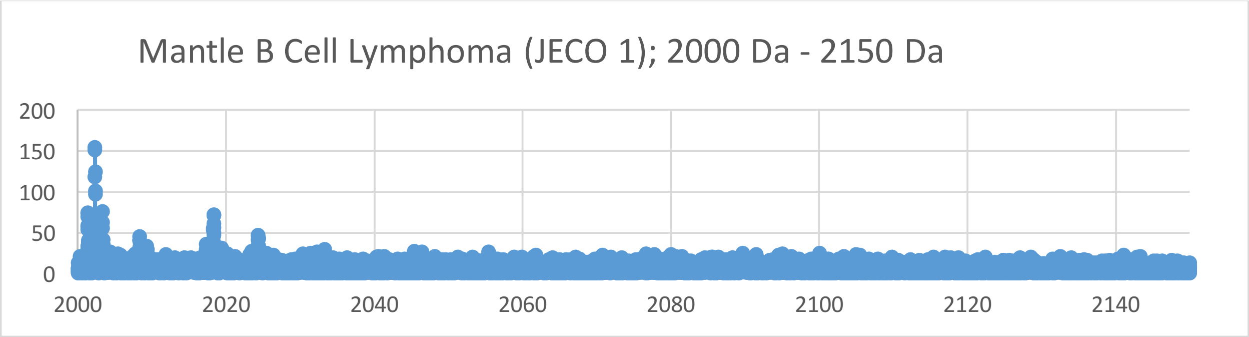
Mantle B Cell Lymphoma (JECO 1); 2000 Da - 2150 Da
200
150
100
50
0
2000
2020
2040
2060
2080
2100
2120
2140

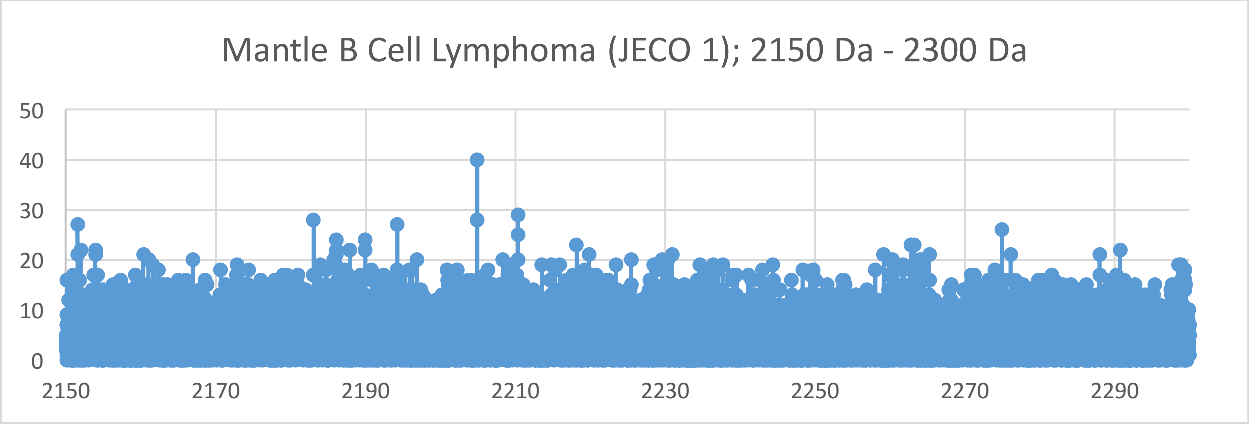
Mantle B Cell Lymphoma (JECO 1); 2150 Da - 2300 Da
50
40
30
20
10
0
2150
2170
2190
2210
2230
2250
2270
2290

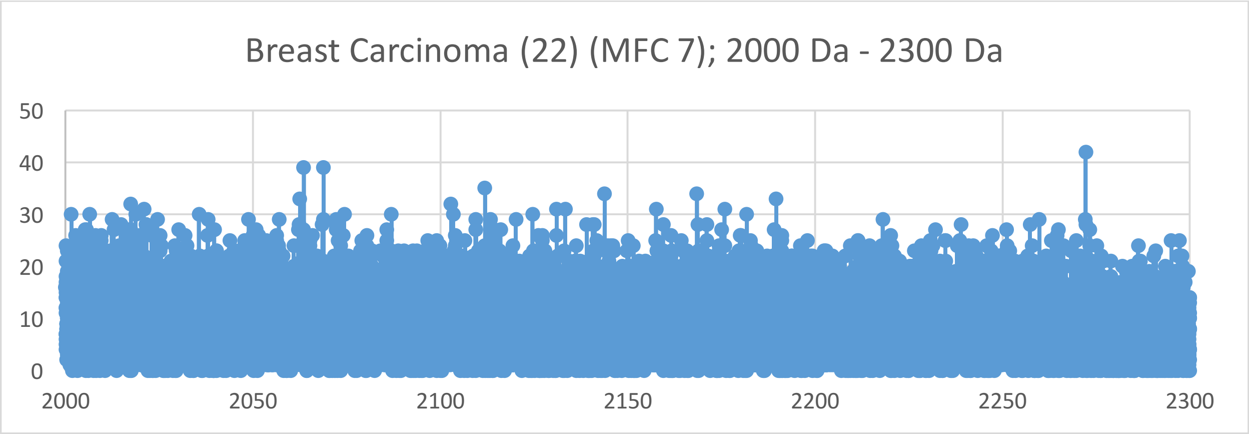
Breast Carcinoma (22) (MFC 7); 2000 Da - 2300 Da
50
40
30
20
10
0
2000
2050
2100
2150
2200
2250
2300

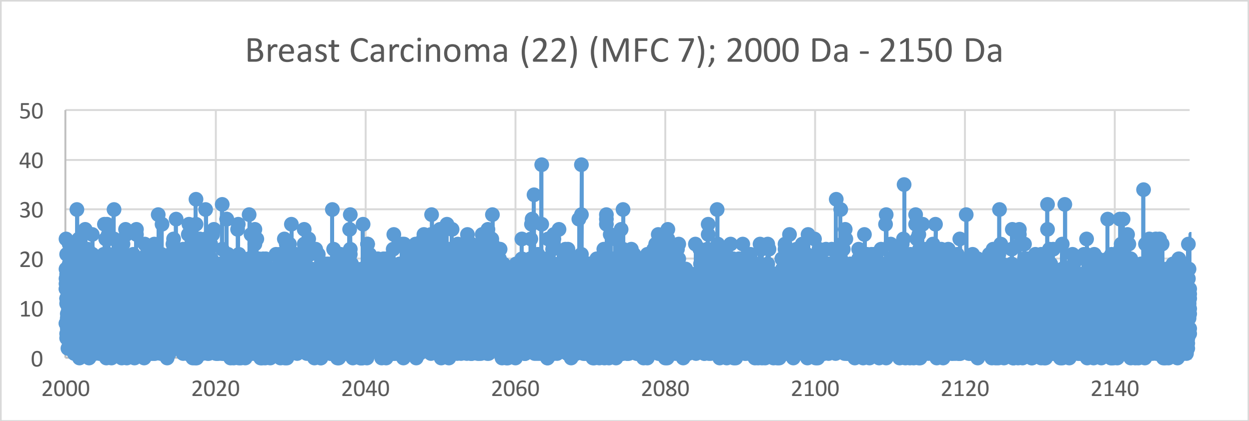
Breast Carcinoma (22) (MFC 7); 2000 Da - 2150 Da
50
40
30
20
10
0
2000
2020
2040
2060
2080
2100
2120
2140

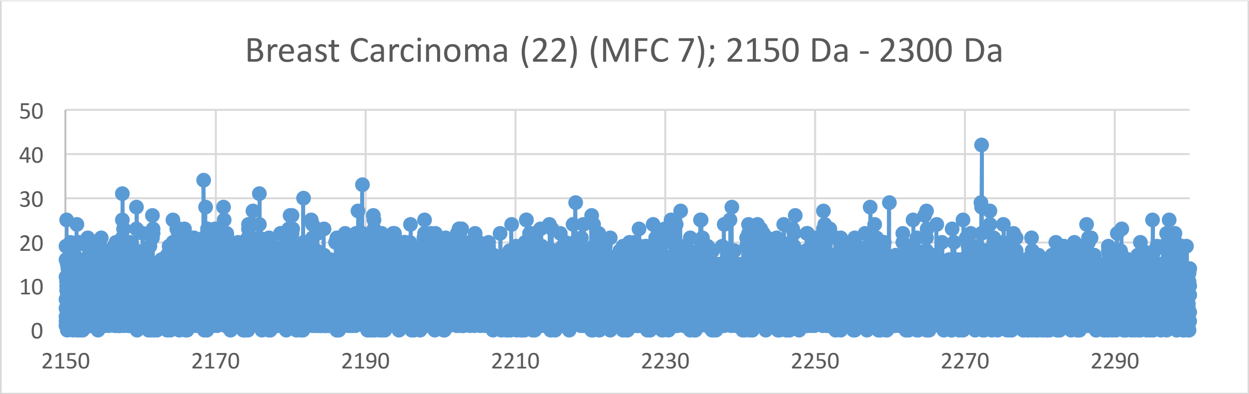
Breast Carcinoma (22) (MFC 7); 2150 Da - 2300 Da
50
40
30
20
10
0
2150
2170
2190
2210
2230
2250
2270
2290

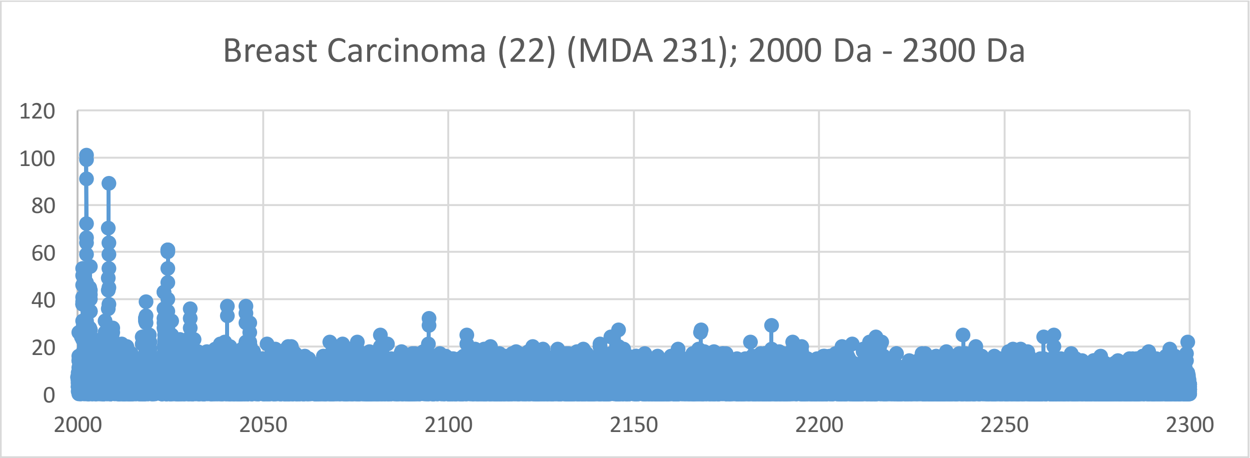

Breast Carcinoma (22) (MDA 231); 2000 Da - 2300 Da
120
100
80
60
40
20
0
2000
2050
2100
2150
2200
2250
2300


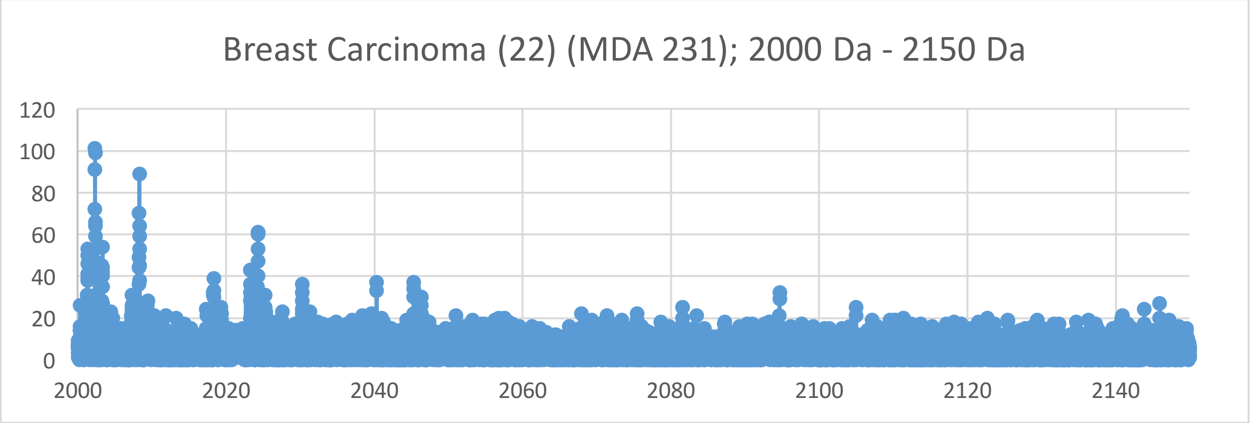

Breast Carcinoma (22) (MDA 231); 2000 Da - 2150 Da
120
100
80
60
40
20
0
2000
2020
2040
2060
2080
2100
2120
2140


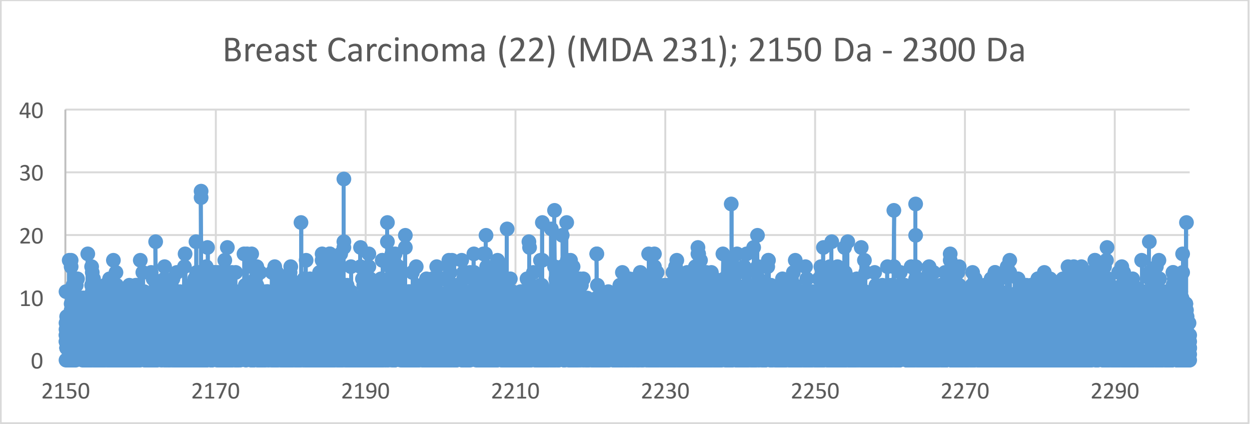

Breast Carcinoma (22) (MDA 231); 2150 Da - 2300 Da
40
30
20
10
0
2150
2170
2190
2210
2230
2250
2270
2290

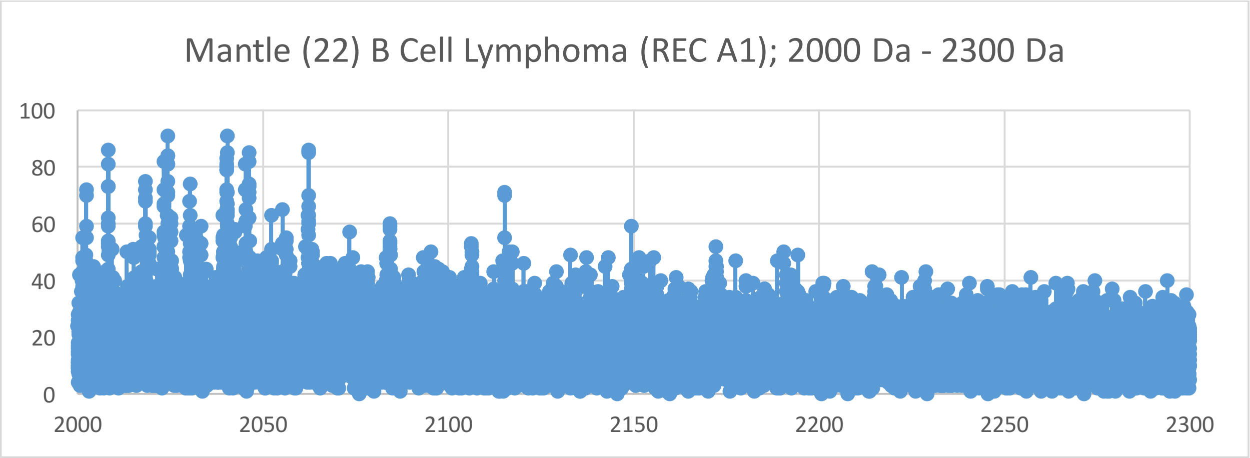

Mantle (22) B Cell Lymphoma (REC A1); 2000 Da - 2300 Da
100
80
60
40
20
0
2000
2050
2100
2150
2200
2250
2300


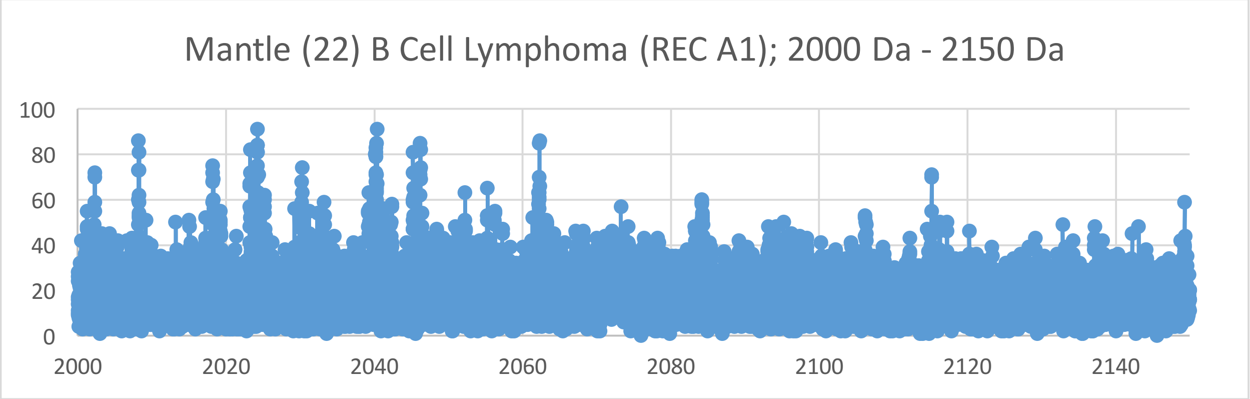

Mantle (22) B Cell Lymphoma (REC A1); 2000 Da - 2150 Da
100
80
60
40
20
0
2000
2020
2040
2060
2080
2100
2120
2140


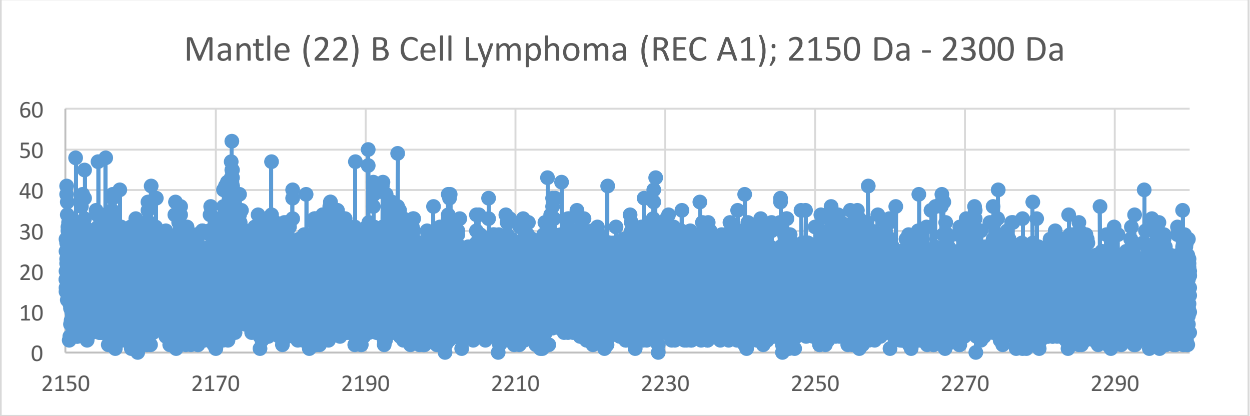

Mantle (22) B Cell Lymphoma (REC A1); 2150 Da - 2300 Da
60
50
40
30
20
10
0
2150
2170
2190
2210
2230
2250
2270
2290

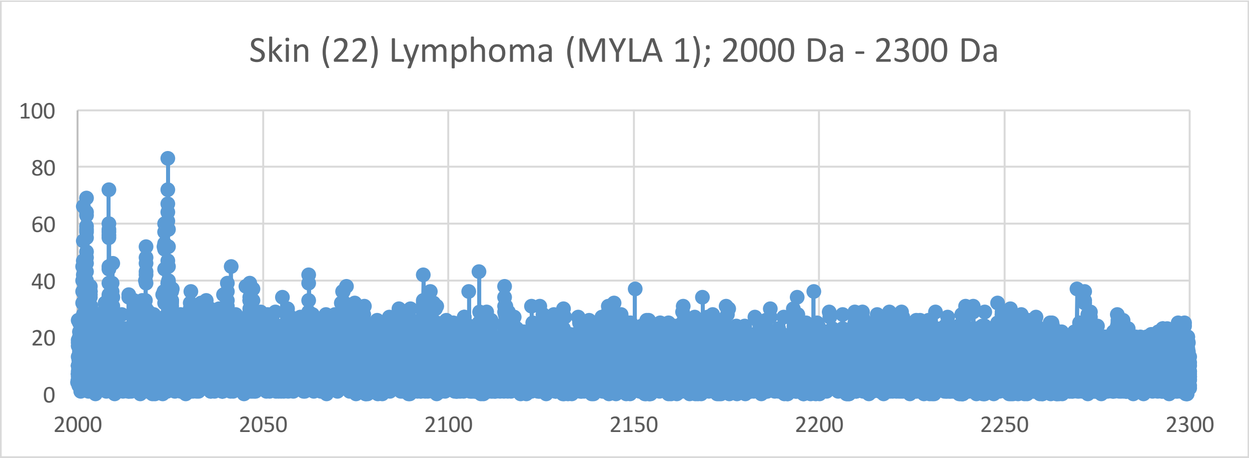
Skin (22) Lymphoma (MYLA 1); 2000 Da - 2300 Da
100
80
60
40
20
0
2000
2050
2100
2150
2200
2250
2300

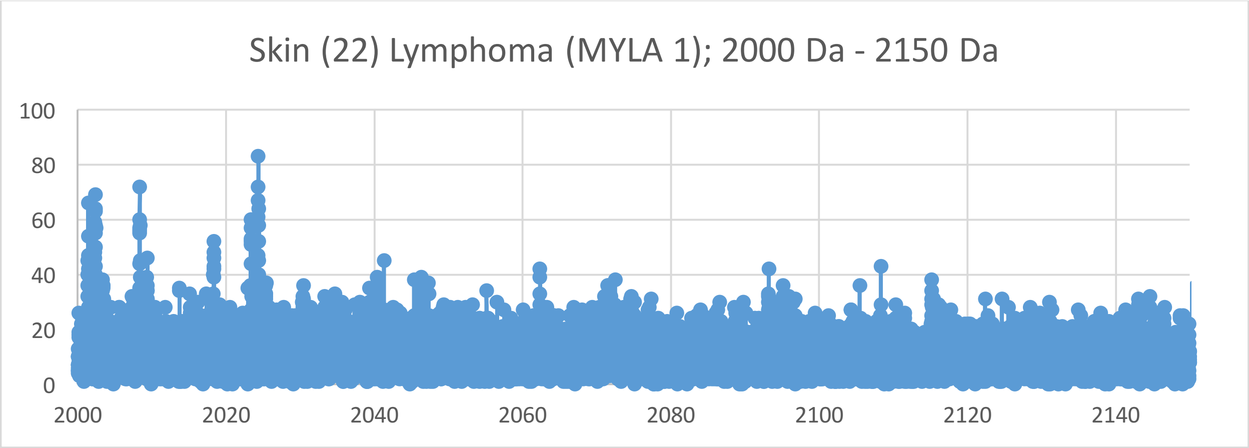
Skin (22) Lymphoma (MYLA 1); 2000 Da - 2150 Da
100
80
60
40
20
0
2000
2020
2040
2060
2080
2100
2120
2140

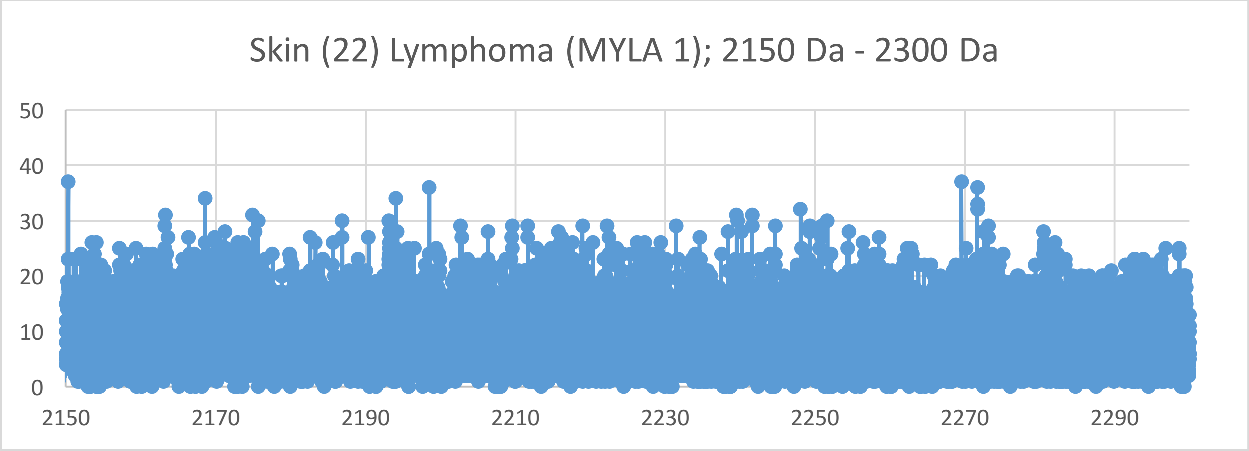
Skin (22) Lymphoma (MYLA 1); 2150 Da - 2300 Da
50
40
30
20
10
0
2150
2170
2190
2210
2230
2250
2270
2290

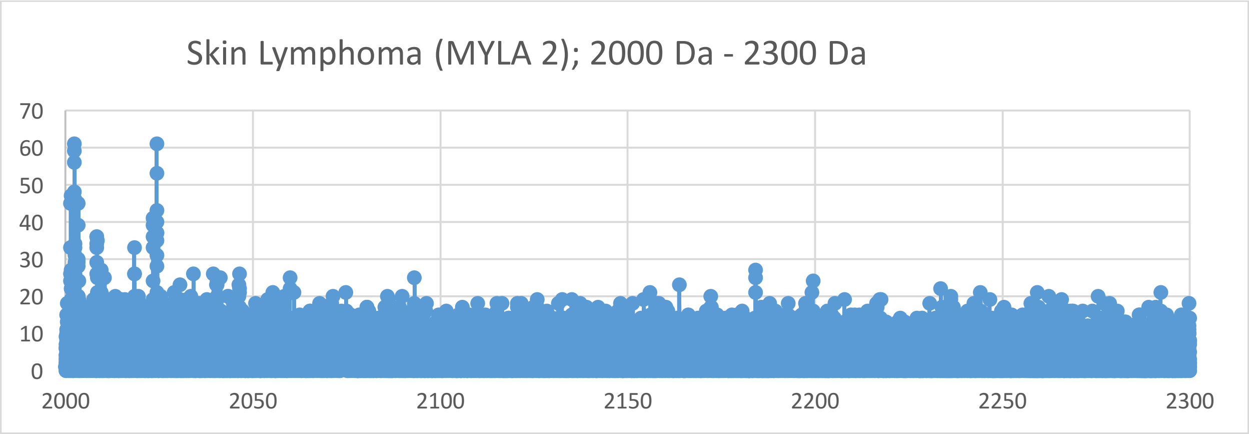
Skin Lymphoma (MYLA 2); 2000 Da - 2300 Da
70
60
50
40
30
20
10
0
2000
2050
2100
2150
2200
2250
2300

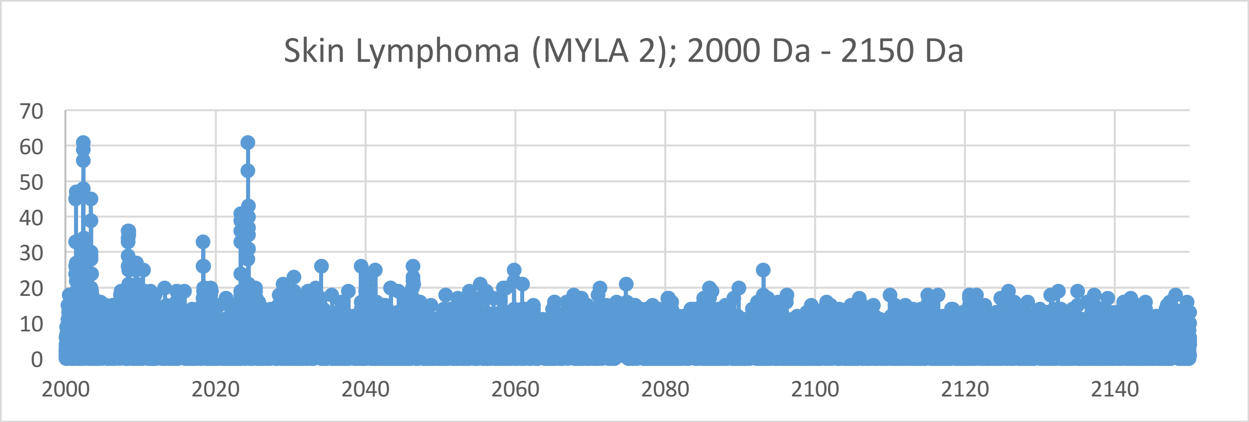
Skin Lymphoma (MYLA 2); 2000 Da - 2150 Da
70
60
50
40
30
20
10
0
2000
2020
2040
2060
2080
2100
2120
2140

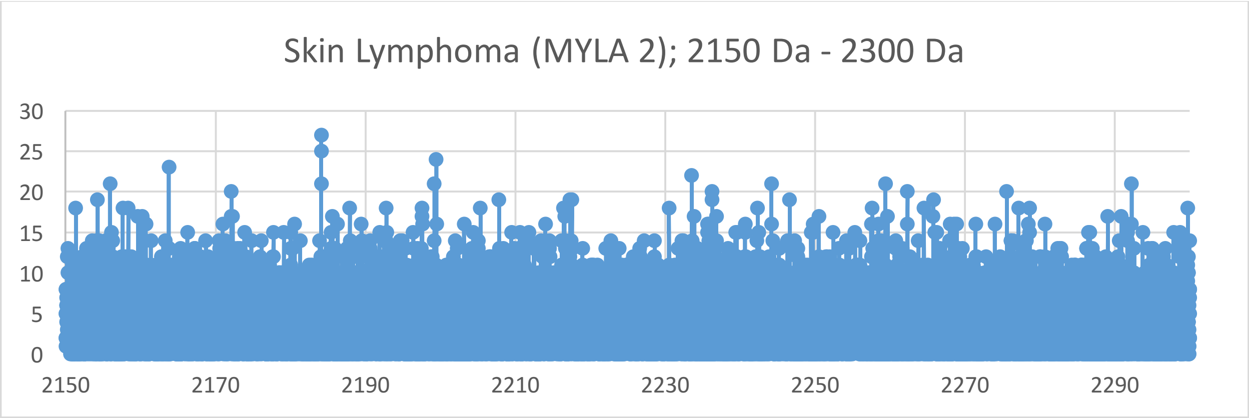
Skin Lymphoma (MYLA 2); 2150 Da - 2300 Da
30
25
20
15
10
5
0
2150
2170
2190
2210
2230
2250
2270
2290

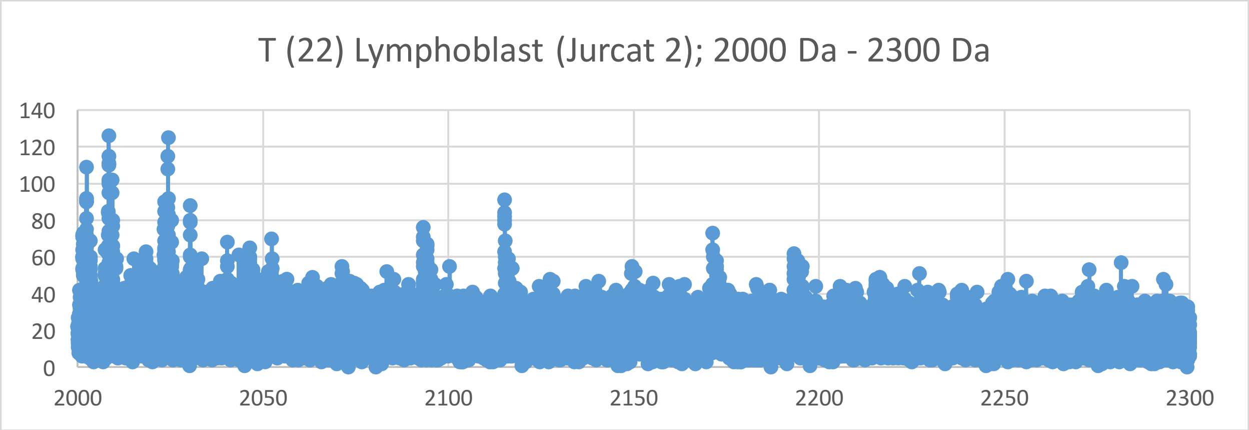
T (22) Lymphoblast (Jurcat 2); 2000 Da - 2300 Da
140
120
100
80
60
40
20
0
2000
2050
2100
2150
2200
2250
2300

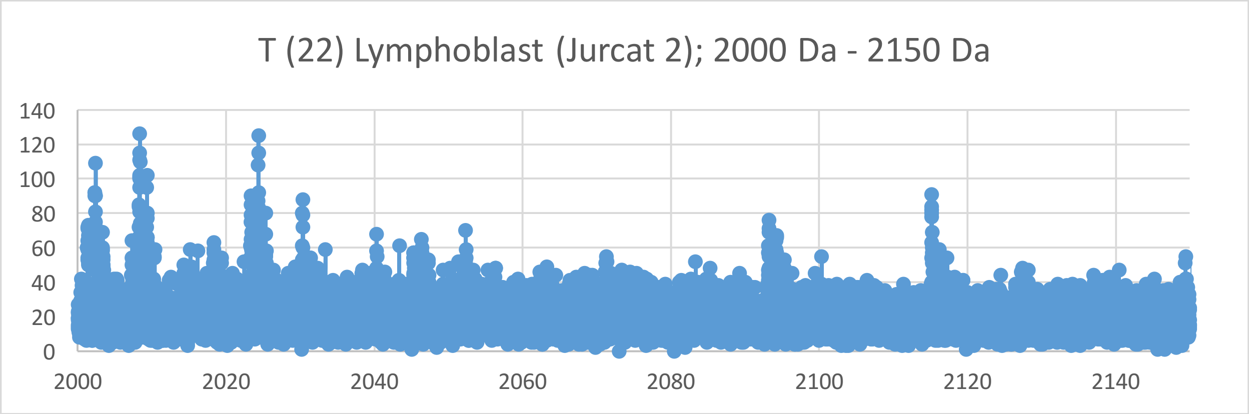
T (22) Lymphoblast (Jurcat 2); 2000 Da - 2150 Da
140
120
100
80
60
40
20
0
2000
2020
2040
2060
2080
2100
2120
2140

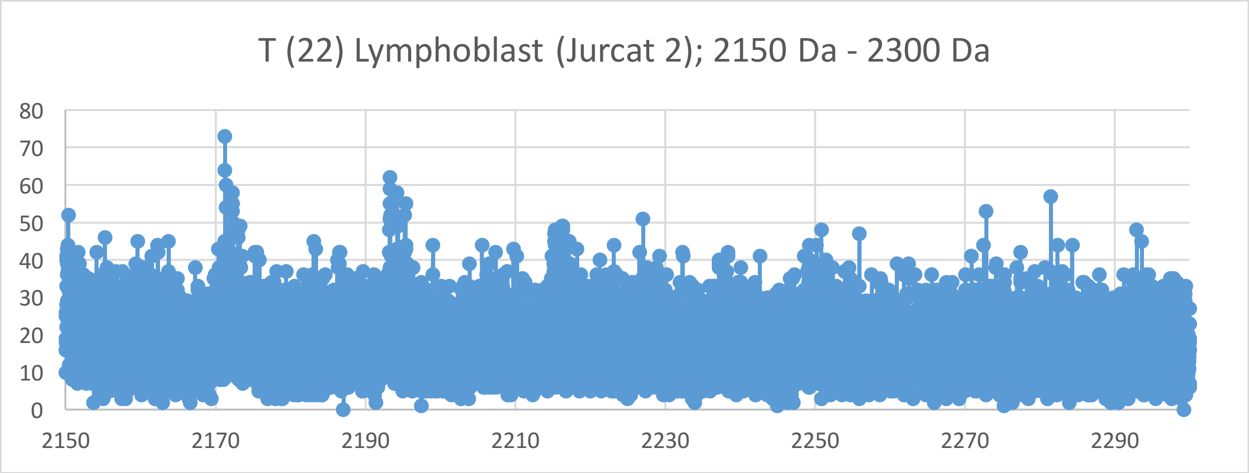
T (22) Lymphoblast (Jurcat 2); 2150 Da - 2300 Da
80
70
60
50
40
30
20
10
0
2150
2170
2190
2210
2230
2250
2270
2290

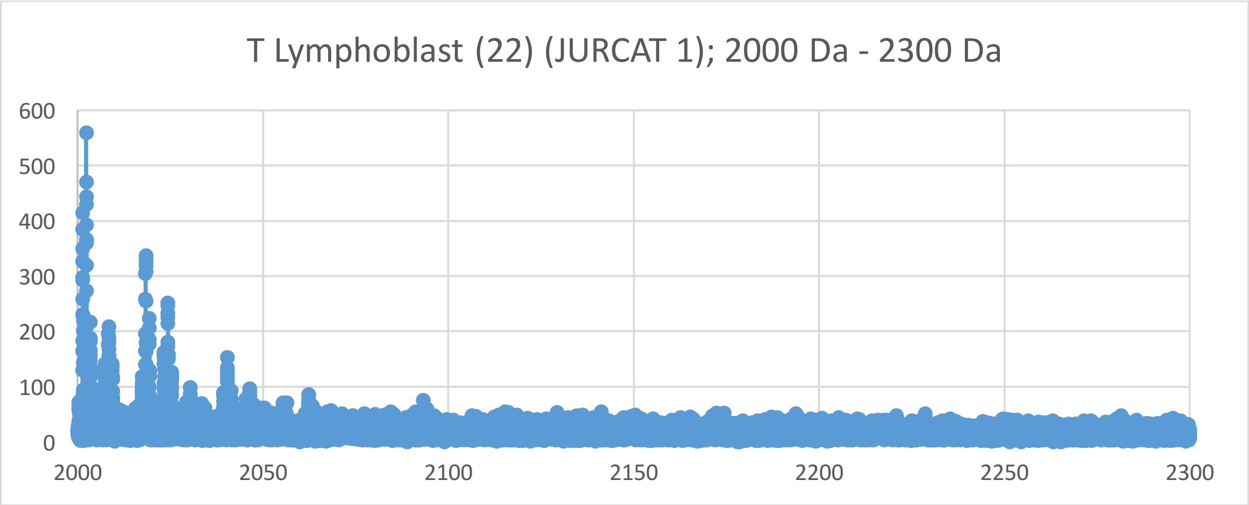
T Lymphoblast (22) (JURCAT 1); 2000 Da - 2300 Da
600
500
400
300
200
100
0
2000
2050
2100
2150
2200
2250
2300

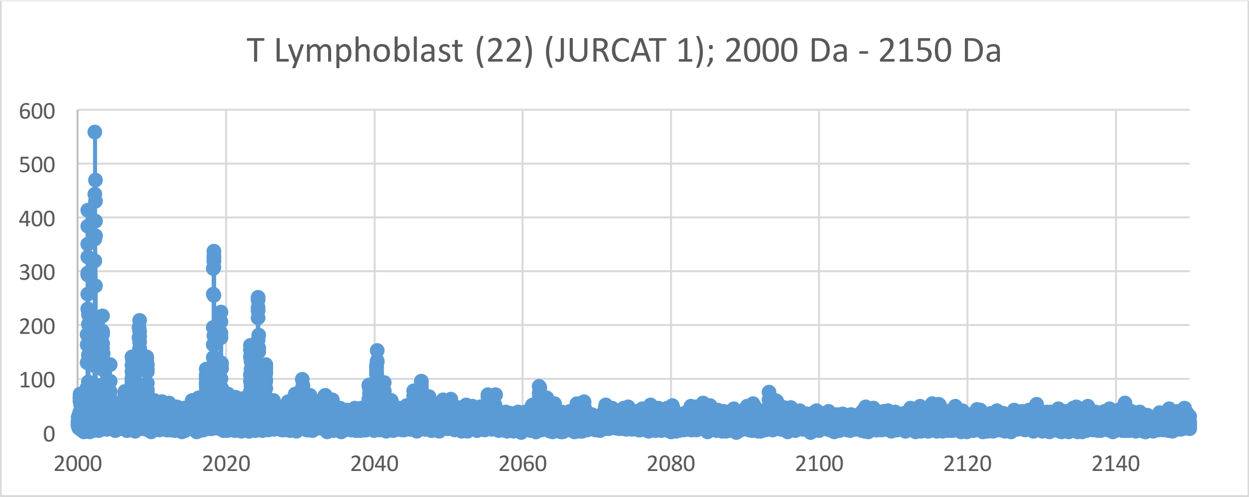
T Lymphoblast (22) (JURCAT 1); 2000 Da - 2150 Da
600
500
400
300
200
100
0
2000
2020
2040
2060
2080
2100
2120
2140

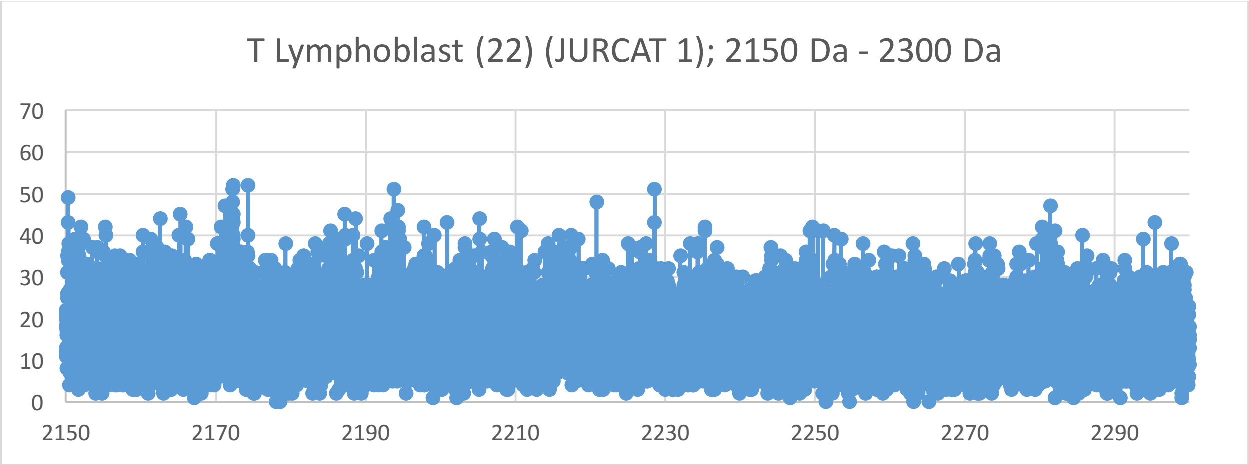
T Lymphoblast (22) (JURCAT 1); 2150 Da - 2300 Da
70
60
50
40
30
20
10
0
2150
2170
2190
2210
2230
2250
2270
2290

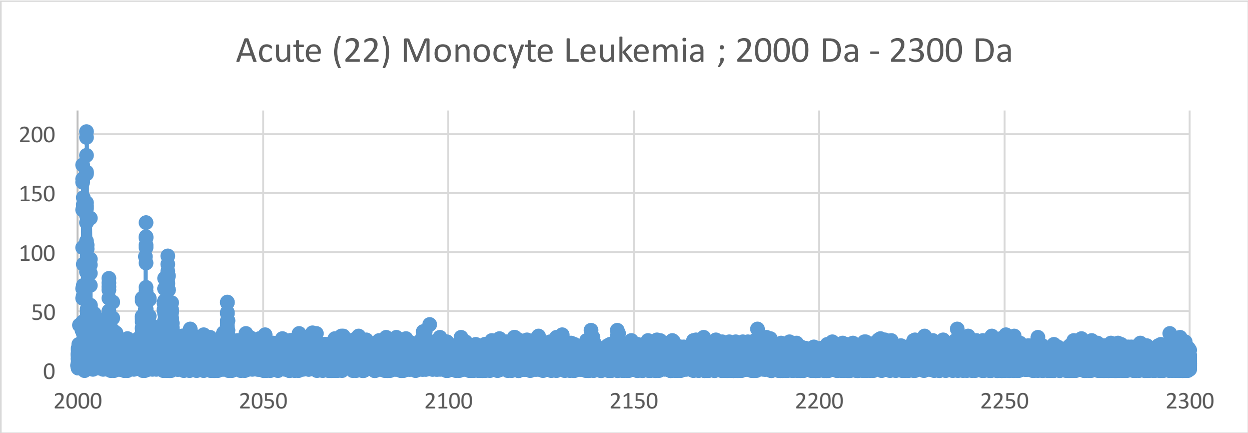
Acute (22) Monocyte Leukemia ; 2000 Da - 2300 Da
200
150
100
50
0
2000
2050
2100
2150
2200
2250
2300

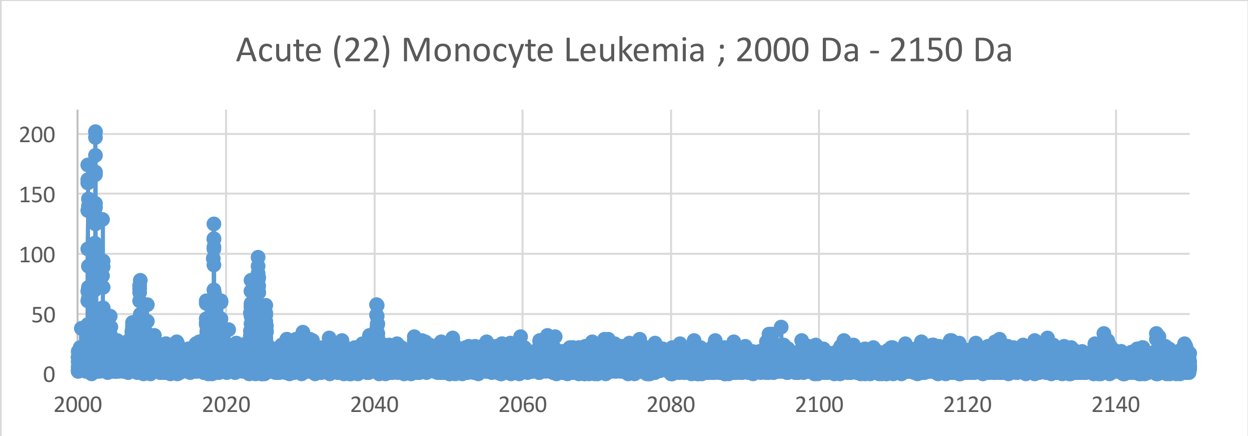
Acute (22) Monocyte Leukemia ; 2000 Da - 2150 Da
200
150
100
50
0
2000
2020
2040
2060
2080
2100
2120
2140

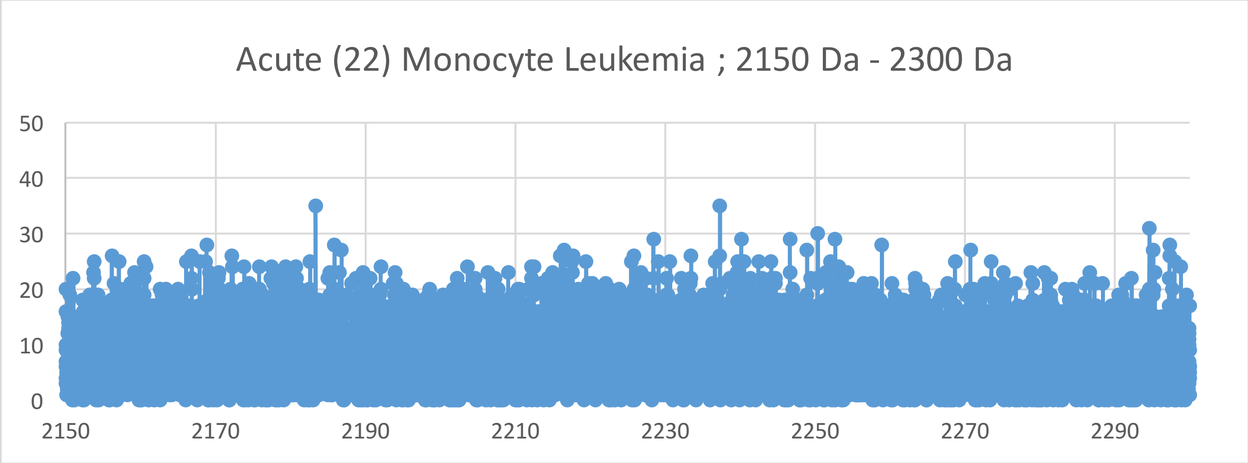
Acute (22) Monocyte Leukemia ; 2150 Da - 2300 Da
50
40
30
20
10
0
2150
2170
2190
2210
2230
2250
2270
2290

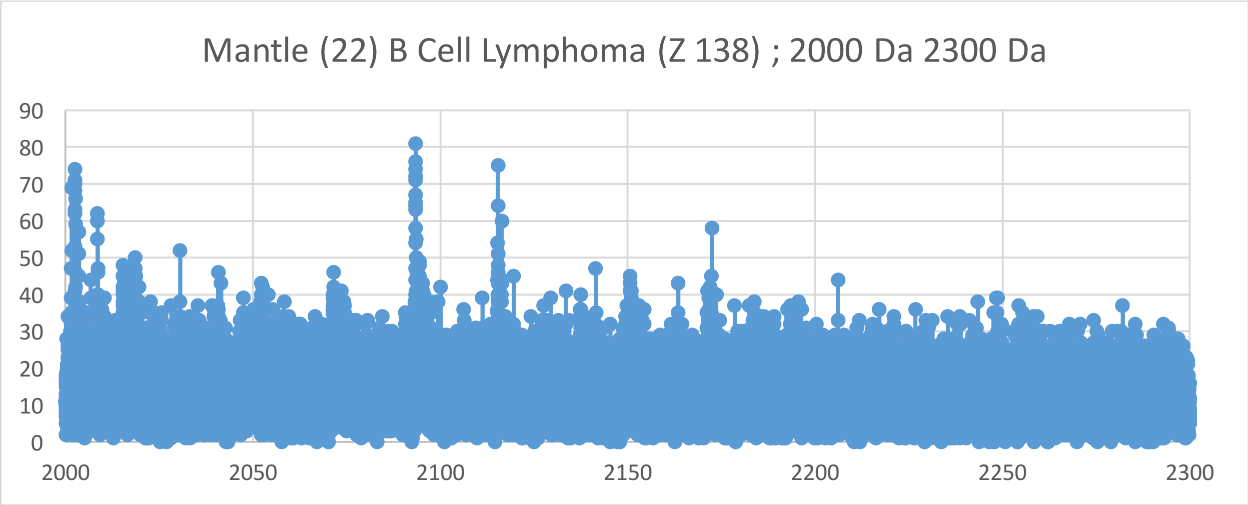

Mantle (22) B Cell Lymphoma (Z 138) ; 2000 Da 2300 Da
90
80
70
60
50
40
30
20
10
0
2000
2050
2100
2150
2200
2250
2300


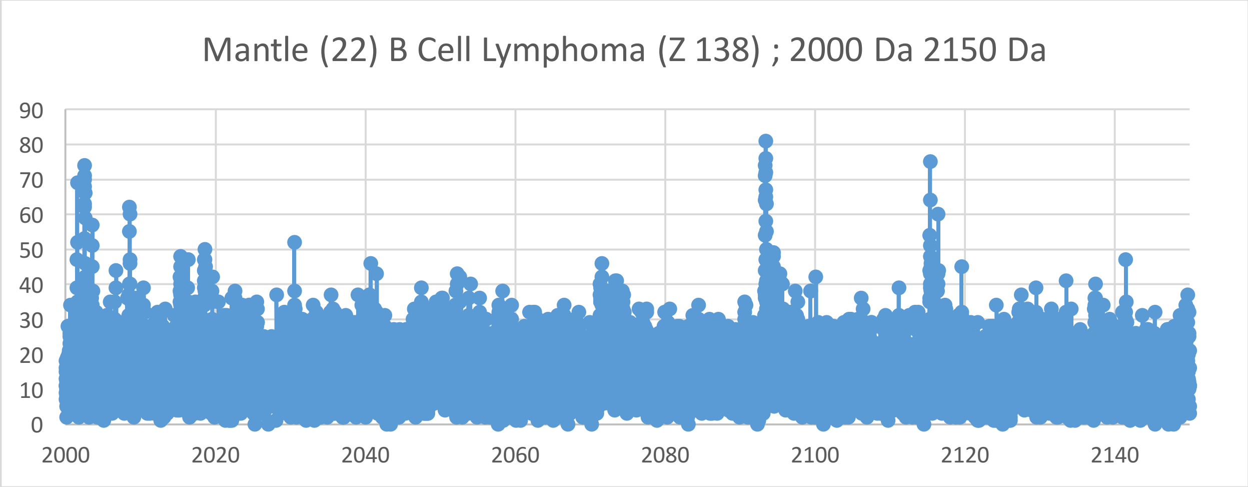

Mantle (22) B Cell Lymphoma (Z 138) ; 2000 Da 2150 Da
90
80
70
60
50
40
30
20
10
0
2000
2020
2040
2060
2080
2100
2120
2140


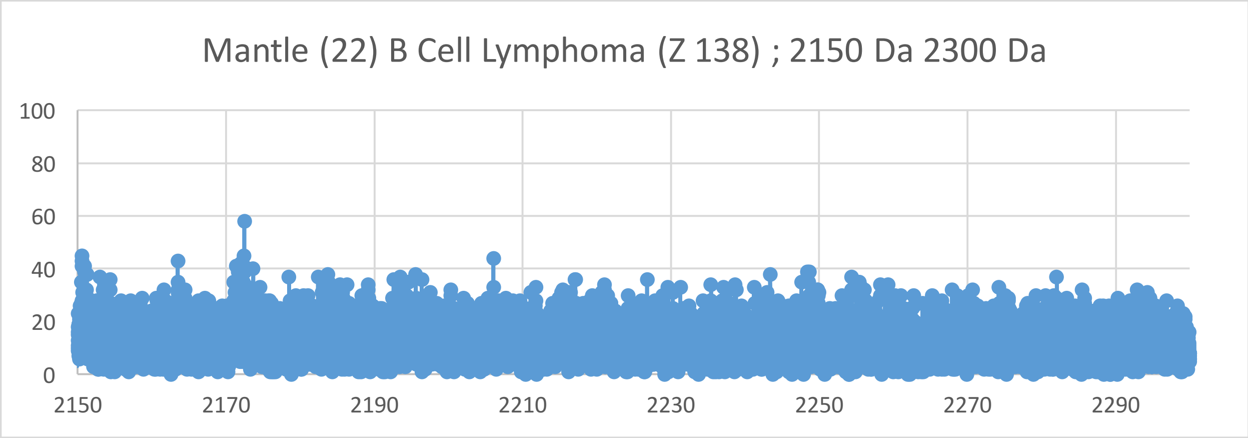

Mantle (22) B Cell Lymphoma (Z 138) ; 2150 Da 2300 Da
100
80
60
40
20
0
2150
2170
2190
2210
2230
2250
2270
2290

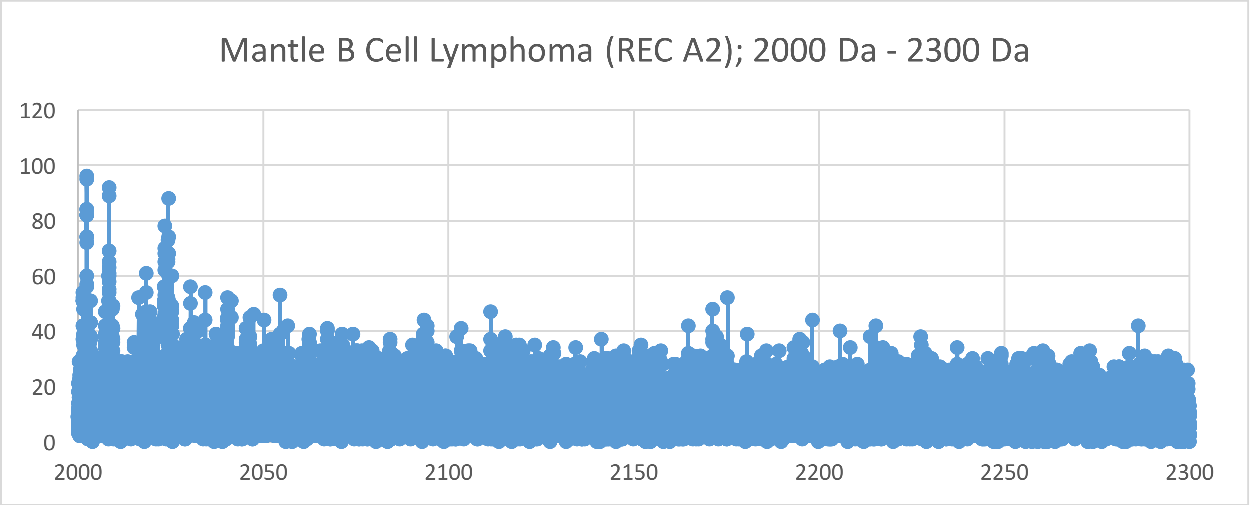
Mantle B Cell Lymphoma (REC A2); 2000 Da - 2300 Da
120
100
80
60
40
20
0
2000
2050
2100
2150
2200
2250
2300

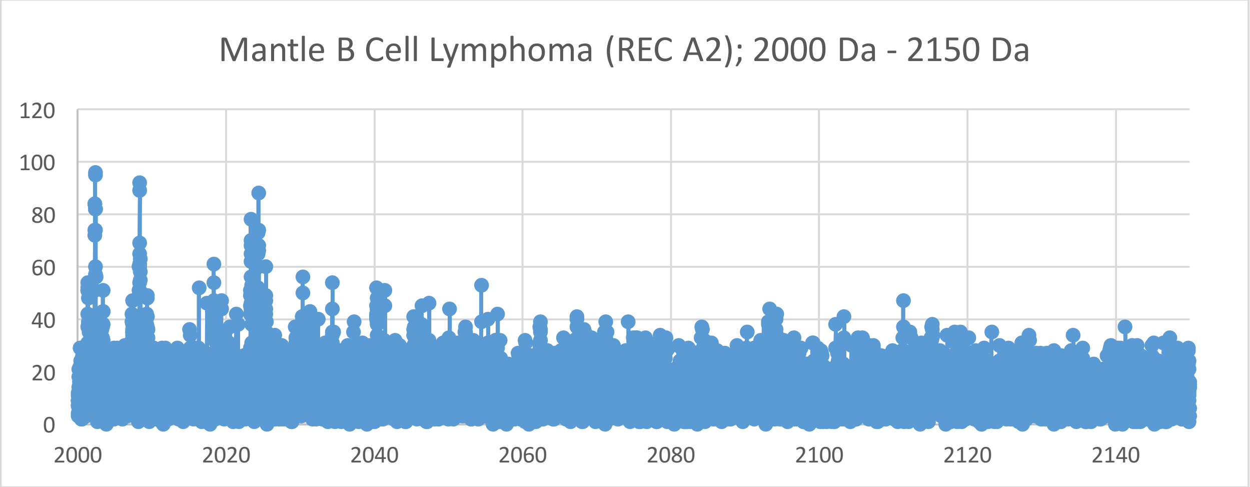
Mantle B Cell Lymphoma (REC A2); 2000 Da - 2150 Da
120
100
80
60
40
20
0
2000
2020
2040
2060
2080
2100
2120
2140

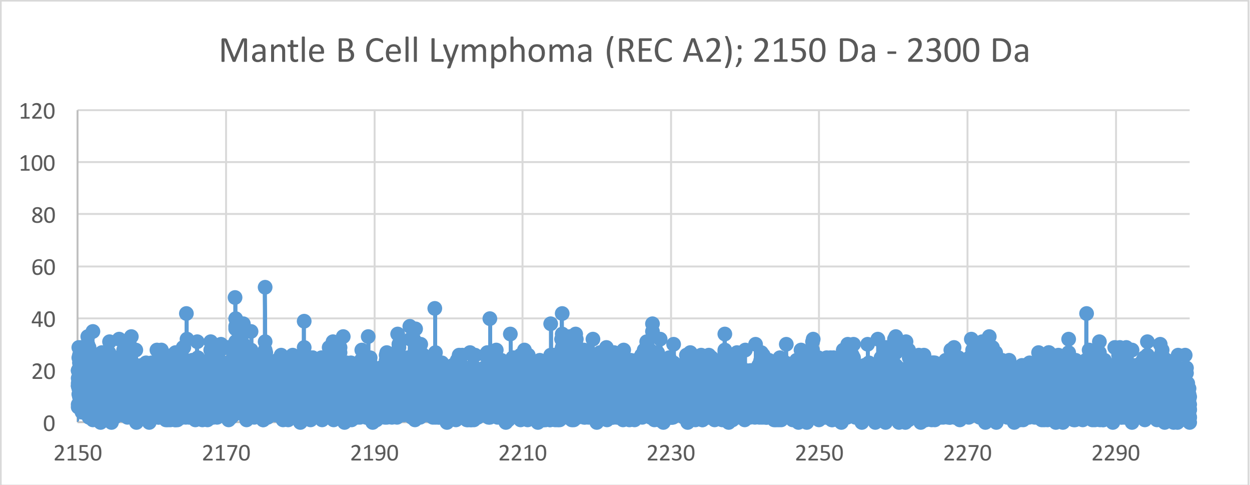
Mantle B Cell Lymphoma (REC A2); 2150 Da - 2300 Da
120
100
80
60
40
20
0
2150
2170
2190
2210
2230
2250
2270
2290

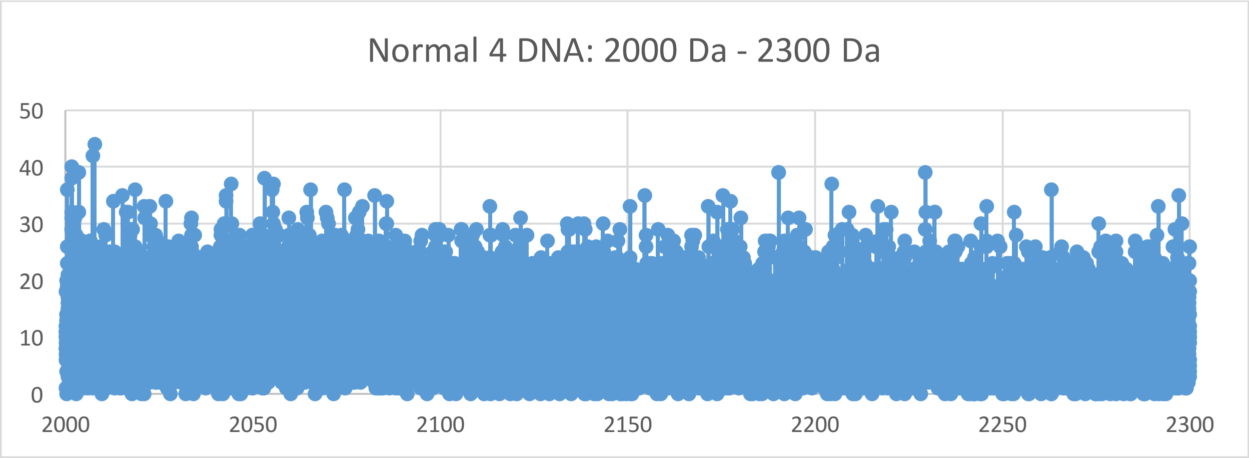
Normal 4 DNA: 2000 Da - 2300 Da
50
40
30
20
10
0
2000
2050
2100
2150
2200
2250
2300

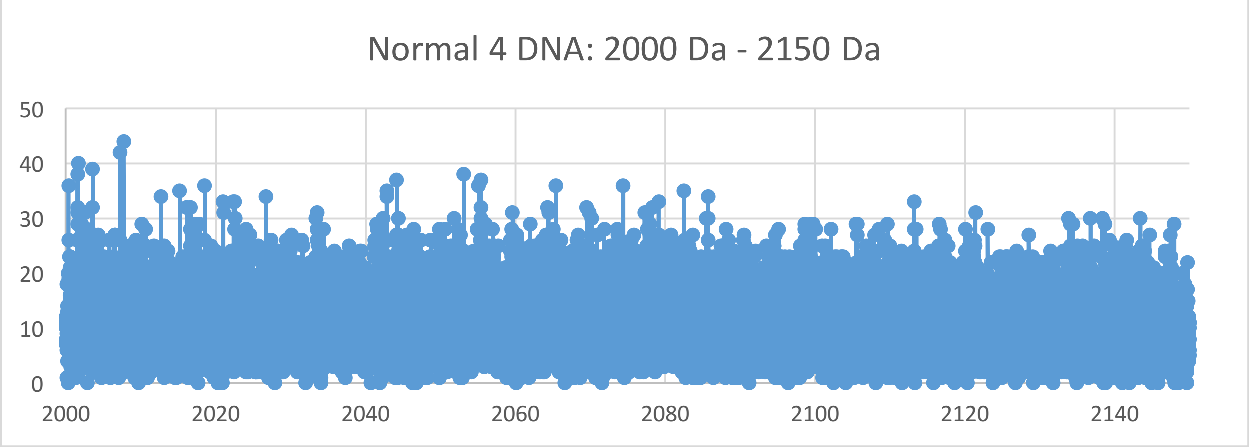
Normal 4 DNA: 2000 Da - 2150 Da
50
40
30
20
10
0
2000
2020
2040
2060
2080
2100
2120
2140

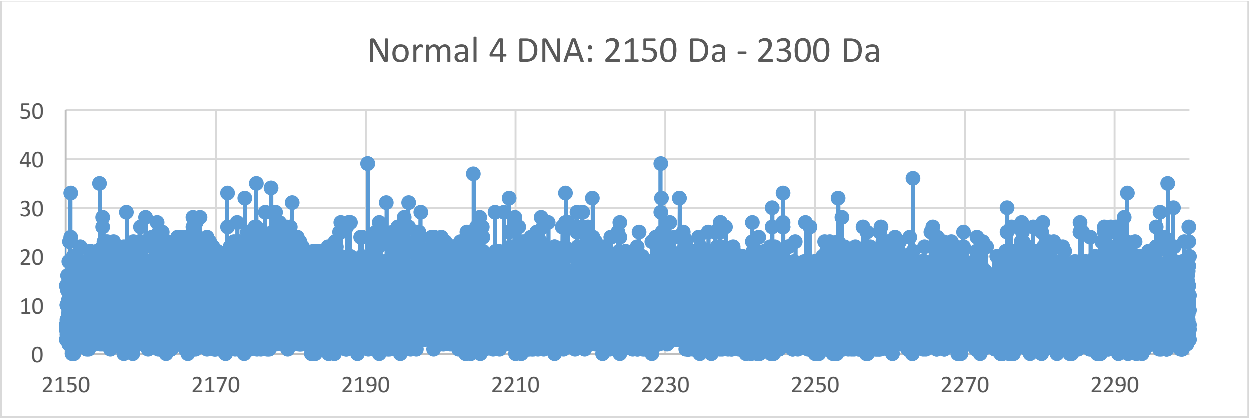
Normal 4 DNA: 2150 Da - 2300 Da
50
40
30
20
10
0
2150
2170
2190
2210
2230
2250
2270
2290

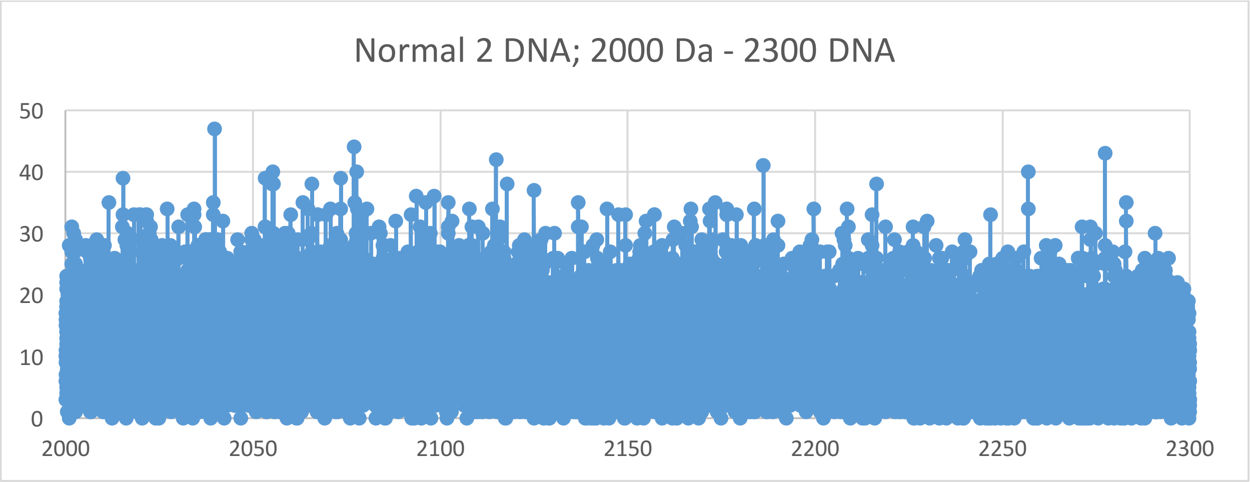
Normal 2 DNA; 2000 Da - 2300 DNA
50
40
30
20
10
0
2000
2050
2100
2150
2200
2250
2300

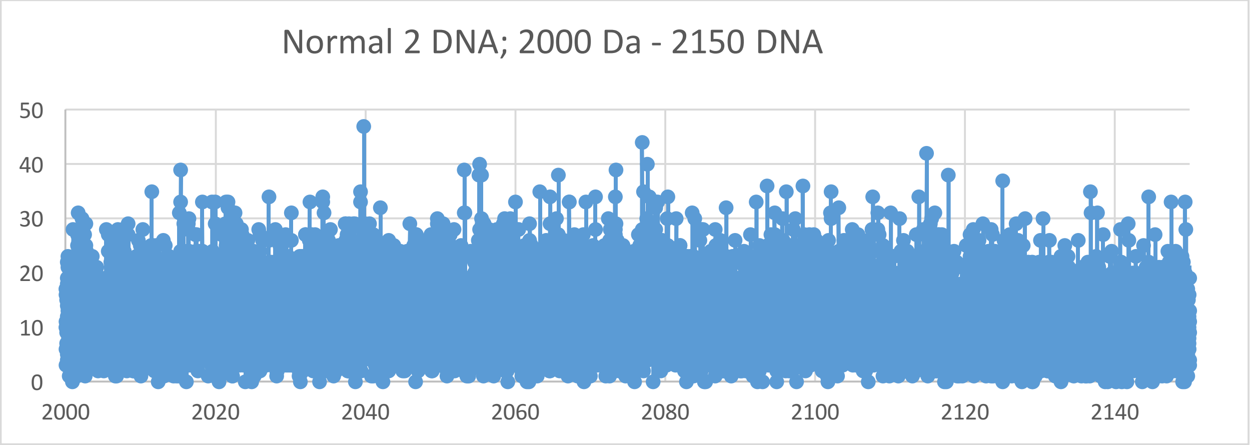
Normal 2 DNA; 2000 Da - 2150 DNA
50
40
30
20
10
0
2000
2020
2040
2060
2080
2100
2120
2140

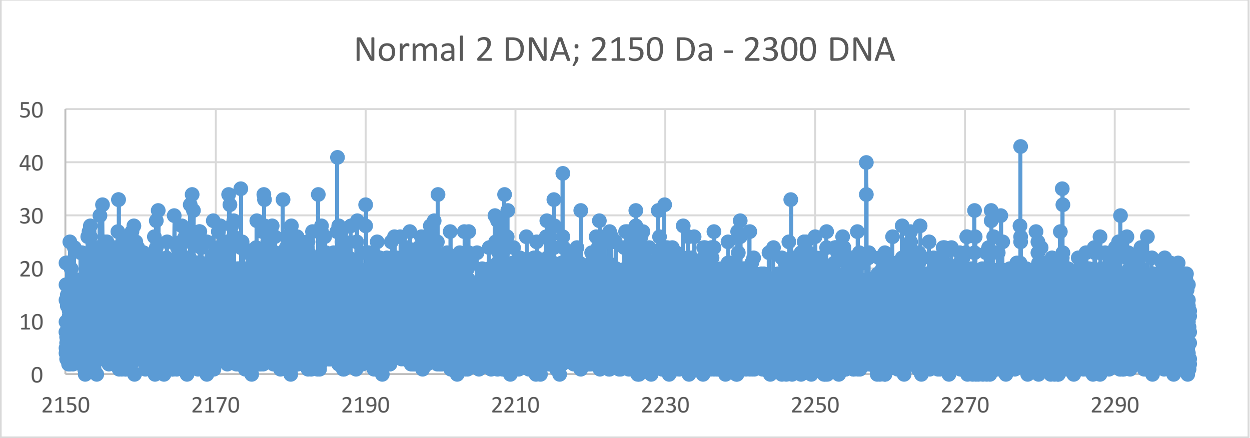
Normal 2 DNA; 2150 Da - 2300 DNA
50
40
30
20
10
0
2150
2170
2190
2210
2230
2250
2270
2290

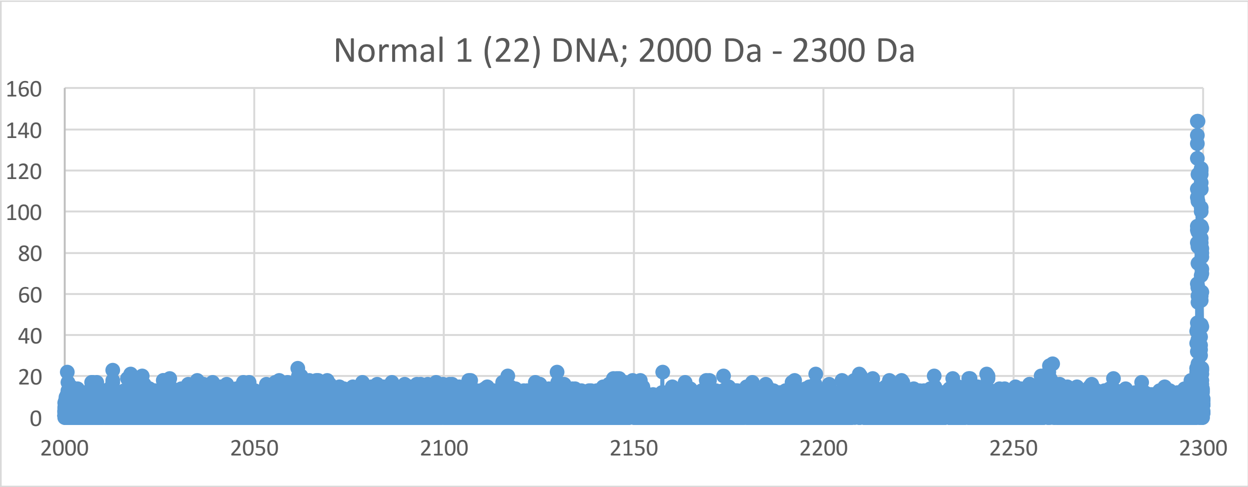
Normal 1 (22) DNA; 2000 Da - 2300 Da
160
140
120
100
80
60
40
20
0
2000
2050
2100
2150
2200
2250
2300

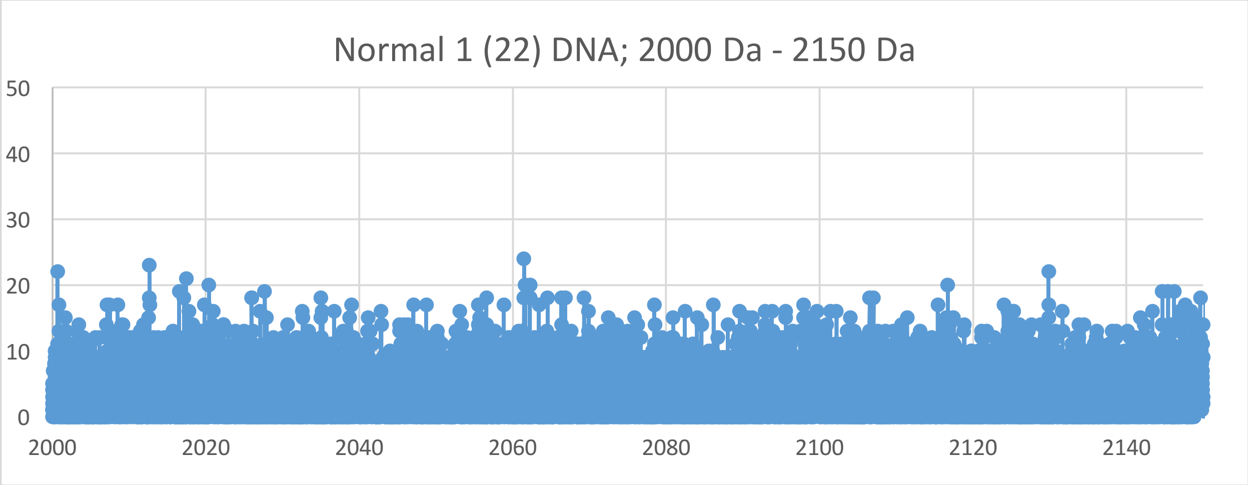
Normal 1 (22) DNA; 2000 Da - 2150 Da
50
40
30
20
10
0
2000
2020
2040
2060
2080
2100
2120
2140

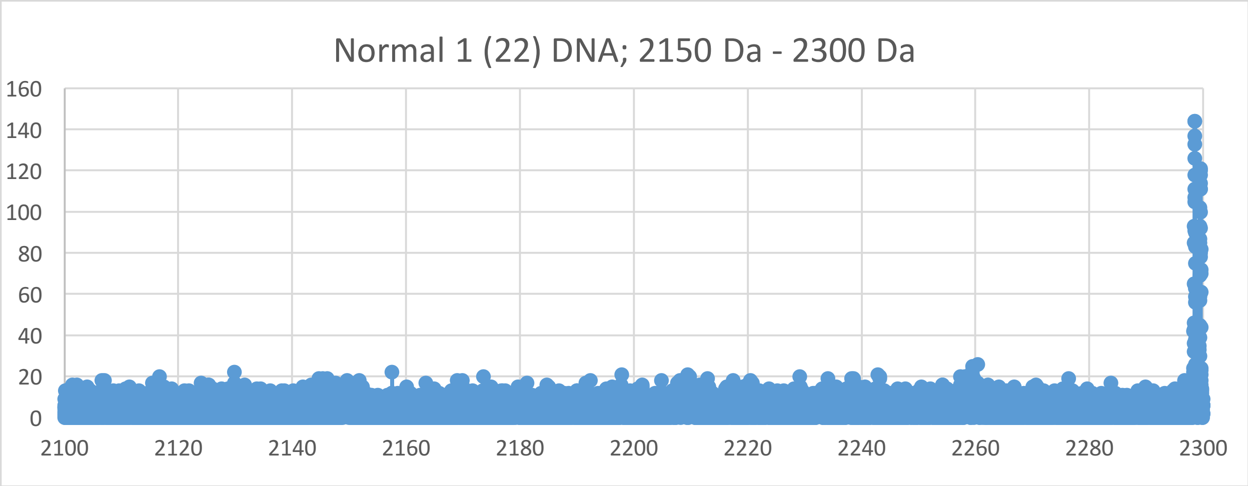
Normal 1 (22) DNA; 2150 Da - 2300 Da
160
140
120
100
80
60
40
20
0
2100
2120
2140
2160
2180
2200
2220
2240
2260
2280
2300

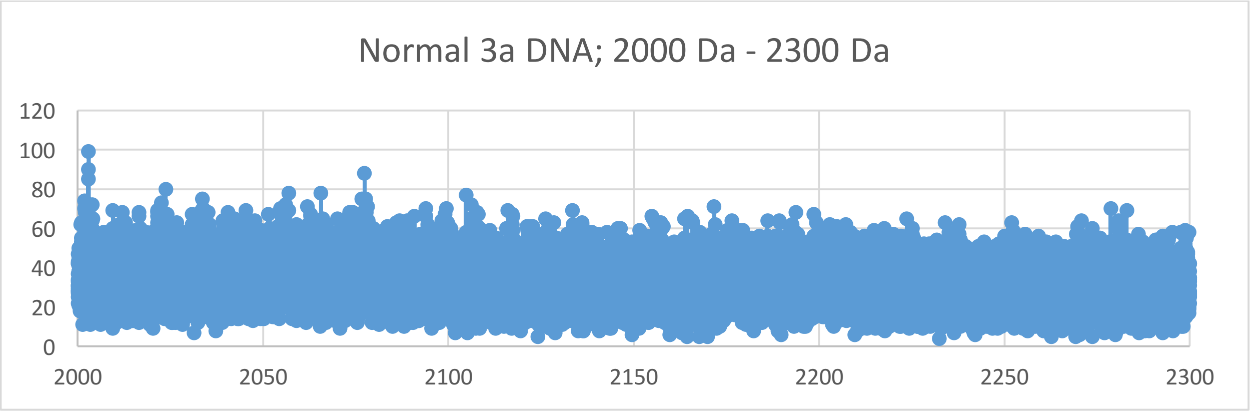
Normal 3a DNA; 2000 Da - 2300 Da
120
100
80
60
40
20
0
2000
2050
2100
2150
2200
2250
2300

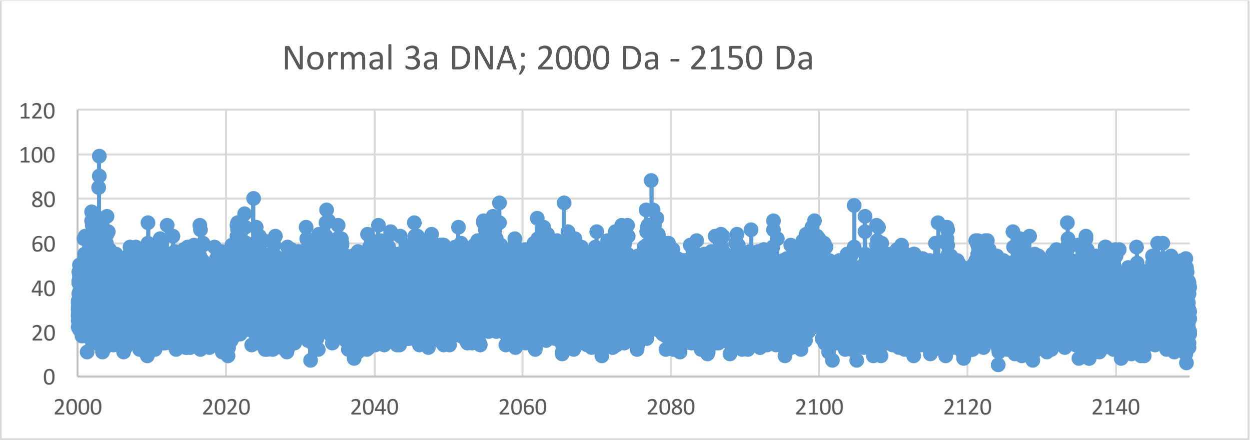
Normal 3a DNA; 2000 Da - 2150 Da
120
100
80
60
40
20
0
2000
2020
2040
2060
2080
2100
2120
2140

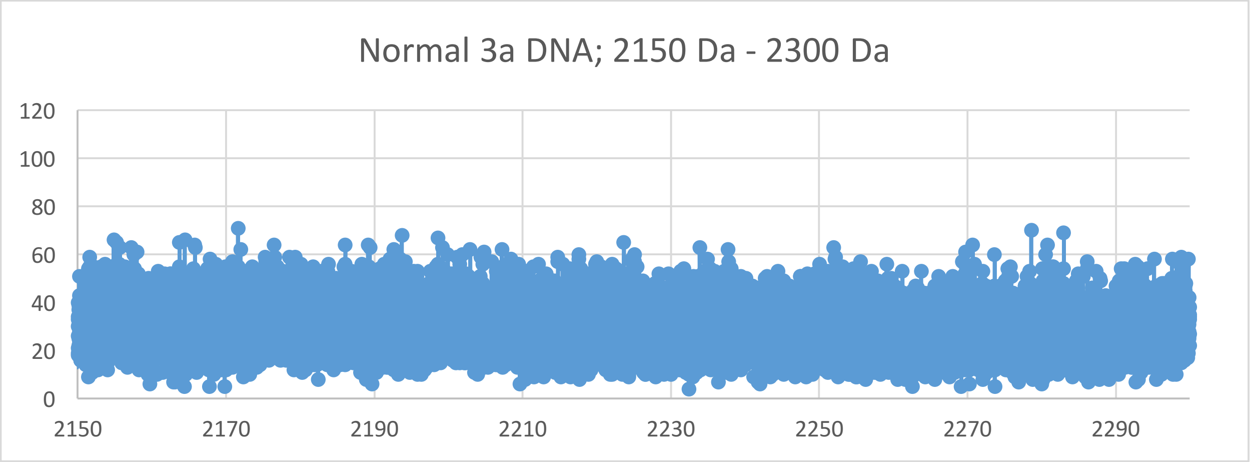
Normal 3a DNA; 2150 Da - 2300 Da
120
100
80
60
40
20
0
2150
2170
2190
2210
2230
2250
2270
2290
