## Supplementary Materials XV for "Oncogenesis and Aging by Isotopic Functionalizations of the Proteins and Nucleic Acids"

XV Supplementary Materials – New Mass Spectra from 2300 Da - 2600 Da

Results for Supplementary 15

So for the pentanucleotides at $(phos)_{15}$, the A, T, G and C peaks for cancer increases in peak intensities and increase in numbers with the $(phos)_{15}$ to $(phos)_{14}$ dephosphorylations for telomeric oligonucleotides. The pentanucleotides dramatically change fragmentation patterns for cancer at $(phos)_{14}$. But as decrease the phosphate by 1 unit, the pentanucleotides change so that the telomeric cancer pentanucleotides more fragment and break with stabilities from the DNA with more number and stabilities for the A, T, C and G for $(phos)_{15}$ to $(phos)_{14}$. The masses range from 2535 Da to 2267 Da. The cancer fragment more intense and stable pentanucleotides in this mass range. See Supplementary Materials XV for graphs. It seems that below the saturation range of some number of phosphates, the telomeres and mating telomeres behave in similar way within cancer and normal DNA. But here the telomeric and mating telomeric pieces behave differently. So this suggests cancer may be favored and distinguished from normal by changing saturated phosphorylations from $(phos)_{15}$ to $(phos)_{14}$. The A, T, C and G pentanucleotide peaks for cancer increase with these dephosphorylations of the pentanucleotides. The dephosphorylations from $(phos)_{15}$ to $(phos)_{14}$ caused the resulting large increase in intensities and stabilities for cancer pentanucleotides for mating telomeric pentanucleotides and decreased isotopic enrichments for mating telomeres. It is important to consider that there are less intense peaks for the pentanucleotides relative to the mononucleotides, dinucleotides, trinucleotides and tetranucleotides for the saturated phosphate units. For the pentanucleotides, the intensities of cancer increased and the intensities of normal fragments decreased from $(phos)_{15}$ to $(phos)_{14}$. The dephosphorylations of telomeric pentanucleotides from $(phos)_{15}$ to $(phos)_{14}$ slightly increased isotopic enrichments of heavy isotopes of $^{15}N$, $^{13}C$ and $^{17}O$. The dephosphorylations from $(phos)_{15}$ to $(phos)_{14}$ also caused the mating telomeric pentanucleotides to be less intense in cancer DNA and less isotopically enriched. It seems the dephosphorylations are related to the isotopic enrichments and depletions; but the telomeres are more intense in cancer and slightly more isotopically enriched. The mating telomeric pentanucleotides of cancer tended to also be less intense and less isotopically enriched by this dephosphorylations; so it seems the cancer stabilities of the telomeric and mating telomeric pentanucleotides are related to their isotopic enrichments from the dephosphorylations from $(phos)_{15}$ to $(phos)_{14}$ and the mutations are purely driven by dephosphorylations and enrichments. Isotopic enrichments strengthen the mating telomeric cancer pentanucleotides while strenthening normal telomeric pentanucleotides. The large intense cancer mating telomeric pentanucleotides were also more isotopically enriched by phosphorylating from $(phos)_{14}$ to $(phos)_{15}$. The greater intense cancer mating telomeric pentanucleotides were slightly more isotopically enriched by heavy isotopes at $(Phos)_{15}$. The dephosphorylations from $(phos)_{15}$ to $(phos)_{14}$ slightly increased the isotopic enrichments of the cancer for the telomeric pentanucleotides. Such seems compatible for replications and transcriptions as stable telomeres seem compatible to unstable mating telomeres for the unravelings and constructions of DNA mating telomeres onto the building templates of the telomeres. But in this case of all normal telomeric and mating telomeric pentanucleotides are negligibly enriched and unstable then the replications, transcriptions and translations cannot occur normally. The dephosphorylations from $(phos)_{15}$ to $(phos)_{14}$ caused demethylations to cause more cancer intensities and stabilities but greater isotopic enrichments for G containing telomeric pentanucleotides. But the demethylations of the T, C and/or A caused negligible alterations of cancer stabilities and isotopic enrichments for mating telomeres. Where isotopic changes occurred the functionalization effects were not negligible. The methylations of the C pentanucleotide negligibly decreased cancer intensities, enrichments and stabilities. It seems from these data that during biological processes the dephosphorylations and stabilizations of normal or vice versa dephosphorylations and stabilizations of cancer are or can be countered by methylations (and isotopic enrichments) and destabilizations of the normal for cancer or vice versa dephosphorylations and destabilizations of the cancer by methylations as this research discovers coupled phosphorylations and methylations (and isotopic enrichments) or coupled dephosphorylations and methylations (and isotopic enrichments) with involved greater enrichments for normal biological functions and explaining cancer! The dephosphorylations from $(phos)_{15}$ to $(phos)_{14}$ caused acetylations to cause more cancer stabilities for A and C with greater isotopic depletions for mating pentanucleotides and much greater normal stabilities for G and T with greater depletions of light isotopic enrichments for telomeric pentanucleotides. The acetylations from $(phos)_{15}$ to $(phos)_{14}$ caused slightly less cancer stabilities and intensities for A and C of mating telomeres with negligibly more isotopic enrichments of light isotopes for the mating telomeric pentanucleotides. This contrast in A, T, C and G from the de-acetylations and dephosphorylations may reflect the developments of cancer as the pentanucleotides fragment but the other pentanucleotides more hold together for damaged DNA for cancer.

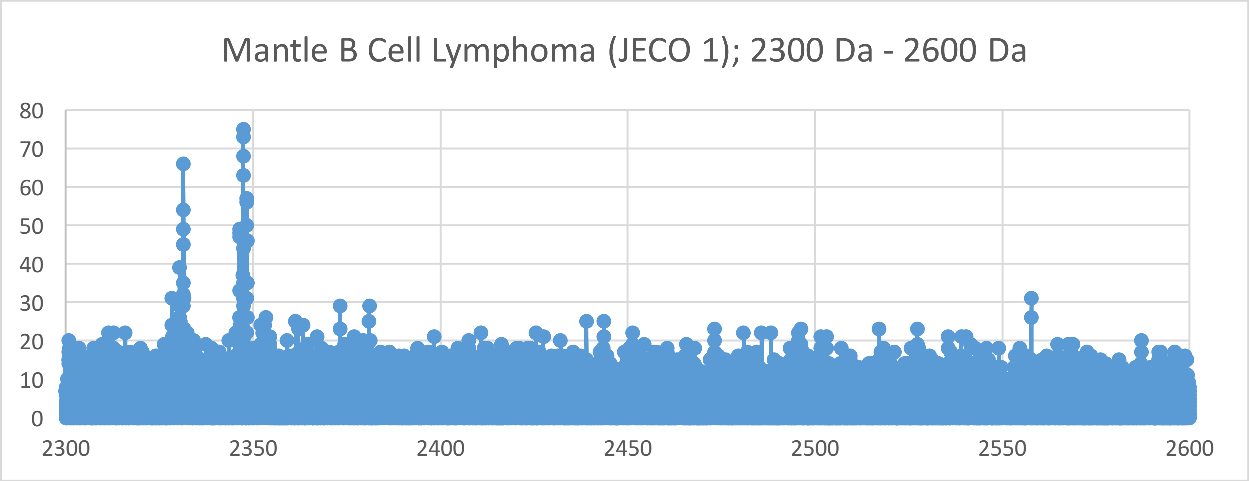
Mantle B Cell Lymphoma (JECO 1); 2300 Da - 2600 Da
80
70
60
50
40
30
20
10
0
2300
2350
2400
2450
2500
2550
2600

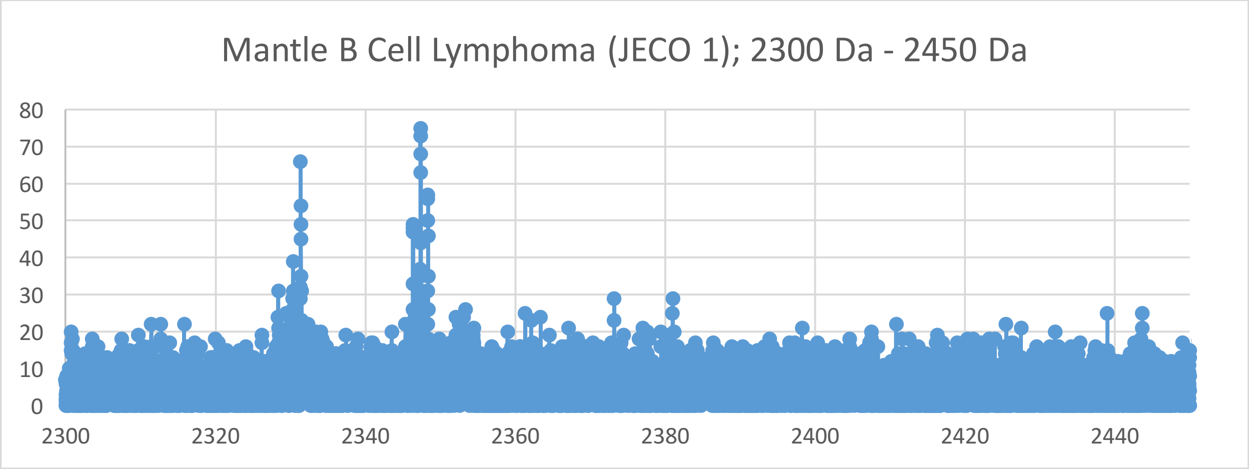
Mantle B Cell Lymphoma (JECO 1); 2300 Da - 2450 Da
80
70
60
50
40
30
20
10
0
2300
2320
2340
2360
2380
2400
2420
2440

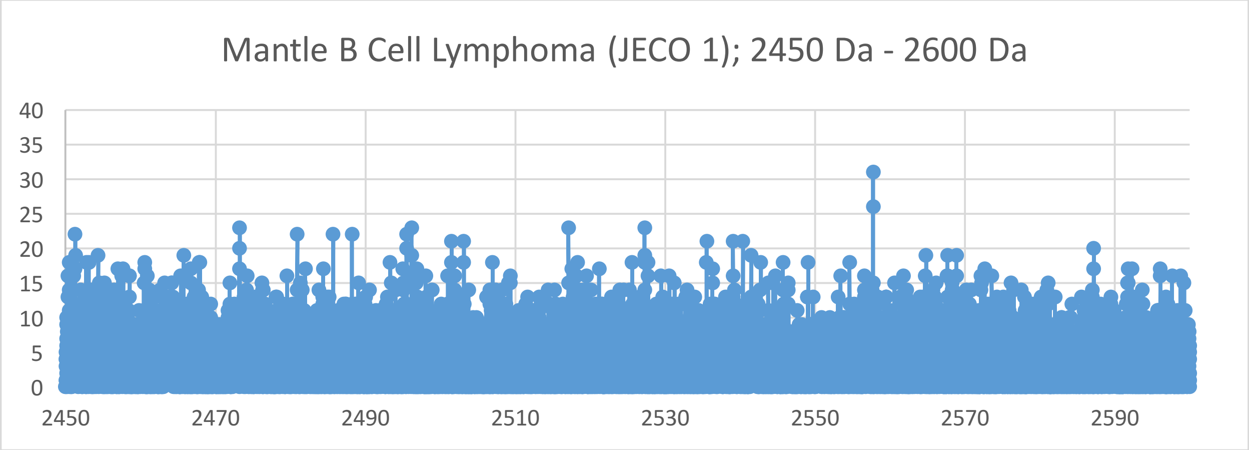
Mantle B Cell Lymphoma (JECO 1); 2450 Da - 2600 Da
40
35
30
25
20
15
10
5
0
2450
2470
2490
2510
2530
2550
2570
2590

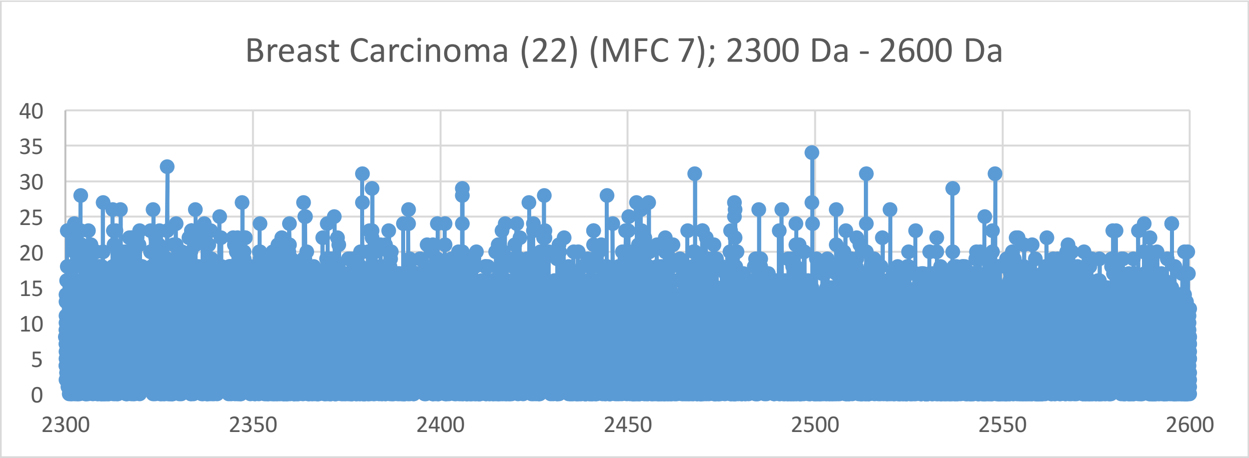
Breast Carcinoma (22) (MFC 7); 2300 Da - 2600 Da
0
5
10
15
20
25
30
35
40
2300
2350
2400
2450
2500
2550
2600

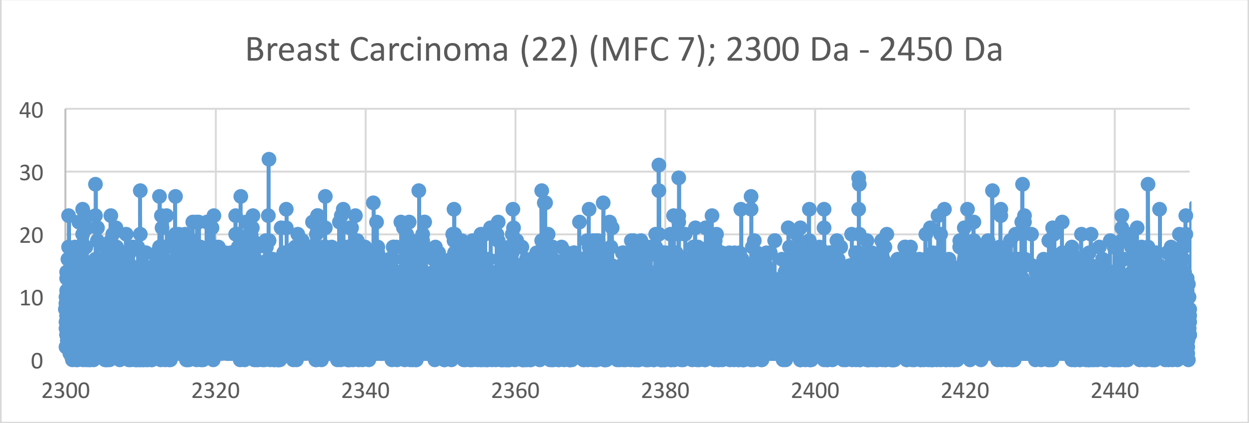
Breast Carcinoma (22) (MFC 7); 2300 Da - 2450 Da
0
10
20
30
40
2300
2320
2340
2360
2380
2400
2420
2440

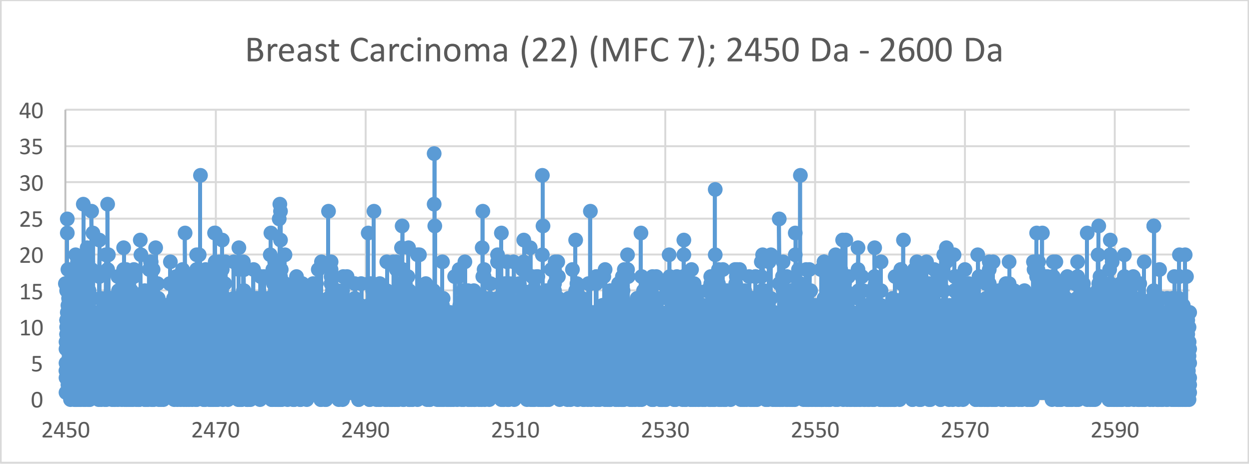
Breast Carcinoma (22) (MFC 7); 2450 Da - 2600 Da
0
5
10
15
20
25
30
35
40
2450
2470
2490
2510
2530
2550
2570
2590

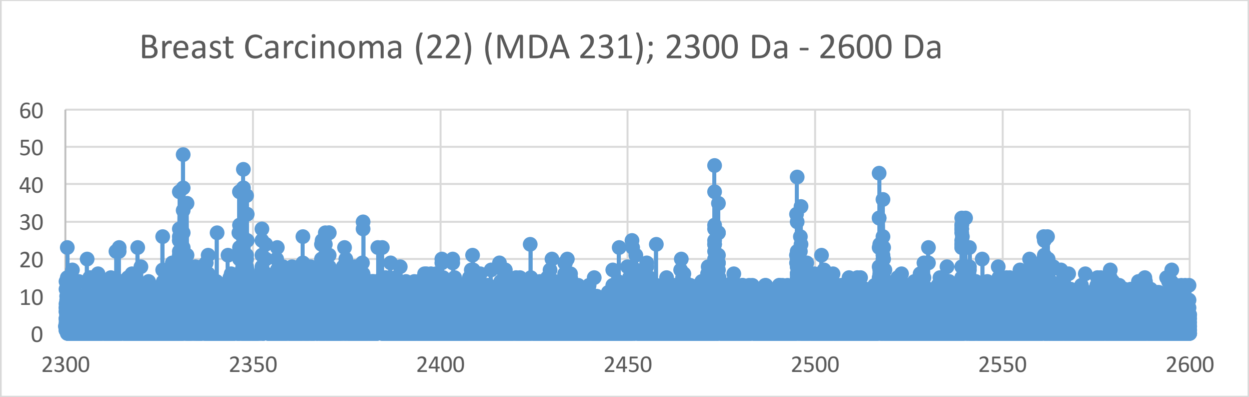
Breast Carcinoma (22) (MDA 231); 2300 Da - 2600 Da
60
50
40
30
20
10
0
2300
2350
2400
2450
2500
2550
2600

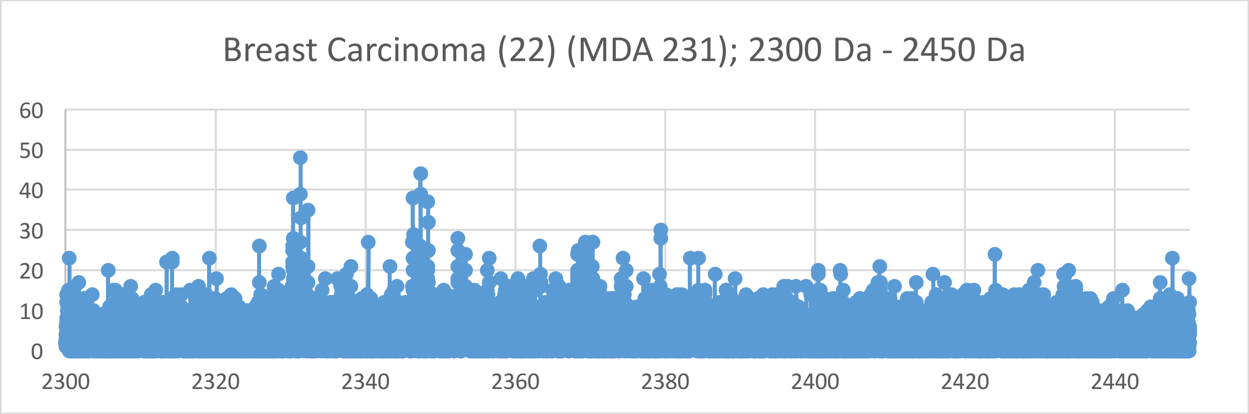
Breast Carcinoma (22) (MDA 231); 2300 Da - 2450 Da
60
50
40
30
20
10
0
2300
2320
2340
2360
2380
2400
2420
2440

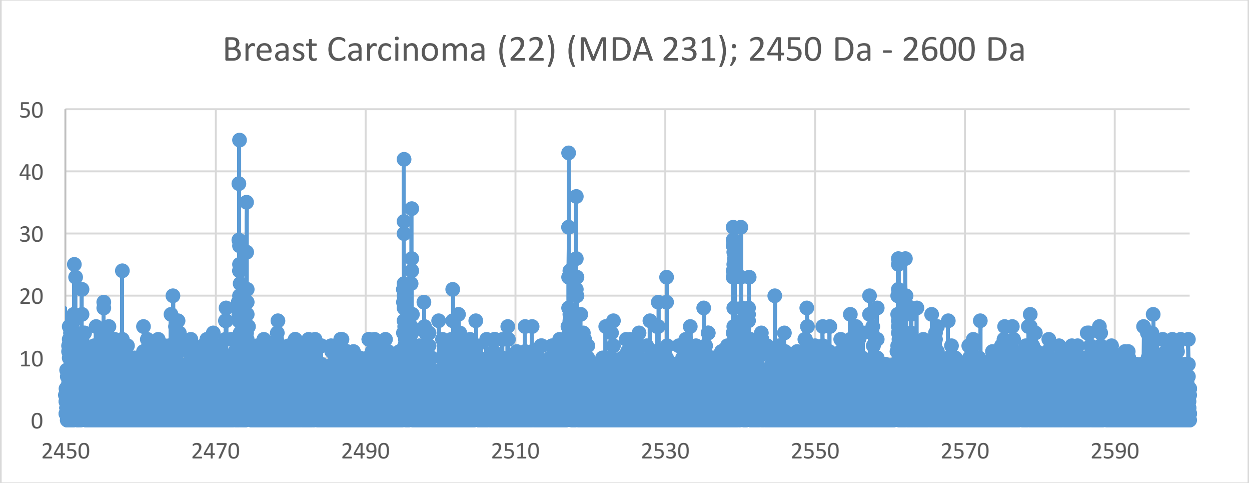
Breast Carcinoma (22) (MDA 231); 2450 Da - 2600 Da
50
40
30
20
10
0
2450
2470
2490
2510
2530
2550
2570
2590

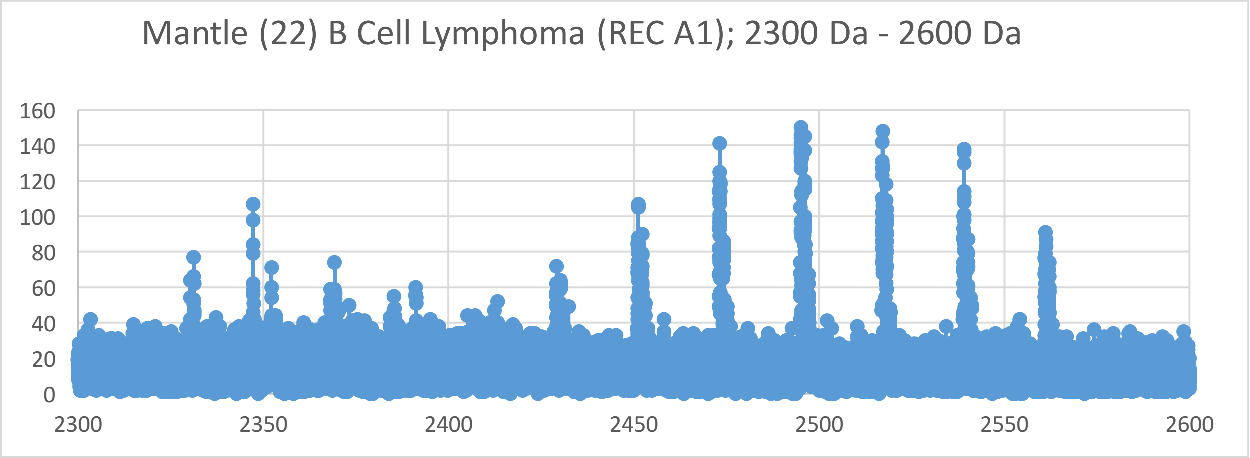
Mantle (22) B Cell Lymphoma (REC A1); 2300 Da - 2600 Da
160
140
120
100
80
60
40
20
0
2300
2350
2400
2450
2500
2550
2600

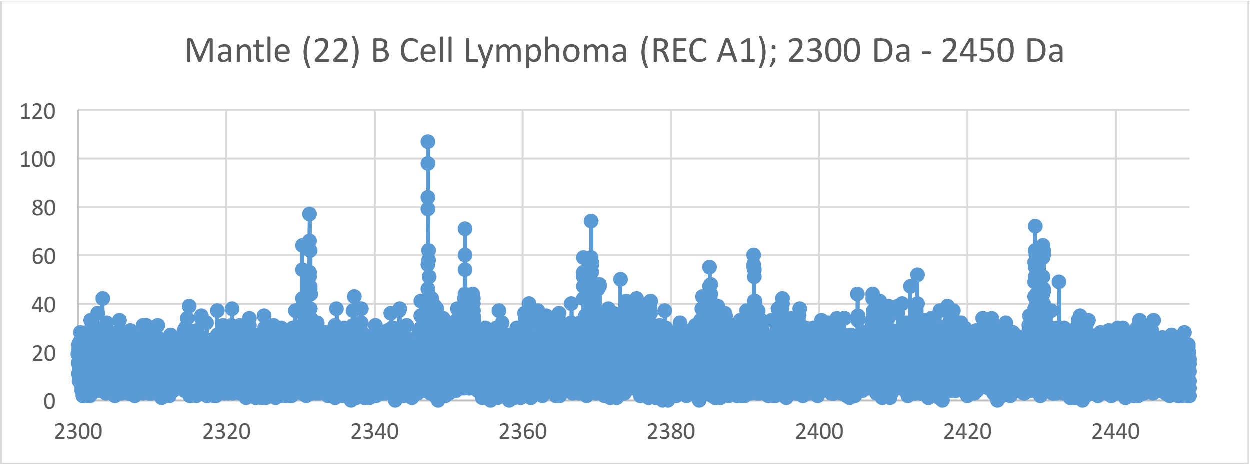
Mantle (22) B Cell Lymphoma (REC A1); 2300 Da - 2450 Da
120
100
80
60
40
20
0
2300
2320
2340
2360
2380
2400
2420
2440

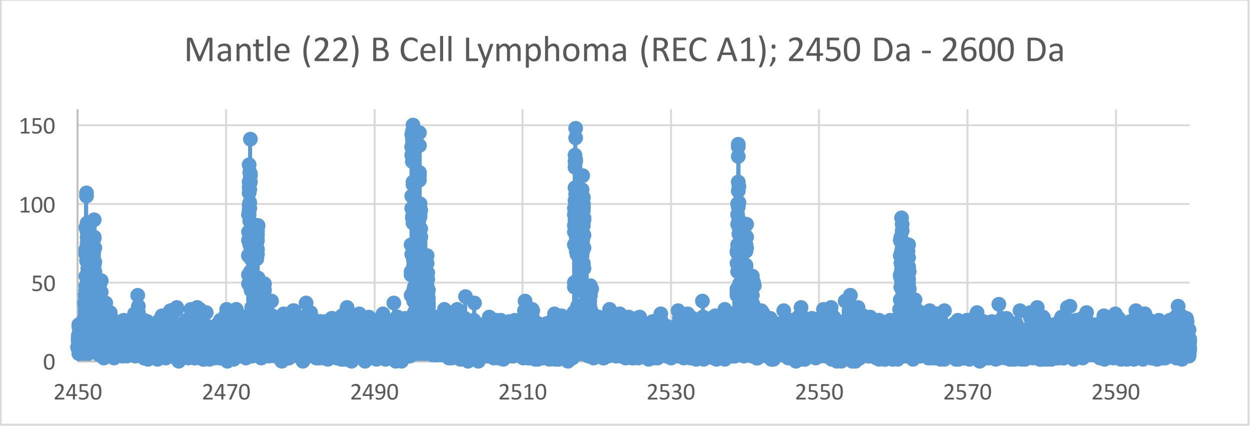
Mantle (22) B Cell Lymphoma (REC A1); 2450 Da - 2600 Da
150
100
50
0
2450
2470
2490
2510
2530
2550
2570
2590

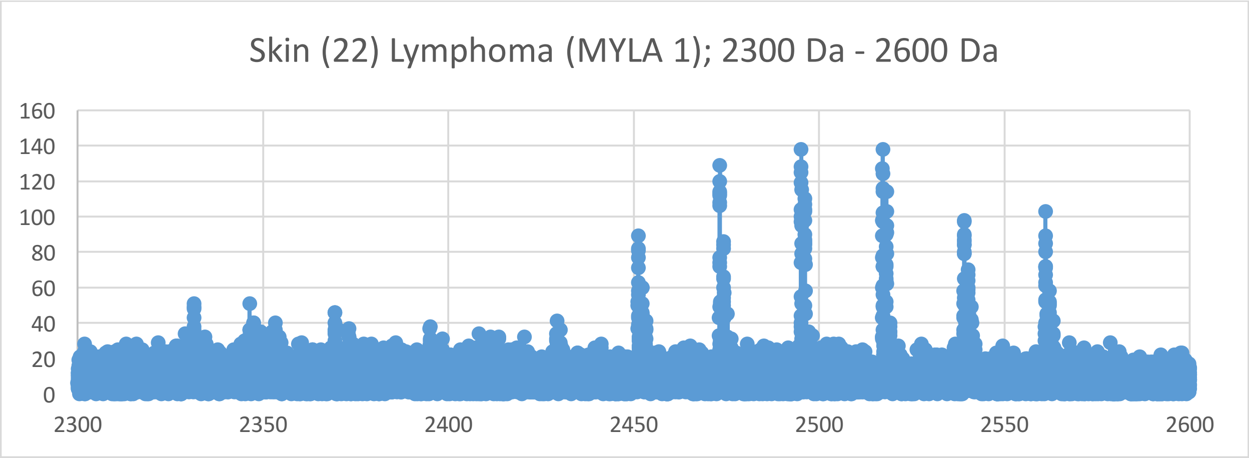
Skin (22) Lymphoma (MYLA 1); 2300 Da - 2600 Da
160
140
120
100
80
60
40
20
0
2300
2350
2400
2450
2500
2550
2600

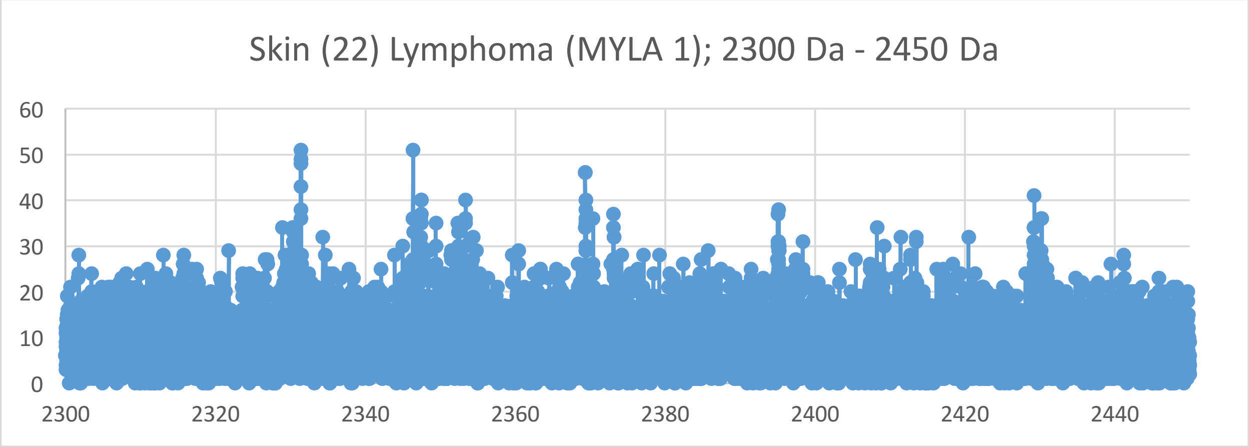
Skin (22) Lymphoma (MYLA 1); 2300 Da - 2450 Da
60
50
40
30
20
10
0
2300
2320
2340
2360
2380
2400
2420
2440

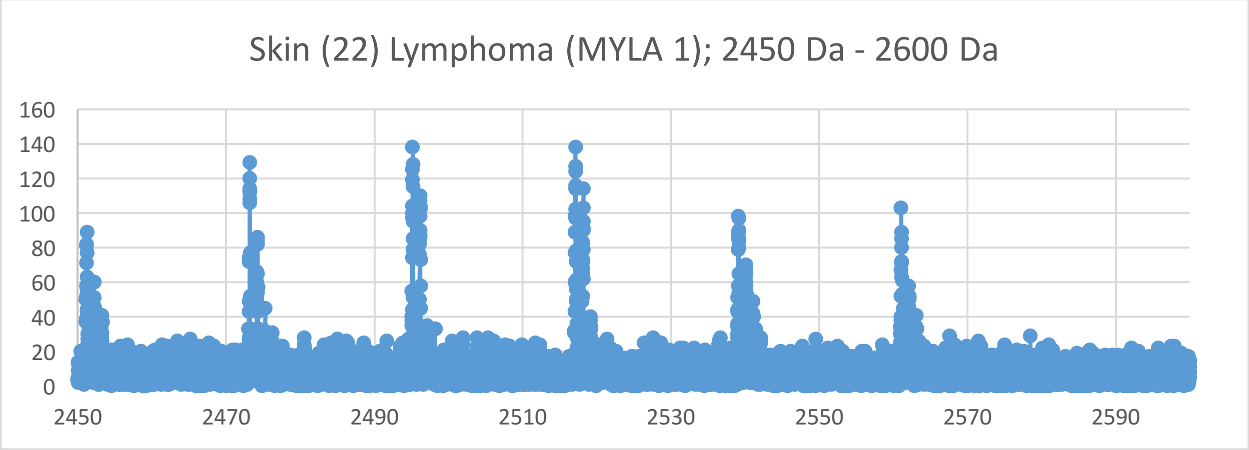
Skin (22) Lymphoma (MYLA 1); 2450 Da - 2600 Da
160
140
120
100
80
60
40
20
0
2450
2470
2490
2510
2530
2550
2570
2590

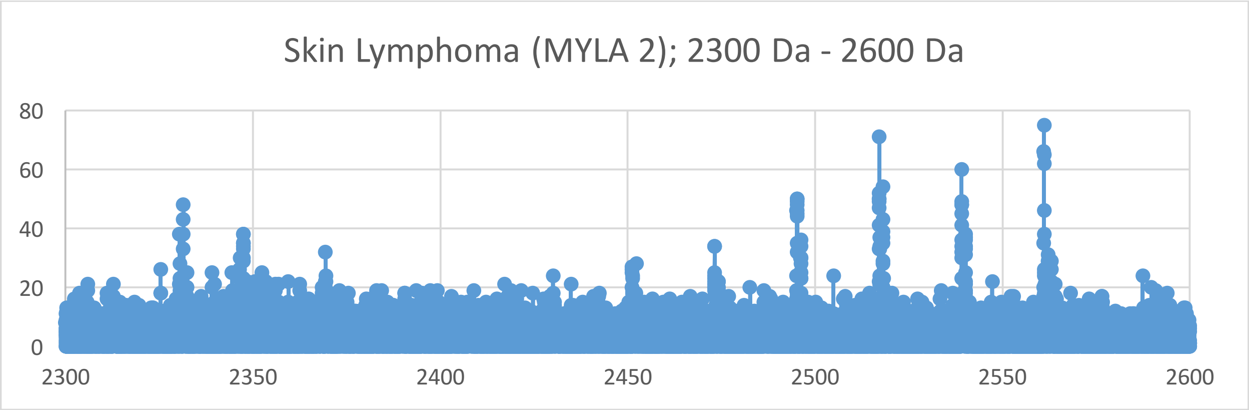
Skin Lymphoma (MYLA 2); 2300 Da - 2600 Da
0
20
40
60
80
2300
2350
2400
2450
2500
2550
2600

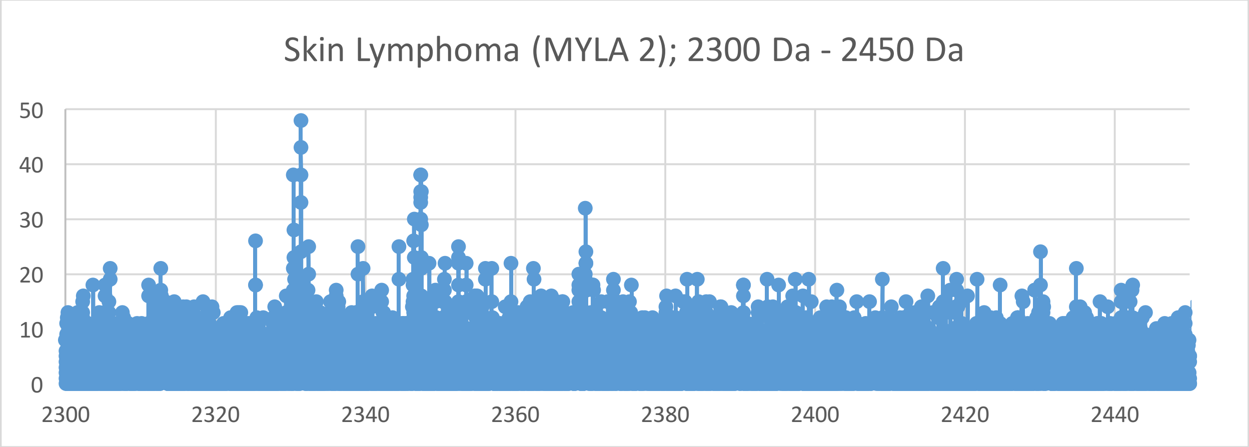
Skin Lymphoma (MYLA 2); 2300 Da - 2450 Da
0
10
20
30
40
50
2300
2320
2340
2360
2380
2400
2420
2440

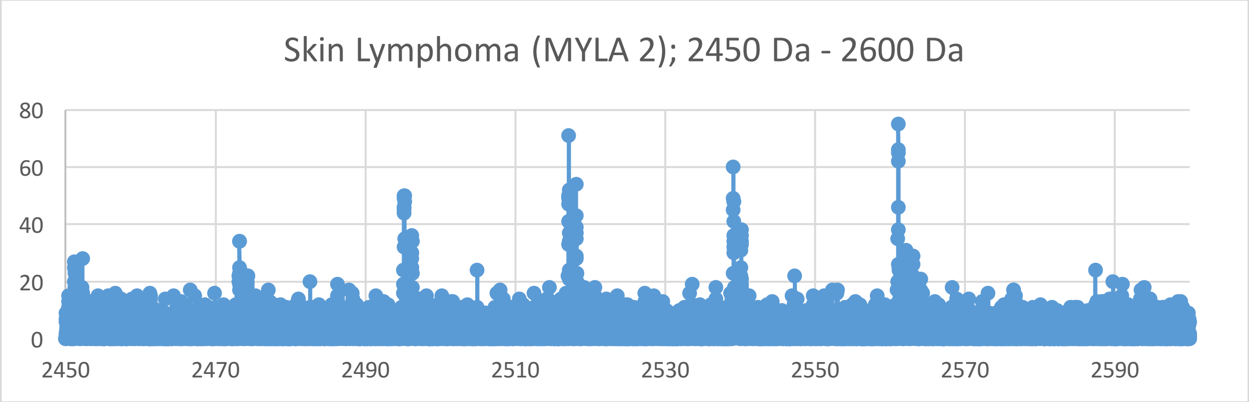
Skin Lymphoma (MYLA 2); 2450 Da - 2600 Da
0
20
40
60
80
2450
2470
2490
2510
2530
2550
2570
2590

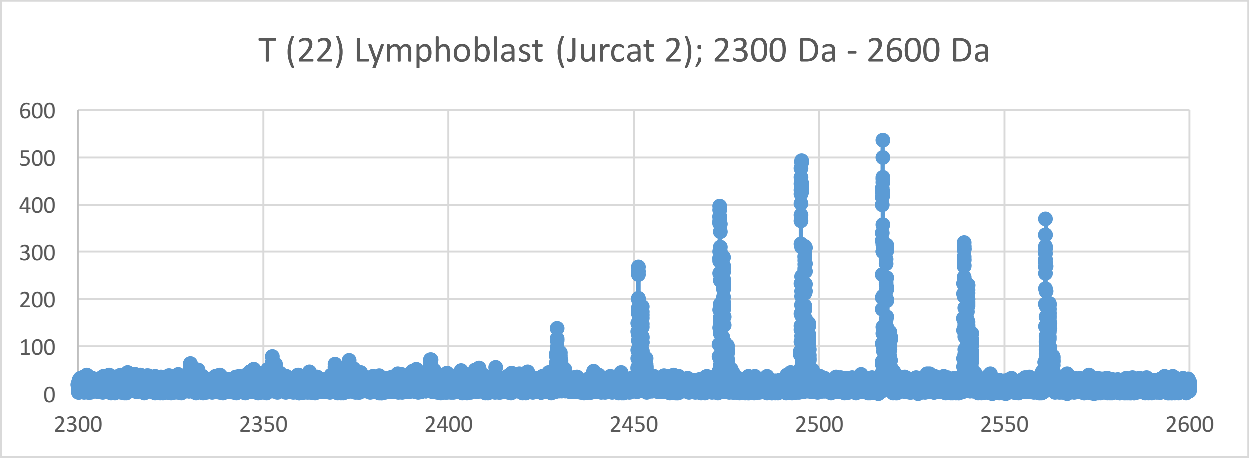
T (22) Lymphoblast (Jurcat 2); 2300 Da - 2600 Da
600
500
400
300
200
100
0
2300
2350
2400
2450
2500
2550
2600

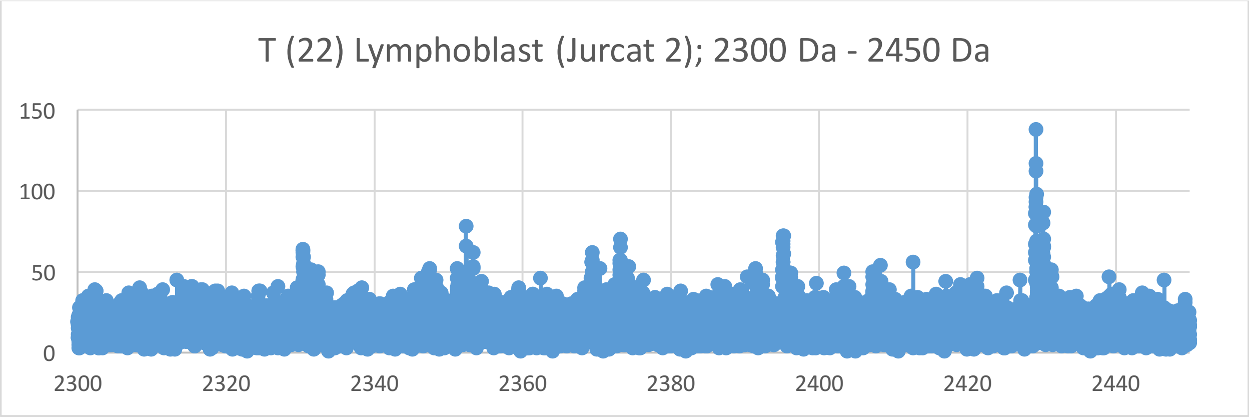
T (22) Lymphoblast (Jurcat 2); 2300 Da - 2450 Da
150
100
50
0
2300
2320
2340
2360
2380
2400
2420
2440

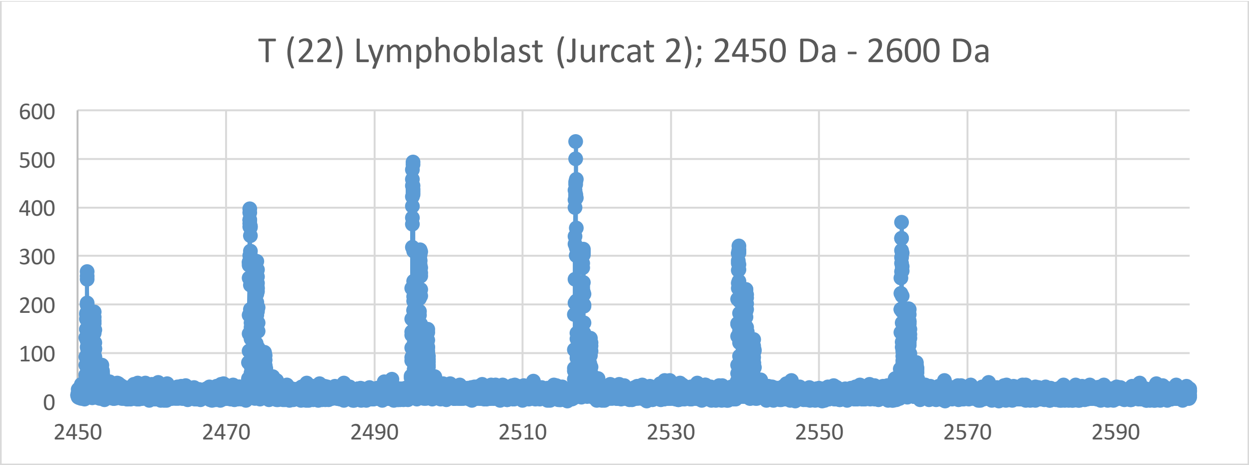
T (22) Lymphoblast (Jurcat 2); 2450 Da - 2600 Da
600
500
400
300
200
100
0
2450
2470
2490
2510
2530
2550
2570
2590

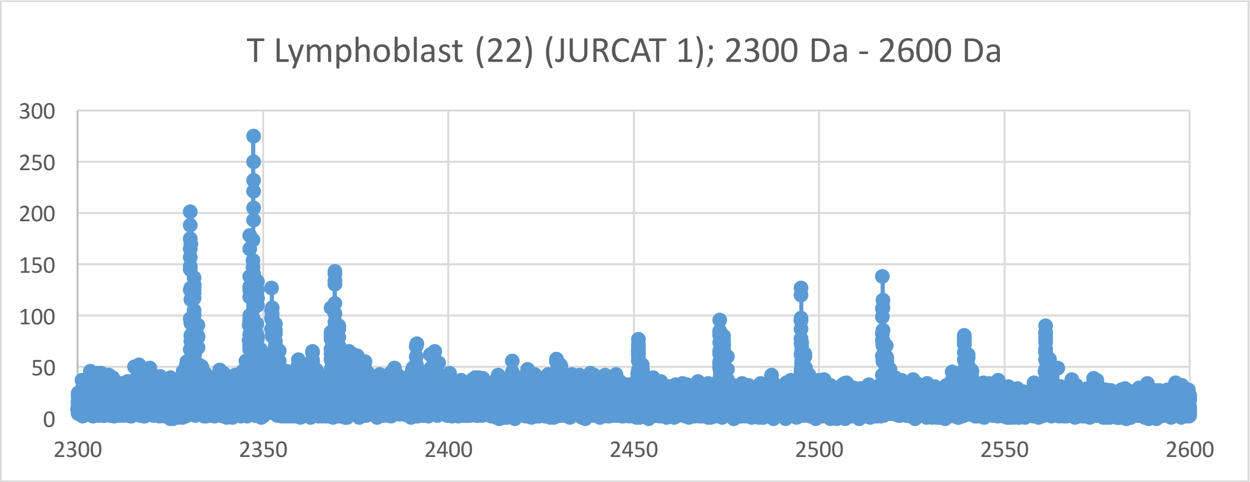
T Lymphoblast (22) (JURCAT 1); 2300 Da - 2600 Da
300
250
200
150
100
50
0
2300
2350
2400
2450
2500
2550
2600

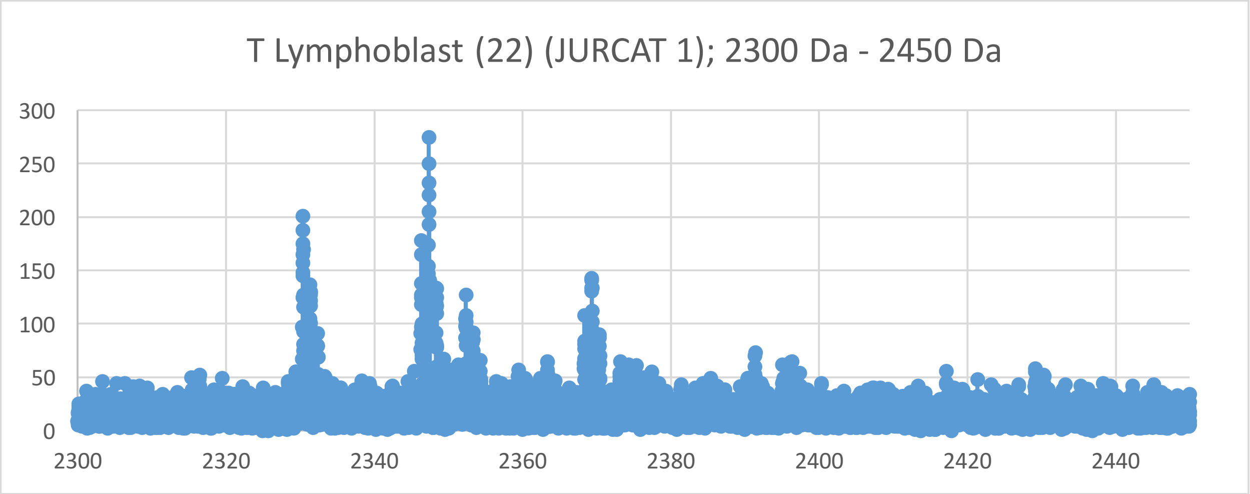
T Lymphoblast (22) (JURCAT 1); 2300 Da - 2450 Da
300
250
200
150
100
50
0
2300
2320
2340
2360
2380
2400
2420
2440

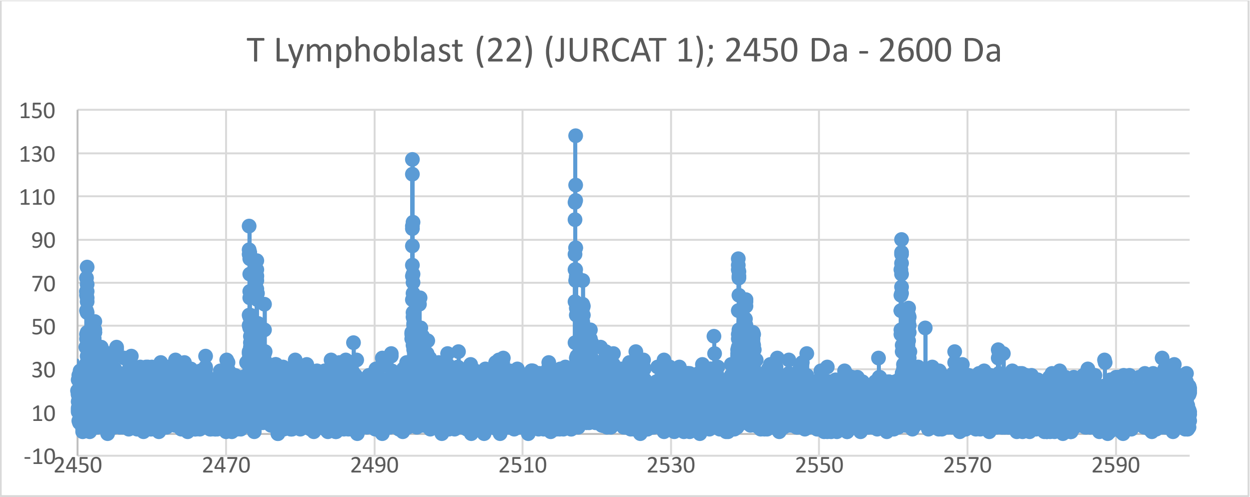
T Lymphoblast (22) (JURCAT 1); 2450 Da - 2600 Da
150
130
110
90
70
50
30
10
-10
2450
2470
2490
2510
2530
2550
2570
2590

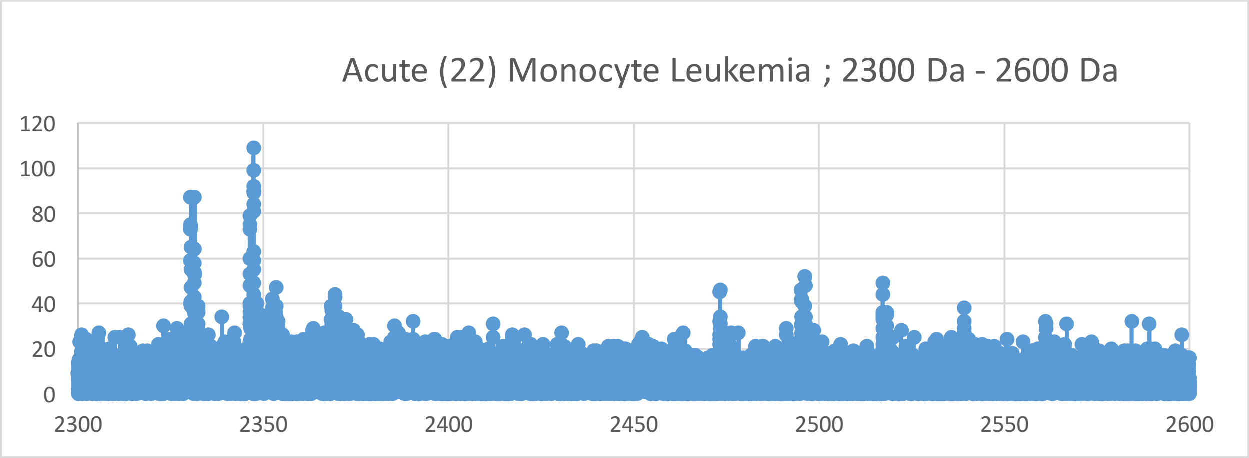
Acute (22) Monocyte Leukemia ; 2300 Da - 2600 Da
120
100
80
60
40
20
0
2300
2350
2400
2450
2500
2550
2600

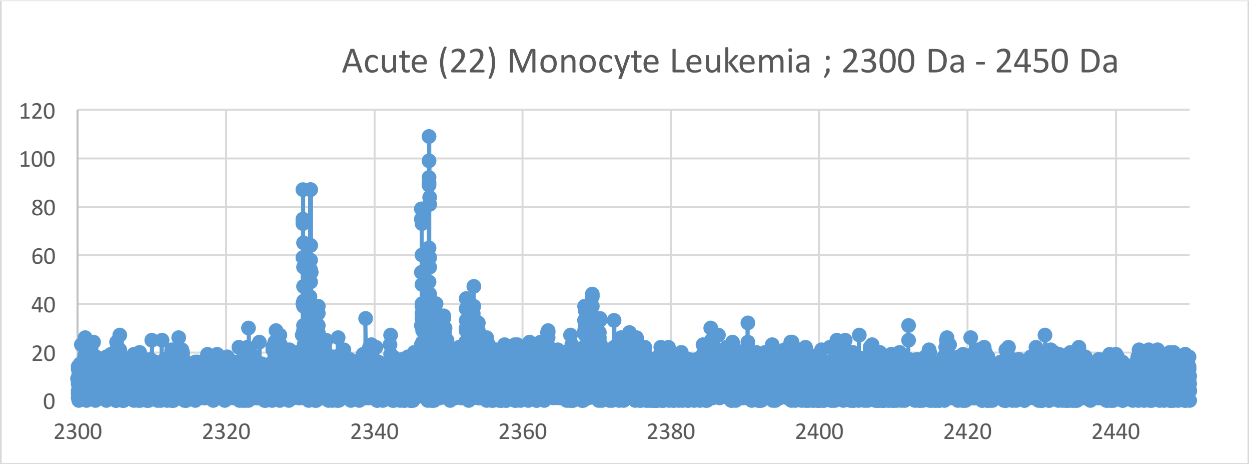
Acute (22) Monocyte Leukemia ; 2300 Da - 2450 Da
120
100
80
60
40
20
0
2300
2320
2340
2360
2380
2400
2420
2440

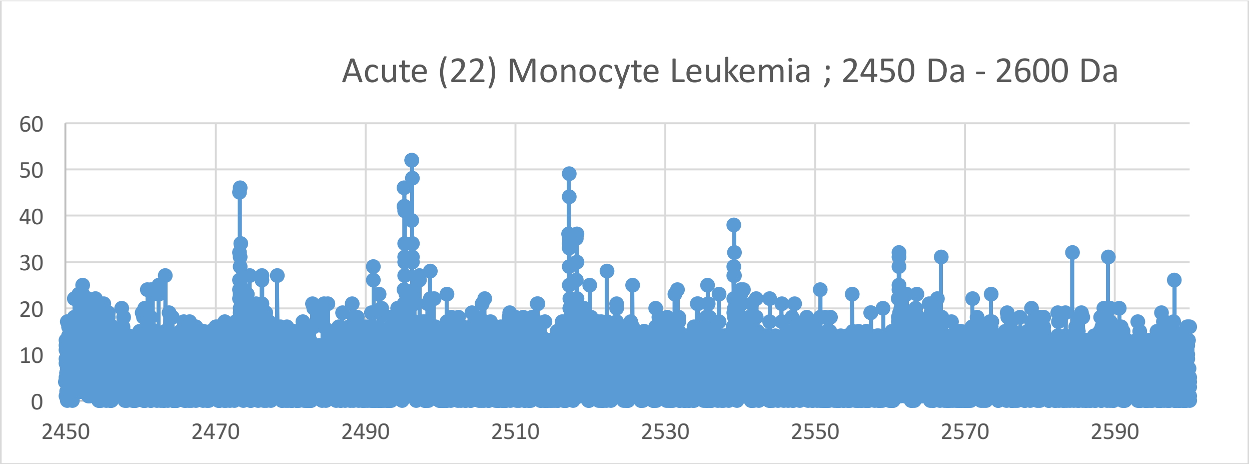
Acute (22) Monocyte Leukemia ; 2450 Da - 2600 Da
60
50
40
30
20
10
0
2450
2470
2490
2510
2530
2550
2570
2590

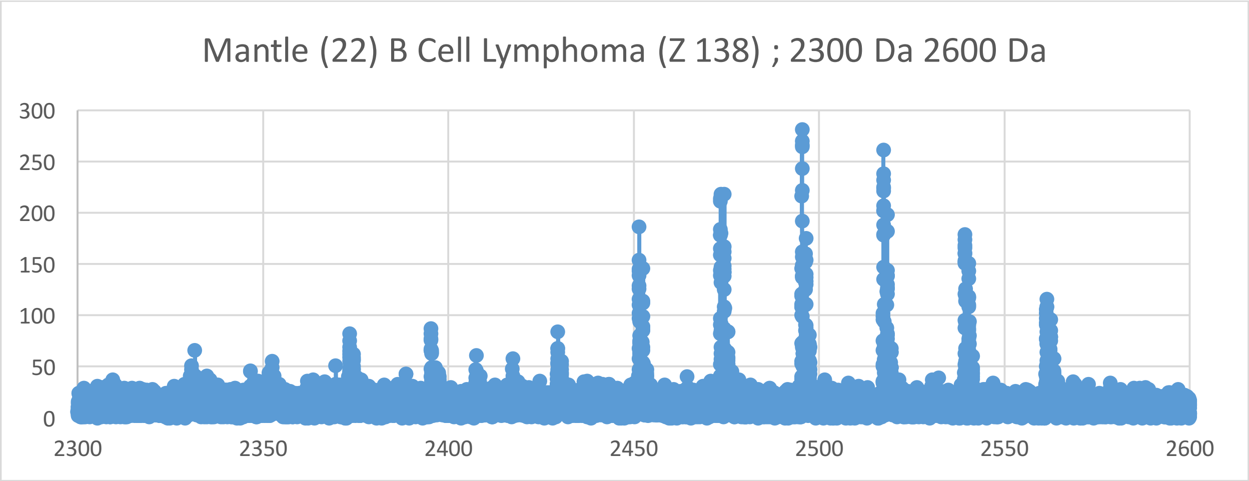
Mantle (22) B Cell Lymphoma (Z 138) ; 2300 Da 2600 Da
300
250
200
150
100
50
0
2300
2350
2400
2450
2500
2550
2600

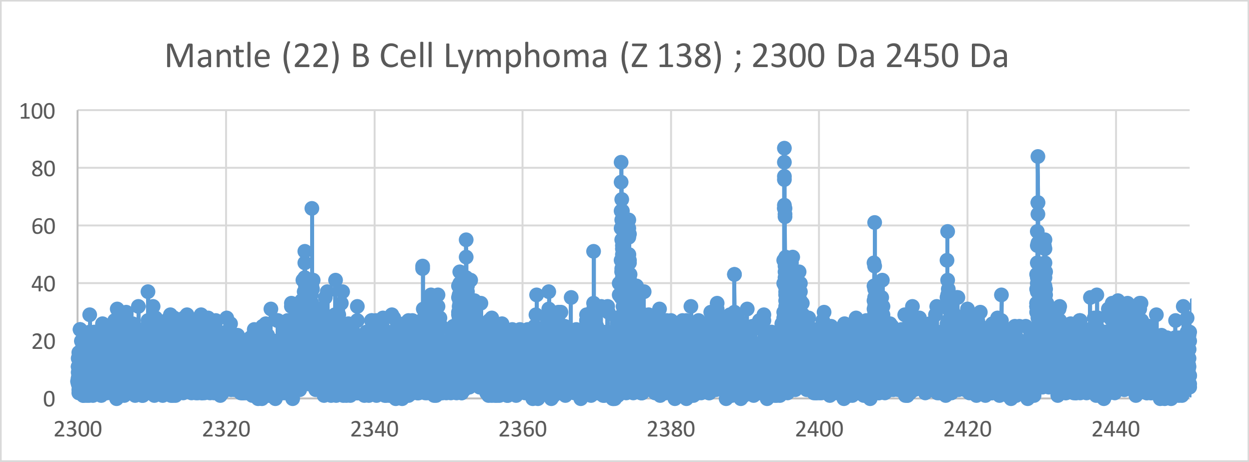
Mantle (22) B Cell Lymphoma (Z 138) ; 2300 Da 2450 Da
100
80
60
40
20
0
2300
2320
2340
2360
2380
2400
2420
2440

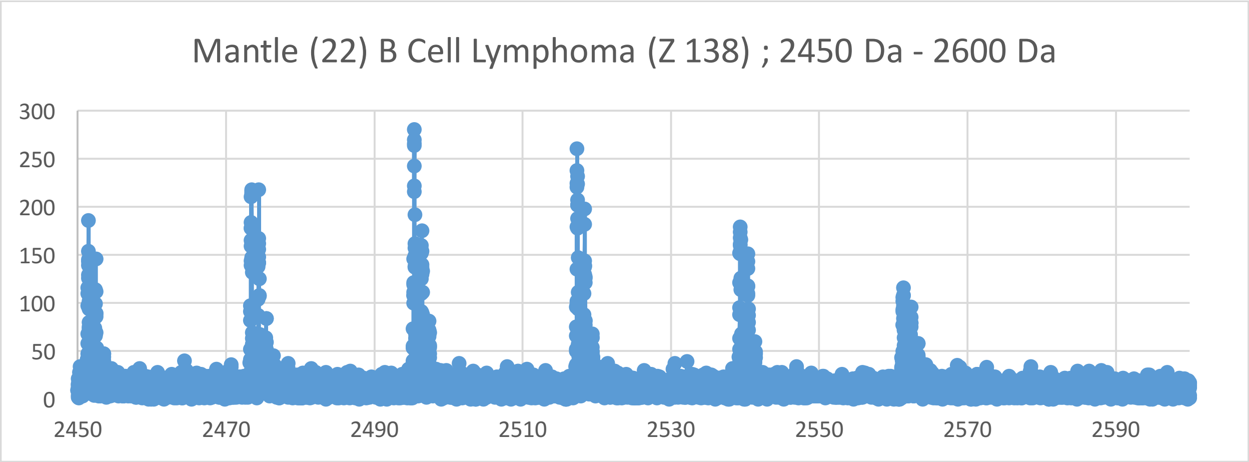
Mantle (22) B Cell Lymphoma (Z 138) ; 2450 Da - 2600 Da
300
250
200
150
100
50
0
2450
2470
2490
2510
2530
2550
2570
2590

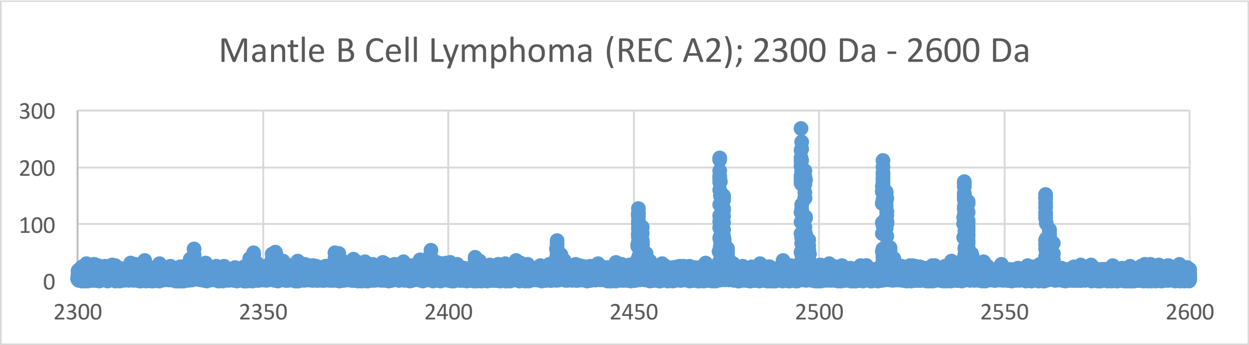
Mantle B Cell Lymphoma (REC A2); 2300 Da - 2600 Da
300
200
100
0
2300
2350
2400
2450
2500
2550
2600

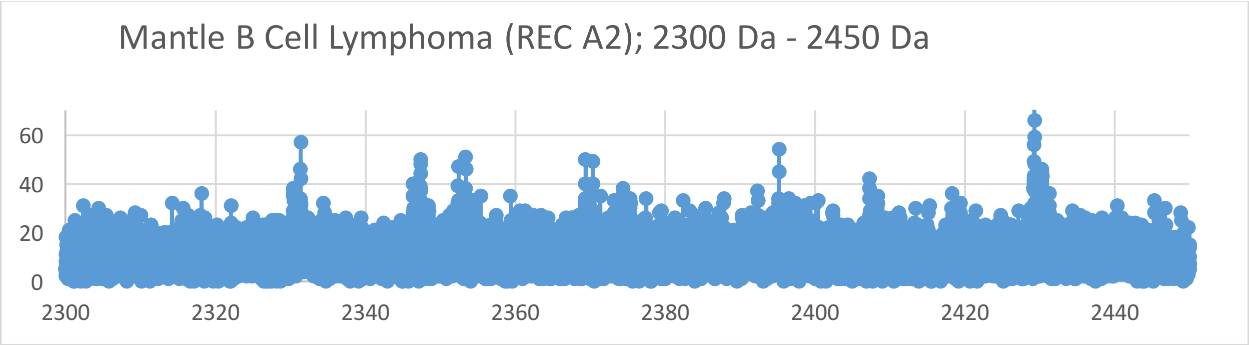
Mantle B Cell Lymphoma (REC A2); 2300 Da - 2450 Da
60
40
20
0
2300
2320
2340
2360
2380
2400
2420
2440

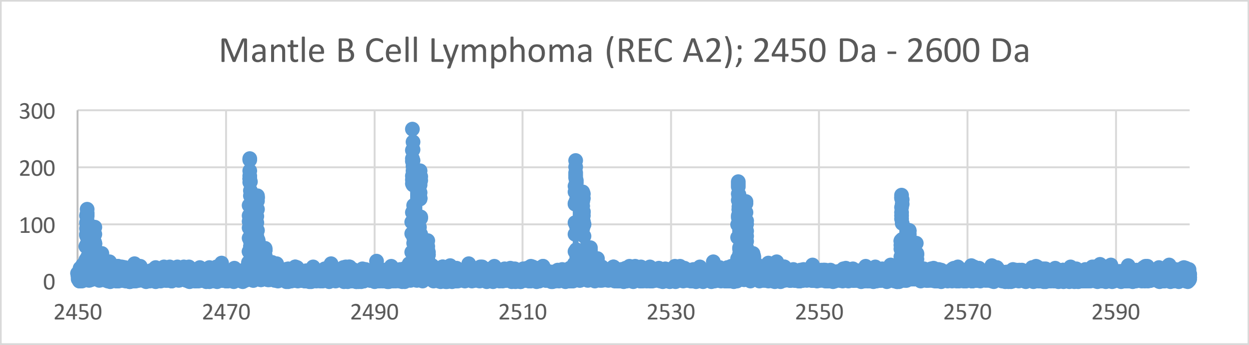
Mantle B Cell Lymphoma (REC A2); 2450 Da - 2600 Da
300
200
100
0
2450
2470
2490
2510
2530
2550
2570
2590

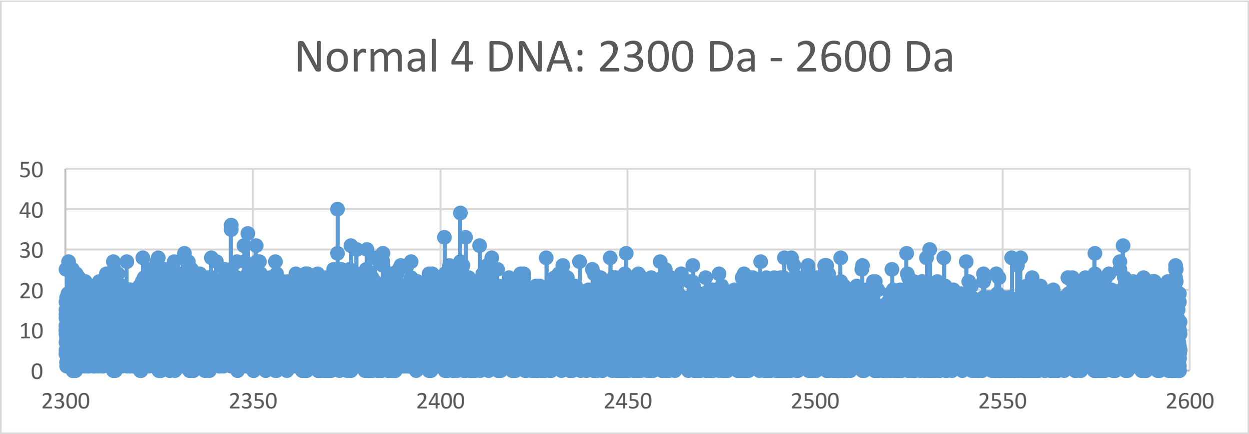
Normal 4 DNA: 2300 Da - 2600 Da
50
40
30
20
10
0
2300
2350
2400
2450
2500
2550
2600

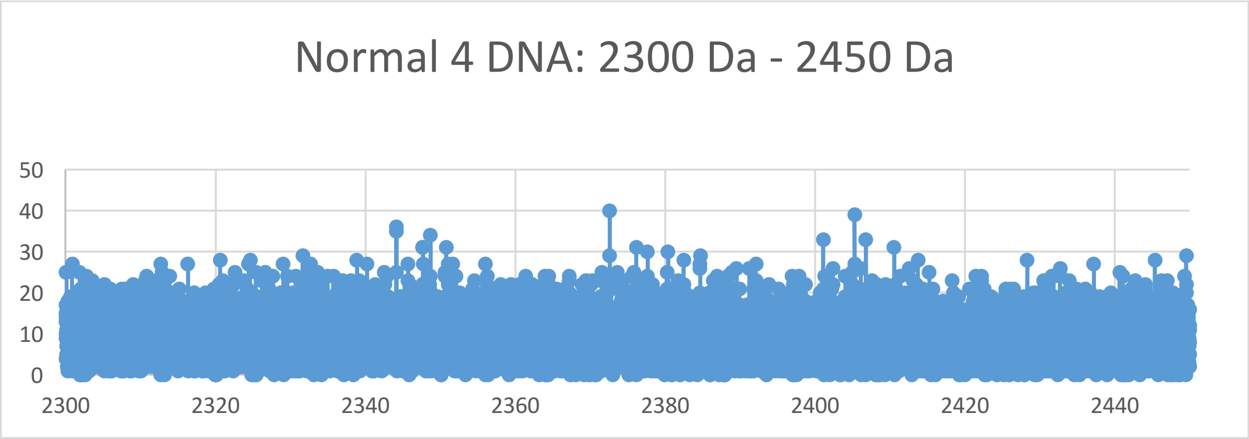
Normal 4 DNA: 2300 Da - 2450 Da
50
40
30
20
10
0
2300
2320
2340
2360
2380
2400
2420
2440

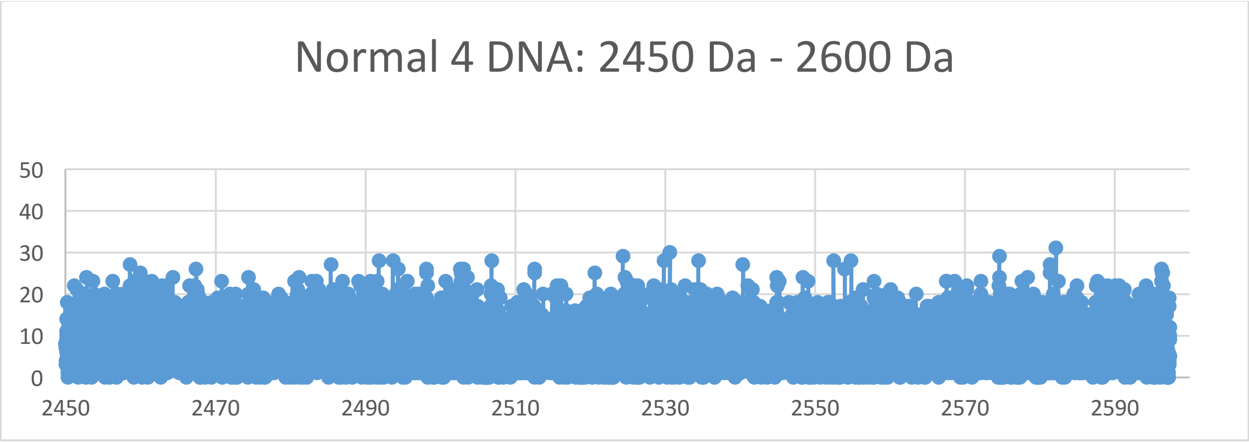
Normal 4 DNA: 2450 Da - 2600 Da
50
40
30
20
10
0
2450
2470
2490
2510
2530
2550
2570
2590

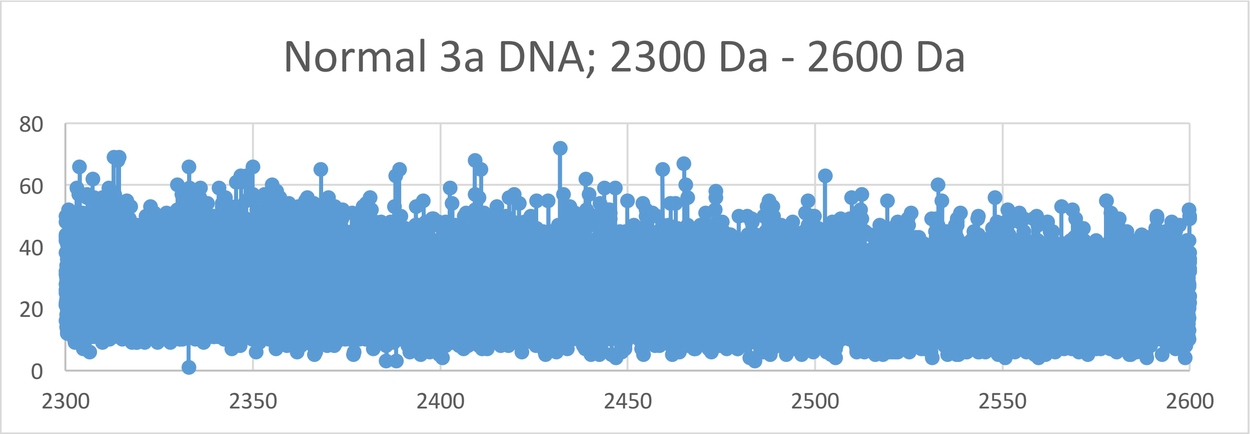
Normal 3a DNA; 2300 Da - 2600 Da
80
60
40
20
0
2300
2350
2400
2450
2500
2550
2600

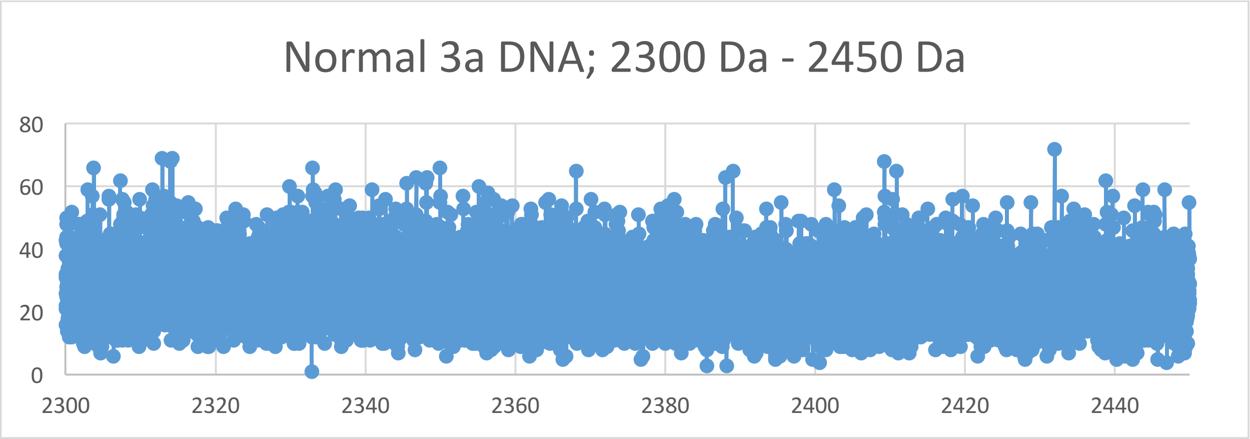
Normal 3a DNA; 2300 Da - 2450 Da
80
60
40
20
0
2300
2320
2340
2360
2380
2400
2420
2440

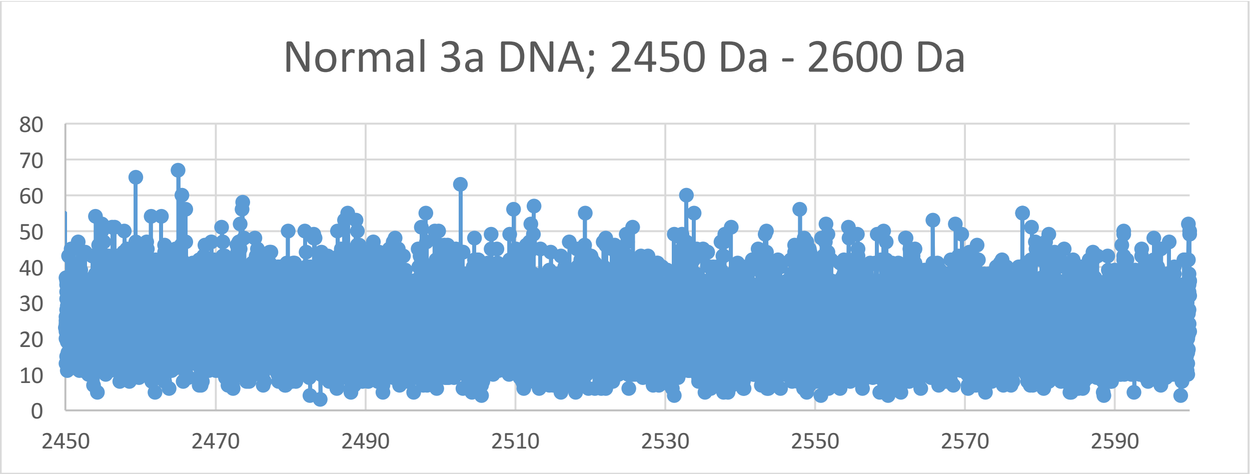
Normal 3a DNA; 2450 Da - 2600 Da
80
70
60
50
40
30
20
10
0
2450
2470
2490
2510
2530
2550
2570
2590

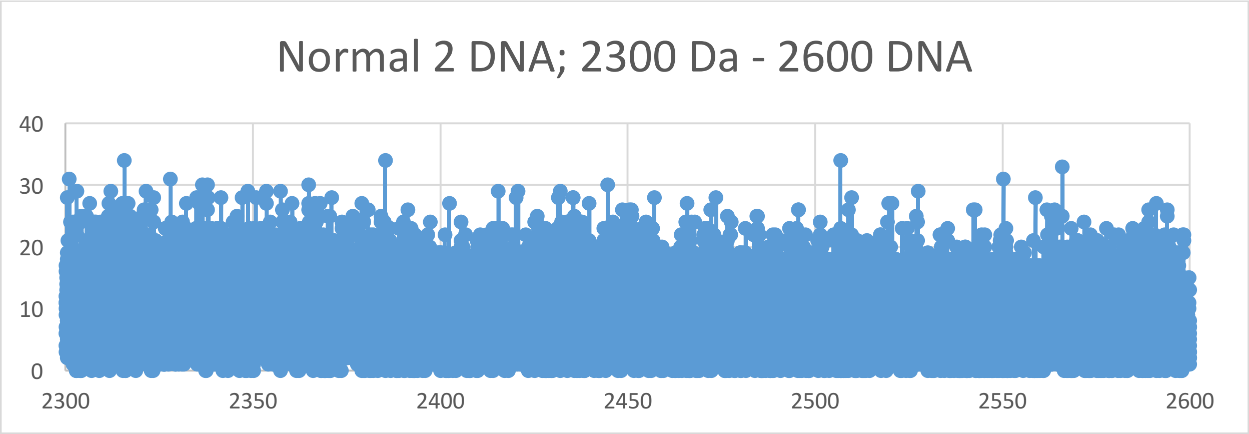

Normal 2 DNA; 2300 Da - 2600 DNA
40
30
20
10
0
2300
2350
2400
2450
2500
2550
2600


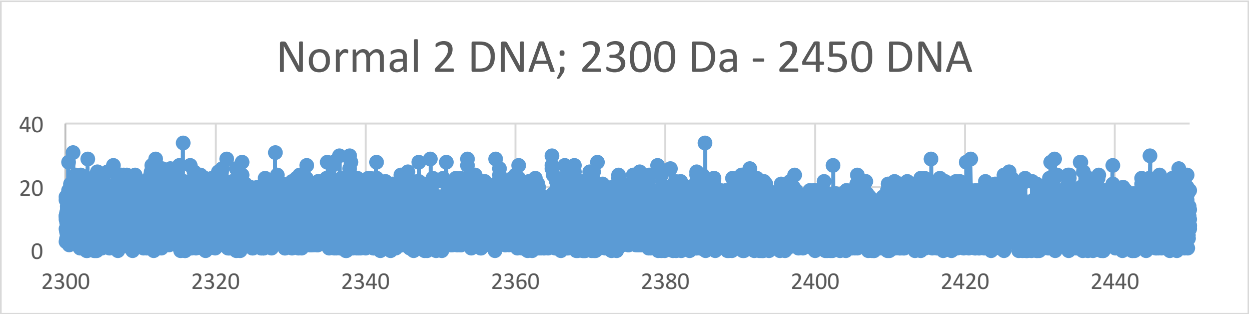

Normal 2 DNA; 2300 Da - 2450 DNA
40
20
0
2300
2320
2340
2360
2380
2400
2420
2440


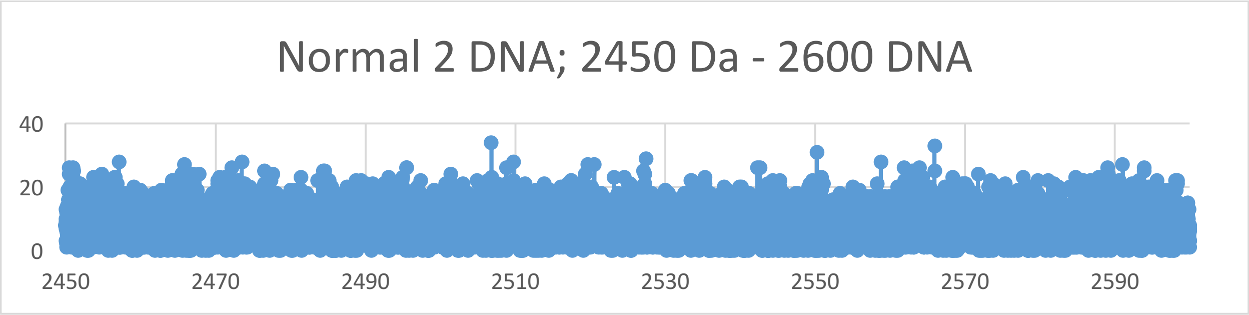

Normal 2 DNA; 2450 Da - 2600 DNA
40
20
0
2450
2470
2490
2510
2530
2550
2570
2590

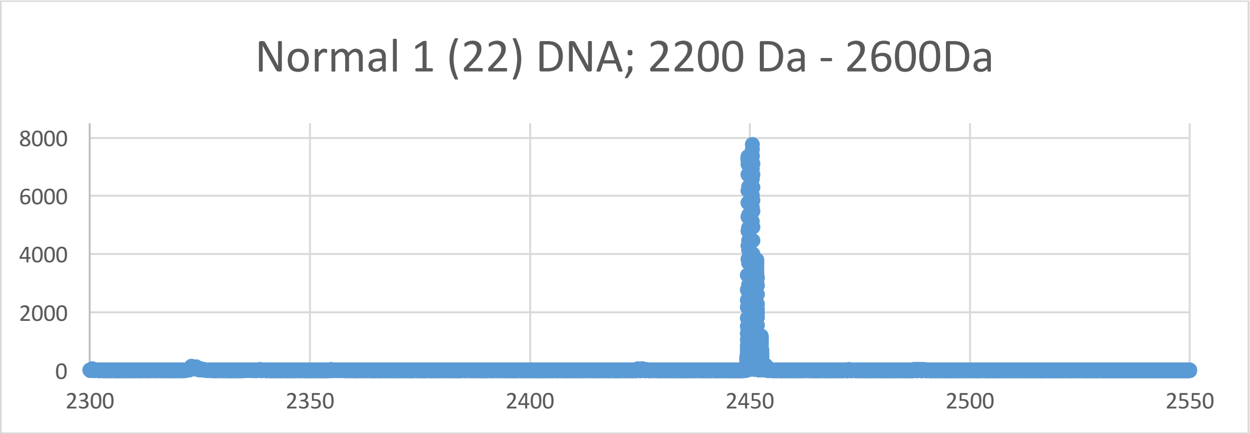
Normal 1 (22) DNA; 2200 Da - 2600Da
8000
6000
4000
2000
0
2300
2350
2400
2450
2500
2550

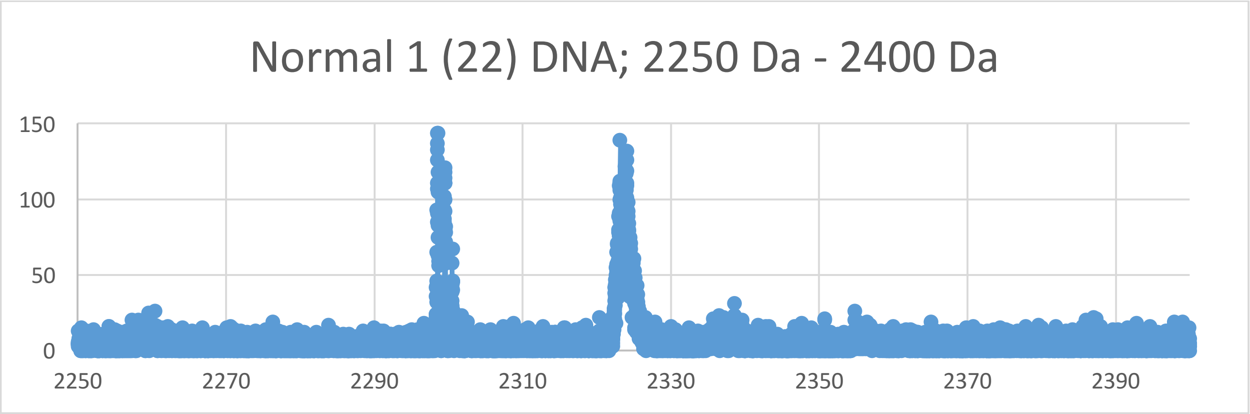
Normal 1 (22) DNA; 2250 Da - 2400 Da
150
100
50
0
2250
2270
2290
2310
2330
2350
2370
2390

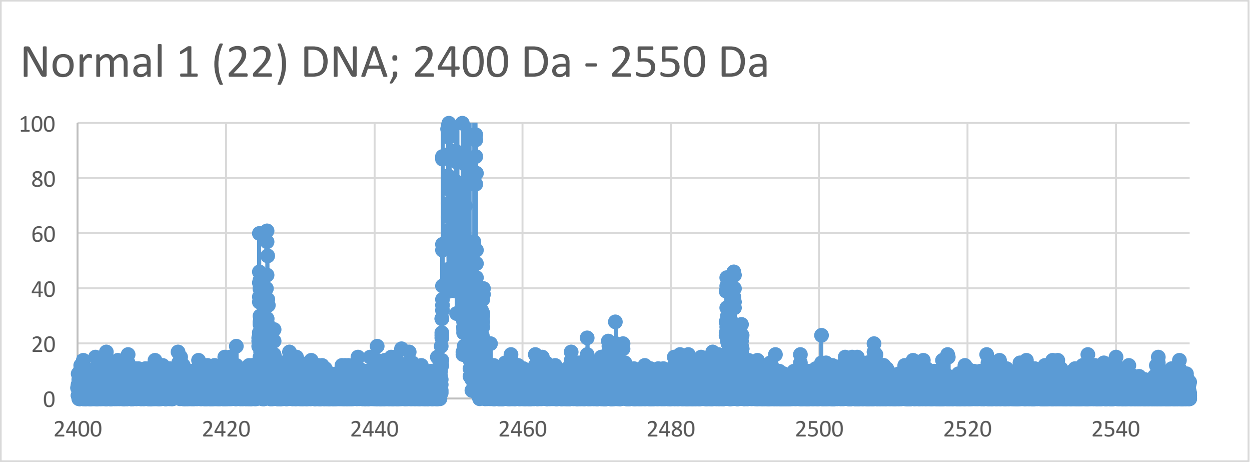
Normal 1 (22) DNA; 2400 Da - 2550 Da
100
80
60
40
20
0
2400
2420
2440
2460
2480
2500
2520
2540
